# Preparation of large-scale digitization samples for automated electron microscopy of tissue and cell ultrastructure

**DOI:** 10.1101/2021.03.02.433512

**Authors:** Carsten Dittmayer, Hans-Hilmar Goebel, Frank L. Heppner, Werner Stenzel, Sebastian Bachmann

## Abstract

Manual selection of targets in experimental or diagnostic samples by transmission electron microscopy (TEM), based on single overview and detail micrographs, has been time-consuming and susceptible to bias. Substantial information and throughput gain may now be achieved by automated acquisition of virtually all structures in a given EM section. Resulting datasets allow convenient pan-and-zoom examination of tissue ultrastructure with preserved microanatomical orientation. The technique is, however, critically sensitive to artifacts in sample preparation. We therefore established a methodology to prepare large-scale digitization samples (LDS) designed to acquire entire sections free of obscuring flaws. For evaluation, we highlight the supreme performance of scanning EM in transmission mode compared to other EM technology. The use of LDS will substantially facilitate access to EM data for a broad range of applications.

## Introduction

Electron microscopy (EM) continues to be a valuable tool in basic research and anatomical pathology. Aside from educated inspection of regions of interest (ROI) by the operator, EM traditionally serves to detect unexpected features in cells and tissues in an “open view” manner ^1^. In research, EM allows flexible recording of structures that may vary in size by three orders of magnitude, delivering excellent resolution down to the nanometer scale. In diagnosis of disease, EM is irreplaceable for muscle or kidney biopsies ^1, 2^. The traditional workflow for transmission electron microscopy (TEM) with an operator interactively screening specimens at low magnification and selecting targets of interest for snapshots at high resolution may, however, be time-consuming, laborious, and sensitive to bias so that its use became diminished over the past decades despite its undisputed values ^3, 4^. Consequent loss of expertise may lead to misinterpretations as recently reported for coronavirus particle structure in SARS-CoV-2 pandemic ^3, 5, 6^.

State-of-the-art EM technology now has made substantial progress to revolutionize the traditional approach. Introducing large-scale digitization of ultrathin sections, i.e. the automated recording of ROIs in an unbiased manner with overlapping, high-resolution TEM images (*mosaic tiling*) has been a major step forward. ROIs may now be as large as the entire ultrathin section. Collected datasets permit software-assisted pan-and-zoom examination independent of the physical location of the facility. This mode has been termed nanotomy, nanoscopy, or virtual EM ^4, 7–9^. A TEM equipped with a computer-driven stage may be used therefore, although with major restrictions ^4, 8, 9^. For a better and more flexible alternative, the traditional scanning electron microscope (SEM) has been modernized. Automated acquisition procedures, extended scanfields, and improved detectors have documented its benefits in large-scale digitization of extended ROIs ^10, 11^. For SEM in backscattered electron detection mode, sections are comfortably collected and stained on stable material such as silicon wafer substrate to be imaged individually for 2D nanotomy or, in series, for array tomography and volume EM ^11^. Inherent stability of the sections favours as well the recording of concomitant light microscopical signals for correlative light-and electron microscopy (CLEM) ^12^. However, high electrical conductivity and contrast are mandatory for these samples ^13^, and image acquisition time in SEM-BSD may be slow, ranging between 30 and 60 h per mm^2^ with 6 to 12 μs beam dwell time per pixel and 7.3 nm pixel size adjusted.

Alternatively, an SEM platform adjusted for scanning TEM mode (SEM-STEM) has proven superior imaging performance and was compatible with conventionally prepared samples ^14–16^. Here, a tenfold accelerated image acquisition at e.g. 1 μs dwell time is easily achievable. Still, the SEM-STEM mode depends on the use of EM grids carrying the sections, whose preparation is technically more demanding compared to silicon wafers. The use of grids notoriously encompasses a number of artifacts, but typically occurring wrinkles and contaminations, which may impair high-end automated image acquisition, visual examination and quantifying approaches substantially, had to be avoided ^14^. A substantial need for technological improvement prevailed to overcome this bottleneck in quality and prepare entire sections with virtual absence of flaws.

We therefore set out to establish a reliable and easy-to-implement workflow to produce sections on large-slot, filmed grids. Resulting large-scale digitization samples (LDS), virtually free of artifacts and with slot dimensions of up to 2 x 1.5 mm, were suitable to be digitized in conventional modern SEM-STEM systems at high throughput and also permitted individual high-end examination using advanced TEM systems. We further address the emerging need for a straightforward data processing pipeline. Resulting bigtif files allow in-depth analysis with the help of tools for annotation and measurement. We illustrate the potential of LDS to improve access to high-quality EM analysis for a broad user community.

## Results

### Requirements for large-scale digitization

Reliable and easy-to-implement methodology for the preparation of ultrathin sections was to be adapted for large-scale digitization with modern SEM-STEM or TEM systems, serving to analyze a broad range of native or experimental tissues and cells. Since artifacts critically impair recording at large scale, a central goal was to prevent formation of wrinkles, stain precipitates, and other contaminations. Therefore, selection and treatment of grids, collection of large-size sections, and staining steps were to be specified. For simplicity, the resulting specimens comprising the filmed slot grid and the large ultrathin or semithin section were termed large-scale digitization samples (LDS). EM recording of LDS had to meet the criteria of high imaging speed, optimal tile size, adequate automation, high structural resolution, and satisfactory signal-to-noise ratio (SNR).

### Preparing support films for LDS

Grids were coated using pioloform film for its established mechanical and thermal stability ^17^. Film had to be placed on the shiny side of the grid to secure a smooth transition between slot and metal surface (Fig. 1a-c). Typical film artifacts such as series of spots of different size and shape, deformations resulting from focal attachments to glass, or variations in thickness could be substantially reduced (Fig. S1 and Fig. S2).

**Fig. 1.**
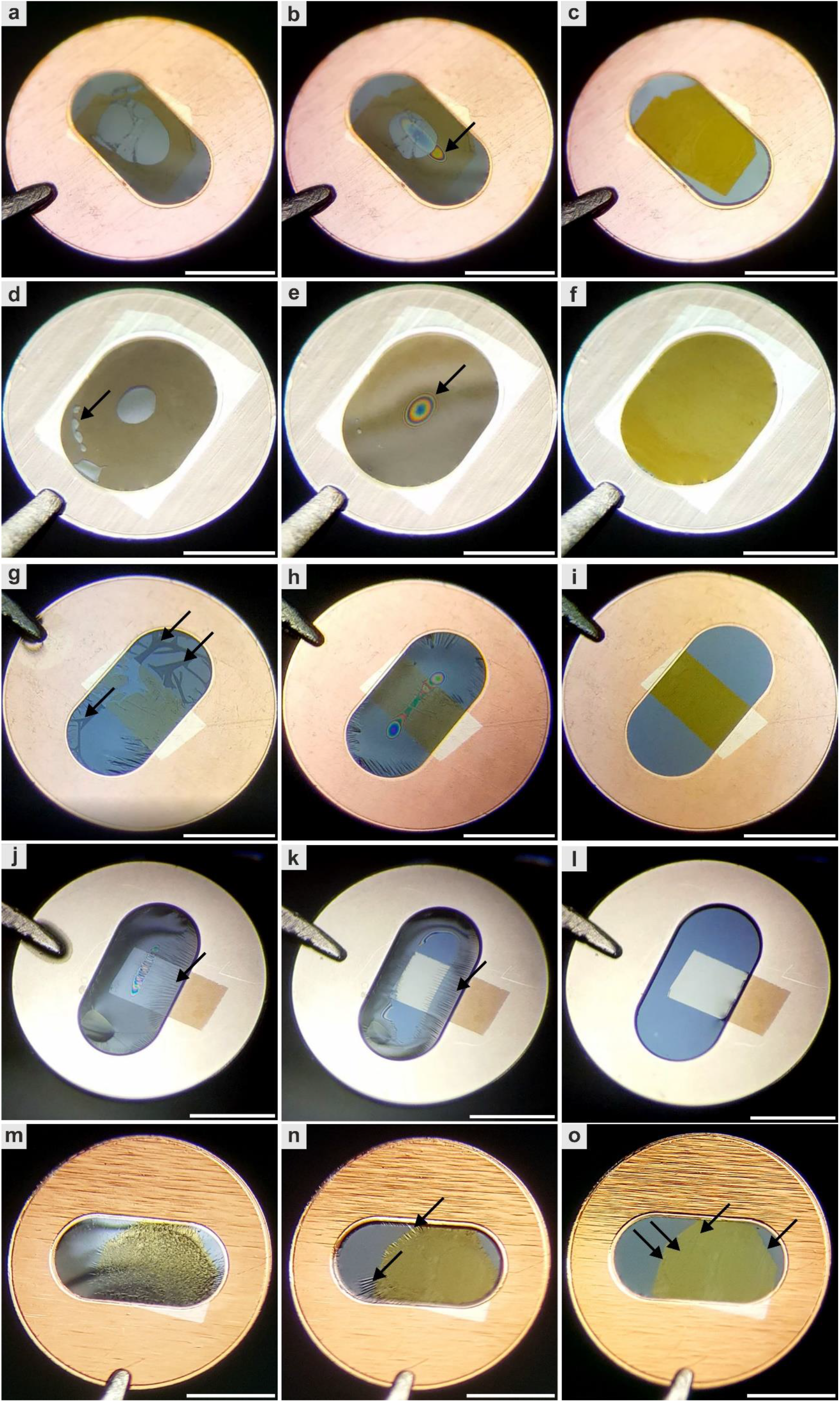
Large-scale digitization sample preparation; attachment and drying of ultrathin sections. **a** to **c**, A pioloform film has been placed on the shiny side of a regular slot grid and the filmed grid glow-discharged and ethanol-smoothened; after collection of a section, its attachment and drying process is demonstrated sequentially. Attachment of the section starts from the periphery, typically indicated by a central, grey meniscus, shrinkage, and formation of oblong Newton rings (arrow in **b**) expanding with evaporation during max. 2 min. The result is a homogeneous yellow appearance of the fully attached section. **d** to **f**, The same sequence was maintained using large-slot grids with 2 x 1.5 mm opening size which allow the use of larger sections; note the rounded Newton ring (arrow in **e**). Tissue components like fat cells (arrow in **d**; kidney) may slow the attachment. **g** to **l**, After glow discharging alone, wrinkles of geographical-appearance (film and section placed on the shiny side of the grid; arrows in **g**) or fine-needle aspect (film and section placed on the dull side of the grid; arrows in **j**,**k**) occurred; although **i** and **l** suggest a resulting smooth attachment of the sections, omission of the ethanol-smoothening step caused persistent wrinkles in both variants which may be visualized stereomicroscopically with tilting the grid, or ultrastructurally. **m** to **o**, Omission glow discharging and the ethanol-smoothening steps results in high hydrophobicity with inadequate water menisci and wrinkles which usually were also prominent after drying (arrows). Scale bars, 1 mm (**a**-**o**).

### Providing hydrophilicity of filmed grids

Hydrophilicity of the support film was a strong requirement to avoid artifacts when mounting sections on the grid. This was achieved by glow-discharging the grid surface using a vacuum evaporator. A current of 6.6 to 6.9 mA and a duration of 10 s were optimal for the process; deviations in time caused wrinkle formation during collection due to over- or underhydration of the support. Prior to collection, grids were individually dipped in ethanol and distilled water. This step was critical to prevent fine needle- or “geographical”- folds (Fig. 1g-l).

### Preparing sections for LDS or silicon wafer

Tissue blocks were trimmed to a slim leading edge, and areas of pure resin were removed for optimal consistency of the samples. Ultrathin sections (60-70 nm) were carefully prepared to avoid folds or knife marks. For electron tomography, semithin sections (200-250 nm) were cut and intensely stretched using xylene vapor to reduce compression. Mechanical vibrations of local or external origin had to be controlled to avoid chatters (Fig. S2).

### Collecting and attaching the sections for LDS or silicon wafers

For imaging with SEM-STEM or TEM, the filmed grids were used. After sectioning, the grid was submersed in the water bath of the diamond knife and the sections directed to the site of attachment at a straight borderline between water and grid surface using an eyelash. Here, the grid was to be held at its edge near the short side of the slot by a forceps to ensure straightness of the borderline (Fig. S3). The section was picked up by lifting the grid out of the water trough. Holding the grid in horizontal position by the forceps, the section was then dried. Quality of the drying process had to be closely monitored. It comprised two different phases, one, the drying of the film’s back side which had to start from the periphery (Fig. 1a), and another, the drying between section and film which had to start from the center as indicated by Newton ring formation (Fig. 1b). Ideally, no folds or wrinkles resulted from this initial drying process. For completion of the process, drying was then extended for another 1 or 2 d to avoid ring-shaped wrinkles during consecutive staining. For comparative SEM-BSD imaging, sections were collected on silicon fragments that had been glow-discharged for 60 to 90 s The size of the fragments allowed to collect multiple sections.

### Staining of sections for LDS or silicon wafers

For staining of sections on grids, aqueous uranyl acetate solution was superior to an ethanolic solution to ensure an adequate drying process after staining that optimally started at the center of the grid (Fig. 1a-c). To finalize LDS preparation, grids were dried again in horizontal position to ensure adequate quality indicated by Newton ring formation, which ideally displaced contaminations towards the periphery of the grids. Sections on silicon substrate were comfortable in handling due to their stable adherence to and solidity of the support, which minimized the risk of wasting samples during staining. Details on sample diversity and a troubleshooting table are provided in the Supplementary Information (Supplementary Table 1-3).

### Technical specification of EM systems available for nanotomy

The use of TEM provided us with 2k and 4k field dimensions which implied an imaging speed ranging near 10 or 15 s, respectively; time was determined by stage movement, camera exposure, and the chosen readout properties. Using SEM-STEM or SEM-BSD, field dimensions of up to 32k or, practically, individual image fields of 50 to 100 μm per side were available. With flexible adaptation of dwell time and signal averaging mode, optimal SNR and imaging velocity provided excellent nanotomy recordings. Atlas 5 software provided the necessary robust and automated control of stage movement as well as autofocus functions (see Supplementary Table 4 for a comparison of the different EM techniques).

### Imaging in TEM

For conventional imaging, minimized preparation artifacts and absence of obscuring bars in LDS permitted a significantly more rapid navigation and improved imaging results as compared to standard samples (Fig. S1). Medium-sized ROIs of muscle, nerve, liver (Fig. S4) or brain and kidney sections (Fig. 2a,d) were digitized in flawless quality using LDS.

**Fig. 2.**
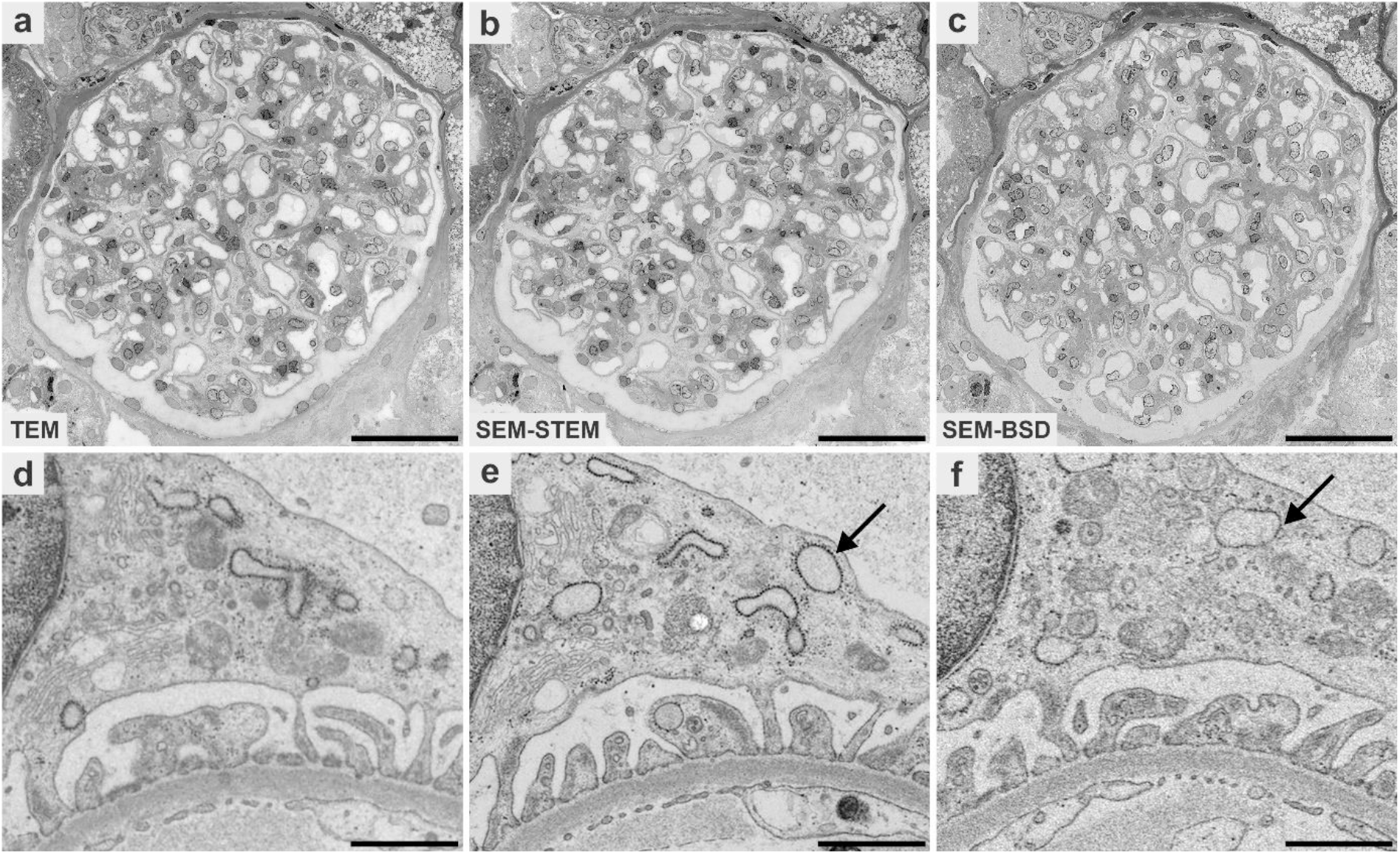
Imaging properties of three different EM systems in comparison using the same diagnostic kidney sample. Images for large-scale digitization were recorded by transmission electron microscopy (TEM; **a**,**d**), scanning transmission electron microscopy (SEM-STEM; **b**,**e**), and backscattered electron detection (SEM-BSD; **c**,**f**). The pixel size of 7.3 nm was selected throughout, and scanning time for SEM-STEM- and SEM-BSD imaging was adjusted to match with TEM imaging time (approximately 1 h). Images were normalized for brightness and contrast using IMOD. **a** to **c**, Overviews demonstrate similar quality. **d** to **f**, At higher digital magnification (same podocyte) the ultrastructural detail in TEM appears mildly blurred in **d** compared to **e** with its more crisp appearance in SEM-STEM. **f**, Quality in SEM-BSD imaging appears similar to SEM-STEM and TEM but reveals lower signal-to-noise-ratio (SNR). Resolution of details such as individual ribosomes is superior with SEM-STEM in **e** as compared to SEM-BSD in **f** (arrows). The displayed images were prepared via TrakEM2 tif export files for highest quality in order to minimize image compression artifacts. Scale bars, 50 μm (**a**-**c**), 1,000 nm (**d**-**f**). See also www.nanotomy.org for internet browser-based pan-and-zoom analysis of the full resolution datasets.

### Digitizing at large-scale in SEM-STEM

To analyze LDS in SEM-STEM, imaging time for a selected medium sized ROI, e.g. a glomerulus, was 1 h at 7.3 nm pixel size and 3 μs dwell time. Resulting images displayed excellent image quality and resolution of detail such as the filtration barrier with diagnostically relevant tubuloreticular inclusions or electron-dense deposits (Fig. 2b,e and Fig.3). Shortening dwell time to 1 μs further reduced the process to 25 min with no major loss in image quality. Advanced automation and throughput, available in state-of-the-art SEM-STEM systems, allowed us to record up to 12 LDS per imaging cycle lasting 6 d at 7.3 nm pixel size and 1 μs dwell time, with each LDS carrying a single ultrathin section sized 2 x 1 mm (Fig. 4 and Fig. 5a). Imaging with very short dwell time still resulted in acceptable SNR (Fig. 6a-c and Fig. S5). Thereafter, LDS were facultatively transferred to the TEM for imaging at highest achievable resolution.

**Fig. 3.**
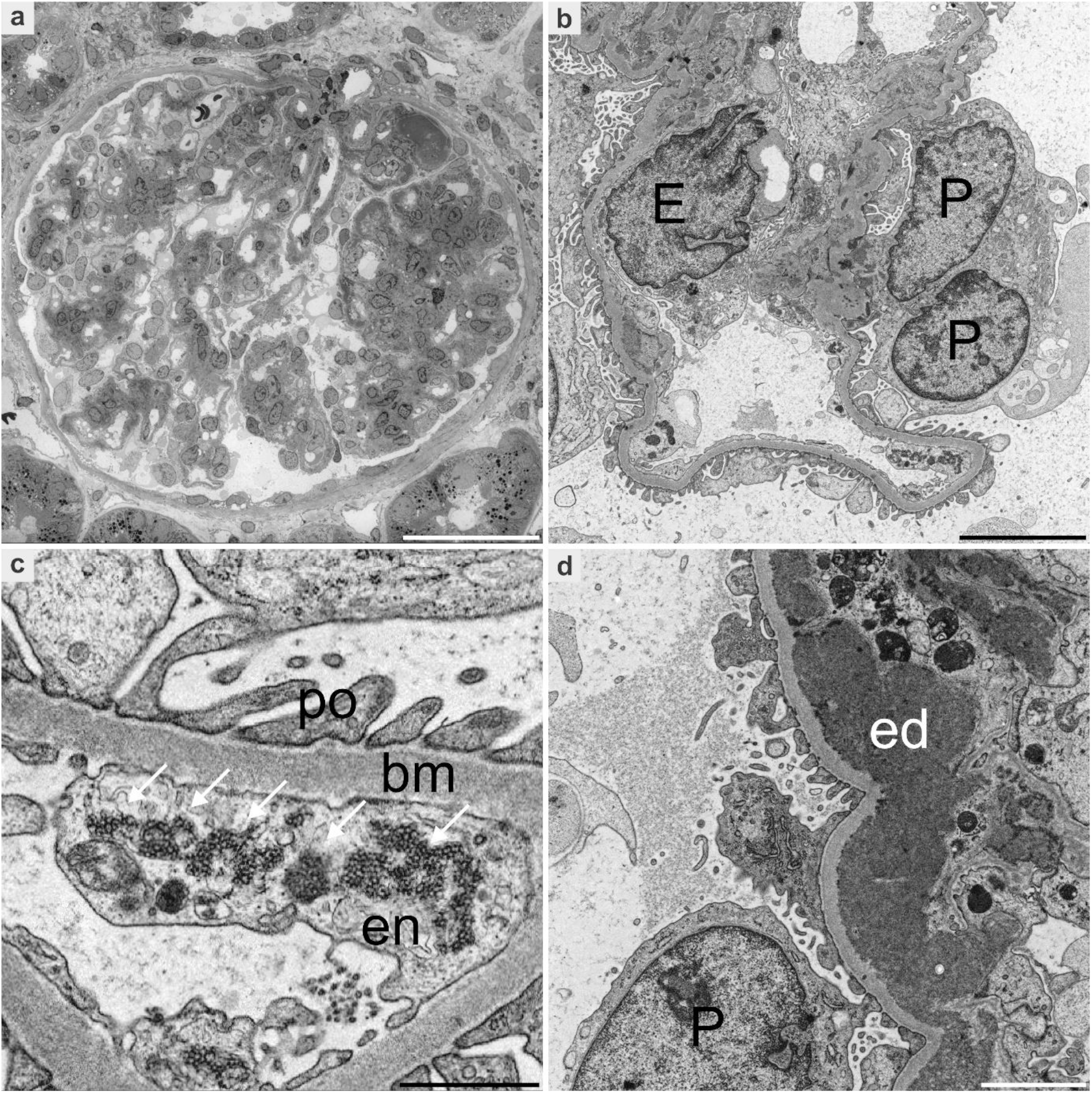
Diagnostic kidney sample with lupus nephritis, recorded by large-scale scanning transmission electron microscopy. **a**, Overview of an entire glomerulus (240 x 240 μm); digitization was performed within 25 min using 1 μs dwell time and 7.3 nm pixel size. **b**, Nuclei of an endothelial cell (E) and a podocyte (P) are readily identified in the coherent, large-scale dataset. **c**,**d**, Ultrastructural detail of the podocyte processes (po), glomerular basement membrane (bm), and endothelium (en) with prominent tubuloreticular inclusions (arrows) or electron dense deposits (ed) are well resolved. Images as shown were prepared from screenshots of an Atlas 5 export dataset. Scale bars, 50 μm (**a**), 5 μm (**b**), 1 μm (**c**), 2 μm (**d**). See also www.nanotomy.org for internet browser-based pan-and-zoom analysis of the full resolution dataset.

**Fig. 4.**
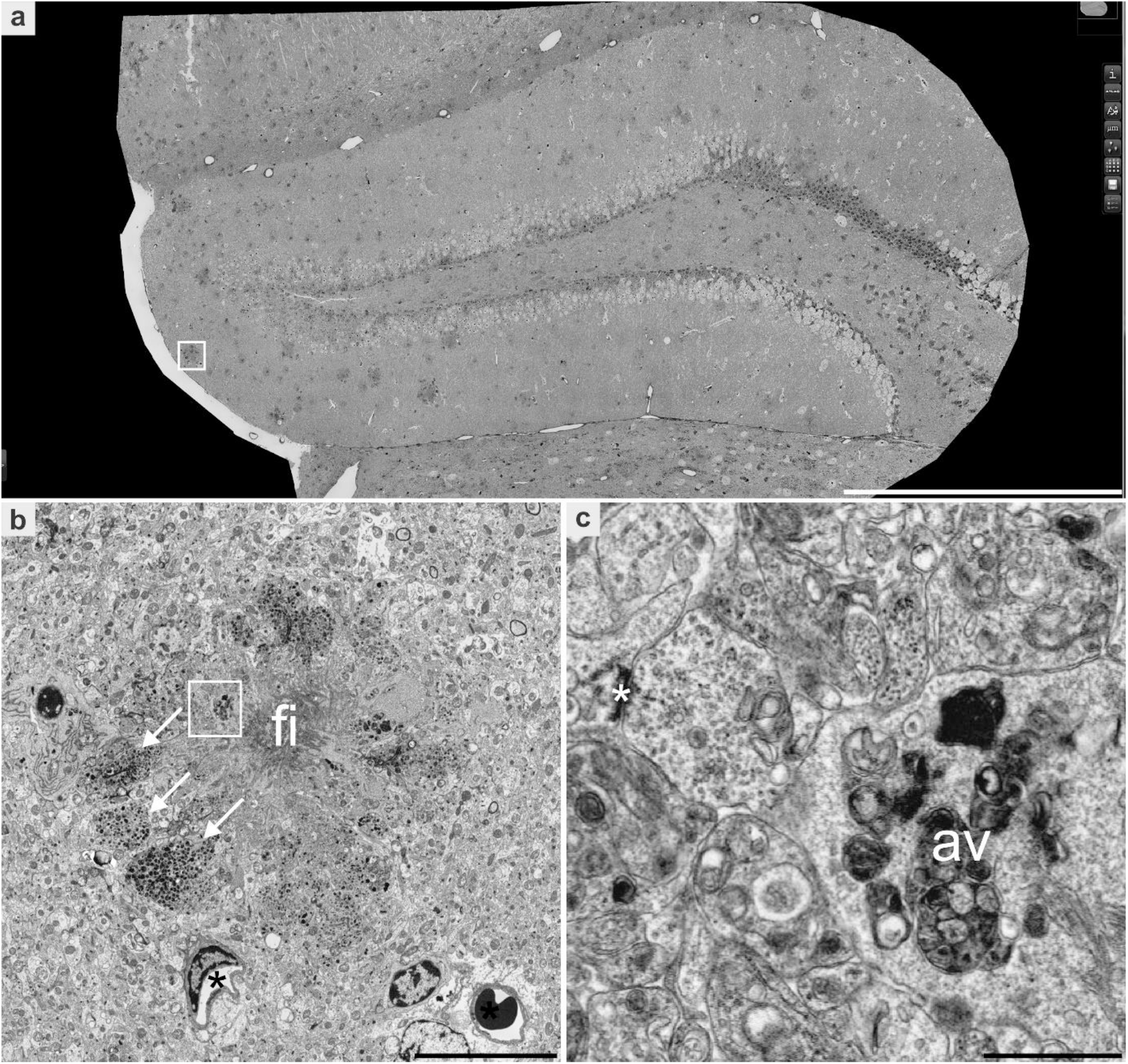
Dentate gyrus of a mouse model for Alzheimer’s disease, digitized by large-scale scanning transmission electron microscopy of an entire ultrathin section. **a**, Overview of the dentate gyrus, measuring 1,800 x 1,000 μm, digitized in about 12.5 h using 1 μs dwell time and 7.3 nm pixel size. Overview shows plaques as electron dense structures. **b**, Digital zooming of the boxed area in **a**; detail of a plaque, shows dystrophic neurites (arrows) and distinct fibrils (fi); neighboring capillaries with empty lumen (asterisk). **c**, Digital zooming of the boxed area in **b**; dystrophic, enlarged neurites are filled with autophagic vacuoles (av); note a synapse (asterisk). Images as shown were prepared by screenshots of an Atlas 5 export dataset. Scale bars, 500 μm (**a**), 10 μm (**b**), 1,000 nm (**c**). See also www.nanotomy.org for internet browser-based pan-and-zoom analysis of the full resolution dataset; see Supplementary Video for a demonstration of this analysis.

**Fig. 5.**
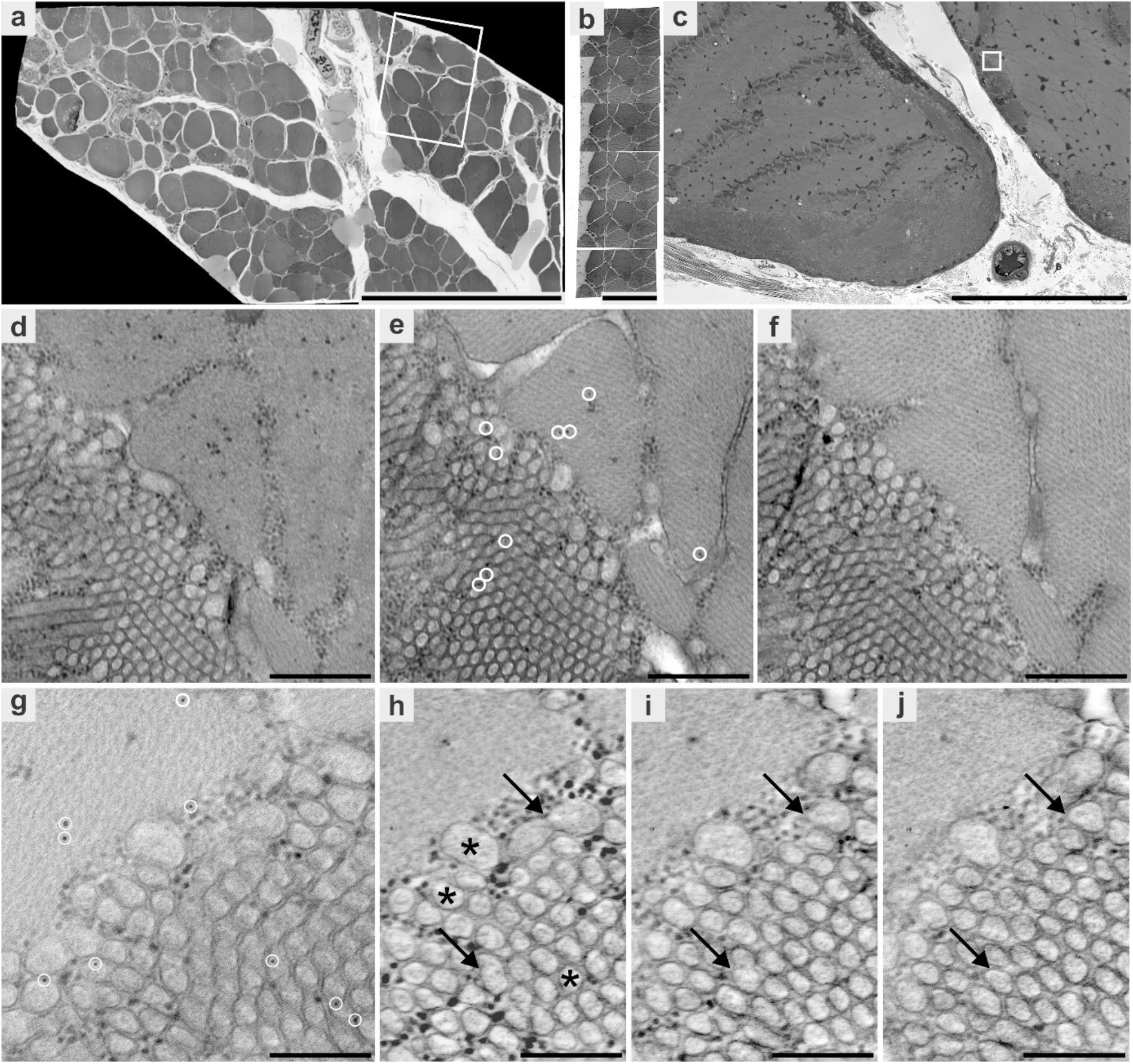
Imaging myopathy with tubular aggregates by consecutive screening with scanning transmission electron microscopy (SEM-STEM) and electron tomography (ET). **a,** An entire ultrathin section was digitized with SEM-STEM to select a region of interest (ROI; box) with well-preserved muscle fibres containing tubular aggregates. **b**, After trimming the ROI, a ribbon of six serial semithin sections was collected on a slot grid, stained, added with fiducial gold particles, and digitized at low resolution with SEM-STEM to locate a region containing tubular aggregates in adequate orientation. **c**, This region was digitized in all of the six sections to evaluate detail of the tubular aggregates as well as number of fiducial particles. **d**-**f**, From the region in **c**(box), a digitally magnified area containing aggregates and adjacent contractile filaments is shown consecutively in three of the serial sections; the middle section in **e** shows the fiducial markers. The grid carrying the serial sections was then transferred to a transmission electron microscope (TEM) to acquire a tilt series for ET. **g**, Standard TEM image in **e** reveals adequate distribution of gold particles to allow for 3D-reconstruction in ET. **h**-**j**, Representative digital slices of the reconstructed volume of the section clearly resolve individual tubular aggregates, demonstrating differences in their size and shape (asterisks) and branching features (arrows in **h**-**j**). Scale bars, 500 μm (**a**), 200 μm (**b**), 20 μm (**c**), 500 nm (**d**-**f**), 250 nm (**g**-**j**). See also www.nanotomy.org for internet browser-based pan-and-zoom analysis of the full resolution datasets of **a** and **c**,**e**.

**Fig. 6.**
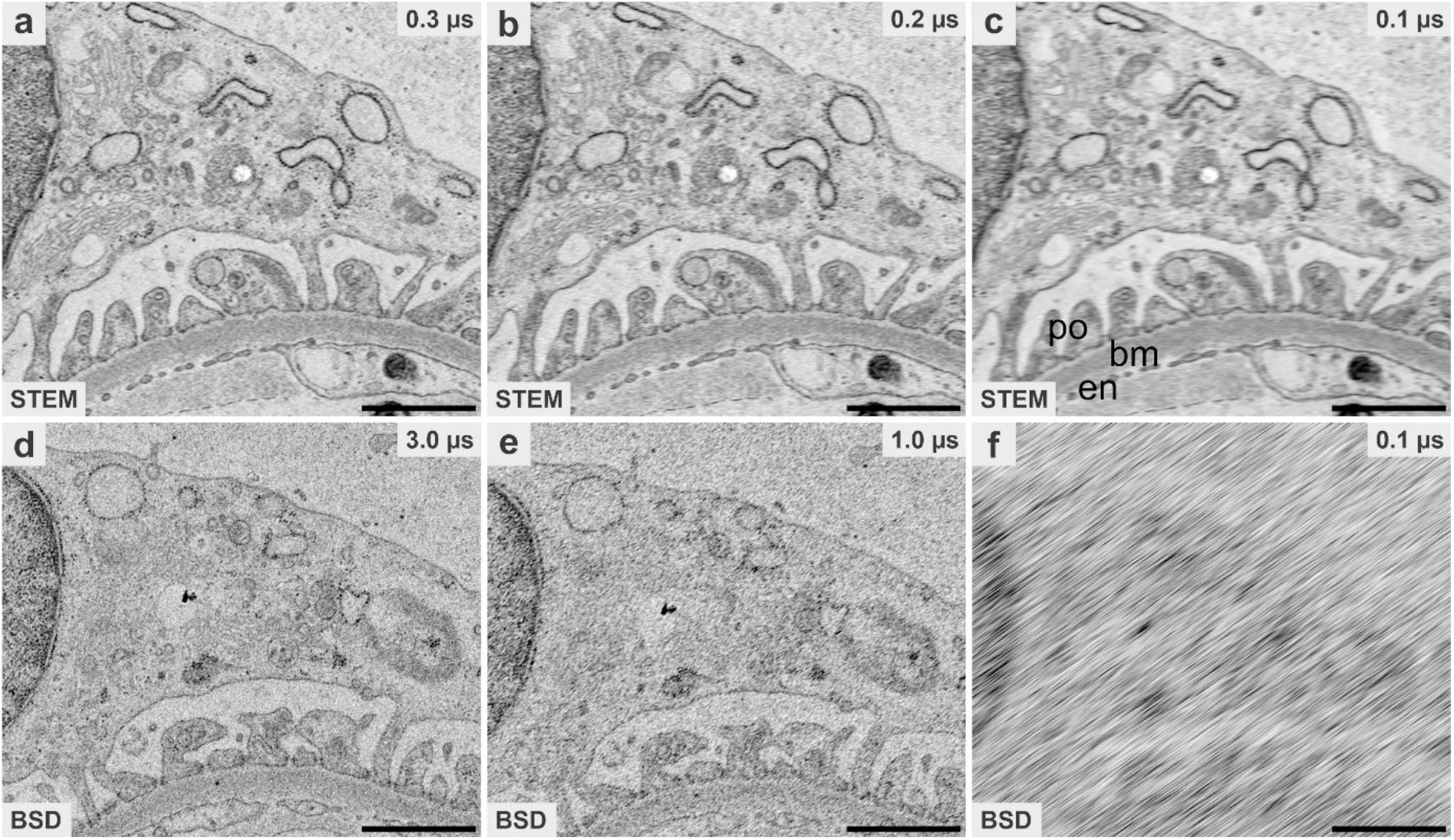
Increased velocity in scanning transmission electron microscopy (SEM-STEM; a-c) compared to backscattered electron detector (SEM-BSD; d-f) imaging mode. **a**, Rapid scanning with 0.3 μs dwell time in SEM-STEM still allows for good resolution of subcellular detail of the kidney filtration barrier, whereas with 0.2 μs in **b** and 0.1 μs in **c**, structure becomes increasingly blurry. Ultrastructural detail of the podocyte processes (po), glomerular basement membrane (bm), and endothelium (en). With SEM-BSD, much longer dwell times (3 μs) are required to achieve comparable image quality; still, signal-to-noise ratio is inferior compared to SEM-STEM (**d** vs. **a**). Faster imaging with SEM-BSD (1.0 μs in **e**, 0.1 μs in **f**) increasingly compromise quality. The displayed images were prepared from tif raw data for highest quality in order to minimize image compression artifacts. Scale bars, 1,000 nm (**a**-**f**). See also www.nanotomy.org for internet browser-based pan-and-zoom analysis of the full resolution datasets.

For electron tomography (ET), the SEM-STEM system was used to record LDS carrying serial semithin sections (Fig. 5b). ROIs from these sections were digitized at high resolution, profiting from the absence of artifacts as well. Preparation of multiple sections allowed to either select a single section with optimal detail for 3D analysis of small volumes or analyze volumes that extend the thickness of single sections by applying ET to the same ROI of consecutive sections in form of serial-section ET. Here, shuttling of LDS to the TEM allowed to apply complementary imaging in tomography mode. Combining ET with the respective overview data generated by SEM-STEM thus preserved contextual orientation of the detail (Fig. 5b,c). A robust basis was thus created to reliably find small ROIs within sections which had been selected for ET before, and the risk of electron beam damage to the section, resulting from extensive live examination, was reduced as well.

### Digitizing at large-scale in SEM-BSD

Analyzing ultrathin sections placed on silicon substrate with SEM-BSD, we adapted imaging time for medium-sized ROIs to meet the duration of 1 h. As for SEM-STEM, we selected 7.3 nm pixel size and 3 μs dwell time for digitization (Fig. 2c,f). Conventionally embedded material required careful adjustment of further imaging parameters such as working distance and accelerating voltage. Keeping the former small (4-5 mm) and setting the latter at 8 kV resulted in satisfactory SNR typically found in lipid-enriched, compact tissue samples like nerve and muscle, which accumulate high amounts of heavy metals during preparation. SNR was further improved by uranyl acetate *en bloc* staining. Samples with lower heavy metal content such as kidney biopsies required setting of the aperture diameter at 60 rather than standard 30 μm. Resulting datasets demonstrated an overall satisfactory image quality, however, the subcellular detail was often obscured by inadequate SNR; 10-fold extended dwell times were required to obtain adequate results comparable to SEM-STEM (Fig. 6).

### Data processing and analysis

Generation of large-scale datasets required the stitching of image tiles to coherent datasets and their conversion into zoomable file format. For TEM, datasets of medium-sized ROIs were suitably generated with TrakEM2. Increasing numbers of image tiles (e.g. 1,000+ 2k images) to be stitched together rendered the procedure vulnerable for misalignments. The stitched datasets were converted into bigtif files to enable pan-and-zoom examination and in-depth analysis with QuPath via annotation and measurements ^18^. For SEM-STEM and SEM-BSD, Atlas 5 software permitted convenient stitching and export, but parameters serving to adjust eventual misalignments were limited, which impaired quality of some of the data. The Atlas 5 datasets for browser-based examination provided pan-and-zoom options, basic tools for analysis and convenient implementation into online repositories such as nanotomy.org. Alternatively, stitching and export to bigtif files were performed as for TEM datasets. Here, larger image tiles permitted efficient batch processing and export of multiple, large datasets in an overnight span.

## Discussion

Limitations of conventional TEM imaging are traditionally based on the need to manually select single ROIs which are then recorded individually in the appropriate ranges of resolution. This time-consuming mode has been technically overcome by automated large-scale image acquisition enabling nanotomy ^4, 9, 15, 19^. Our aim was to obtain EM samples suitable for this destination. ^14, 15^ We have presented methodology to prepare sections mounted on large-slot grids, termed LDS for their property of minimized artifacts and potential for high-end data acquisition. We have furthermore compared evaluation of LDS in TEM and SEM-STEM systems and discussed the alternative SEM-BSD approach. A robust data processing pipeline, based on open-source software, was proposed in order to create bigtif files which enabled in-depth analysis.

The search for causes and elimination strategies of artifacts in EM sections such as folds, staining precipitates and other contaminations, which obscure their inspection, has been challenging for decades ^20–22^. Quality of the support film has been another source for artifacts; the delicate structure of commonly used support films led to the search of alternatives in materials and their processing, albeit without a breakthrough ^23, 24^. Large-scale digitization of sections required further refinement in sample preparation to achieve adequate standards ^14, 15^. The measures we took in order to produce reliable support films with minimal amounts of artifacts have been successful (Fig. S2). Standard slot grids with large opening sizes (2 x 1 or 2 x 1.5 mm) thus carried a pioloform film which also did not require additional carbon coating ^9^. Thus, entire sections prepared from resin blocks in commonly used sizes could be collected for unobscured examination in the EM ^7, 19^.

Optimizing the collection step of the sections onto the large-slot grids, control of the hydrophilicity of the grid surface was essential, since focal drying points confounded regular attachment of the sections to the pioloform film. Inadequate hydrophilicity of the grids was indeed problematic, since overhydration caused an oversized bulk of water on the grid during collection, resulting in improper attachment of sections, whereas underhydration led to inhomogeneous wetting of the film; both extremes inevitably led to artifacts.

Glow discharging of the filmed grids prior to collection of the sections essentially solved the problem, providing adequate hydrophilicity in a highly controllable manner. This step has formerly been in use to spread fluids on filmed grids in special applications, or to reduce wrinkles when collecting sections on silicon or plastic substrate ^13, 25–27^. Reducing surface tension of the film by an ethanolic smoothening step prior to collection of the sections was essential as well. Monitoring the subsequent drying process of the sections on the film served as an important quality control.

Staining of sections was necessary, since we envisaged optimal SNR and imaging speed with conventionally embedded samples. Specific protocols for high-contrast *en bloc* staining were thus not applicable ^13^, so that the risk of staining artifacts had to be accepted. A known pitfall is the potential reaction between uranyl acetate and lead citrate, resulting in massive contaminations ^22^. This may be related with a vulnerable surface texture resulting from knife marks, folds, or cracks ^22, 28^, which were minimized by LDS. Detachment artifacts of sections, forming ring-shaped wrinkles, may occur as well during the staining procedure. Similar artifacts have been described for semithin sections ^29^, possibly caused by reaction of free epoxy groups with water. To overcome this problem, we simply extended the drying period of sections for LDS after collection for up to two days, based on the assumption that free epoxy groups may be inactivated by oxidation.

The resulting, refined LDS produced excellent results in conventional imaging and large-scale digitization of medium-sized ROIs using a TEM, thus demonstrating viability in routine research and pathology applications. The main goal of our methodology was, however, to confirm whether LDS were reliably applicable for digitization of entire sections at a high throughput. Here, even modern TEM systems reach their limitations owing to small image fields and limited sample-holder capacities ^9, 14^. Therefore, SEM-STEM was the optimal choice for its flexible capacities in automated digitization, with the additional advantage of minimal operator involvement ^14^. The general benefits of SEM-STEM systems have been previously described, but it was also acknowledged that their performance was limited by flaws on the samples ^14, 15^. This drawback has been essentially overcome by the present LDS methodology. Using LDS, imaging parameters such as dwell time and pixel size were varied to critically analyze structural detail in various tissues at intermediate resolution. High-resolution imaging using pixel sizes between 2 and 5 nm has been accomplished as well, albeit at the cost of long acquisition time ^15^.

Selecting the SEM-BSD system as an alternative approach facilitates the collection of large sections singly or in series using silicon wafers ^11^. Absence of fragile support films as well as an unobscured imaging potential for a larger tissue context favour this approach ^30^. Using a standard acquisition speed of 3 μs dwell time, image quality of conventionally embedded samples was not sufficient in SEM-BSD mode owing to poor SNR, and structural detail like components of the glomerular filtration barrier was insufficiently resolved. Acquisition times ranging up to tenfold that of the SEM-STEM mode were required for similar outcomes which clearly limits the use of SEM-BSD. Imaging with SEM-BSD may be improved with advanced *en bloc* staining protocols, although other shortcomings related to lipid content of the respective tissues may be introduced hereby ^13, 30^. Another drawback would be the resulting exclusion of already embedded tissues from routine pathology archives for research ^16^.

While the SEM-STEM mode provided superior imaging performance in 2D nanotomy using LDS, collection of serial sections for 3D analysis was cumbersome due to the fragile nature of the support film. Here, the silicon substrate used in the SEM-BSD system delivered better performance with sections stably adhered. This approach is therefore preferable for 3D array tomography ^11^. Stable adherence to substrate also favours correlative light-EM (CLEM) by reducing distortion of sections in consecutive imaging procedures ^12^. Imaging quality of hydrophilic resin sections imaged by SEM-BSD may, however, be limited by low contrast. On the other hand, LDS prepared with hydrophilic resin sections stained for immunogold may be favorable with SEM-STEM for better contrast ^31^. Likewise, LDS may be helpful in localizing electron dense polymers generated via genetically encoded tags that preserve the microanatomical context ^32^.

Routine application of large-scale digitization using LDS may ideally be achieved in specialized EM facilities to make nanotomy available to a broad usership, matching current trends to centralize state-of-the-art technology for the sake of cost reduction and improved quality management ^3, 33^. In research applications, digitization of entire sections will substantially improve analyses of cultured cells, organoids, animal models and human tissues with more detail and at higher throughput ^7–9, 15, 34^. Medical diagnostic applications will be improved as well by the use of LDS, since for instance, about 20 entire renal glomeruli may be digitized overnight with SEM-STEM for optimal diagnostic accuracy ^1, 3^. Future innovation in SEM- and TEM technology will allow faster imaging in conventional EM systems that may equally profit from LDS methodology ^35, 36^.

In sum, we have presented a detailed workflow to prepare LDS carrying ultrathin sections free of limiting artifacts and ready for routine application of nanotomy. Perspectively, the use of LDS will greatly facilitate quantifying EM and consolidate a novel mode of expert consultation on EM data, based on comprehensive, digitized information. Smooth access to EM in multiple fields of cell biology and pathology becomes available. Online repositories will serve to share the respective information for data mining, translational approaches, and teaching. Technological innovation will make SEM- and TEM systems faster to further improve their routine application of LDS in the future.

## Online methods

### Fixation and embedding of tissue samples

Diagnostic muscle, nerve and kidney samples were used. Tissues were dissected (2 x 2 mm blocks) and fixed in 2.5% glutaraldehyde buffered with 0.1 M Na-cacodylate, at 4°C overnight, then washed 3 x 10 min in 0.1 M cacodylate buffer and osmicated with 1% OsO4 in 0.05 M Na-cacodylate (muscle, nerve; overnight) or 0.1 M phosphate buffer (kidney; 30 min) at room temperature. Samples were then washed again in buffer and dehydrated in a graded acetone series in an EMP-5160 automated tissue processor (RMC). The 70% step was used for *en bloc* staining with 1% uranyl acetate and 0.1% phosphotungstic acid, dissolved in 70% acetone (60 min). For resin embedding, tissue blocks were transferred to a 1:1 mixture of 100% acetone and Renlam resin (Serva, containing accelerator), left in a fume hood overnight with the cap open for evaporation of acetone, then embedded in fresh Renlam (4 h). Renlam was also used to prepare the embedding molds; in the molds, tissue blocks were oriented and polymerized at 60°C for 72 h. Kidney blocks were dehydrated in a graded methanol series, infiltrated with Epon resin using propylene oxide as an intermedium, and polymerized.

Mice and rats were anesthetized (Nembutal) and perfusion-fixed via the abdominal aorta using 2% glutaraldehyde buffered in 0.1 M sodium cacodylate buffer, facultatively added with 1% hydroxyethyl starch. Kidney or brain tissue blocks were post-fixed overnight at 4°C, rinsed in 0.1 M sodium cacodylate buffer, osmicated for 2 h, rinsed, dehydrated in a graded ethanol series, infiltrated in Epon/propylene oxide, and embedded in Epon. Alternatively, the Renlam protocol was used (v.s.).

### Semithin sections for light microscopy

Semithin sections (300 to 500 nm) were cut with an ultramicrotome (Ultracut E, Reichert-Jung) and a histo-diamond knife (Diatome), transferred to and stretched on a microscope slide (120°C), dried (80°C), stabilized (120°C, 15 min), and stained with Richardson‘s solution (1% azure II in distilled water mixed 1:1 with 1% methylene blue plus 1% Na-borate in distilled water; 80°C, 1 min).

### Preparation of support films for LDS

Support films were prepared by slide stripping as previously described ^37^, with several adaptations to reduce artifacts and provide adequate quality for digitization of entire ultrathin sections (Fig. S1). Restrictions from mesh grids with obscuring grid bars were avoided by the use of filmed slot grids for unrestricted examination of entire sections prepared as LDS. Briefly, copper or nickel slot grids were ultrasound-cleaned in consecutive steps using acetone, pure ethanol, and distilled water (each 3 times for 2 min). Conventional microscope glass slides were cleaned with warm water and dishwashing detergent, rinsed with 70% denatured ethanol, dipped in pure ethanol, and dried with a Kimtech wipe. The dried slides were coated on their top sides with dry curd soap to ensure later detachment of the pioloform film. The curd soap was evenly distributed by intense rubbing in longitudinal and transverse directions with cotton wool for about 3 min. Individual slides were then dipped into a filtered 0.7% pioloform solution in chloroform (100 ml in a larger cylinder) with a clamp, lifted smoothly, and held above the fluid for 15 s. The latter time span was facultatively adjusted to prepare a medium to heavy silver film interference color during floating. The slide was then removed and air-dried in dust-free atmosphere (3 min). After drying, the edges of the slide were gently scratched using a stiff razor blade to improve detachment. The pioloform film was then floated onto a water trough and slot grids with standard size (2 x 1 mm aperture) and extra-large slot grids (2 x 1.5 mm aperture) were then placed on the film with their shiny side down. A parafilm stripe was lowered onto the grids and removed with the grids attached. Grids were dried in a glass petri dish (2 d) to stabilize the film. Prior to cutting sections, grids were removed from parafilm and placed on a new sheet of parafilm, the shiny side up, and hydrophilized by glow-discharging in a MED020 sputter coater within the visible plasma zone for 10 s using argon gas at 1.1 to 1.3×10 ^−1^ mbar and 6.6 to 6.9 mA. The hydrophilized grids were used within 2 to 6 h for collection of sections.

### Ultramicrotomy

For 2D EM, ultrathin sections (60 to 70 nm), sized up to ~ 2 x 1.5 mm, were cut with an Ultracut E microtome (Reichert-Jung) or PowerTome (RMC) using an ultra 35° diamond knife for minimal compression of sections ^38^ (Diatome) and stretched using xylene vapor. The grids were picked at their short edge with a fine forceps, dipped into absolute ethanol, then 10 times into distilled water to achieve smoothening of the pioloform film, and then inserted into the water trough of the diamond knife (Fig. S3). Sections were placed on the pioloform film by attachment of a section at the water-grid borderline and gently removing the grid from the water. Subsequent drying was controlled and documented via the stereomicroscope. For electron tomography, ribbons of multiple semithin sections (200-350 nm) were cut using an ultrasemi diamond knife (Diatome), stretched, and collected on grids. For optimal ribbon formation, the resin block was trimmed with a 20° diamond trim tool (Diatome) to provide parallel edges ^39^. To release ribbons from the knife, section thickness was reduced to 5 nm for 1 to 2 cutting movements. For SEM-BSD imaging, ultrathin sections were placed on freshly glow-discharged silicon wafers as substrates ^11^.

### Contrasting for LDS

Prior to heavy metal staining, grids carrying the ultrathin sections were incubated in 1% aqueous EDTA solution (4 min) to reduce formation of embedding pepper ^21^, then stained with 5% aqueous uranyl acetate (8 min) and Reynolds’ lead citrate (3 min). Between these steps, grids were washed with distilled water by moving them up and down gently for more than 20 times consecutively in three 25 ml glass beakers. Petri dishes for lead citrate staining were kept in carbon dioxide-reduced atmosphere by placing NaOH pellets next to the grids to prevent the formation of spherical precipitates during incubation. Solutions were filtered prior to use (0.22 μm filter; Millipore) and drops pipetted on parafilm stripes attached to the bottom of glass Petri dishes. Grids were stained within the droplets, the section side up, and dried horizontally, held by the forceps. Grids carrying the semithin sections for electron tomography were stained with lead citrate alone (7 min); fiducial gold particles were then added by incubation on droplets for 3 min for each side, then dried with filter paper.

### TEM imaging

A TEM 906 (Zeiss), equipped with a 2k CCD camera (TRS), and a TEM Tecnai G2 (FEI), equipped with a 4k CCD camera (Eagle), were used for standard examination and large-scale digitization. Large-scale digitization with both systems was performed largely as established earlier ^4, 8, 9^. Briefly, both TEM systems provided a computer-driven stage to automatically acquire overlapping images in a serpentine pattern, digitizing selected regions of interest (ROI). With TEM 906, columns and rows were selected manually using ImageSP software for square or rectangular datasets, while the TEM Tecnai G2 offered a freehand selection tool based on an acquired overview image with FEI photomontage software, allowing more individually shaped ROIs. Beam and focus were calibrated for the respective magnification used in the automated image acquisition process, and a background image correction tool was applied to reach uniform brightness. Automated acquisition of up to about 1,000 images was accomplished with TEM 906, while the Tecnai G2 in its basic adjustment permitted acquisition of only 500 images maximally; a modified photomontage software (collaboration with Max Otten, FEI) allowed the acquisition of up to 5,000 images. Facultatively, an array of 4 x 4 images with an overlap of 60% was acquired after digitization to correct for lens distortions during subsequent data processing. Images were saved as 8- or 16-bit tif files. For electron tomography, a 2k CCD camera (Veleta) or alternatively the 4k camera were used for tilt series acquisition with the Tecnai G2 at 200 kV. Tilt series were automatically acquired at 29,000x or 50,000x magnification within a range from −60° to +60° in steps of 1-2°.

### SEM-STEM imaging

A Gemini 300 SEM (Zeiss) equipped with a scanning transmission electron microscopy (STEM) detector was used with SmartSEM software to adjust the electron beam and with Atlas 5 (Fibics) to perform automated large-scale digitization of ROIs or entire sections. The workflow of Kuipers et al. ^15^ was modified and adjusted to allow for automated pre-irradiation and imaging of up to 12 entire LDS with intermedium to high resolution, using a pixel size of 7.3 or 9 nm, and increased imaging speed. Briefly, the following steps were followed:

1. Transfer of LDS: For large-scale digitization of multiple entire sections in a high-throughput manner, up to 12 LDS with sections of different blocks were transferred into the vacuum chamber using the STEM-sample holder.
2. Quick sample check: Using 29 kV accelerating voltage, 3.3 mm working distance and SmartSEM software, all LDS were briefly checked at a magnification of 50-100x to verify adequate quality for later large-scale digitization. Usually, LDS were stored in the vacuum overnight to allow for outgassing.
3. Plasma cleaning: On the next day, a short (3 min) plasma cleaning cycle was initiated to clean the chamber.
4. Pre-irradiation: A 120 μm aperture was used for pre-irradiation at 29 kV, carried out with a retracted STEM detector. For single sections or medium-sized ROIs, the stage was adjusted manually and pre-irradiation was carried out for about 30 min (ROI of 300 x 300 μm) or 60-120 min (entire section of about 2 x 1 mm). For automated pre-irradiation, we manually prepared a macro file (see Supplementary Information for parameters) with SmartSEM, based on coordinates for each section with stage position in X, Y and Z as well as stage rotation, scan rotation and delay (in s). Scan speed and reduced window were adjusted for scan cycles of about 1 s, and de-focus of 1.5 mm was applied to blurr the image. A test run was performed with 30 s delay to check for correct position of each section and then started with 3,600 s to 7,200 s delay.
5. Overviews: Using Atlas 5, overview images of all LDS were automatically acquired at 250 nm pixel size to define the borders of the sections, followed by overview images at 100 nm pixel size for higher image quality.
6. Beam alignments: Focus and correction for astigmatism were checked at 40,000 to 60,000x, then, the beam was wobbled to center the aperture.
7. Setting parameters: For most datasets, a pixel size of 7.3 nm or 9 nm was used with a tile size of 8k to 12k, resulting in image fields of 50-90 μm. Autofocus settings were adjusted for either automated digitization of multiple sections (using a range of about 10 μm) or digitization of single sections (using a range of 4-5 μm). Usually, 300-500% pixel ratio, 1-3 μs dwell time and pixel averaging (= line averaging 1) and autostigmation with 1% allowed reliable performance. However, in case of very large areas with insufficient amount of structural details, such as lumina of kidney tubules and lung alveoles, a pixel ratio of 1,000% was used in conjunction with an increased range of about 15 μm and a minimum dwell time of 3 μs. Autofocus and autostigmation were performed on the previous tiles. Brightness and contrast of each section or ROI were adjusted manually prior to imaging using small ROIs on representative tissue areas.
8. Final adjustments: Next to the first image tile, beam alignments were checked manually using SmartSEM.
9. Large-scale digitization: All ROIs or sections for large-scale digitization were selected, manually checked for focus and adjusted for brightness and contrast using the previously defined values (step 7.).
10. Stitching and export: Image tiles were stitched and exported using Atlas 5, allowing convenient pan-and-zoom examination of the resulting datasets with the Atlas 5 browser-based viewer. Alternatively, stitching and export to bigtif files were performed using Fiji software with TrakEM2 plugin and nip 2 software (see below as well as step-by-step protocols in the Supplementary Information).

### SEM-BSD imaging

The same SEM platform that was used for SEM-STEM imaging was also used for imaging with a backscattered electron detector (SEM-BSD). We used an accelerating voltage of 4-8 kV, a working distance of 4-5 mm, a standard aperture of 30 μm for samples with high contrast, a large aperture of 60 μm for samples with low contrast and a dwell time of 3-12 μs.

### Data processing

A HP Z840 workstation was used for processing of most datasets (128 GB working storage, 2x Xeon E5-2667 with 8 cores each and 3.20 GHz, NVIDIA quadro 4000 with 8 GB), with a BenQ SW2700PT (27”) screen. The screen was calibrated using a Spyder 4 elite CCD sensor.

Detailed step-by-step protocols are provided in the Supplementary Information. Briefly, for stitching of TEM large-scale datasets, the images were renamed using bulk rename utility software (001.tif, 002.tif, etc.). The images for the ROI and the lens correction were then adjusted for brightness and contrast using Fiji, exported to 8-bit tif files and imported to TrakEM2. In case of major differences in brightness and contrast between the individual images, an image filter (“normalize local contrast”) was applied for temporary compensation. Images were then stitched by multiple consecutive alignment steps including values of the lens correction dataset. Temporary image filters were then removed and differences in brightness between the individual stitched images were facultatively permanently reduced with the “match intensities” tool. For export of small datasets, the “make flat image” tool was used to prepare a single standard tif file. For large datasets that exceed the pixel limit of standard tif files, a modified CATMAID export script by Stephan Saalfeld was used to export the ROI into multiple non-overlapping tiles with pixel dimensions of 25,000 x 25,000 pixels (see “https://github.com/axtimwalde/fiji-scripts/blob/master/TrakEM2/catmaid-export2.bsh”, last accessed 29.10.2018). These tiles were then processed to one coherent large bigtif file using nip2.

For STEM- and BSD data, a similar workflow for processing was used. A text file for import of image tiles to TrakEM2 of up to about 10 entire digitized sections (up to about 4,000 image tiles) was prepared by calculation of image pixel coordinates using an Excel file, as positions of the tiles within each dataset are implemented in the image tile names. A template of this excel file, filled with data of several datasets and a brief documentation for its usage, is provided in the Supplementary Information. An Excel macro was used to extract numbers (see “https://www.extendoffice.com/excel/1622-excel-extract-number-from-string.html?page_comment=6”, last accessed 03.11.2020). For increased data processing speed, all processing steps were performed using a solid-state-drive (SSD). To reduce the size of the TrakEM2-project, jpg-format for mipmap generation was selected (see project properties in TrakEM2). Image tiles with very low amounts of structural information at the periphery of datasets were deleted manually. Only one alignment step with adjusted image parameters was performed (usually with minimum image size of “600” and maximum image size of “1600”), without introducing temporary image filters or lens correction. In case of intensity matching, a black background of image tiles was changed to white using photoshop with batch processing (replace color command, used with “0” tolerance to ensure only a switch of the artificially black background to white). For bigtif export, non-overlapping tif tiles were prepared as for TEM-datasets, however, up to 10 different macros were run in parallel for automated export for several h or overnight. Bigtif files allowed examination and in-depth analysis using QuPath open-source software ^18^.

For 3D reconstruction of tilt images using fiducial gold particles, IMOD software package including etomo was used ^40^.

## Supporting information

Excel Template

Zoom-in Video

## Acknowledgements

The authors wish to thank Matthias Ochs for critical discussion of the work; Sara Timm, Petra Schrade, Cordula zum Bruch, Carola Geiler, Agnieszka Münster-Wandowski, and Imre Vida (all: Charité) and Christoph Fahrenson (ZELMI, TU Berlin) for expert help in sample preparation and imaging; David Mastronarde (CU Boulder) for valuable help in data processing with IMOD. We thank the Core Facility for Electron Microscopy of the Charité for support in data acquisition. This work was supported by the Charité foundation (C.D.; Max Rubner award 2016); Charité CRC 1365 (S.B.; S01 and C04); the Zeiss Gemini 300 SEM was acquired by Charité and DFG 596-1 FUGG, §91 Program (S.B. and consortium).

## Author contributions

C.D. developed the improved preparation and carried out the experiments including section preparation, support film preparation, imaging and data processing as well as wrote the manuscript. S.B. performed design of the study, wrote and edited the manuscript. H.H.G., F.L.H. and W.S. edited the manuscript.

## Data and materials availability

All data are available in the main text and the Supplementary Information, or are available upon request. Selected large-scale datasets are provided for open access pan-and-zoom analysis via www.nanotomy.org.

## Ethics statement

For this study, archive resin blocks from diagnostic muscle, nerve and kidney samples as well as experimental rodent tissues, embedded according to standard protocols, were used. The study was approved by the local ethics committee of the Charité - Universitätsmedizin Berlin (EA2/107/14) and done in accordance with the declaration of Helsinki. Following routine clinical diagnostics, samples from patients were anonymized and thereafter included in the study. Animal experiments were approved by the German Animal Welfare Regulation Authorities on the protection of animals used for scientific purpose (Berlin Senate, G0148/18, O0298-17).

## Additional information

A list of Supplementary Information is provided in a separate pdf.

## Graphical abstract

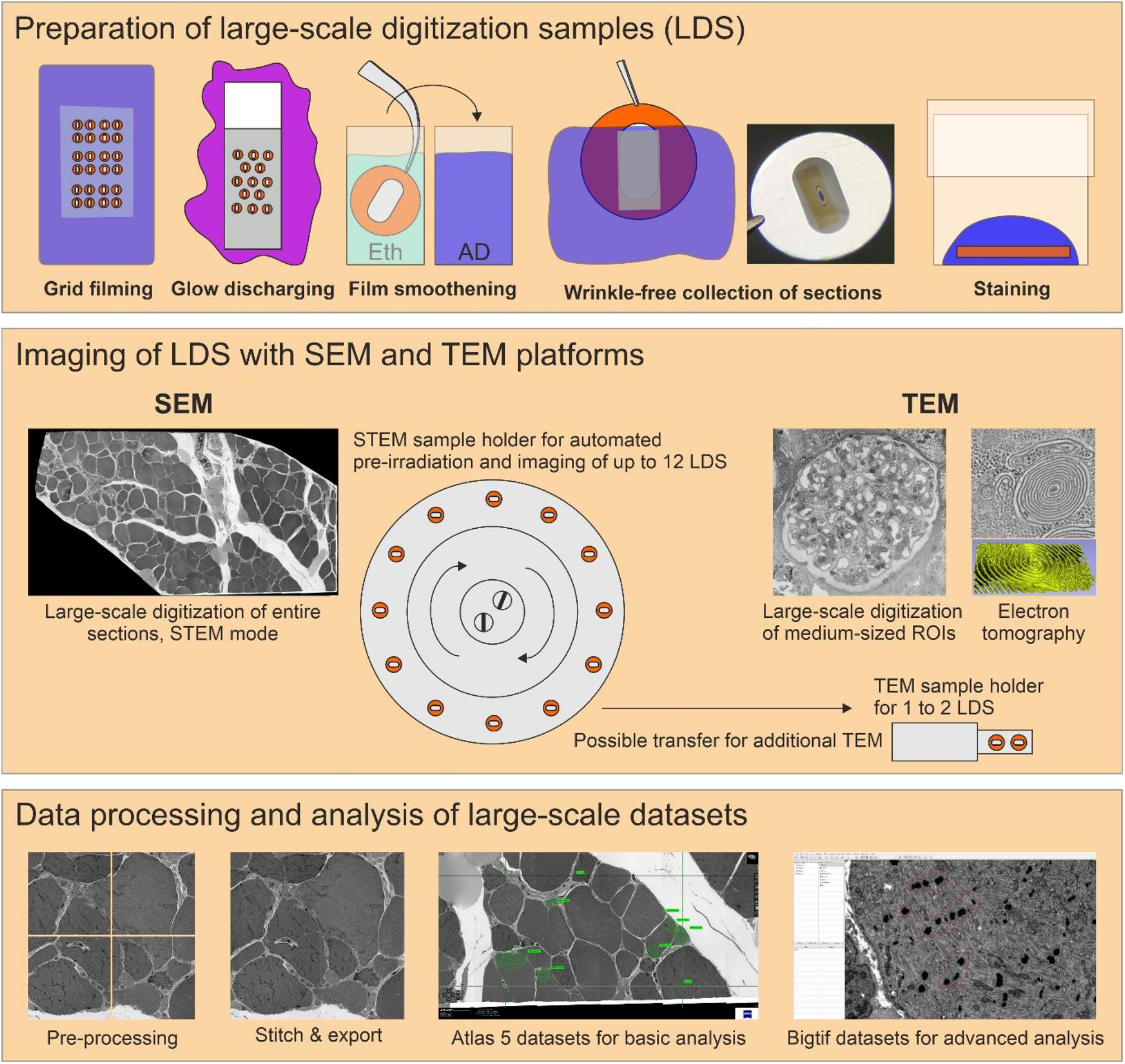

Large-scale digitization samples (LDS) with virtual absence of limiting artifacts are prepared for advanced data acquisition. Scanning electron microscope (SEM) and transmission electron microscope (TEM) systems are compared for efficient imaging. A refined protocol is required to generate support films with reduced artifacts. Reliable, wrinkle-free collection of sections is achieved by glow discharging and smoothening of the film under control of Newton ring formation. Contrasting is optimized by a smooth section surface and refinements of the staining protocol. An SEM operated in transmission mode (STEM) provides automation to reduce operator involvement. This enables efficient recording of large ROIs up to entire sections in high-throughput. Comparingly, TEM imaging is restricted to small- or medium-sized ROIs, but appropriate for additional electron tomography, high-end resolution, or conventional imaging. Processing of data includes stitching of image tiles to mosaics. Resulting large datasets comprise the context between overview and highly resolved detail. Export of data to bigtif files facilitates an improved analysis by annotations or quantification.

