## Supplementary Information for "Preparation of large-scale digitization samples for automated electron microscopy of tissue and cell ultrastructure"

Supplementary Figures 1-5 with legends and Supplementary Tables 1-4

Joined pdf files of step-by-step protocols for data processing:

Data processing steps for 1) Stitching and lens correction of overlapping TEM image tiles to a coherent dataset with TrakEM2, including export to non-overlapping tif tiles as a basis for bigtif file generation; 2A) Semiautomated generation of a text file containing coordinates (X,Y and Z) of overlapping STEM image tiles from two large datasets for import into TrakEM2 [1] using Excel; 2B) import and stitching of these tiles to coherent datasets with TrakEM2; 2C) export to non-overlapping tiles; 2D) bigtif generation using nip2; 2E) import in QuPath as a basis for in-depth analysis using measurements and annotations

Screenshot of the protocol for automated pre-irradiation (electron beam shower) for 12 grids using SmartSEM Software

Separate files

ExcelTemplate: Excel file for calculation of image tile coordinates, filled with exemplary data and a brief documentation how to use the file

Pan_and_zoom_video_Hippocampus: A zoom-in video to the large-scale dataset of the dentate gyrus, prepared with SEM-STEM

**Online repository datasets**: Selected large-scale datasets, as indicated in Supplementary Table 2, are accessible for online pan-and-zoom analysis via www.nanotomy.org

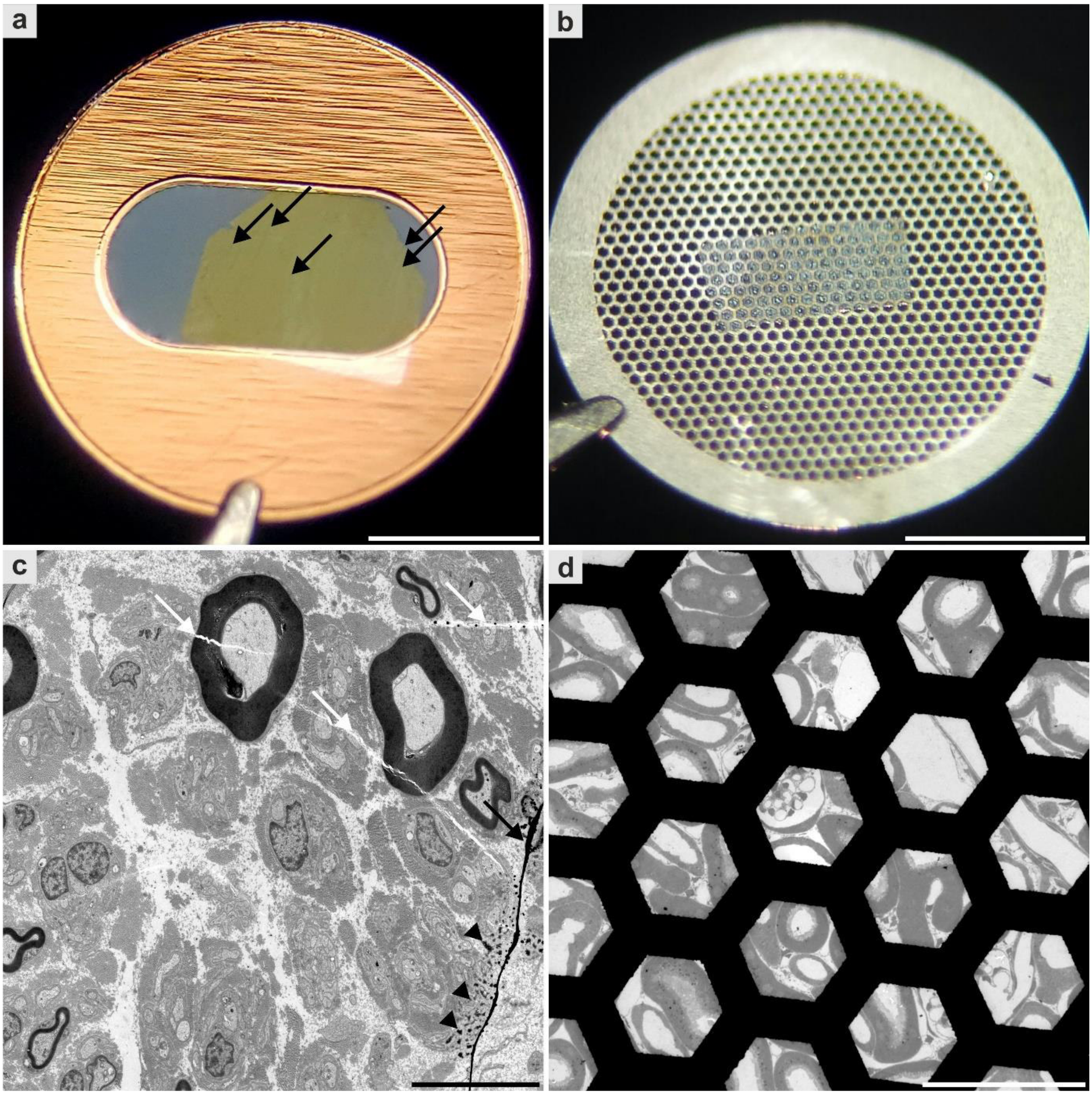


**Fig. S1. Common limitations of slot and mesh grids**. **a,c**, Slot grid provide a large observable area, allowing entire ultrathin sections to be viewed without limiting mesh bars. However, the delicate film is vulnerable to film imperfections, wrinkles (black arrows), stain precipitates (arrowheads in **c**), other contamination and section cracks (white arrows in **c**). **b,d**, The use of mesh grids on the other hand, allows preparation without additional film artifacts. **d**, This often results in cleaner samples, but viewing is impaired by mesh bars as well as decreased section stability. **c**, Biopsy of a peripheral nerve; hereditary neuropathy. **d**, Perfusion-fixed mouse kidney. Scale bars, 1 mm (**a,b**), 10 µm (**c**), 100 µm (**d**).

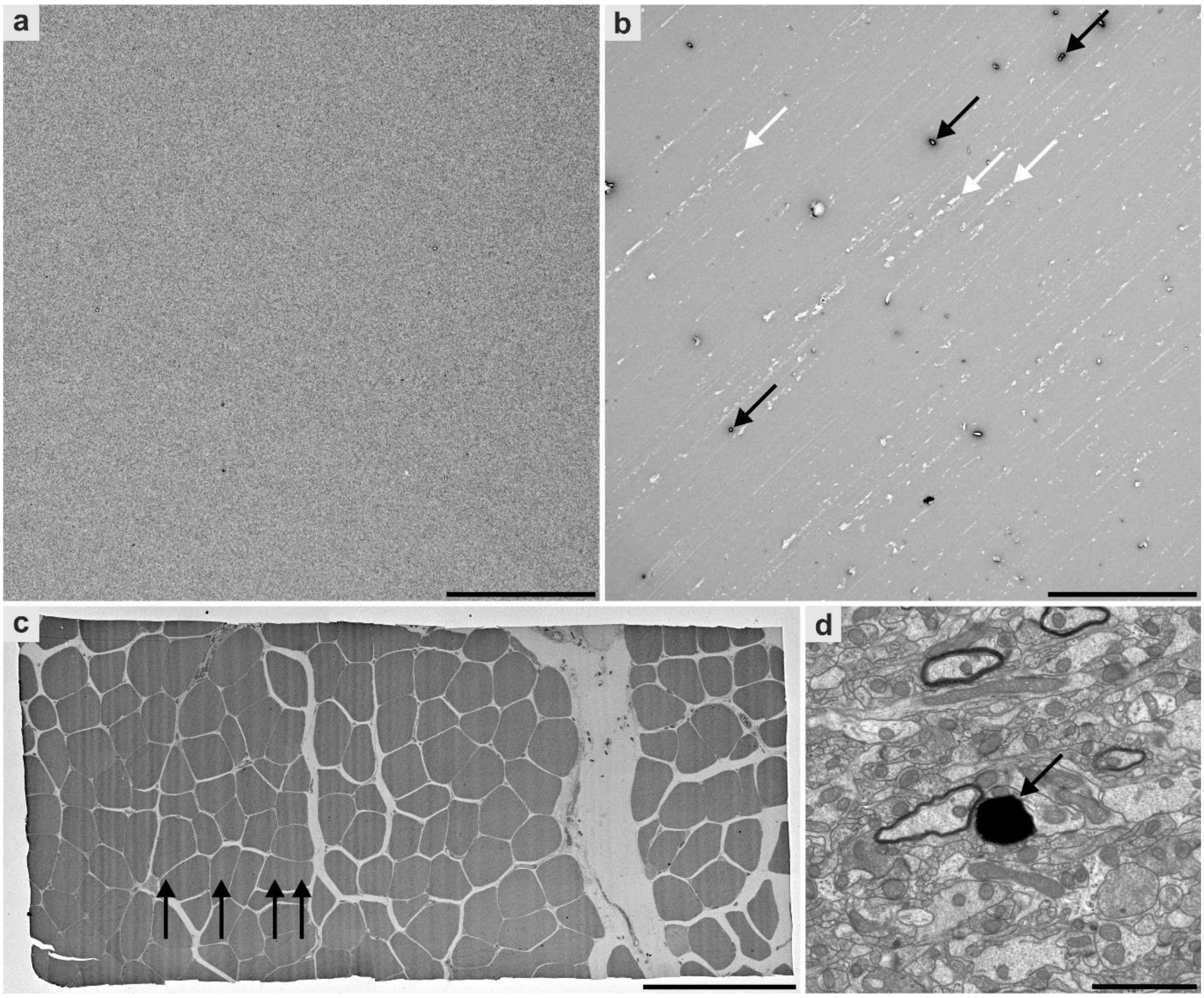


**Fig. S2. Different artifacts of ultrathin sections from LDS**. Images acquired by transmission electron microscopy. **a**, A good pioloform film shows almost no flaws as compared to a pioloform film in **b** with numerous imperfections. **b**, Water contamination leads to holes (black arrows), and improper smoothening of the curd soap causes smear artifacts (white arrows). **c**, Chatter (arrows) became easily visible at low magnification overview images such as this overview of an entire section. Here, cutting was performed on the 5th floor of a building, resulting in chatter due to low frequency vibrations. **d**, Contamination (arrow) likely due to the glow discharge procedure in a MED020 sputter coater that is used for carbon coating as well. **c**, Diagnostic muscle biopsy. **d**, Rodent brain, fixed by perfusion fixation. Scale bars, 50 µm (**a,b**), 200 µm (**c**), 2 µm (**d**).

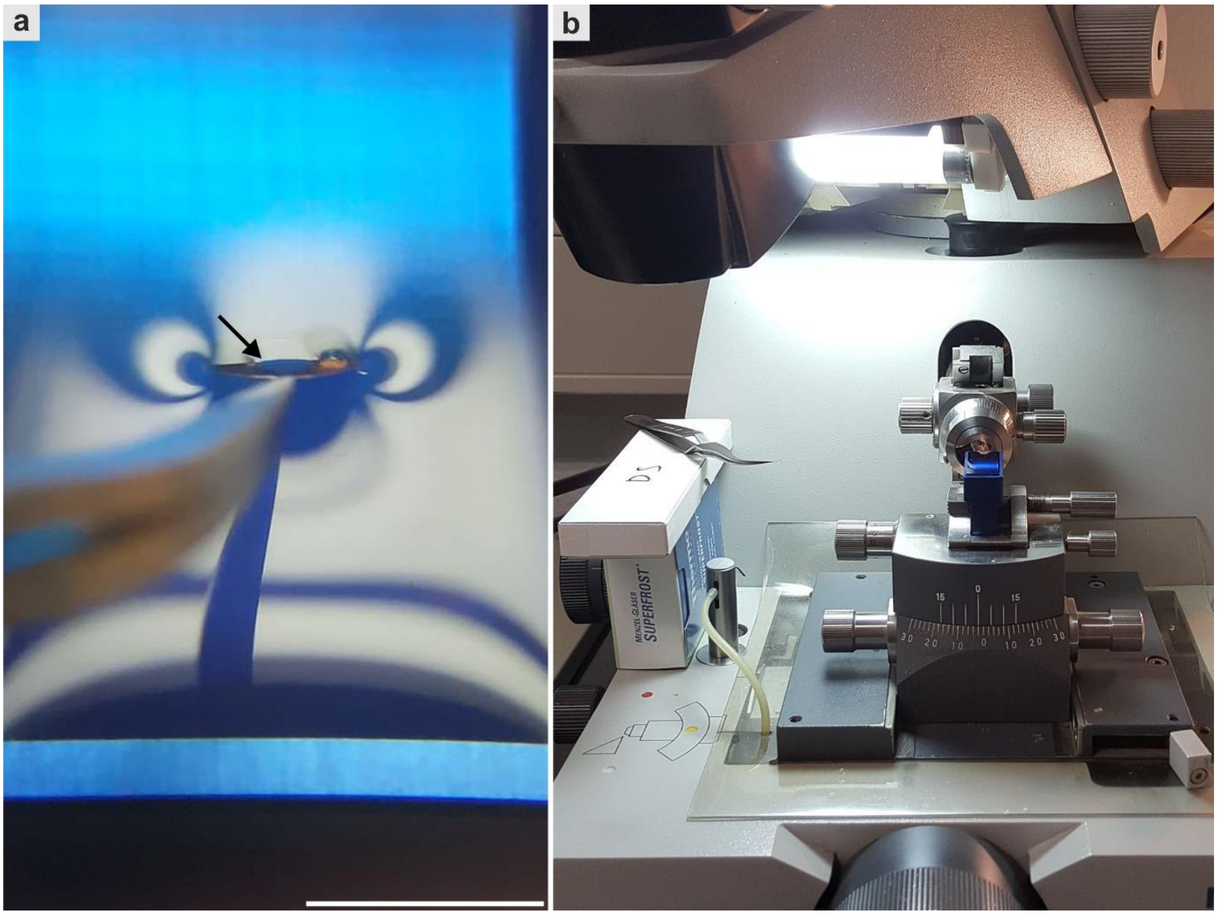


**Fig. S3. Preparing LDS; collection and drying of a section using a glow-discharged and ethanol-smoothened pioloform-coated slot grid**. **a**, The grid is inserted at the "6 o'clock position" to follow the section during the collection process. Ideal hydrophilicity of grids is indicated by a sharp and smooth water-grid borderline (arrow) which facilitates attachment of a section at the borderline and allows to collect the section by raising the grid out of the water. Note that the borderline is crossing the pioloform film; alternatively, the borderline may be placed on the metal surface directly underneath the forceps to compensate for mildly insufficient hydrophilicity (see Supplementary Table 3 for troubleshooting). **b**, After collection of a section, the self-closing forceps is placed next to the diamond knife, allowing examination of the attachment and drying processes. Scale bar, 5 mm (**a**).

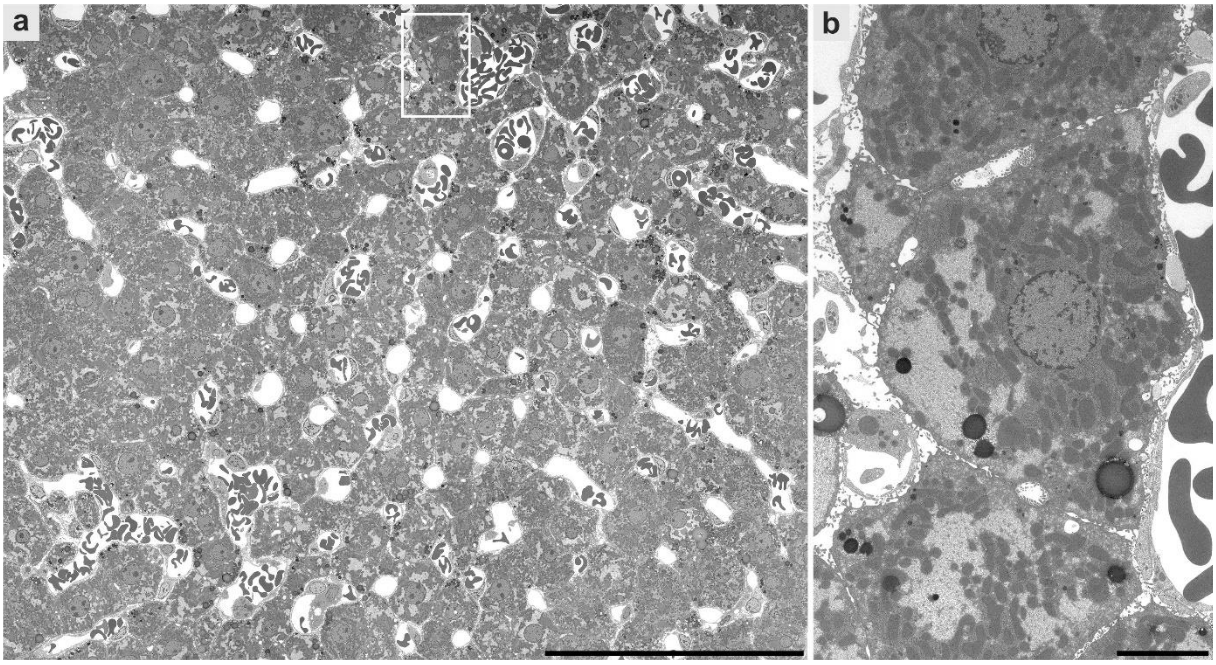


**Fig. S4. Large-scale digitization of a medium-sized region of interest within a section of rodent liver using transmission electron microscopy**. An intermediate magnification of 4646x was used, resulting in a pixel size of 9.3 nm. **a**, Microanatomical overview of parenchyma. **b**, Digitally magnified region from **a** shows hepatocytes with well-resolved glycogen, mitochondria, lysosomes, perisinusoidal space and intercellular bile ducts. Scale bars, 100 µm (**a**), 5 µm (**b**). See also www.nanotomy.org for download of the full resolution dataset.

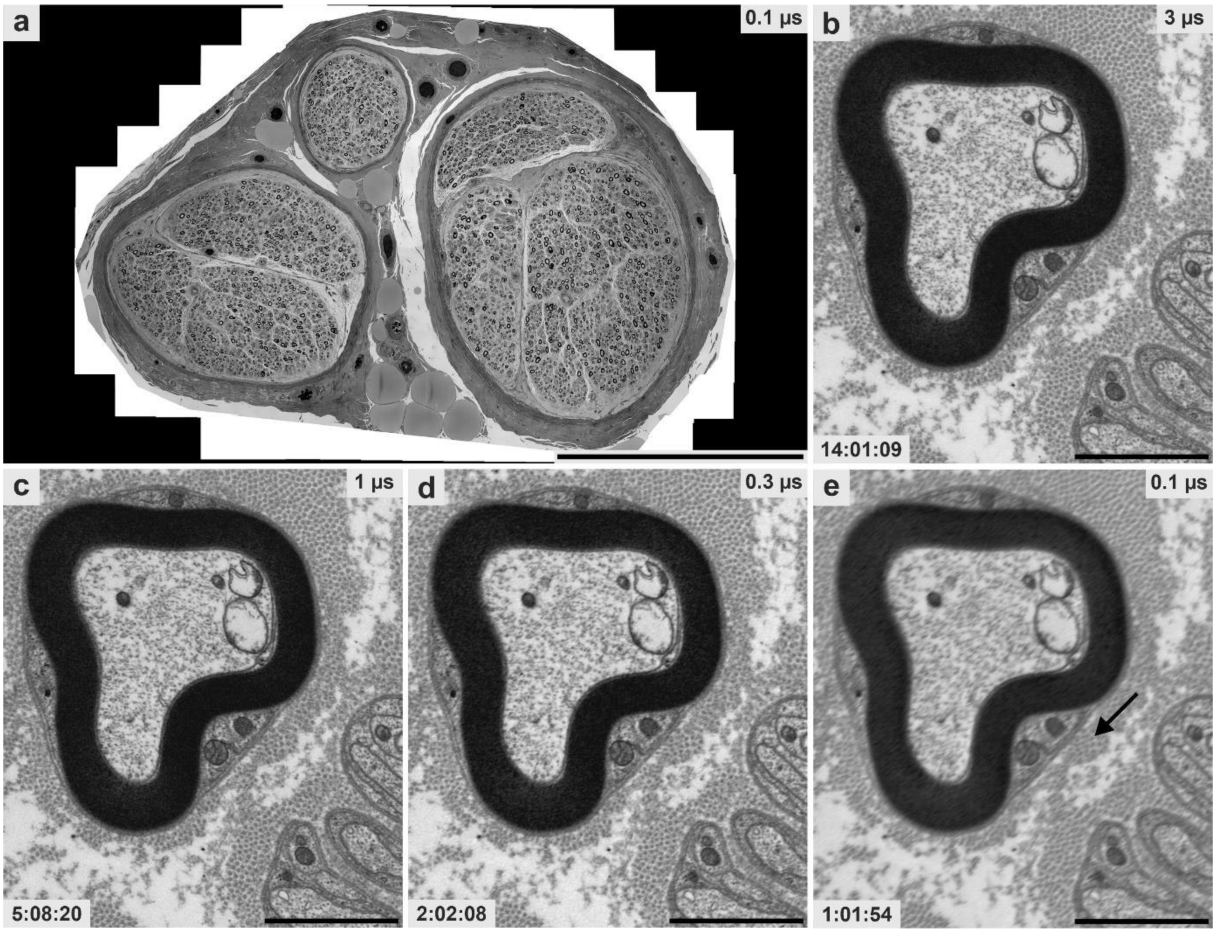


**Fig. S5. Large-scale digitization of a diagnostic nerve sample using scanning transmission electron microscopy.** **a**, An entire ultrathin section was digitized at 9 nm pixel size using different dwell times to analyze imaging time and quality. **b** to **d**, Image quality only moderately decreases with increased imaging speed using 3 µs, 1 µs and 0.3 µs dwell time, while imaging time is drastically decreasing from 14 to 2 h. Quality in **d** allows to clearly resolve diagnostically relevant structures such as collagen pockets (see step-by-step protocol 2E; QuPath analysis). **e**, At 0.1 µs dwell time, still individual collagen fibrils are resolved (arrow), but the image quality clearly appears blurred. Scale bars, 500 µm (**a**), 2 µm (**b-e**). See also www.nanotomy.org for internet browser-based pan-and-zoom analysis of the full resolution datasets of **c,d** and **e**.

Supplementary Table 1. Information on tissue blocks illustrated in this work*

| Fig. | Tissue | Pathology | Species | Resin | Osmication | UA en bloc staining |
|---|---|---|---|---|---|---|
| 2,6 | Kidney | Chronic hypokalemic nephropathy | Human | Epon | 30 min 1% in 0.1 M phosphate buffer | - |
| 3 | Kidney | Lupus nephritis | Human | Epon | 30 min 1% in 0.1 M phosphate buffer | - |
| 4 | Brain | Alzheimer's disease | Mouse | Renlam | Overnight 1% in 0.5 M cacodylate buffer | + |
| ga,5 | Muscle | Tubular aggregates | Human | Renlam | Overnight 1% in 0.5 M cacodylate buffer | + |
| S4 | Liver | - | Rat | Epon | 2 h 2% in distilled water | - |
| S5 | Nerve | Neuropathy | Human | Renlam | Overnight 1% in 0.5 M cacodylate buffer | + |

*In total, we prepared ultrathin sections of 100 different tissue blocks from different research projects to establish the LDS protocol. However, early preparations were less sophisticated and demonstrated higher amounts of preparation flaws so that more sections were needed to prepare one high-quality section [2, 3]. ga, graphical abstract; UA uranyl acetate.

Supplementary Table 2. Information on large-scale datasets illustrated in this work*

| Fig. | Imaging system | Pixel size | Dwell time | Tile size, p | Mosaic size | Tile number | Acquisition time | Repository number |
|---|---|---|---|---|---|---|---|---|
| 2a,d | TEM | 7.3 nm | - | 2,048 | 254 x 254 µm | 400 | 1:06:40 | 2 |
| 2b,e | SEM-STEM | 7.3 nm | 3 µs | 12,288 | 245 x 229 µm | 9 | 01:09:37 | 3 |
| 2c,f | SEM-BSD | 7.3 nm | 3 µs | 12,288 | 269 x 269 µm | 16 | 01:35:32 | 4 |
| 3a-d | SEM-STEM | 7.3 nm | 1 µs | 8,192 | 239 x 239 µm | 16 | 00:24:51 | 5 |
| 4a-c | SEM-STEM | 7.3 nm | 1 µs | 8,192 | 1,794 x 1,017 µm | 510 | 12:32:09 | 6 |
| 5a** | SEM-STEM | 9 nm | 1 µs | 10,240 | 1,383 x 760 µm | 127 | 04:40:54 | 1 |
| 5b | SEM-STEM | 100 nm | 1 µs | 2,048 | 1,236 x 432 µm | 21 | 00:02:25 | |
| 5c-f | SEM-STEM | 4 nm | 3 µs*** | 24,576 | 94 x 49 µm | 1 | 00:50:03**** | 7 |
| 6a | SEM-STEM | 7.3 nm | 0.3 µs | 12,288 | 245 x 229 µm | 9 | 00:09:18 | 9 |
| 6b | SEM-STEM | 7.3 nm | 0.2 µs | 20,480 | 245 x 229 µm | 9 | 00:07:07 | 10 |
| 6c | SEM-STEM | 7.3 nm | 0.1 µs | 12,288 | 245 x 229 µm | 9 | 00:03:21 | 11 |
| 6d | SEM-BSD | 7.3 nm | 3.0 µs | | | | | 12 |
| 6e | SEM-BSD | 7.3 nm | 1.0 µs | | | | | 13 |
| 6f | SEM-BSD | 7.3 nm | 0.1 µs | | | | | 14 |
| S4a,b | TEM | 9.3 nm | - | 2,048 | 380 x 240 µm | 360 | 01:00:00 | 8 |
| S5a,e | SEM-STEM | 9 nm | 0.1 µs | 10,240 | 1,347 x 917 µm | 155 | 01:01:54 | 15 |
| S5b | SEM-STEM | 9 nm | 3.0 µs | 10,240 | 1,347 x 917 µm | 155 | 14:01:09 | |
| S5c | SEM-STEM | 9 nm | 1.0 µs | 10,240 | 1,347 x 917 µm | 155 | 05:08:20 | 17 |
| S5d | SEM-STEM | 9 nm | 0.3 µs | 10,240 | 1,347 x 917 µm | 155 | 02:02:08 | 16 |

*In total, we digitized large areas (entire ultrathin sections) from 92 different samples; 85 were diagnostic (76x skeletal muscle, 4x brain, 2x lung, 2x tumor, 1x nerve) and 10 experimental samples (5x kidney, 1x liver, 2x brain, 1x lung, 1x cell culture). Medium sized ROIs came from 5 different blocks of diagnostic samples (5x kidney). ** Same dataset as in graphical abstract; *** including line averaging of 3 x 3 µs; **** for each of the 6 regions of interest. p, pixels.

Supplementary Table 3. Critical steps in preparation of large-scale digitization samples (troubleshooting)

| Category | Artifact pattern | How to avoid them |
|---|---|---|
| Support film and grids | | |
| Cleaning | Large spots on pioloform film | Cleaning sequence of first using acetone, then ethanol, then distilled water must be respected |
| Curd soap coating | Smears on pioloform film | Ensure homogeneous distribution of curd soap |
| Pioloform | Small dots and holes of pioloform film | Open the bottle with pioloform solution warmed to room temperature to avoid water contamination |
| Hydrophilization | | |
| Glow discharging | Pioloform film destroyed in vacuum | Transfer grids to a separate parafilm strip to allow air circulation, thus avoiding damage of the pioloform film |
| Glow discharging | Overhydration; no sharp water-pioloform borderline during collection of sections<br>Underhydration; sharp water-pioloform borderline, but wrinkles appear | Grids should only be moderately hydrophilic; during collection of sections from the knife's water trough, the water-grid borderline has to be sharp<br>When overhydrated, reduce glow discharging time (Fig. S3)<br>Conversely, sections may be attached on the metal surface next to the forceps to stabilize water within the slot area in case of underhydration |
| Glow discharging | Electron-dense contaminations on pioloform film | Avoid e.g. carbon contamination in the sputter coater, if it is also used for carbon coating (Fig. S2) |
| Glow discharging | Underhydration | Ideally, use glow-discharged grids within 1 to 2 h |
| Grid storage | Inhomogeneous wetting or wrinkles | Let freshly filmed grids dry for 2 d to ensure homogeneous wetting when collecting the sections and avoid use of grids older than 2 to 3 months to ensure adequate tension of the film |
| Ultramicrotomy | | |
| Water borderline | Overhydration of pioloform film (no sharp borderline) | Place the forceps as far as possible at the grid periphery<br>After ethanol smoothening, insert grid only to 1/3 to 1/2 into the water of the knife's trough and wait for 10 s to generate a sharp and stable water-grid borderline<br>Do not fully submerge the grid when inserting it into the water trough |
| Water borderline | Underhydration of pioloform film | Prepare another batch of glow-discharged grids; do not perform additional glow discharging of the same grids since this may result in overhydration |
| Drying process | No or displaced Newton ring formation, wrinkles, section movement | Ensure a horizontal position of the grids held within forceps<br>Avoid or remove compression or folds in the sections by adjusting sectioning parameters and using xylene or chloroform vapor to ensure smoothness<br>Prepare straight block edges to fix the section onto the water-grid borderline during collection |
| Staining | | |
| Embedding pepper | Electron dense contamination | Treat grids with 1% EDTA in distilled water prior to staining |
| Precipitates | Needle-like (uranyl acetate) or spherical (lead citrate) contamination | Use aqueous instead of ethanolic uranyl acetate solution<br>Place NaOH pellets next to lead citrate in closed Petri dish |
| Other contamination | Amorphous electron-dense contamination | Rinse intensely after staining with uranyl acetate to avoid excess contamination and let the sections dry in a horizontal position for adequate Newton ring formation |
| Detachment artifacts | Small or medium sized, ring- or comma-shaped wrinkles, probably due to water-induced swelling of the sections<br>Some areas, e.g. strongly fixed structures or areas with pure resin, e.g. cell culture with low cell density, seem to be prone to these artifacts | Trim block areas of pure resin or overfixed areas away<br>Ensure adequate drying of the sections after collection at room temperature for 48 h<br>Stabilize sections, fixed individually by forceps, at 120°C in an oven for 15-30 min prior to staining |

Supplementary Table 4. Comparison of large-scale digitization using TEM, STEM and BSD

<table>
<tr><th></th><th>TEM</th><th>SEM-STEM</th><th>SEM-BSD</th></tr>
<tr><td>Substrate</td><td>Slot grid</td><td>Slot grid</td><td>Silicon wafer</td></tr>
<tr><td>Imaging speed</td><td>Fixed, about 3 µs*</td><td>Variable, about 1 µs</td><td>Variable, 6 to 12 µs</td></tr>
<tr><td>Image tiles (field dimensions)</td><td>2,048 or 4,096 pixels per side</td><td colspan="2">Variable, up to 32,768 pixels per side</td></tr>
<tr><td>Field size</td><td>15 µm or 30 µm (at 7.3 nm pixel size)</td><td colspan="2">Variable, 50 to 100 µm</td></tr>
<tr><td>Autofocus</td><td>Not available with our systems</td><td colspan="2">Autofocus/ autostigmation in center of image tiles</td></tr>
<tr><td>Stitching</td><td>TrakEM2</td><td colspan="2">Atlas 5 or TrakEM2</td></tr>
<tr><td>Zoomable dataset</td><td>bigtif</td><td colspan="2">Atlas 5 or bigtif</td></tr>
<tr><td>Limitations</td><td>Fixed imaging speed, Small-size image tiles**), pre-irradiation recommended</td><td>Beam damage (support film required)<br>Pre-irradiation required</td><td>Slow imaging speed<br>Special preparation for improved results<br>Pre-irradiation sometimes necessary</td></tr>
<tr><td></td><td colspan="2">Vulnerable sections</td><td></td></tr>
<tr><td>Advantages</td><td>Good resolution and SNR, compatible with many old or conventional TEM systems</td><td>Good resolution and SNR, high imaging speed. Up to 12 grids in the sample holder</td><td>Stable preparation</td></tr>
<tr><td></td><td></td><td colspan="2">Choice of adjustable imaging parameters such as dwell time, tile and pixel size, averaging, autofocus and autostigmator. Low-grade lens distortions, high degree of automation.</td></tr>
<tr><td></td><td colspan="2">Grids interchangeable (TEM, SEM-STEM; shuttle workflow)</td><td></td></tr>
</table>

* µs per pixel, 2k CCD camera. TEM, transmission electron microscopy; SEM, scanning electron microscope; STEM, scanning transmission electron microscopy; BSD, backscattered electron detection.

**1)** Import of region of interest (ROI) and lens correction (LC)

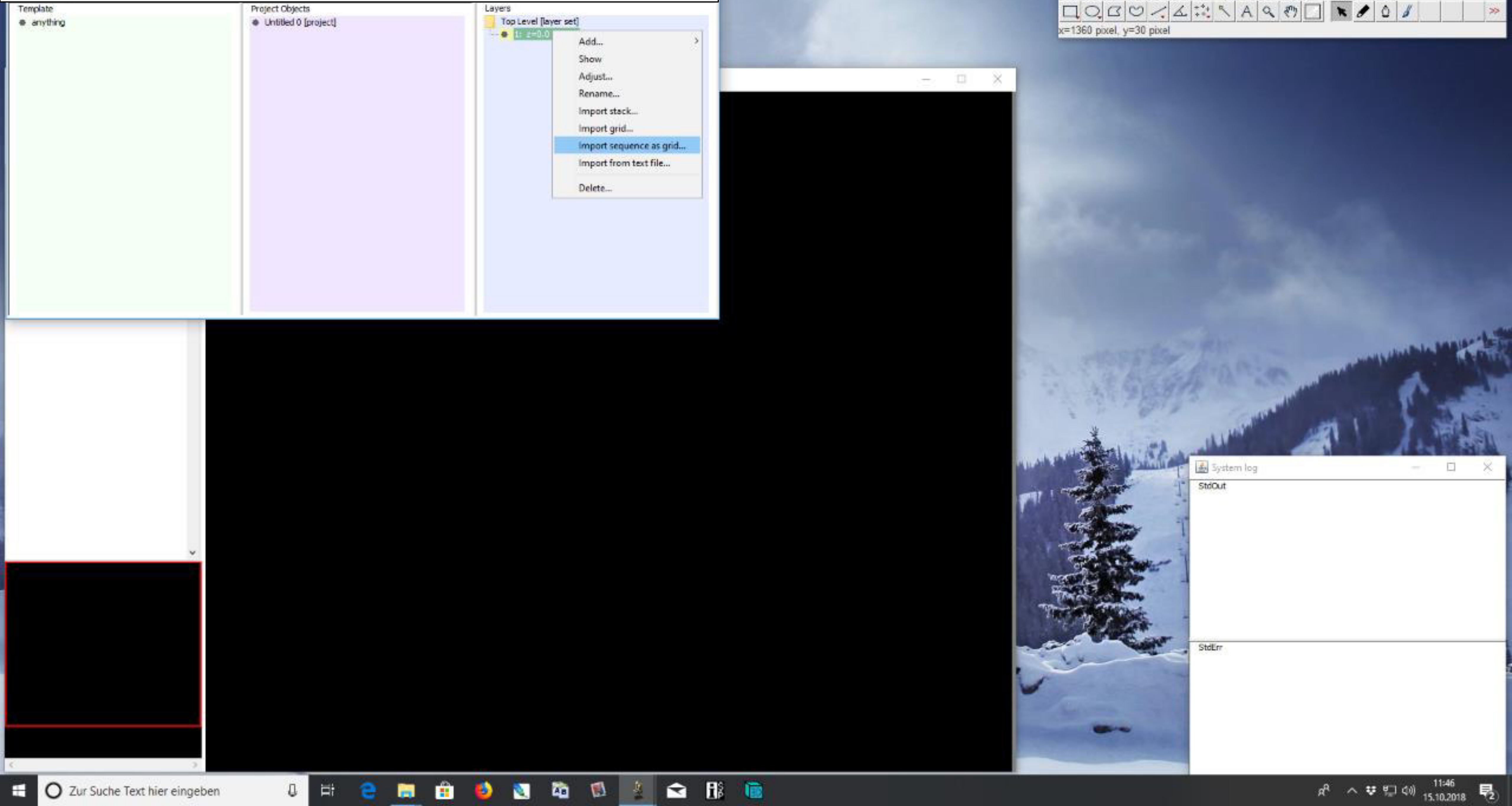

### 1) Import of ROI and LC

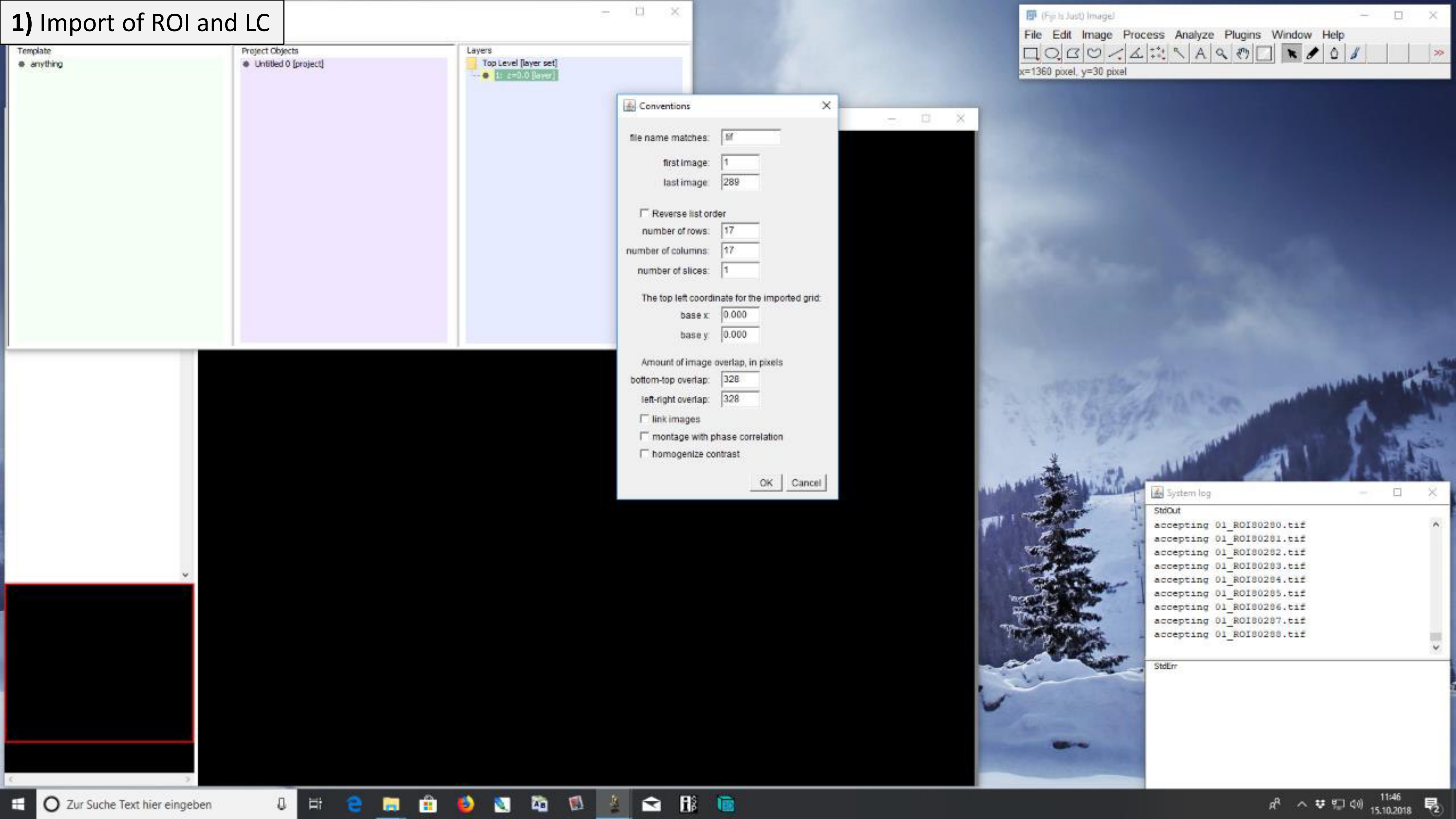

### **1)** Import of ROI and LC

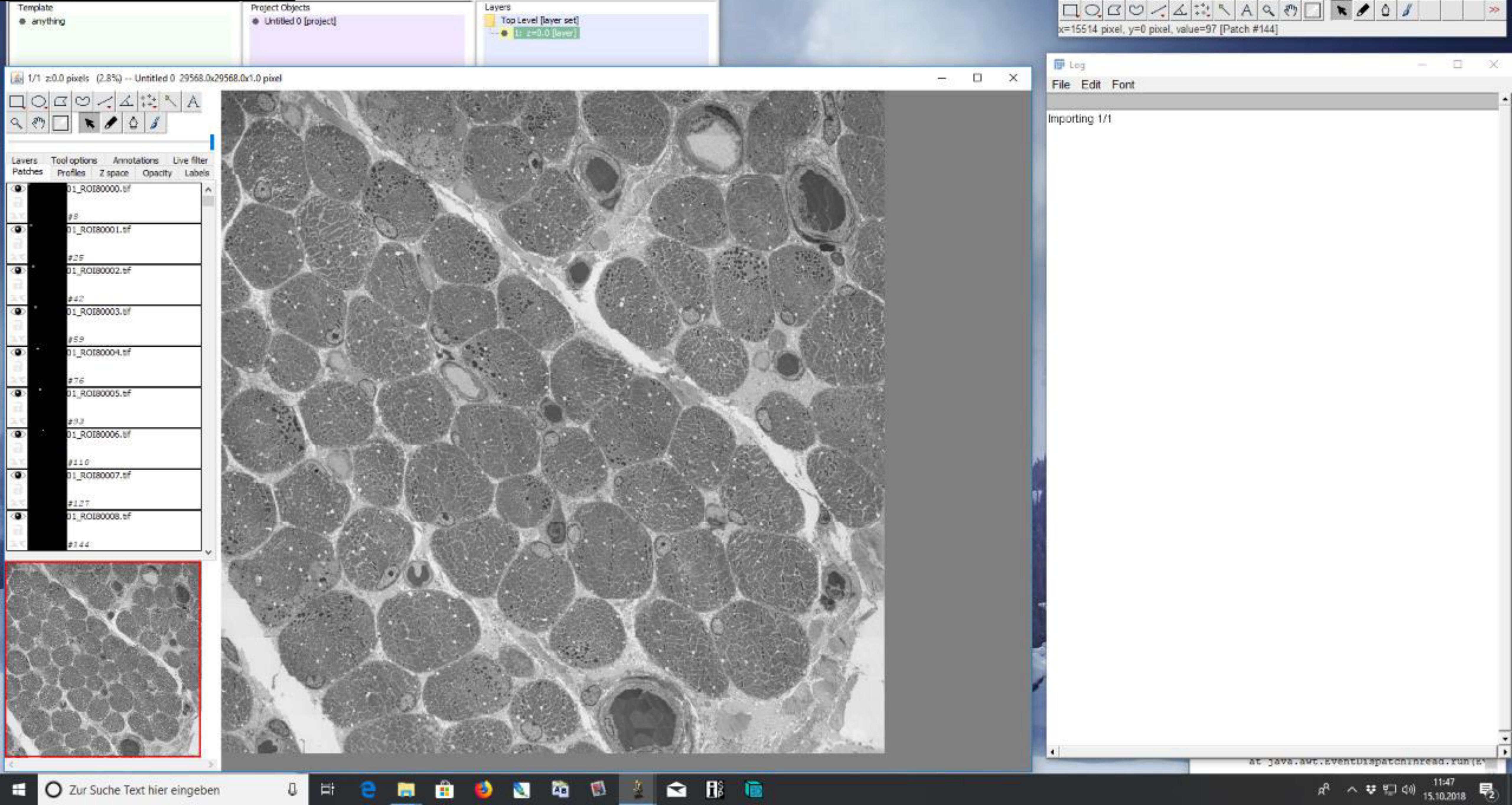

**1)** Import of ROI and LC

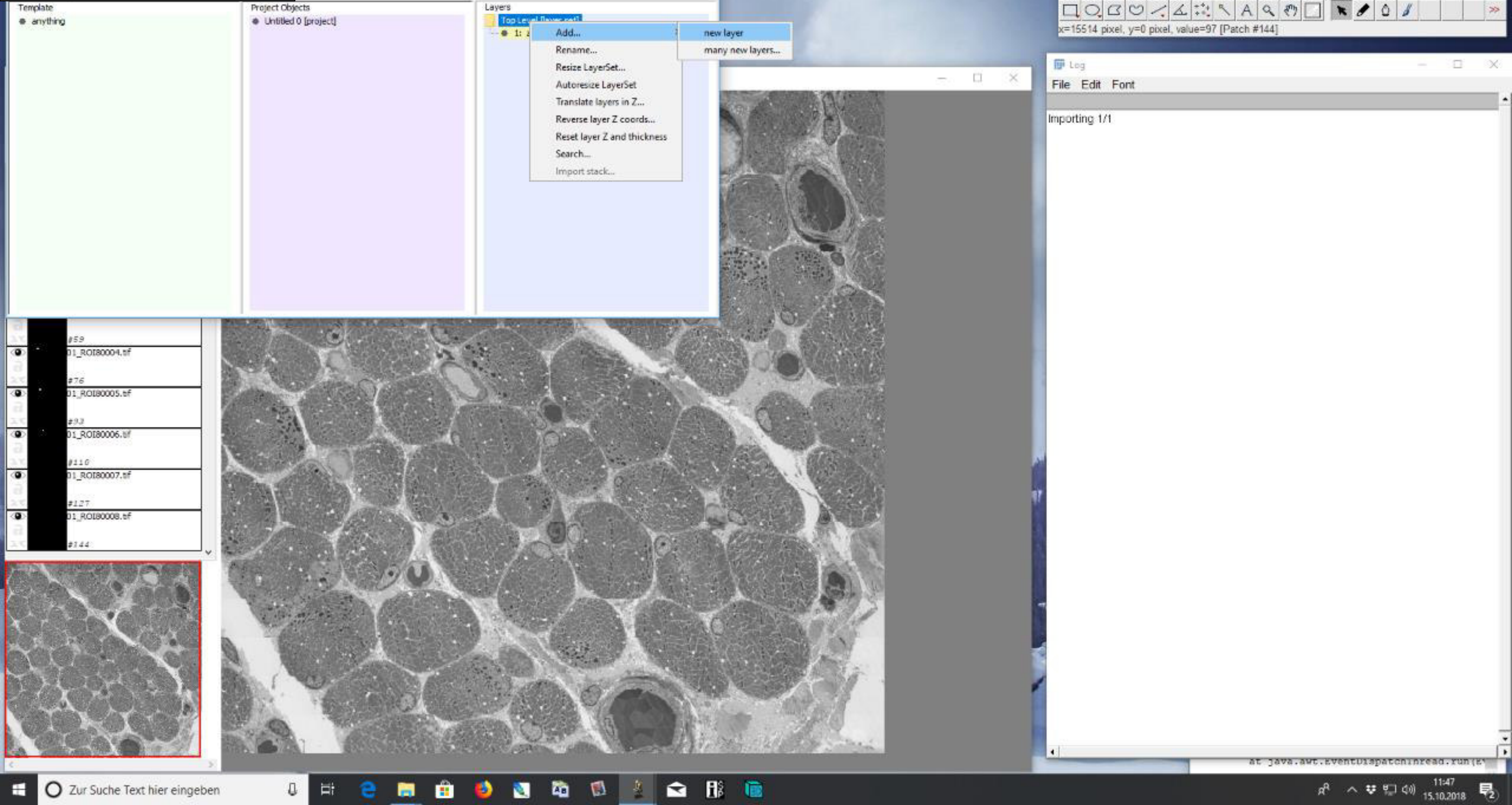

### 1) Import of ROI and LC

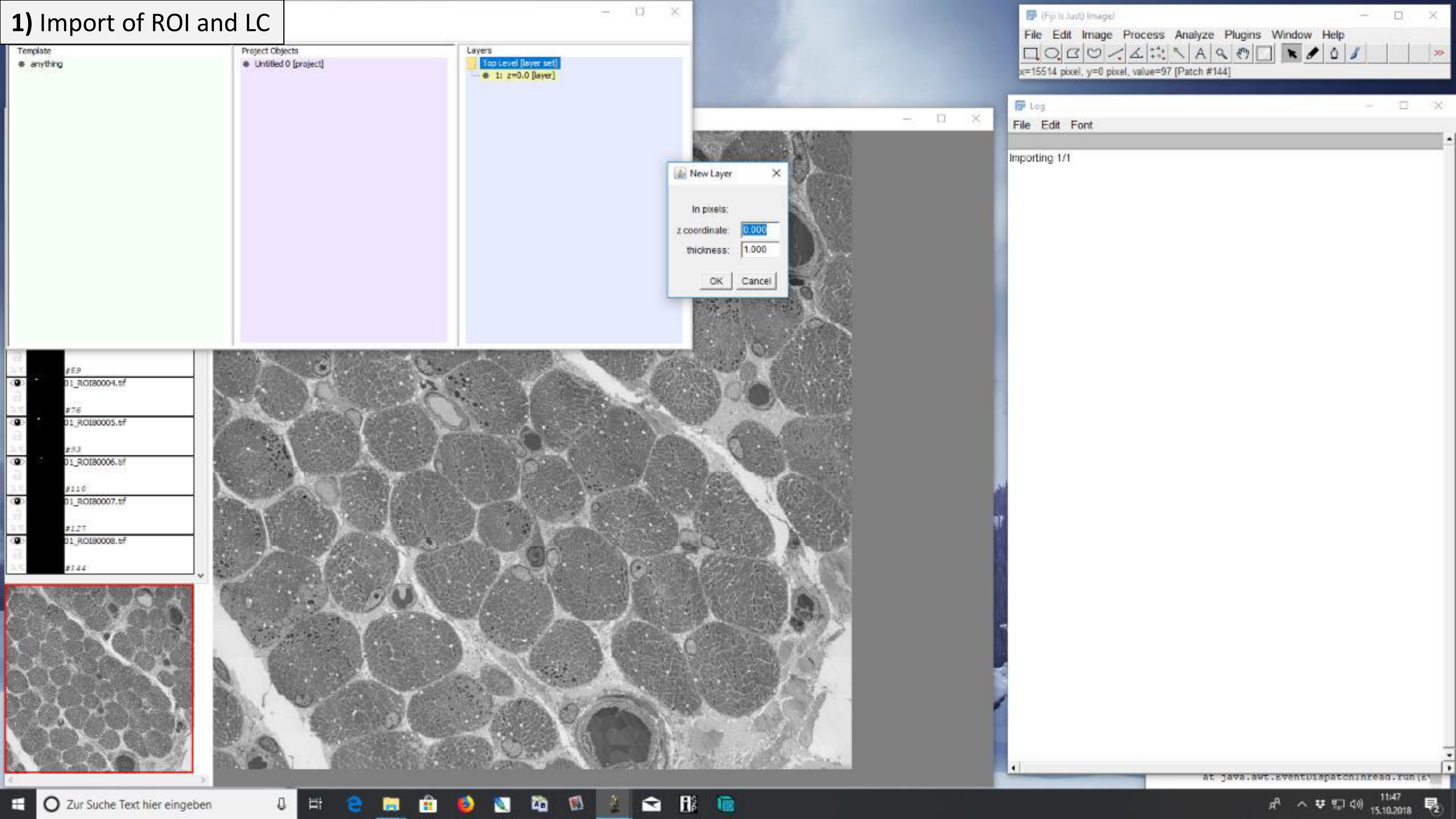

### 1) Import of ROI and LC

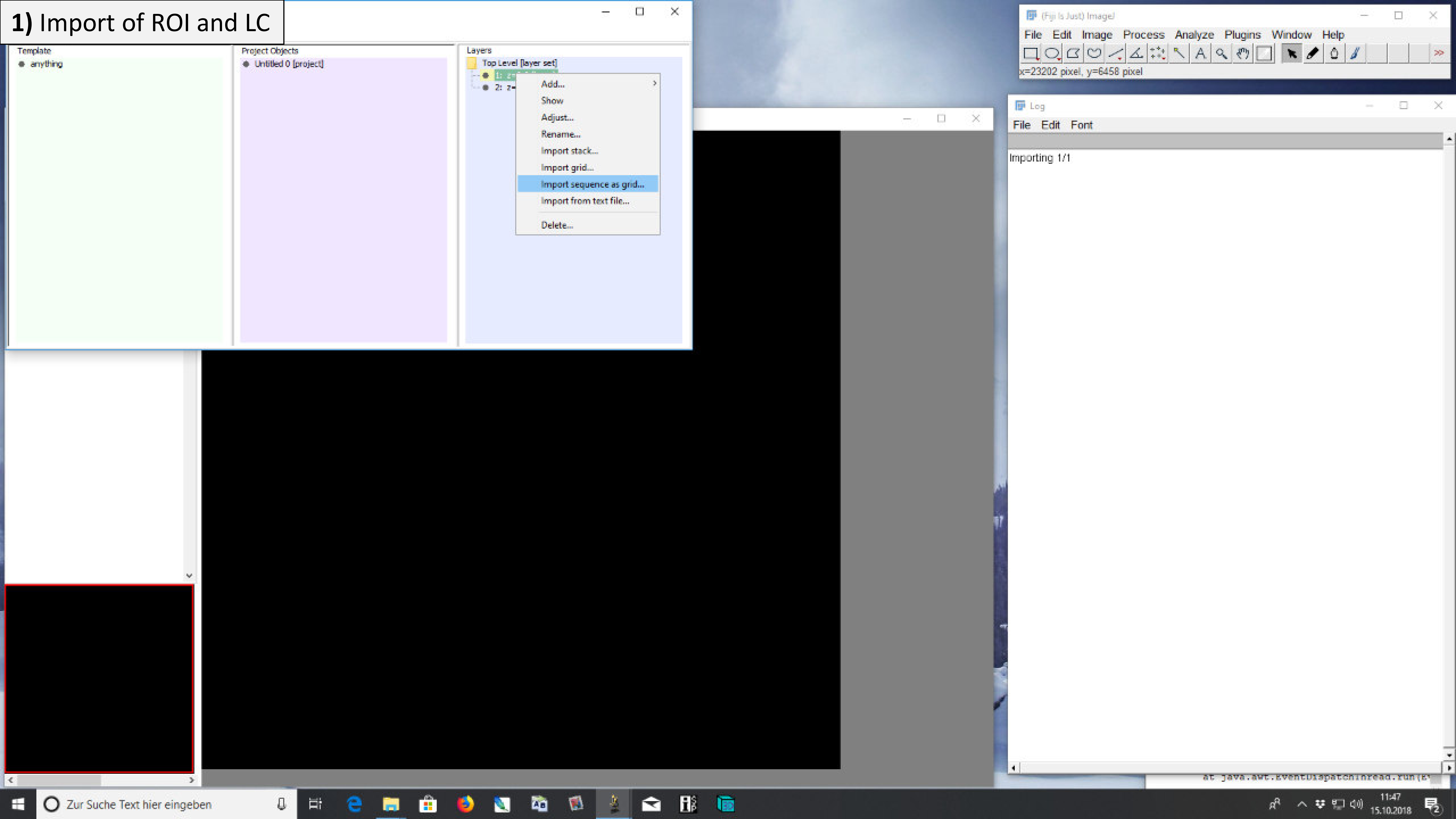

### 1) Import of ROI and LC

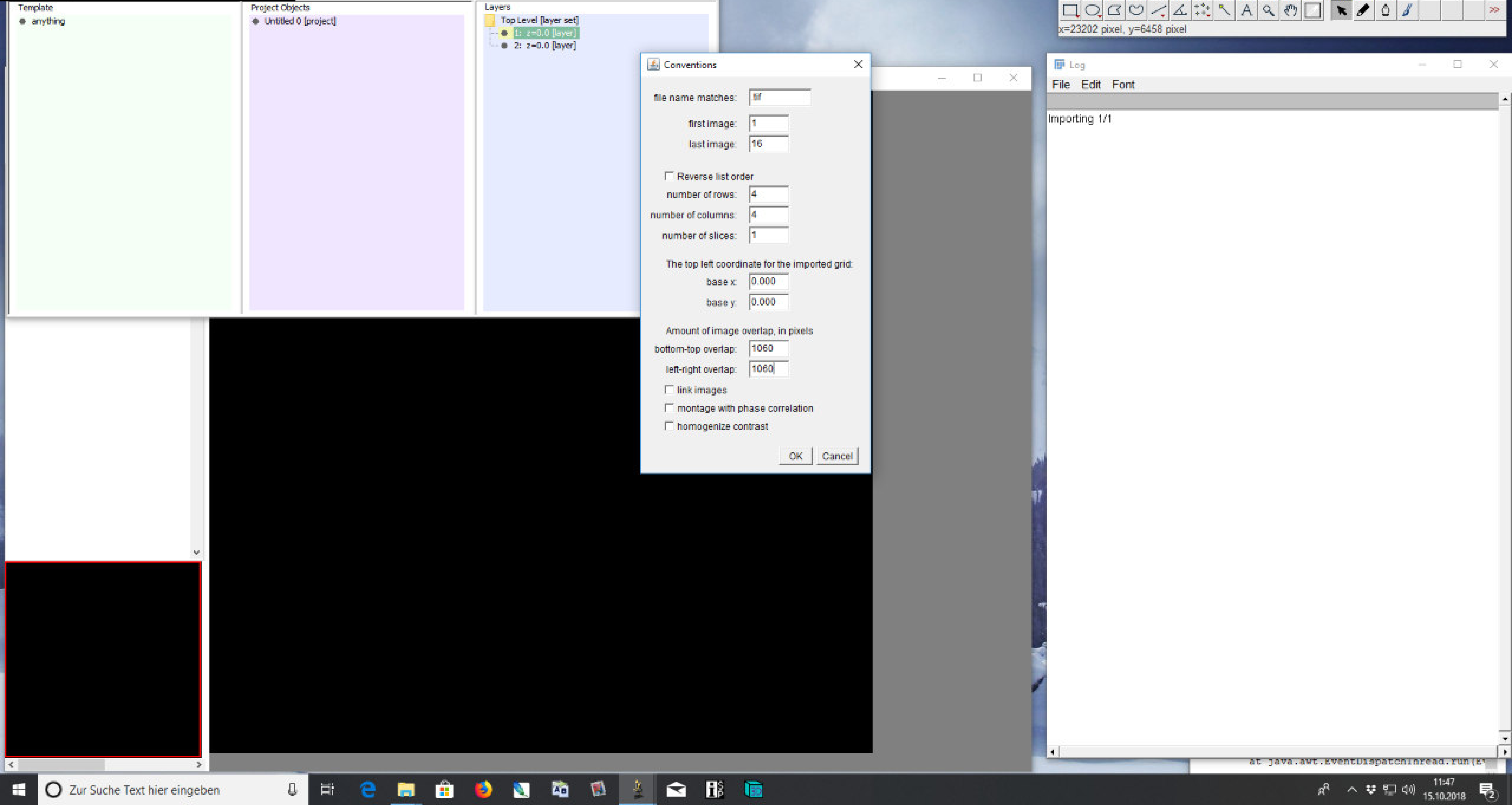

### 1) Import of ROI and LC

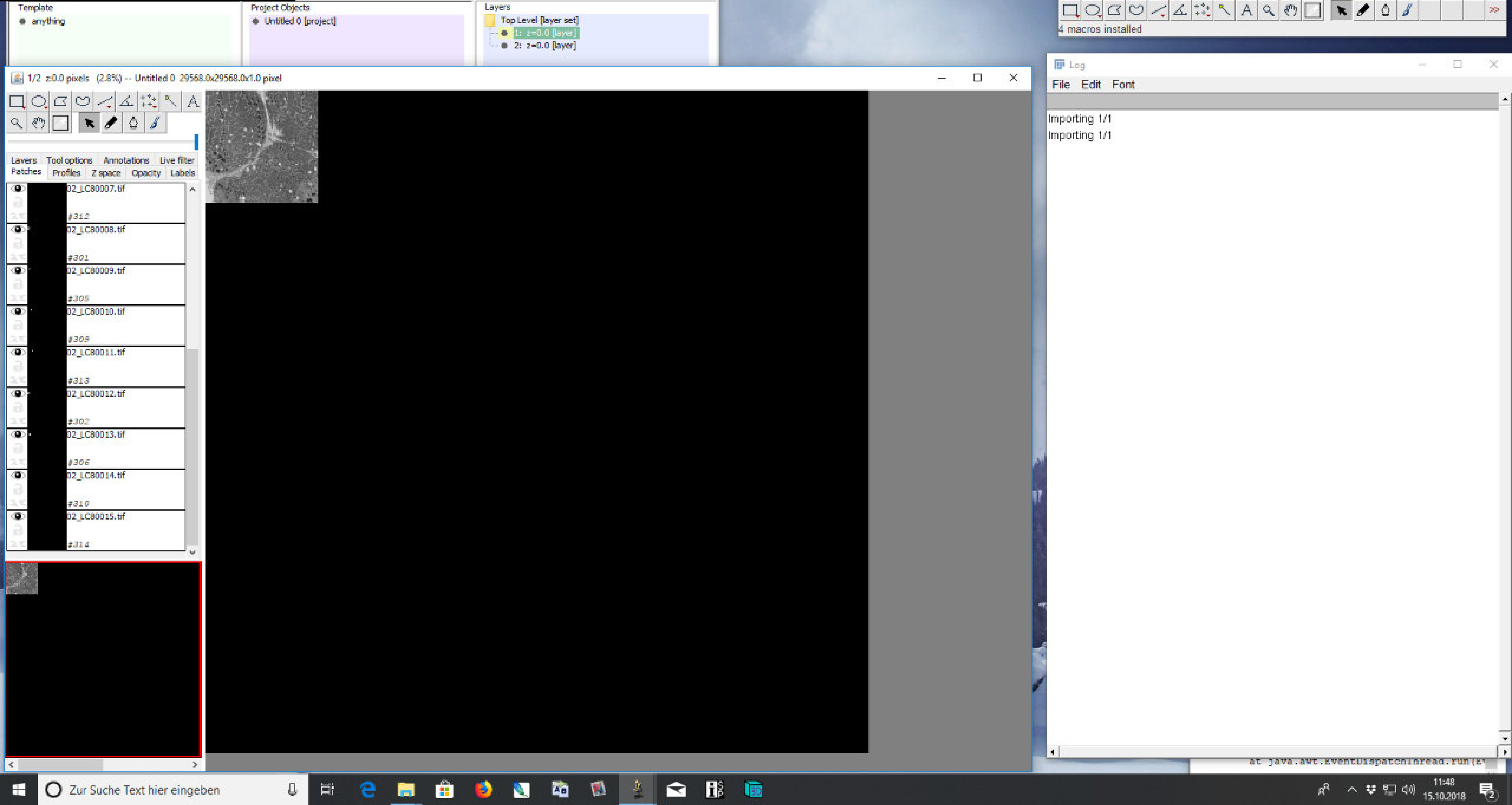

### 2) Apply filter

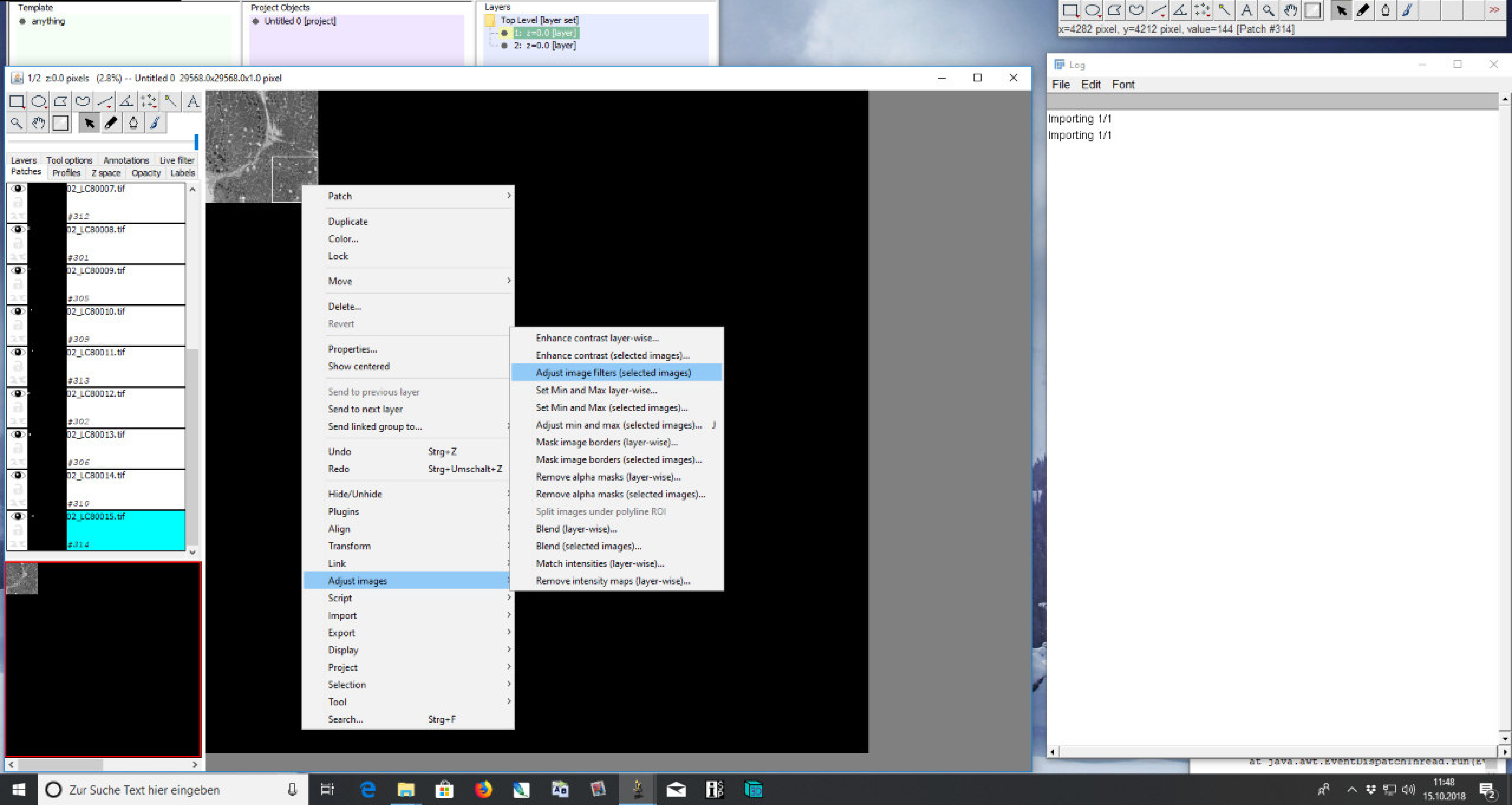

### 2) Apply filter

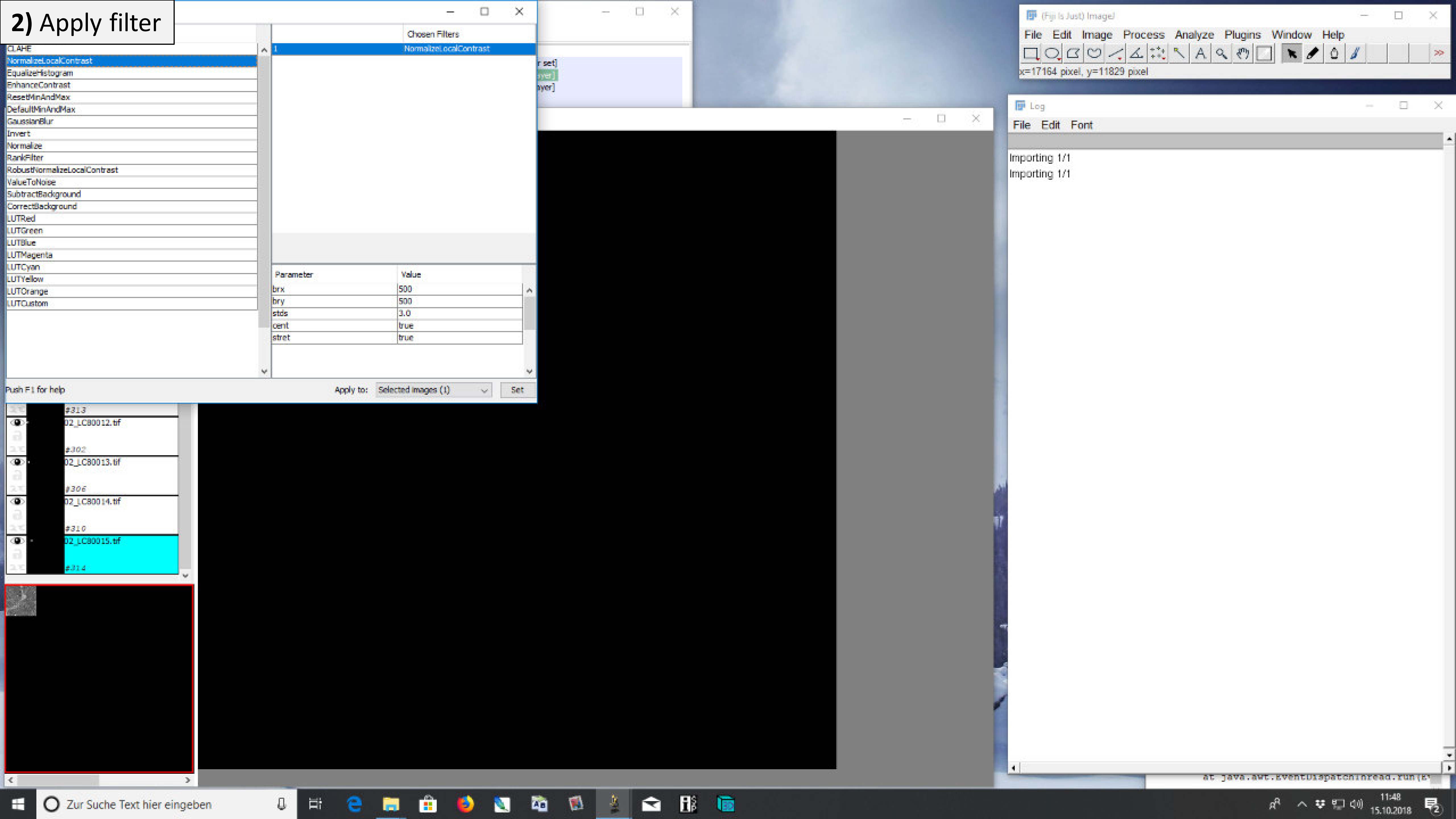

### 2) Apply filter

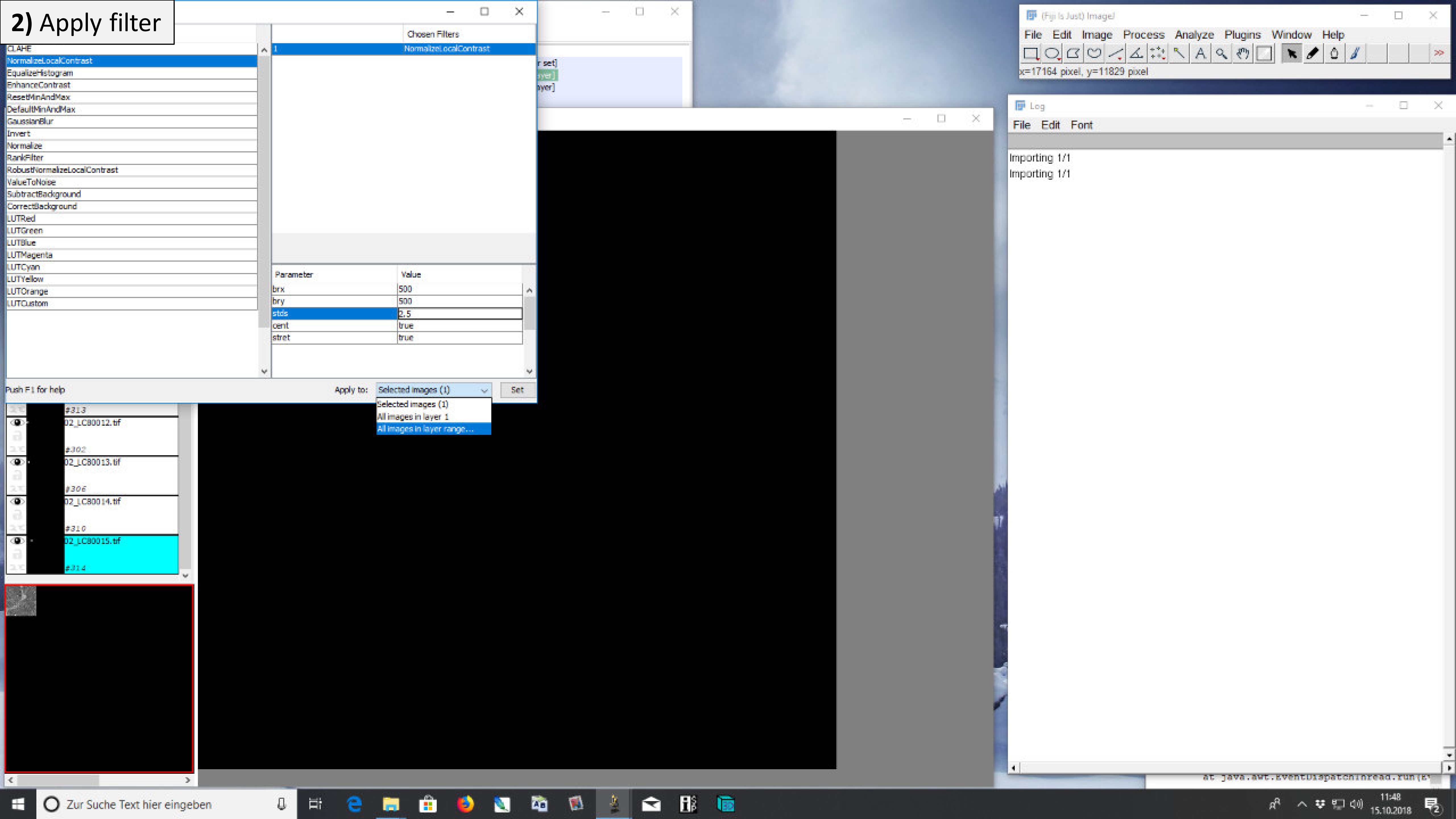

#### 2) Apply filter

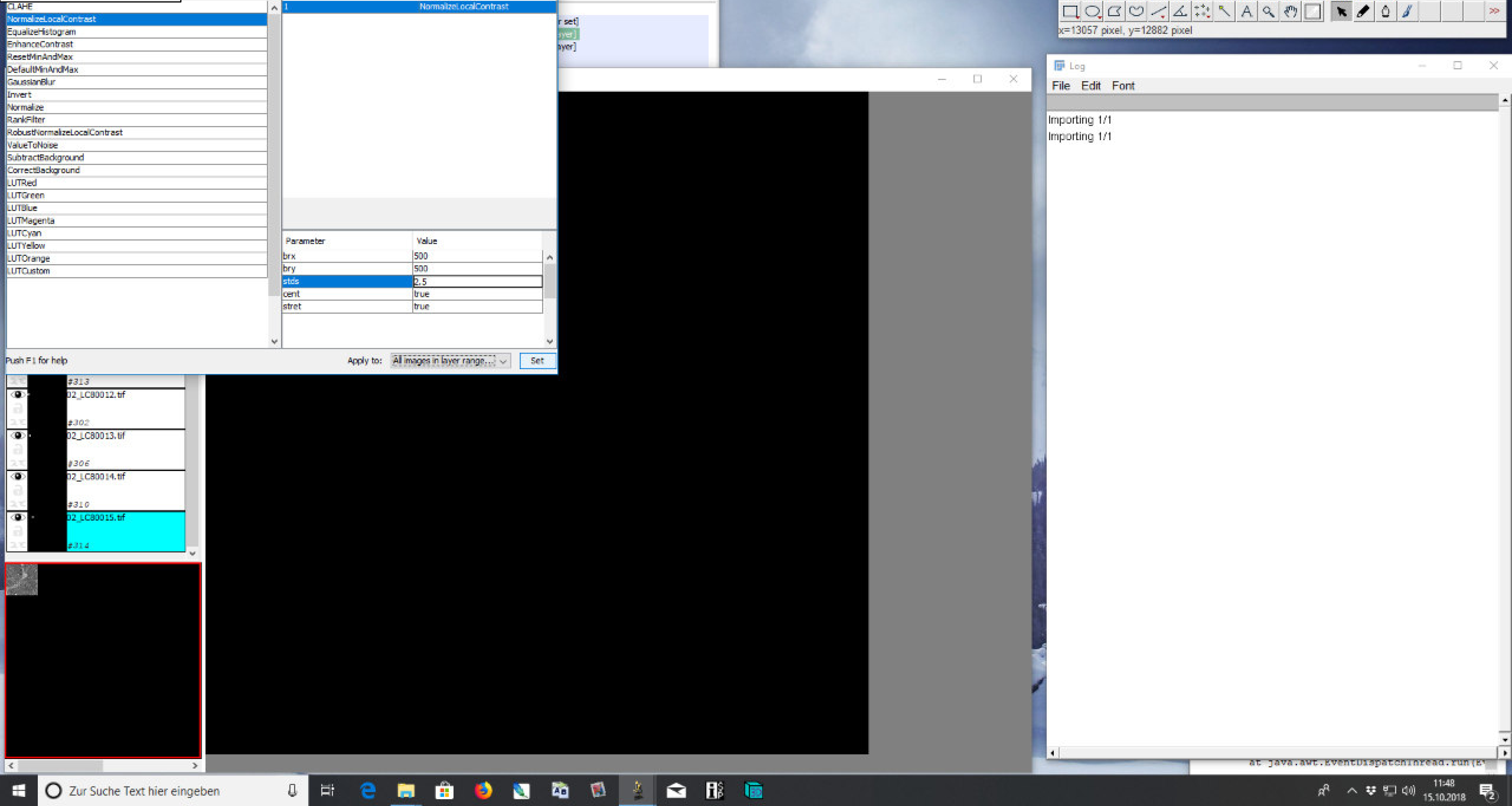

### 2) Apply filter

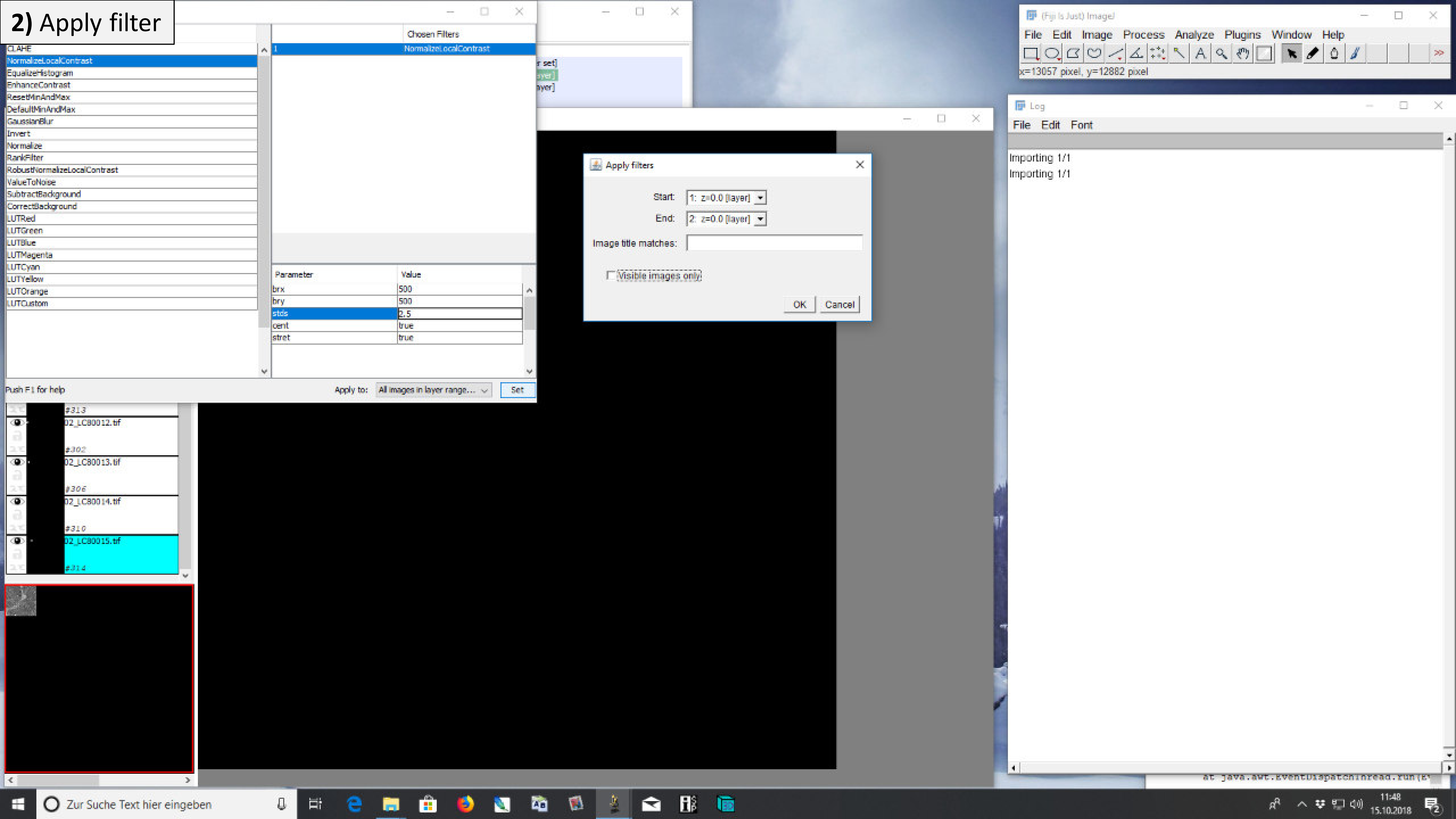

#### 2) Apply filter

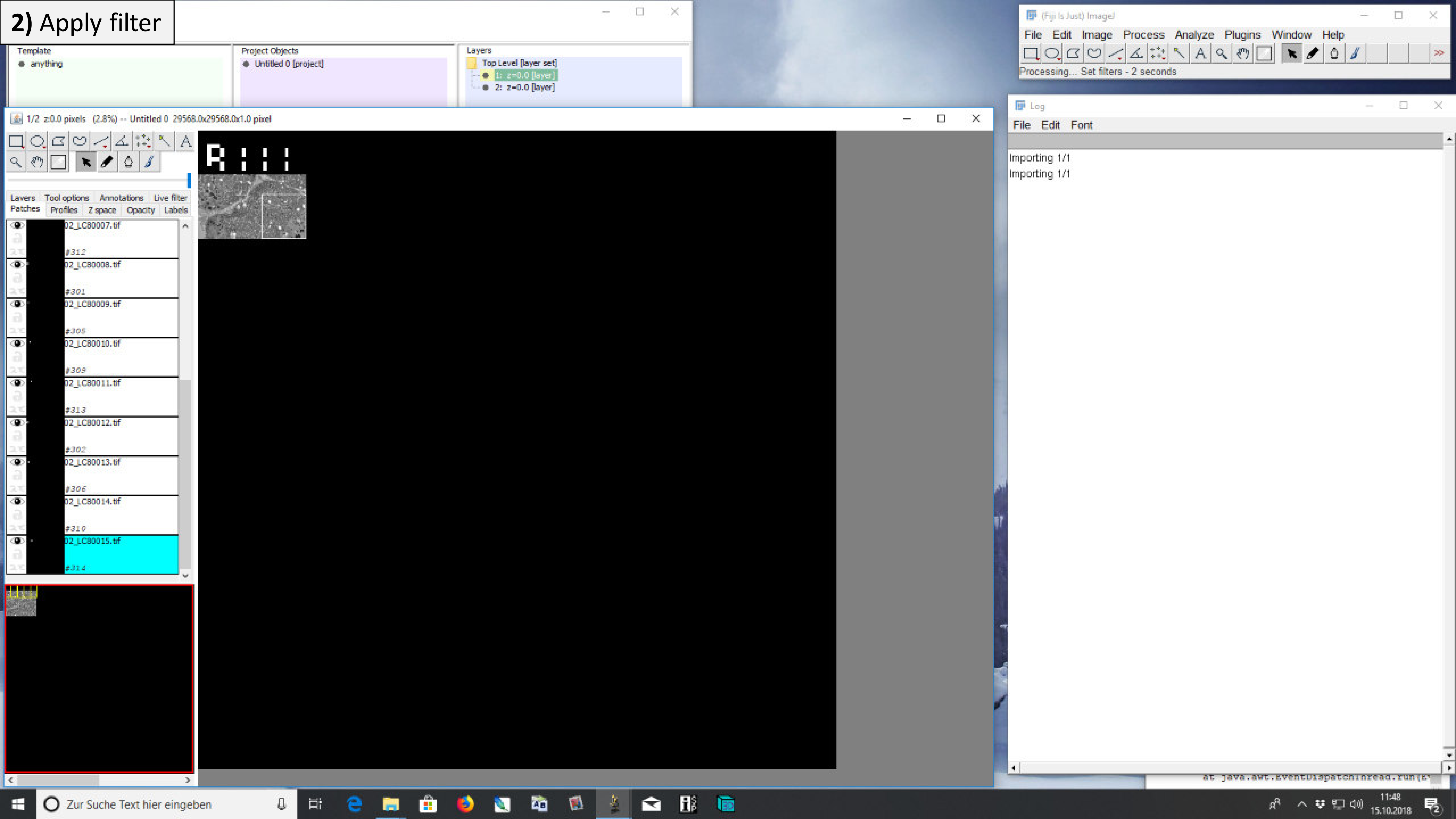

### 2) Apply filter

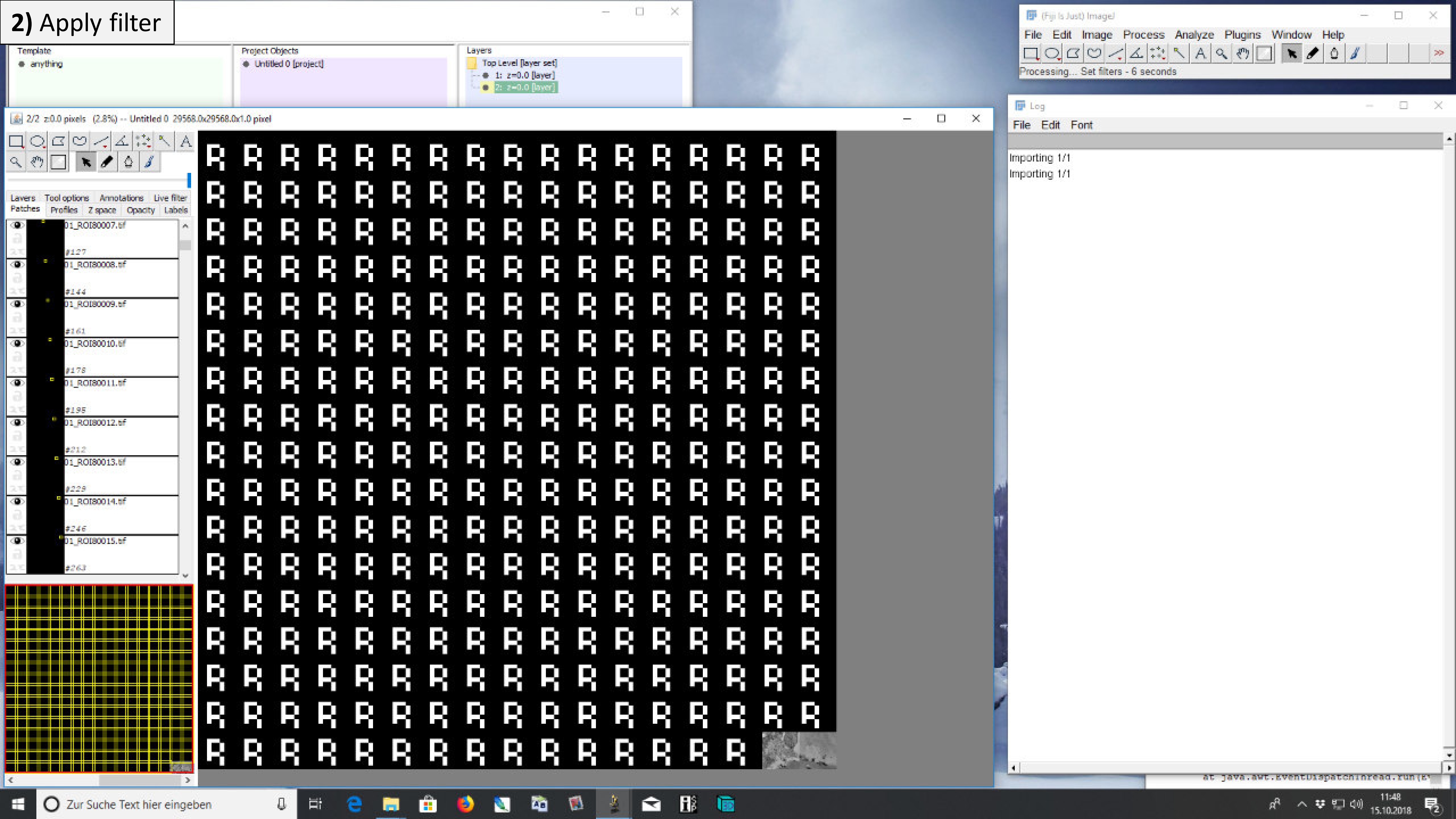

#### 2) Apply filter

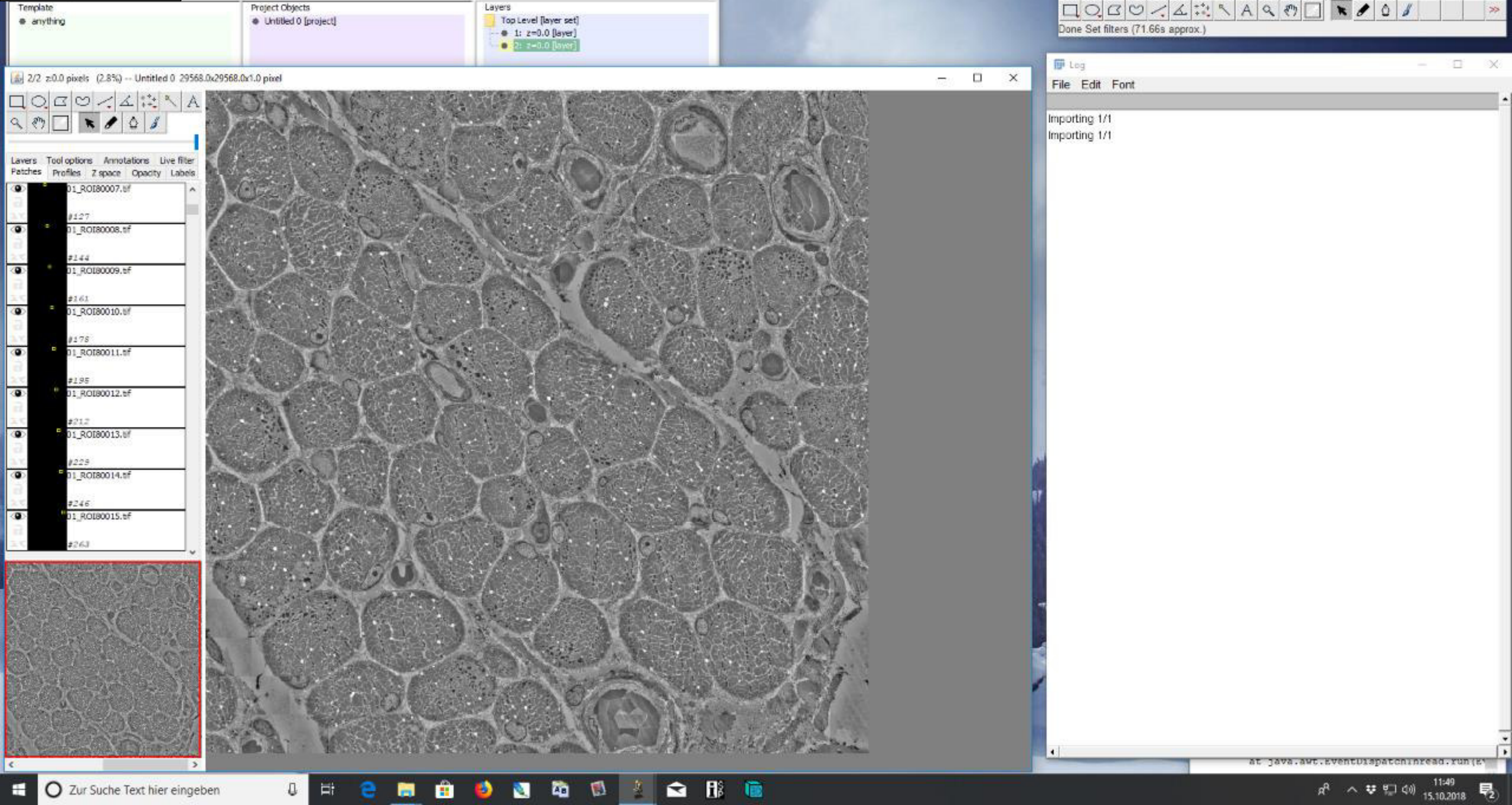

### 3) Alignment of ROI and LC

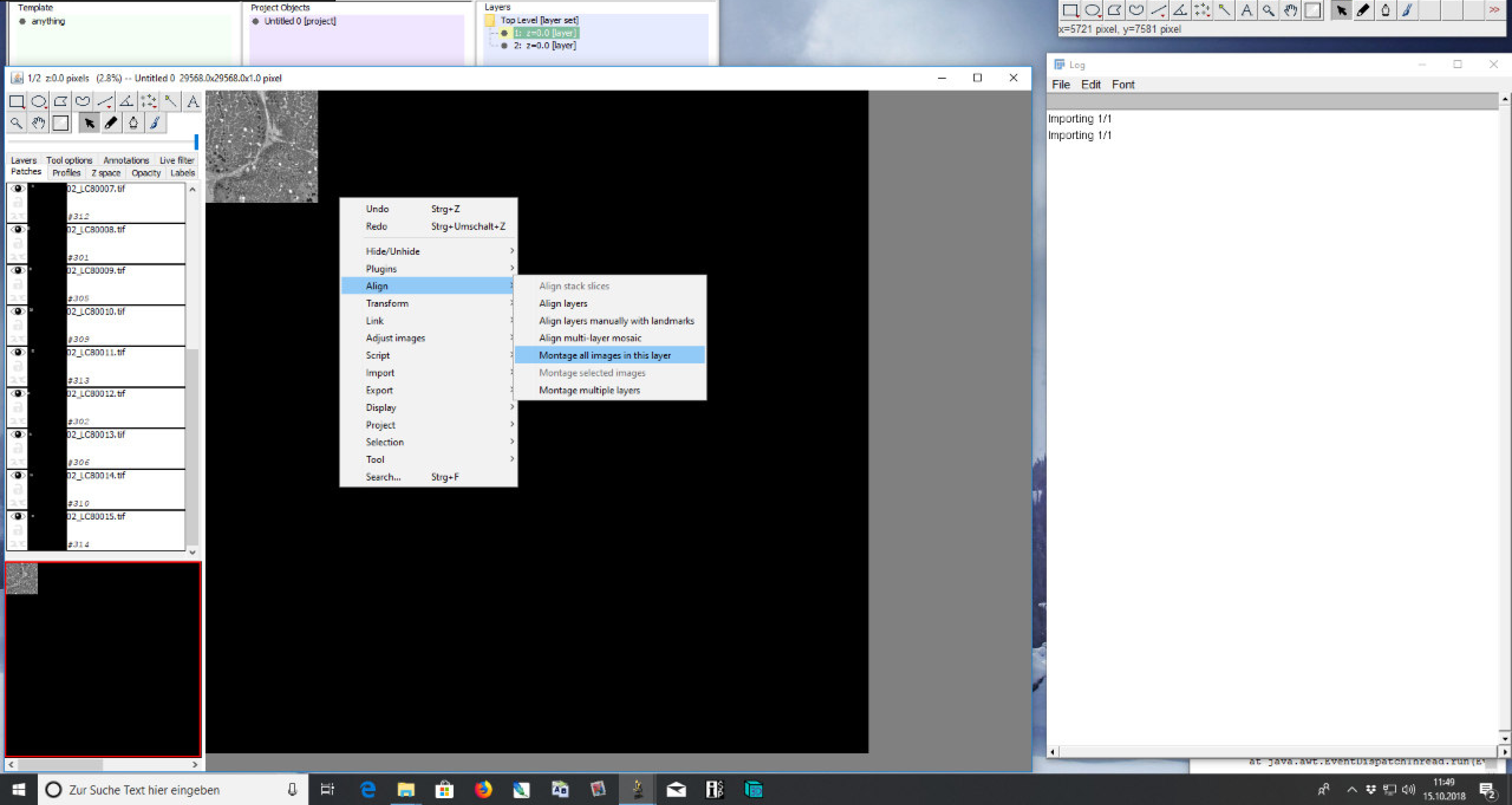

### **3)** Alignment of ROI and LC

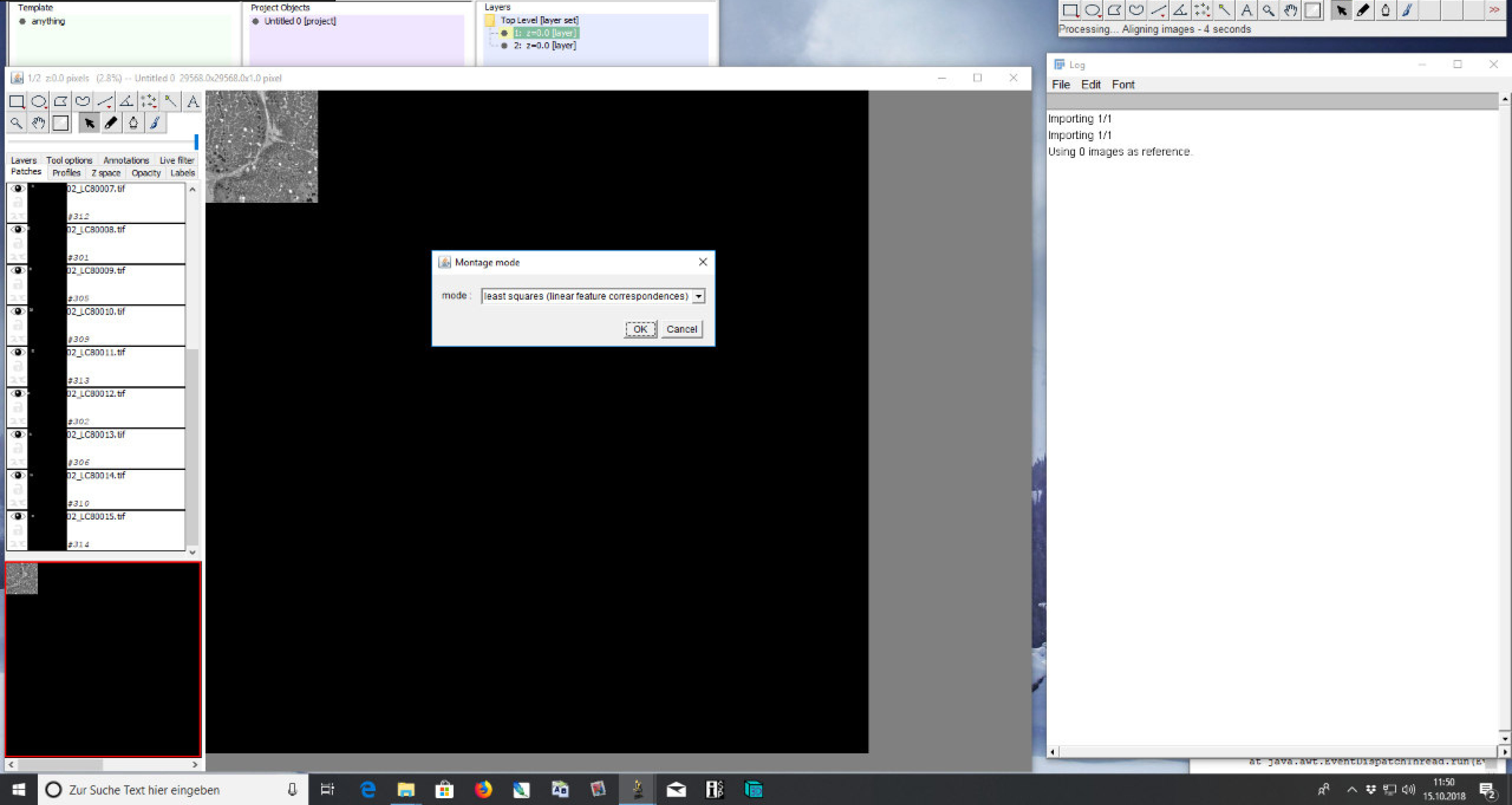

### 3) Alignment of ROI and LC

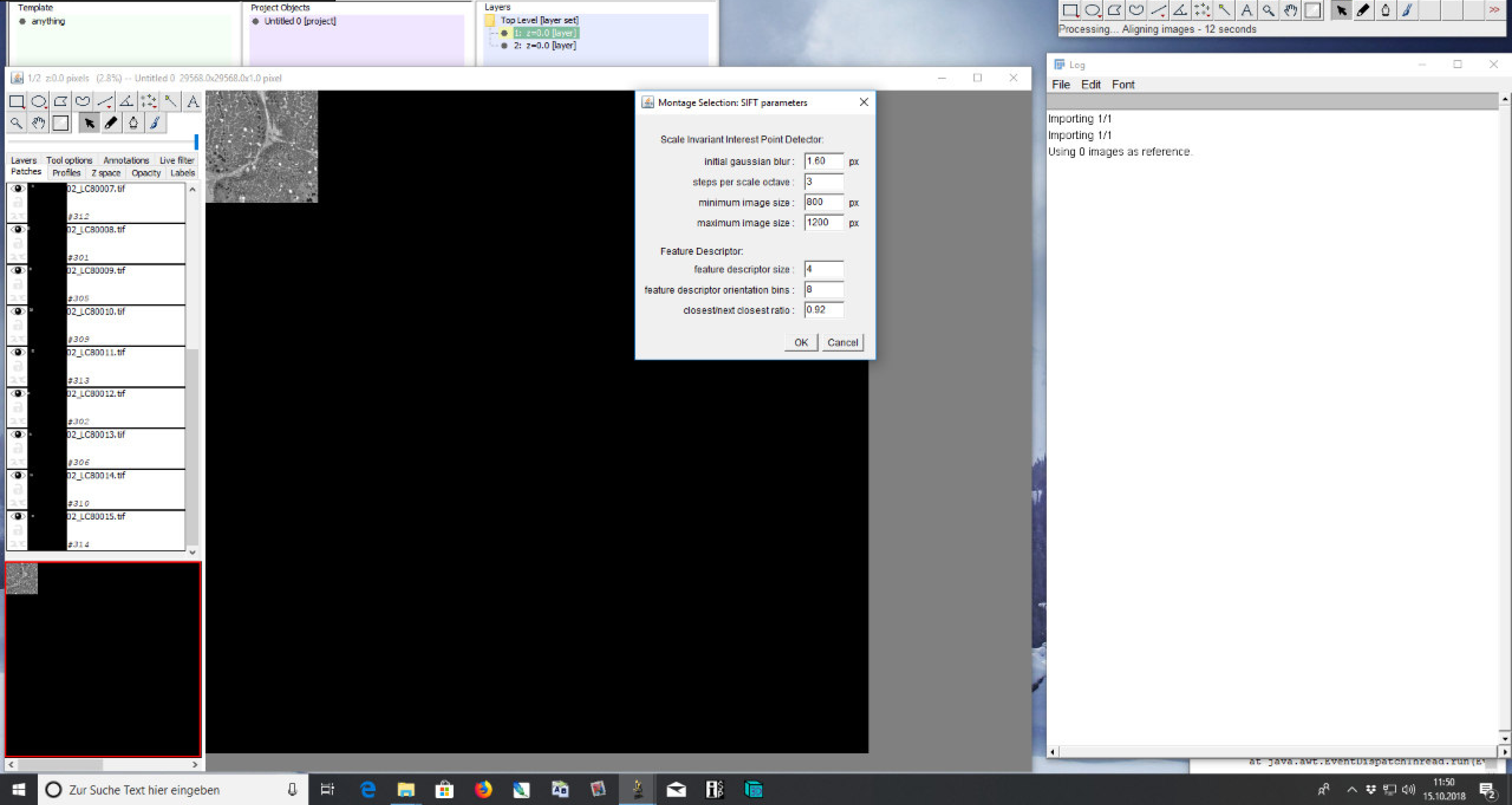

### **3)** Alignment of ROI and LC

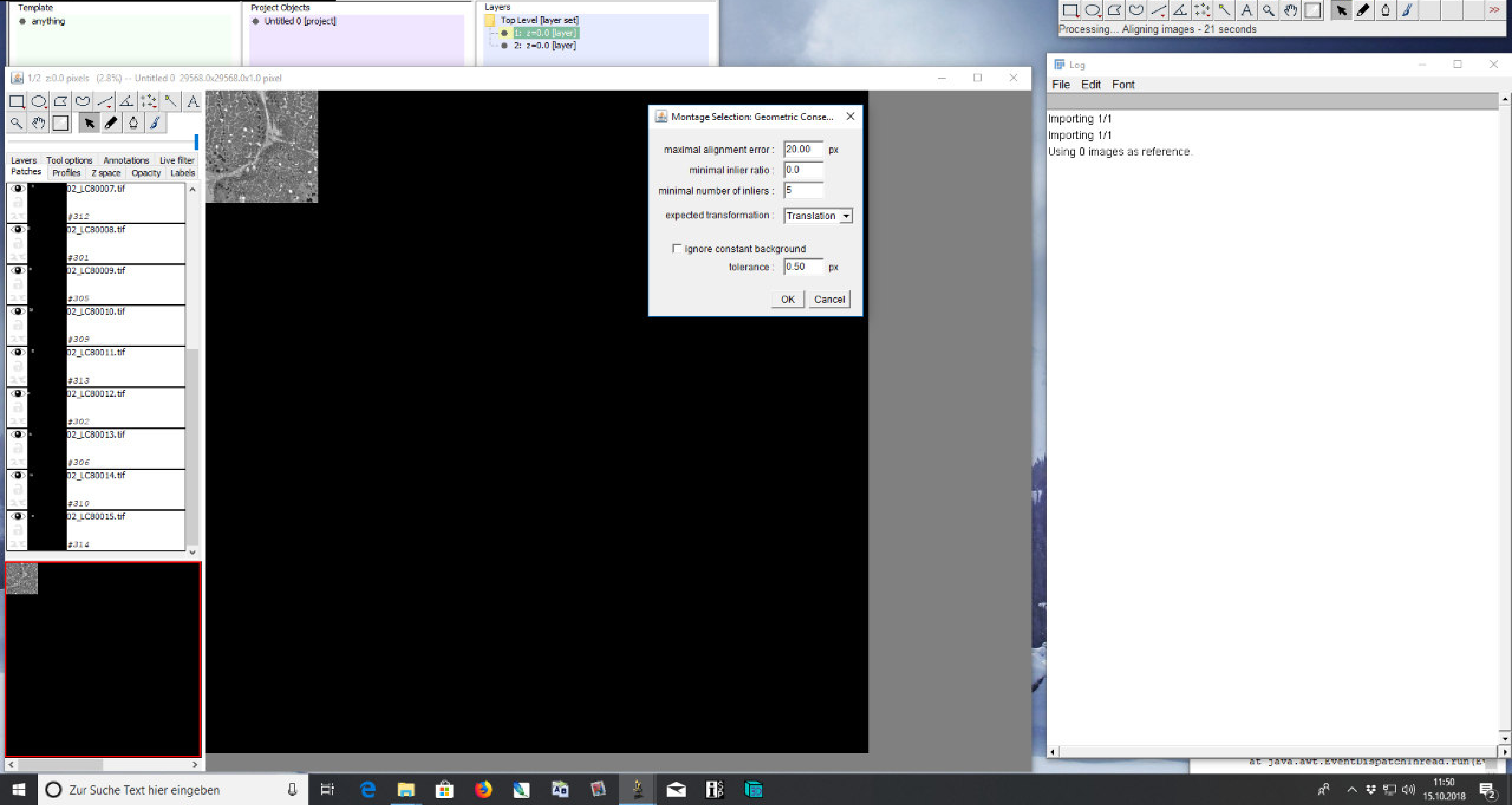

### 3) Alignment of ROI and LC

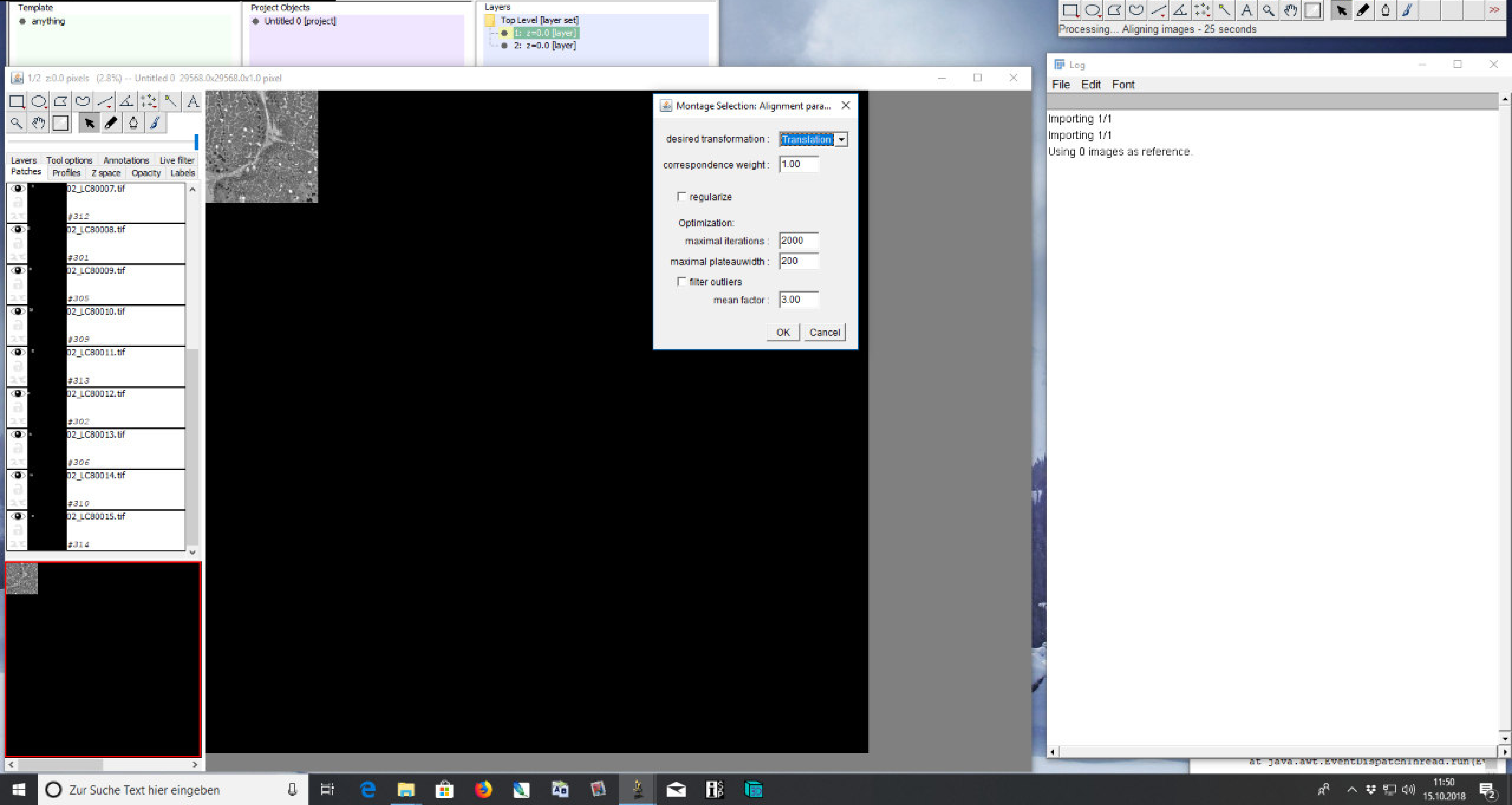

### 3) Alignment of ROI and LC

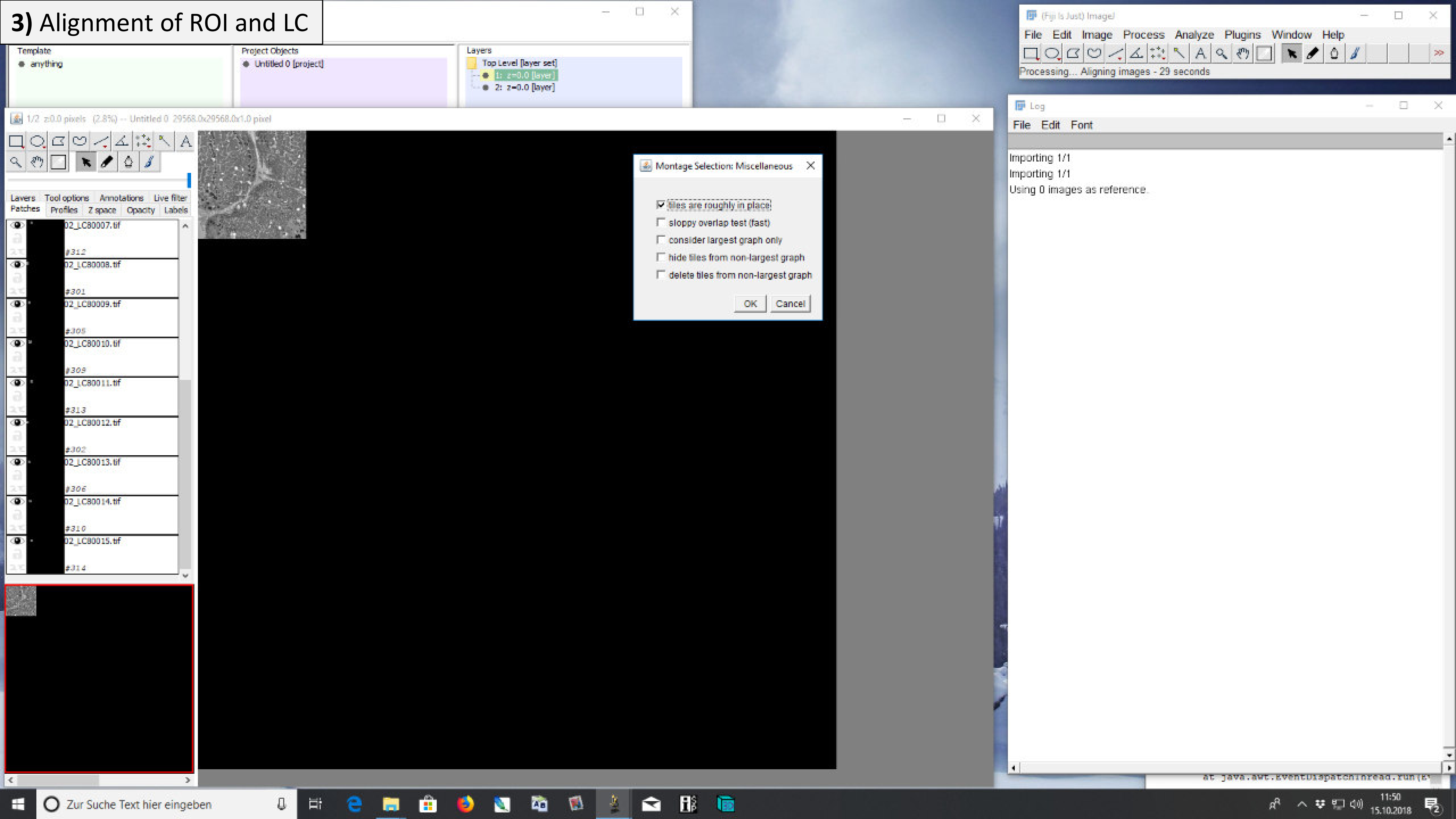

### **3)** Alignment of ROI and LC

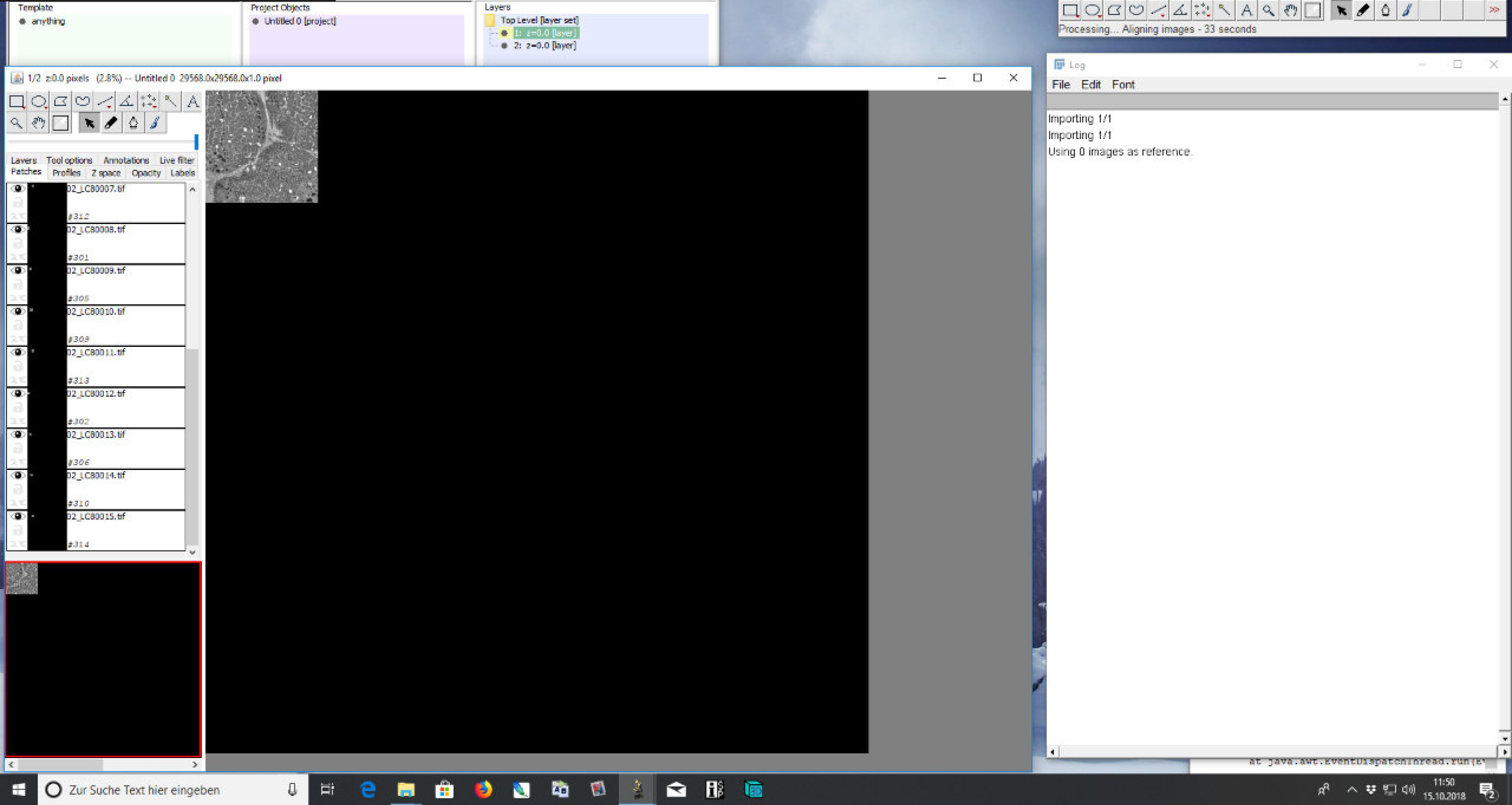

### 3) Alignment of ROI and LC

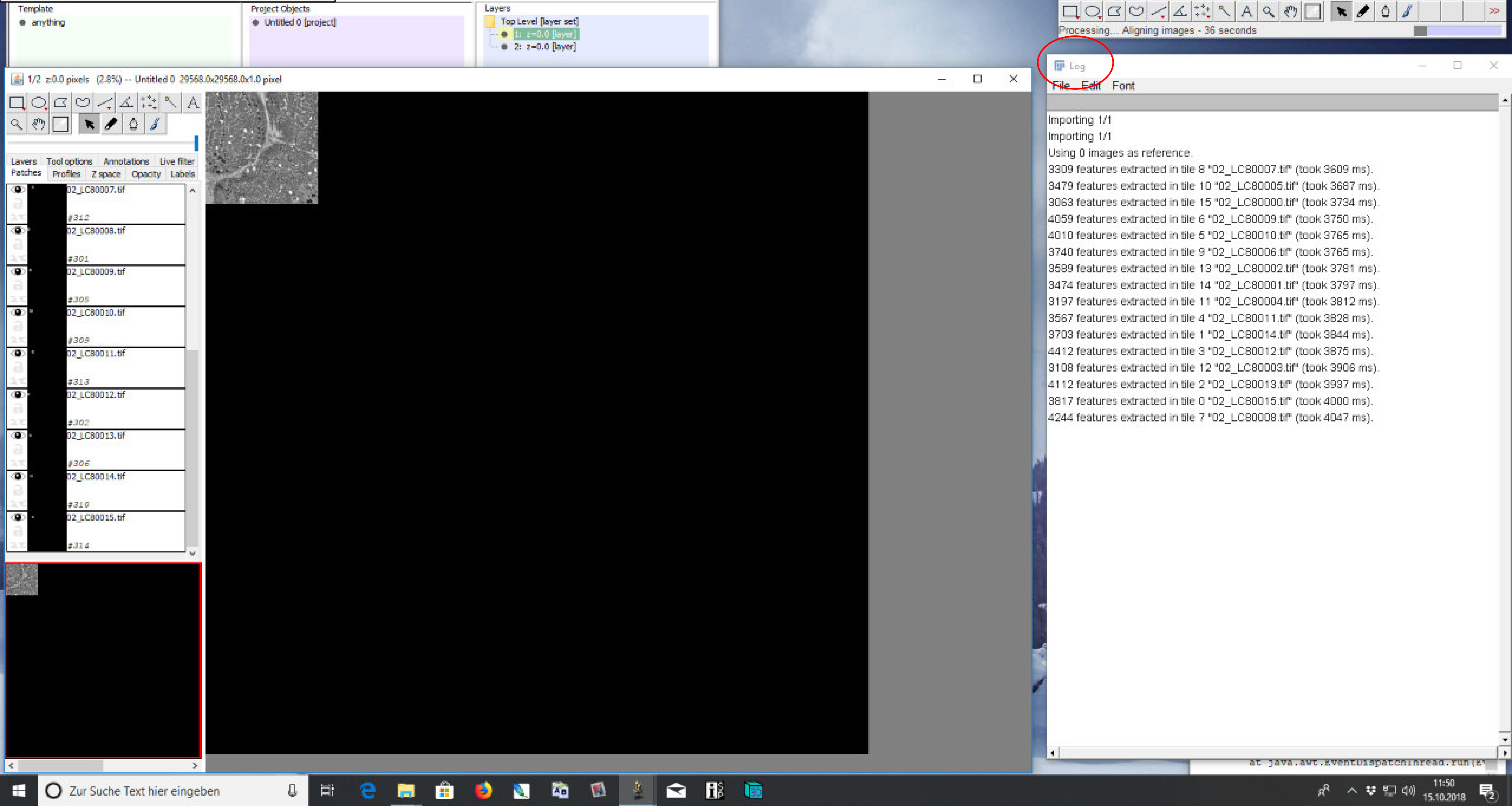

### 3) Alignment of ROI and LC

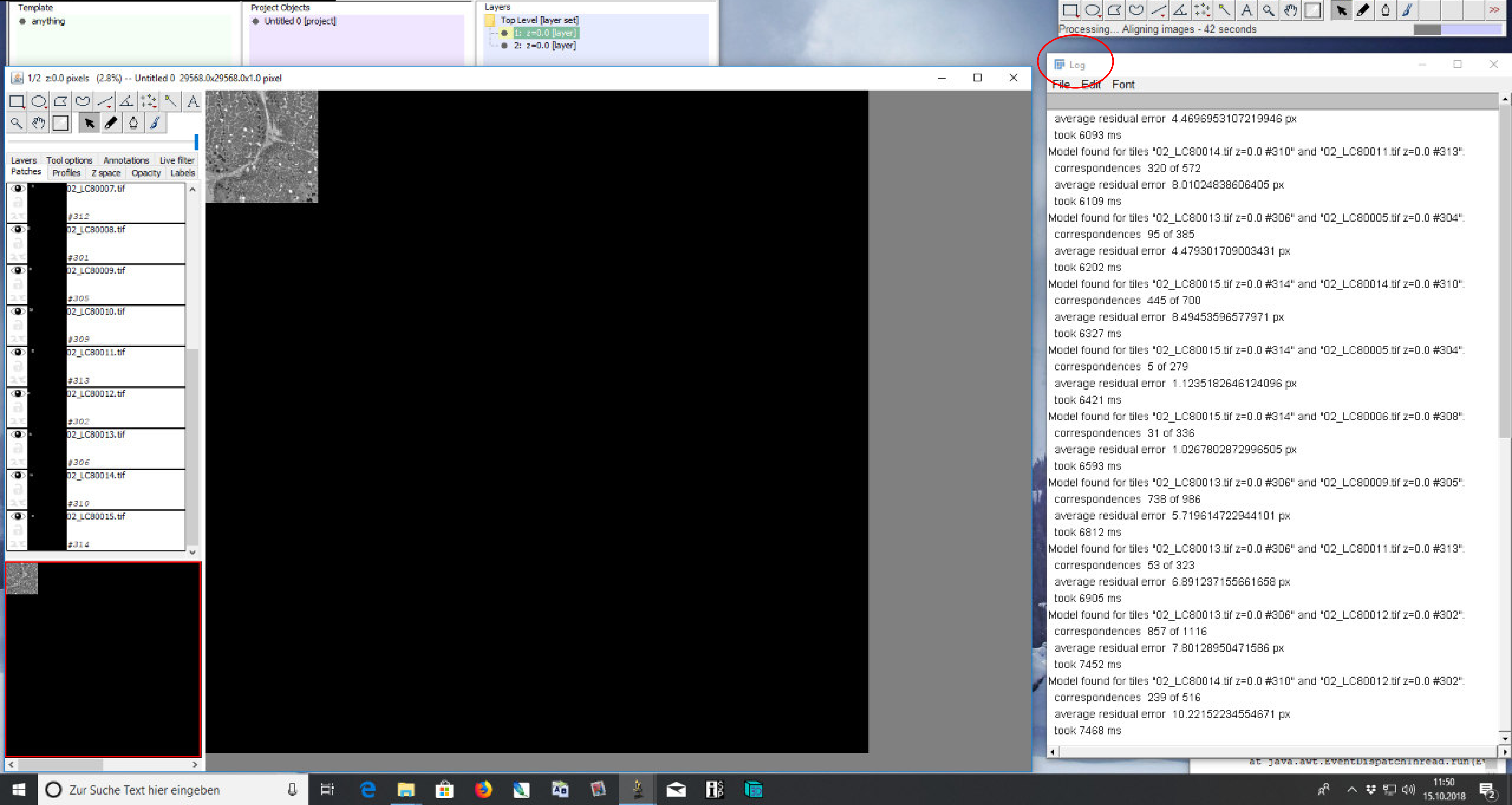

### 3) Alignment of ROI and LC

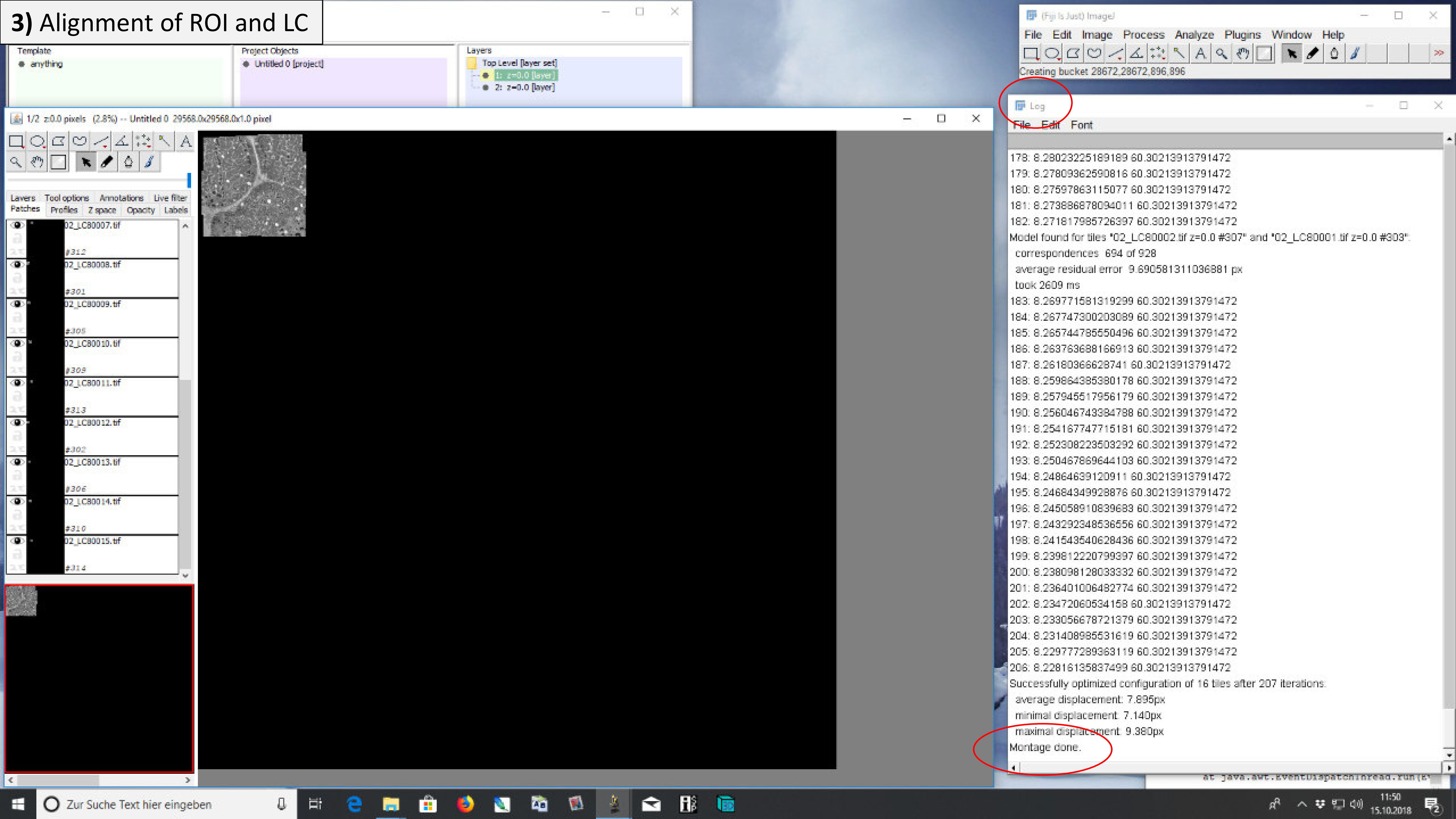

### 3) Alignment of ROI and LC

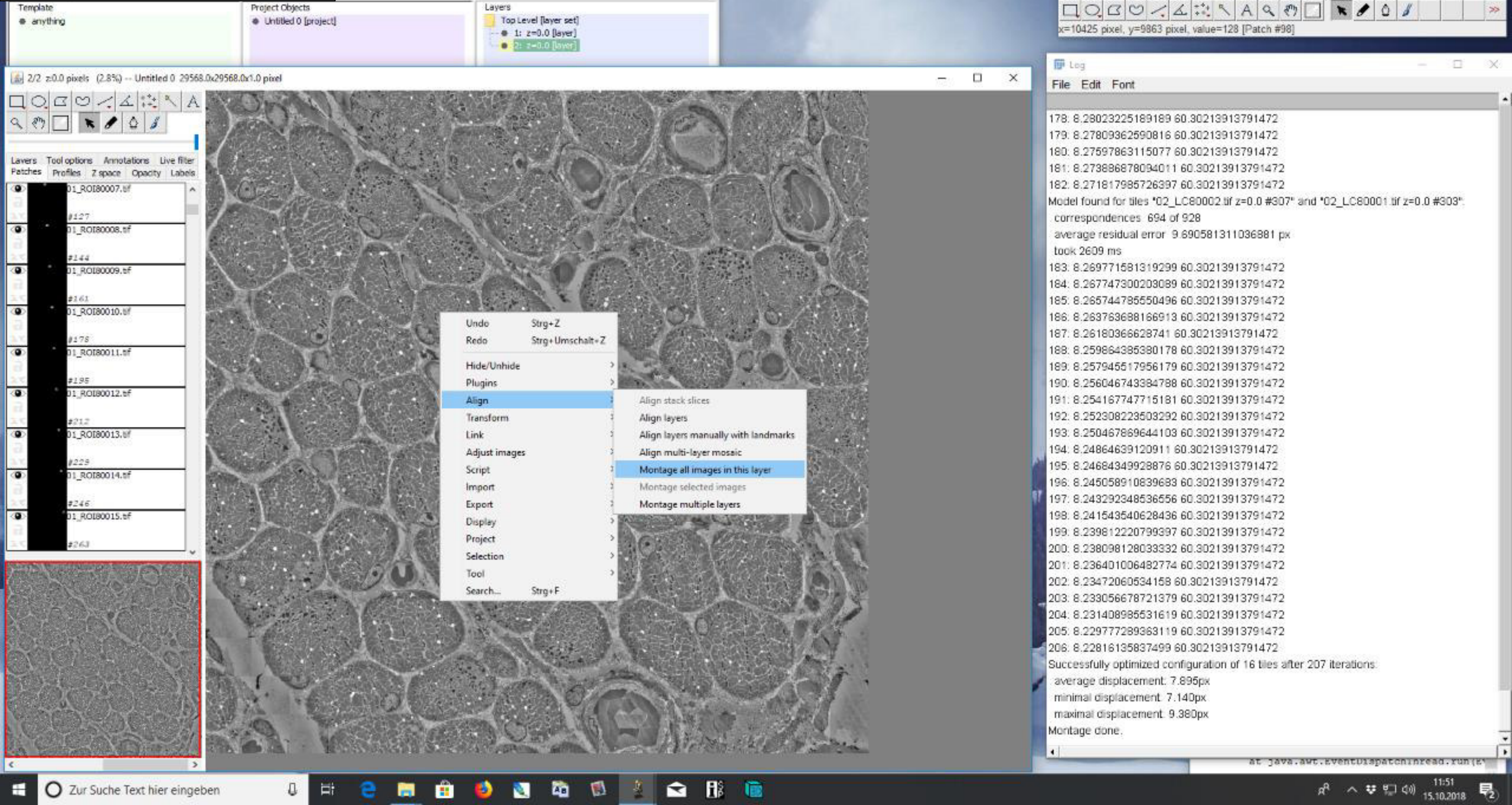

### **3)** Alignment of ROI and LC

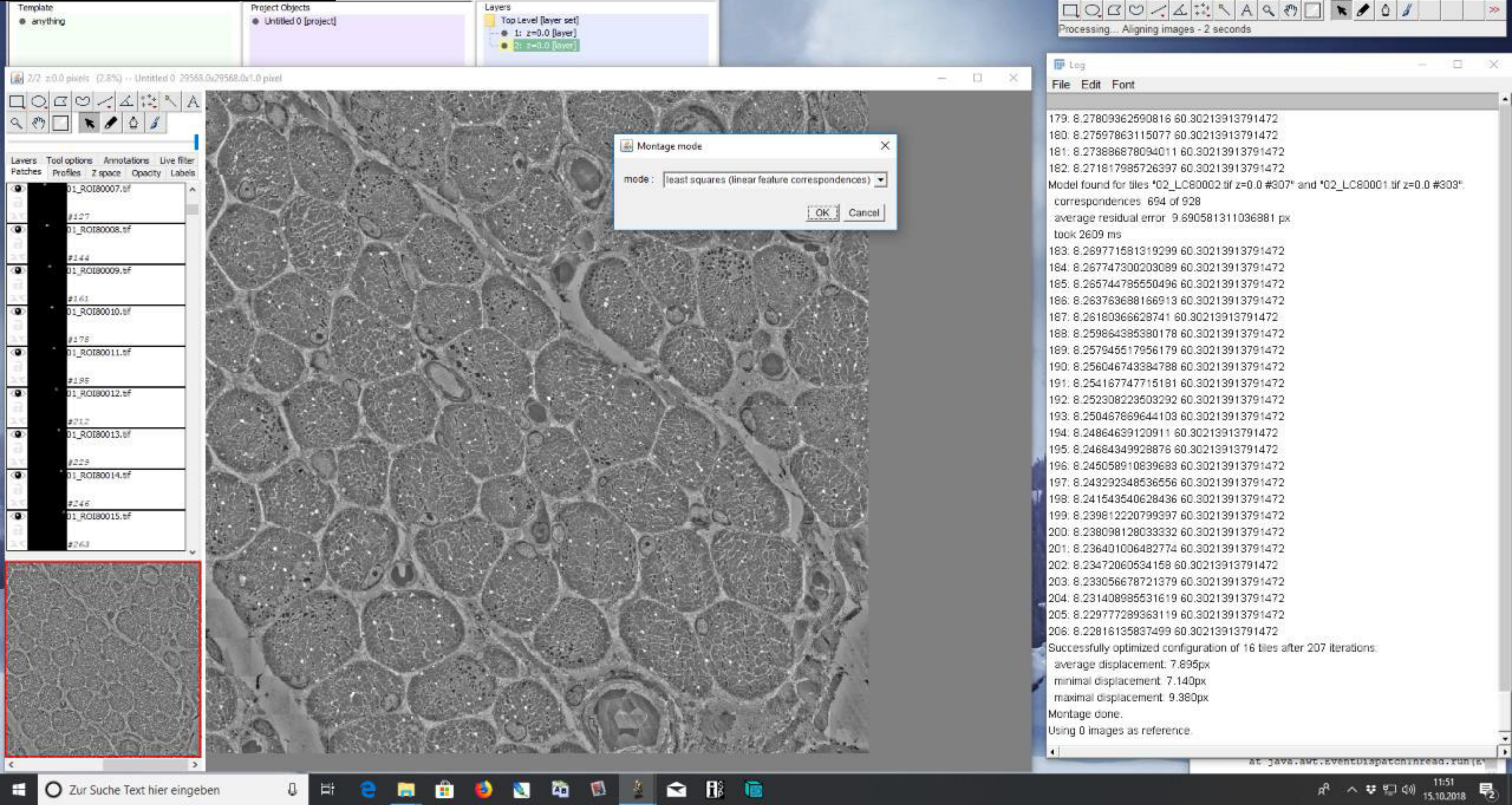

**3)** Alignment of ROI and LC

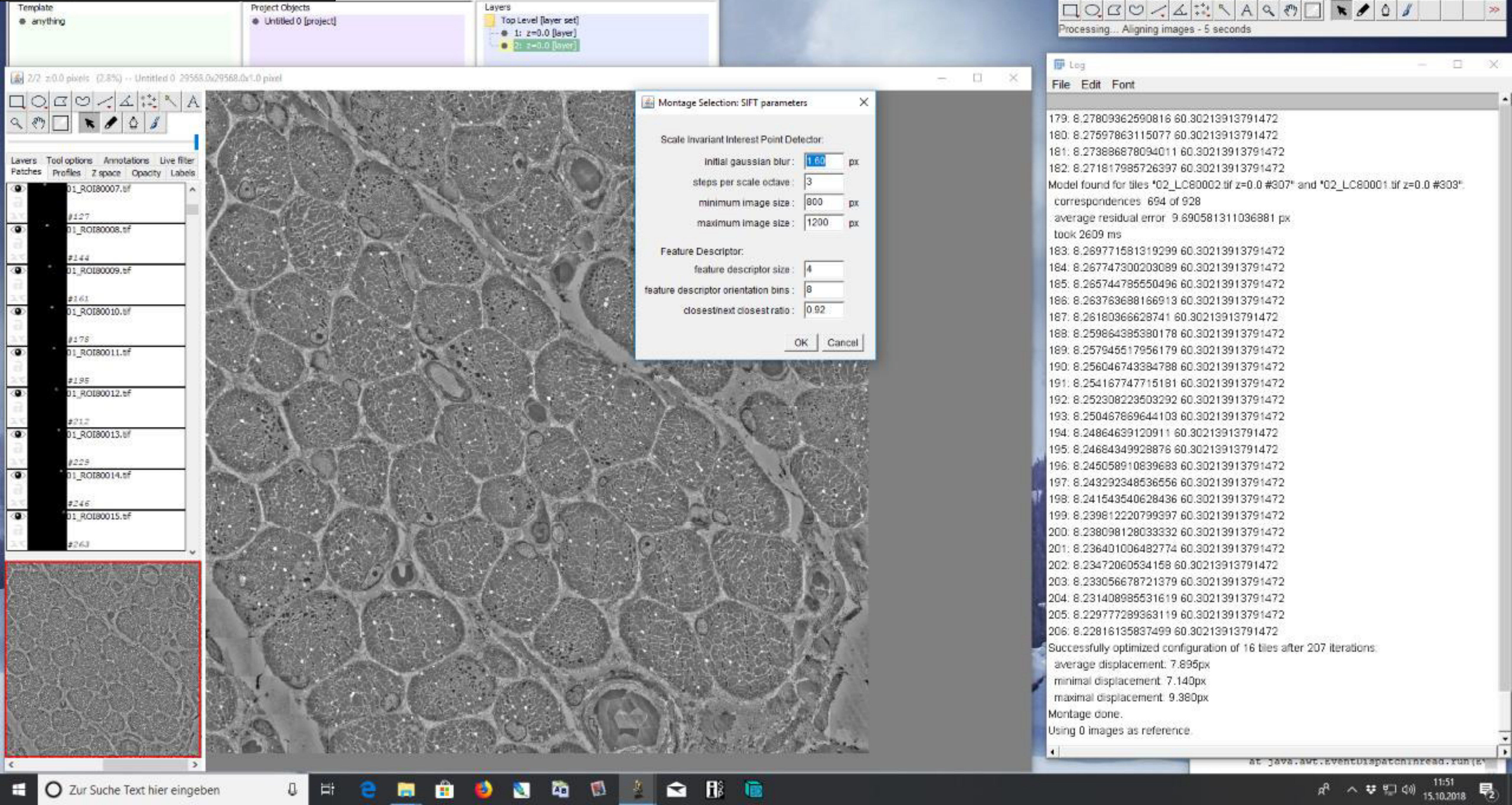

### **3)** Alignment of ROI and LC

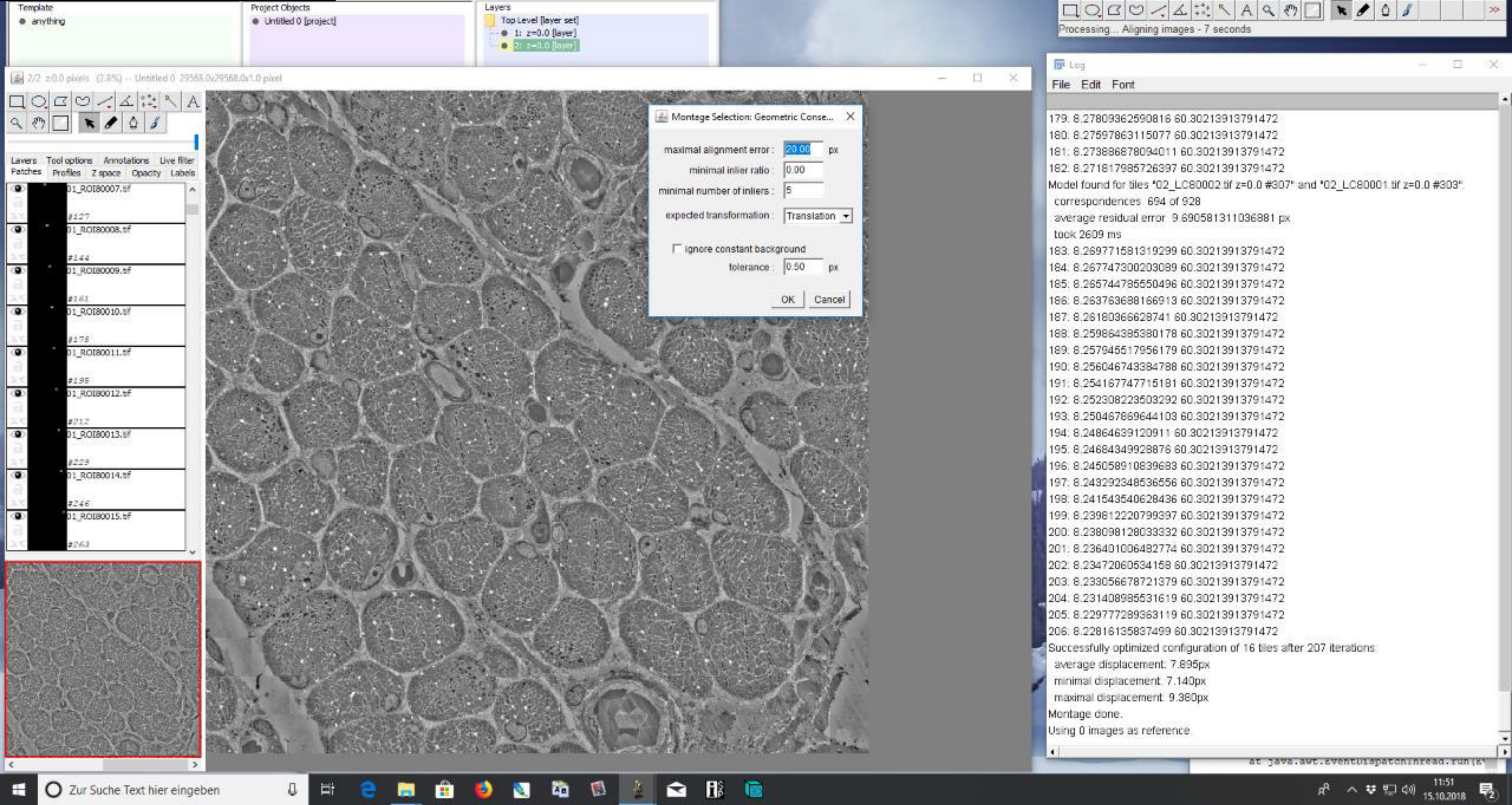

### **3)** Alignment of ROI and LC

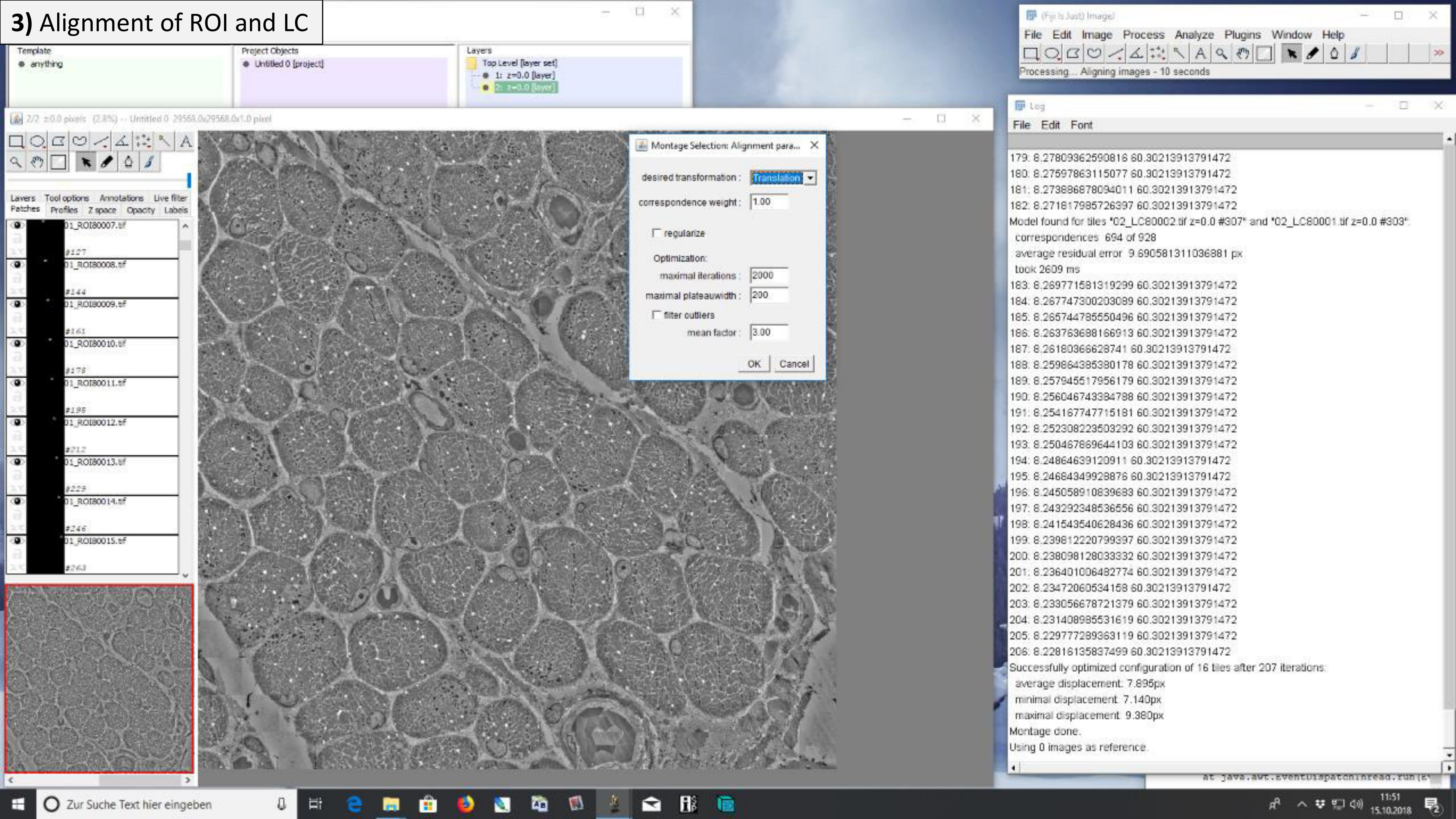

### 3) Alignment of ROI and LC

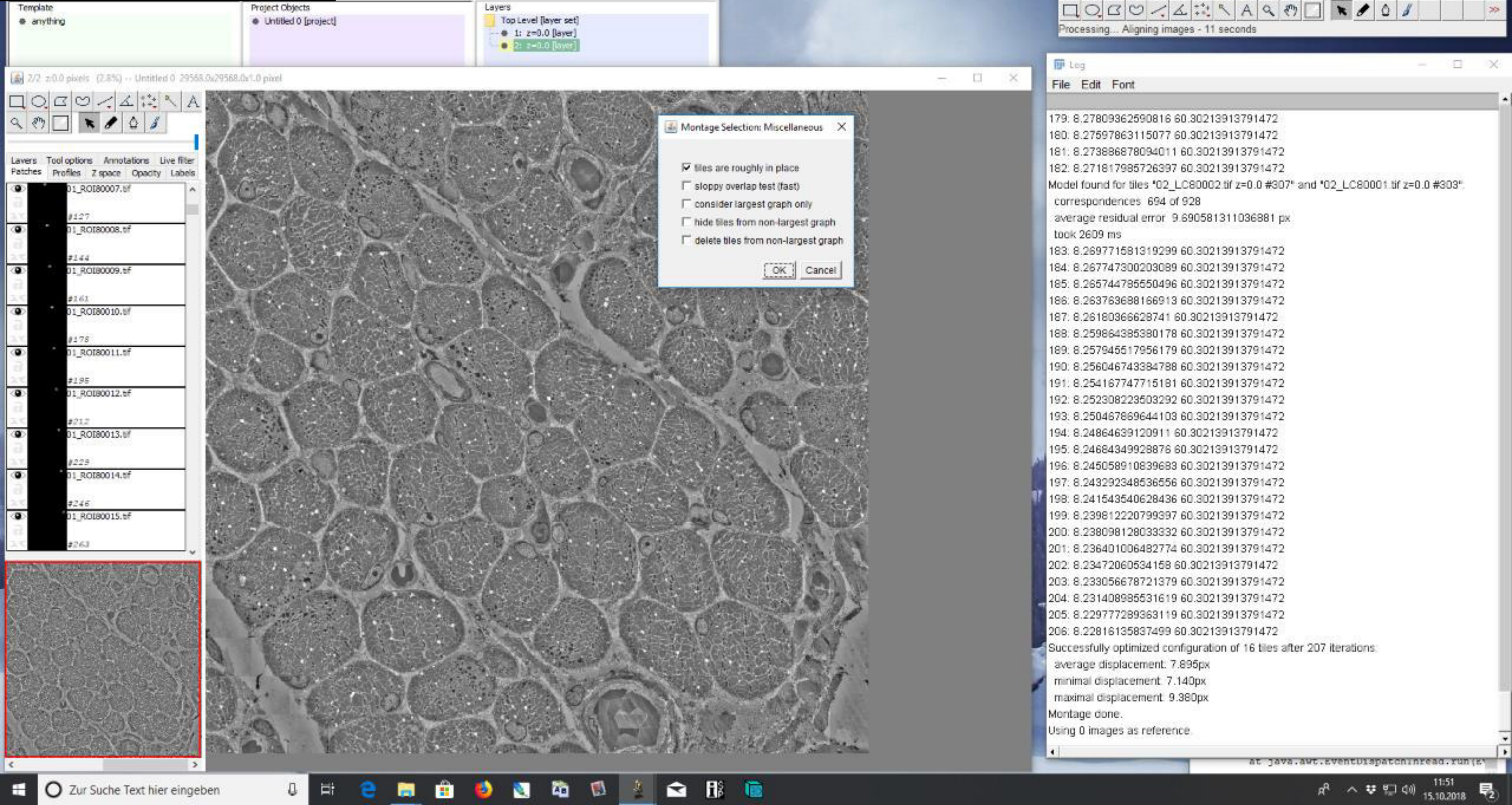

#### **3)** Alignment of ROI and LC

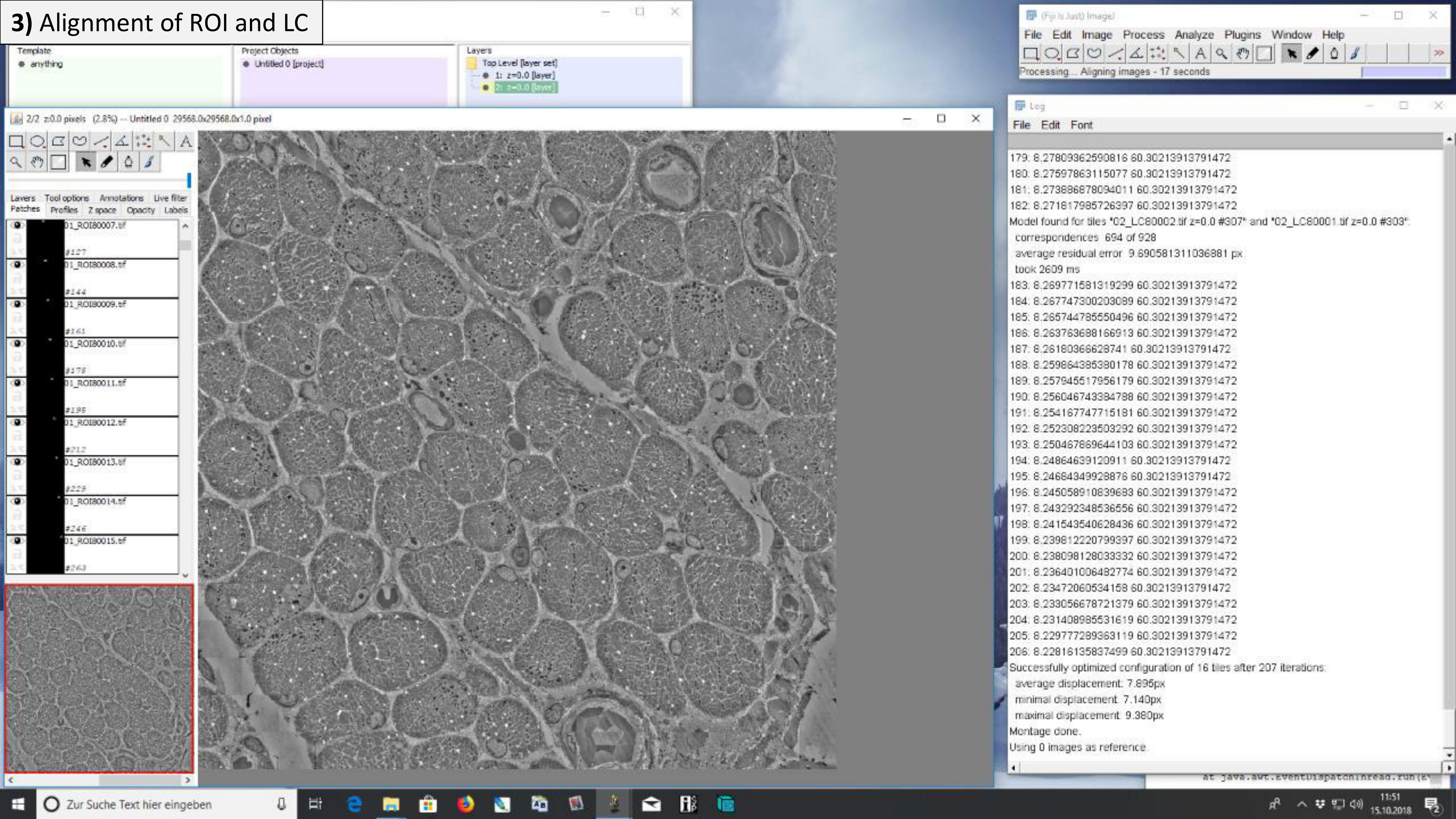

### **3)** Alignment of ROI and LC

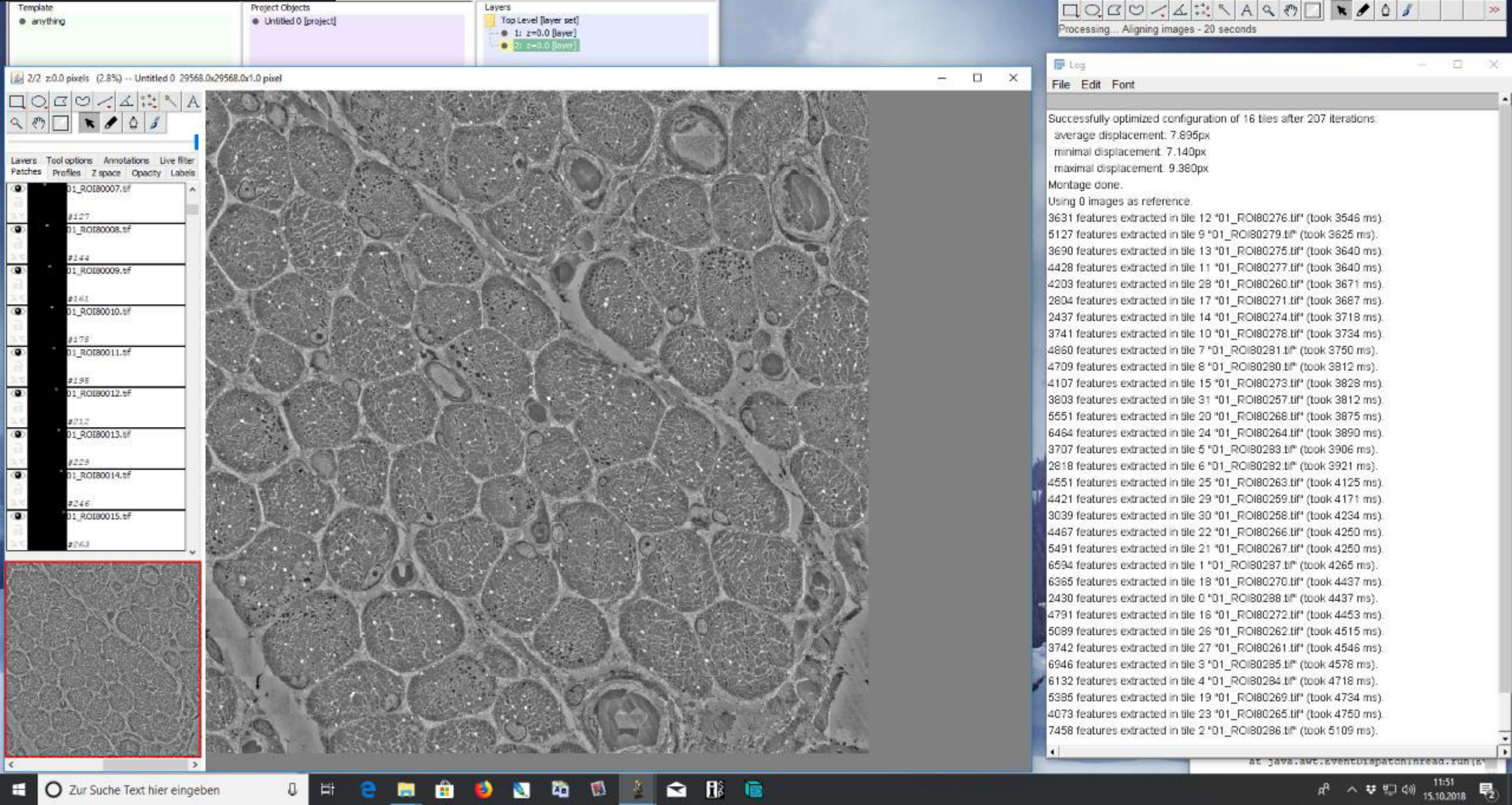

### **3)** Alignment of ROI and LC

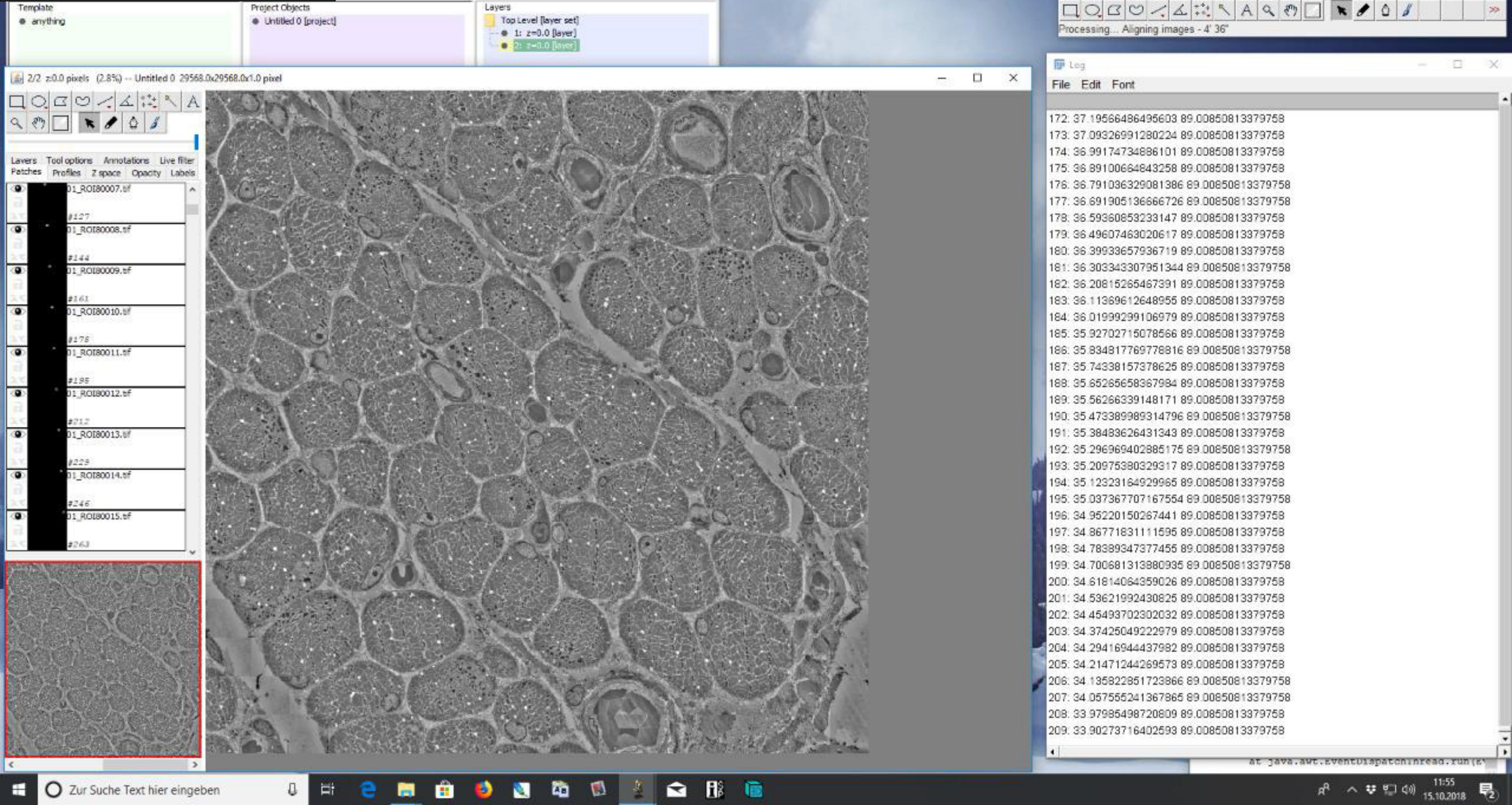

### 3) Alignment of ROI and LC

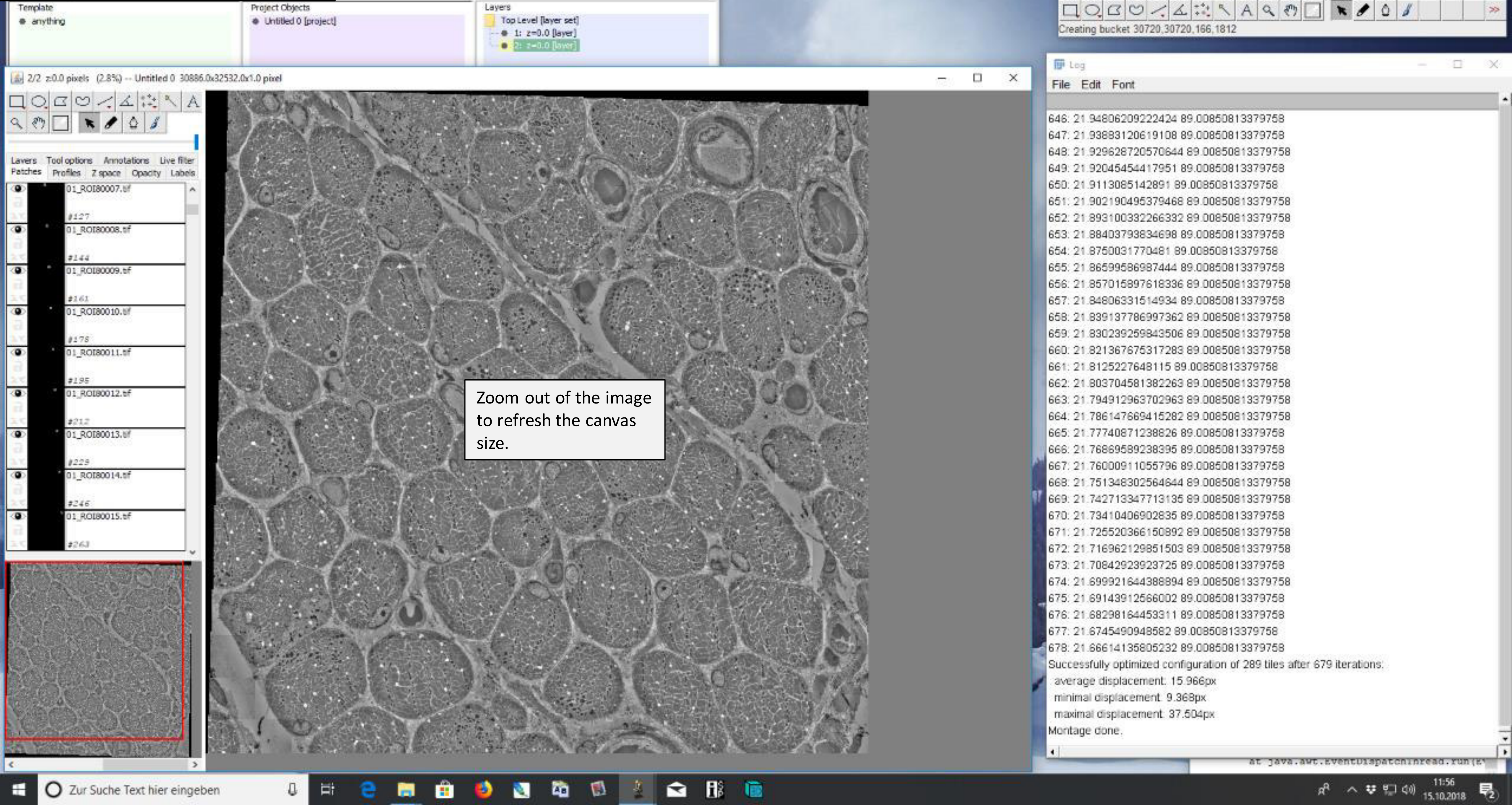


Zoom out of the image to refresh the canvas size.

### 3) Alignment of ROI and LC

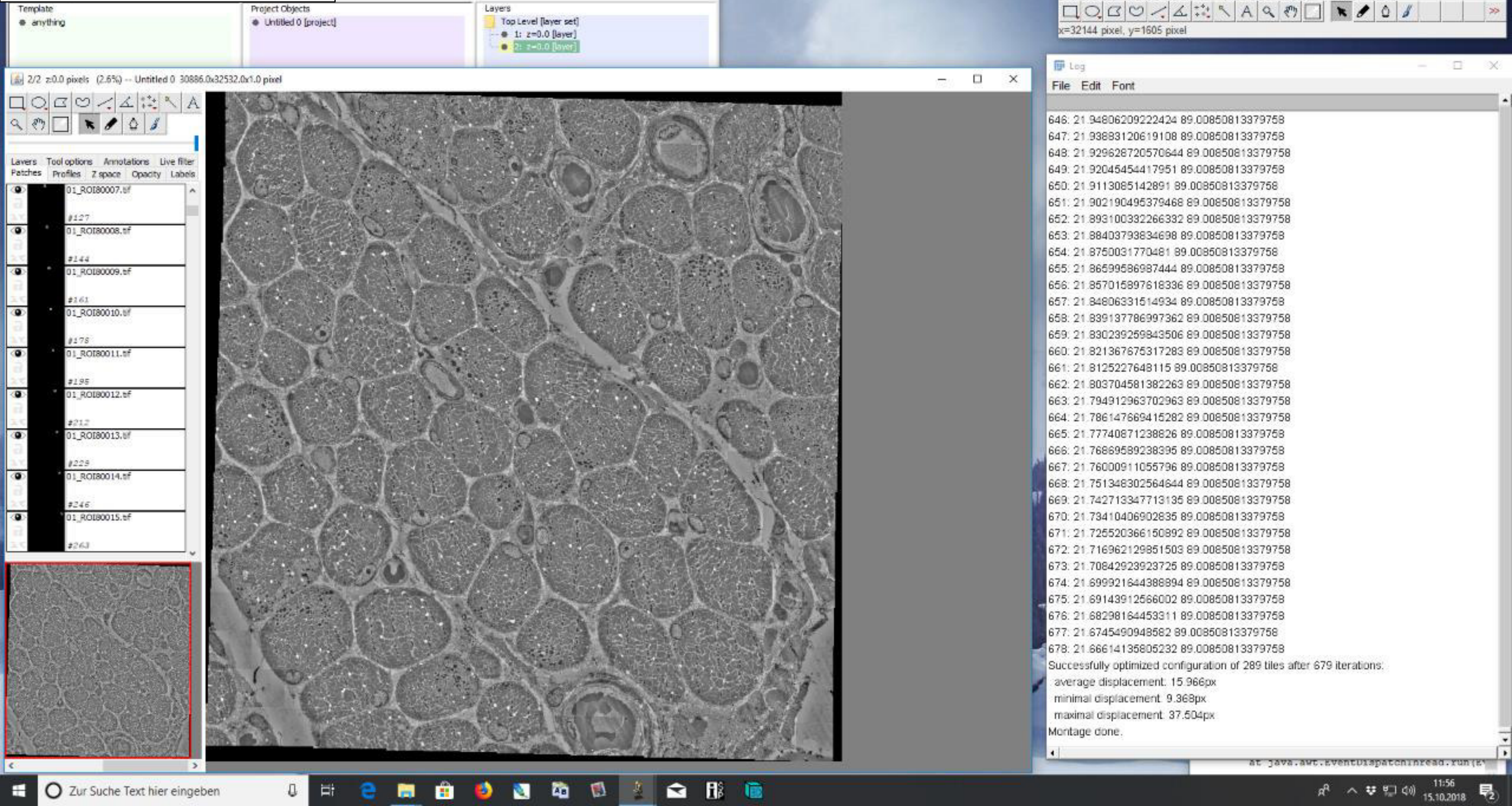

### 4) Lens correction

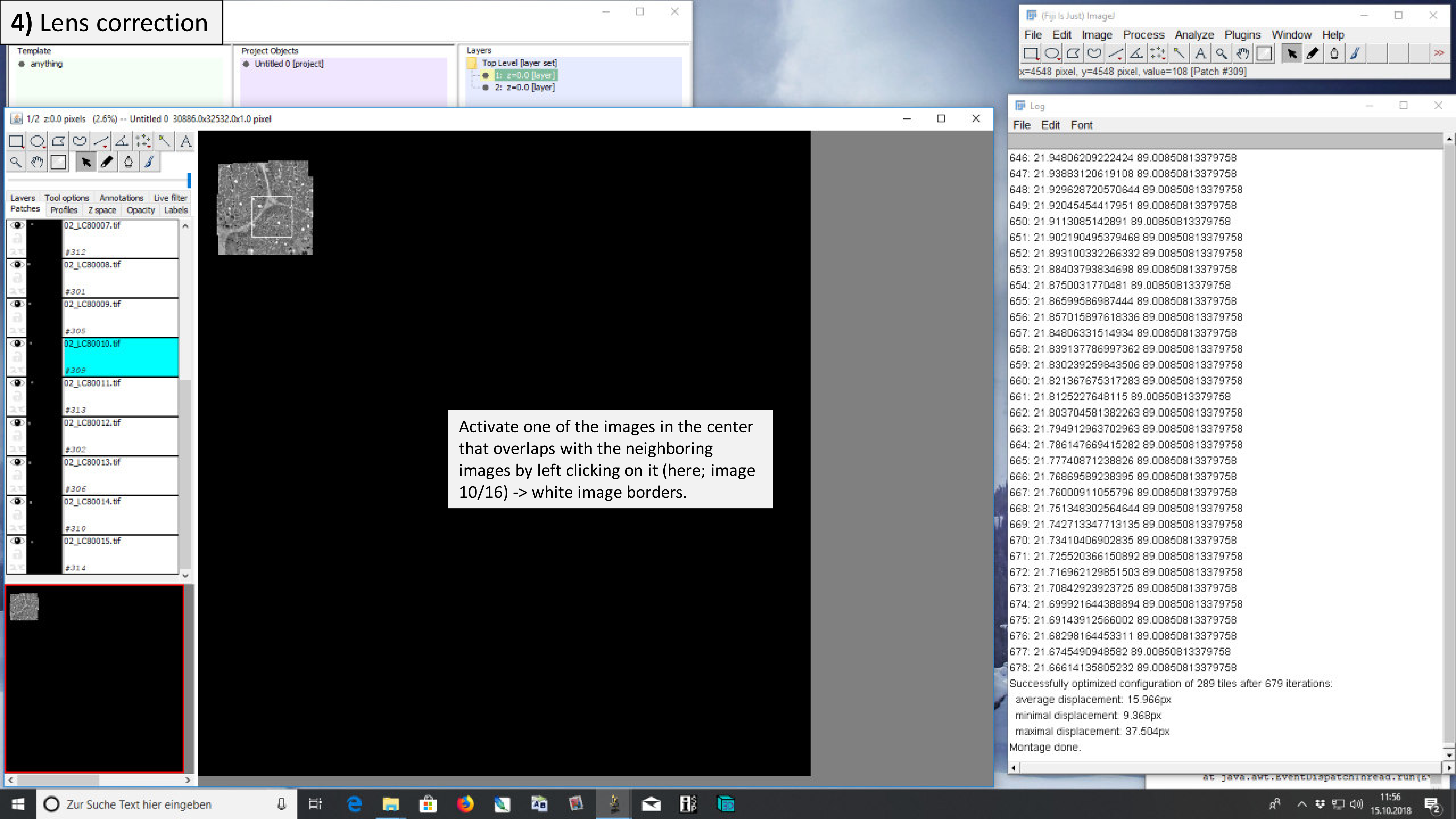


Activate one of the images in the center that overlaps with the neighboring images by left clicking on it (here; image 10/16) -> white image borders.

### 4) Lens correction

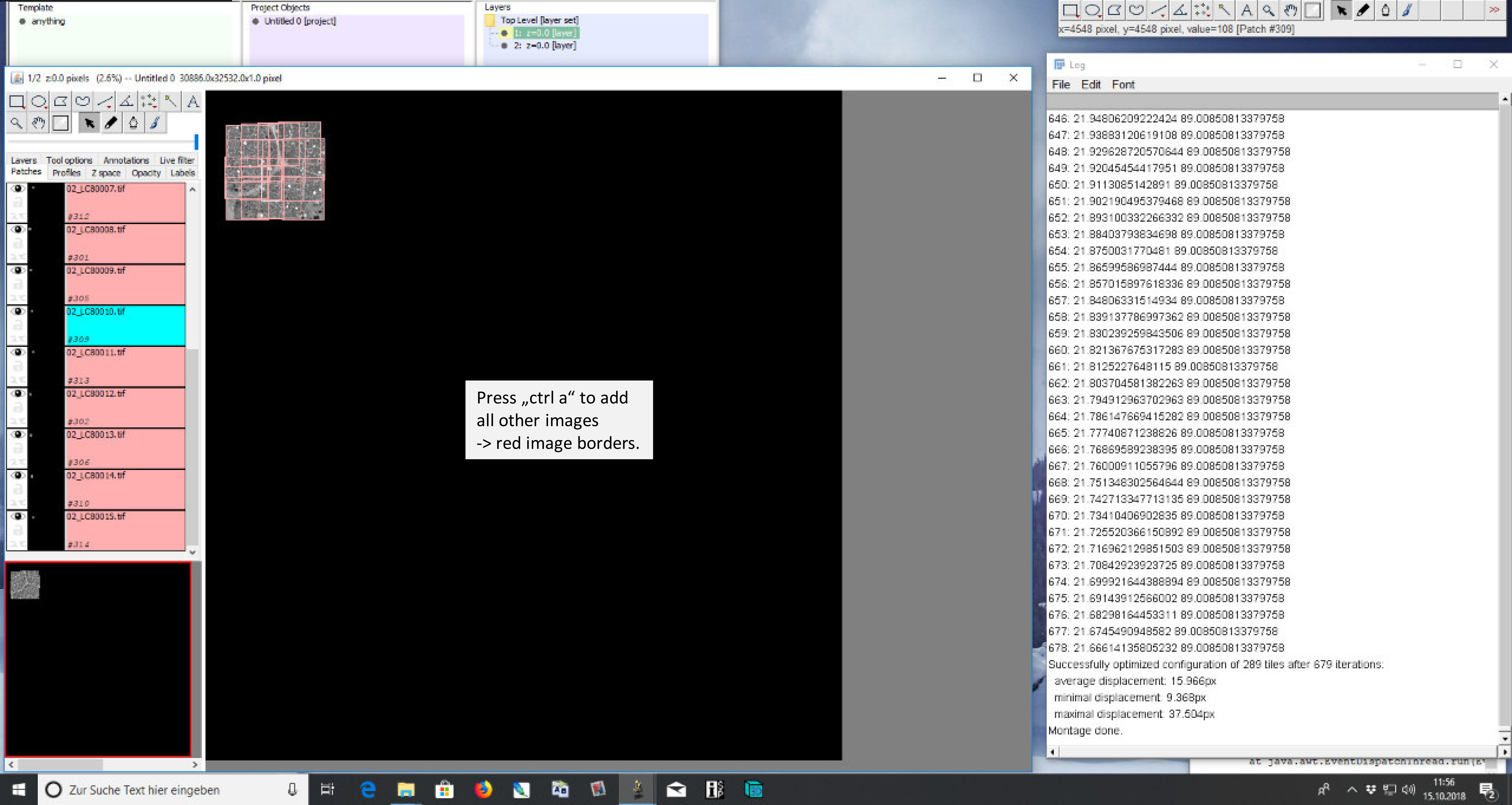

### 4) Lens correction

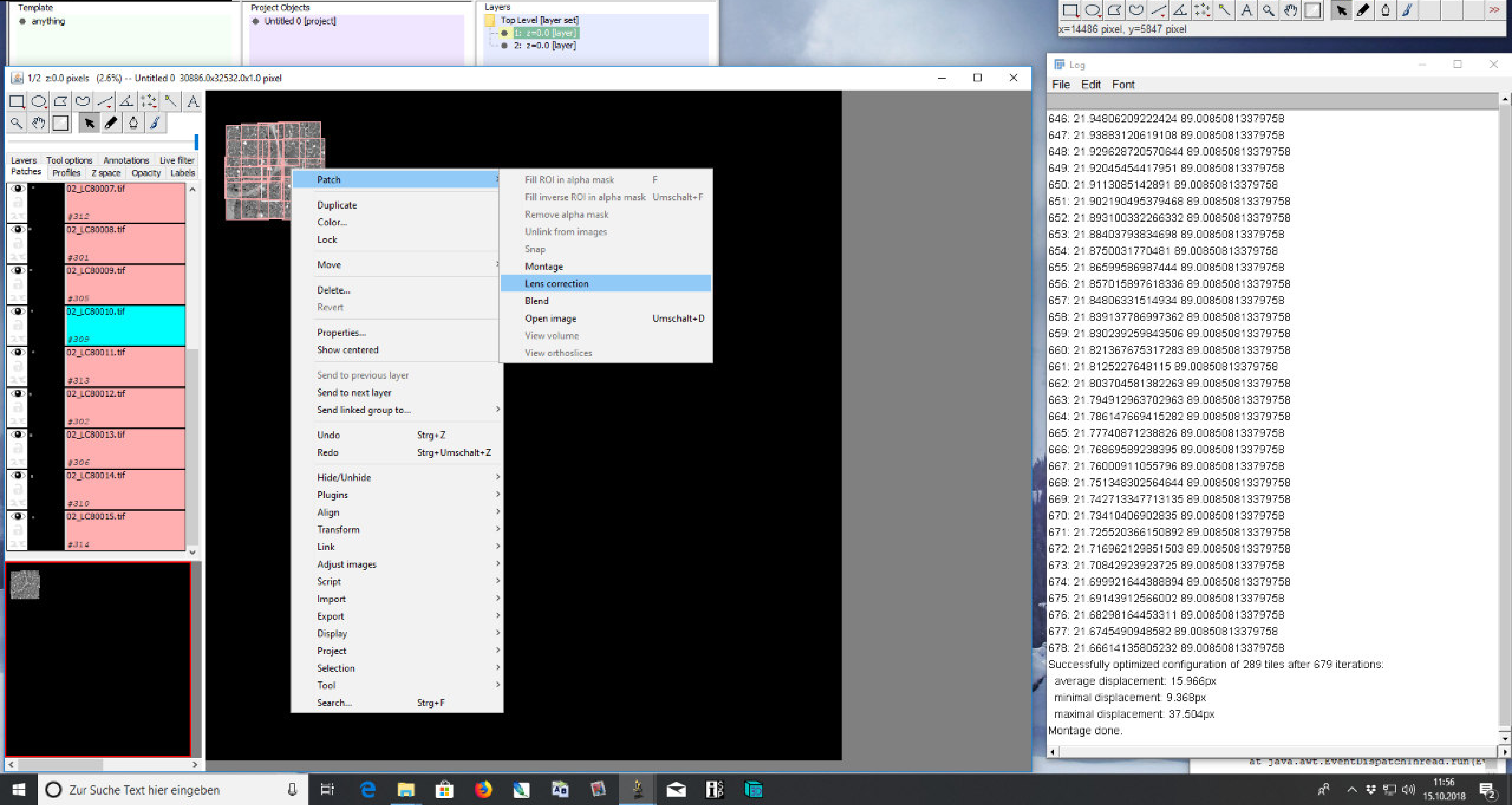

### 4) Lens correction

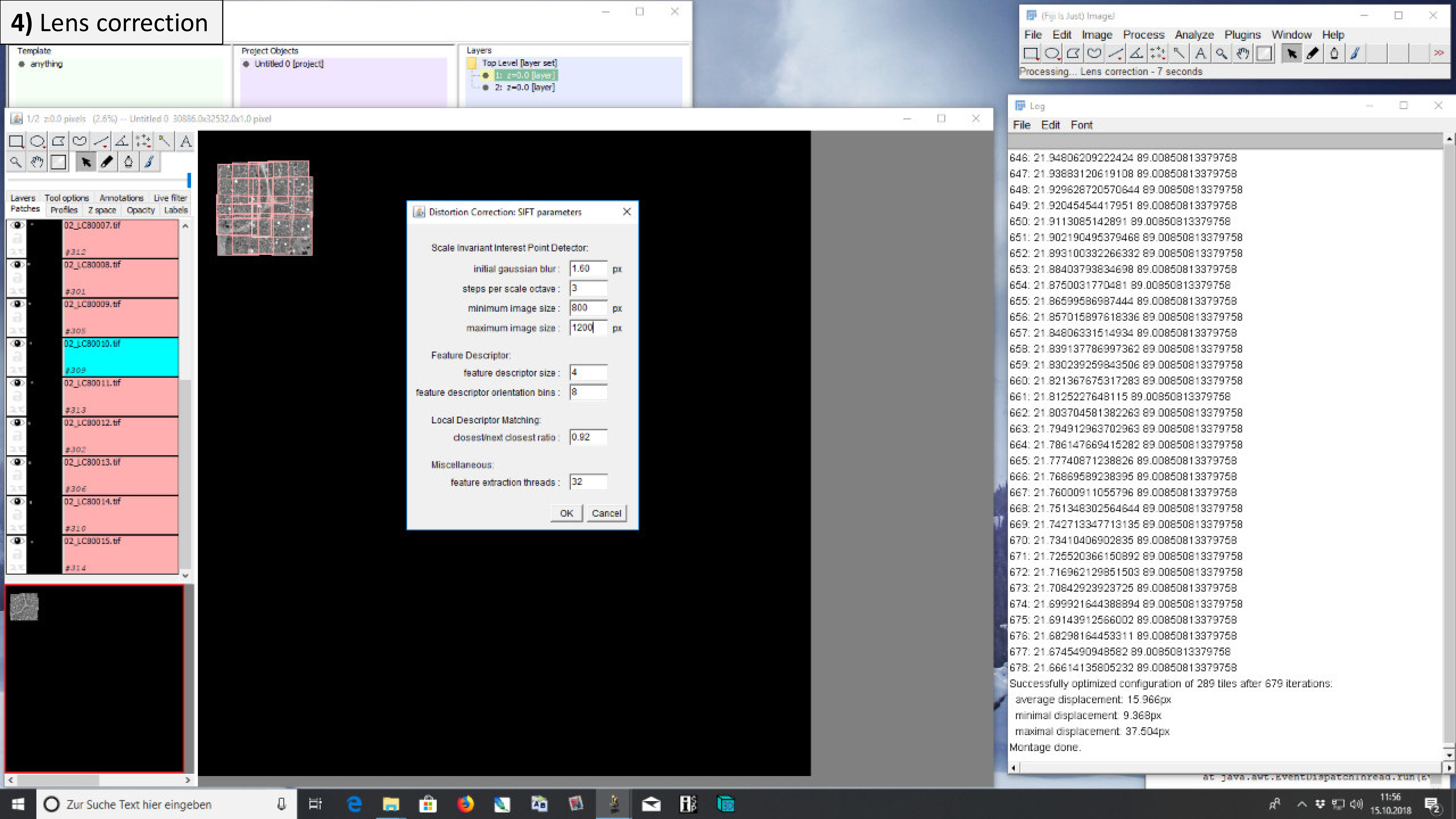

### 4) Lens correction

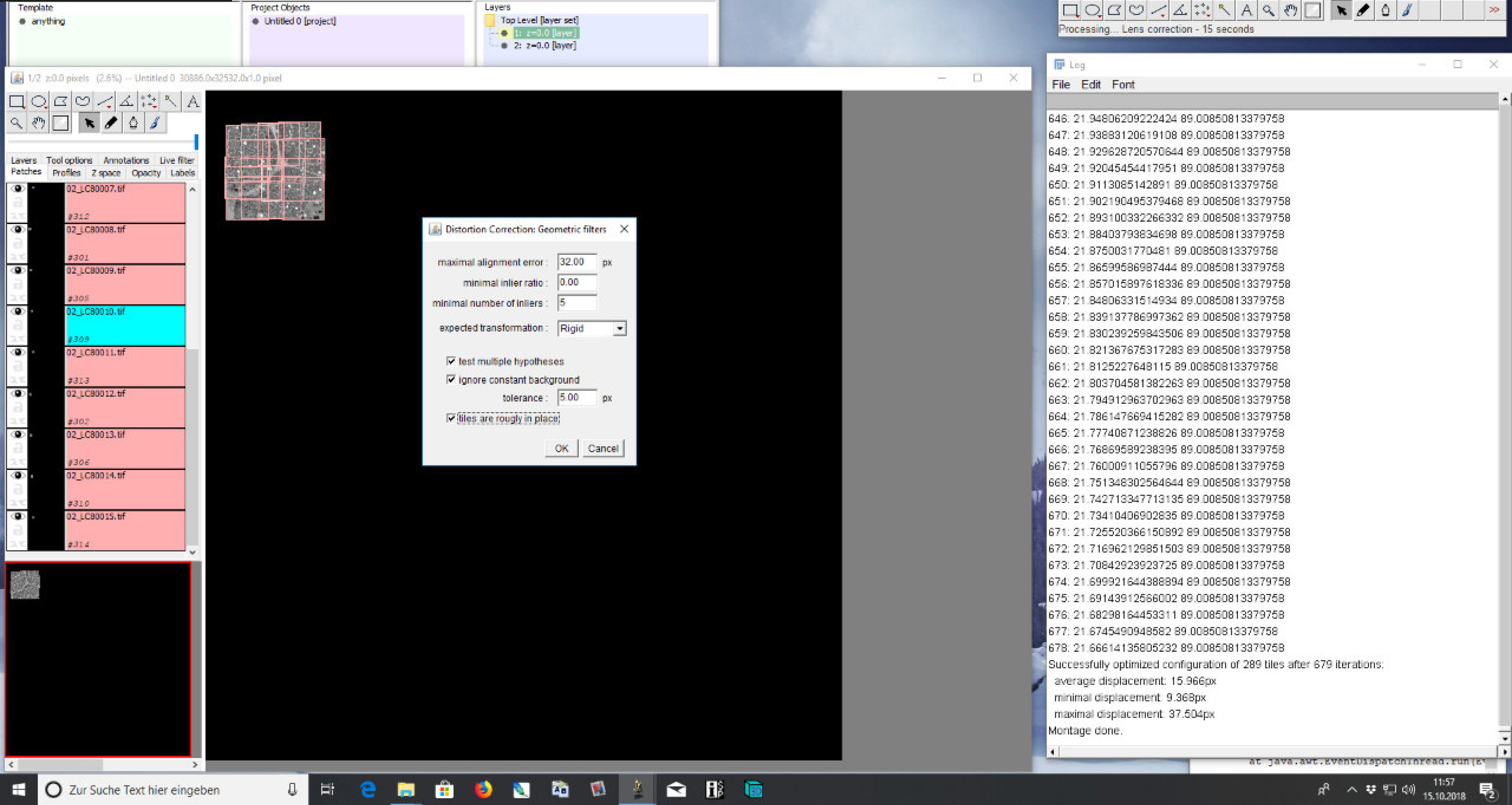

### 4) Lens correction

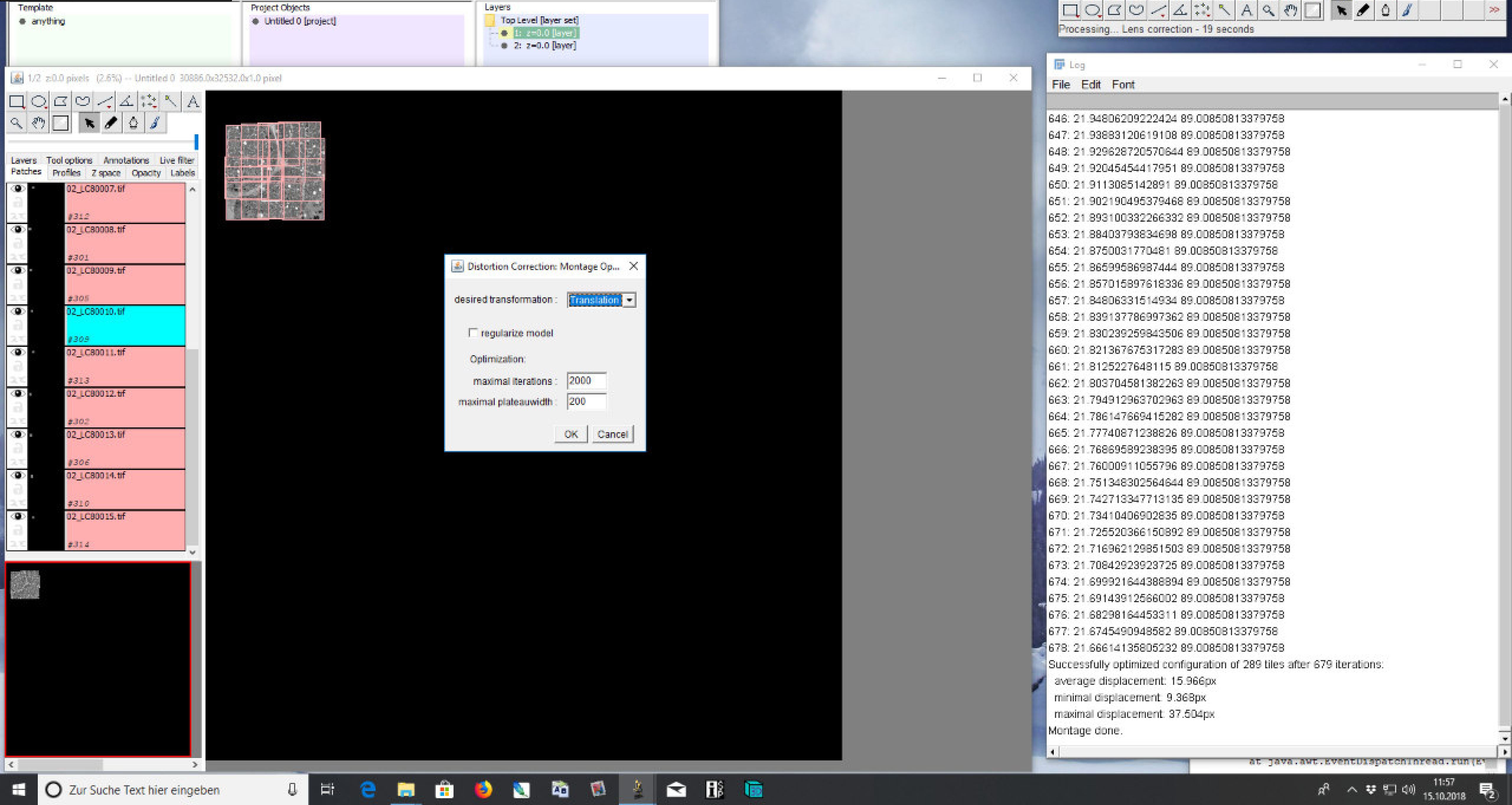

### 4) Lens correction

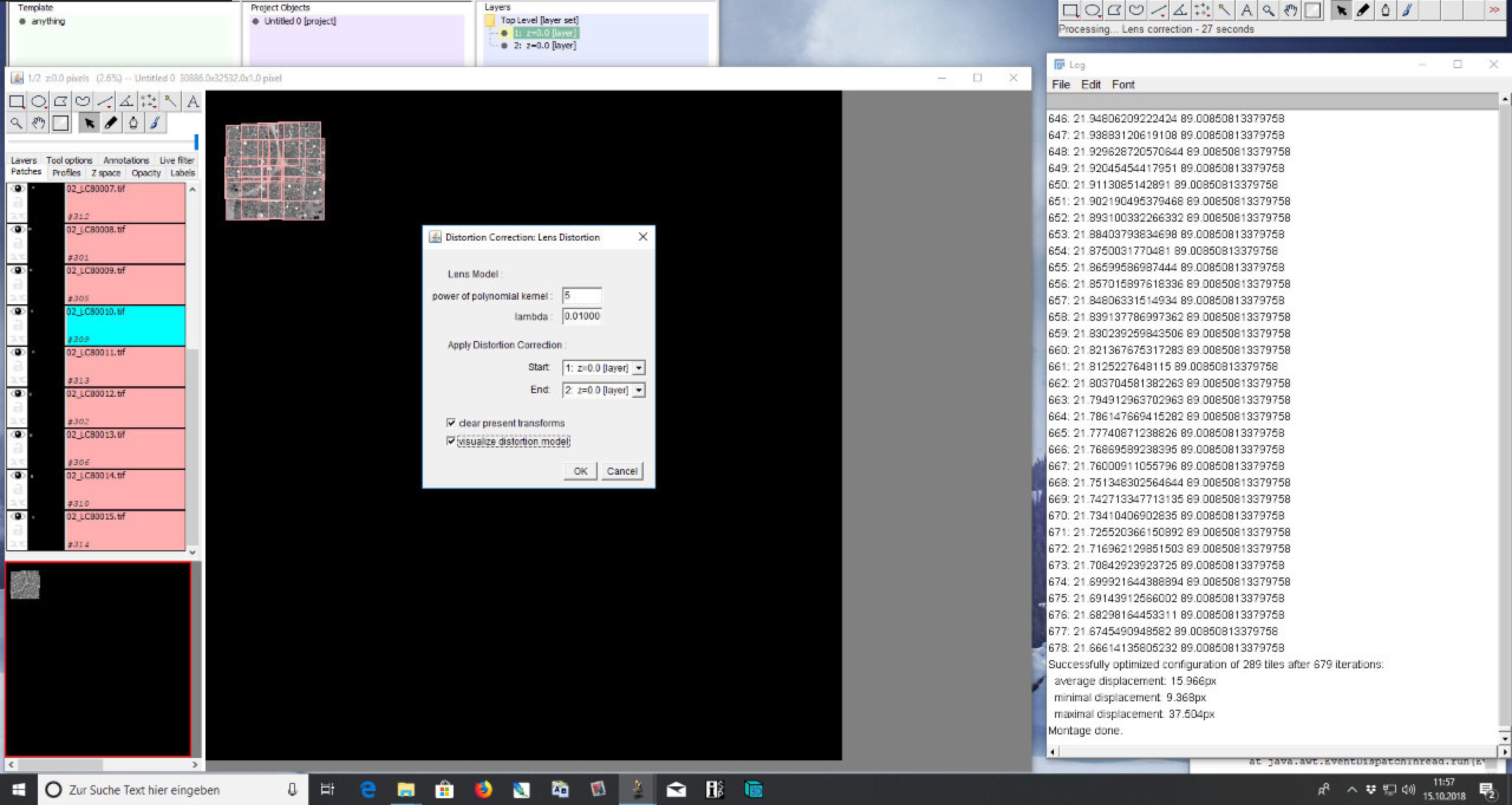

### 4) Lens correction

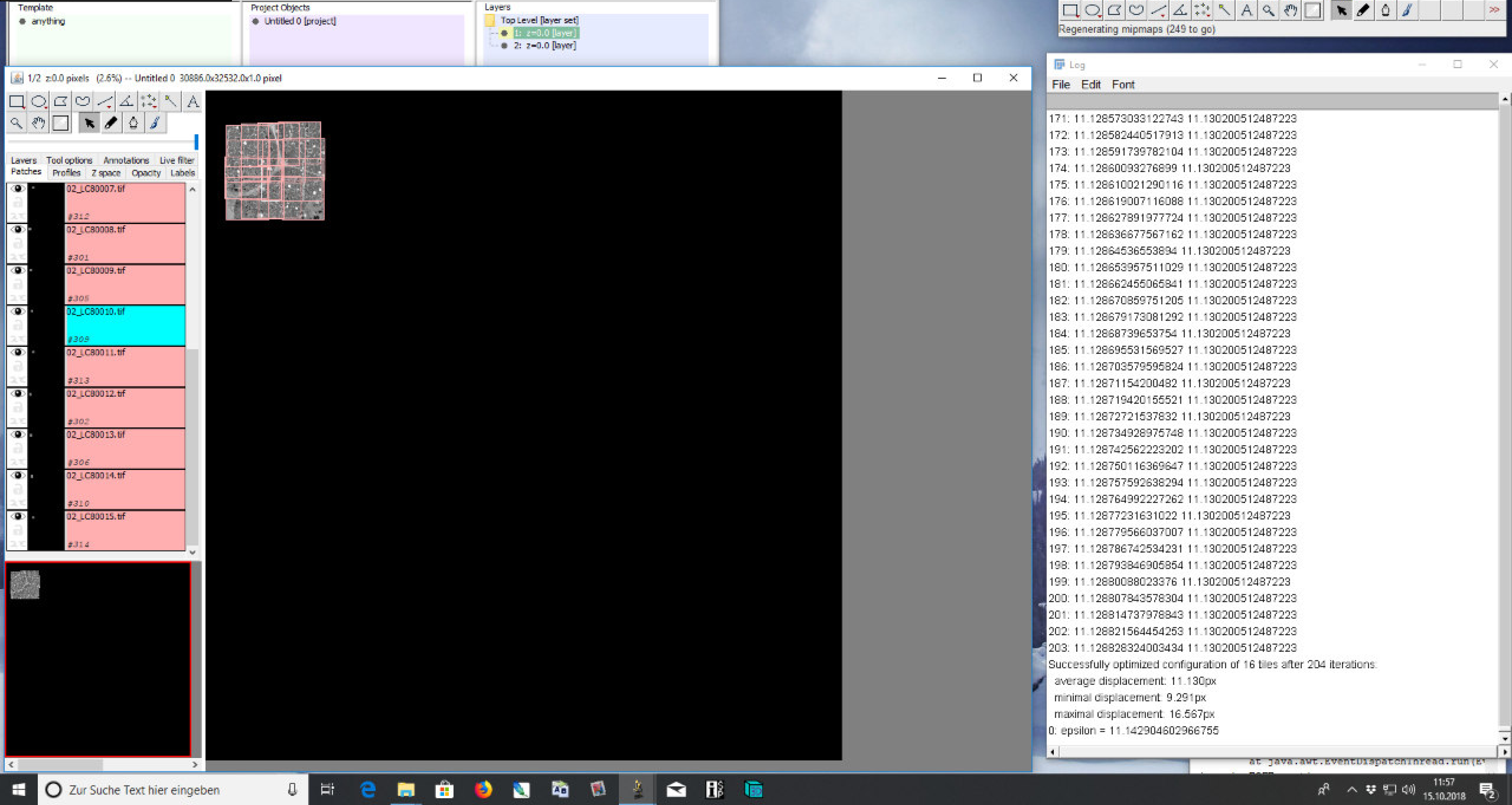

### 4) Lens correction

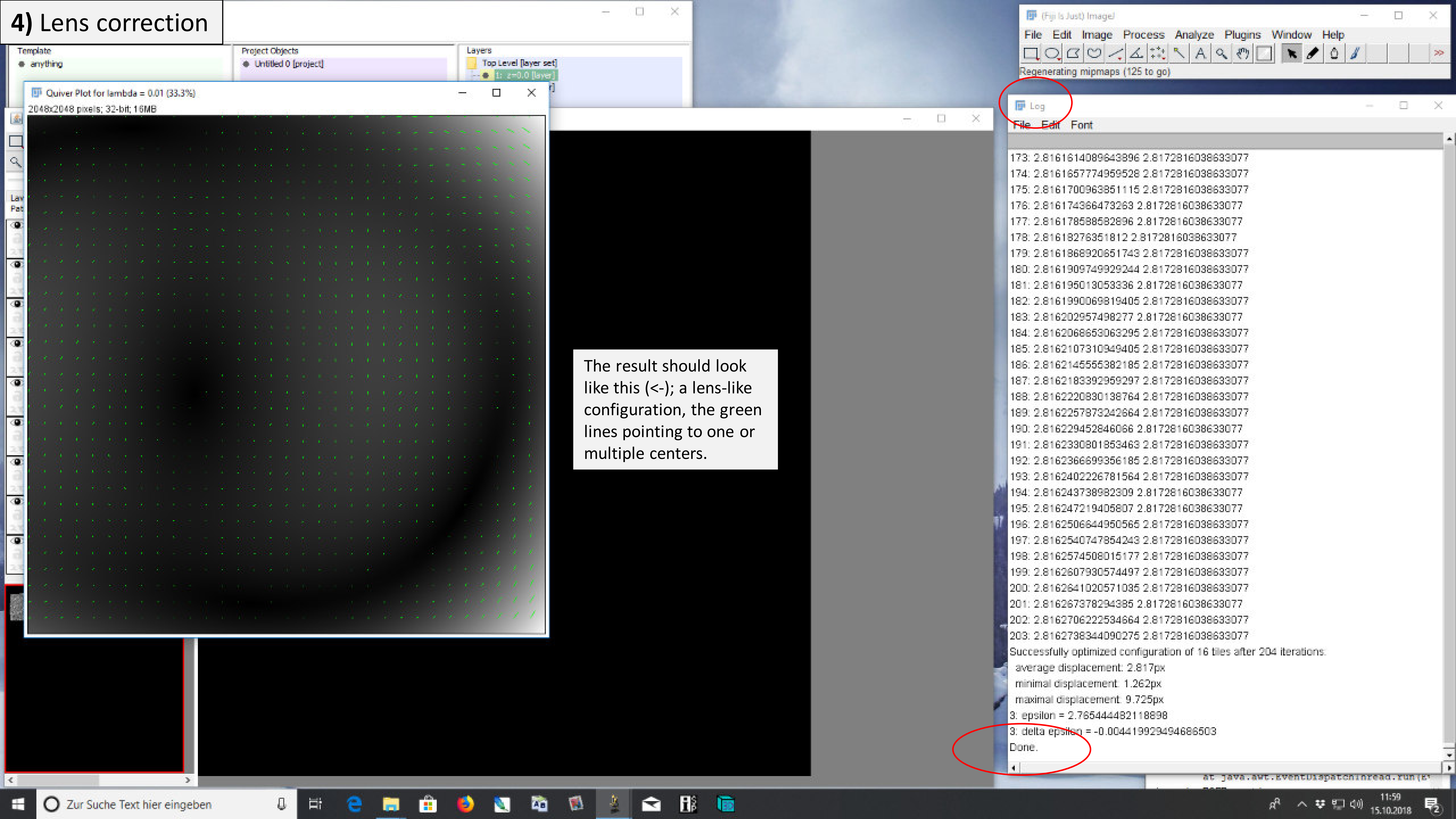


The result should look like this (<-); a lens-like configuration, the green lines pointing to one or multiple centers.

### 4) Lens correction

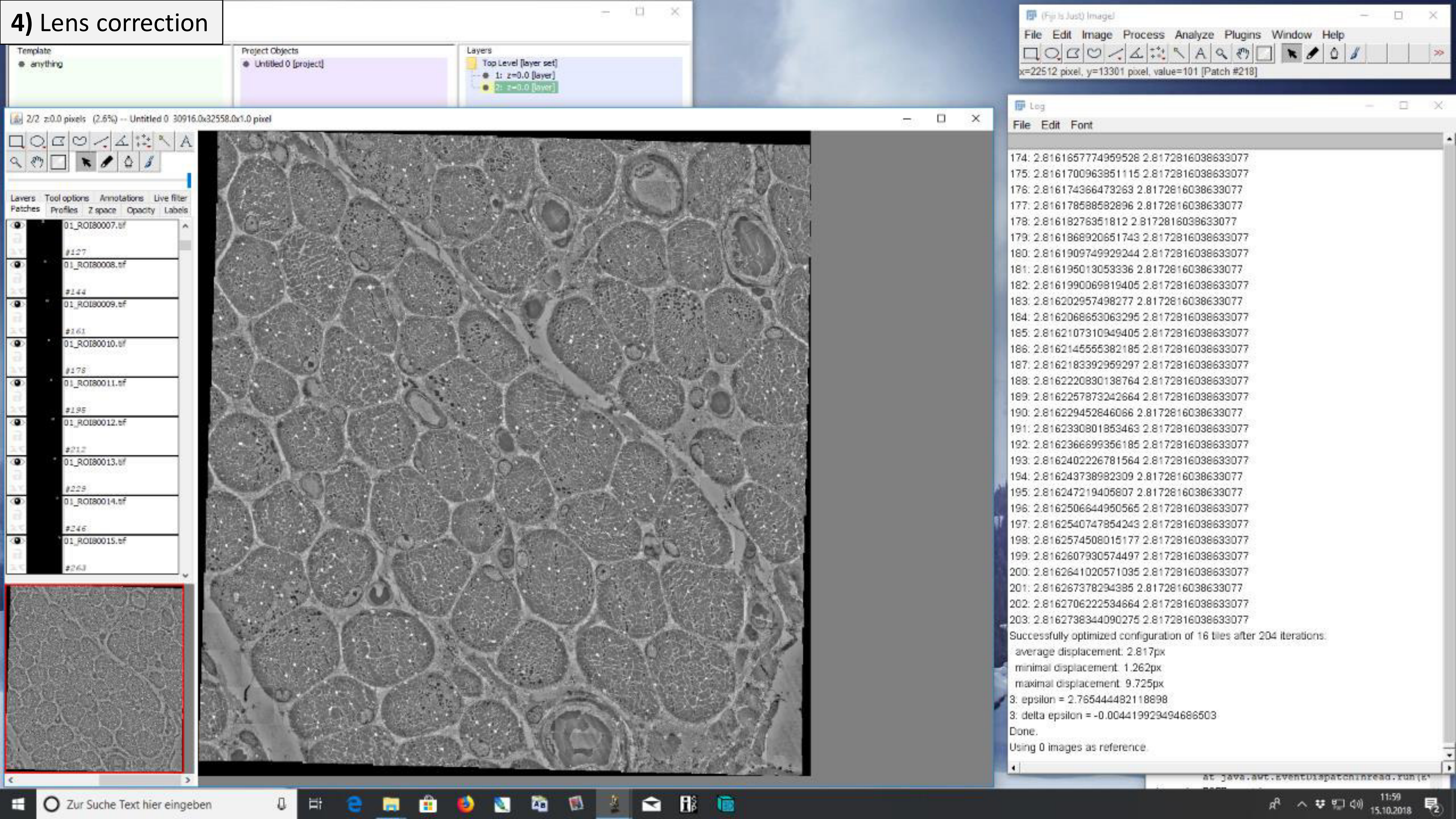

### 4) Lens correction

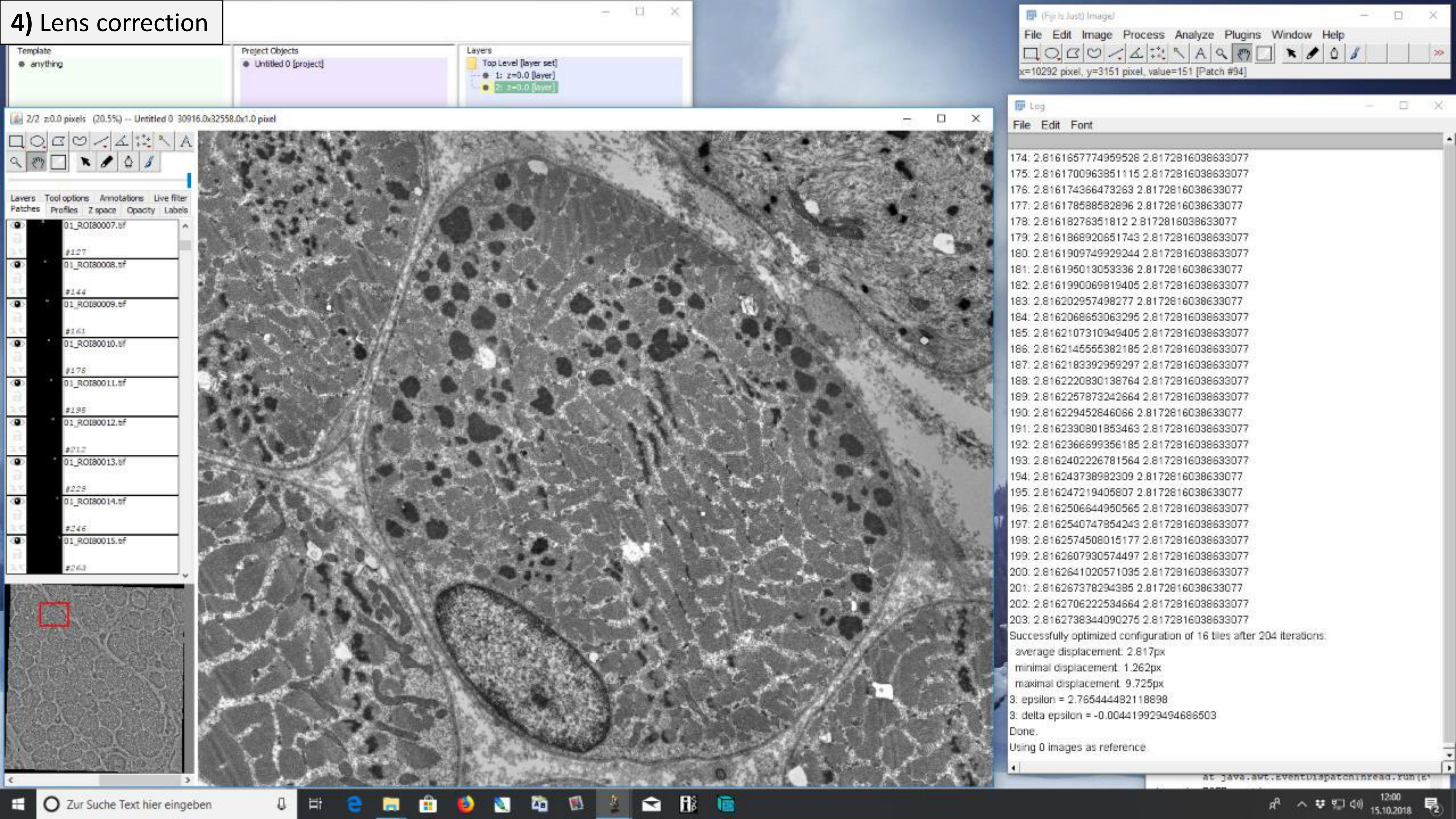

### 4) Lens correction

After lens correction, the correction must be applied by another alignment.

Before: Bad alignment

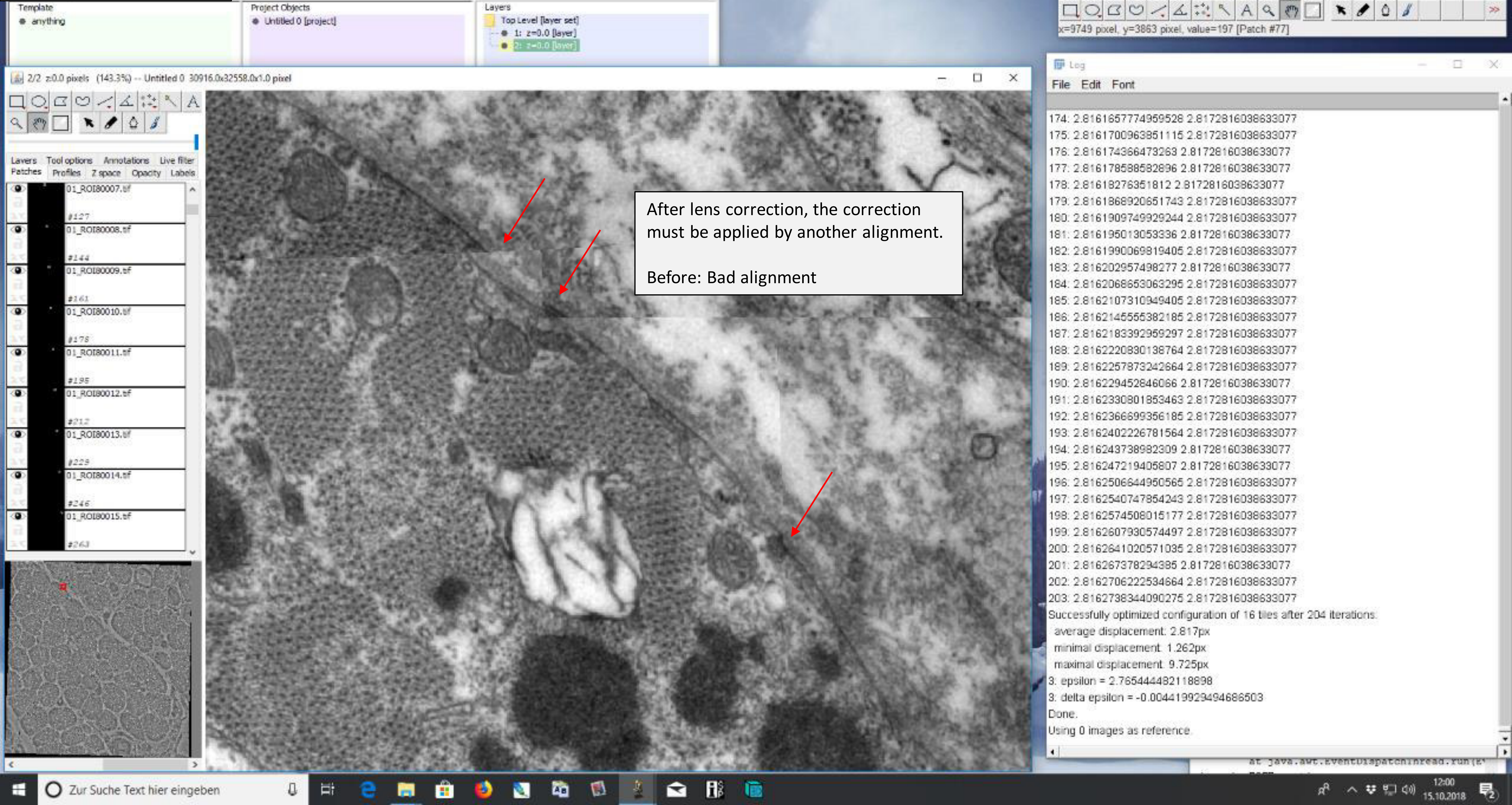

### 5) Alignment after lens correction

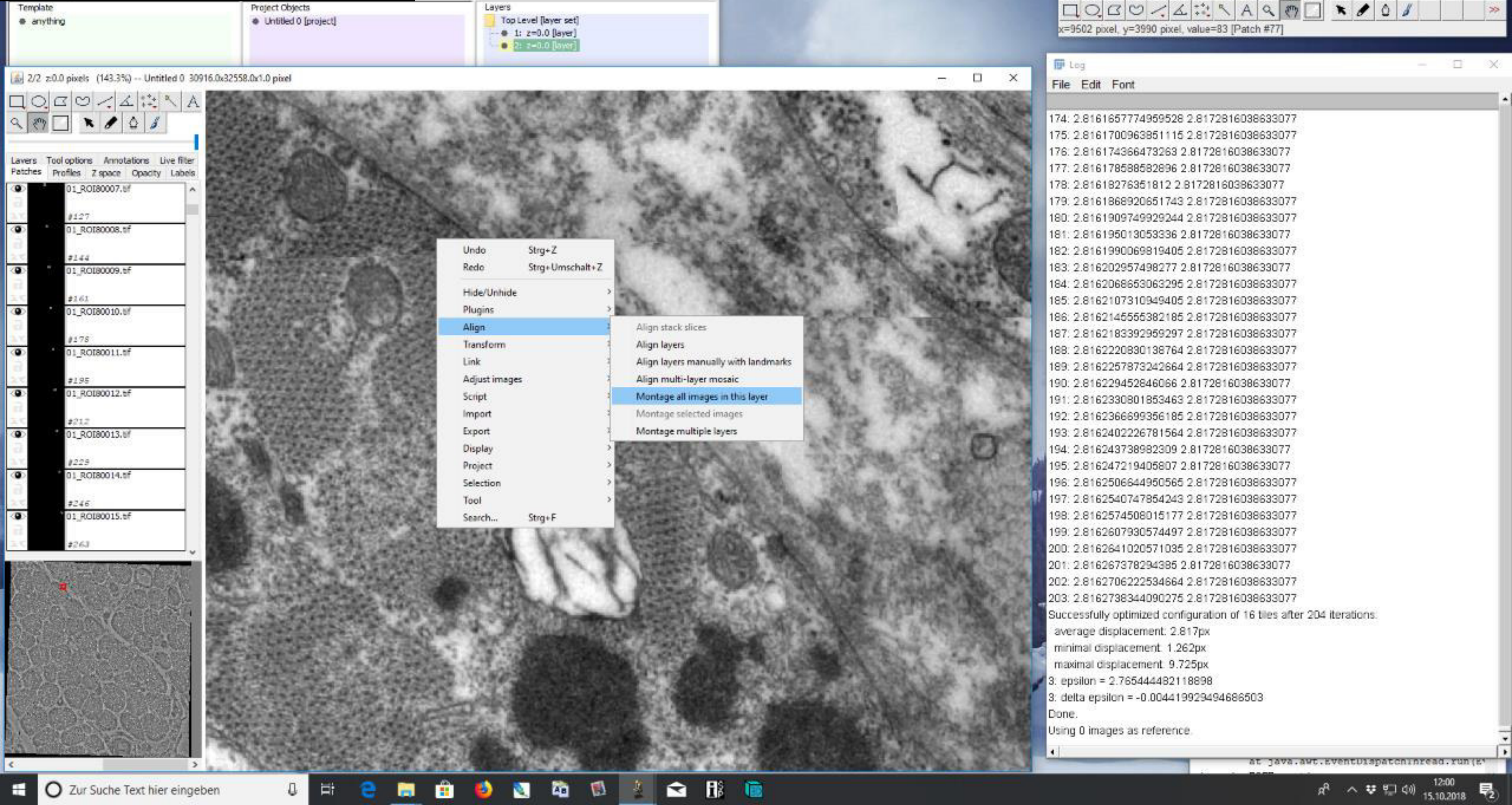

#### 5) Alignment after lens correction

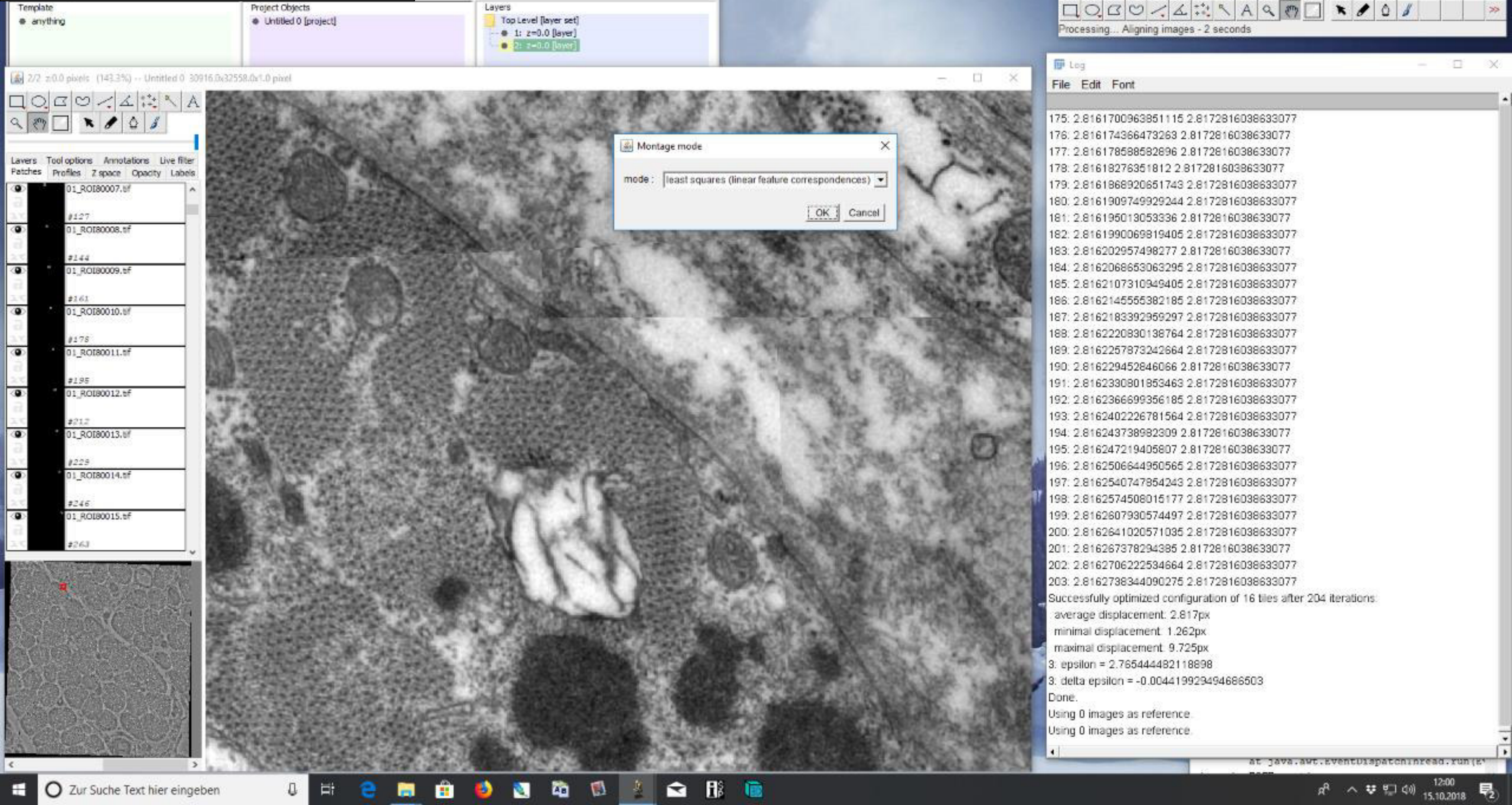

### 5) Alignment after lens correction

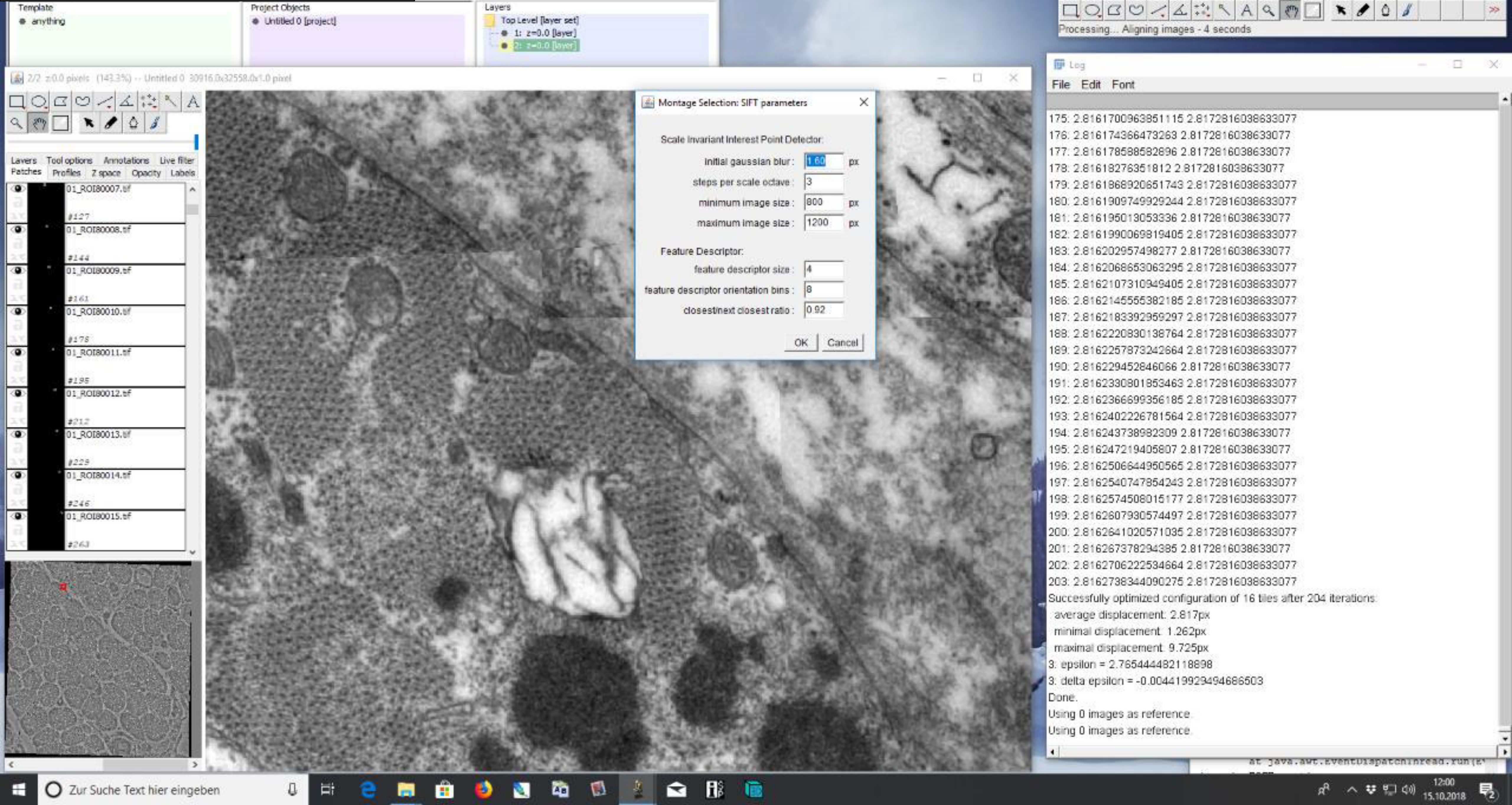

### **5)** Alignment after lens correction

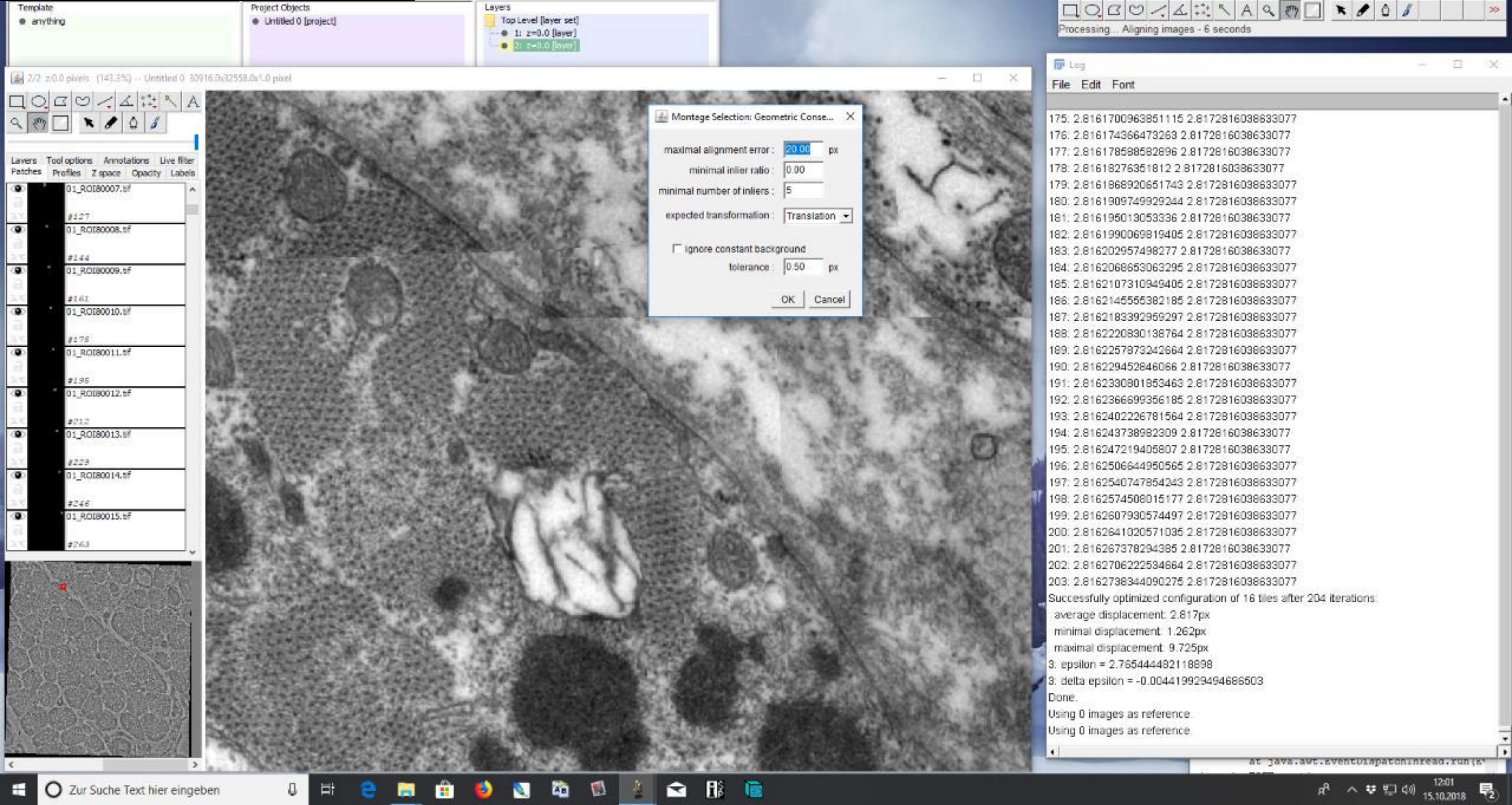

### 5) Alignment after lens correction

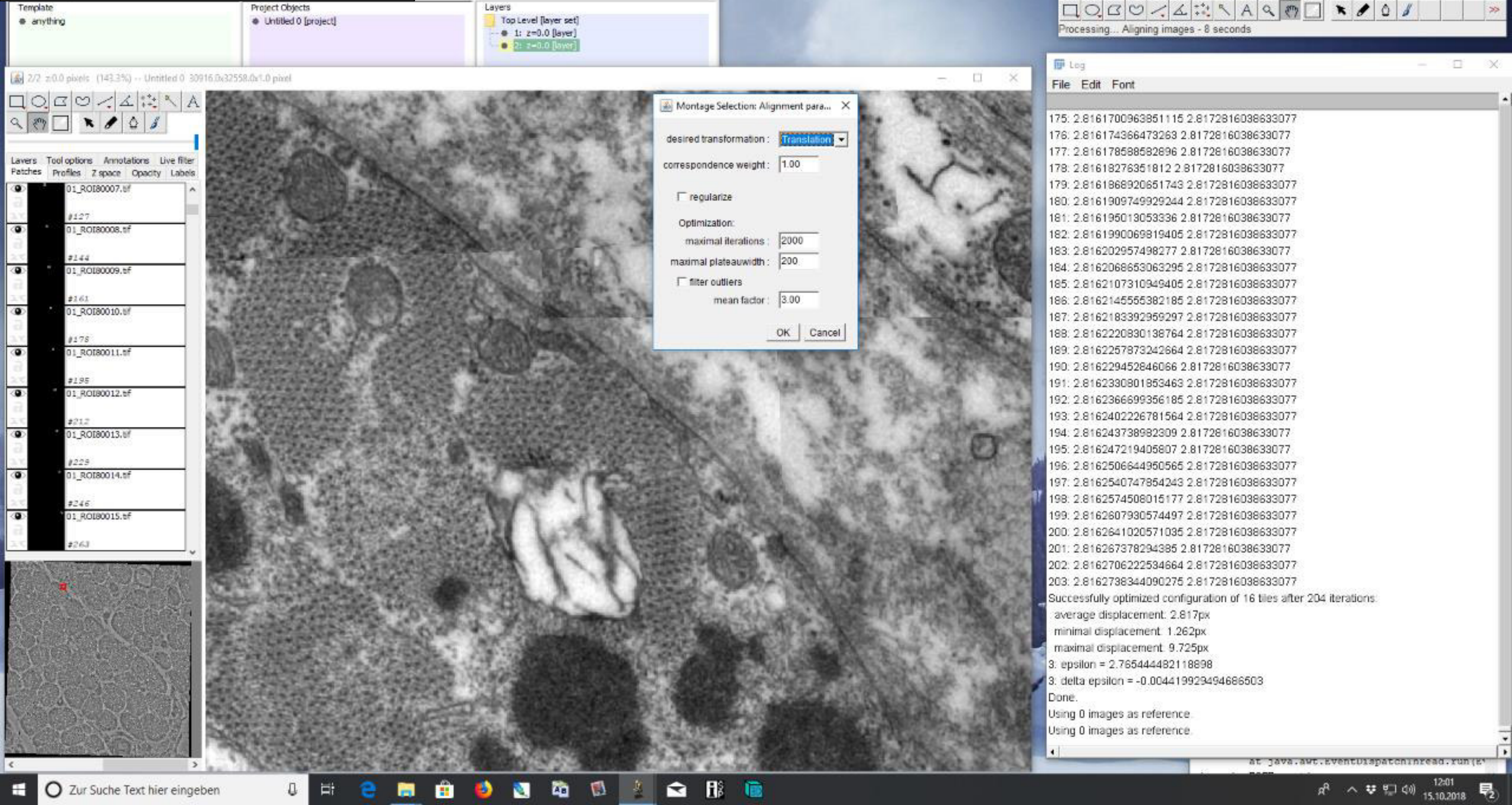

### 5) Alignment after lens correction

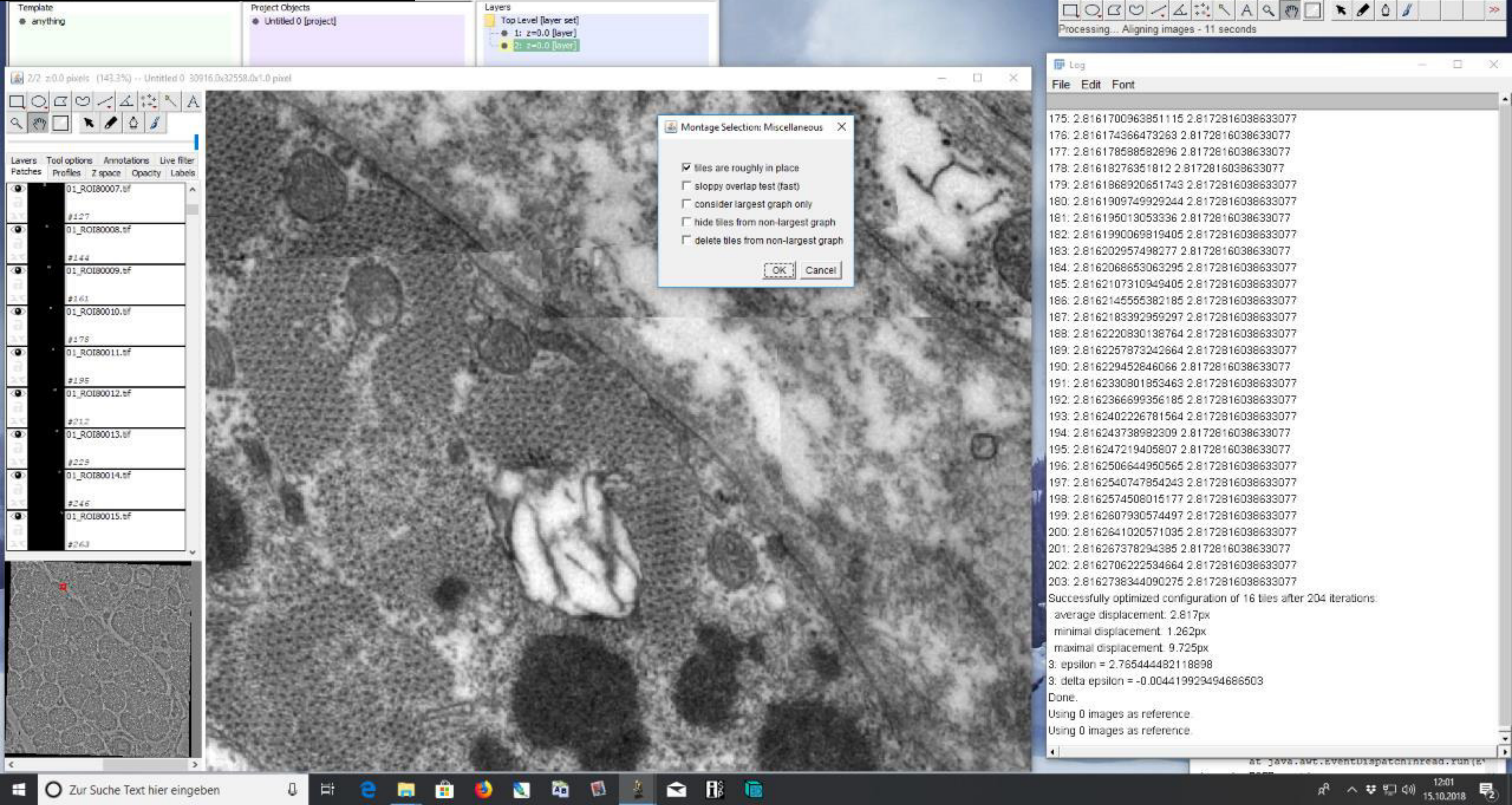

### 5) Alignment after lens correction

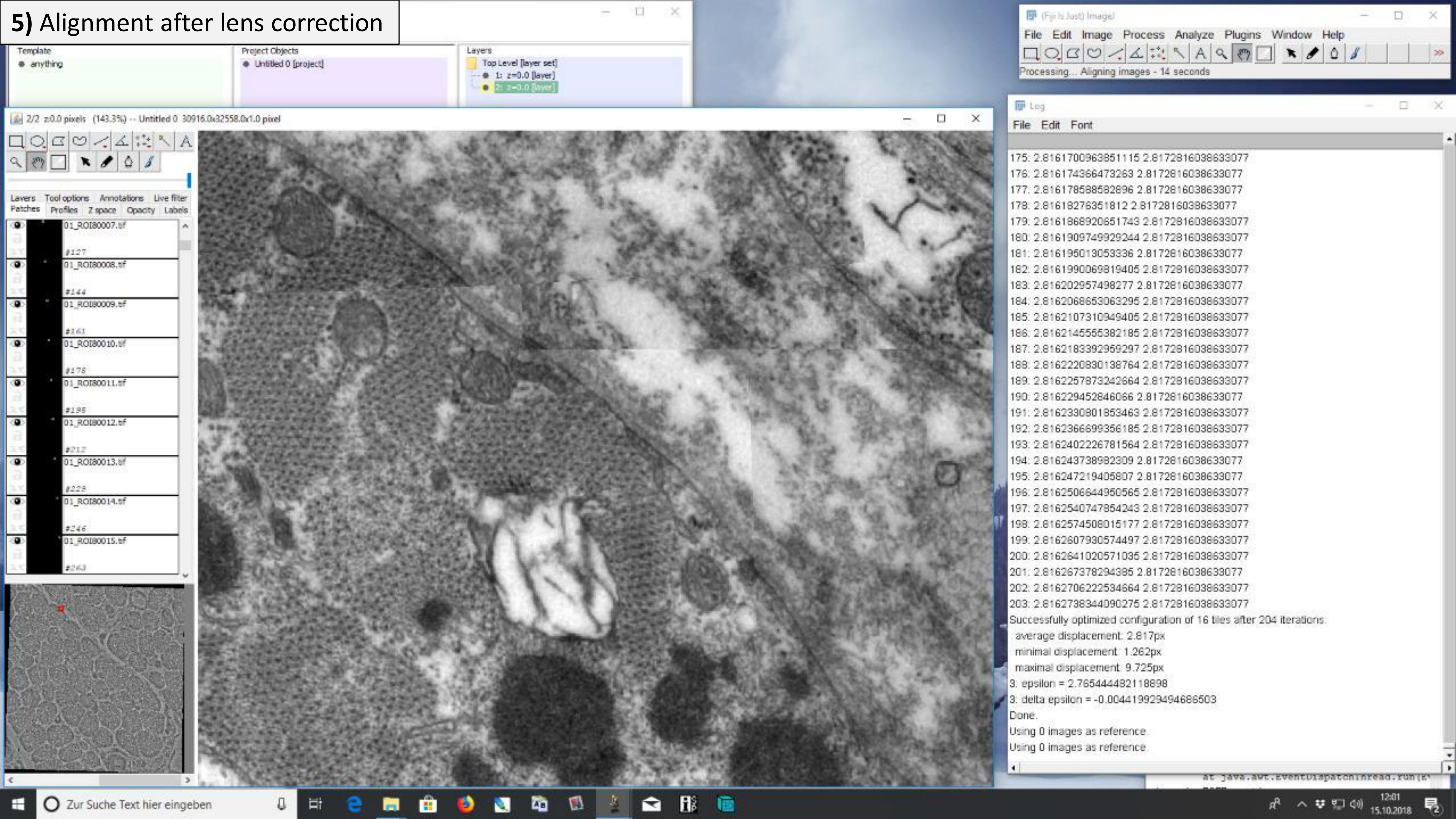

### 5) Alignment after lens correction

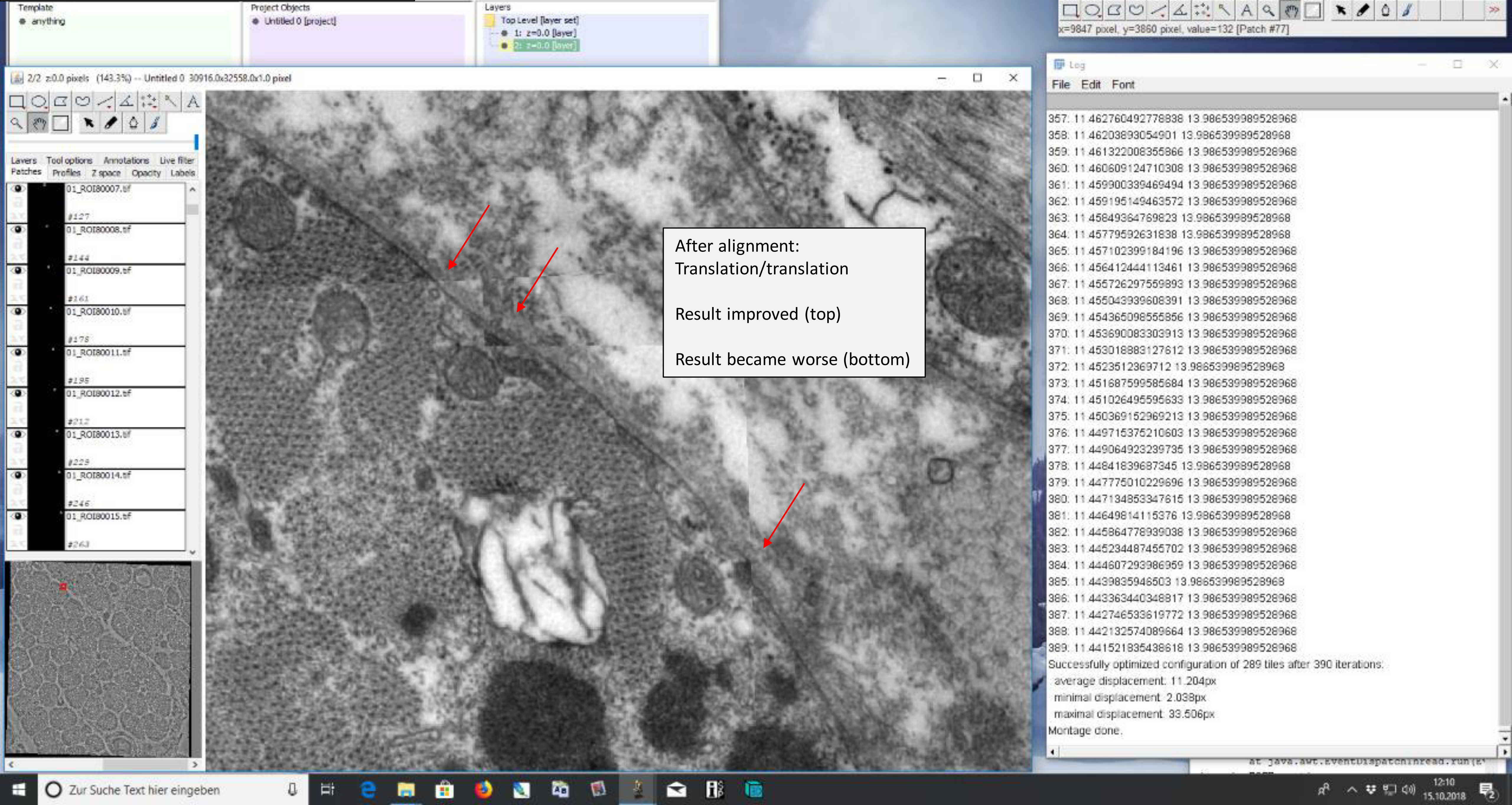

### **5)** Alignment after lens correction

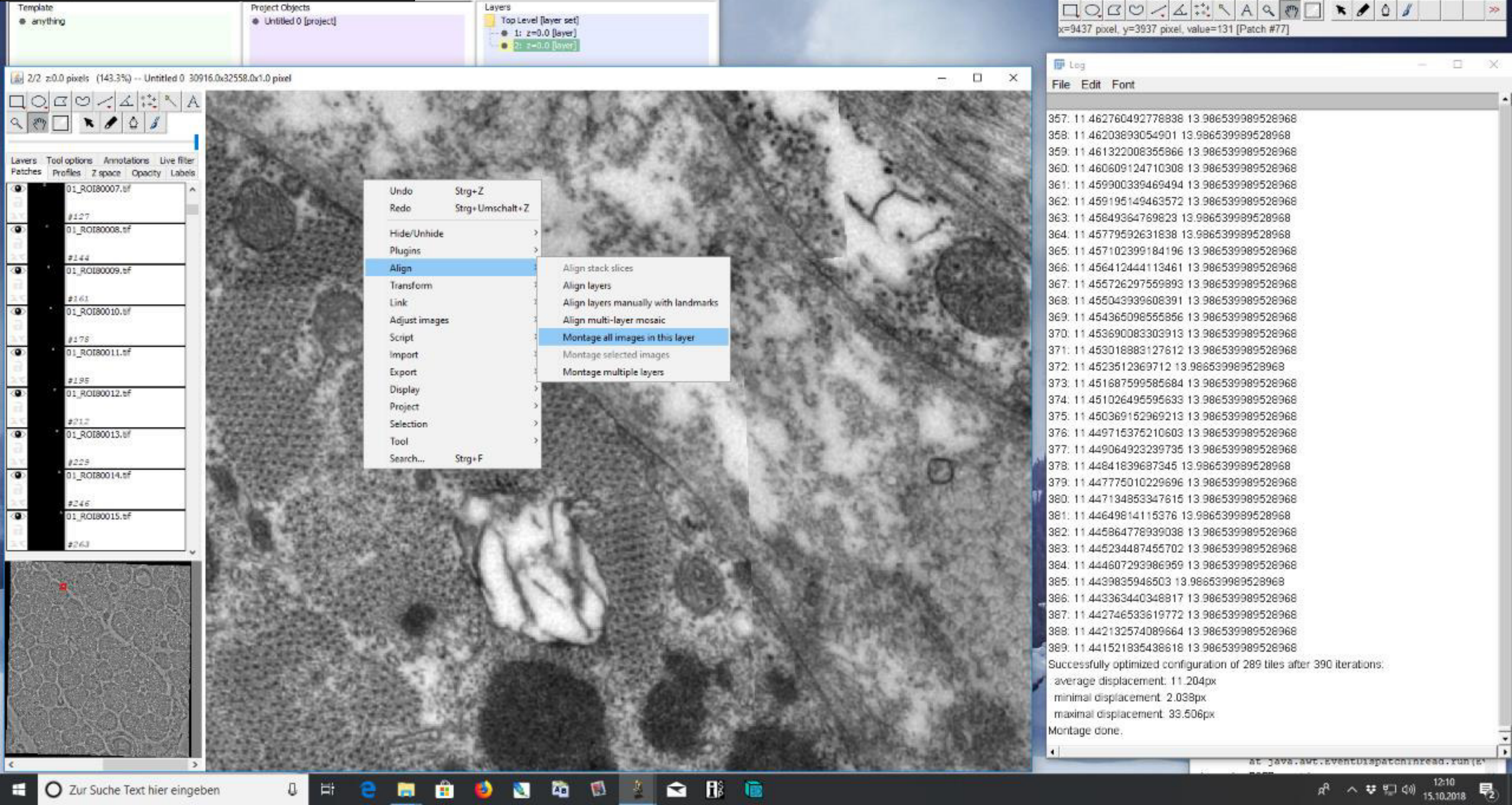

### **5)** Alignment after lens correction

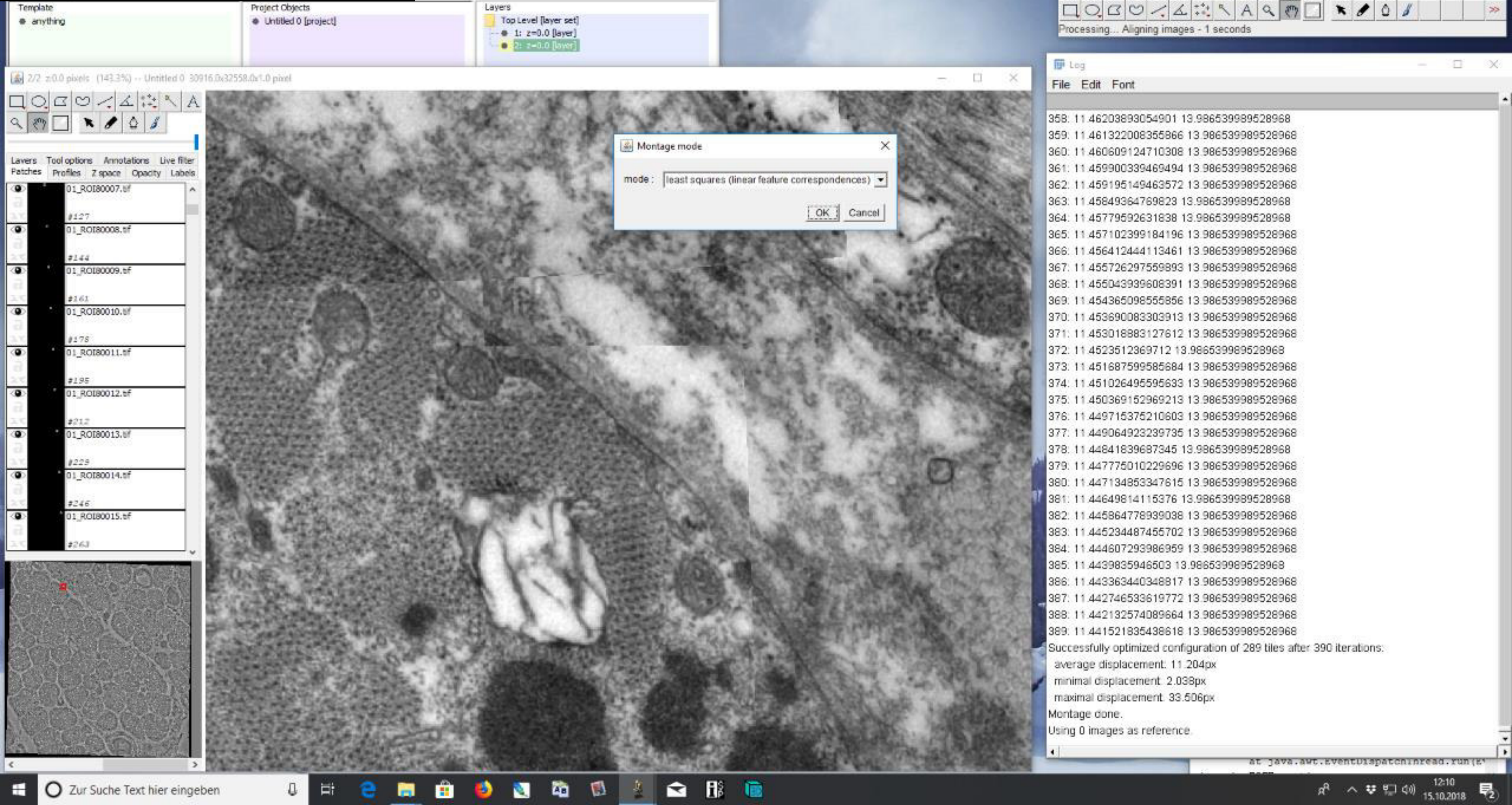

### 5) Alignment after lens correction

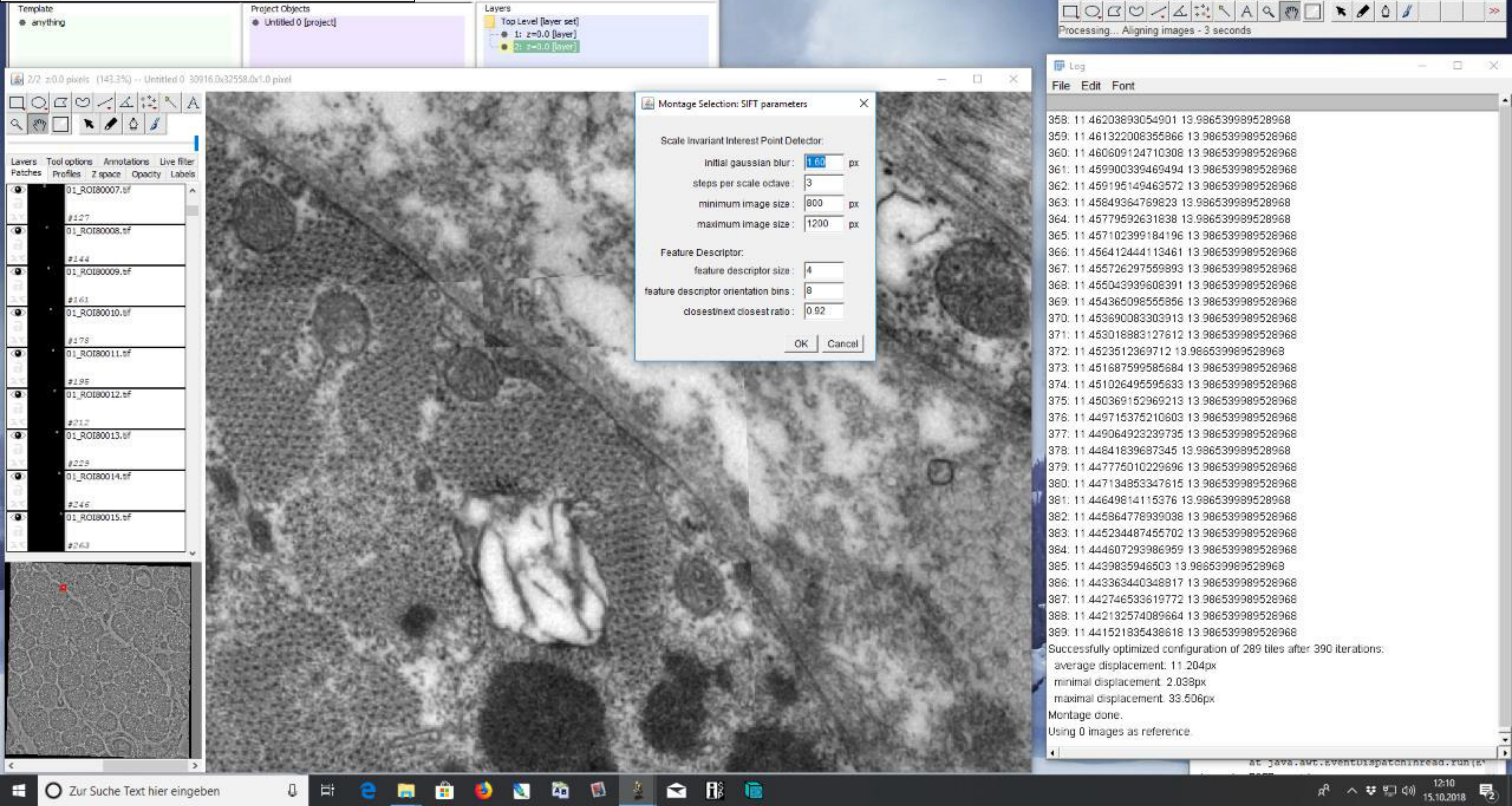

### 5) Alignment after lens correction

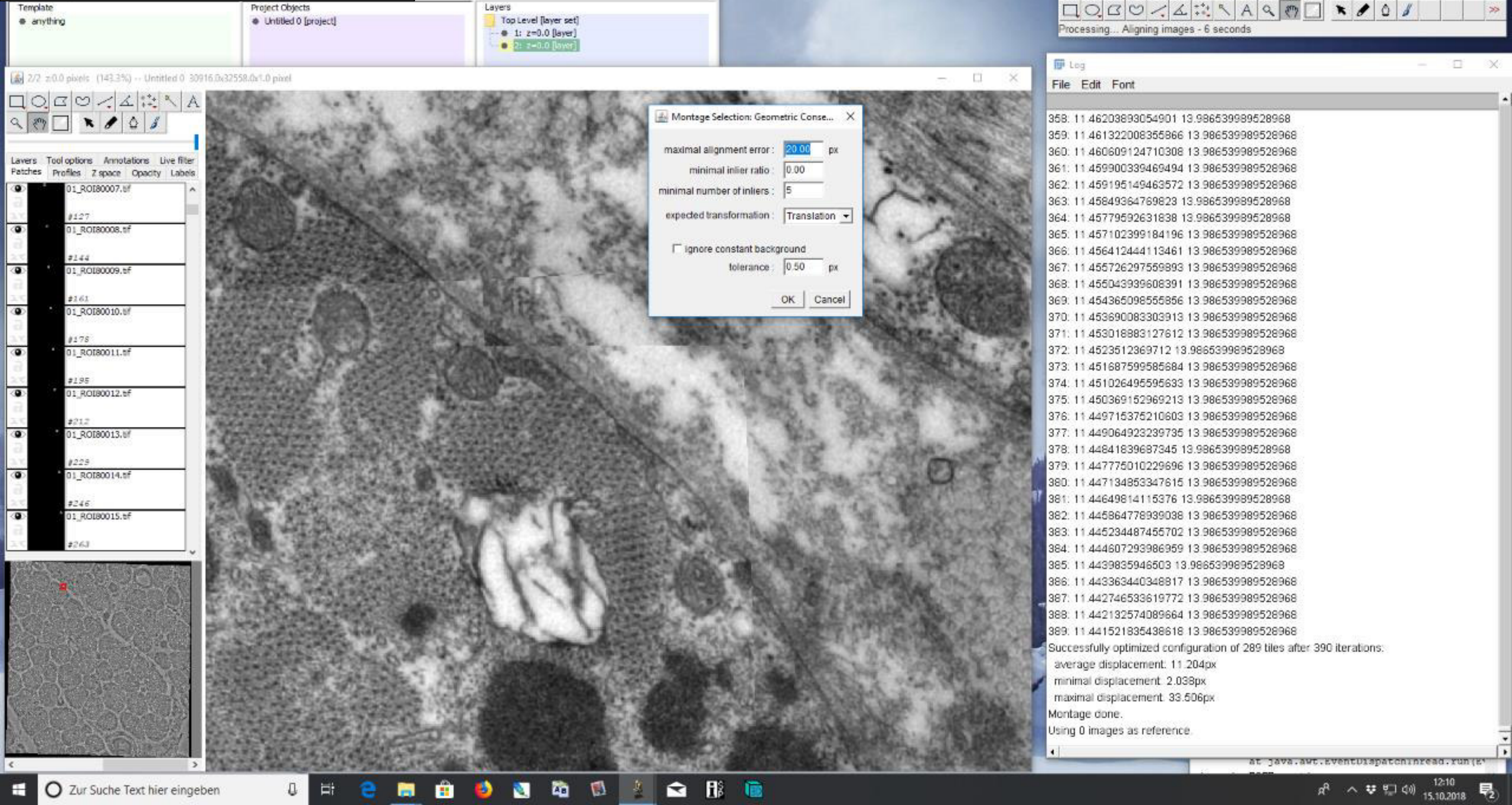

### 5) Alignment after lens correction

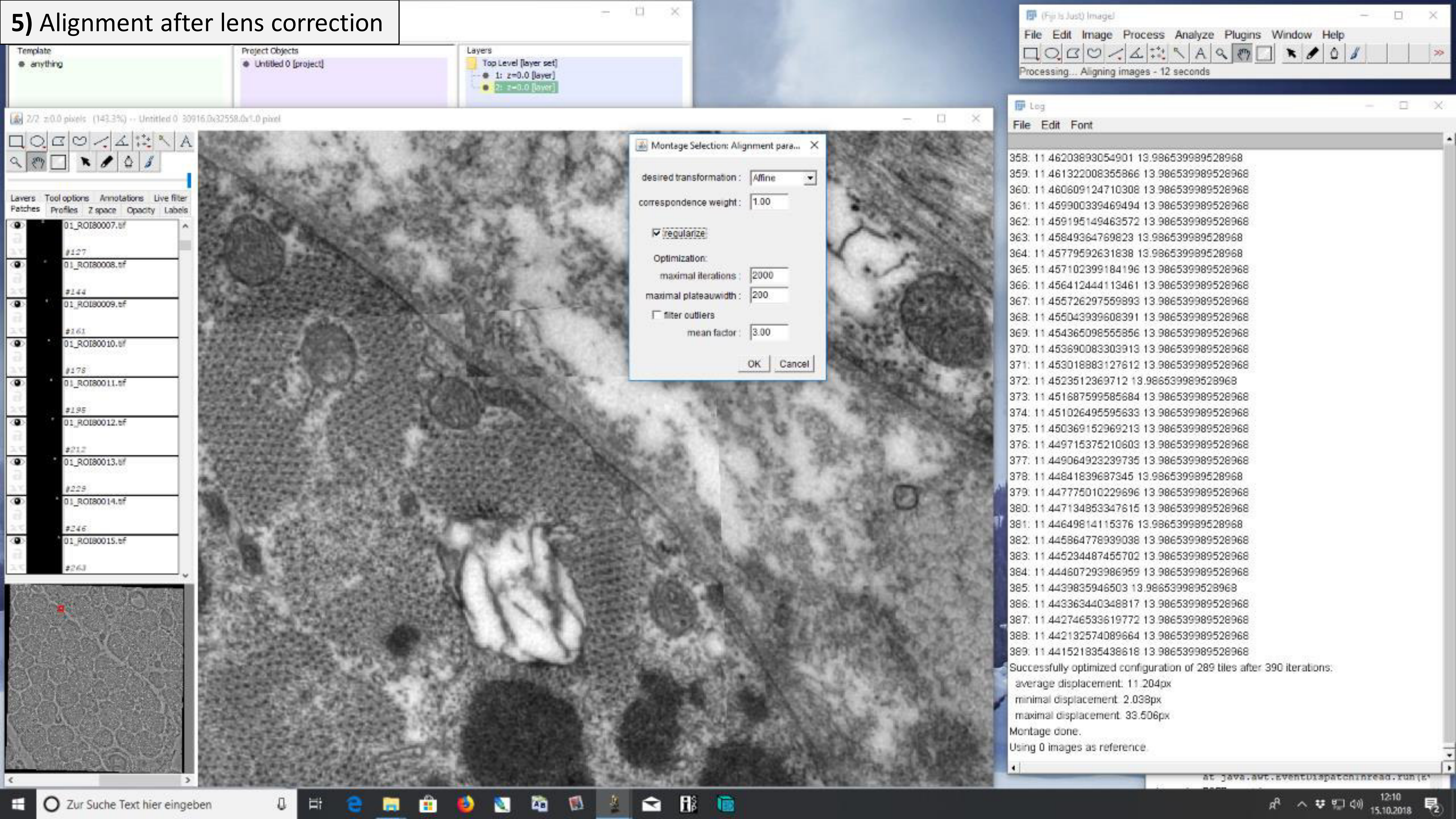

### 5) Alignment after lens correction

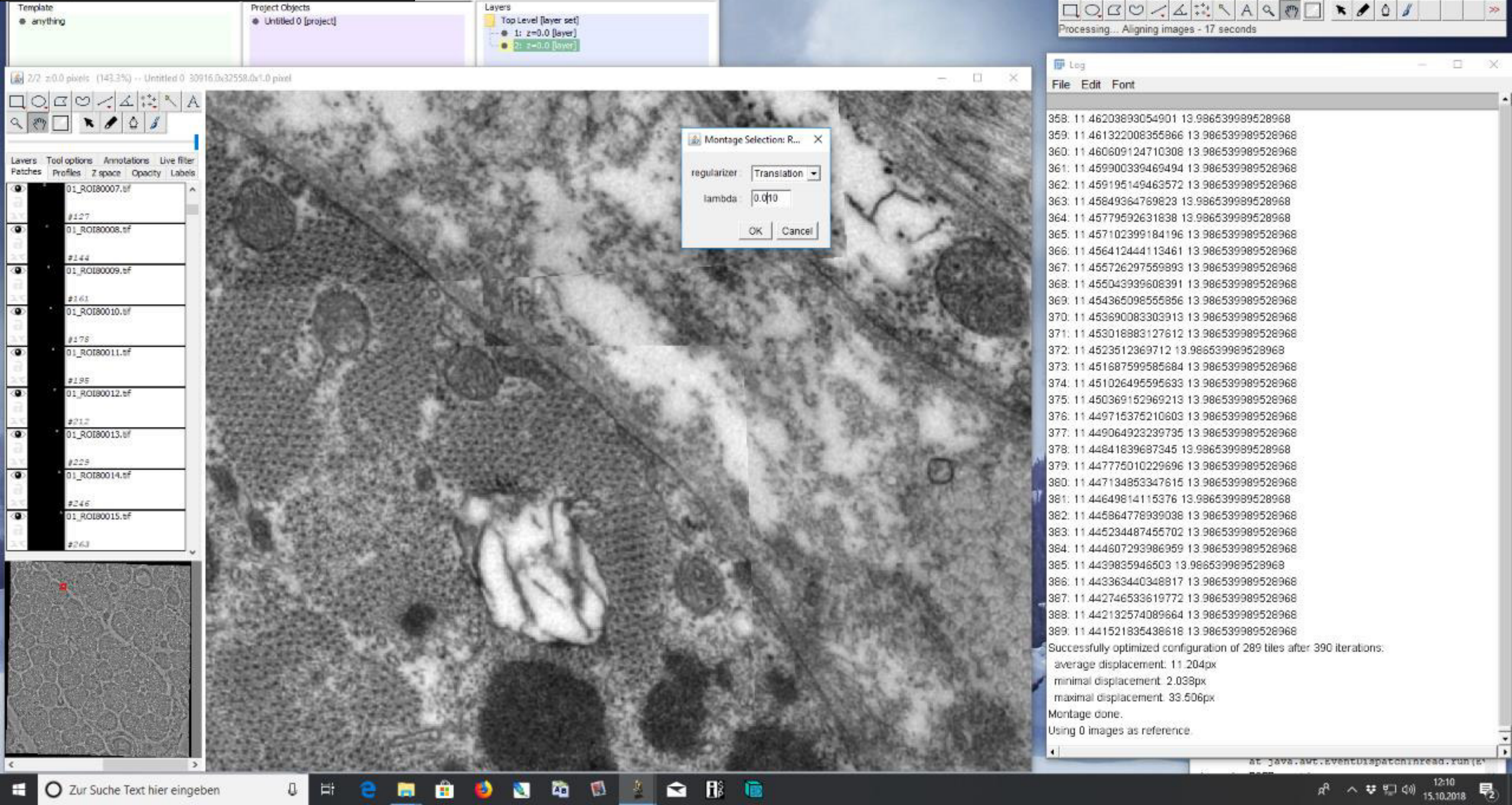

### 5) Alignment after lens correction

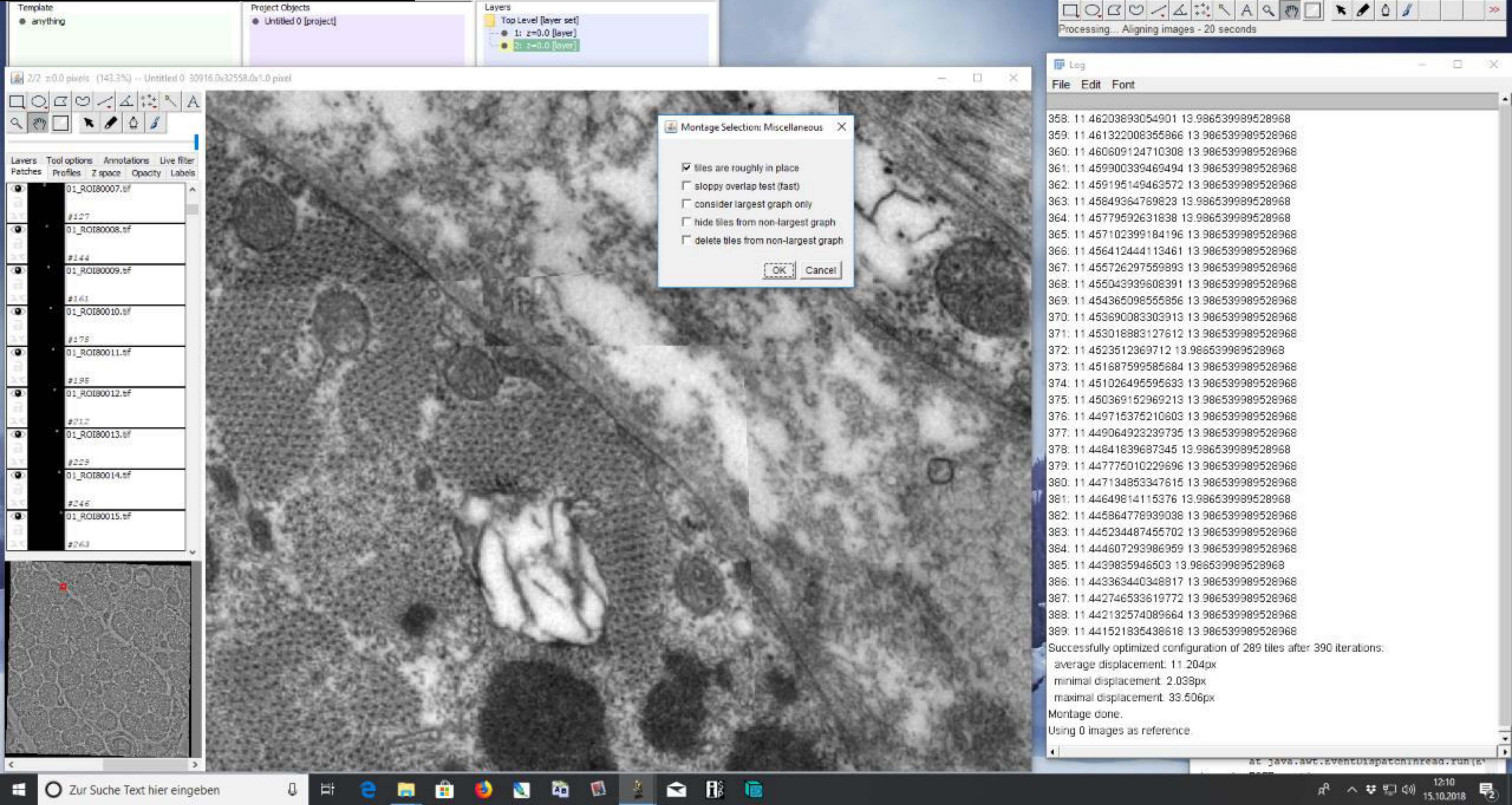

### 5) Alignment after lens correction

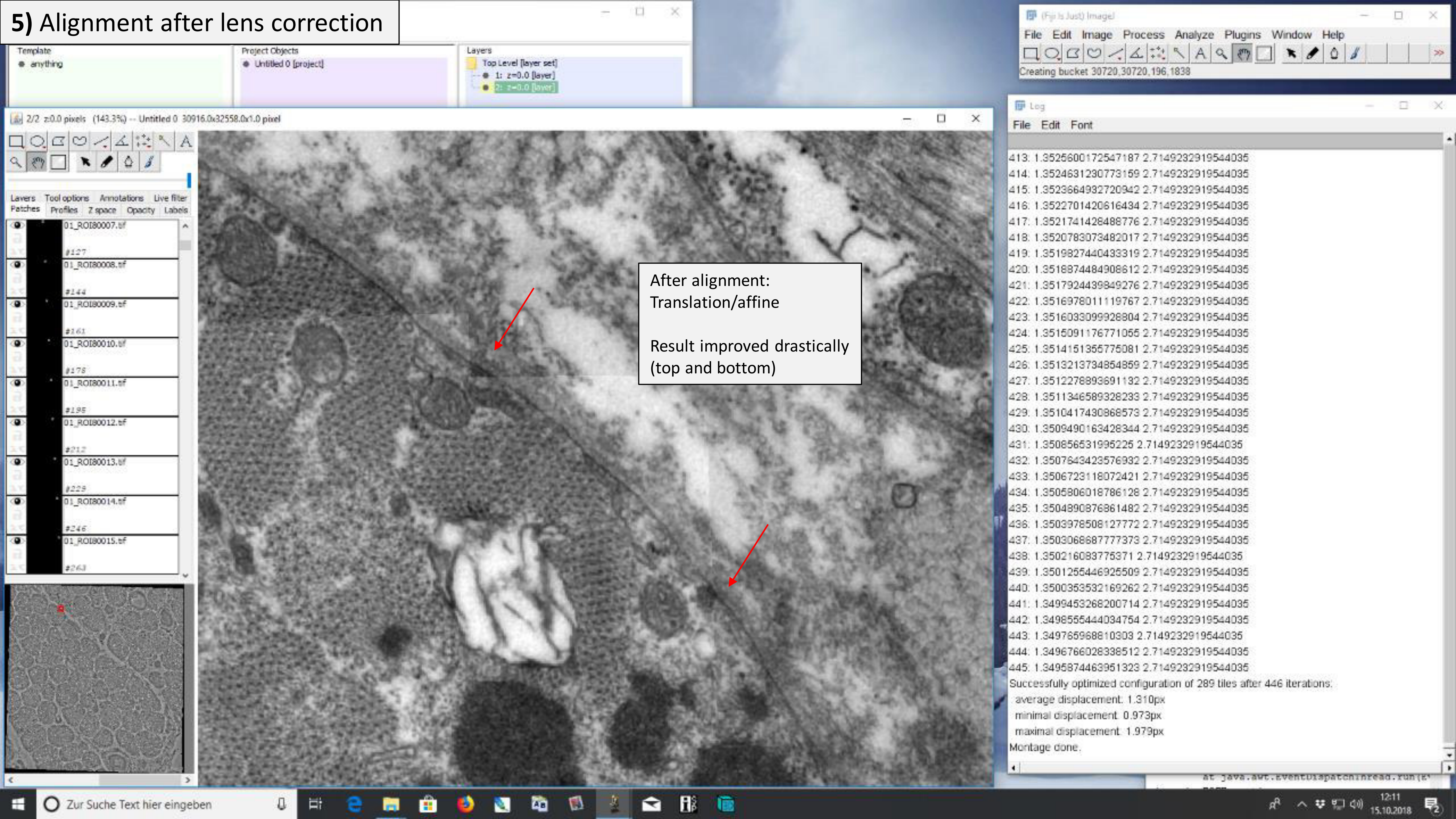

### 6) Remove filter

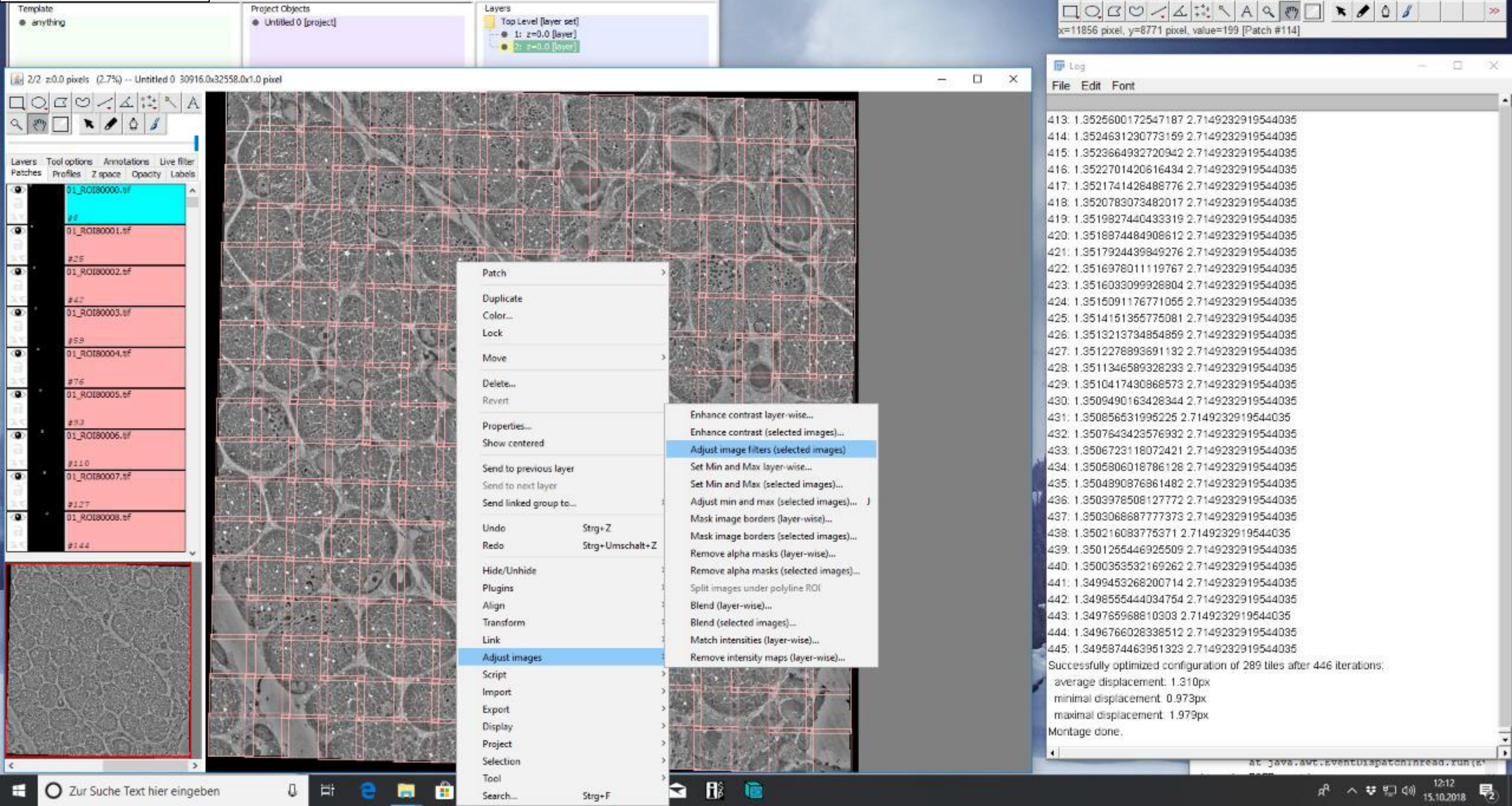

### 6) Remove filter

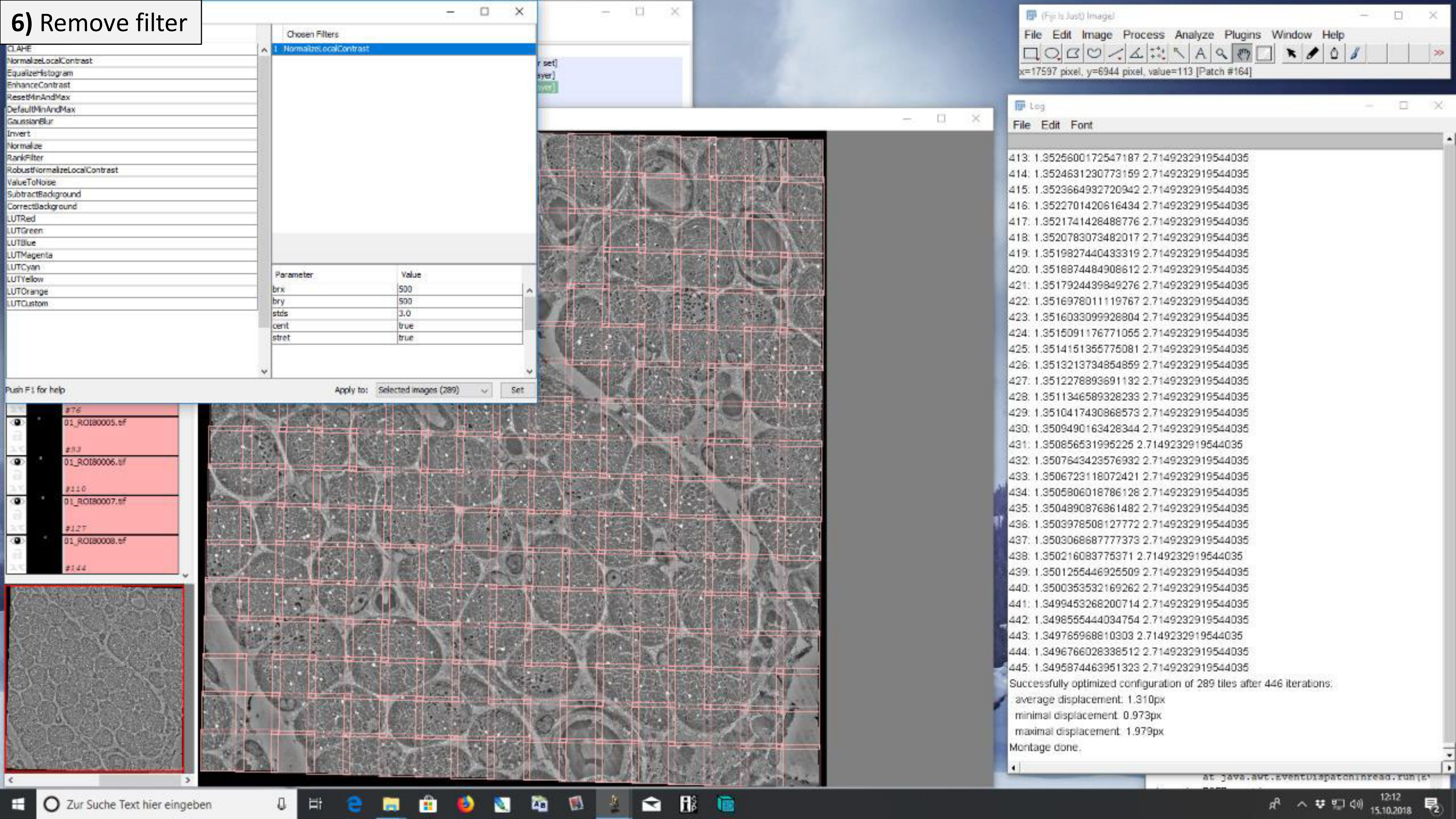

### 6) Remove filter

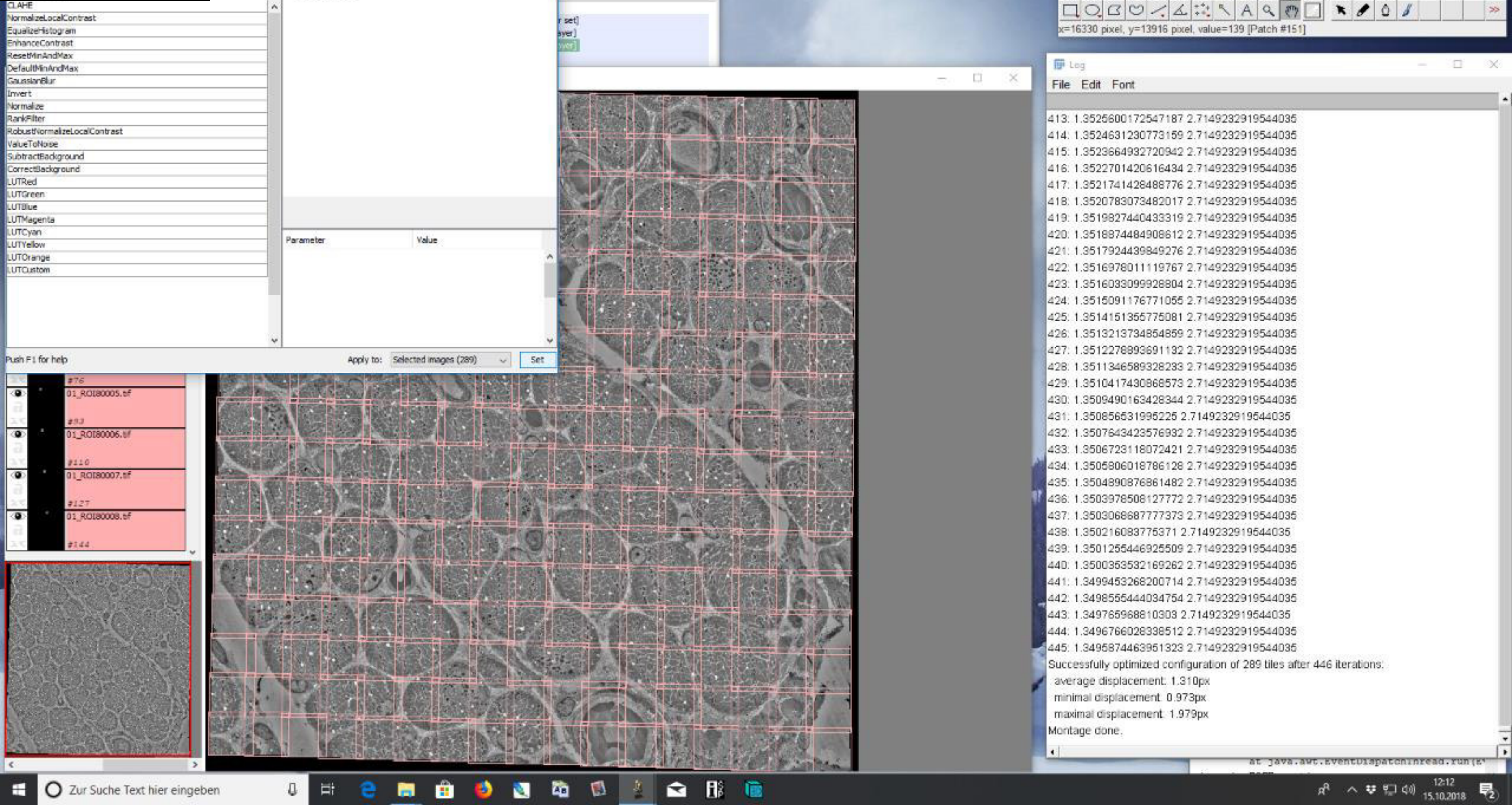

### 6) Remove filter

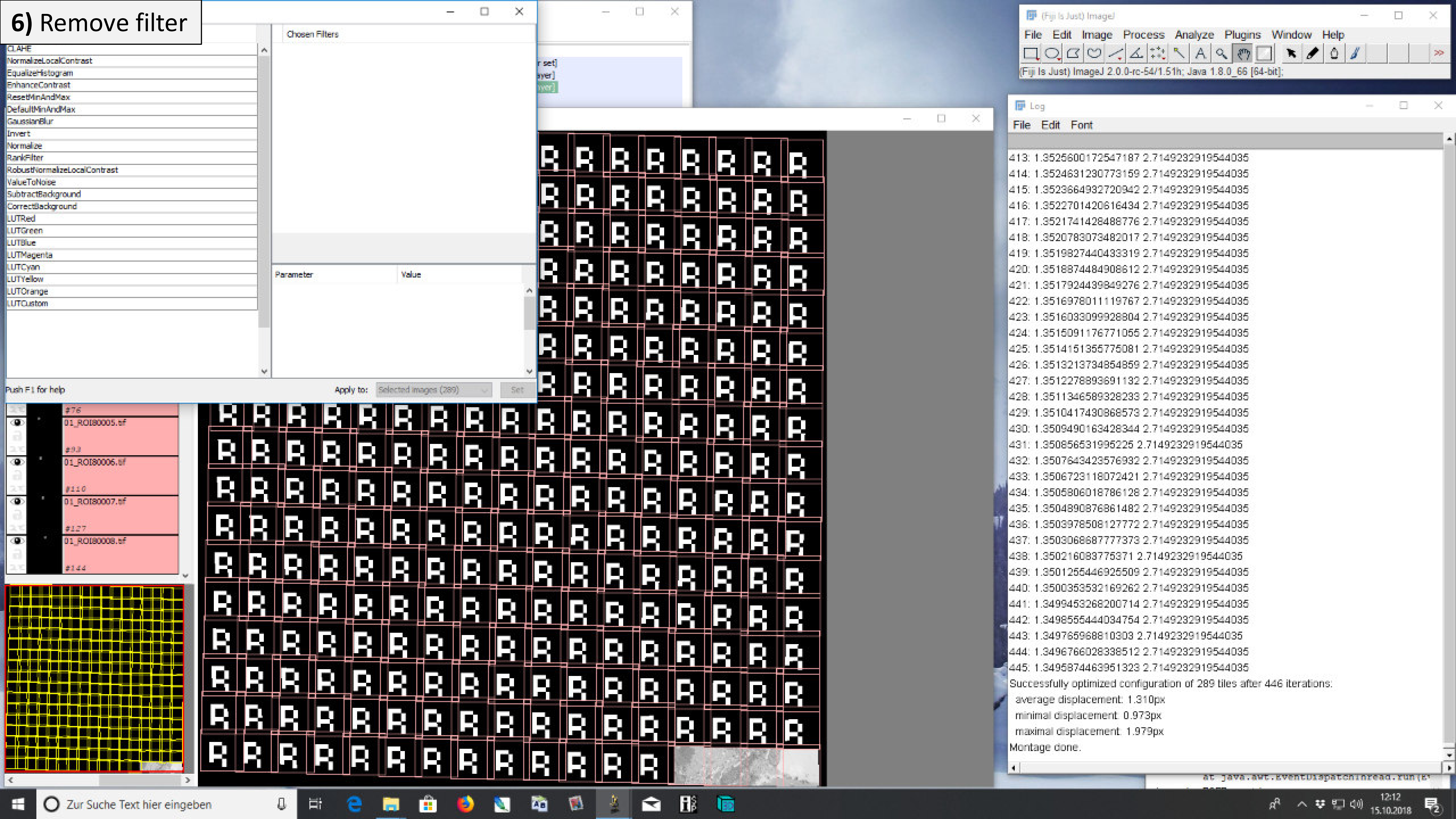

### **6)** Remove filter

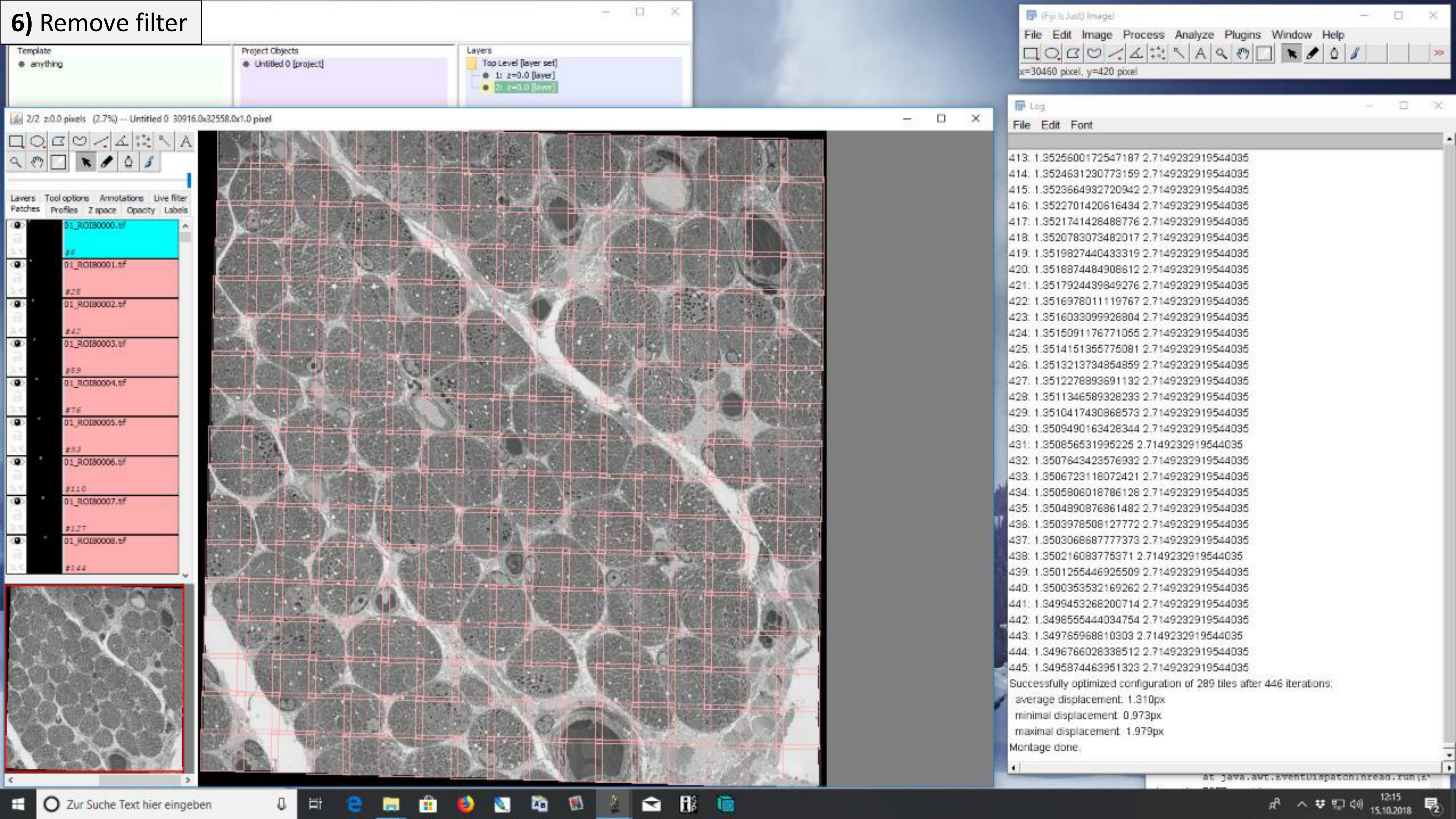

### **6)** Remove filter

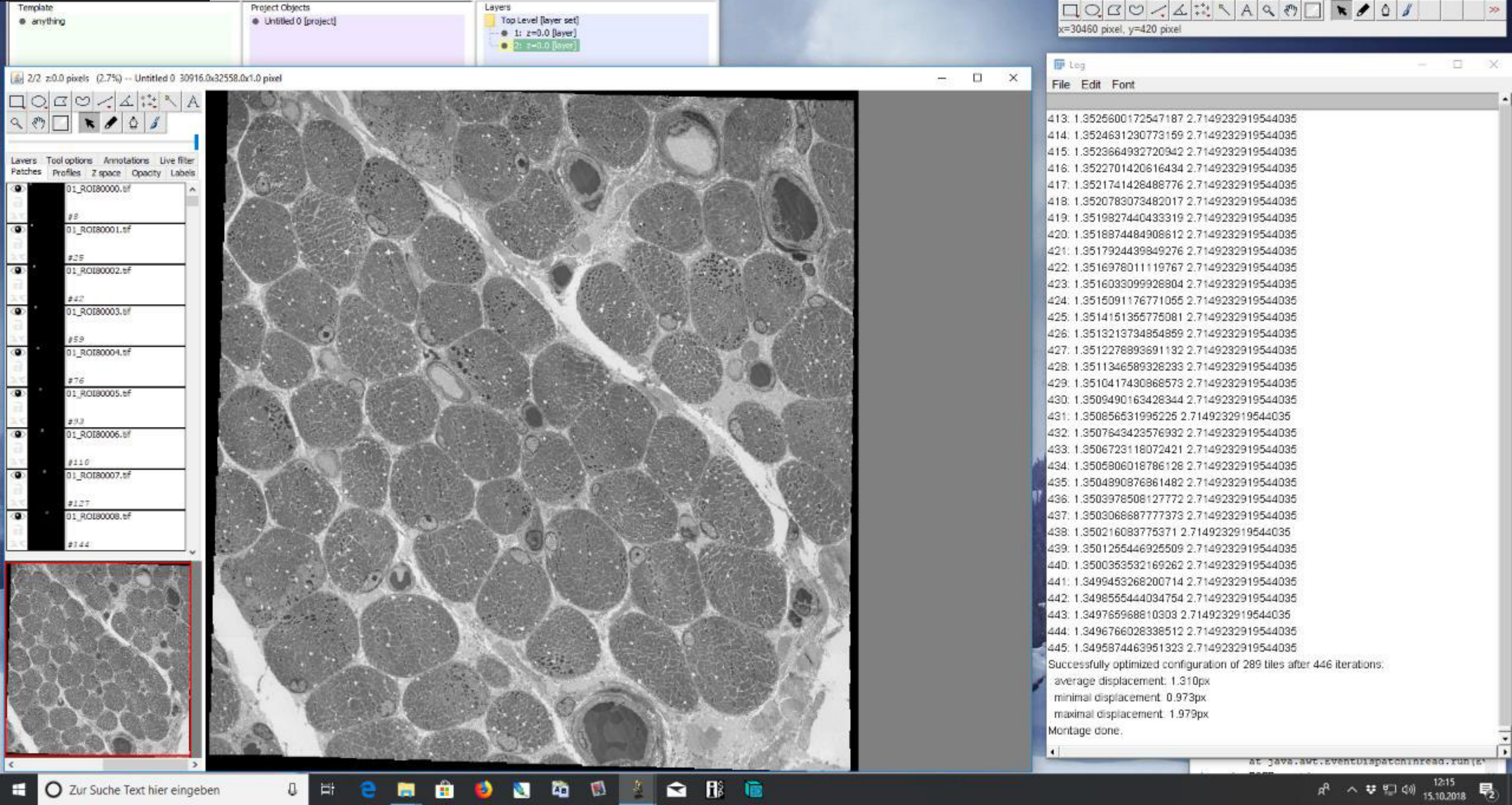

### 7) Match intensities

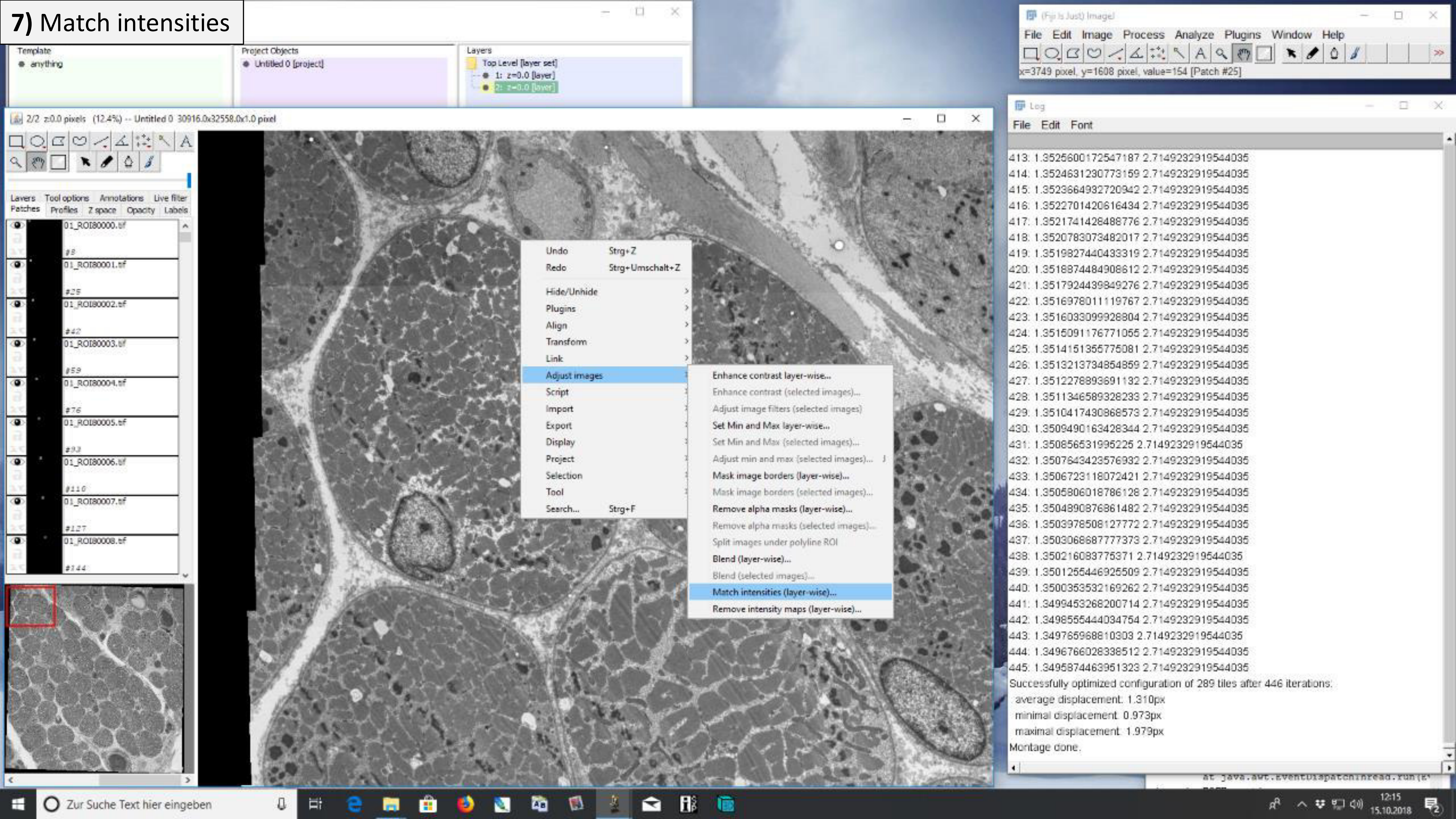

### 7) Match intensities

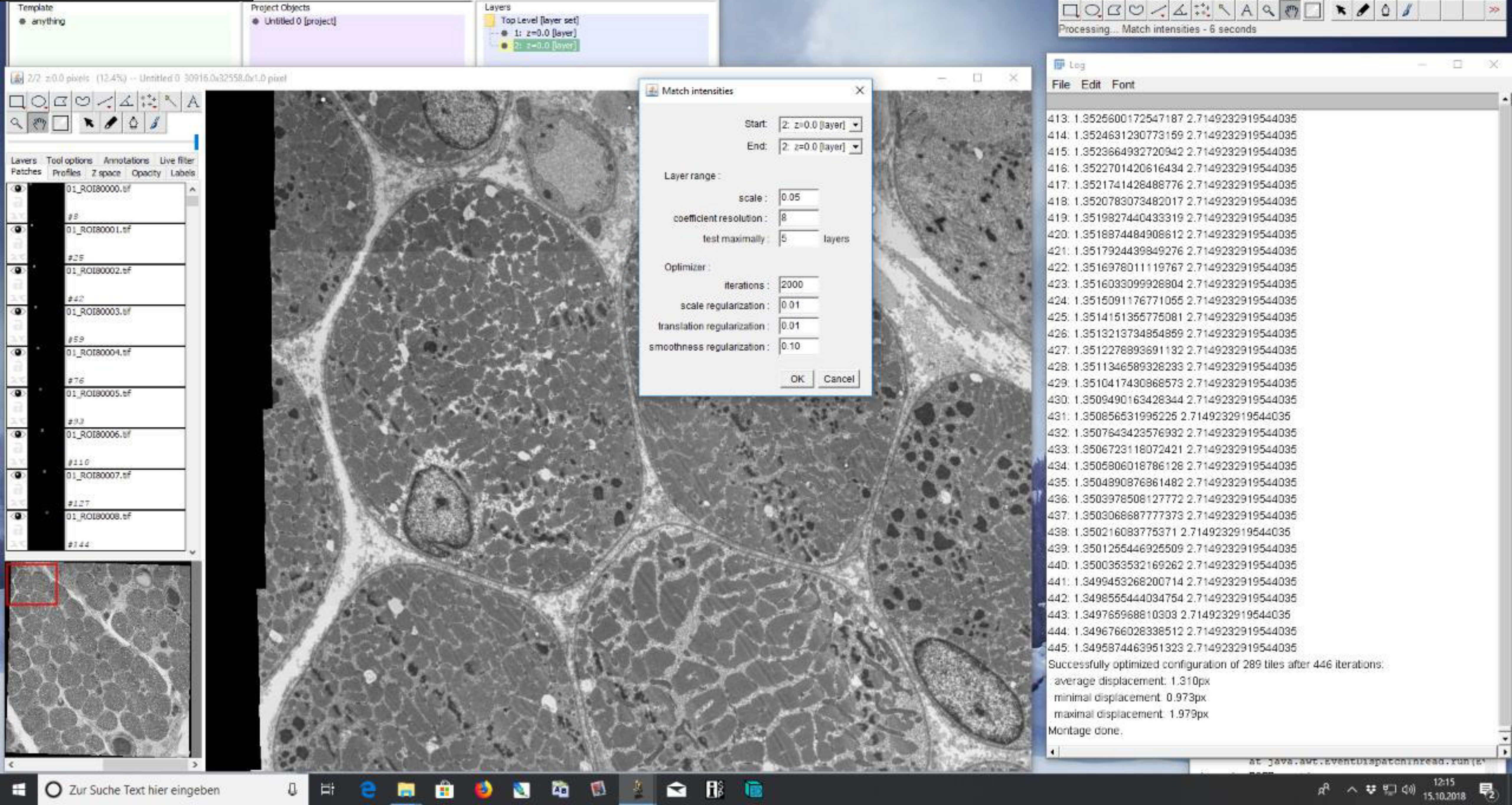

### 7) Match intensities

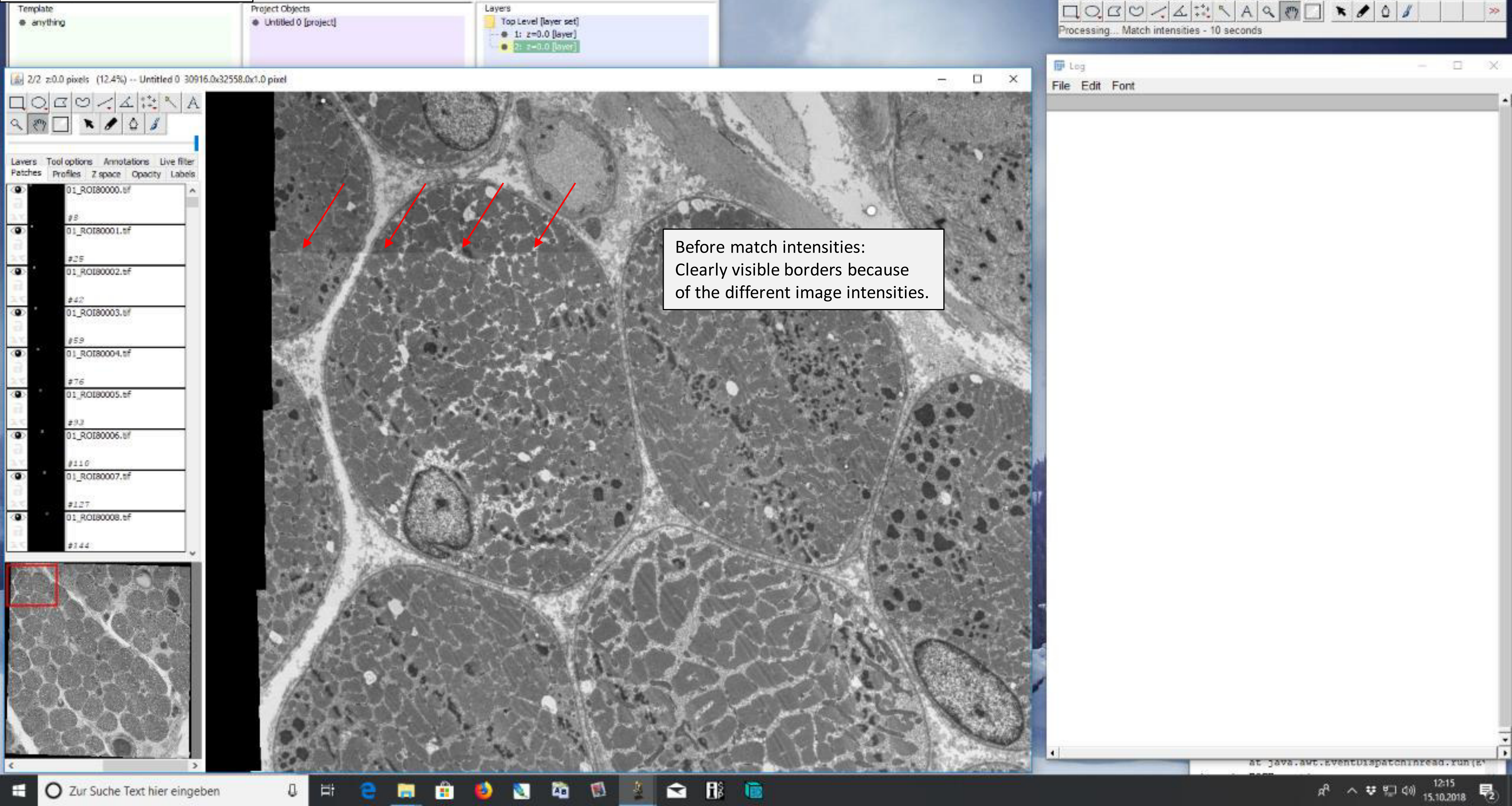

### 7) Match intensities

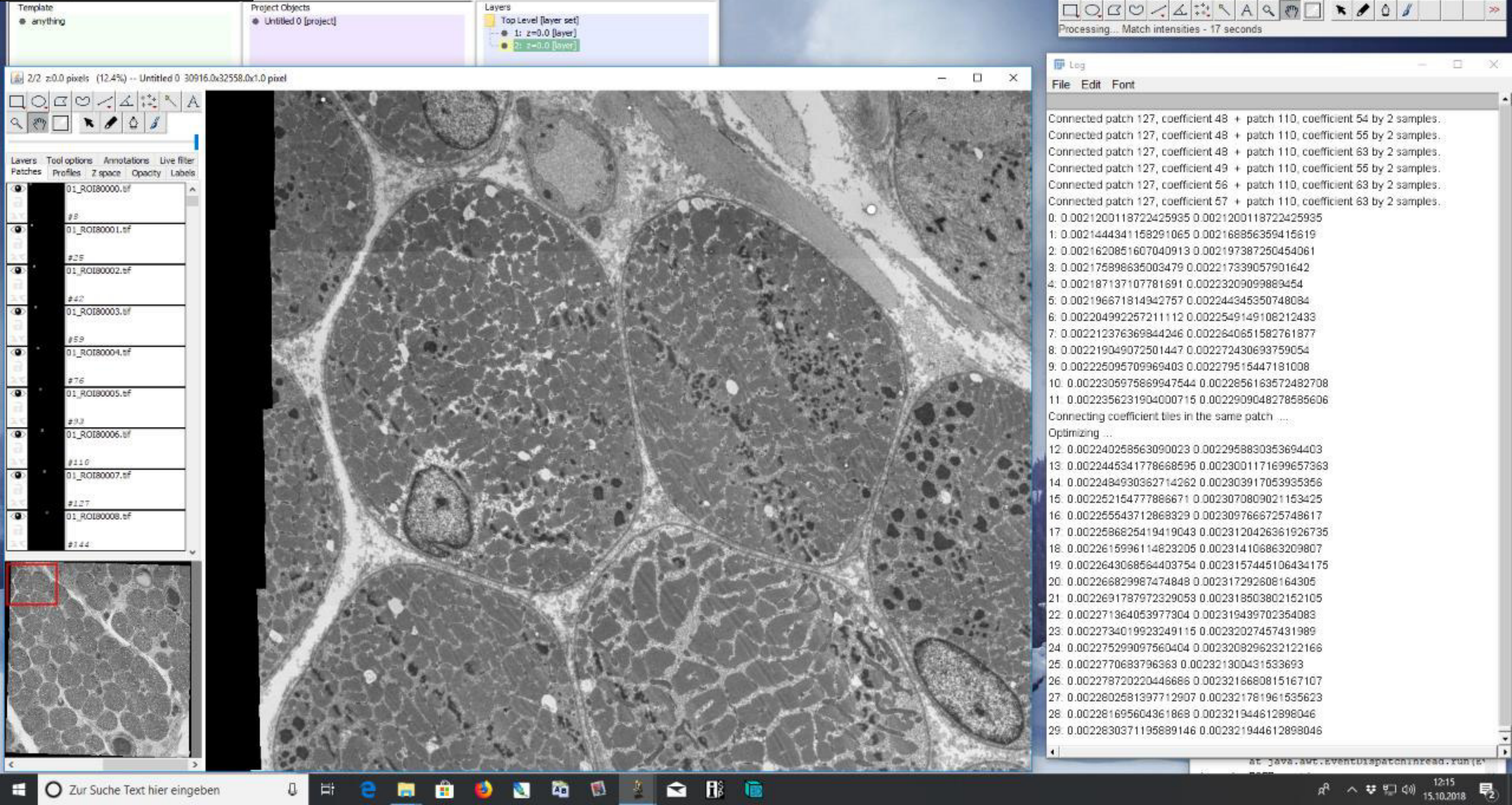

### 7) Match intensities

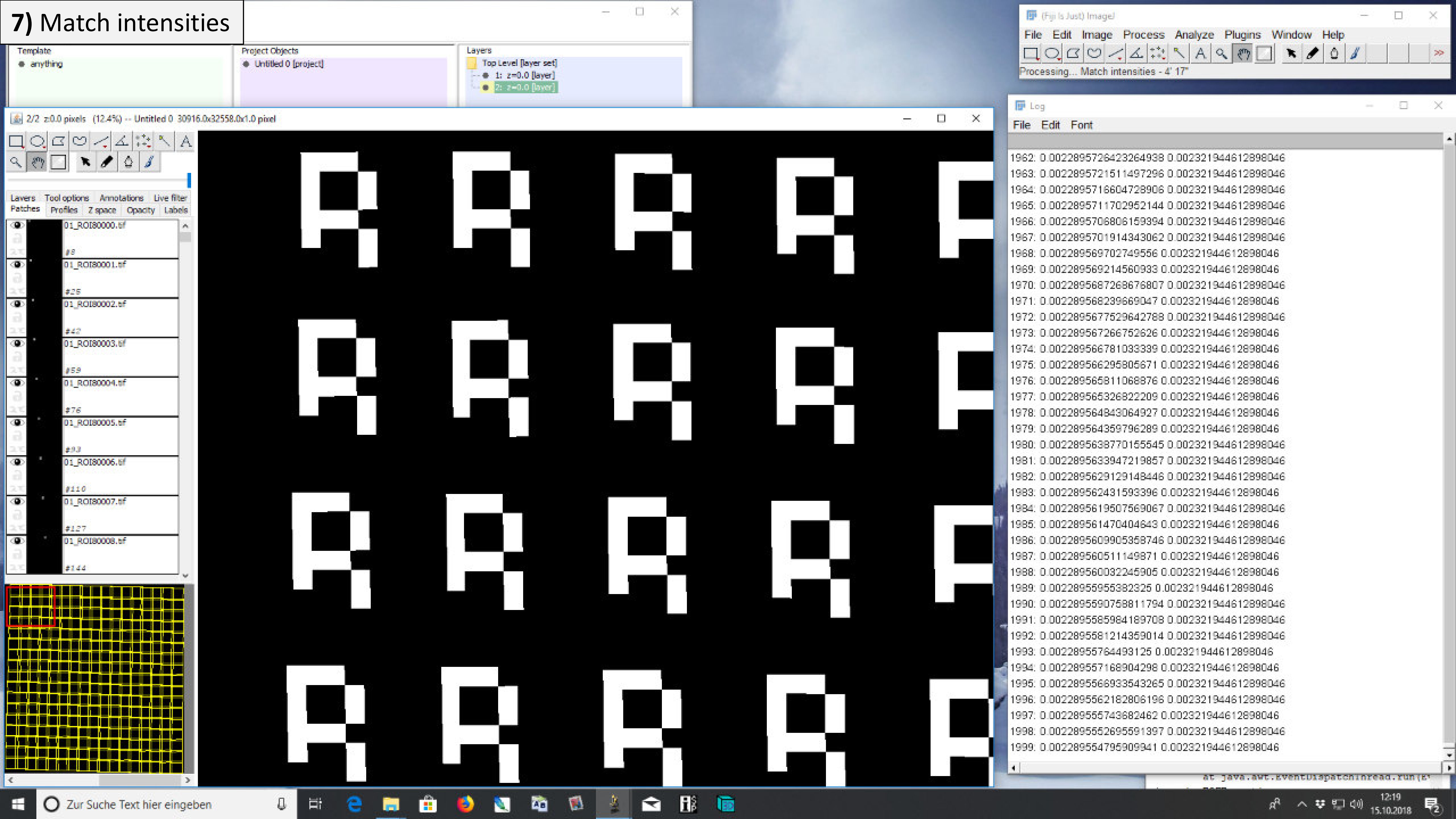

### 7) Match intensities

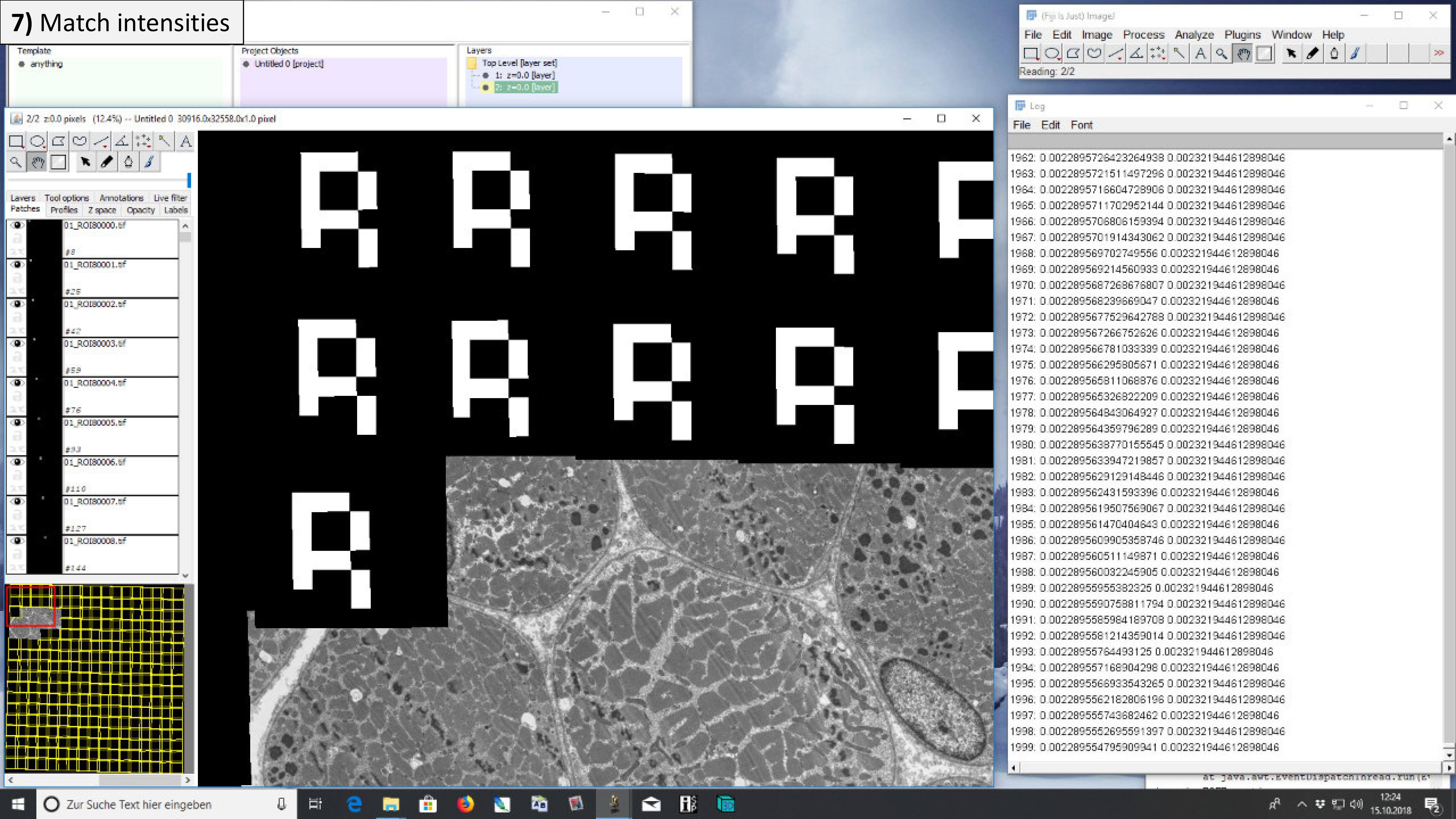

### 7) Match intensities

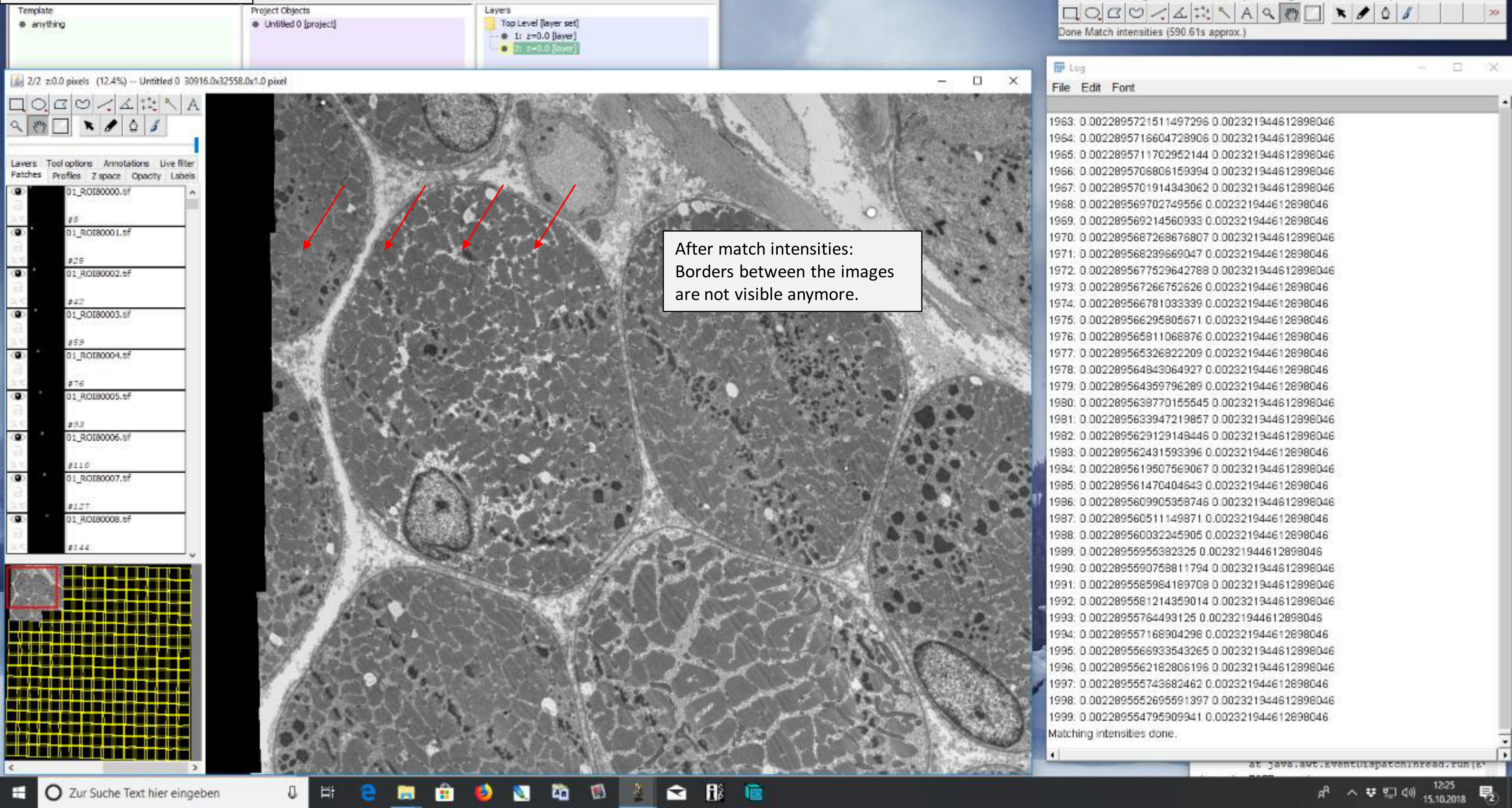


After match intensities:
Borders between the images
are not visible anymore.

**8)** Rotate dataset

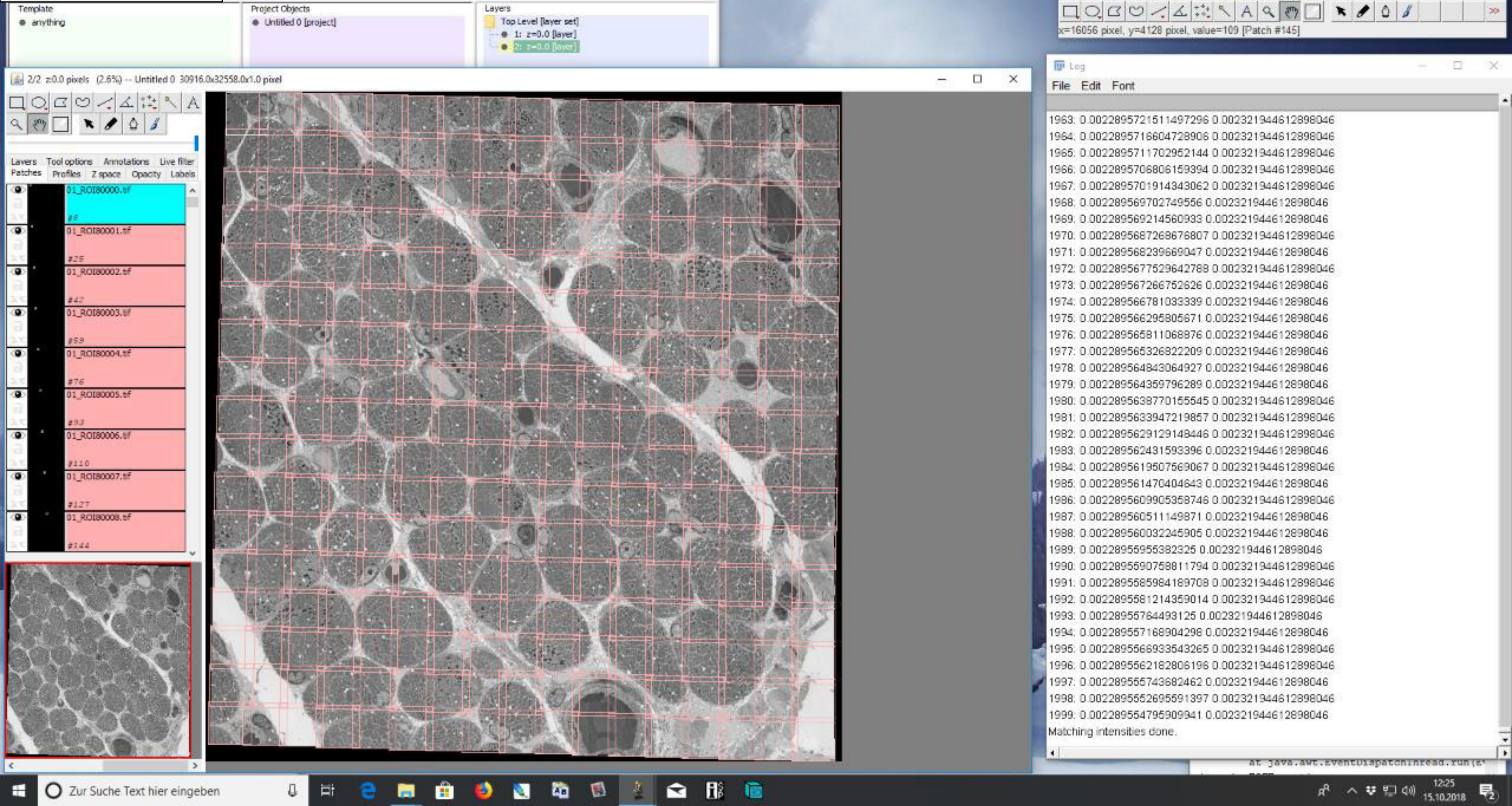

### 8) Rotate dataset

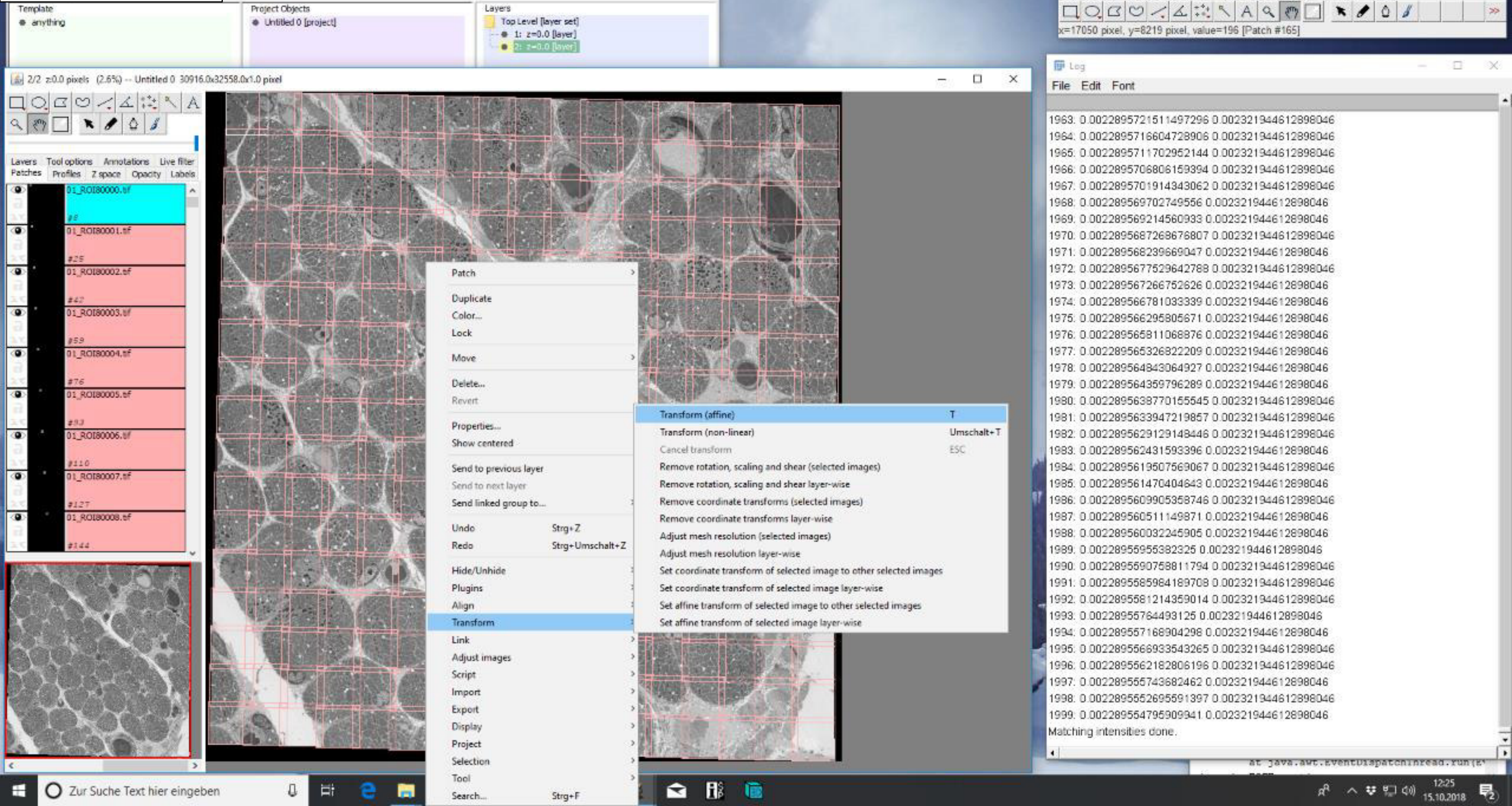

### 8) Rotate dataset

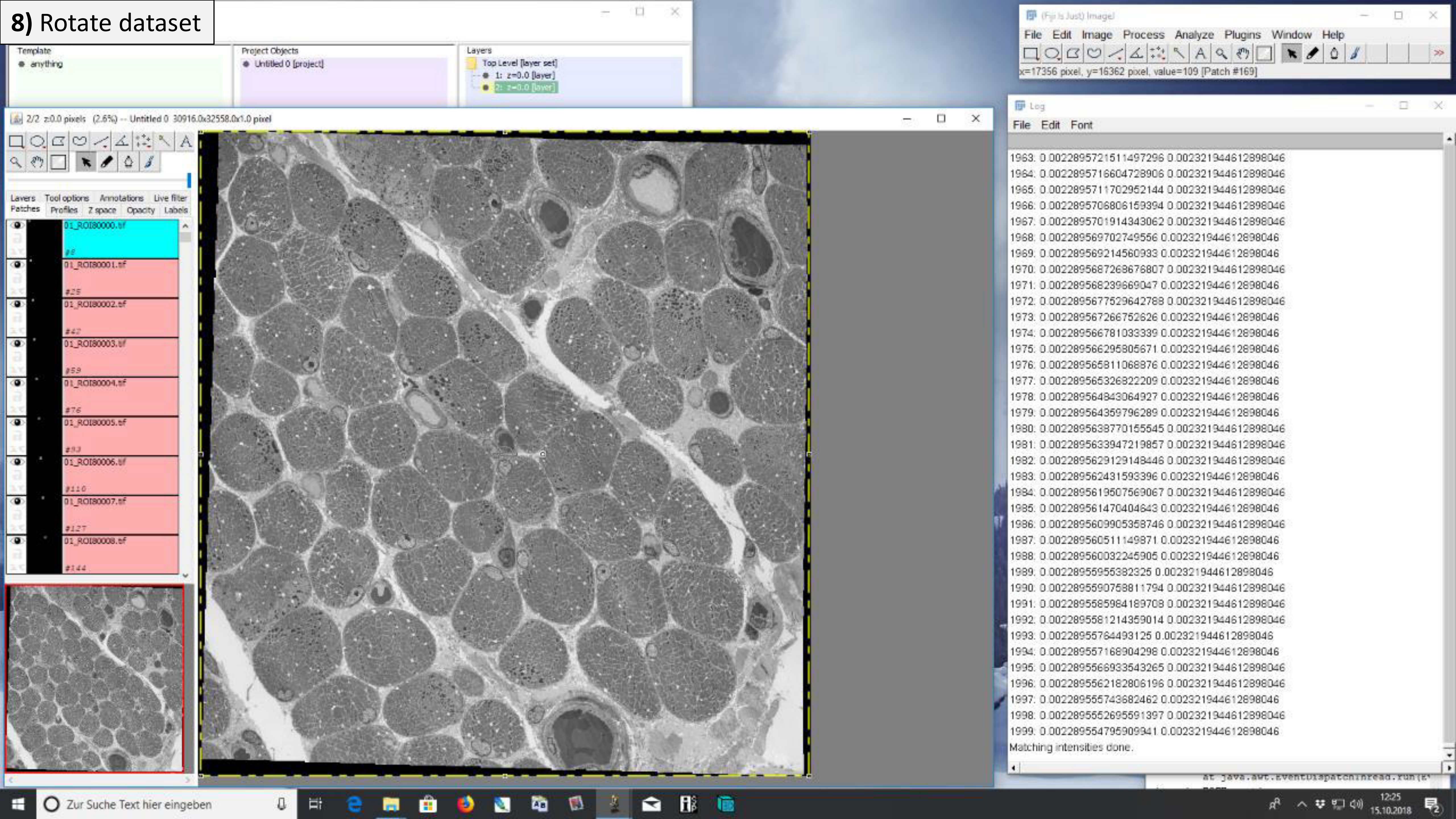

### 8) Rotate dataset

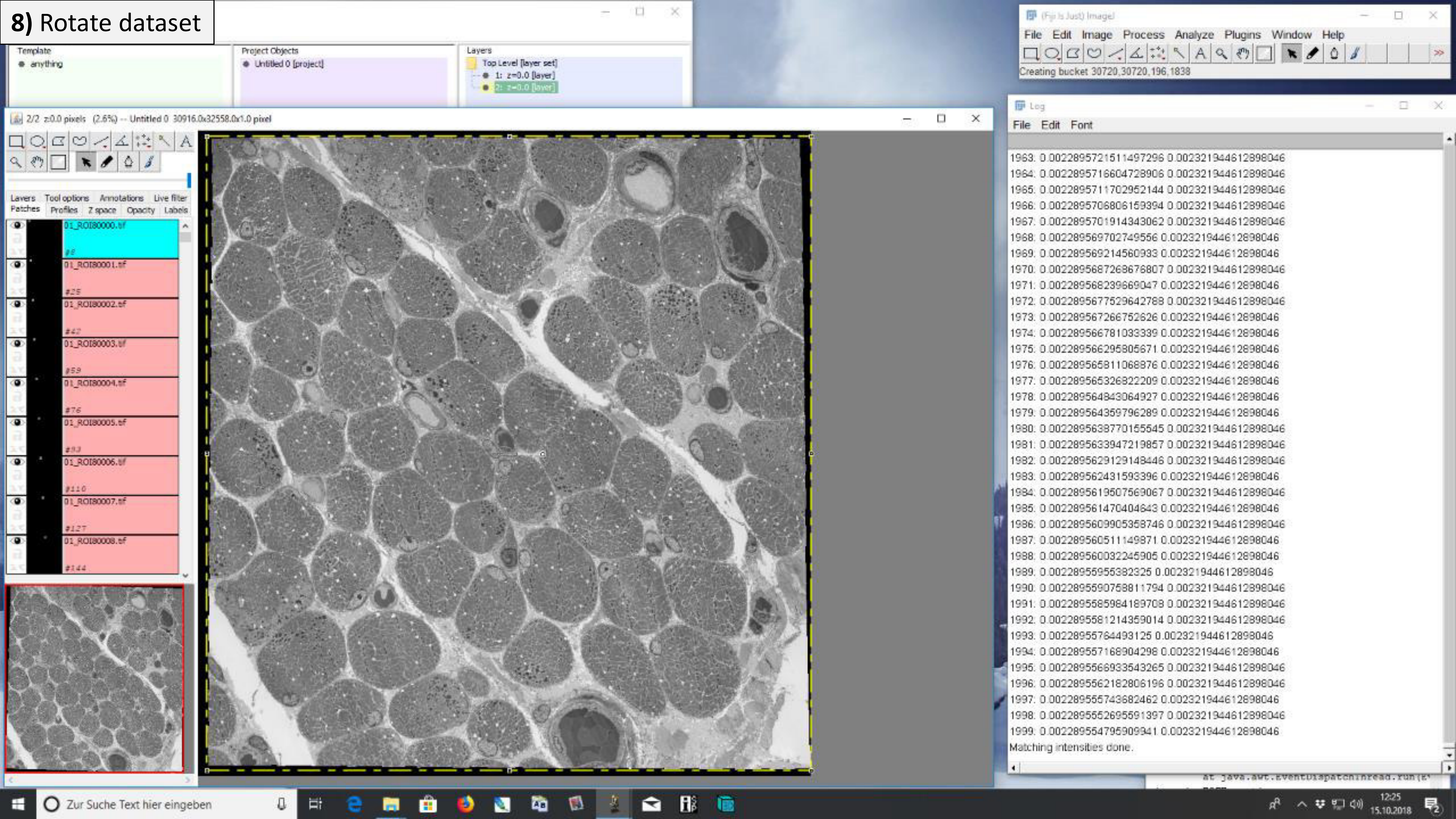

### 8) Rotate dataset

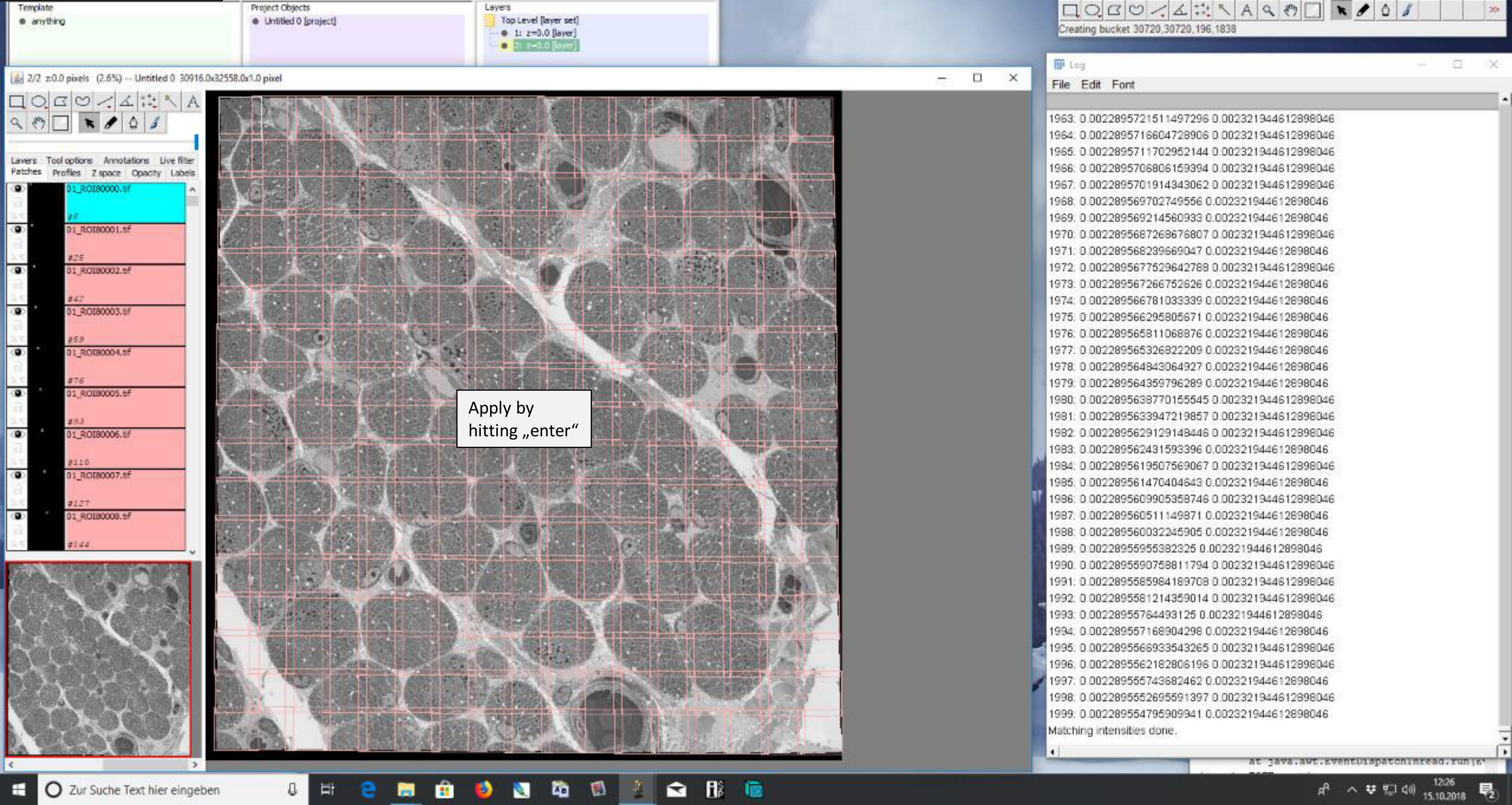

### **8)** Rotate dataset

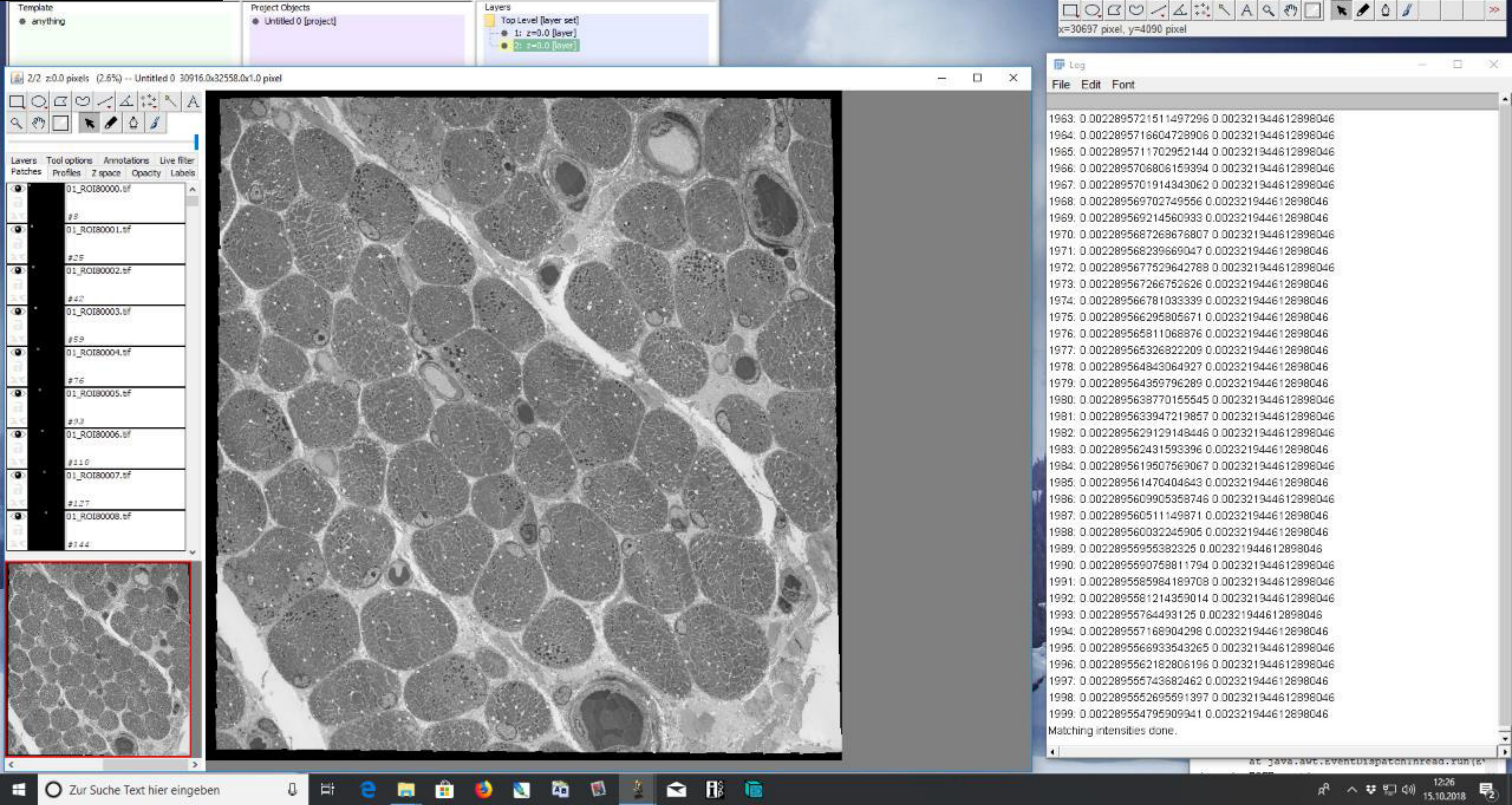

### 9) Standard export

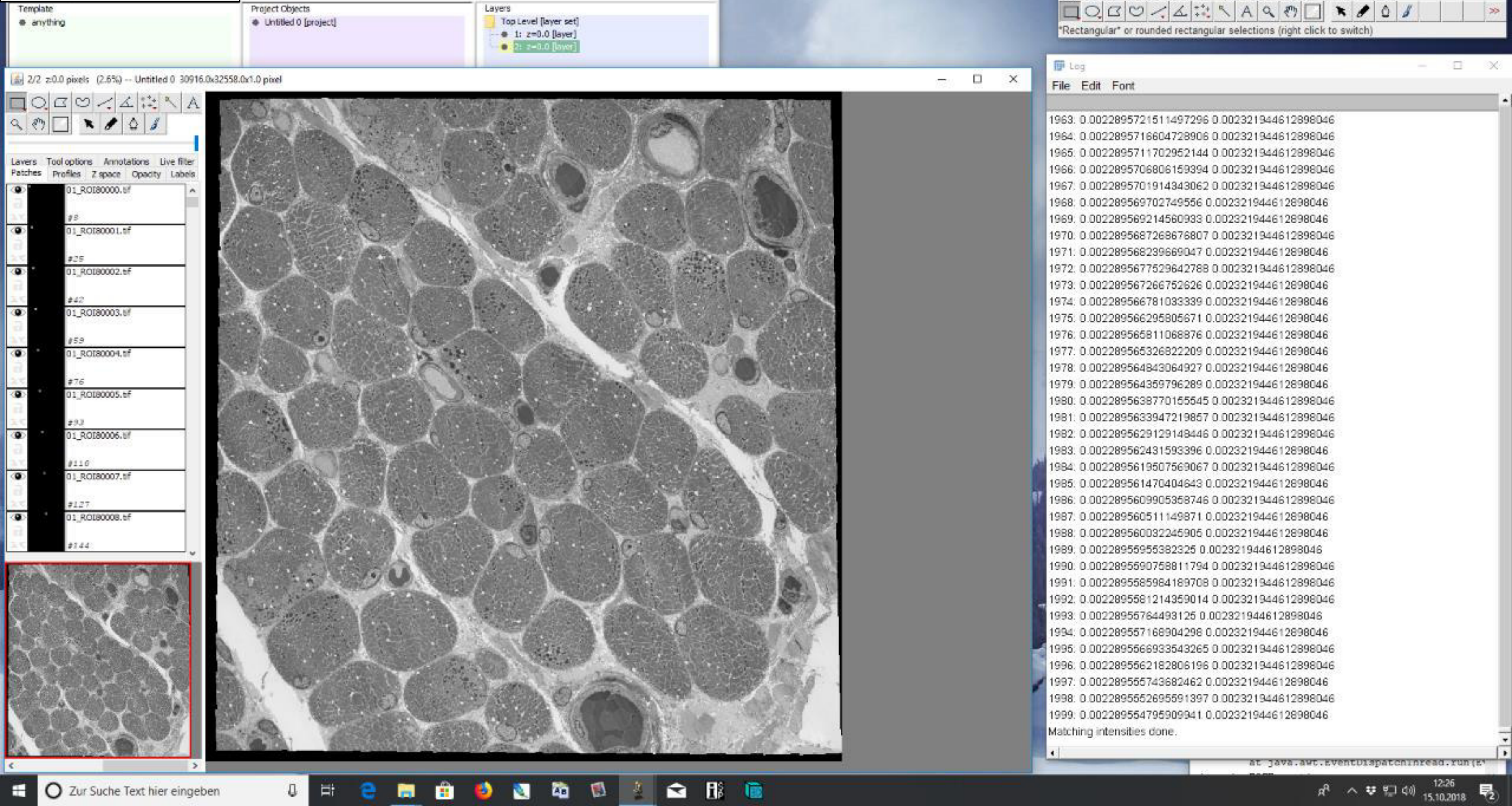

### 9) Standard export

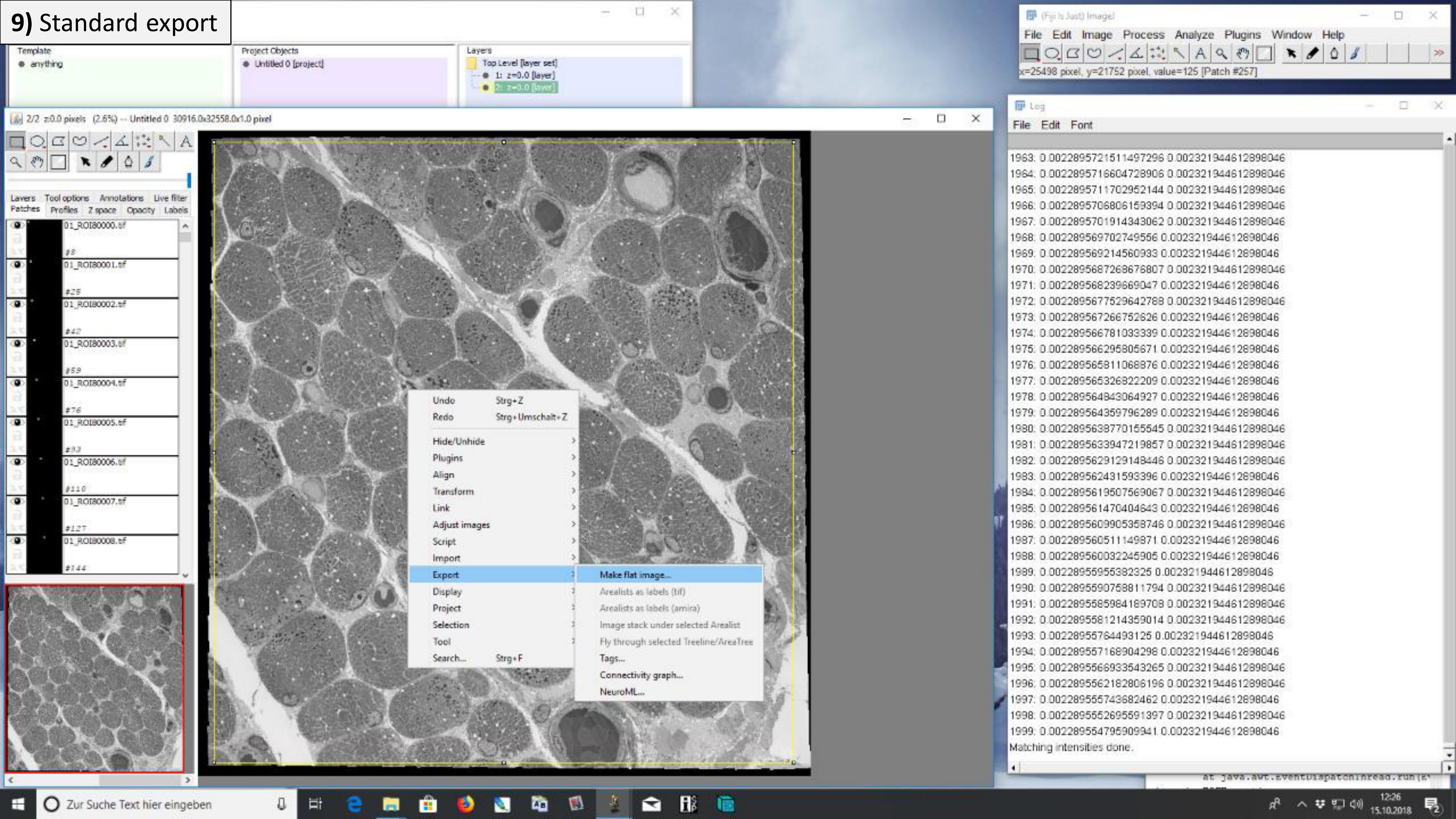

### 9) Standard export

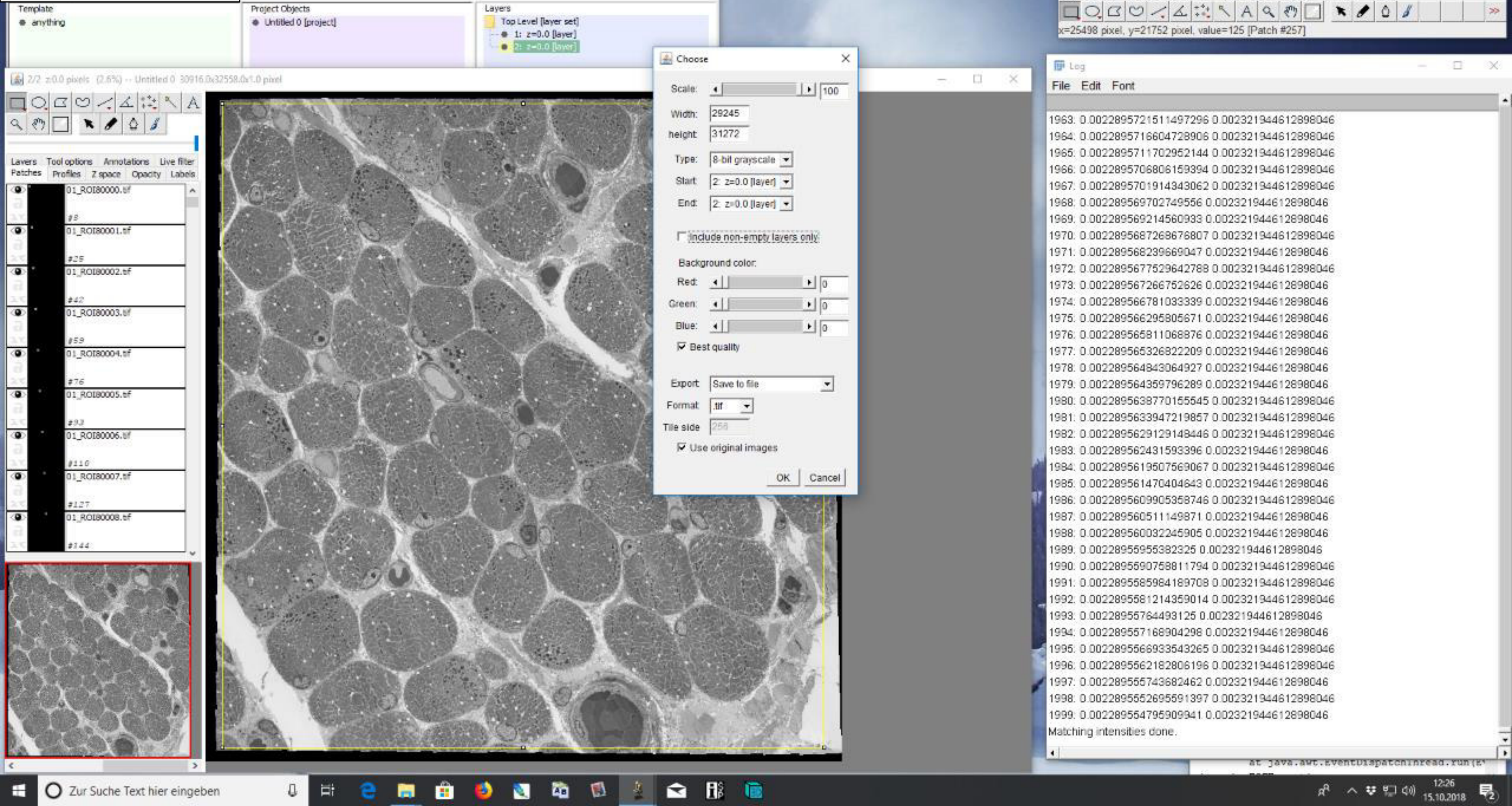

### 9) Standard export

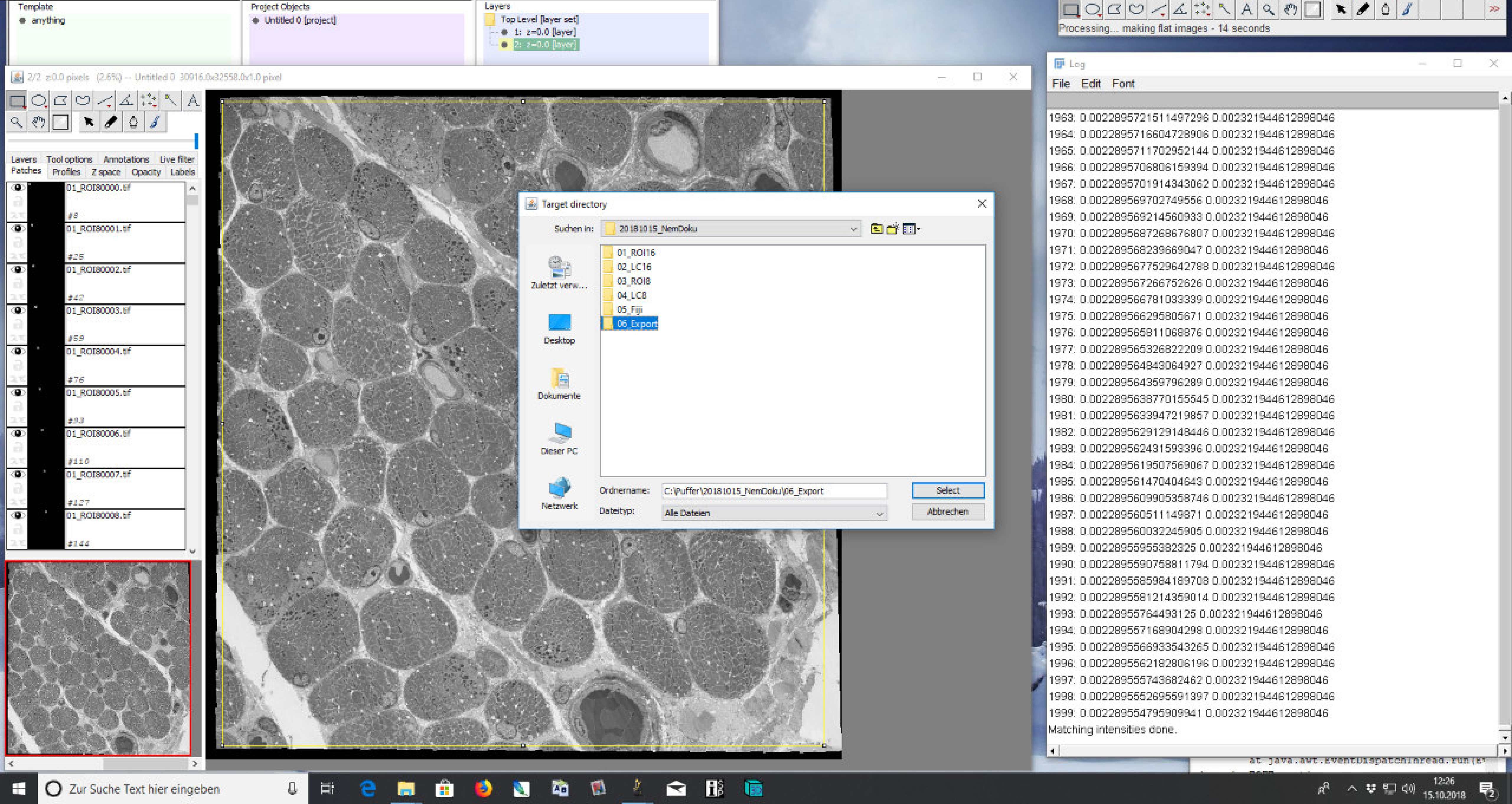

### 9) Standard export

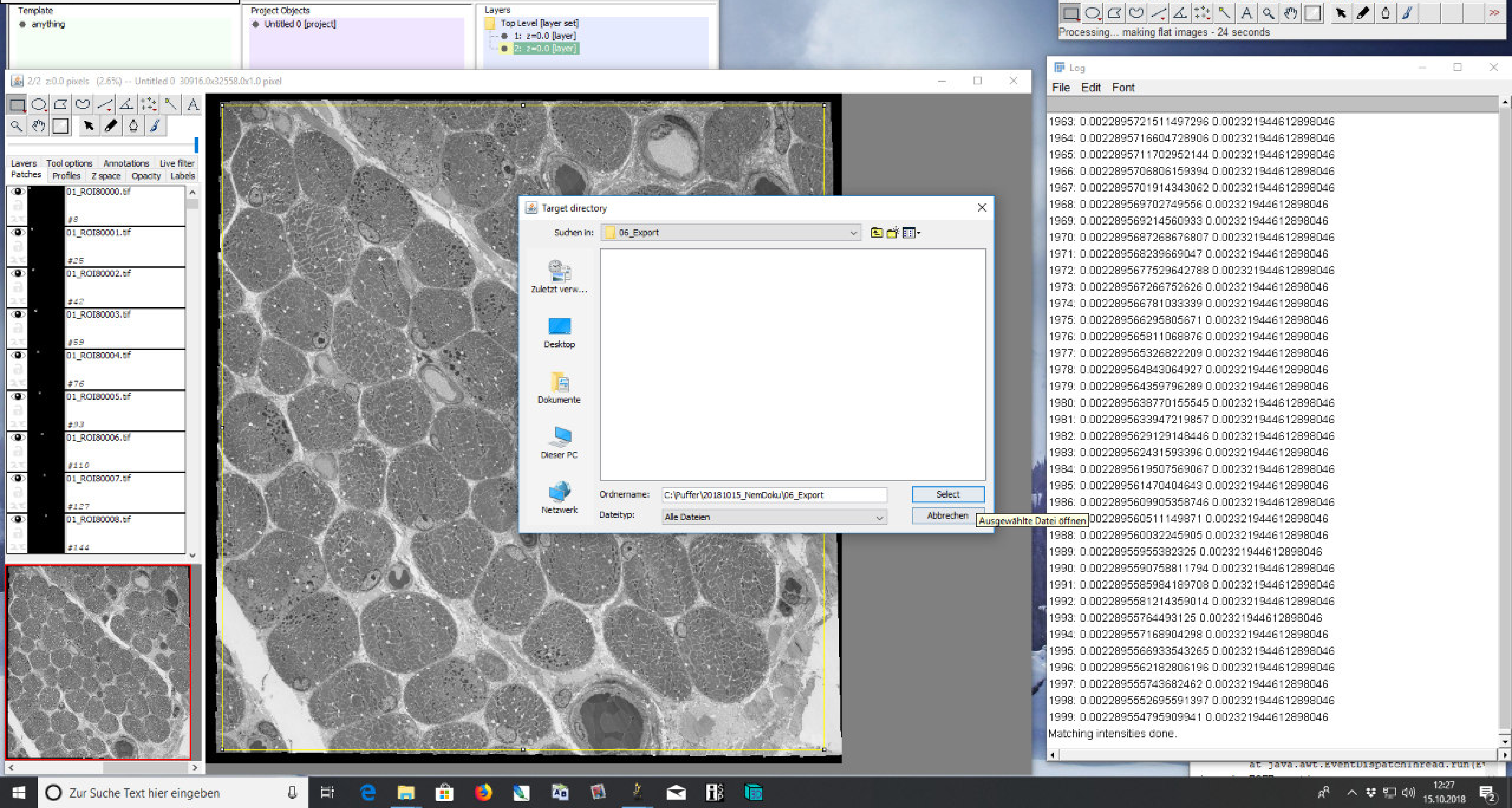

### 9) Standard export

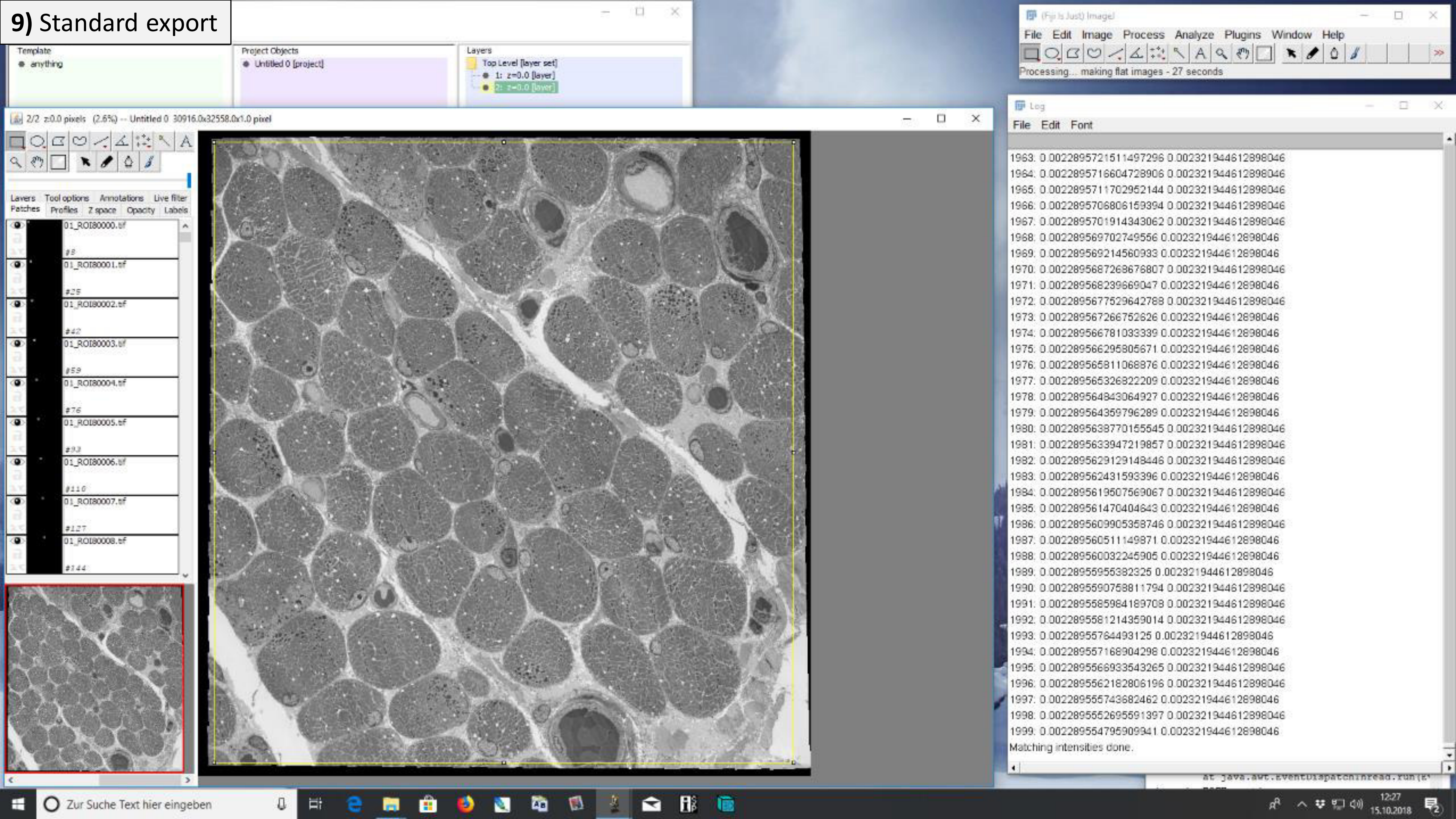

### 9) Standard export

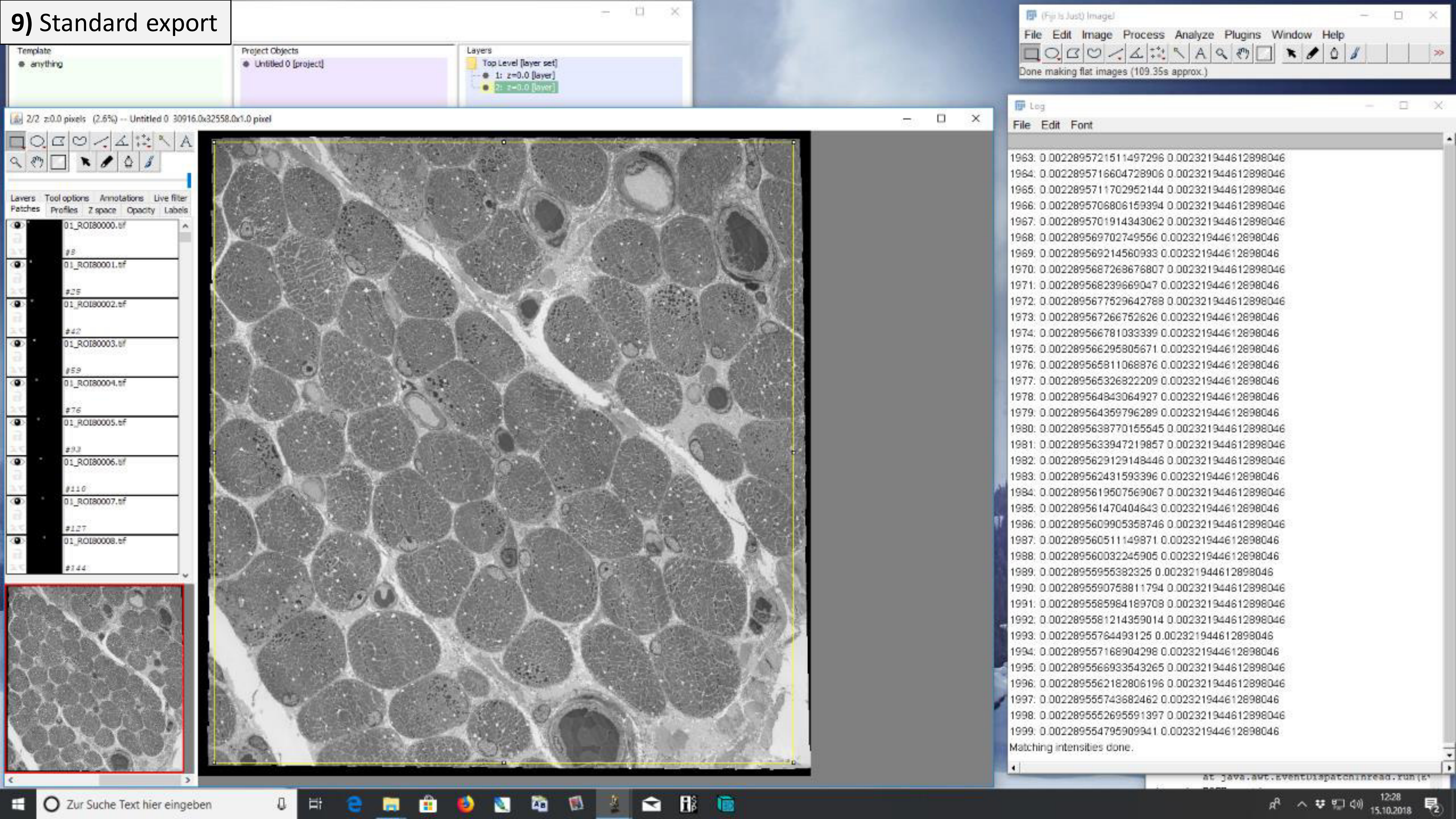

### 10) CATMAID tile export

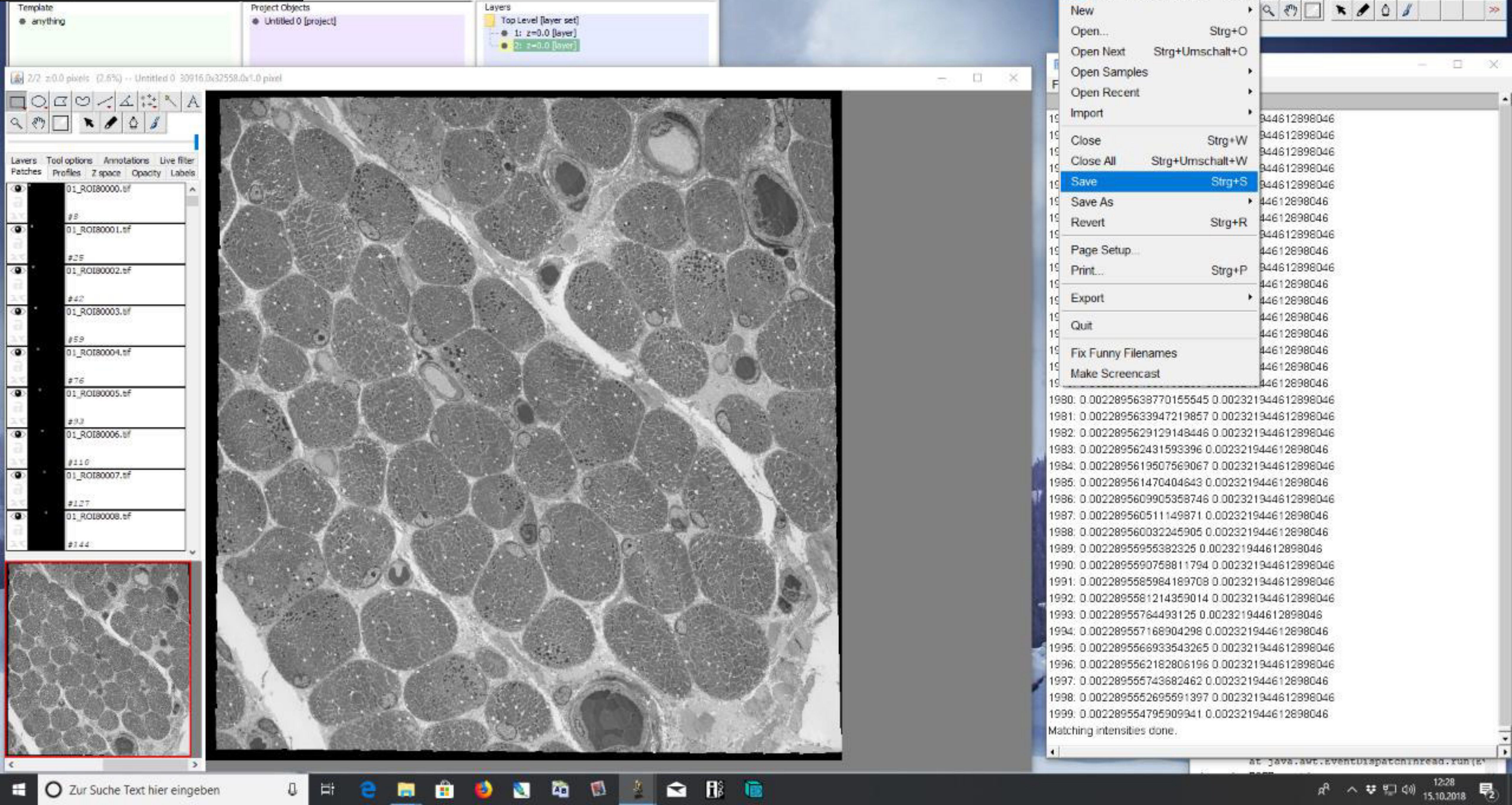

### **10)** CATMAID tile export

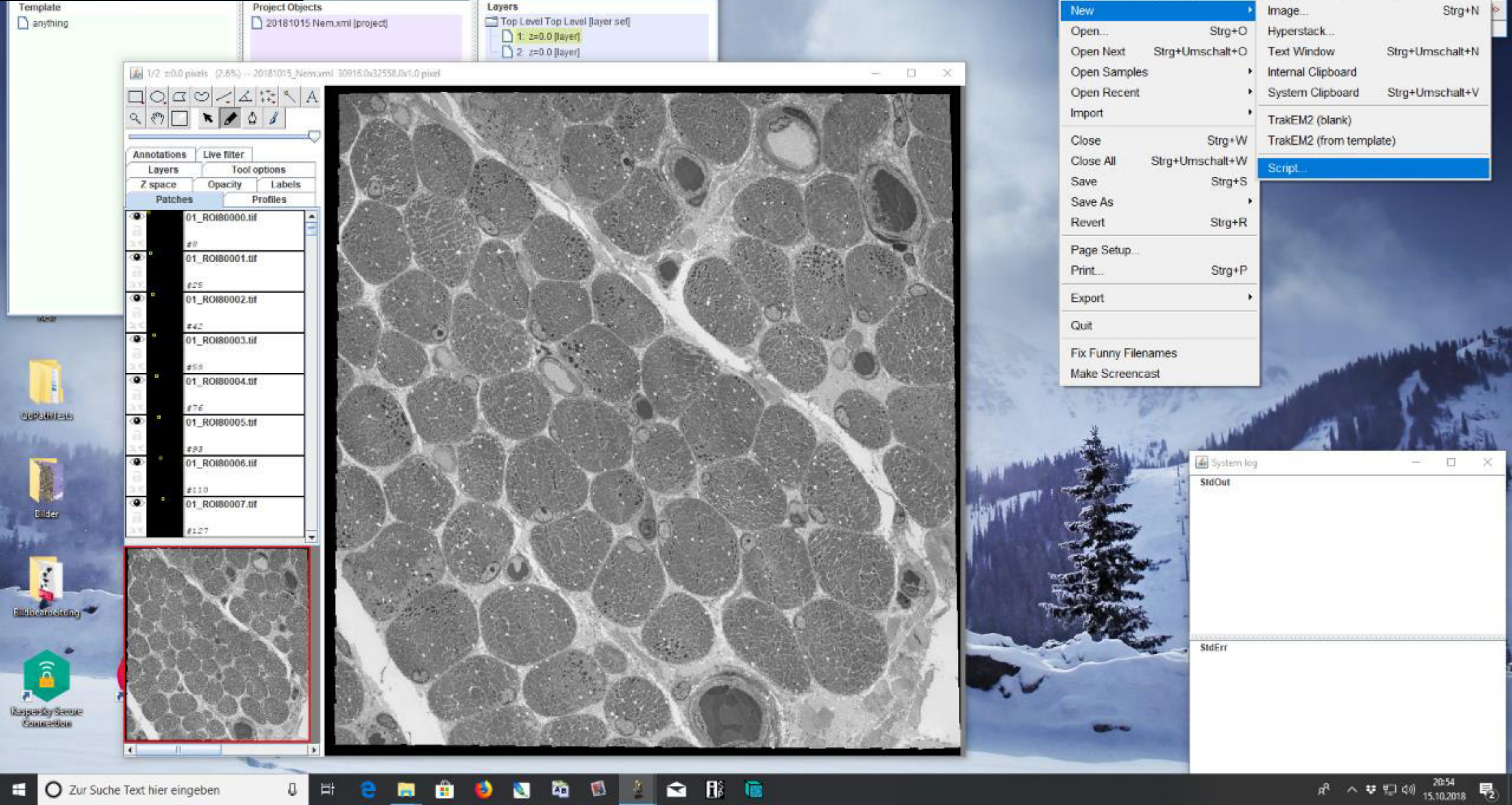

### 10) CATMAID tile export

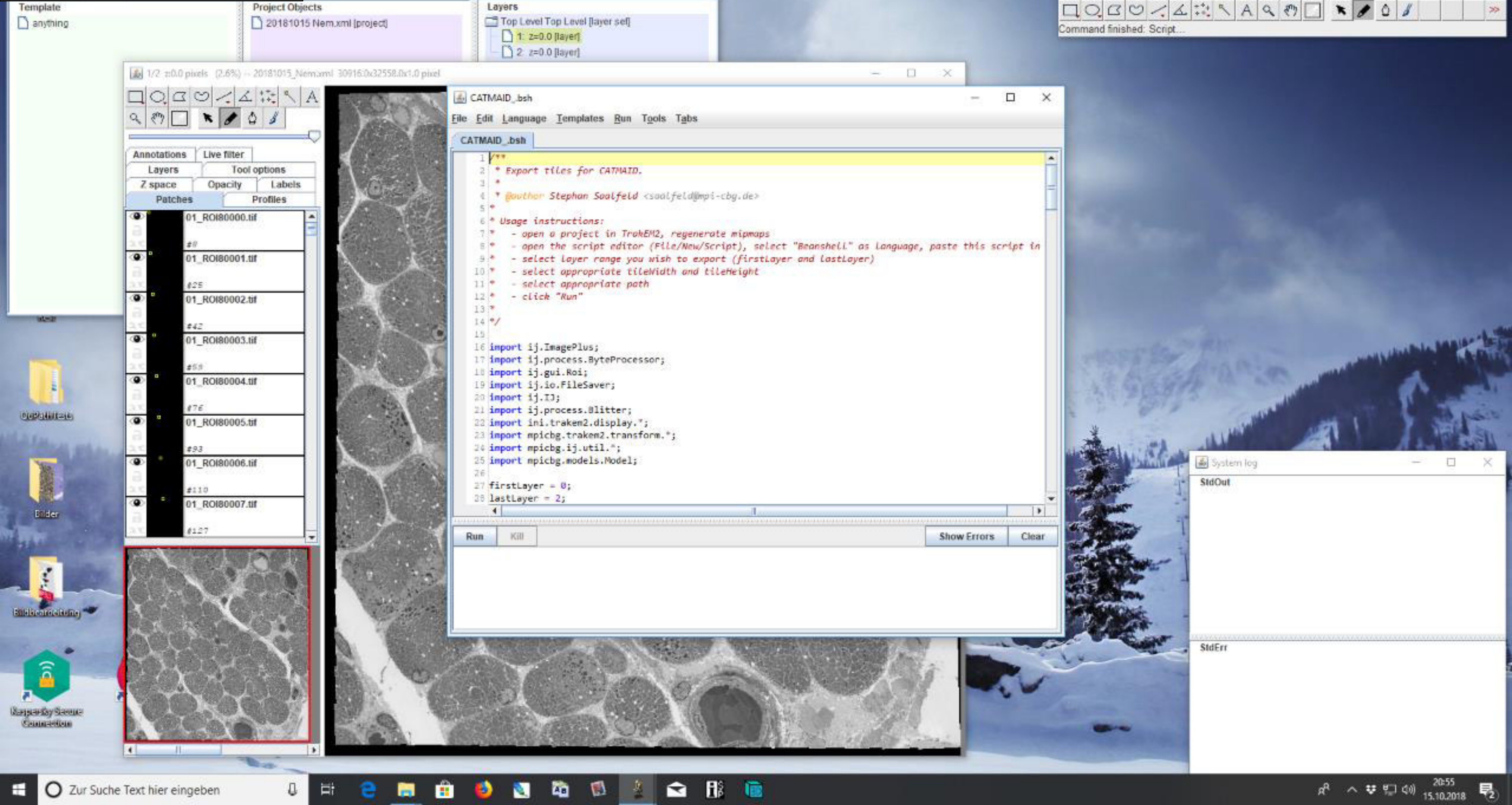

### 10) CATMAID tile export

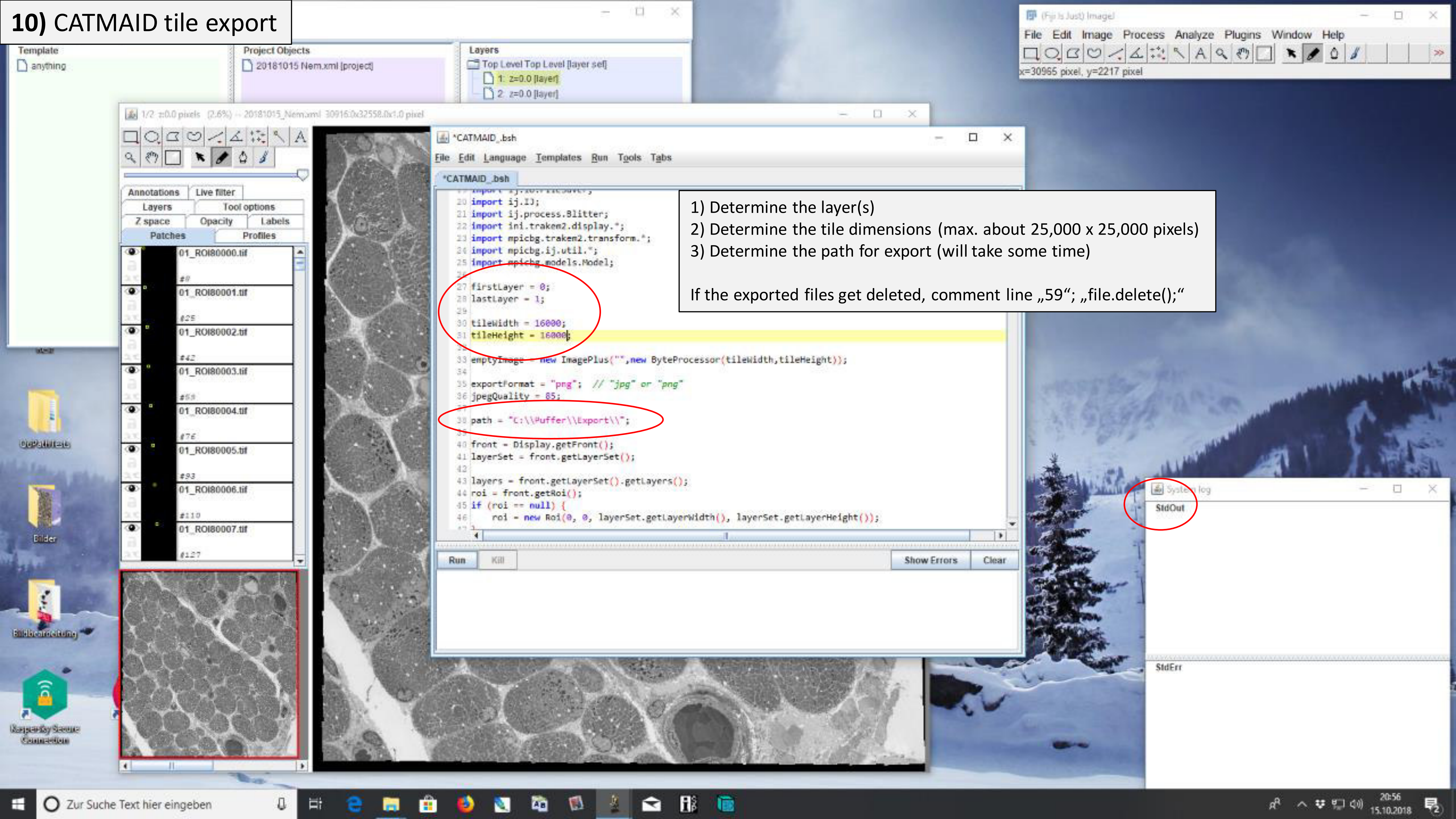


1) Determine the layer(s)
2) Determine the tile dimensions (max. about 25,000 x 25,000 pixels)
3) Determine the path for export (will take some time)

If the exported files get deleted, comment line „59"; „file.delete();"

### 10) CATMAID tile export

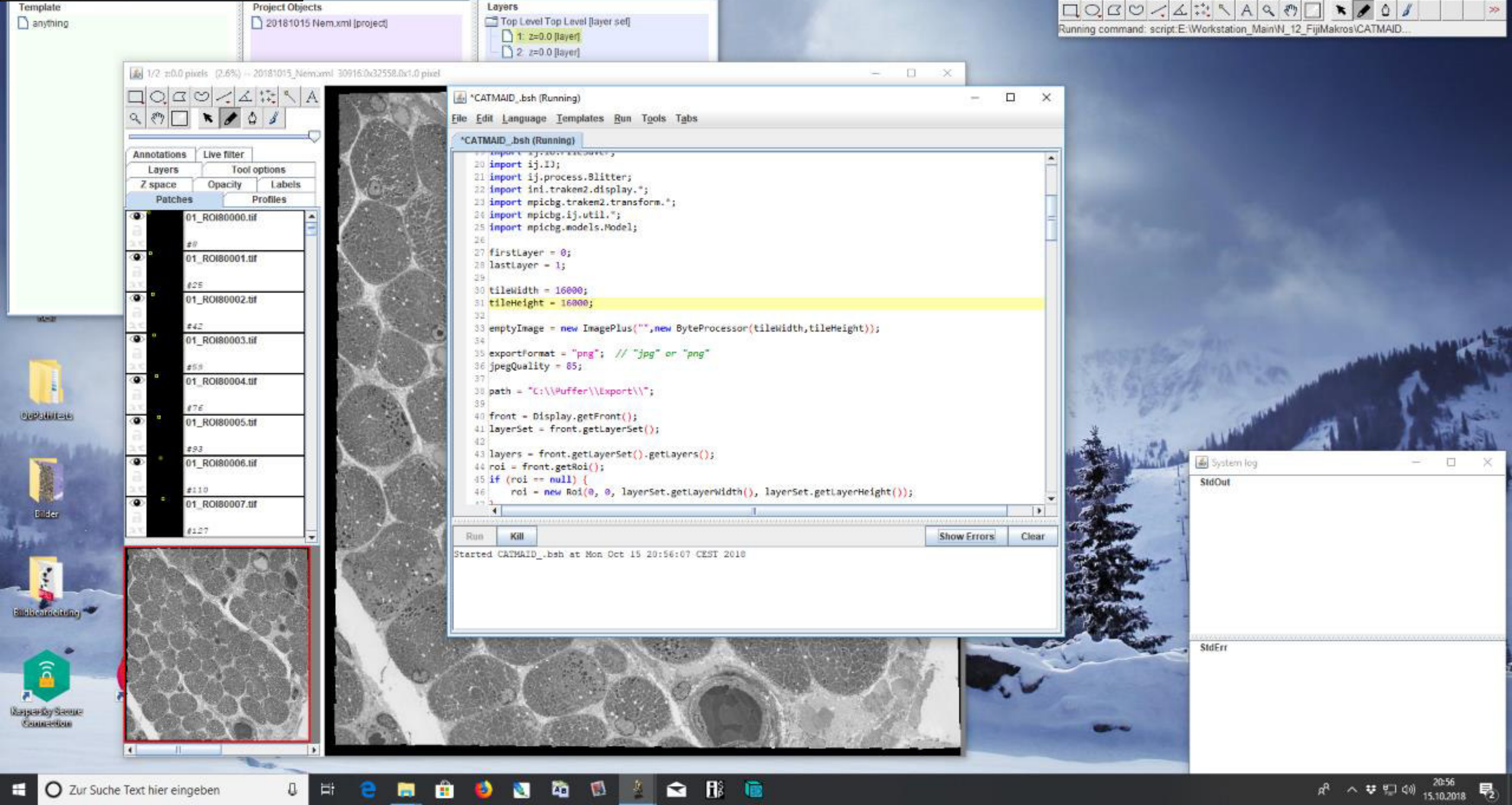

**step-by-step protocol 2 (pdf files A-E)**

Processing of STEM image tiles; stitching and bigtif generation (the process is scalable for up to more than 10 large datasets)

A) Excel; semiautomated generation of a text file containing coordinates (X, Y and Z) of overlapping STEM image tiles

B) Fiji/ TrakEM2 plugin; import of overlapping STEM image tiles and stitching of these into coherent datasets

C) Fiji/ TrakEM2 plugin; export of stitched datasets into non-overlapping tif tiles

D) Nip2; import of non-overlapping tif tiles and export into a coherent bigtif file

E) QuPath; basic workflow for import, pan-and-zoom analysis, annotation, measurement, saving

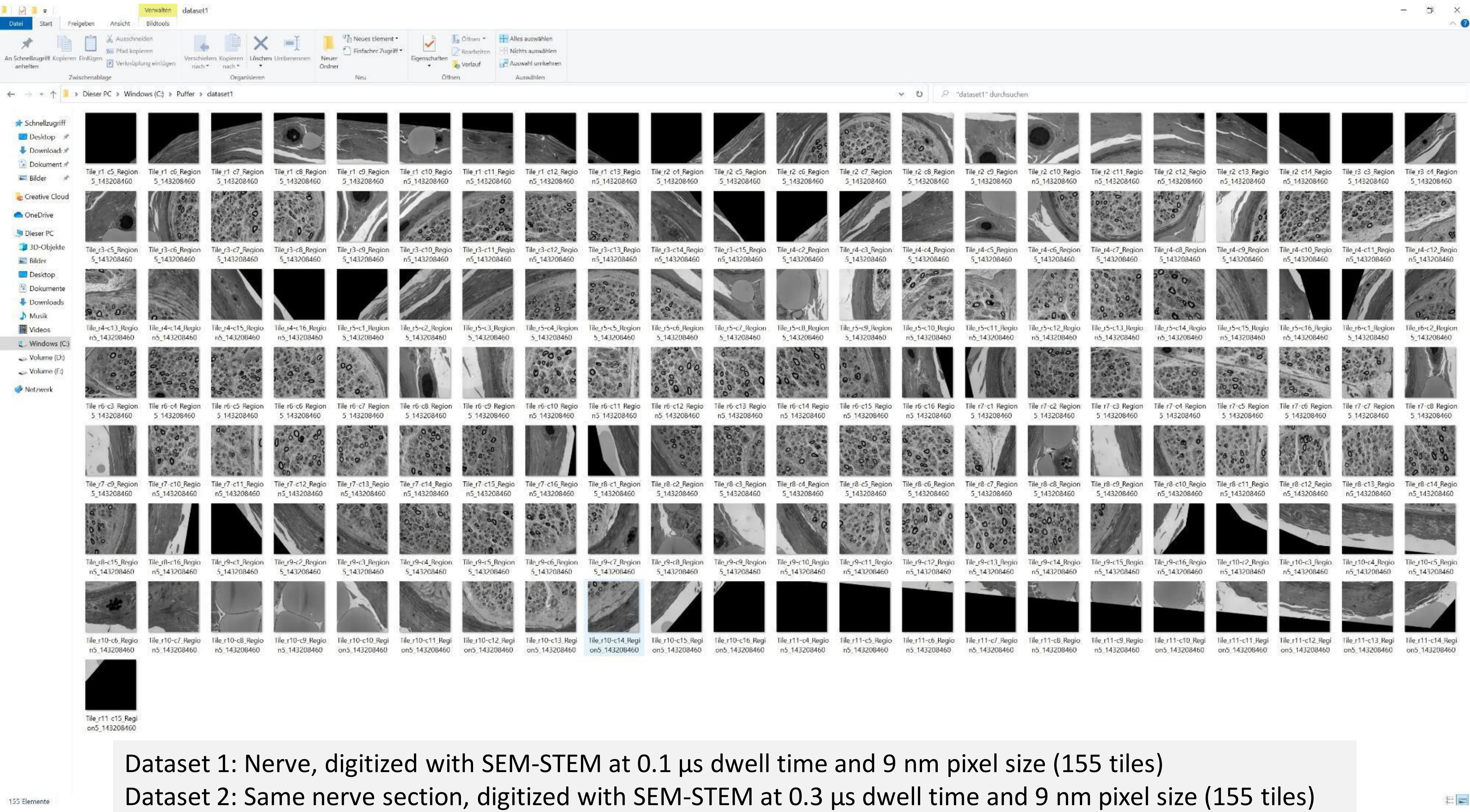


Dataset 1: Nerve, digitized with SEM-STEM at 0.1 µs dwell time and 9 nm pixel size (155 tiles)

Dataset 2: Same nerve section, digitized with SEM-STEM at 0.3 µs dwell time and 9 nm pixel size (155 tiles)

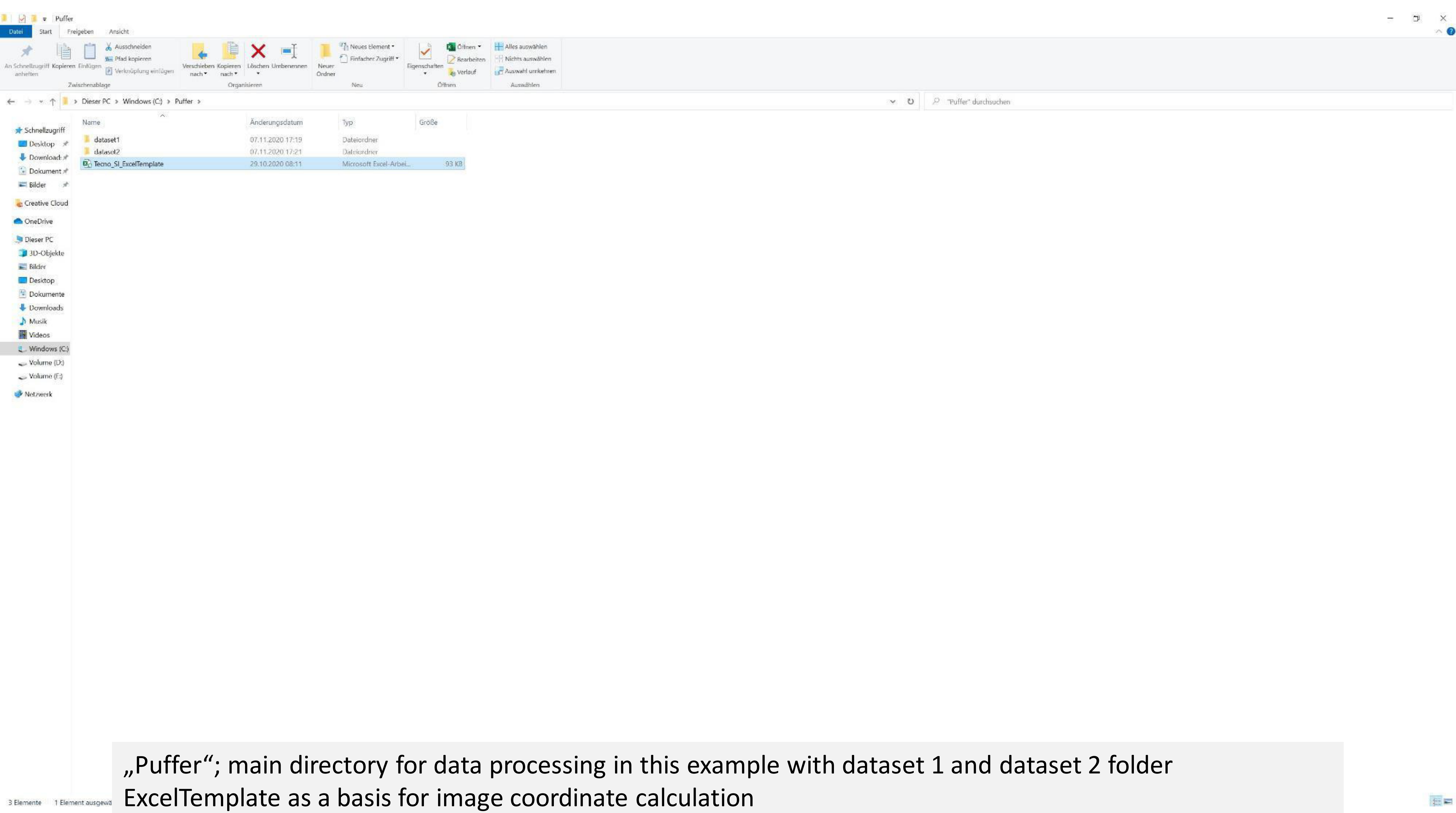


„Puffer"; main directory for data processing in this example with dataset 1 and dataset 2 folder
ExcelTemplate as a basis for image coordinate calculation

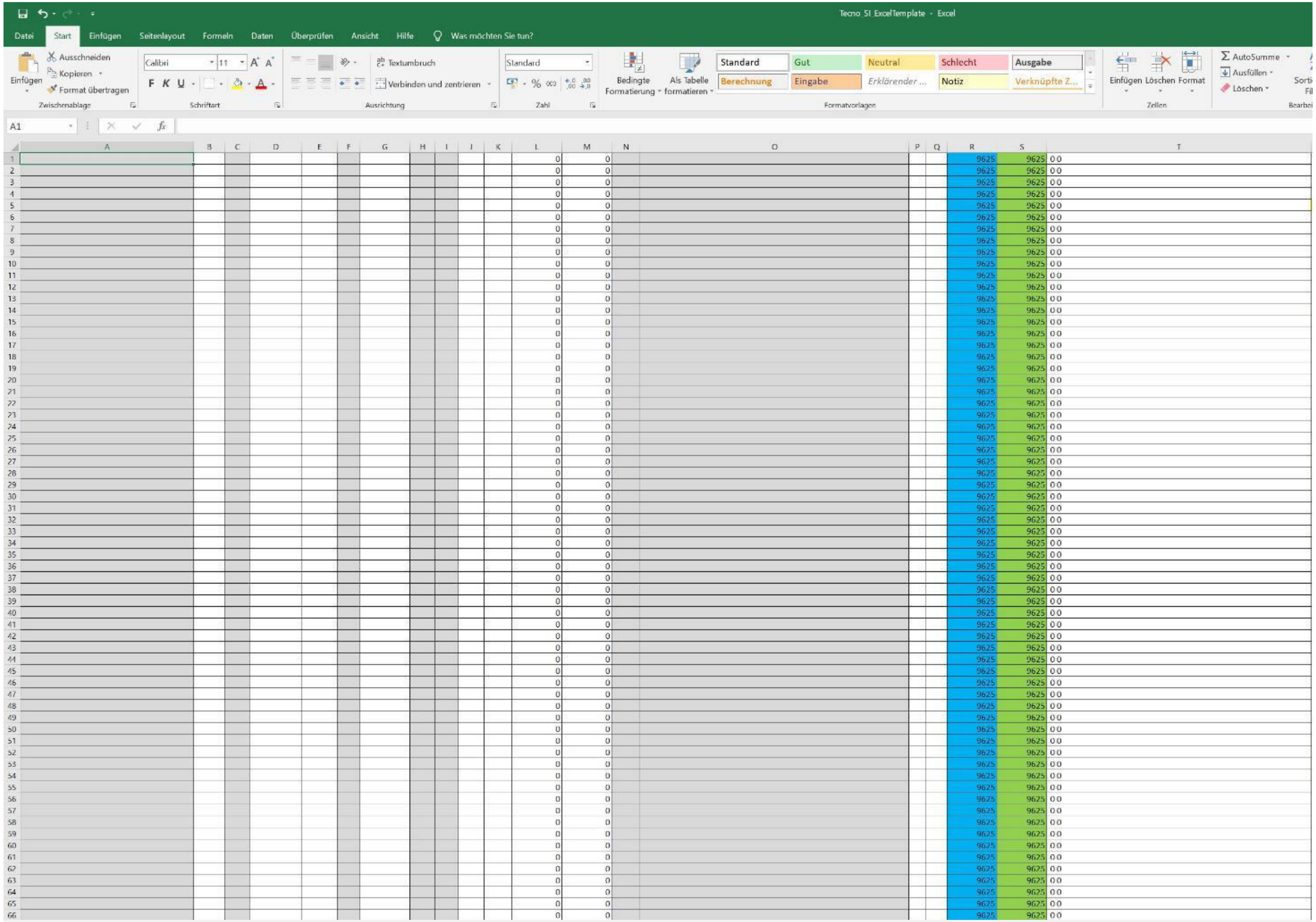

| | L | M | R | S | T |
|---|---|---|---|---|---|
| 1 | 0 | 0 | 9625 | 9625 | 0 0 |
| 2 | 0 | 0 | 9625 | 9625 | 0 0 |
| 3 | 0 | 0 | 9625 | 9625 | 0 0 |
| 4 | 0 | 0 | 9625 | 9625 | 0 0 |
| 5 | 0 | 0 | 9625 | 9625 | 0 0 |
| 6 | 0 | 0 | 9625 | 9625 | 0 0 |
| 7 | 0 | 0 | 9625 | 9625 | 0 0 |
| 8 | 0 | 0 | 9625 | 9625 | 0 0 |
| 9 | 0 | 0 | 9625 | 9625 | 0 0 |
| 10 | 0 | 0 | 9625 | 9625 | 0 0 |
| 11 | 0 | 0 | 9625 | 9625 | 0 0 |
| 12 | 0 | 0 | 9625 | 9625 | 0 0 |
| 13 | 0 | 0 | 9625 | 9625 | 0 0 |
| 14 | 0 | 0 | 9625 | 9625 | 0 0 |
| 15 | 0 | 0 | 9625 | 9625 | 0 0 |
| 16 | 0 | 0 | 9625 | 9625 | 0 0 |
| 17 | 0 | 0 | 9625 | 9625 | 0 0 |
| 18 | 0 | 0 | 9625 | 9625 | 0 0 |
| 19 | 0 | 0 | 9625 | 9625 | 0 0 |
| 20 | 0 | 0 | 9625 | 9625 | 0 0 |
| 21 | 0 | 0 | 9625 | 9625 | 0 0 |
| 22 | 0 | 0 | 9625 | 9625 | 0 0 |
| 23 | 0 | 0 | 9625 | 9625 | 0 0 |
| 24 | 0 | 0 | 9625 | 9625 | 0 0 |
| 25 | 0 | 0 | 9625 | 9625 | 0 0 |
| 26 | 0 | 0 | 9625 | 9625 | 0 0 |
| 27 | 0 | 0 | 9625 | 9625 | 0 0 |
| 28 | 0 | 0 | 9625 | 9625 | 0 0 |
| 29 | 0 | 0 | 9625 | 9625 | 0 0 |
| 30 | 0 | 0 | 9625 | 9625 | 0 0 |
| 31 | 0 | 0 | 9625 | 9625 | 0 0 |
| 32 | 0 | 0 | 9625 | 9625 | 0 0 |
| 33 | 0 | 0 | 9625 | 9625 | 0 0 |
| 34 | 0 | 0 | 9625 | 9625 | 0 0 |
| 35 | 0 | 0 | 9625 | 9625 | 0 0 |
| 36 | 0 | 0 | 9625 | 9625 | 0 0 |
| 37 | 0 | 0 | 9625 | 9625 | 0 0 |
| 38 | 0 | 0 | 9625 | 9625 | 0 0 |
| 39 | 0 | 0 | 9625 | 9625 | 0 0 |
| 40 | 0 | 0 | 9625 | 9625 | 0 0 |
| 41 | 0 | 0 | 9625 | 9625 | 0 0 |
| 42 | 0 | 0 | 9625 | 9625 | 0 0 |
| 43 | 0 | 0 | 9625 | 9625 | 0 0 |
| 44 | 0 | 0 | 9625 | 9625 | 0 0 |
| 45 | 0 | 0 | 9625 | 9625 | 0 0 |
| 46 | 0 | 0 | 9625 | 9625 | 0 0 |
| 47 | 0 | 0 | 9625 | 9625 | 0 0 |
| 48 | 0 | 0 | 9625 | 9625 | 0 0 |
| 49 | 0 | 0 | 9625 | 9625 | 0 0 |
| 50 | 0 | 0 | 9625 | 9625 | 0 0 |
| 51 | 0 | 0 | 9625 | 9625 | 0 0 |
| 52 | 0 | 0 | 9625 | 9625 | 0 0 |
| 53 | 0 | 0 | 9625 | 9625 | 0 0 |
| 54 | 0 | 0 | 9625 | 9625 | 0 0 |
| 55 | 0 | 0 | 9625 | 9625 | 0 0 |
| 56 | 0 | 0 | 9625 | 9625 | 0 0 |
| 57 | 0 | 0 | 9625 | 9625 | 0 0 |
| 58 | 0 | 0 | 9625 | 9625 | 0 0 |
| 59 | 0 | 0 | 9625 | 9625 | 0 0 |
| 60 | 0 | 0 | 9625 | 9625 | 0 0 |
| 61 | 0 | 0 | 9625 | 9625 | 0 0 |
| 62 | 0 | 0 | 9625 | 9625 | 0 0 |
| 63 | 0 | 0 | 9625 | 9625 | 0 0 |
| 64 | 0 | 0 | 9625 | 9625 | 0 0 |
| 65 | 0 | 0 | 9625 | 9625 | 0 0 |
| 66 | 0 | 0 | 9625 | 9625 | 0 0 |

Details are described in the Excel file (Description)

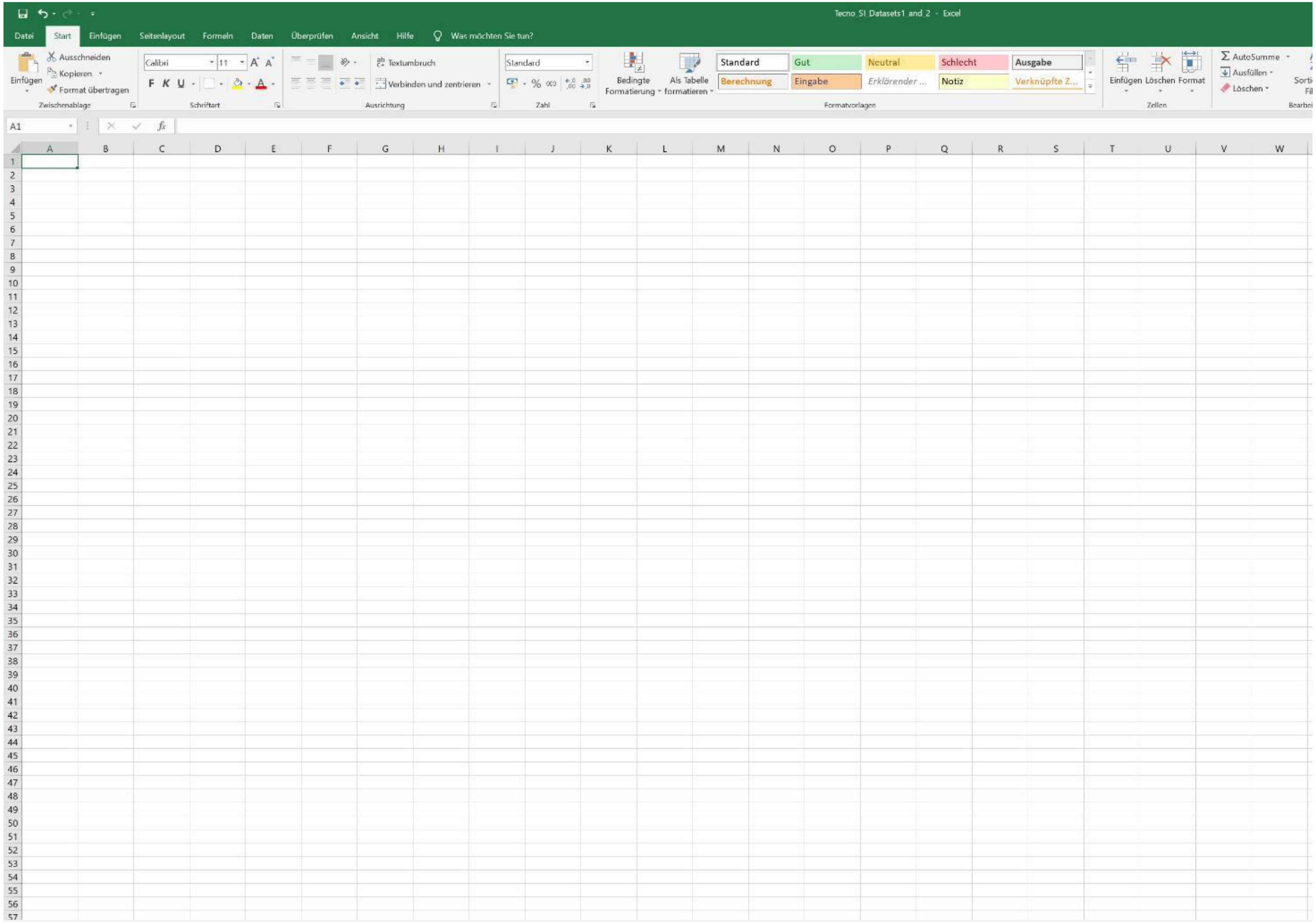
Tecno SI Datasets1 and 2 - Excel
Datei
Start
Einfügen
Seitenlayout
Formeln
Daten
Überprüfen
Ansicht
Hilfe
Was möchten Sie tun?
Ausschneiden
Kopieren
Format übertragen
Einfügen
Zwischenablage
Calibri
11
F K U
Schriftart
Textumbruch
Verbinden und zentrieren
Ausrichtung
Standard
Zahl
Bedingte Formatierung
Als Tabelle formatieren
Standard
Gut
Neutral
Schlecht
Ausgabe
Berechnung
Eingabe
Erklärender ...
Notiz
Verknüpfte Z...
Formatvorlagen
Einfügen
Löschen
Format
Zellen
AutoSumme
Ausfüllen
Löschen
A1

New page

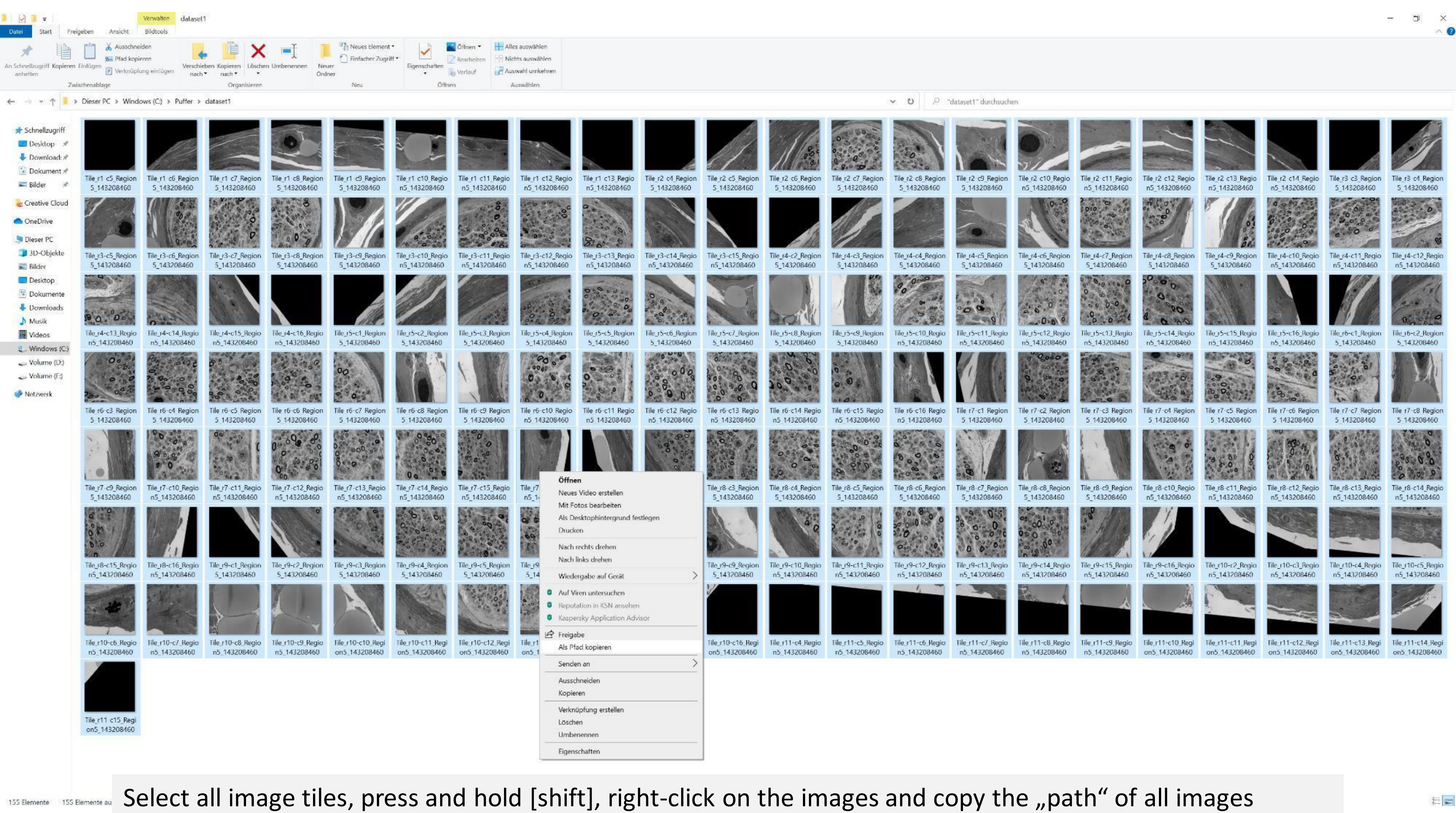


Select all image tiles, press and hold [shift], right-click on the images and copy the „path“ of all images

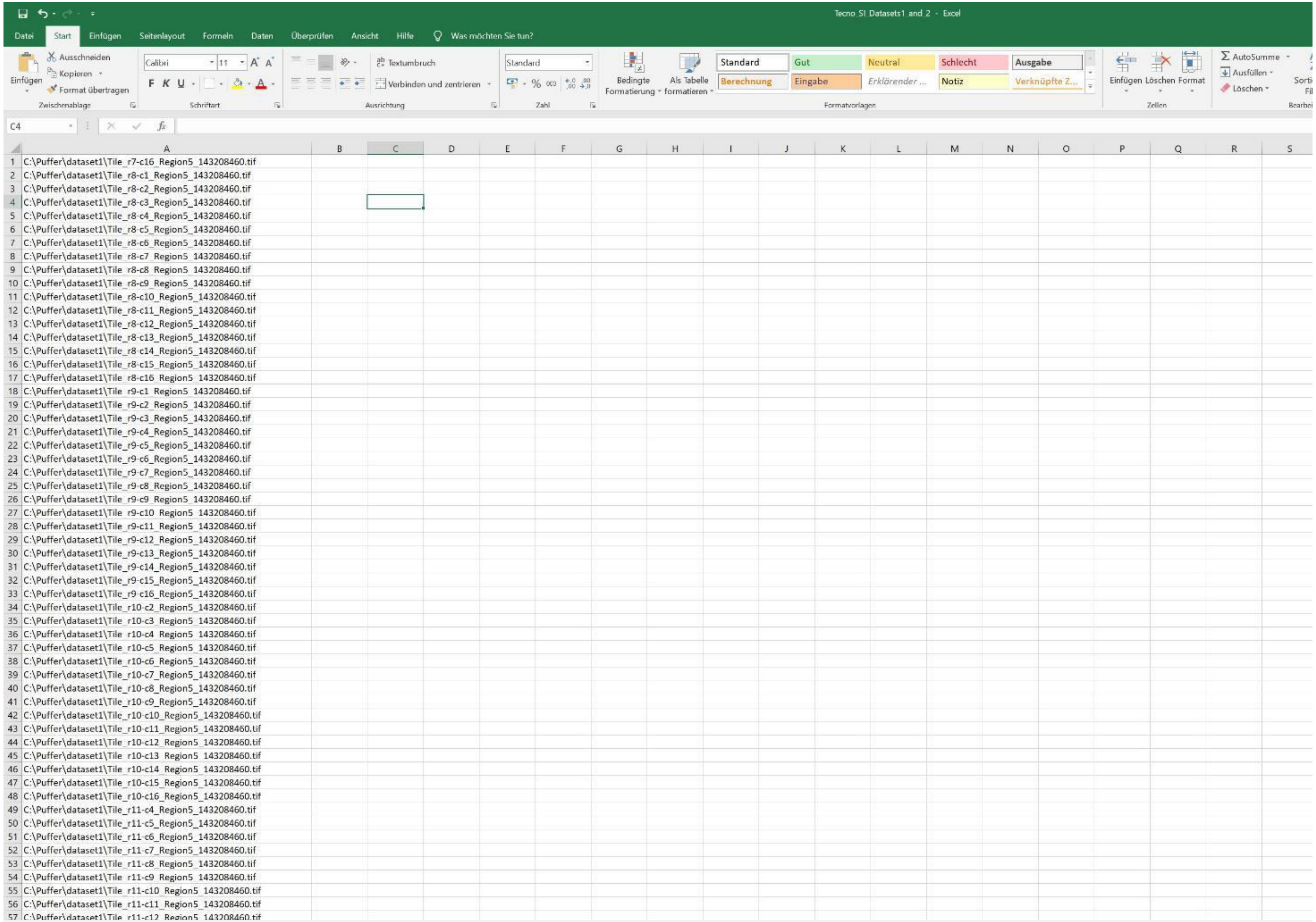


Copy the „path" of all images into the new Excel page

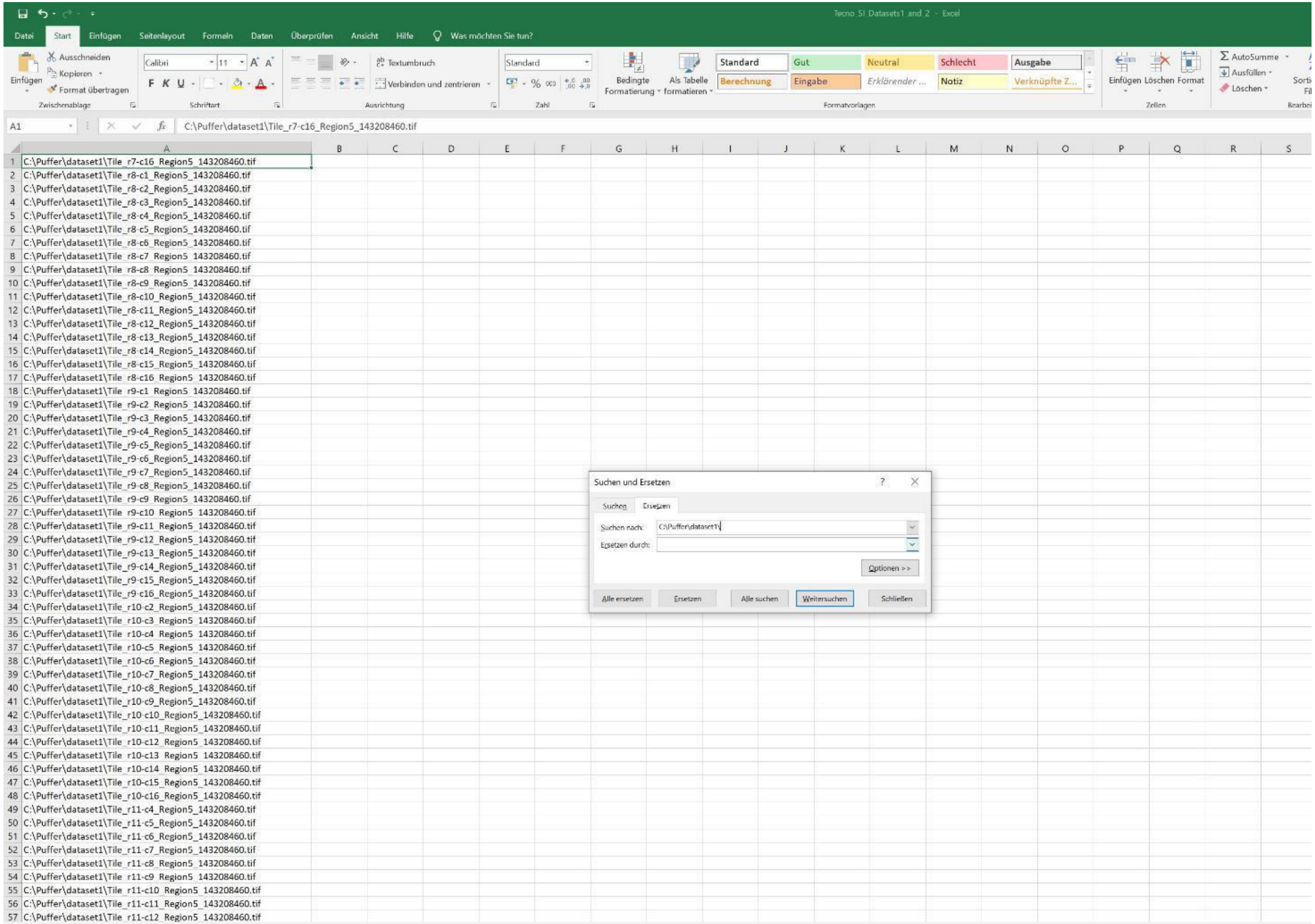


Exchange the „path“ by „nothing“, thus, only the image tile names will remain

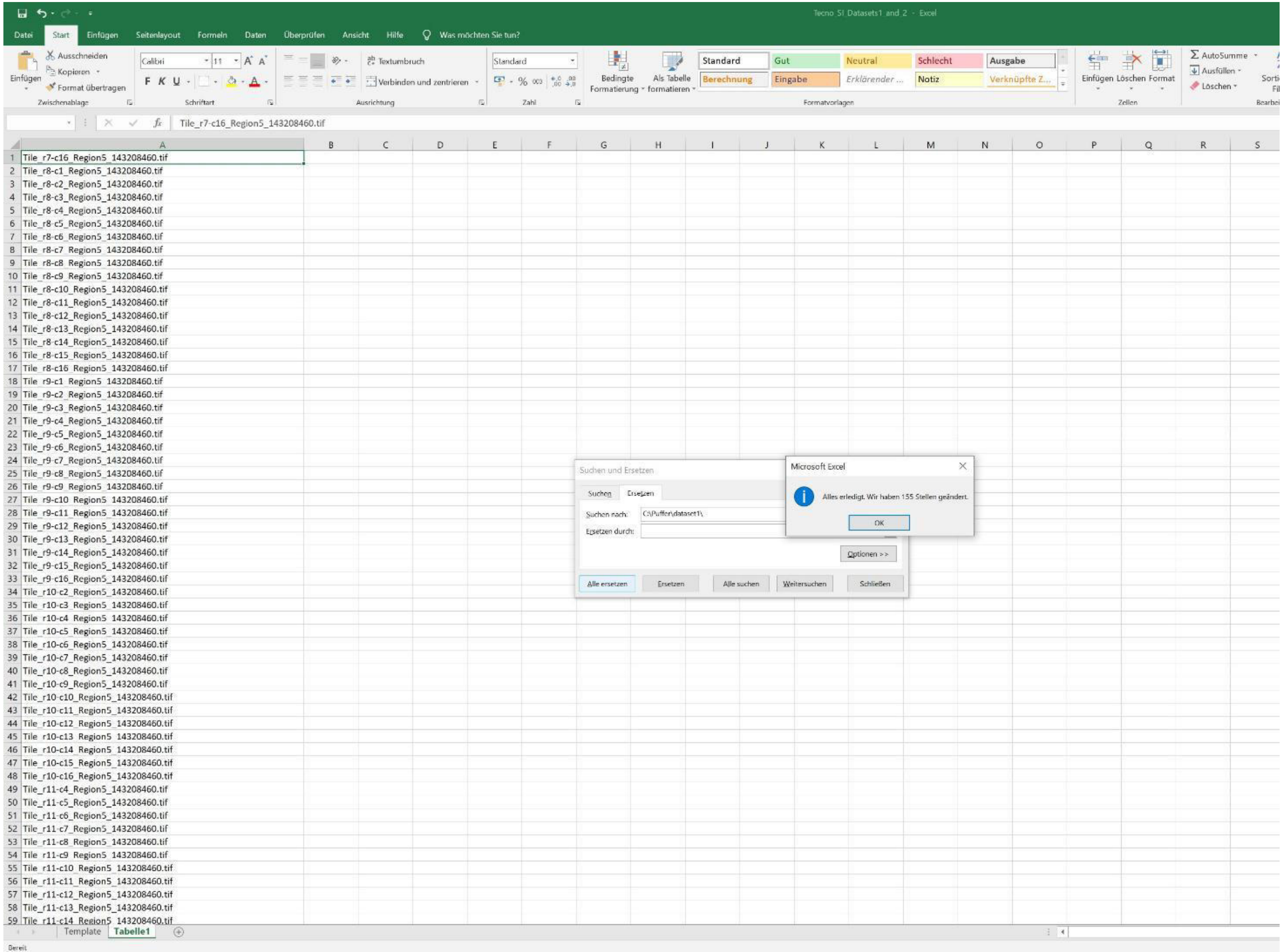
Tecno SI Datasets1 and 2 - Excel
Datei
Start
Einfügen
Seitenlayout
Formeln
Daten
Überprüfen
Ansicht
Hilfe
Was möchten Sie tun?
Ausschneiden
Kopieren
Format übertragen
Einfügen
Zwischenablage
Calibri
11
Schriftart
Textumbruch
Verbinden und zentrieren
Ausrichtung
Standard
Zahl
Bedingte Formatierung
Als Tabelle formatieren
Standard
Gut
Neutral
Schlecht
Ausgabe
Berechnung
Eingabe
Erklärender ...
Notiz
Verknüpfte Z...
Formatvorlagen
Einfügen
Löschen
Format
Zellen
AutoSumme
Ausfüllen
Löschen
Tile_r7-c16_Region5_143208460.tif
Tile_r8-c1_Region5_143208460.tif
Tile_r8-c2_Region5_143208460.tif
Tile_r8-c3_Region5_143208460.tif
Tile_r8-c4_Region5_143208460.tif
Tile_r8-c5_Region5_143208460.tif
Tile_r8-c6_Region5_143208460.tif
Tile_r8-c7_Region5_143208460.tif
Tile_r8-c8_Region5_143208460.tif
Tile_r8-c9_Region5_143208460.tif
Tile_r8-c10_Region5_143208460.tif
Tile_r8-c11_Region5_143208460.tif
Tile_r8-c12_Region5_143208460.tif
Tile_r8-c13_Region5_143208460.tif
Tile_r8-c14_Region5_143208460.tif
Tile_r8-c15_Region5_143208460.tif
Tile_r8-c16_Region5_143208460.tif
Tile_r9-c1_Region5_143208460.tif
Tile_r9-c2_Region5_143208460.tif
Tile_r9-c3_Region5_143208460.tif
Tile_r9-c4_Region5_143208460.tif
Tile_r9-c5_Region5_143208460.tif
Tile_r9-c6_Region5_143208460.tif
Tile_r9-c7_Region5_143208460.tif
Tile_r9-c8_Region5_143208460.tif
Tile_r9-c9_Region5_143208460.tif
Tile_r9-c10_Region5_143208460.tif
Tile_r9-c11_Region5_143208460.tif
Tile_r9-c12_Region5_143208460.tif
Tile_r9-c13_Region5_143208460.tif
Tile_r9-c14_Region5_143208460.tif
Tile_r9-c15_Region5_143208460.tif
Tile_r9-c16_Region5_143208460.tif
Tile_r10-c2_Region5_143208460.tif
Tile_r10-c3_Region5_143208460.tif
Tile_r10-c4_Region5_143208460.tif
Tile_r10-c5_Region5_143208460.tif
Tile_r10-c6_Region5_143208460.tif
Tile_r10-c7_Region5_143208460.tif
Tile_r10-c8_Region5_143208460.tif
Tile_r10-c9_Region5_143208460.tif
Tile_r10-c10_Region5_143208460.tif
Tile_r10-c11_Region5_143208460.tif
Tile_r10-c12_Region5_143208460.tif
Tile_r10-c13_Region5_143208460.tif
Tile_r10-c14_Region5_143208460.tif
Tile_r10-c15_Region5_143208460.tif
Tile_r10-c16_Region5_143208460.tif
Tile_r11-c4_Region5_143208460.tif
Tile_r11-c5_Region5_143208460.tif
Tile_r11-c6_Region5_143208460.tif
Tile_r11-c7_Region5_143208460.tif
Tile_r11-c8_Region5_143208460.tif
Tile_r11-c9_Region5_143208460.tif
Tile_r11-c10_Region5_143208460.tif
Tile_r11-c11_Region5_143208460.tif
Tile_r11-c12_Region5_143208460.tif
Tile_r11-c13_Region5_143208460.tif
Tile_r11-c14_Region5_143208460.tif
Suchen und Ersetzen
Suchen
Ersetzen
Suchen nach:
C:\Puffer\dataset1\
Ersetzen durch:
Optionen >>
Alle ersetzen
Ersetzen
Alle suchen
Weitersuchen
Schließen
Microsoft Excel
Alles erledigt. Wir haben 155 Stellen geändert.
OK
Template
Tabelle1
Bereit

Tecno SI Datasets1 and 2 - Excel

A1 | $f_x$ | Tile_r7-c16_Region5_143208460.tif

| | A | B | E | L | M | R | S | T |
|---|---|---|---|---|---|---|---|---|
| 1 | Tile_r7-c16_Region5_143208460.tif | 7- | c16 | 0 | 0 | 9625 | 9625 | 0 0 |
| 2 | Tile_r8-c1_Region5_143208460.tif | 8- | c1_R | 0 | 0 | 9625 | 9625 | 0 0 |
| 3 | Tile_r8-c2_Region5_143208460.tif | 8- | c2_R | 0 | 0 | 9625 | 9625 | 0 0 |
| 4 | Tile_r8-c3_Region5_143208460.tif | 8- | c3_R | 0 | 0 | 9625 | 9625 | 0 0 |
| 5 | Tile_r8-c4_Region5_143208460.tif | 8- | c4_R | 0 | 0 | 9625 | 9625 | 0 0 |
| 6 | Tile_r8-c5_Region5_143208460.tif | 8- | c5_R | 0 | 0 | 9625 | 9625 | 0 0 |
| 7 | Tile_r8-c6_Region5_143208460.tif | 8- | c6_R | 0 | 0 | 9625 | 9625 | 0 0 |
| 8 | Tile_r8-c7_Region5_143208460.tif | 8- | c7_R | 0 | 0 | 9625 | 9625 | 0 0 |
| 9 | Tile_r8-c8_Region5_143208460.tif | 8- | c8_R | 0 | 0 | 9625 | 9625 | 0 0 |
| 10 | Tile_r8-c9_Region5_143208460.tif | 8- | c9_R | 0 | 0 | 9625 | 9625 | 0 0 |
| 11 | Tile_r8-c10_Region5_143208460.tif | 8- | c10_ | 0 | 0 | 9625 | 9625 | 0 0 |
| 12 | Tile_r8-c11_Region5_143208460.tif | 8- | c11_ | 0 | 0 | 9625 | 9625 | 0 0 |
| 13 | Tile_r8-c12_Region5_143208460.tif | 8- | c12_ | 0 | 0 | 9625 | 9625 | 0 0 |
| 14 | Tile_r8-c13_Region5_143208460.tif | 8- | c13_ | 0 | 0 | 9625 | 9625 | 0 0 |
| 15 | Tile_r8-c14_Region5_143208460.tif | 8- | c14_ | 0 | 0 | 9625 | 9625 | 0 0 |
| 16 | Tile_r8-c15_Region5_143208460.tif | 8- | c15_ | 0 | 0 | 9625 | 9625 | 0 0 |
| 17 | Tile_r8-c16_Region5_143208460.tif | 8- | c16_ | 0 | 0 | 9625 | 9625 | 0 0 |
| 18 | Tile_r9-c1_Region5_143208460.tif | 9- | c1_R | 0 | 0 | 9625 | 9625 | 0 0 |
| 19 | Tile_r9-c2_Region5_143208460.tif | 9- | c2_R | 0 | 0 | 9625 | 9625 | 0 0 |
| 20 | Tile_r9-c3_Region5_143208460.tif | 9- | c3_R | 0 | 0 | 9625 | 9625 | 0 0 |
| 21 | Tile_r9-c4_Region5_143208460.tif | 9- | c4_R | 0 | 0 | 9625 | 9625 | 0 0 |
| 22 | Tile_r9-c5_Region5_143208460.tif | 9- | c5_R | 0 | 0 | 9625 | 9625 | 0 0 |
| 23 | Tile_r9-c6_Region5_143208460.tif | 9 | c6_R | 0 | 0 | 9625 | 9625 | 0 0 |
| 24 | Tile_r9-c7_Region5_143208460.tif | 9 | c7_R | 0 | 0 | 9625 | 9625 | 0 0 |
| 25 | Tile_r9-c8_Region5_143208460.tif | 9 | c8_R | 0 | 0 | 9625 | 9625 | 0 0 |
| 26 | Tile_r9-c9_Region5_143208460.tif | 9 | c9_R | 0 | 0 | 9625 | 9625 | 0 0 |
| 27 | Tile_r9-c10_Region5_143208460.tif | 9 | c10_ | 0 | 0 | 9625 | 9625 | 0 0 |
| 28 | Tile_r9-c11_Region5_143208460.tif | 9 | c11_ | 0 | 0 | 9625 | 9625 | 0 0 |
| 29 | Tile_r9-c12_Region5_143208460.tif | 9- | c12_ | 0 | 0 | 9625 | 9625 | 0 0 |
| 30 | Tile_r9-c13_Region5_143208460.tif | 9- | c13_ | 0 | 0 | 9625 | 9625 | 0 0 |
| 31 | Tile_r9-c14_Region5_143208460.tif | 9- | c14_ | 0 | 0 | 9625 | 9625 | 0 0 |
| 32 | Tile_r9-c15_Region5_143208460.tif | 9- | c15_ | 0 | 0 | 9625 | 9625 | 0 0 |
| 33 | Tile_r9-c16_Region5_143208460.tif | 9- | c16_ | 0 | 0 | 9625 | 9625 | 0 0 |
| 34 | Tile_r10-c2_Region5_143208460.tif | 10 | -c2_ | 0 | 0 | 9625 | 9625 | 0 0 |
| 35 | Tile_r10-c3_Region5_143208460.tif | 10 | -c3_ | 0 | 0 | 9625 | 9625 | 0 0 |
| 36 | Tile_r10-c4_Region5_143208460.tif | 10 | -c4_ | 0 | 0 | 9625 | 9625 | 0 0 |
| 37 | Tile_r10-c5_Region5_143208460.tif | 10 | -c5_ | 0 | 0 | 9625 | 9625 | 0 0 |
| 38 | Tile_r10-c6_Region5_143208460.tif | 10 | -c6_ | 0 | 0 | 9625 | 9625 | 0 0 |
| 39 | Tile_r10-c7_Region5_143208460.tif | 10 | -c7 | 0 | 0 | 9625 | 9625 | 0 0 |
| 40 | Tile_r10-c8_Region5_143208460.tif | 10 | -c8 | 0 | 0 | 9625 | 9625 | 0 0 |
| 41 | Tile_r10-c9_Region5_143208460.tif | 10 | -c9 | 0 | 0 | 9625 | 9625 | 0 0 |
| 42 | Tile_r10-c10_Region5_143208460.tif | 10 | -c10 | 0 | 0 | 9625 | 9625 | 0 0 |
| 43 | Tile_r10-c11_Region5_143208460.tif | 10 | -c11 | 0 | 0 | 9625 | 9625 | 0 0 |
| 44 | Tile_r10-c12_Region5_143208460.tif | 10 | -c12 | 0 | 0 | 9625 | 9625 | 0 0 |
| 45 | Tile_r10-c13_Region5_143208460.tif | 10 | -c13 | 0 | 0 | 9625 | 9625 | 0 0 |
| 46 | Tile_r10-c14_Region5_143208460.tif | 10 | -c14 | 0 | 0 | 9625 | 9625 | 0 0 |
| 47 | Tile_r10-c15_Region5_143208460.tif | 10 | -c15 | 0 | 0 | 9625 | 9625 | 0 0 |
| 48 | Tile_r10-c16_Region5_143208460.tif | 10 | -c16 | 0 | 0 | 9625 | 9625 | 0 0 |
| 49 | Tile_r11-c4_Region5_143208460.tif | 11 | -c4_ | 0 | 0 | 9625 | 9625 | 0 0 |
| 50 | Tile_r11-c5_Region5_143208460.tif | 11 | -c5_ | 0 | 0 | 9625 | 9625 | 0 0 |
| 51 | Tile_r11-c6_Region5_143208460.tif | 11 | -c6_ | 0 | 0 | 9625 | 9625 | 0 0 |
| 52 | Tile_r11-c7_Region5_143208460.tif | 11 | -c7_ | 0 | 0 | 9625 | 9625 | 0 0 |
| 53 | Tile_r11-c8_Region5_143208460.tif | 11 | -c8_ | 0 | 0 | 9625 | 9625 | 0 0 |
| 54 | Tile_r11-c9_Region5_143208460.tif | 11 | -c9_ | 0 | 0 | 9625 | 9625 | 0 0 |
| 55 | Tile_r11-c10_Region5_143208460.tif | 11 | -c10 | 0 | 0 | 9625 | 9625 | 0 0 |
| 56 | Tile_r11-c11_Region5_143208460.tif | 11 | -c11 | 0 | 0 | 9625 | 9625 | 0 0 |
| 57 | Tile_r11-c12_Region5_143208460.tif | 11 | -c12 | 0 | 0 | 9625 | 9625 | 0 0 |
| 58 | Tile_r11-c13_Region5_143208460.tif | 11 | -c13 | 0 | 0 | 9625 | 9625 | 0 0 |
| 59 | Tile_r11-c14_Region5_143208460.tif | 11 | -c14 | 0 | 0 | 9625 | 9625 | 0 0 |
| 60 | Tile_r11-c15_Region5_143208460.tif | 11 | -c15 | 0 | 0 | 9625 | 9625 | 0 0 |
| 61 | Tile_r1-c5_Region5_143208460.tif | 1- | c5_R | 0 | 0 | 9625 | 9625 | 0 0 |
| 62 | Tile_r1-c6_Region5_143208460.tif | 1- | c6_R | 0 | 0 | 9625 | 9625 | 0 0 |
| 63 | Tile_r1-c7_Region5_143208460.tif | 1- | c7_R | 0 | 0 | 9625 | 9625 | 0 0 |
| 64 | Tile_r1-c8_Region5_143208460.tif | 1- | c8_R | 0 | 0 | 9625 | 9625 | 0 0 |
| 65 | Tile_r1-c9_Region5_143208460.tif | 1 | c9_R | 0 | 0 | 9625 | 9625 | 0 0 |
| 66 | Tile_r1-c10_Region5_143208460.tif | 1 | c10_ | 0 | 0 | 9625 | 9625 | 0 0 |

Copy the image tile names (of dataset 1) into the Template (first clear all gray areas)

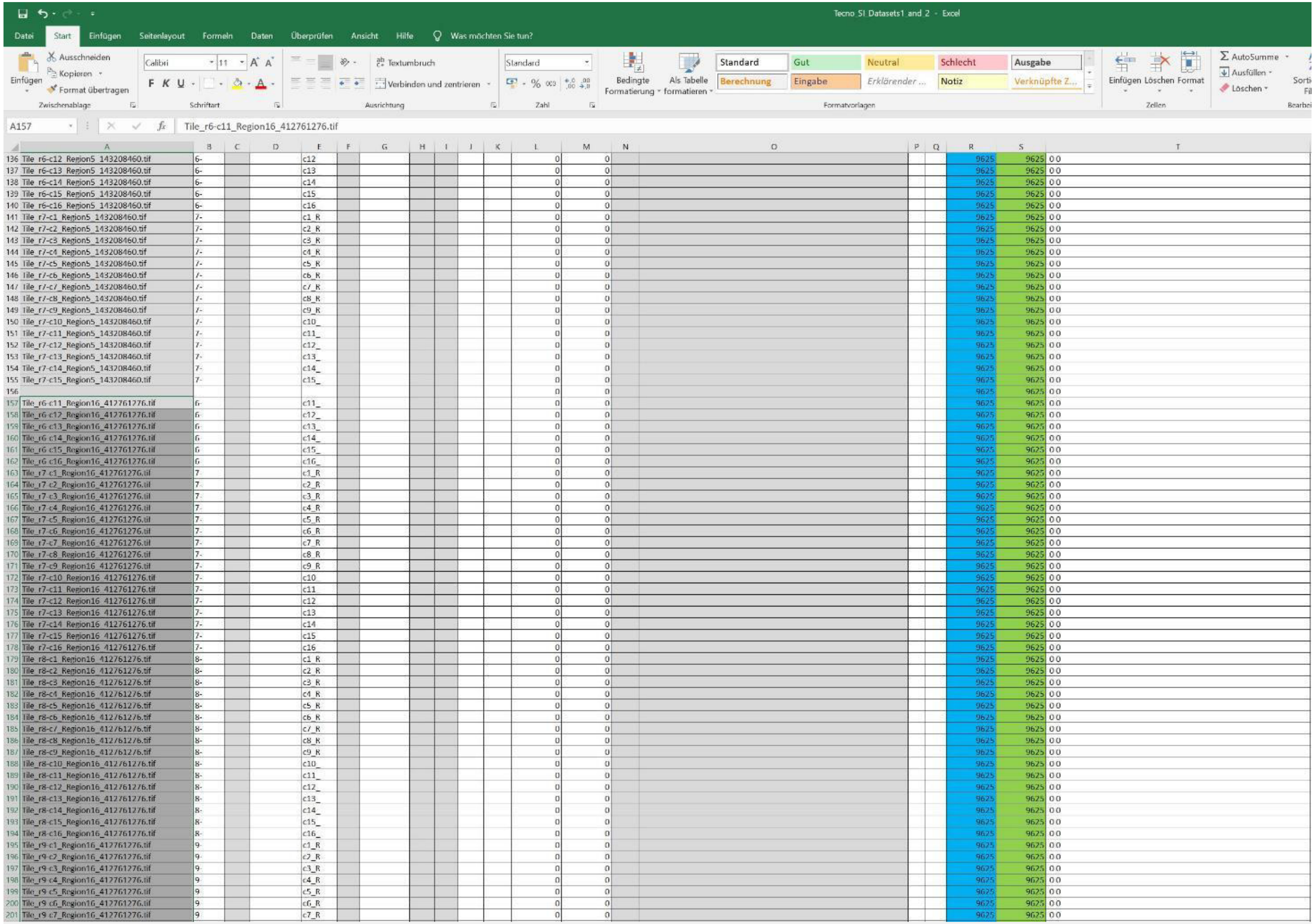


| | A | B | E | L | M | R | S | T |
|---|---|---|---|---|---|---|---|---|
| 136 | Tile_r6-c12_Region5_143208460.tif | 6- | c12 | 0 | 0 | 9625 | 9625 | 0 0 |
| 137 | Tile_r6-c13_Region5_143208460.tif | 6- | c13 | 0 | 0 | 9625 | 9625 | 0 0 |
| 138 | Tile_r6-c14_Region5_143208460.tif | 6- | c14 | 0 | 0 | 9625 | 9625 | 0 0 |
| 139 | Tile_r6-c15_Region5_143208460.tif | 6- | c15_ | 0 | 0 | 9625 | 9625 | 0 0 |
| 140 | Tile_r6-c16_Region5_143208460.tif | 6- | c16_ | 0 | 0 | 9625 | 9625 | 0 0 |
| 141 | Tile_r7-c1_Region5_143208460.tif | 7- | c1_R | 0 | 0 | 9625 | 9625 | 0 0 |
| 142 | Tile_r7-c2_Region5_143208460.tif | 7- | c2_R | 0 | 0 | 9625 | 9625 | 0 0 |
| 143 | Tile_r7-c3_Region5_143208460.tif | 7- | c3_R | 0 | 0 | 9625 | 9625 | 0 0 |
| 144 | Tile_r7-c4_Region5_143208460.tif | 7- | c4_R | 0 | 0 | 9625 | 9625 | 0 0 |
| 145 | Tile_r7-c5_Region5_143208460.tif | 7- | c5_R | 0 | 0 | 9625 | 9625 | 0 0 |
| 146 | Tile_r7-c6_Region5_143208460.tif | 7- | c6_R | 0 | 0 | 9625 | 9625 | 0 0 |
| 147 | Tile_r7-c7_Region5_143208460.tif | 7- | c7_R | 0 | 0 | 9625 | 9625 | 0 0 |
| 148 | Tile_r7-c8_Region5_143208460.tif | 7- | c8_R | 0 | 0 | 9625 | 9625 | 0 0 |
| 149 | Tile_r7-c9_Region5_143208460.tif | 7- | c9_R | 0 | 0 | 9625 | 9625 | 0 0 |
| 150 | Tile_r7-c10_Region5_143208460.tif | 7- | c10_ | 0 | 0 | 9625 | 9625 | 0 0 |
| 151 | Tile_r7-c11_Region5_143208460.tif | 7- | c11_ | 0 | 0 | 9625 | 9625 | 0 0 |
| 152 | Tile_r7-c12_Region5_143208460.tif | 7- | c12_ | 0 | 0 | 9625 | 9625 | 0 0 |
| 153 | Tile_r7-c13_Region5_143208460.tif | 7- | c13_ | 0 | 0 | 9625 | 9625 | 0 0 |
| 154 | Tile_r7-c14_Region5_143208460.tif | 7- | c14_ | 0 | 0 | 9625 | 9625 | 0 0 |
| 155 | Tile_r7-c15_Region5_143208460.tif | 7- | c15_ | 0 | 0 | 9625 | 9625 | 0 0 |
| 156 | | | | 0 | 0 | 9625 | 9625 | 0 0 |
| 157 | Tile_r6-c11_Region16_412761276.tif | 6- | c11_ | 0 | 0 | 9625 | 9625 | 0 0 |
| 158 | Tile_r6-c12_Region16_412761276.tif | 6- | c12_ | 0 | 0 | 9625 | 9625 | 0 0 |
| 159 | Tile_r6-c13_Region16_412761276.tif | 6- | c13_ | 0 | 0 | 9625 | 9625 | 0 0 |
| 160 | Tile_r6-c14_Region16_412761276.tif | 6- | c14_ | 0 | 0 | 9625 | 9625 | 0 0 |
| 161 | Tile_r6-c15_Region16_412761276.tif | 6- | c15_ | 0 | 0 | 9625 | 9625 | 0 0 |
| 162 | Tile_r6-c16_Region16_412761276.tif | 6- | c16_ | 0 | 0 | 9625 | 9625 | 0 0 |
| 163 | Tile_r7-c1_Region16_412761276.tif | 7- | c1_R | 0 | 0 | 9625 | 9625 | 0 0 |
| 164 | Tile_r7-c2_Region16_412761276.tif | 7- | c2_R | 0 | 0 | 9625 | 9625 | 0 0 |
| 165 | Tile_r7-c3_Region16_412761276.tif | 7- | c3_R | 0 | 0 | 9625 | 9625 | 0 0 |
| 166 | Tile_r7-c4_Region16_412761276.tif | 7- | c4_R | 0 | 0 | 9625 | 9625 | 0 0 |
| 167 | Tile_r7-c5_Region16_412761276.tif | 7- | c5_R | 0 | 0 | 9625 | 9625 | 0 0 |
| 168 | Tile_r7-c6_Region16_412761276.tif | 7- | c6_R | 0 | 0 | 9625 | 9625 | 0 0 |
| 169 | Tile_r7-c7_Region16_412761276.tif | 7- | c7_R | 0 | 0 | 9625 | 9625 | 0 0 |
| 170 | Tile_r7-c8_Region16_412761276.tif | 7- | c8_R | 0 | 0 | 9625 | 9625 | 0 0 |
| 171 | Tile_r7-c9_Region16_412761276.tif | 7- | c9_R | 0 | 0 | 9625 | 9625 | 0 0 |
| 172 | Tile_r7-c10_Region16_412761276.tif | 7- | c10_ | 0 | 0 | 9625 | 9625 | 0 0 |
| 173 | Tile_r7-c11_Region16_412761276.tif | 7- | c11_ | 0 | 0 | 9625 | 9625 | 0 0 |
| 174 | Tile_r7-c12_Region16_412761276.tif | 7- | c12_ | 0 | 0 | 9625 | 9625 | 0 0 |
| 175 | Tile_r7-c13_Region16_412761276.tif | 7- | c13_ | 0 | 0 | 9625 | 9625 | 0 0 |
| 176 | Tile_r7-c14_Region16_412761276.tif | 7- | c14_ | 0 | 0 | 9625 | 9625 | 0 0 |
| 177 | Tile_r7-c15_Region16_412761276.tif | 7- | c15_ | 0 | 0 | 9625 | 9625 | 0 0 |
| 178 | Tile_r7-c16_Region16_412761276.tif | 7- | c16_ | 0 | 0 | 9625 | 9625 | 0 0 |
| 179 | Tile_r8-c1_Region16_412761276.tif | 8- | c1_R | 0 | 0 | 9625 | 9625 | 0 0 |
| 180 | Tile_r8-c2_Region16_412761276.tif | 8- | c2_R | 0 | 0 | 9625 | 9625 | 0 0 |
| 181 | Tile_r8-c3_Region16_412761276.tif | 8- | c3_R | 0 | 0 | 9625 | 9625 | 0 0 |
| 182 | Tile_r8-c4_Region16_412761276.tif | 8- | c4_R | 0 | 0 | 9625 | 9625 | 0 0 |
| 183 | Tile_r8-c5_Region16_412761276.tif | 8- | c5_R | 0 | 0 | 9625 | 9625 | 0 0 |
| 184 | Tile_r8-c6_Region16_412761276.tif | 8- | c6_R | 0 | 0 | 9625 | 9625 | 0 0 |
| 185 | Tile_r8-c7_Region16_412761276.tif | 8- | c7_R | 0 | 0 | 9625 | 9625 | 0 0 |
| 186 | Tile_r8-c8_Region16_412761276.tif | 8- | c8_R | 0 | 0 | 9625 | 9625 | 0 0 |
| 187 | Tile_r8-c9_Region16_412761276.tif | 8- | c9_R | 0 | 0 | 9625 | 9625 | 0 0 |
| 188 | Tile_r8-c10_Region16_412761276.tif | 8- | c10_ | 0 | 0 | 9625 | 9625 | 0 0 |
| 189 | Tile_r8-c11_Region16_412761276.tif | 8- | c11_ | 0 | 0 | 9625 | 9625 | 0 0 |
| 190 | Tile_r8-c12_Region16_412761276.tif | 8- | c12_ | 0 | 0 | 9625 | 9625 | 0 0 |
| 191 | Tile_r8-c13_Region16_412761276.tif | 8- | c13_ | 0 | 0 | 9625 | 9625 | 0 0 |
| 192 | Tile_r8-c14_Region16_412761276.tif | 8- | c14_ | 0 | 0 | 9625 | 9625 | 0 0 |
| 193 | Tile_r8-c15_Region16_412761276.tif | 8- | c15_ | 0 | 0 | 9625 | 9625 | 0 0 |
| 194 | Tile_r8-c16_Region16_412761276.tif | 8- | c16_ | 0 | 0 | 9625 | 9625 | 0 0 |
| 195 | Tile_r9-c1_Region16_412761276.tif | 9- | c1_R | 0 | 0 | 9625 | 9625 | 0 0 |
| 196 | Tile_r9-c2_Region16_412761276.tif | 9- | c2_R | 0 | 0 | 9625 | 9625 | 0 0 |
| 197 | Tile_r9-c3_Region16_412761276.tif | 9- | c3_R | 0 | 0 | 9625 | 9625 | 0 0 |
| 198 | Tile_r9-c4_Region16_412761276.tif | 9 | c4_R | 0 | 0 | 9625 | 9625 | 0 0 |
| 199 | Tile_r9-c5_Region16_412761276.tif | 9 | c5_R | 0 | 0 | 9625 | 9625 | 0 0 |
| 200 | Tile_r9-c6_Region16_412761276.tif | 9 | c6_R | 0 | 0 | 9625 | 9625 | 0 0 |
| 201 | Tile_r9-c7_Region16_412761276.tif | 9 | c7_R | 0 | 0 | 9625 | 9625 | 0 0 |

Prepare the image tile names of dataset 2 and copy these into the Template

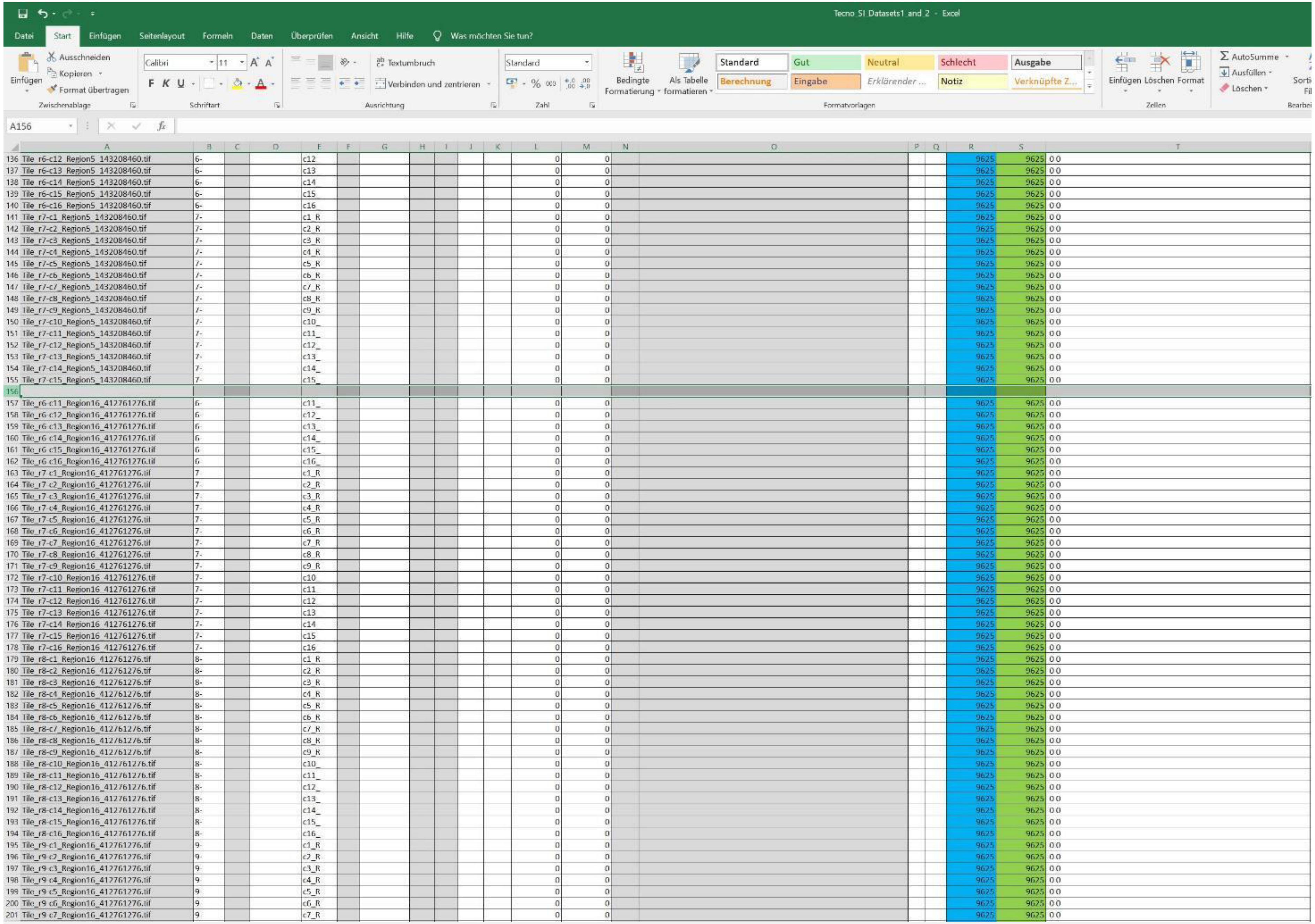


| | A | B | E | L | M | R | S | T |
|---|---|---|---|---|---|---|---|---|
| 136 | Tile_r6-c12_Region5_143208460.tif | 6- | c12 | 0 | 0 | 9625 | 9625 | 0 0 |
| 137 | Tile_r6-c13_Region5_143208460.tif | 6- | c13 | 0 | 0 | 9625 | 9625 | 0 0 |
| 138 | Tile_r6-c14_Region5_143208460.tif | 6- | c14 | 0 | 0 | 9625 | 9625 | 0 0 |
| 139 | Tile_r6-c15_Region5_143208460.tif | 6- | c15_ | 0 | 0 | 9625 | 9625 | 0 0 |
| 140 | Tile_r6-c16_Region5_143208460.tif | 6- | c16_ | 0 | 0 | 9625 | 9625 | 0 0 |
| 141 | Tile_r7-c1_Region5_143208460.tif | 7- | c1_R | 0 | 0 | 9625 | 9625 | 0 0 |
| 142 | Tile_r7-c2_Region5_143208460.tif | 7- | c2_R | 0 | 0 | 9625 | 9625 | 0 0 |
| 143 | Tile_r7-c3_Region5_143208460.tif | 7- | c3_R | 0 | 0 | 9625 | 9625 | 0 0 |
| 144 | Tile_r7-c4_Region5_143208460.tif | 7- | c4_R | 0 | 0 | 9625 | 9625 | 0 0 |
| 145 | Tile_r7-c5_Region5_143208460.tif | 7- | c5_R | 0 | 0 | 9625 | 9625 | 0 0 |
| 146 | Tile_r7-c6_Region5_143208460.tif | 7- | c6_R | 0 | 0 | 9625 | 9625 | 0 0 |
| 147 | Tile_r7-c7_Region5_143208460.tif | 7- | c7_R | 0 | 0 | 9625 | 9625 | 0 0 |
| 148 | Tile_r7-c8_Region5_143208460.tif | 7- | c8_R | 0 | 0 | 9625 | 9625 | 0 0 |
| 149 | Tile_r7-c9_Region5_143208460.tif | 7- | c9_R | 0 | 0 | 9625 | 9625 | 0 0 |
| 150 | Tile_r7-c10_Region5_143208460.tif | 7- | c10_ | 0 | 0 | 9625 | 9625 | 0 0 |
| 151 | Tile_r7-c11_Region5_143208460.tif | 7- | c11_ | 0 | 0 | 9625 | 9625 | 0 0 |
| 152 | Tile_r7-c12_Region5_143208460.tif | 7- | c12_ | 0 | 0 | 9625 | 9625 | 0 0 |
| 153 | Tile_r7-c13_Region5_143208460.tif | 7- | c13_ | 0 | 0 | 9625 | 9625 | 0 0 |
| 154 | Tile_r7-c14_Region5_143208460.tif | 7- | c14_ | 0 | 0 | 9625 | 9625 | 0 0 |
| 155 | Tile_r7-c15_Region5_143208460.tif | 7- | c15_ | 0 | 0 | 9625 | 9625 | 0 0 |
| 156 | | | | | | | | |
| 157 | Tile_r6-c11_Region16_412761276.tif | 6- | c11_ | 0 | 0 | 9625 | 9625 | 0 0 |
| 158 | Tile_r6-c12_Region16_412761276.tif | 6- | c12_ | 0 | 0 | 9625 | 9625 | 0 0 |
| 159 | Tile_r6-c13_Region16_412761276.tif | 6- | c13_ | 0 | 0 | 9625 | 9625 | 0 0 |
| 160 | Tile_r6-c14_Region16_412761276.tif | 6- | c14_ | 0 | 0 | 9625 | 9625 | 0 0 |
| 161 | Tile_r6-c15_Region16_412761276.tif | 6- | c15_ | 0 | 0 | 9625 | 9625 | 0 0 |
| 162 | Tile_r6-c16_Region16_412761276.tif | 6- | c16_ | 0 | 0 | 9625 | 9625 | 0 0 |
| 163 | Tile_r7-c1_Region16_412761276.tif | 7- | c1_R | 0 | 0 | 9625 | 9625 | 0 0 |
| 164 | Tile_r7-c2_Region16_412761276.tif | 7- | c2_R | 0 | 0 | 9625 | 9625 | 0 0 |
| 165 | Tile_r7-c3_Region16_412761276.tif | 7- | c3_R | 0 | 0 | 9625 | 9625 | 0 0 |
| 166 | Tile_r7-c4_Region16_412761276.tif | 7- | c4_R | 0 | 0 | 9625 | 9625 | 0 0 |
| 167 | Tile_r7-c5_Region16_412761276.tif | 7- | c5_R | 0 | 0 | 9625 | 9625 | 0 0 |
| 168 | Tile_r7-c6_Region16_412761276.tif | 7- | c6_R | 0 | 0 | 9625 | 9625 | 0 0 |
| 169 | Tile_r7-c7_Region16_412761276.tif | 7- | c7_R | 0 | 0 | 9625 | 9625 | 0 0 |
| 170 | Tile_r7-c8_Region16_412761276.tif | 7- | c8_R | 0 | 0 | 9625 | 9625 | 0 0 |
| 171 | Tile_r7-c9_Region16_412761276.tif | 7- | c9_R | 0 | 0 | 9625 | 9625 | 0 0 |
| 172 | Tile_r7-c10_Region16_412761276.tif | 7- | c10_ | 0 | 0 | 9625 | 9625 | 0 0 |
| 173 | Tile_r7-c11_Region16_412761276.tif | 7- | c11_ | 0 | 0 | 9625 | 9625 | 0 0 |
| 174 | Tile_r7-c12_Region16_412761276.tif | 7- | c12_ | 0 | 0 | 9625 | 9625 | 0 0 |
| 175 | Tile_r7-c13_Region16_412761276.tif | 7- | c13 | 0 | 0 | 9625 | 9625 | 0 0 |
| 176 | Tile_r7-c14_Region16_412761276.tif | 7- | c14 | 0 | 0 | 9625 | 9625 | 0 0 |
| 177 | Tile_r7-c15_Region16_412761276.tif | 7- | c15 | 0 | 0 | 9625 | 9625 | 0 0 |
| 178 | Tile_r7-c16_Region16_412761276.tif | 7- | c16_ | 0 | 0 | 9625 | 9625 | 0 0 |
| 179 | Tile_r8-c1_Region16_412761276.tif | 8- | c1_R | 0 | 0 | 9625 | 9625 | 0 0 |
| 180 | Tile_r8-c2_Region16_412761276.tif | 8- | c2_R | 0 | 0 | 9625 | 9625 | 0 0 |
| 181 | Tile_r8-c3_Region16_412761276.tif | 8- | c3_R | 0 | 0 | 9625 | 9625 | 0 0 |
| 182 | Tile_r8-c4_Region16_412761276.tif | 8- | c4_R | 0 | 0 | 9625 | 9625 | 0 0 |
| 183 | Tile_r8-c5_Region16_412761276.tif | 8- | c5_R | 0 | 0 | 9625 | 9625 | 0 0 |
| 184 | Tile_r8-c6_Region16_412761276.tif | 8- | c6_R | 0 | 0 | 9625 | 9625 | 0 0 |
| 185 | Tile_r8-c7_Region16_412761276.tif | 8- | c7_R | 0 | 0 | 9625 | 9625 | 0 0 |
| 186 | Tile_r8-c8_Region16_412761276.tif | 8- | c8_R | 0 | 0 | 9625 | 9625 | 0 0 |
| 187 | Tile_r8-c9_Region16_412761276.tif | 8- | c9_R | 0 | 0 | 9625 | 9625 | 0 0 |
| 188 | Tile_r8-c10_Region16_412761276.tif | 8- | c10_ | 0 | 0 | 9625 | 9625 | 0 0 |
| 189 | Tile_r8-c11_Region16_412761276.tif | 8- | c11_ | 0 | 0 | 9625 | 9625 | 0 0 |
| 190 | Tile_r8-c12_Region16_412761276.tif | 8- | c12_ | 0 | 0 | 9625 | 9625 | 0 0 |
| 191 | Tile_r8-c13_Region16_412761276.tif | 8- | c13_ | 0 | 0 | 9625 | 9625 | 0 0 |
| 192 | Tile_r8-c14_Region16_412761276.tif | 8- | c14_ | 0 | 0 | 9625 | 9625 | 0 0 |
| 193 | Tile_r8-c15_Region16_412761276.tif | 8- | c15_ | 0 | 0 | 9625 | 9625 | 0 0 |
| 194 | Tile_r8-c16_Region16_412761276.tif | 8- | c16_ | 0 | 0 | 9625 | 9625 | 0 0 |
| 195 | Tile_r9-c1_Region16_412761276.tif | 9- | c1_R | 0 | 0 | 9625 | 9625 | 0 0 |
| 196 | Tile_r9-c2_Region16_412761276.tif | 9- | c2_R | 0 | 0 | 9625 | 9625 | 0 0 |
| 197 | Tile_r9-c3_Region16_412761276.tif | 9- | c3_R | 0 | 0 | 9625 | 9625 | 0 0 |
| 198 | Tile_r9-c4_Region16_412761276.tif | 9- | c4_R | 0 | 0 | 9625 | 9625 | 0 0 |
| 199 | Tile_r9-c5_Region16_412761276.tif | 9- | c5_R | 0 | 0 | 9625 | 9625 | 0 0 |
| 200 | Tile_r9-c6_Region16_412761276.tif | 9 | c6_R | 0 | 0 | 9625 | 9625 | 0 0 |
| 201 | Tile_r9-c7_Region16_412761276.tif | 9 | c7_R | 0 | 0 | 9625 | 9625 | 0 0 |

Clear the space between the names of dataset 1 and dataset 2

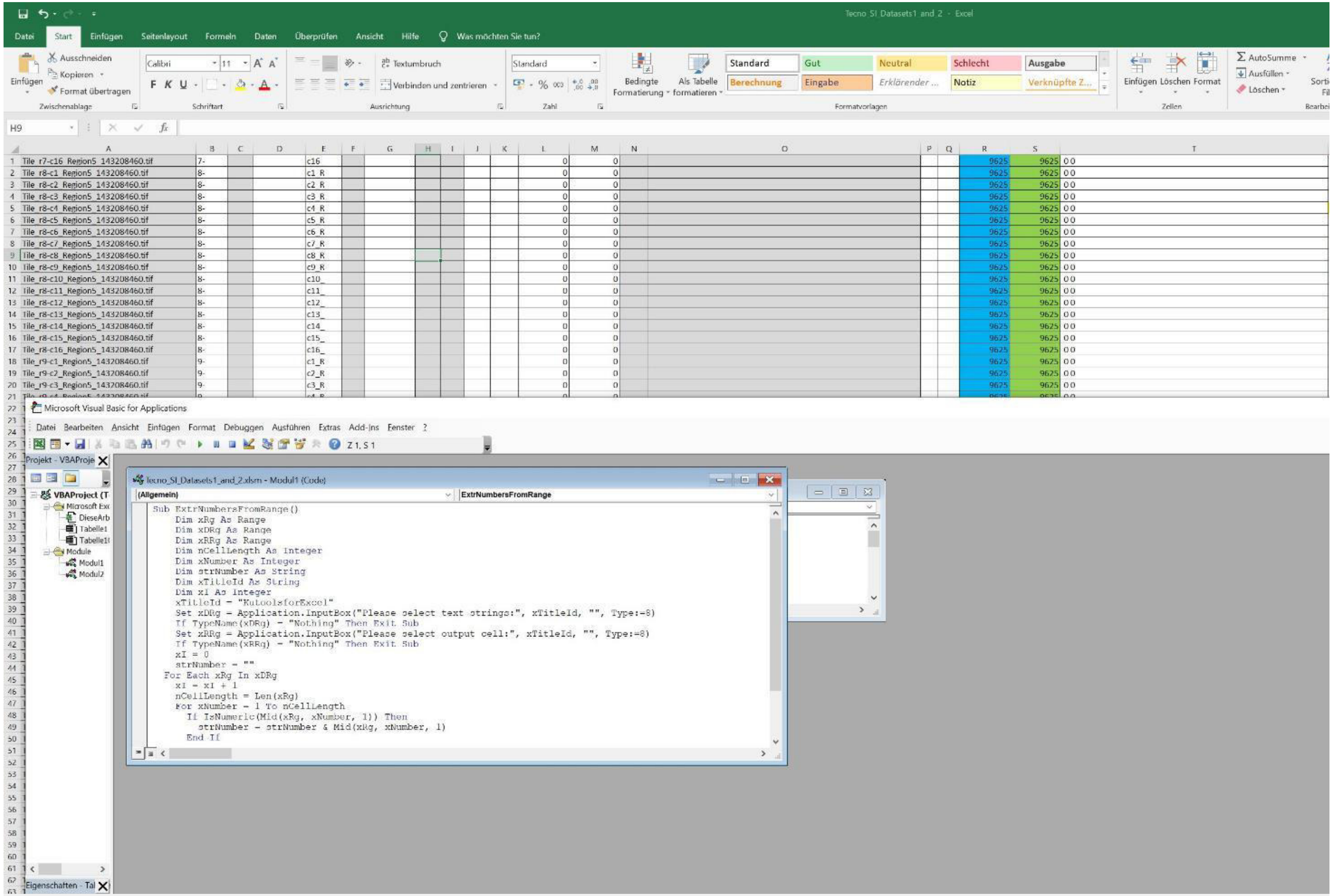


Press [Alt] and [F11] to open the macro file that is included in the Excel file
(see online methods for source)

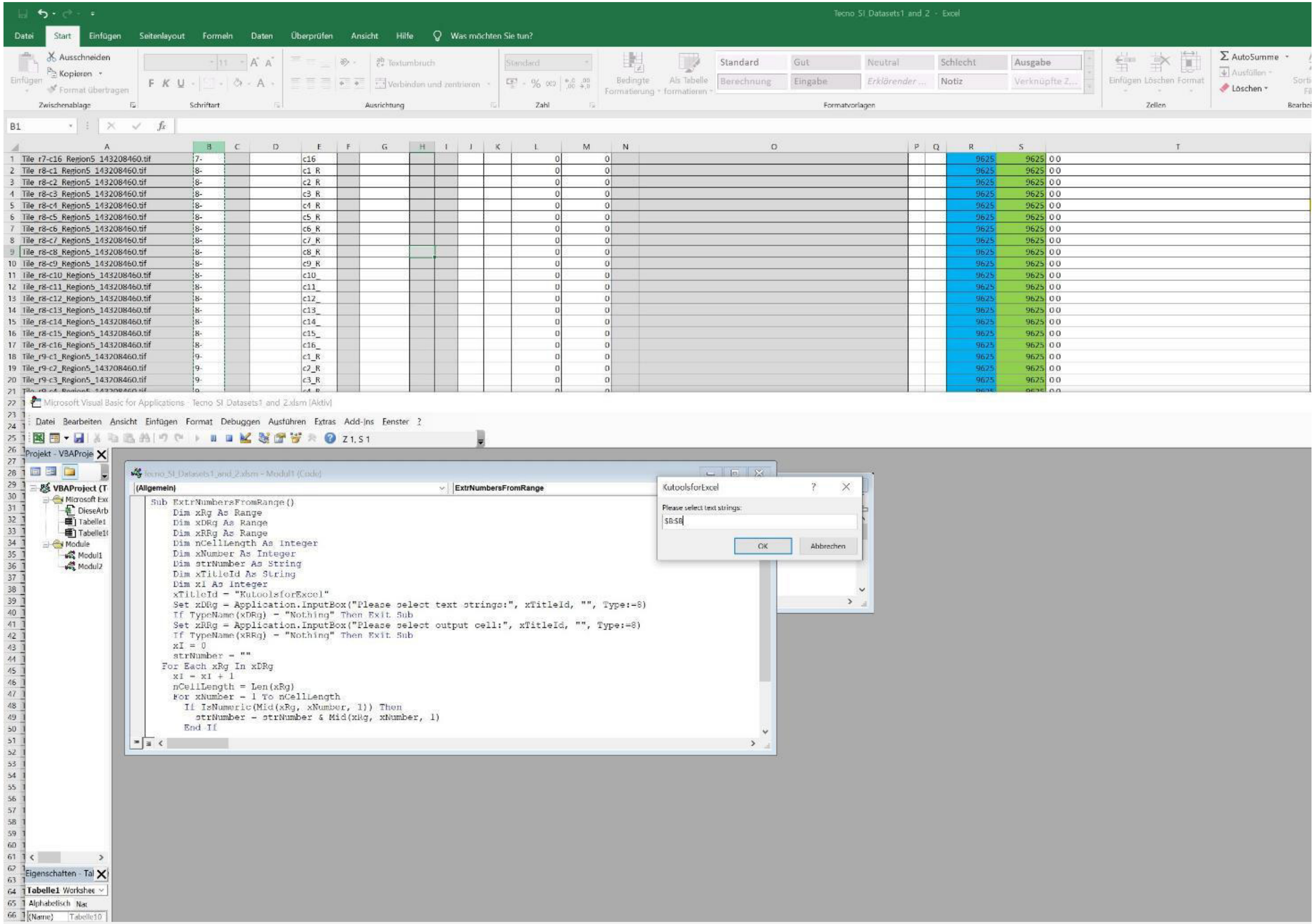


Press [F5] and select the entire column B as import

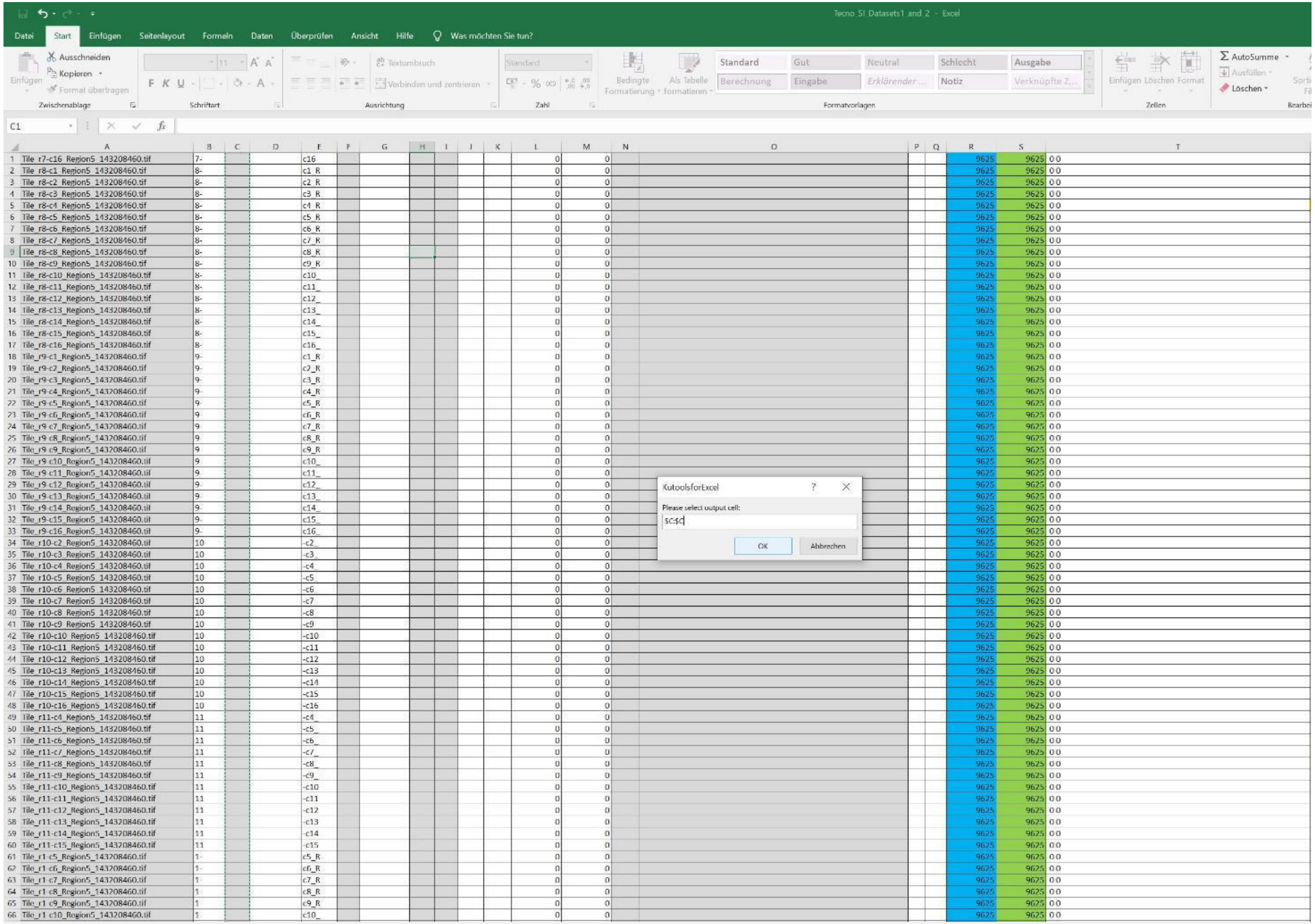


Select the entire column C as export, press „ok“

| | A | B | C | E | L | M | R | S | T |
|---|---|---|---|---|---|---|---|---|---|
| 1 | Tile_r7-c16_Region5_143208460.tif | 7- | 7 | c16 | 0 | 0 | 9625 | 9625 | 0 0 |
| 2 | Tile_r8-c1_Region5_143208460.tif | 8- | 8 | c1_R | 0 | 0 | 9625 | 9625 | 0 0 |
| 3 | Tile_r8-c2_Region5_143208460.tif | 8- | 8 | c2_R | 0 | 0 | 9625 | 9625 | 0 0 |
| 4 | Tile_r8-c3_Region5_143208460.tif | 8- | 8 | c3_R | 0 | 0 | 9625 | 9625 | 0 0 |
| 5 | Tile_r8-c4_Region5_143208460.tif | 8- | 8 | c4_R | 0 | 0 | 9625 | 9625 | 0 0 |
| 6 | Tile_r8-c5_Region5_143208460.tif | 8- | 8 | c5_R | 0 | 0 | 9625 | 9625 | 0 0 |
| 7 | Tile_r8-c6_Region5_143208460.tif | 8- | 8 | c6_R | 0 | 0 | 9625 | 9625 | 0 0 |
| 8 | Tile_r8-c7_Region5_143208460.tif | 8- | 8 | c7_R | 0 | 0 | 9625 | 9625 | 0 0 |
| 9 | Tile_r8-c8_Region5_143208460.tif | 8- | 8 | c8_R | 0 | 0 | 9625 | 9625 | 0 0 |
| 10 | Tile_r8-c9_Region5_143208460.tif | 8- | 8 | c9_R | 0 | 0 | 9625 | 9625 | 0 0 |
| 11 | Tile_r8-c10_Region5_143208460.tif | 8- | 8 | c10_ | 0 | 0 | 9625 | 9625 | 0 0 |
| 12 | Tile_r8-c11_Region5_143208460.tif | 8- | 8 | c11_ | 0 | 0 | 9625 | 9625 | 0 0 |
| 13 | Tile_r8-c12_Region5_143208460.tif | 8- | 8 | c12_ | 0 | 0 | 9625 | 9625 | 0 0 |
| 14 | Tile_r8-c13_Region5_143208460.tif | 8- | 8 | c13_ | 0 | 0 | 9625 | 9625 | 0 0 |
| 15 | Tile_r8-c14_Region5_143208460.tif | 8- | 8 | c14_ | 0 | 0 | 9625 | 9625 | 0 0 |
| 16 | Tile_r8-c15_Region5_143208460.tif | 8- | 8 | c15_ | 0 | 0 | 9625 | 9625 | 0 0 |
| 17 | Tile_r8-c16_Region5_143208460.tif | 8- | 8 | c16_ | 0 | 0 | 9625 | 9625 | 0 0 |
| 18 | Tile_r9-c1_Region5_143208460.tif | 9- | 9 | c1_R | 0 | 0 | 9625 | 9625 | 0 0 |
| 19 | Tile_r9-c2_Region5_143208460.tif | 9- | 9 | c2_R | 0 | 0 | 9625 | 9625 | 0 0 |
| 20 | Tile_r9-c3_Region5_143208460.tif | 9- | 9 | c3_R | 0 | 0 | 9625 | 9625 | 0 0 |
| 21 | Tile_r9-c4_Region5_143208460.tif | 9- | 9 | c4_R | 0 | 0 | 9625 | 9625 | 0 0 |
| 22 | Tile_r9-c5_Region5_143208460.tif | 9- | 9 | c5_R | 0 | 0 | 9625 | 9625 | 0 0 |
| 23 | Tile_r9-c6_Region5_143208460.tif | 9- | 9 | c6_R | 0 | 0 | 9625 | 9625 | 0 0 |
| 24 | Tile_r9-c7_Region5_143208460.tif | 9- | 9 | c7_R | 0 | 0 | 9625 | 9625 | 0 0 |
| 25 | Tile_r9-c8_Region5_143208460.tif | 9- | 9 | c8_R | 0 | 0 | 9625 | 9625 | 0 0 |
| 26 | Tile_r9-c9_Region5_143208460.tif | 9- | 9 | c9_R | 0 | 0 | 9625 | 9625 | 0 0 |
| 27 | Tile_r9-c10_Region5_143208460.tif | 9- | 9 | c10_ | 0 | 0 | 9625 | 9625 | 0 0 |
| 28 | Tile_r9-c11_Region5_143208460.tif | 9- | 9 | c11_ | 0 | 0 | 9625 | 9625 | 0 0 |
| 29 | Tile_r9-c12_Region5_143208460.tif | 9- | 9 | c12_ | 0 | 0 | 9625 | 9625 | 0 0 |
| 30 | Tile_r9-c13_Region5_143208460.tif | 9- | 9 | c13_ | 0 | 0 | 9625 | 9625 | 0 0 |
| 31 | Tile_r9-c14_Region5_143208460.tif | 9- | 9 | c14_ | 0 | 0 | 9625 | 9625 | 0 0 |
| 32 | Tile_r9-c15_Region5_143208460.tif | 9- | 9 | c15_ | 0 | 0 | 9625 | 9625 | 0 0 |
| 33 | Tile_r9-c16_Region5_143208460.tif | 9- | 9 | c16_ | 0 | 0 | 9625 | 9625 | 0 0 |
| 34 | Tile_r10-c2_Region5_143208460.tif | 10 | 10 | -c2_ | 0 | 0 | 9625 | 9625 | 0 0 |
| 35 | Tile_r10-c3_Region5_143208460.tif | 10 | 10 | -c3_ | 0 | 0 | 9625 | 9625 | 0 0 |
| 36 | Tile_r10-c4_Region5_143208460.tif | 10 | 10 | -c4_ | 0 | 0 | 9625 | 9625 | 0 0 |
| 37 | Tile_r10-c5_Region5_143208460.tif | 10 | 10 | -c5_ | 0 | 0 | 9625 | 9625 | 0 0 |
| 38 | Tile_r10-c6_Region5_143208460.tif | 10 | 10 | -c6_ | 0 | 0 | 9625 | 9625 | 0 0 |
| 39 | Tile_r10-c7_Region5_143208460.tif | 10 | 10 | -c7_ | 0 | 0 | 9625 | 9625 | 0 0 |
| 40 | Tile_r10-c8_Region5_143208460.tif | 10 | 10 | -c8_ | 0 | 0 | 9625 | 9625 | 0 0 |
| 41 | Tile_r10-c9_Region5_143208460.tif | 10 | 10 | -c9_ | 0 | 0 | 9625 | 9625 | 0 0 |
| 42 | Tile_r10-c10_Region5_143208460.tif | 10 | 10 | -c10 | 0 | 0 | 9625 | 9625 | 0 0 |
| 43 | Tile_r10-c11_Region5_143208460.tif | 10 | 10 | -c11 | 0 | 0 | 9625 | 9625 | 0 0 |
| 44 | Tile_r10-c12_Region5_143208460.tif | 10 | 10 | -c12 | 0 | 0 | 9625 | 9625 | 0 0 |
| 45 | Tile_r10-c13_Region5_143208460.tif | 10 | 10 | -c13 | 0 | 0 | 9625 | 9625 | 0 0 |
| 46 | Tile_r10-c14_Region5_143208460.tif | 10 | 10 | -c14 | 0 | 0 | 9625 | 9625 | 0 0 |
| 47 | Tile_r10-c15_Region5_143208460.tif | 10 | 10 | -c15 | 0 | 0 | 9625 | 9625 | 0 0 |
| 48 | Tile_r10-c16_Region5_143208460.tif | 10 | 10 | -c16 | 0 | 0 | 9625 | 9625 | 0 0 |
| 49 | Tile_r11-c4_Region5_143208460.tif | 11 | 11 | -c4_ | 0 | 0 | 9625 | 9625 | 0 0 |
| 50 | Tile_r11-c5_Region5_143208460.tif | 11 | 11 | -c5_ | 0 | 0 | 9625 | 9625 | 0 0 |
| 51 | Tile_r11-c6_Region5_143208460.tif | 11 | 11 | -c6_ | 0 | 0 | 9625 | 9625 | 0 0 |
| 52 | Tile_r11-c7_Region5_143208460.tif | 11 | 11 | -c7_ | 0 | 0 | 9625 | 9625 | 0 0 |
| 53 | Tile_r11-c8_Region5_143208460.tif | 11 | 11 | -c8_ | 0 | 0 | 9625 | 9625 | 0 0 |
| 54 | Tile_r11-c9_Region5_143208460.tif | 11 | 11 | -c9_ | 0 | 0 | 9625 | 9625 | 0 0 |
| 55 | Tile_r11-c10_Region5_143208460.tif | 11 | 11 | -c10 | 0 | 0 | 9625 | 9625 | 0 0 |
| 56 | Tile_r11-c11_Region5_143208460.tif | 11 | 11 | -c11 | 0 | 0 | 9625 | 9625 | 0 0 |
| 57 | Tile_r11-c12_Region5_143208460.tif | 11 | 11 | -c12 | 0 | 0 | 9625 | 9625 | 0 0 |
| 58 | Tile_r11-c13_Region5_143208460.tif | 11 | 11 | -c13 | 0 | 0 | 9625 | 9625 | 0 0 |
| 59 | Tile_r11-c14_Region5_143208460.tif | 11 | 11 | -c14 | 0 | 0 | 9625 | 9625 | 0 0 |
| 60 | Tile_r11-c15_Region5_143208460.tif | 11 | 11 | -c15 | 0 | 0 | 9625 | 9625 | 0 0 |
| 61 | Tile_r1-c5_Region5_143208460.tif | 1- | 1 | c5_R | 0 | 0 | 9625 | 9625 | 0 0 |
| 62 | Tile_r1-c6_Region5_143208460.tif | 1- | 1 | c6_R | 0 | 0 | 9625 | 9625 | 0 0 |
| 63 | Tile_r1-c7_Region5_143208460.tif | 1- | 1 | c7_R | 0 | 0 | 9625 | 9625 | 0 0 |
| 64 | Tile_r1-c8_Region5_143208460.tif | 1- | 1 | c8_R | 0 | 0 | 9625 | 9625 | 0 0 |
| 65 | Tile_r1-c9_Region5_143208460.tif | 1- | 1 | c9_R | 0 | 0 | 9625 | 9625 | 0 0 |
| 66 | Tile_r1-c10_Region5_143208460.tif | 1- | 1 | c10_ | 0 | 0 | 9625 | 9625 | 0 0 |

The numbers of column B are extracted and placed in column C

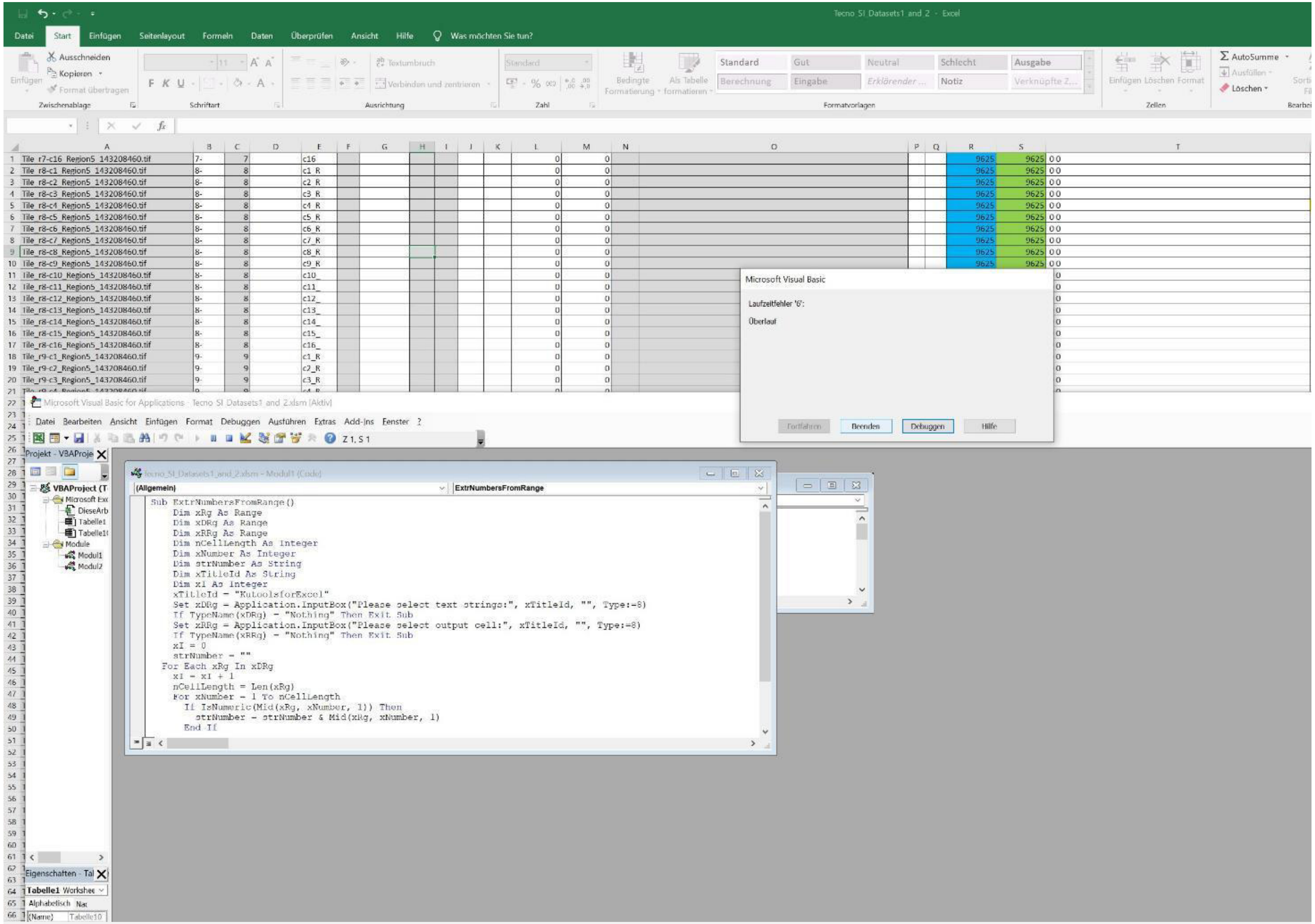


An error might occur, but also then, the extraction usually worked

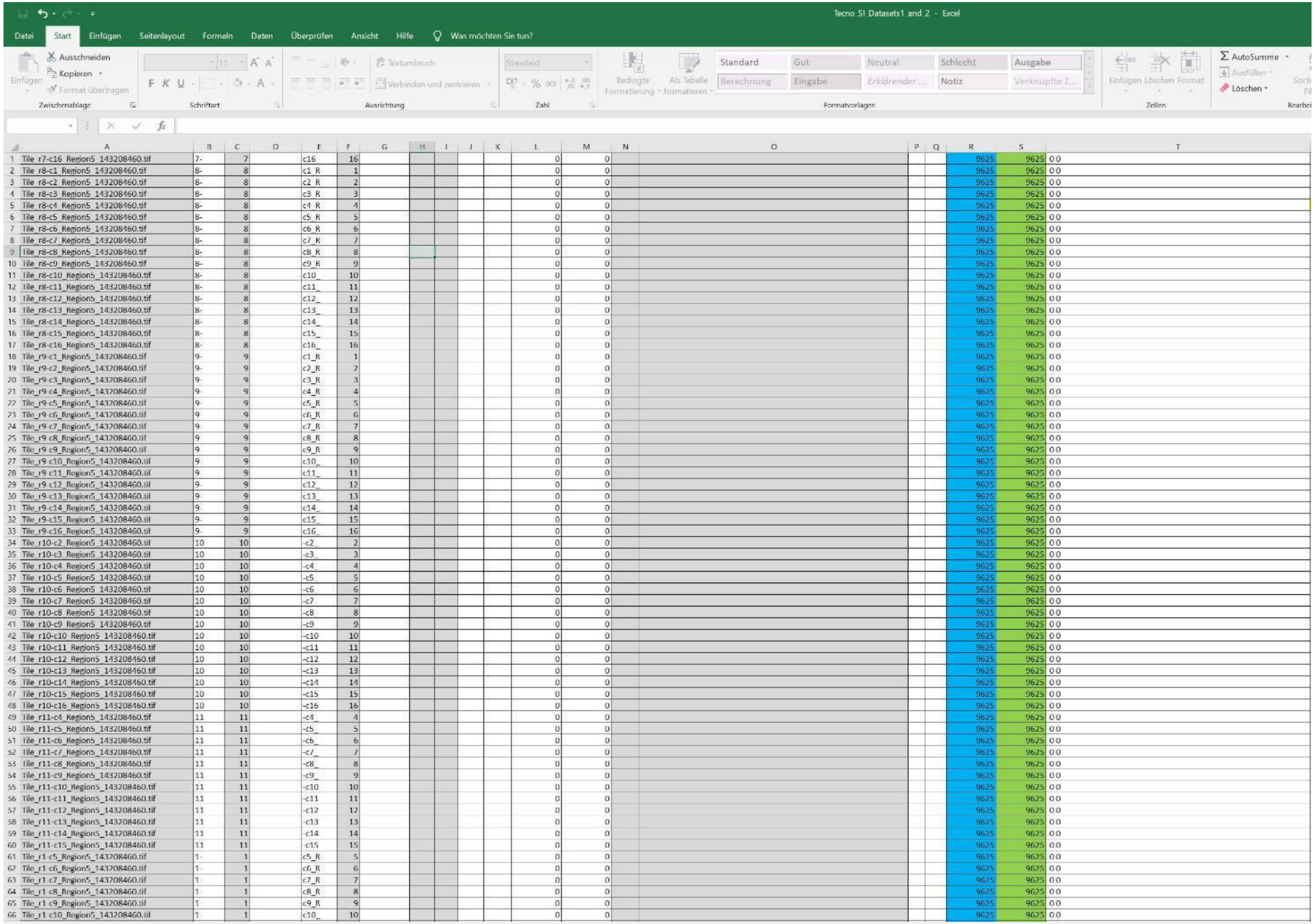


| | A | B | C | E | F | L | M | R | S | T |
|---|---|---|---|---|---|---|---|---|---|---|
| 1 | Tile_r7-c16_Region5_143208460.tif | 7- | 7 | c16 | 16 | 0 | 0 | 9625 | 9625 | 0 0 |
| 2 | Tile_r8-c1_Region5_143208460.tif | 8- | 8 | c1_R | 1 | 0 | 0 | 9625 | 9625 | 0 0 |
| 3 | Tile_r8-c2_Region5_143208460.tif | 8- | 8 | c2_R | 2 | 0 | 0 | 9625 | 9625 | 0 0 |
| 4 | Tile_r8-c3_Region5_143208460.tif | 8- | 8 | c3_R | 3 | 0 | 0 | 9625 | 9625 | 0 0 |
| 5 | Tile_r8-c4_Region5_143208460.tif | 8- | 8 | c4_R | 4 | 0 | 0 | 9625 | 9625 | 0 0 |
| 6 | Tile_r8-c5_Region5_143208460.tif | 8- | 8 | c5_R | 5 | 0 | 0 | 9625 | 9625 | 0 0 |
| 7 | Tile_r8-c6_Region5_143208460.tif | 8- | 8 | c6_R | 6 | 0 | 0 | 9625 | 9625 | 0 0 |
| 8 | Tile_r8-c7_Region5_143208460.tif | 8- | 8 | c7_R | 7 | 0 | 0 | 9625 | 9625 | 0 0 |
| 9 | Tile_r8-c8_Region5_143208460.tif | 8- | 8 | c8_R | 8 | 0 | 0 | 9625 | 9625 | 0 0 |
| 10 | Tile_r8-c9_Region5_143208460.tif | 8- | 8 | c9_R | 9 | 0 | 0 | 9625 | 9625 | 0 0 |
| 11 | Tile_r8-c10_Region5_143208460.tif | 8- | 8 | c10_ | 10 | 0 | 0 | 9625 | 9625 | 0 0 |
| 12 | Tile_r8-c11_Region5_143208460.tif | 8- | 8 | c11_ | 11 | 0 | 0 | 9625 | 9625 | 0 0 |
| 13 | Tile_r8-c12_Region5_143208460.tif | 8- | 8 | c12_ | 12 | 0 | 0 | 9625 | 9625 | 0 0 |
| 14 | Tile_r8-c13_Region5_143208460.tif | 8- | 8 | c13_ | 13 | 0 | 0 | 9625 | 9625 | 0 0 |
| 15 | Tile_r8-c14_Region5_143208460.tif | 8- | 8 | c14_ | 14 | 0 | 0 | 9625 | 9625 | 0 0 |
| 16 | Tile_r8-c15_Region5_143208460.tif | 8- | 8 | c15_ | 15 | 0 | 0 | 9625 | 9625 | 0 0 |
| 17 | Tile_r8-c16_Region5_143208460.tif | 8- | 8 | c16_ | 16 | 0 | 0 | 9625 | 9625 | 0 0 |
| 18 | Tile_r9-c1_Region5_143208460.tif | 9- | 9 | c1_R | 1 | 0 | 0 | 9625 | 9625 | 0 0 |
| 19 | Tile_r9-c2_Region5_143208460.tif | 9- | 9 | c2_R | 2 | 0 | 0 | 9625 | 9625 | 0 0 |
| 20 | Tile_r9-c3_Region5_143208460.tif | 9- | 9 | c3_R | 3 | 0 | 0 | 9625 | 9625 | 0 0 |
| 21 | Tile_r9-c4_Region5_143208460.tif | 9- | 9 | c4_R | 4 | 0 | 0 | 9625 | 9625 | 0 0 |
| 22 | Tile_r9-c5_Region5_143208460.tif | 9- | 9 | c5_R | 5 | 0 | 0 | 9625 | 9625 | 0 0 |
| 23 | Tile_r9-c6_Region5_143208460.tif | 9- | 9 | c6_R | 6 | 0 | 0 | 9625 | 9625 | 0 0 |
| 24 | Tile_r9-c7_Region5_143208460.tif | 9- | 9 | c7_R | 7 | 0 | 0 | 9625 | 9625 | 0 0 |
| 25 | Tile_r9-c8_Region5_143208460.tif | 9- | 9 | c8_R | 8 | 0 | 0 | 9625 | 9625 | 0 0 |
| 26 | Tile_r9-c9_Region5_143208460.tif | 9- | 9 | c9_R | 9 | 0 | 0 | 9625 | 9625 | 0 0 |
| 27 | Tile_r9-c10_Region5_143208460.tif | 9- | 9 | c10_ | 10 | 0 | 0 | 9625 | 9625 | 0 0 |
| 28 | Tile_r9-c11_Region5_143208460.tif | 9- | 9 | c11_ | 11 | 0 | 0 | 9625 | 9625 | 0 0 |
| 29 | Tile_r9-c12_Region5_143208460.tif | 9- | 9 | c12_ | 12 | 0 | 0 | 9625 | 9625 | 0 0 |
| 30 | Tile_r9-c13_Region5_143208460.tif | 9- | 9 | c13_ | 13 | 0 | 0 | 9625 | 9625 | 0 0 |
| 31 | Tile_r9-c14_Region5_143208460.tif | 9- | 9 | c14_ | 14 | 0 | 0 | 9625 | 9625 | 0 0 |
| 32 | Tile_r9-c15_Region5_143208460.tif | 9- | 9 | c15_ | 15 | 0 | 0 | 9625 | 9625 | 0 0 |
| 33 | Tile_r9-c16_Region5_143208460.tif | 9- | 9 | c16_ | 16 | 0 | 0 | 9625 | 9625 | 0 0 |
| 34 | Tile_r10-c2_Region5_143208460.tif | 10 | 10 | -c2_ | 2 | 0 | 0 | 9625 | 9625 | 0 0 |
| 35 | Tile_r10-c3_Region5_143208460.tif | 10 | 10 | -c3_ | 3 | 0 | 0 | 9625 | 9625 | 0 0 |
| 36 | Tile_r10-c4_Region5_143208460.tif | 10 | 10 | -c4_ | 4 | 0 | 0 | 9625 | 9625 | 0 0 |
| 37 | Tile_r10-c5_Region5_143208460.tif | 10 | 10 | -c5_ | 5 | 0 | 0 | 9625 | 9625 | 0 0 |
| 38 | Tile_r10-c6_Region5_143208460.tif | 10 | 10 | -c6_ | 6 | 0 | 0 | 9625 | 9625 | 0 0 |
| 39 | Tile_r10-c7_Region5_143208460.tif | 10 | 10 | -c7 | 7 | 0 | 0 | 9625 | 9625 | 0 0 |
| 40 | Tile_r10-c8_Region5_143208460.tif | 10 | 10 | -c8 | 8 | 0 | 0 | 9625 | 9625 | 0 0 |
| 41 | Tile_r10-c9_Region5_143208460.tif | 10 | 10 | -c9 | 9 | 0 | 0 | 9625 | 9625 | 0 0 |
| 42 | Tile_r10-c10_Region5_143208460.tif | 10 | 10 | -c10 | 10 | 0 | 0 | 9625 | 9625 | 0 0 |
| 43 | Tile_r10-c11_Region5_143208460.tif | 10 | 10 | -c11 | 11 | 0 | 0 | 9625 | 9625 | 0 0 |
| 44 | Tile_r10-c12_Region5_143208460.tif | 10 | 10 | -c12 | 12 | 0 | 0 | 9625 | 9625 | 0 0 |
| 45 | Tile_r10-c13_Region5_143208460.tif | 10 | 10 | -c13 | 13 | 0 | 0 | 9625 | 9625 | 0 0 |
| 46 | Tile_r10-c14_Region5_143208460.tif | 10 | 10 | -c14 | 14 | 0 | 0 | 9625 | 9625 | 0 0 |
| 47 | Tile_r10-c15_Region5_143208460.tif | 10 | 10 | -c15 | 15 | 0 | 0 | 9625 | 9625 | 0 0 |
| 48 | Tile_r10-c16_Region5_143208460.tif | 10 | 10 | -c16 | 16 | 0 | 0 | 9625 | 9625 | 0 0 |
| 49 | Tile_r11-c4_Region5_143208460.tif | 11 | 11 | -c4_ | 4 | 0 | 0 | 9625 | 9625 | 0 0 |
| 50 | Tile_r11-c5_Region5_143208460.tif | 11 | 11 | -c5_ | 5 | 0 | 0 | 9625 | 9625 | 0 0 |
| 51 | Tile_r11-c6_Region5_143208460.tif | 11 | 11 | -c6_ | 6 | 0 | 0 | 9625 | 9625 | 0 0 |
| 52 | Tile_r11-c7_Region5_143208460.tif | 11 | 11 | -c7_ | 7 | 0 | 0 | 9625 | 9625 | 0 0 |
| 53 | Tile_r11-c8_Region5_143208460.tif | 11 | 11 | -c8_ | 8 | 0 | 0 | 9625 | 9625 | 0 0 |
| 54 | Tile_r11-c9_Region5_143208460.tif | 11 | 11 | -c9_ | 9 | 0 | 0 | 9625 | 9625 | 0 0 |
| 55 | Tile_r11-c10_Region5_143208460.tif | 11 | 11 | -c10 | 10 | 0 | 0 | 9625 | 9625 | 0 0 |
| 56 | Tile_r11-c11_Region5_143208460.tif | 11 | 11 | -c11 | 11 | 0 | 0 | 9625 | 9625 | 0 0 |
| 57 | Tile_r11-c12_Region5_143208460.tif | 11 | 11 | -c12 | 12 | 0 | 0 | 9625 | 9625 | 0 0 |
| 58 | Tile_r11-c13_Region5_143208460.tif | 11 | 11 | -c13 | 13 | 0 | 0 | 9625 | 9625 | 0 0 |
| 59 | Tile_r11-c14_Region5_143208460.tif | 11 | 11 | -c14 | 14 | 0 | 0 | 9625 | 9625 | 0 0 |
| 60 | Tile_r11-c15_Region5_143208460.tif | 11 | 11 | -c15 | 15 | 0 | 0 | 9625 | 9625 | 0 0 |
| 61 | Tile_r1-c5_Region5_143208460.tif | 1- | 1 | c5_R | 5 | 0 | 0 | 9625 | 9625 | 0 0 |
| 62 | Tile_r1-c6_Region5_143208460.tif | 1- | 1 | c6_R | 6 | 0 | 0 | 9625 | 9625 | 0 0 |
| 63 | Tile_r1-c7_Region5_143208460.tif | 1- | 1 | c7_R | 7 | 0 | 0 | 9625 | 9625 | 0 0 |
| 64 | Tile_r1-c8_Region5_143208460.tif | 1- | 1 | c8_R | 8 | 0 | 0 | 9625 | 9625 | 0 0 |
| 65 | Tile_r1-c9_Region5_143208460.tif | 1- | 1 | c9_R | 9 | 0 | 0 | 9625 | 9625 | 0 0 |
| 66 | Tile_r1-c10_Region5_143208460.tif | 1- | 1 | c10_ | 10 | 0 | 0 | 9625 | 9625 | 0 0 |

Extract the numbers in column E to column F the same way

Tecno SI Datasets1 and 2 - Excel

| | A | B | C | E | F | H | I | L | M | R | S | T |
|---|---|---|---|---|---|---|---|---|---|---|---|---|
| 1 | Tile_r7-c16_Region5_143208460.tif | 7- | 7 | c16 | 16 | 7 | 16 | 154000 | 67375 | 9625 | 9625 | 154000 67375 |
| 2 | Tile_r8-c1_Region5_143208460.tif | 8- | 8 | c1_R | 1 | 8 | 1 | 9625 | 77000 | 9625 | 9625 | 9625 77000 |
| 3 | Tile_r8-c2_Region5_143208460.tif | 8- | 8 | c2_R | 2 | 8 | 2 | 19250 | 77000 | 9625 | 9625 | 19250 77000 |
| 4 | Tile_r8-c3_Region5_143208460.tif | 8- | 8 | c3_R | 3 | 8 | 3 | 28875 | 77000 | 9625 | 9625 | 28875 77000 |
| 5 | Tile_r8-c4_Region5_143208460.tif | 8- | 8 | c4_R | 4 | 8 | 4 | 38500 | 77000 | 9625 | 9625 | 38500 77000 |
| 6 | Tile_r8-c5_Region5_143208460.tif | 8- | 8 | c5_R | 5 | 8 | 5 | 48125 | 77000 | 9625 | 9625 | 48125 77000 |
| 7 | Tile_r8-c6_Region5_143208460.tif | 8- | 8 | c6_R | 6 | 8 | 6 | 57750 | 77000 | 9625 | 9625 | 57750 77000 |
| 8 | Tile_r8-c7_Region5_143208460.tif | 8- | 8 | c7_R | 7 | 8 | 7 | 67375 | 77000 | 9625 | 9625 | 67375 77000 |
| 9 | Tile_r8-c8_Region5_143208460.tif | 8- | 8 | c8_R | 8 | 8 | 8 | 77000 | 77000 | 9625 | 9625 | 77000 77000 |
| 10 | Tile_r8-c9_Region5_143208460.tif | 8- | 8 | c9_R | 9 | 8 | 9 | 86625 | 77000 | 9625 | 9625 | 86625 77000 |
| 11 | Tile_r8-c10_Region5_143208460.tif | 8- | 8 | c10_ | 10 | 8 | 10 | 96250 | 77000 | 9625 | 9625 | 96250 77000 |
| 12 | Tile_r8-c11_Region5_143208460.tif | 8- | 8 | c11_ | 11 | 8 | 11 | 105875 | 77000 | 9625 | 9625 | 105875 77000 |
| 13 | Tile_r8-c12_Region5_143208460.tif | 8- | 8 | c12_ | 12 | 8 | 12 | 115500 | 77000 | 9625 | 9625 | 115500 77000 |
| 14 | Tile_r8-c13_Region5_143208460.tif | 8- | 8 | c13_ | 13 | 8 | 13 | 125125 | 77000 | 9625 | 9625 | 125125 77000 |
| 15 | Tile_r8-c14_Region5_143208460.tif | 8- | 8 | c14_ | 14 | 8 | 14 | 134750 | 77000 | 9625 | 9625 | 134750 77000 |
| 16 | Tile_r8-c15_Region5_143208460.tif | 8- | 8 | c15_ | 15 | 8 | 15 | 144375 | 77000 | 9625 | 9625 | 144375 77000 |
| 17 | Tile_r8-c16_Region5_143208460.tif | 8- | 8 | c16_ | 16 | 8 | 16 | 154000 | 77000 | 9625 | 9625 | 154000 77000 |
| 18 | Tile_r9-c1_Region5_143208460.tif | 9- | 9 | c1_R | 1 | 9 | 1 | 9625 | 86625 | 9625 | 9625 | 9625 86625 |
| 19 | Tile_r9-c2_Region5_143208460.tif | 9- | 9 | c2_R | 2 | 9 | 2 | 19250 | 86625 | 9625 | 9625 | 19250 86625 |
| 20 | Tile_r9-c3_Region5_143208460.tif | 9- | 9 | c3_R | 3 | 9 | 3 | 28875 | 86625 | 9625 | 9625 | 28875 86625 |
| 21 | Tile_r9-c4_Region5_143208460.tif | 9- | 9 | c4_R | 4 | 9 | 4 | 38500 | 86625 | 9625 | 9625 | 38500 86625 |
| 22 | Tile_r9-c5_Region5_143208460.tif | 9- | 9 | c5_R | 5 | 9 | 5 | 48125 | 86625 | 9625 | 9625 | 48125 86625 |
| 23 | Tile_r9-c6_Region5_143208460.tif | 9 | 9 | c6_R | 6 | 9 | 6 | 57750 | 86625 | 9625 | 9625 | 57750 86625 |
| 24 | Tile_r9-c7_Region5_143208460.tif | 9 | 9 | c7_R | 7 | 9 | 7 | 67375 | 86625 | 9625 | 9625 | 67375 86625 |
| 25 | Tile_r9-c8_Region5_143208460.tif | 9 | 9 | c8_R | 8 | 9 | 8 | 77000 | 86625 | 9625 | 9625 | 77000 86625 |
| 26 | Tile_r9-c9_Region5_143208460.tif | 9 | 9 | c9_R | 9 | 9 | 9 | 86625 | 86625 | 9625 | 9625 | 86625 86625 |
| 27 | Tile_r9-c10_Region5_143208460.tif | 9 | 9 | c10_ | 10 | 9 | 10 | 96250 | 86625 | 9625 | 9625 | 96250 86625 |
| 28 | Tile_r9-c11_Region5_143208460.tif | 9 | 9 | c11_ | 11 | 9 | 11 | 105875 | 86625 | 9625 | 9625 | 105875 86625 |
| 29 | Tile_r9-c12_Region5_143208460.tif | 9- | 9 | c12_ | 12 | 9 | 12 | 115500 | 86625 | 9625 | 9625 | 115500 86625 |
| 30 | Tile_r9-c13_Region5_143208460.tif | 9- | 9 | c13_ | 13 | 9 | 13 | 125125 | 86625 | 9625 | 9625 | 125125 86625 |
| 31 | Tile_r9-c14_Region5_143208460.tif | 9- | 9 | c14_ | 14 | 9 | 14 | 134750 | 86625 | 9625 | 9625 | 134750 86625 |
| 32 | Tile_r9-c15_Region5_143208460.tif | 9- | 9 | c15_ | 15 | 9 | 15 | 144375 | 86625 | 9625 | 9625 | 144375 86625 |
| 33 | Tile_r9-c16_Region5_143208460.tif | 9- | 9 | c16_ | 16 | 9 | 16 | 154000 | 86625 | 9625 | 9625 | 154000 86625 |
| 34 | Tile_r10-c2_Region5_143208460.tif | 10 | 10 | -c2_ | 2 | 10 | 2 | 19250 | 96250 | 9625 | 9625 | 19250 96250 |
| 35 | Tile_r10-c3_Region5_143208460.tif | 10 | 10 | -c3_ | 3 | 10 | 3 | 28875 | 96250 | 9625 | 9625 | 28875 96250 |
| 36 | Tile_r10-c4_Region5_143208460.tif | 10 | 10 | -c4_ | 4 | 10 | 4 | 38500 | 96250 | 9625 | 9625 | 38500 96250 |
| 37 | Tile_r10-c5_Region5_143208460.tif | 10 | 10 | -c5_ | 5 | 10 | 5 | 48125 | 96250 | 9625 | 9625 | 48125 96250 |
| 38 | Tile_r10-c6_Region5_143208460.tif | 10 | 10 | -c6 | 6 | 10 | 6 | 57750 | 96250 | 9625 | 9625 | 57750 96250 |
| 39 | Tile_r10-c7_Region5_143208460.tif | 10 | 10 | -c7 | 7 | 10 | 7 | 67375 | 96250 | 9625 | 9625 | 67375 96250 |
| 40 | Tile_r10-c8_Region5_143208460.tif | 10 | 10 | -c8 | 8 | 10 | 8 | 77000 | 96250 | 9625 | 9625 | 77000 96250 |
| 41 | Tile_r10-c9_Region5_143208460.tif | 10 | 10 | -c9 | 9 | 10 | 9 | 86625 | 96250 | 9625 | 9625 | 86625 96250 |
| 42 | Tile_r10-c10_Region5_143208460.tif | 10 | 10 | -c10 | 10 | 10 | 10 | 96250 | 96250 | 9625 | 9625 | 96250 96250 |
| 43 | Tile_r10-c11_Region5_143208460.tif | 10 | 10 | -c11 | 11 | 10 | 11 | 105875 | 96250 | 9625 | 9625 | 105875 96250 |
| 44 | Tile_r10-c12_Region5_143208460.tif | 10 | 10 | -c12 | 12 | 10 | 12 | 115500 | 96250 | 9625 | 9625 | 115500 96250 |
| 45 | Tile_r10-c13_Region5_143208460.tif | 10 | 10 | -c13 | 13 | 10 | 13 | 125125 | 96250 | 9625 | 9625 | 125125 96250 |
| 46 | Tile_r10-c14_Region5_143208460.tif | 10 | 10 | -c14 | 14 | 10 | 14 | 134750 | 96250 | 9625 | 9625 | 134750 96250 |
| 47 | Tile_r10-c15_Region5_143208460.tif | 10 | 10 | -c15 | 15 | 10 | 15 | 144375 | 96250 | 9625 | 9625 | 144375 96250 |
| 48 | Tile_r10-c16_Region5_143208460.tif | 10 | 10 | -c16 | 16 | 10 | 16 | 154000 | 96250 | 9625 | 9625 | 154000 96250 |
| 49 | Tile_r11-c4_Region5_143208460.tif | 11 | 11 | -c4_ | 4 | 11 | 4 | 38500 | 105875 | 9625 | 9625 | 38500 105875 |
| 50 | Tile_r11-c5_Region5_143208460.tif | 11 | 11 | -c5_ | 5 | 11 | 5 | 48125 | 105875 | 9625 | 9625 | 48125 105875 |
| 51 | Tile_r11-c6_Region5_143208460.tif | 11 | 11 | -c6_ | 6 | 11 | 6 | 57750 | 105875 | 9625 | 9625 | 57750 105875 |
| 52 | Tile_r11-c7_Region5_143208460.tif | 11 | 11 | -c7_ | 7 | 11 | 7 | 67375 | 105875 | 9625 | 9625 | 67375 105875 |
| 53 | Tile_r11-c8_Region5_143208460.tif | 11 | 11 | -c8_ | 8 | 11 | 8 | 77000 | 105875 | 9625 | 9625 | 77000 105875 |
| 54 | Tile_r11-c9_Region5_143208460.tif | 11 | 11 | -c9_ | 9 | 11 | 9 | 86625 | 105875 | 9625 | 9625 | 86625 105875 |
| 55 | Tile_r11-c10_Region5_143208460.tif | 11 | 11 | -c10 | 10 | 11 | 10 | 96250 | 105875 | 9625 | 9625 | 96250 105875 |
| 56 | Tile_r11-c11_Region5_143208460.tif | 11 | 11 | -c11 | 11 | 11 | 11 | 105875 | 105875 | 9625 | 9625 | 105875 105875 |
| 57 | Tile_r11-c12_Region5_143208460.tif | 11 | 11 | -c12 | 12 | 11 | 12 | 115500 | 105875 | 9625 | 9625 | 115500 105875 |
| 58 | Tile_r11-c13_Region5_143208460.tif | 11 | 11 | -c13 | 13 | 11 | 13 | 125125 | 105875 | 9625 | 9625 | 125125 105875 |
| 59 | Tile_r11-c14_Region5_143208460.tif | 11 | 11 | -c14 | 14 | 11 | 14 | 134750 | 105875 | 9625 | 9625 | 134750 105875 |
| 60 | Tile_r11-c15_Region5_143208460.tif | 11 | 11 | -c15 | 15 | 11 | 15 | 144375 | 105875 | 9625 | 9625 | 144375 105875 |
| 61 | Tile_r1-c5_Region5_143208460.tif | 1- | 1 | c5_R | 5 | 1 | 5 | 48125 | 9625 | 9625 | 9625 | 48125 9625 |
| 62 | Tile_r1-c6_Region5_143208460.tif | 1- | 1 | c6_R | 6 | 1 | 6 | 57750 | 9625 | 9625 | 9625 | 57750 9625 |
| 63 | Tile_r1-c7_Region5_143208460.tif | 1- | 1 | c7_R | 7 | 1 | 7 | 67375 | 9625 | 9625 | 9625 | 67375 9625 |
| 64 | Tile_r1-c8_Region5_143208460.tif | 1- | 1 | c8_R | 8 | 1 | 8 | 77000 | 9625 | 9625 | 9625 | 77000 9625 |
| 65 | Tile_r1-c9_Region5_143208460.tif | 1 | 1 | c9_R | 9 | 1 | 9 | 86625 | 9625 | 9625 | 9625 | 86625 9625 |
| 66 | Tile_r1-c10_Region5_143208460.tif | 1 | 1 | c10_ | 10 | 1 | 10 | 96250 | 9625 | 9625 | 9625 | 96250 9625 |

Copy all numbers of column C and column F to column H and column I

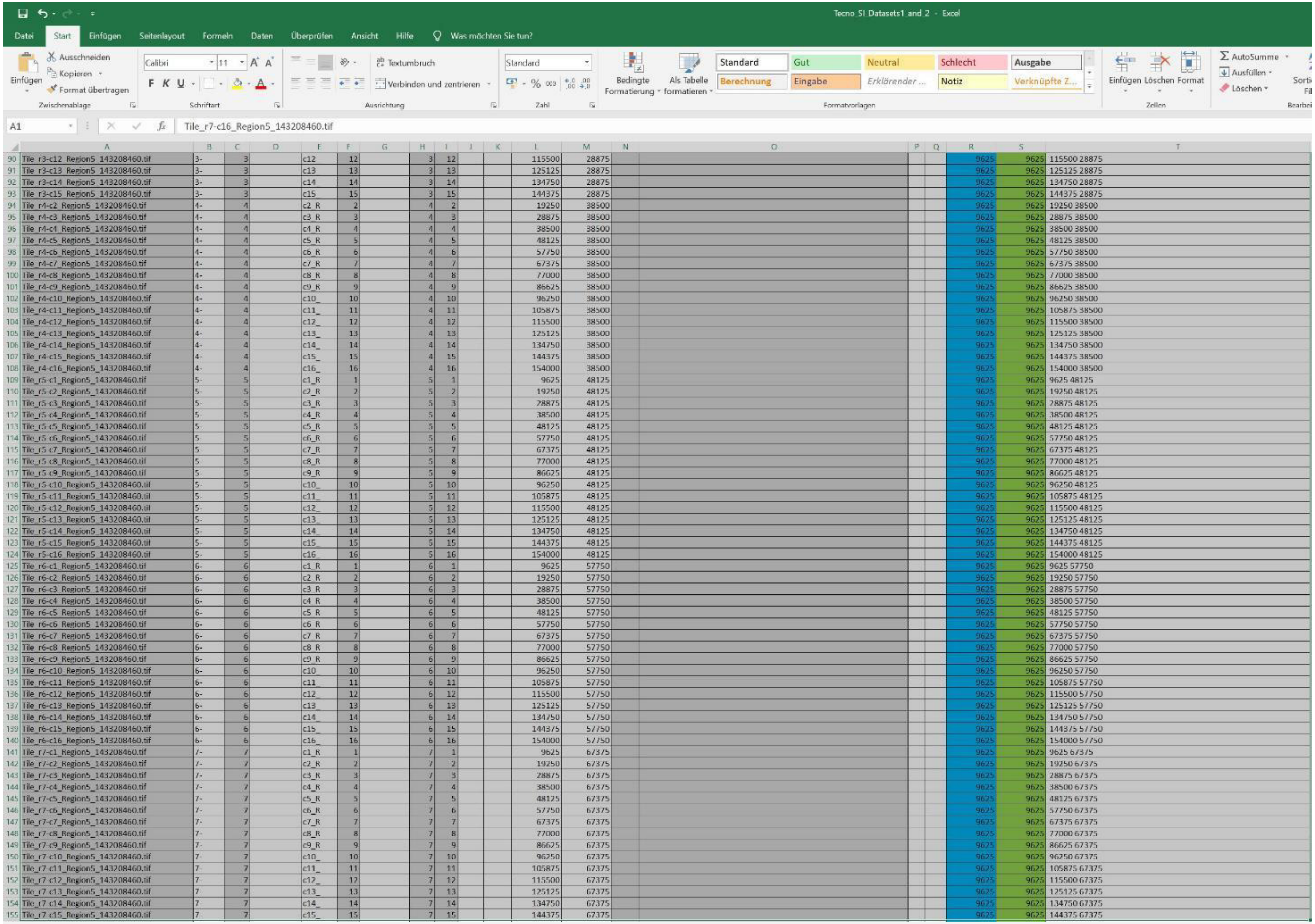


Select all data as shown of dataset 1

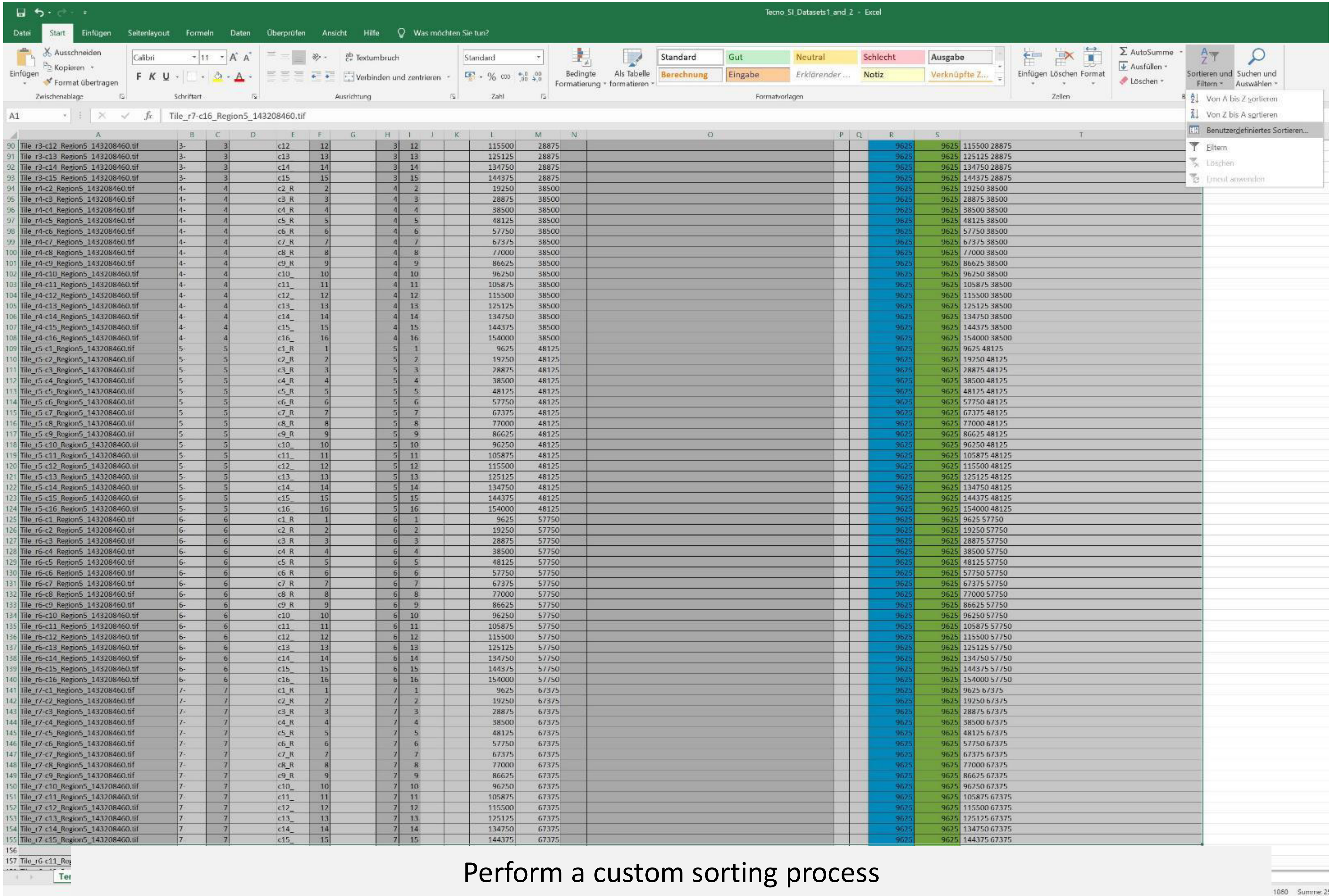


Perform a custom sorting process

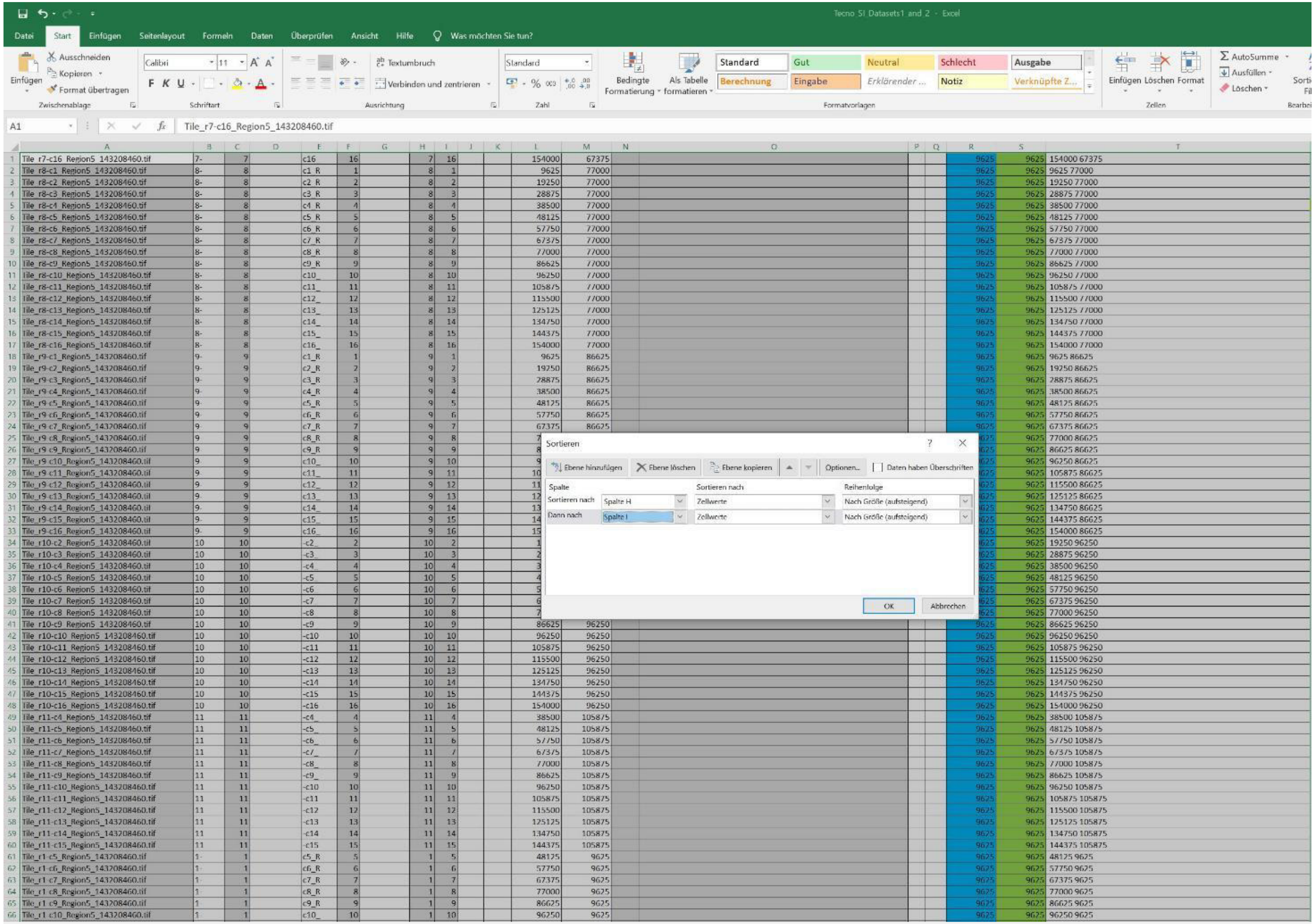


Include column H and column I

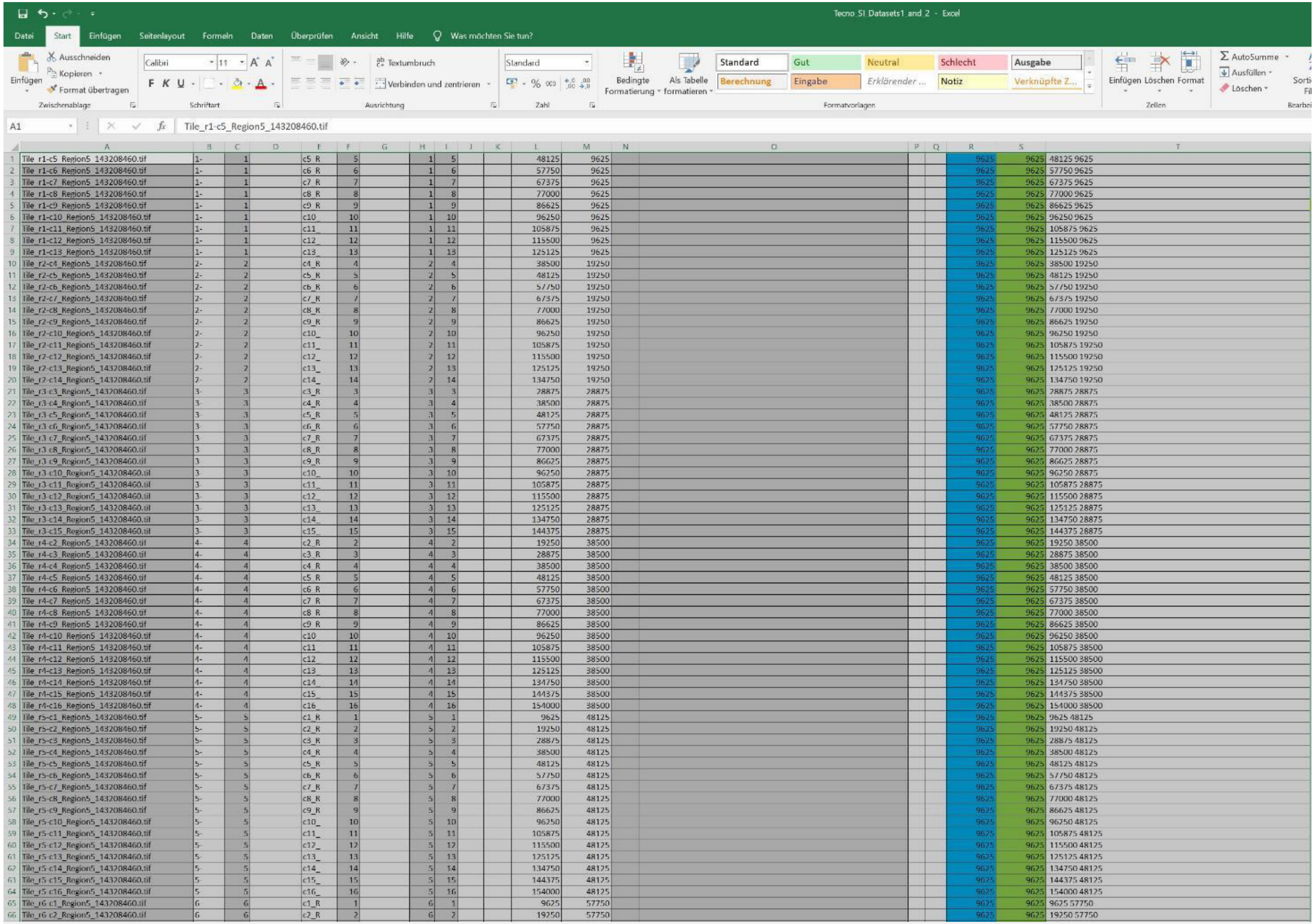

| | A | B | C | E | F | H | I | L | M | R | S | T |
|---|---|---|---|---|---|---|---|---|---|---|---|---|
| 1 | Tile_r1-c5_Region5_143208460.tif | 1- | 1 | c5_R | 5 | 1 | 5 | 48125 | 9625 | 9625 | 9625 | 48125 9625 |
| 2 | Tile_r1-c6_Region5_143208460.tif | 1- | 1 | c6_R | 6 | 1 | 6 | 57750 | 9625 | 9625 | 9625 | 57750 9625 |
| 3 | Tile_r1-c7_Region5_143208460.tif | 1- | 1 | c7_R | 7 | 1 | 7 | 67375 | 9625 | 9625 | 9625 | 67375 9625 |
| 4 | Tile_r1-c8_Region5_143208460.tif | 1- | 1 | c8_R | 8 | 1 | 8 | 77000 | 9625 | 9625 | 9625 | 77000 9625 |
| 5 | Tile_r1-c9_Region5_143208460.tif | 1- | 1 | c9_R | 9 | 1 | 9 | 86625 | 9625 | 9625 | 9625 | 86625 9625 |
| 6 | Tile_r1-c10_Region5_143208460.tif | 1- | 1 | c10_ | 10 | 1 | 10 | 96250 | 9625 | 9625 | 9625 | 96250 9625 |
| 7 | Tile_r1-c11_Region5_143208460.tif | 1- | 1 | c11_ | 11 | 1 | 11 | 105875 | 9625 | 9625 | 9625 | 105875 9625 |
| 8 | Tile_r1-c12_Region5_143208460.tif | 1- | 1 | c12_ | 12 | 1 | 12 | 115500 | 9625 | 9625 | 9625 | 115500 9625 |
| 9 | Tile_r1-c13_Region5_143208460.tif | 1- | 1 | c13_ | 13 | 1 | 13 | 125125 | 9625 | 9625 | 9625 | 125125 9625 |
| 10 | Tile_r2-c4_Region5_143208460.tif | 2- | 2 | c4_R | 4 | 2 | 4 | 38500 | 19250 | 9625 | 9625 | 38500 19250 |
| 11 | Tile_r2-c5_Region5_143208460.tif | 2- | 2 | c5_R | 5 | 2 | 5 | 48125 | 19250 | 9625 | 9625 | 48125 19250 |
| 12 | Tile_r2-c6_Region5_143208460.tif | 2- | 2 | c6_R | 6 | 2 | 6 | 57750 | 19250 | 9625 | 9625 | 57750 19250 |
| 13 | Tile_r2-c7_Region5_143208460.tif | 2- | 2 | c7_R | 7 | 2 | 7 | 67375 | 19250 | 9625 | 9625 | 67375 19250 |
| 14 | Tile_r2-c8_Region5_143208460.tif | 2- | 2 | c8_R | 8 | 2 | 8 | 77000 | 19250 | 9625 | 9625 | 77000 19250 |
| 15 | Tile_r2-c9_Region5_143208460.tif | 2- | 2 | c9_R | 9 | 2 | 9 | 86625 | 19250 | 9625 | 9625 | 86625 19250 |
| 16 | Tile_r2-c10_Region5_143208460.tif | 2- | 2 | c10_ | 10 | 2 | 10 | 96250 | 19250 | 9625 | 9625 | 96250 19250 |
| 17 | Tile_r2-c11_Region5_143208460.tif | 2- | 2 | c11_ | 11 | 2 | 11 | 105875 | 19250 | 9625 | 9625 | 105875 19250 |
| 18 | Tile_r2-c12_Region5_143208460.tif | 2- | 2 | c12_ | 12 | 2 | 12 | 115500 | 19250 | 9625 | 9625 | 115500 19250 |
| 19 | Tile_r2-c13_Region5_143208460.tif | 2- | 2 | c13_ | 13 | 2 | 13 | 125125 | 19250 | 9625 | 9625 | 125125 19250 |
| 20 | Tile_r2-c14_Region5_143208460.tif | 2- | 2 | c14_ | 14 | 2 | 14 | 134750 | 19250 | 9625 | 9625 | 134750 19250 |
| 21 | Tile_r3-c3_Region5_143208460.tif | 3- | 3 | c3_R | 3 | 3 | 3 | 28875 | 28875 | 9625 | 9625 | 28875 28875 |
| 22 | Tile_r3-c4_Region5_143208460.tif | 3- | 3 | c4_R | 4 | 3 | 4 | 38500 | 28875 | 9625 | 9625 | 38500 28875 |
| 23 | Tile_r3-c5_Region5_143208460.tif | 3- | 3 | c5_R | 5 | 3 | 5 | 48125 | 28875 | 9625 | 9625 | 48125 28875 |
| 24 | Tile_r3-c6_Region5_143208460.tif | 3- | 3 | c6_R | 6 | 3 | 6 | 57750 | 28875 | 9625 | 9625 | 57750 28875 |
| 25 | Tile_r3-c7_Region5_143208460.tif | 3 | 3 | c7_R | 7 | 3 | 7 | 67375 | 28875 | 9625 | 9625 | 67375 28875 |
| 26 | Tile_r3-c8_Region5_143208460.tif | 3 | 3 | c8_R | 8 | 3 | 8 | 77000 | 28875 | 9625 | 9625 | 77000 28875 |
| 27 | Tile_r3-c9_Region5_143208460.tif | 3 | 3 | c9_R | 9 | 3 | 9 | 86625 | 28875 | 9625 | 9625 | 86625 28875 |
| 28 | Tile_r3-c10_Region5_143208460.tif | 3- | 3 | c10_ | 10 | 3 | 10 | 96250 | 28875 | 9625 | 9625 | 96250 28875 |
| 29 | Tile_r3-c11_Region5_143208460.tif | 3- | 3 | c11_ | 11 | 3 | 11 | 105875 | 28875 | 9625 | 9625 | 105875 28875 |
| 30 | Tile_r3-c12_Region5_143208460.tif | 3- | 3 | c12_ | 12 | 3 | 12 | 115500 | 28875 | 9625 | 9625 | 115500 28875 |
| 31 | Tile_r3-c13_Region5_143208460.tif | 3- | 3 | c13_ | 13 | 3 | 13 | 125125 | 28875 | 9625 | 9625 | 125125 28875 |
| 32 | Tile_r3-c14_Region5_143208460.tif | 3- | 3 | c14_ | 14 | 3 | 14 | 134750 | 28875 | 9625 | 9625 | 134750 28875 |
| 33 | Tile_r3-c15_Region5_143208460.tif | 3- | 3 | c15_ | 15 | 3 | 15 | 144375 | 28875 | 9625 | 9625 | 144375 28875 |
| 34 | Tile_r4-c2_Region5_143208460.tif | 4- | 4 | c2_R | 2 | 4 | 2 | 19250 | 38500 | 9625 | 9625 | 19250 38500 |
| 35 | Tile_r4-c3_Region5_143208460.tif | 4- | 4 | c3_R | 3 | 4 | 3 | 28875 | 38500 | 9625 | 9625 | 28875 38500 |
| 36 | Tile_r4-c4_Region5_143208460.tif | 4- | 4 | c4_R | 4 | 4 | 4 | 38500 | 38500 | 9625 | 9625 | 38500 38500 |
| 37 | Tile_r4-c5_Region5_143208460.tif | 4- | 4 | c5_R | 5 | 4 | 5 | 48125 | 38500 | 9625 | 9625 | 48125 38500 |
| 38 | Tile_r4-c6_Region5_143208460.tif | 4- | 4 | c6_R | 6 | 4 | 6 | 57750 | 38500 | 9625 | 9625 | 57750 38500 |
| 39 | Tile_r4-c7_Region5_143208460.tif | 4- | 4 | c7_R | 7 | 4 | 7 | 67375 | 38500 | 9625 | 9625 | 67375 38500 |
| 40 | Tile_r4-c8_Region5_143208460.tif | 4- | 4 | c8_R | 8 | 4 | 8 | 77000 | 38500 | 9625 | 9625 | 77000 38500 |
| 41 | Tile_r4-c9_Region5_143208460.tif | 4- | 4 | c9_R | 9 | 4 | 9 | 86625 | 38500 | 9625 | 9625 | 86625 38500 |
| 42 | Tile_r4-c10_Region5_143208460.tif | 4- | 4 | c10_ | 10 | 4 | 10 | 96250 | 38500 | 9625 | 9625 | 96250 38500 |
| 43 | Tile_r4-c11_Region5_143208460.tif | 4- | 4 | c11_ | 11 | 4 | 11 | 105875 | 38500 | 9625 | 9625 | 105875 38500 |
| 44 | Tile_r4-c12_Region5_143208460.tif | 4- | 4 | c12_ | 12 | 4 | 12 | 115500 | 38500 | 9625 | 9625 | 115500 38500 |
| 45 | Tile_r4-c13_Region5_143208460.tif | 4- | 4 | c13_ | 13 | 4 | 13 | 125125 | 38500 | 9625 | 9625 | 125125 38500 |
| 46 | Tile_r4-c14_Region5_143208460.tif | 4- | 4 | c14_ | 14 | 4 | 14 | 134750 | 38500 | 9625 | 9625 | 134750 38500 |
| 47 | Tile_r4-c15_Region5_143208460.tif | 4- | 4 | c15_ | 15 | 4 | 15 | 144375 | 38500 | 9625 | 9625 | 144375 38500 |
| 48 | Tile_r4-c16_Region5_143208460.tif | 4- | 4 | c16_ | 16 | 4 | 16 | 154000 | 38500 | 9625 | 9625 | 154000 38500 |
| 49 | Tile_r5-c1_Region5_143208460.tif | 5- | 5 | c1_R | 1 | 5 | 1 | 9625 | 48125 | 9625 | 9625 | 9625 48125 |
| 50 | Tile_r5-c2_Region5_143208460.tif | 5- | 5 | c2_R | 2 | 5 | 2 | 19250 | 48125 | 9625 | 9625 | 19250 48125 |
| 51 | Tile_r5-c3_Region5_143208460.tif | 5- | 5 | c3_R | 3 | 5 | 3 | 28875 | 48125 | 9625 | 9625 | 28875 48125 |
| 52 | Tile_r5-c4_Region5_143208460.tif | 5- | 5 | c4_R | 4 | 5 | 4 | 38500 | 48125 | 9625 | 9625 | 38500 48125 |
| 53 | Tile_r5-c5_Region5_143208460.tif | 5- | 5 | c5_R | 5 | 5 | 5 | 48125 | 48125 | 9625 | 9625 | 48125 48125 |
| 54 | Tile_r5-c6_Region5_143208460.tif | 5- | 5 | c6_R | 6 | 5 | 6 | 57750 | 48125 | 9625 | 9625 | 57750 48125 |
| 55 | Tile_r5-c7_Region5_143208460.tif | 5- | 5 | c7_R | 7 | 5 | 7 | 67375 | 48125 | 9625 | 9625 | 67375 48125 |
| 56 | Tile_r5-c8_Region5_143208460.tif | 5- | 5 | c8_R | 8 | 5 | 8 | 77000 | 48125 | 9625 | 9625 | 77000 48125 |
| 57 | Tile_r5-c9_Region5_143208460.tif | 5- | 5 | c9_R | 9 | 5 | 9 | 86625 | 48125 | 9625 | 9625 | 86625 48125 |
| 58 | Tile_r5-c10_Region5_143208460.tif | 5- | 5 | c10_ | 10 | 5 | 10 | 96250 | 48125 | 9625 | 9625 | 96250 48125 |
| 59 | Tile_r5-c11_Region5_143208460.tif | 5- | 5 | c11_ | 11 | 5 | 11 | 105875 | 48125 | 9625 | 9625 | 105875 48125 |
| 60 | Tile_r5-c12_Region5_143208460.tif | 5- | 5 | c12_ | 12 | 5 | 12 | 115500 | 48125 | 9625 | 9625 | 115500 48125 |
| 61 | Tile_r5-c13_Region5_143208460.tif | 5- | 5 | c13_ | 13 | 5 | 13 | 125125 | 48125 | 9625 | 9625 | 125125 48125 |
| 62 | Tile_r5-c14_Region5_143208460.tif | 5- | 5 | c14_ | 14 | 5 | 14 | 134750 | 48125 | 9625 | 9625 | 134750 48125 |
| 63 | Tile_r5-c15_Region5_143208460.tif | 5- | 5 | c15_ | 15 | 5 | 15 | 144375 | 48125 | 9625 | 9625 | 144375 48125 |
| 64 | Tile_r5-c16_Region5_143208460.tif | 5- | 5 | c16_ | 16 | 5 | 16 | 154000 | 48125 | 9625 | 9625 | 154000 48125 |
| 65 | Tile_r6-c1_Region5_143208460.tif | 6 | 6 | c1_R | 1 | 6 | 1 | 9625 | 57750 | 9625 | 9625 | 9625 57750 |
| 66 | Tile_r6-c2_Region5_143208460.tif | 6 | 6 | c2_R | 2 | 6 | 2 | 19250 | 57750 | 9625 | 9625 | 19250 57750 |

As a result, all rows will be sorted in the order of STEM image tile acqusition

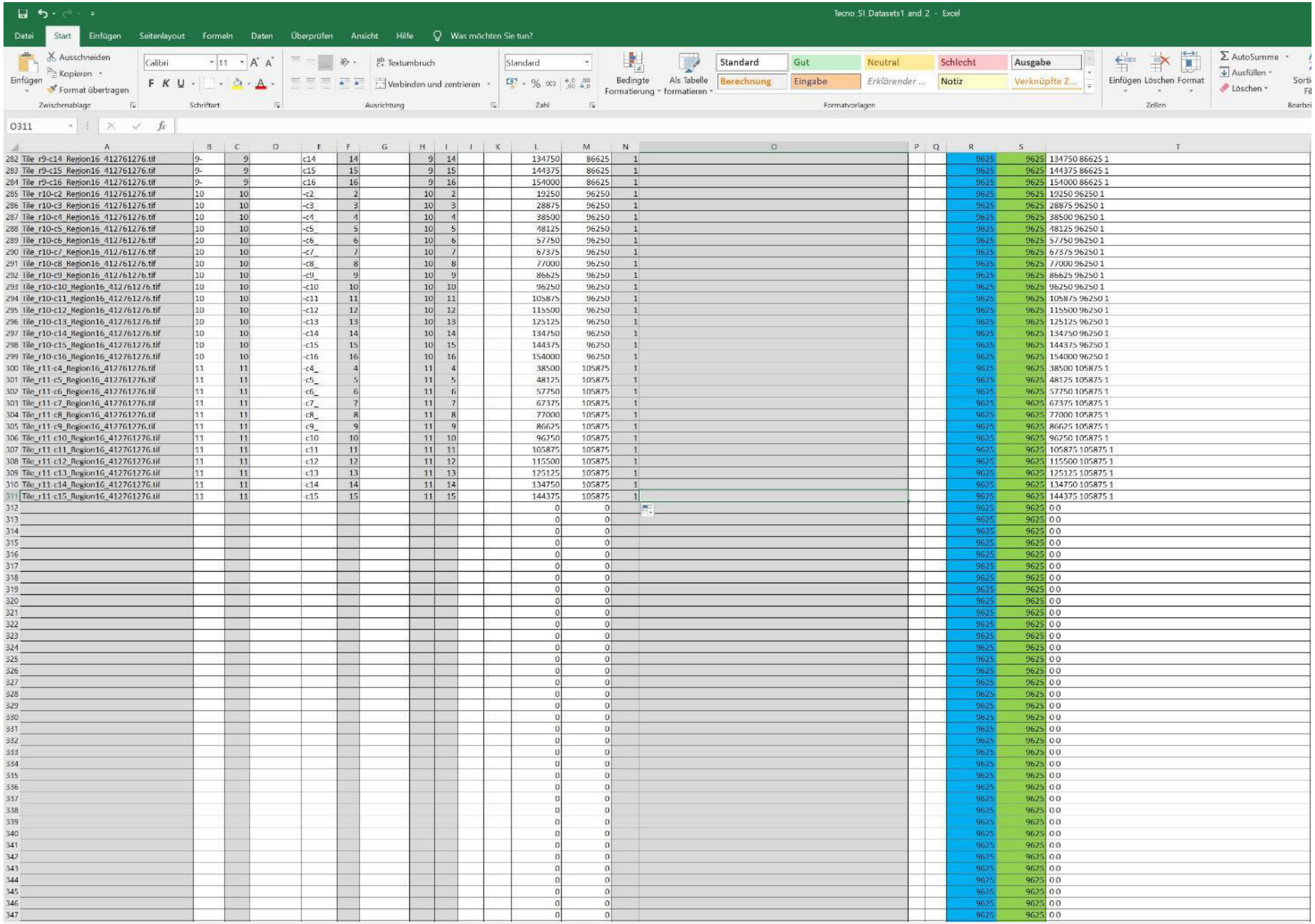


| | A | B | C | D | E | F | G | H | I | J | K | L | M | N | O | P | Q | R | S | T |
|---|---|---|---|---|---|---|---|---|---|---|---|---|---|---|---|---|---|---|---|---|
| 282 | Tile_r9-c14_Region16_412761276.tif | 9- | 9 | | c14 | 14 | | 9 | 14 | | | 134750 | 86625 | 1 | | | | 9625 | 9625 | 134750 86625 1 |
| 283 | Tile_r9-c15_Region16_412761276.tif | 9- | 9 | | c15 | 15 | | 9 | 15 | | | 144375 | 86625 | 1 | | | | 9625 | 9625 | 144375 86625 1 |
| 284 | Tile_r9-c16_Region16_412761276.tif | 9- | 9 | | c16 | 16 | | 9 | 16 | | | 154000 | 86625 | 1 | | | | 9625 | 9625 | 154000 86625 1 |
| 285 | Tile_r10-c2_Region16_412761276.tif | 10 | 10 | | -c2_ | 2 | | 10 | 2 | | | 19250 | 96250 | 1 | | | | 9625 | 9625 | 19250 96250 1 |
| 286 | Tile_r10-c3_Region16_412761276.tif | 10 | 10 | | -c3_ | 3 | | 10 | 3 | | | 28875 | 96250 | 1 | | | | 9625 | 9625 | 28875 96250 1 |
| 287 | Tile_r10-c4_Region16_412761276.tif | 10 | 10 | | -c4_ | 4 | | 10 | 4 | | | 38500 | 96250 | 1 | | | | 9625 | 9625 | 38500 96250 1 |
| 288 | Tile_r10-c5_Region16_412761276.tif | 10 | 10 | | -c5_ | 5 | | 10 | 5 | | | 48125 | 96250 | 1 | | | | 9625 | 9625 | 48125 96250 1 |
| 289 | Tile_r10-c6_Region16_412761276.tif | 10 | 10 | | -c6_ | 6 | | 10 | 6 | | | 57750 | 96250 | 1 | | | | 9625 | 9625 | 57750 96250 1 |
| 290 | Tile_r10-c7_Region16_412761276.tif | 10 | 10 | | -c7_ | 7 | | 10 | 7 | | | 67375 | 96250 | 1 | | | | 9625 | 9625 | 67375 96250 1 |
| 291 | Tile_r10-c8_Region16_412761276.tif | 10 | 10 | | -c8_ | 8 | | 10 | 8 | | | 77000 | 96250 | 1 | | | | 9625 | 9625 | 77000 96250 1 |
| 292 | Tile_r10-c9_Region16_412761276.tif | 10 | 10 | | -c9_ | 9 | | 10 | 9 | | | 86625 | 96250 | 1 | | | | 9625 | 9625 | 86625 96250 1 |
| 293 | Tile_r10-c10_Region16_412761276.tif | 10 | 10 | | -c10 | 10 | | 10 | 10 | | | 96250 | 96250 | 1 | | | | 9625 | 9625 | 96250 96250 1 |
| 294 | Tile_r10-c11_Region16_412761276.tif | 10 | 10 | | -c11 | 11 | | 10 | 11 | | | 105875 | 96250 | 1 | | | | 9625 | 9625 | 105875 96250 1 |
| 295 | Tile_r10-c12_Region16_412761276.tif | 10 | 10 | | -c12 | 12 | | 10 | 12 | | | 115500 | 96250 | 1 | | | | 9625 | 9625 | 115500 96250 1 |
| 296 | Tile_r10-c13_Region16_412761276.tif | 10 | 10 | | -c13 | 13 | | 10 | 13 | | | 125125 | 96250 | 1 | | | | 9625 | 9625 | 125125 96250 1 |
| 297 | Tile_r10-c14_Region16_412761276.tif | 10 | 10 | | -c14 | 14 | | 10 | 14 | | | 134750 | 96250 | 1 | | | | 9625 | 9625 | 134750 96250 1 |
| 298 | Tile_r10-c15_Region16_412761276.tif | 10 | 10 | | -c15 | 15 | | 10 | 15 | | | 144375 | 96250 | 1 | | | | 9625 | 9625 | 144375 96250 1 |
| 299 | Tile_r10-c16_Region16_412761276.tif | 10 | 10 | | -c16 | 16 | | 10 | 16 | | | 154000 | 96250 | 1 | | | | 9625 | 9625 | 154000 96250 1 |
| 300 | Tile_r11-c4_Region16_412761276.tif | 11 | 11 | | -c4_ | 4 | | 11 | 4 | | | 38500 | 105875 | 1 | | | | 9625 | 9625 | 38500 105875 1 |
| 301 | Tile_r11-c5_Region16_412761276.tif | 11 | 11 | | -c5_ | 5 | | 11 | 5 | | | 48125 | 105875 | 1 | | | | 9625 | 9625 | 48125 105875 1 |
| 302 | Tile_r11-c6_Region16_412761276.tif | 11 | 11 | | -c6_ | 6 | | 11 | 6 | | | 57750 | 105875 | 1 | | | | 9625 | 9625 | 57750 105875 1 |
| 303 | Tile_r11-c7_Region16_412761276.tif | 11 | 11 | | -c7_ | 7 | | 11 | 7 | | | 67375 | 105875 | 1 | | | | 9625 | 9625 | 67375 105875 1 |
| 304 | Tile_r11-c8_Region16_412761276.tif | 11 | 11 | | -c8_ | 8 | | 11 | 8 | | | 77000 | 105875 | 1 | | | | 9625 | 9625 | 77000 105875 1 |
| 305 | Tile_r11-c9_Region16_412761276.tif | 11 | 11 | | -c9_ | 9 | | 11 | 9 | | | 86625 | 105875 | 1 | | | | 9625 | 9625 | 86625 105875 1 |
| 306 | Tile_r11-c10_Region16_412761276.tif | 11 | 11 | | -c10 | 10 | | 11 | 10 | | | 96250 | 105875 | 1 | | | | 9625 | 9625 | 96250 105875 1 |
| 307 | Tile_r11-c11_Region16_412761276.tif | 11 | 11 | | -c11 | 11 | | 11 | 11 | | | 105875 | 105875 | 1 | | | | 9625 | 9625 | 105875 105875 1 |
| 308 | Tile_r11-c12_Region16_412761276.tif | 11 | 11 | | -c12 | 12 | | 11 | 12 | | | 115500 | 105875 | 1 | | | | 9625 | 9625 | 115500 105875 1 |
| 309 | Tile_r11-c13_Region16_412761276.tif | 11 | 11 | | -c13 | 13 | | 11 | 13 | | | 125125 | 105875 | 1 | | | | 9625 | 9625 | 125125 105875 1 |
| 310 | Tile_r11-c14_Region16_412761276.tif | 11 | 11 | | -c14 | 14 | | 11 | 14 | | | 134750 | 105875 | 1 | | | | 9625 | 9625 | 134750 105875 1 |
| 311 | Tile_r11-c15_Region16_412761276.tif | 11 | 11 | | -c15 | 15 | | 11 | 15 | | | 144375 | 105875 | 1 | | | | 9625 | 9625 | 144375 105875 1 |
| 312 | | | | | | | | | | | | 0 | 0 | | | | | 9625 | 9625 | 0 0 |
| 313 | | | | | | | | | | | | 0 | 0 | | | | | 9625 | 9625 | 0 0 |
| 314 | | | | | | | | | | | | 0 | 0 | | | | | 9625 | 9625 | 0 0 |
| 315 | | | | | | | | | | | | 0 | 0 | | | | | 9625 | 9625 | 0 0 |
| 316 | | | | | | | | | | | | 0 | 0 | | | | | 9625 | 9625 | 0 0 |
| 317 | | | | | | | | | | | | 0 | 0 | | | | | 9625 | 9625 | 0 0 |
| 318 | | | | | | | | | | | | 0 | 0 | | | | | 9625 | 9625 | 0 0 |
| 319 | | | | | | | | | | | | 0 | 0 | | | | | 9625 | 9625 | 0 0 |
| 320 | | | | | | | | | | | | 0 | 0 | | | | | 9625 | 9625 | 0 0 |
| 321 | | | | | | | | | | | | 0 | 0 | | | | | 9625 | 9625 | 0 0 |
| 322 | | | | | | | | | | | | 0 | 0 | | | | | 9625 | 9625 | 0 0 |
| 323 | | | | | | | | | | | | 0 | 0 | | | | | 9625 | 9625 | 0 0 |
| 324 | | | | | | | | | | | | 0 | 0 | | | | | 9625 | 9625 | 0 0 |
| 325 | | | | | | | | | | | | 0 | 0 | | | | | 9625 | 9625 | 0 0 |
| 326 | | | | | | | | | | | | 0 | 0 | | | | | 9625 | 9625 | 0 0 |
| 327 | | | | | | | | | | | | 0 | 0 | | | | | 9625 | 9625 | 0 0 |
| 328 | | | | | | | | | | | | 0 | 0 | | | | | 9625 | 9625 | 0 0 |
| 329 | | | | | | | | | | | | 0 | 0 | | | | | 9625 | 9625 | 0 0 |
| 330 | | | | | | | | | | | | 0 | 0 | | | | | 9625 | 9625 | 0 0 |
| 331 | | | | | | | | | | | | 0 | 0 | | | | | 9625 | 9625 | 0 0 |
| 332 | | | | | | | | | | | | 0 | 0 | | | | | 9625 | 9625 | 0 0 |
| 333 | | | | | | | | | | | | 0 | 0 | | | | | 9625 | 9625 | 0 0 |
| 334 | | | | | | | | | | | | 0 | 0 | | | | | 9625 | 9625 | 0 0 |
| 335 | | | | | | | | | | | | 0 | 0 | | | | | 9625 | 9625 | 0 0 |
| 336 | | | | | | | | | | | | 0 | 0 | | | | | 9625 | 9625 | 0 0 |
| 337 | | | | | | | | | | | | 0 | 0 | | | | | 9625 | 9625 | 0 0 |
| 338 | | | | | | | | | | | | 0 | 0 | | | | | 9625 | 9625 | 0 0 |
| 339 | | | | | | | | | | | | 0 | 0 | | | | | 9625 | 9625 | 0 0 |
| 340 | | | | | | | | | | | | 0 | 0 | | | | | 9625 | 9625 | 0 0 |
| 341 | | | | | | | | | | | | 0 | 0 | | | | | 9625 | 9625 | 0 0 |
| 342 | | | | | | | | | | | | 0 | 0 | | | | | 9625 | 9625 | 0 0 |
| 343 | | | | | | | | | | | | 0 | 0 | | | | | 9625 | 9625 | 0 0 |
| 344 | | | | | | | | | | | | 0 | 0 | | | | | 9625 | 9625 | 0 0 |
| 345 | | | | | | | | | | | | 0 | 0 | | | | | 9625 | 9625 | 0 0 |
| 346 | | | | | | | | | | | | 0 | 0 | | | | | 9625 | 9625 | 0 0 |
| 347 | | | | | | | | | | | | 0 | 0 | | | | | 9625 | 9625 | 0 0 |

Include the „z“ coordinate manually in column N; „0“ for dataset 1 and „1“ for dataset 2

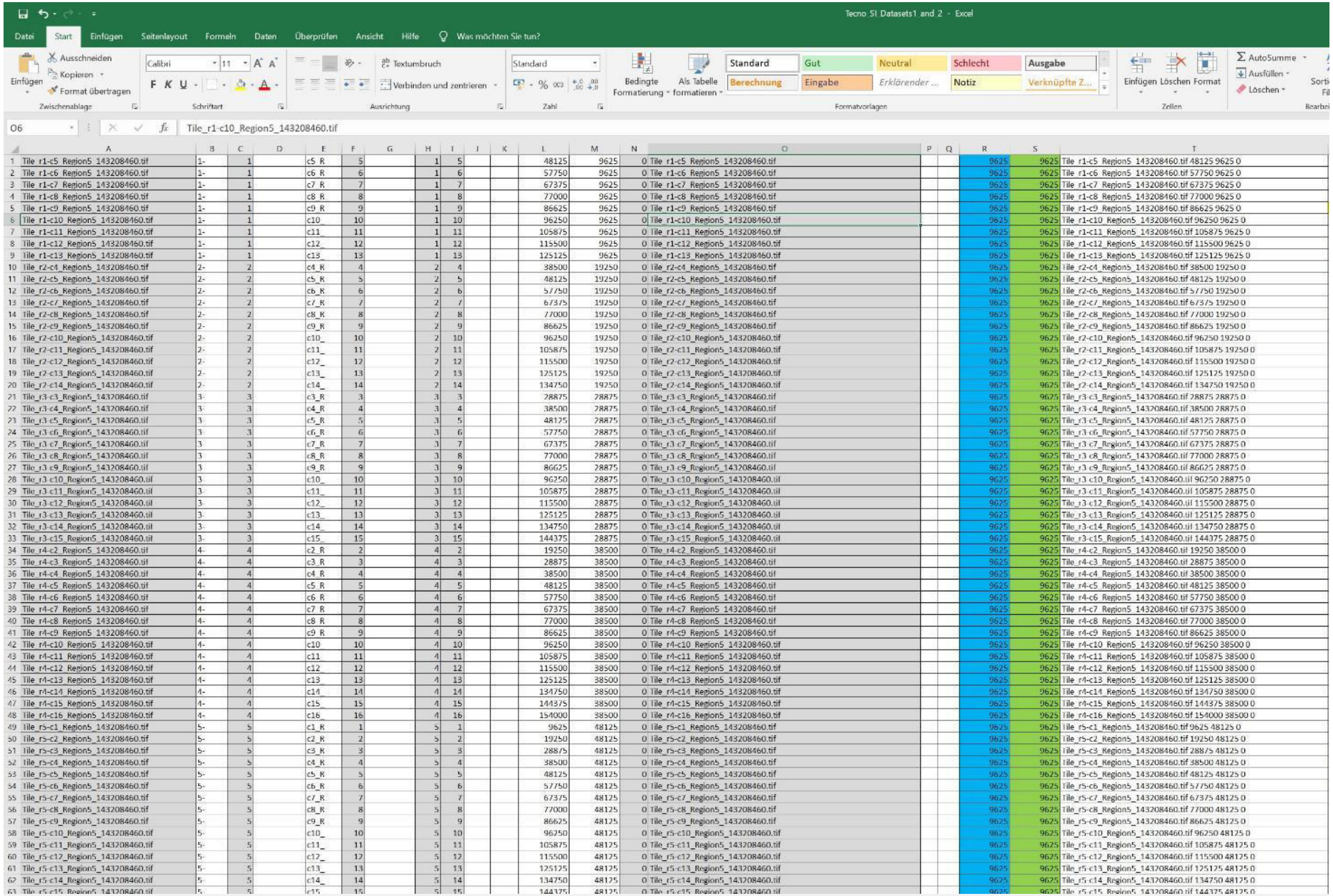


| | A | B | C | D | E | F | G | H | I | J | K | L | M | N | O | P | Q | R | S | T |
|---|---|---|---|---|---|---|---|---|---|---|---|---|---|---|---|---|---|---|---|---|
| 1 | Tile_r1-c5_Region5_143208460.tif | 1- | 1 | | c5_R | 5 | | 1 | 5 | | | 48125 | 9625 | 0 | Tile_r1-c5_Region5_143208460.tif | | | 9625 | 9625 | Tile_r1-c5_Region5_143208460.tif 48125 9625 0 |
| 2 | Tile_r1-c6_Region5_143208460.tif | 1- | 1 | | c6_R | 6 | | 1 | 6 | | | 57750 | 9625 | 0 | Tile_r1-c6_Region5_143208460.tif | | | 9625 | 9625 | Tile_r1-c6_Region5_143208460.tif 57750 9625 0 |
| 3 | Tile_r1-c7_Region5_143208460.tif | 1- | 1 | | c7_R | 7 | | 1 | 7 | | | 67375 | 9625 | 0 | Tile_r1-c7_Region5_143208460.tif | | | 9625 | 9625 | Tile_r1-c7_Region5_143208460.tif 67375 9625 0 |
| 4 | Tile_r1-c8_Region5_143208460.tif | 1- | 1 | | c8_R | 8 | | 1 | 8 | | | 77000 | 9625 | 0 | Tile_r1-c8_Region5_143208460.tif | | | 9625 | 9625 | Tile_r1-c8_Region5_143208460.tif 77000 9625 0 |
| 5 | Tile_r1-c9_Region5_143208460.tif | 1- | 1 | | c9_R | 9 | | 1 | 9 | | | 86625 | 9625 | 0 | Tile_r1-c9_Region5_143208460.tif | | | 9625 | 9625 | Tile_r1-c9_Region5_143208460.tif 86625 9625 0 |
| 6 | Tile_r1-c10_Region5_143208460.tif | 1- | 1 | | c10_ | 10 | | 1 | 10 | | | 96250 | 9625 | 0 | Tile_r1-c10_Region5_143208460.tif | | | 9625 | 9625 | Tile_r1-c10_Region5_143208460.tif 96250 9625 0 |
| 7 | Tile_r1-c11_Region5_143208460.tif | 1- | 1 | | c11_ | 11 | | 1 | 11 | | | 105875 | 9625 | 0 | Tile_r1-c11_Region5_143208460.tif | | | 9625 | 9625 | Tile_r1-c11_Region5_143208460.tif 105875 9625 0 |
| 8 | Tile_r1-c12_Region5_143208460.tif | 1- | 1 | | c12_ | 12 | | 1 | 12 | | | 115500 | 9625 | 0 | Tile_r1-c12_Region5_143208460.tif | | | 9625 | 9625 | Tile_r1-c12_Region5_143208460.tif 115500 9625 0 |
| 9 | Tile_r1-c13_Region5_143208460.tif | 1- | 1 | | c13_ | 13 | | 1 | 13 | | | 125125 | 9625 | 0 | Tile_r1-c13_Region5_143208460.tif | | | 9625 | 9625 | Tile_r1-c13_Region5_143208460.tif 125125 9625 0 |
| 10 | Tile_r2-c4_Region5_143208460.tif | 2- | 2 | | c4_R | 4 | | 2 | 4 | | | 38500 | 19250 | 0 | Tile_r2-c4_Region5_143208460.tif | | | 9625 | 9625 | Tile_r2-c4_Region5_143208460.tif 38500 19250 0 |
| 11 | Tile_r2-c5_Region5_143208460.tif | 2- | 2 | | c5_R | 5 | | 2 | 5 | | | 48125 | 19250 | 0 | Tile_r2-c5_Region5_143208460.tif | | | 9625 | 9625 | Tile_r2-c5_Region5_143208460.tif 48125 19250 0 |
| 12 | Tile_r2-c6_Region5_143208460.tif | 2- | 2 | | c6_R | 6 | | 2 | 6 | | | 57750 | 19250 | 0 | Tile_r2-c6_Region5_143208460.tif | | | 9625 | 9625 | Tile_r2-c6_Region5_143208460.tif 57750 19250 0 |
| 13 | Tile_r2-c7_Region5_143208460.tif | 2- | 2 | | c7_R | 7 | | 2 | 7 | | | 67375 | 19250 | 0 | Tile_r2-c7_Region5_143208460.tif | | | 9625 | 9625 | Tile_r2-c7_Region5_143208460.tif 67375 19250 0 |
| 14 | Tile_r2-c8_Region5_143208460.tif | 2- | 2 | | c8_R | 8 | | 2 | 8 | | | 77000 | 19250 | 0 | Tile_r2-c8_Region5_143208460.tif | | | 9625 | 9625 | Tile_r2-c8_Region5_143208460.tif 77000 19250 0 |
| 15 | Tile_r2-c9_Region5_143208460.tif | 2- | 2 | | c9_R | 9 | | 2 | 9 | | | 86625 | 19250 | 0 | Tile_r2-c9_Region5_143208460.tif | | | 9625 | 9625 | Tile_r2-c9_Region5_143208460.tif 86625 19250 0 |
| 16 | Tile_r2-c10_Region5_143208460.tif | 2- | 2 | | c10_ | 10 | | 2 | 10 | | | 96250 | 19250 | 0 | Tile_r2-c10_Region5_143208460.tif | | | 9625 | 9625 | Tile_r2-c10_Region5_143208460.tif 96250 19250 0 |
| 17 | Tile_r2-c11_Region5_143208460.tif | 2- | 2 | | c11_ | 11 | | 2 | 11 | | | 105875 | 19250 | 0 | Tile_r2-c11_Region5_143208460.tif | | | 9625 | 9625 | Tile_r2-c11_Region5_143208460.tif 105875 19250 0 |
| 18 | Tile_r2-c12_Region5_143208460.tif | 2- | 2 | | c12_ | 12 | | 2 | 12 | | | 115500 | 19250 | 0 | Tile_r2-c12_Region5_143208460.tif | | | 9625 | 9625 | Tile_r2-c12_Region5_143208460.tif 115500 19250 0 |
| 19 | Tile_r2-c13_Region5_143208460.tif | 2- | 2 | | c13_ | 13 | | 2 | 13 | | | 125125 | 19250 | 0 | Tile_r2-c13_Region5_143208460.tif | | | 9625 | 9625 | Tile_r2-c13_Region5_143208460.tif 125125 19250 0 |
| 20 | Tile_r2-c14_Region5_143208460.tif | 2- | 2 | | c14_ | 14 | | 2 | 14 | | | 134750 | 19250 | 0 | Tile_r2-c14_Region5_143208460.tif | | | 9625 | 9625 | Tile_r2-c14_Region5_143208460.tif 134750 19250 0 |
| 21 | Tile_r3-c3_Region5_143208460.tif | 3- | 3 | | c3_R | 3 | | 3 | 3 | | | 28875 | 28875 | 0 | Tile_r3-c3_Region5_143208460.tif | | | 9625 | 9625 | Tile_r3-c3_Region5_143208460.tif 28875 28875 0 |
| 22 | Tile_r3-c4_Region5_143208460.tif | 3- | 3 | | c4_R | 4 | | 3 | 4 | | | 38500 | 28875 | 0 | Tile_r3-c4_Region5_143208460.tif | | | 9625 | 9625 | Tile_r3-c4_Region5_143208460.tif 38500 28875 0 |
| 23 | Tile_r3-c5_Region5_143208460.tif | 3 | 3 | | c5_R | 5 | | 3 | 5 | | | 48125 | 28875 | 0 | Tile_r3-c5_Region5_143208460.tif | | | 9625 | 9625 | Tile_r3-c5_Region5_143208460.tif 48125 28875 0 |
| 24 | Tile_r3-c6_Region5_143208460.tif | 3 | 3 | | c6_R | 6 | | 3 | 6 | | | 57750 | 28875 | 0 | Tile_r3-c6_Region5_143208460.tif | | | 9625 | 9625 | Tile_r3-c6_Region5_143208460.tif 57750 28875 0 |
| 25 | Tile_r3-c7_Region5_143208460.tif | 3 | 3 | | c7_R | 7 | | 3 | 7 | | | 67375 | 28875 | 0 | Tile_r3-c7_Region5_143208460.tif | | | 9625 | 9625 | Tile_r3-c7_Region5_143208460.tif 67375 28875 0 |
| 26 | Tile_r3-c8_Region5_143208460.tif | 3 | 3 | | c8_R | 8 | | 3 | 8 | | | 77000 | 28875 | 0 | Tile_r3-c8_Region5_143208460.tif | | | 9625 | 9625 | Tile_r3-c8_Region5_143208460.tif 77000 28875 0 |
| 27 | Tile_r3-c9_Region5_143208460.tif | 3 | 3 | | c9_R | 9 | | 3 | 9 | | | 86625 | 28875 | 0 | Tile_r3-c9_Region5_143208460.tif | | | 9625 | 9625 | Tile_r3-c9_Region5_143208460.tif 86625 28875 0 |
| 28 | Tile_r3-c10_Region5_143208460.tif | 3- | 3 | | c10_ | 10 | | 3 | 10 | | | 96250 | 28875 | 0 | Tile_r3-c10_Region5_143208460.tif | | | 9625 | 9625 | Tile_r3-c10_Region5_143208460.tif 96250 28875 0 |
| 29 | Tile_r3-c11_Region5_143208460.tif | 3- | 3 | | c11_ | 11 | | 3 | 11 | | | 105875 | 28875 | 0 | Tile_r3-c11_Region5_143208460.tif | | | 9625 | 9625 | Tile_r3-c11_Region5_143208460.tif 105875 28875 0 |
| 30 | Tile_r3-c12_Region5_143208460.tif | 3- | 3 | | c12_ | 12 | | 3 | 12 | | | 115500 | 28875 | 0 | Tile_r3-c12_Region5_143208460.tif | | | 9625 | 9625 | Tile_r3-c12_Region5_143208460.tif 115500 28875 0 |
| 31 | Tile_r3-c13_Region5_143208460.tif | 3- | 3 | | c13_ | 13 | | 3 | 13 | | | 125125 | 28875 | 0 | Tile_r3-c13_Region5_143208460.tif | | | 9625 | 9625 | Tile_r3-c13_Region5_143208460.tif 125125 28875 0 |
| 32 | Tile_r3-c14_Region5_143208460.tif | 3- | 3 | | c14_ | 14 | | 3 | 14 | | | 134750 | 28875 | 0 | Tile_r3-c14_Region5_143208460.tif | | | 9625 | 9625 | Tile_r3-c14_Region5_143208460.tif 134750 28875 0 |
| 33 | Tile_r3-c15_Region5_143208460.tif | 3- | 3 | | c15_ | 15 | | 3 | 15 | | | 144375 | 28875 | 0 | Tile_r3-c15_Region5_143208460.tif | | | 9625 | 9625 | Tile_r3-c15_Region5_143208460.tif 144375 28875 0 |
| 34 | Tile_r4-c2_Region5_143208460.tif | 4- | 4 | | c2_R | 2 | | 4 | 2 | | | 19250 | 38500 | 0 | Tile_r4-c2_Region5_143208460.tif | | | 9625 | 9625 | Tile_r4-c2_Region5_143208460.tif 19250 38500 0 |
| 35 | Tile_r4-c3_Region5_143208460.tif | 4- | 4 | | c3_R | 3 | | 4 | 3 | | | 28875 | 38500 | 0 | Tile_r4-c3_Region5_143208460.tif | | | 9625 | 9625 | Tile_r4-c3_Region5_143208460.tif 28875 38500 0 |
| 36 | Tile_r4-c4_Region5_143208460.tif | 4- | 4 | | c4_R | 4 | | 4 | 4 | | | 38500 | 38500 | 0 | Tile_r4-c4_Region5_143208460.tif | | | 9625 | 9625 | Tile_r4-c4_Region5_143208460.tif 38500 38500 0 |
| 37 | Tile_r4-c5_Region5_143208460.tif | 4- | 4 | | c5_R | 5 | | 4 | 5 | | | 48125 | 38500 | 0 | Tile_r4-c5_Region5_143208460.tif | | | 9625 | 9625 | Tile_r4-c5_Region5_143208460.tif 48125 38500 0 |
| 38 | Tile_r4-c6_Region5_143208460.tif | 4- | 4 | | c6_R | 6 | | 4 | 6 | | | 57750 | 38500 | 0 | Tile_r4-c6_Region5_143208460.tif | | | 9625 | 9625 | Tile_r4-c6_Region5_143208460.tif 57750 38500 0 |
| 39 | Tile_r4-c7_Region5_143208460.tif | 4- | 4 | | c7_R | 7 | | 4 | 7 | | | 67375 | 38500 | 0 | Tile_r4-c7_Region5_143208460.tif | | | 9625 | 9625 | Tile_r4-c7_Region5_143208460.tif 67375 38500 0 |
| 40 | Tile_r4-c8_Region5_143208460.tif | 4- | 4 | | c8_R | 8 | | 4 | 8 | | | 77000 | 38500 | 0 | Tile_r4-c8_Region5_143208460.tif | | | 9625 | 9625 | Tile_r4-c8_Region5_143208460.tif 77000 38500 0 |
| 41 | Tile_r4-c9_Region5_143208460.tif | 4- | 4 | | c9_R | 9 | | 4 | 9 | | | 86625 | 38500 | 0 | Tile_r4-c9_Region5_143208460.tif | | | 9625 | 9625 | Tile_r4-c9_Region5_143208460.tif 86625 38500 0 |
| 42 | Tile_r4-c10_Region5_143208460.tif | 4- | 4 | | c10 | 10 | | 4 | 10 | | | 96250 | 38500 | 0 | Tile_r4-c10_Region5_143208460.tif | | | 9625 | 9625 | Tile_r4-c10_Region5_143208460.tif 96250 38500 0 |
| 43 | Tile_r4-c11_Region5_143208460.tif | 4- | 4 | | c11 | 11 | | 4 | 11 | | | 105875 | 38500 | 0 | Tile_r4-c11_Region5_143208460.tif | | | 9625 | 9625 | Tile_r4-c11_Region5_143208460.tif 105875 38500 0 |
| 44 | Tile_r4-c12_Region5_143208460.tif | 4- | 4 | | c12_ | 12 | | 4 | 12 | | | 115500 | 38500 | 0 | Tile_r4-c12_Region5_143208460.tif | | | 9625 | 9625 | Tile_r4-c12_Region5_143208460.tif 115500 38500 0 |
| 45 | Tile_r4-c13_Region5_143208460.tif | 4- | 4 | | c13_ | 13 | | 4 | 13 | | | 125125 | 38500 | 0 | Tile_r4-c13_Region5_143208460.tif | | | 9625 | 9625 | Tile_r4-c13_Region5_143208460.tif 125125 38500 0 |
| 46 | Tile_r4-c14_Region5_143208460.tif | 4- | 4 | | c14_ | 14 | | 4 | 14 | | | 134750 | 38500 | 0 | Tile_r4-c14_Region5_143208460.tif | | | 9625 | 9625 | Tile_r4-c14_Region5_143208460.tif 134750 38500 0 |
| 47 | Tile_r4-c15_Region5_143208460.tif | 4- | 4 | | c15_ | 15 | | 4 | 15 | | | 144375 | 38500 | 0 | Tile_r4-c15_Region5_143208460.tif | | | 9625 | 9625 | Tile_r4-c15_Region5_143208460.tif 144375 38500 0 |
| 48 | Tile_r4-c16_Region5_143208460.tif | 4- | 4 | | c16_ | 16 | | 4 | 16 | | | 154000 | 38500 | 0 | Tile_r4-c16_Region5_143208460.tif | | | 9625 | 9625 | Tile_r4-c16_Region5_143208460.tif 154000 38500 0 |
| 49 | Tile_r5-c1_Region5_143208460.tif | 5- | 5 | | c1_R | 1 | | 5 | 1 | | | 9625 | 48125 | 0 | Tile_r5-c1_Region5_143208460.tif | | | 9625 | 9625 | Tile_r5-c1_Region5_143208460.tif 9625 48125 0 |
| 50 | Tile_r5-c2_Region5_143208460.tif | 5- | 5 | | c2_R | 2 | | 5 | 2 | | | 19250 | 48125 | 0 | Tile_r5-c2_Region5_143208460.tif | | | 9625 | 9625 | Tile_r5-c2_Region5_143208460.tif 19250 48125 0 |
| 51 | Tile_r5-c3_Region5_143208460.tif | 5- | 5 | | c3_R | 3 | | 5 | 3 | | | 28875 | 48125 | 0 | Tile_r5-c3_Region5_143208460.tif | | | 9625 | 9625 | Tile_r5-c3_Region5_143208460.tif 28875 48125 0 |
| 52 | Tile_r5-c4_Region5_143208460.tif | 5- | 5 | | c4_R | 4 | | 5 | 4 | | | 38500 | 48125 | 0 | Tile_r5-c4_Region5_143208460.tif | | | 9625 | 9625 | Tile_r5-c4_Region5_143208460.tif 38500 48125 0 |
| 53 | Tile_r5-c5_Region5_143208460.tif | 5- | 5 | | c5_R | 5 | | 5 | 5 | | | 48125 | 48125 | 0 | Tile_r5-c5_Region5_143208460.tif | | | 9625 | 9625 | Tile_r5-c5_Region5_143208460.tif 48125 48125 0 |
| 54 | Tile_r5-c6_Region5_143208460.tif | 5- | 5 | | c6_R | 6 | | 5 | 6 | | | 57750 | 48125 | 0 | Tile_r5-c6_Region5_143208460.tif | | | 9625 | 9625 | Tile_r5-c6_Region5_143208460.tif 57750 48125 0 |
| 55 | Tile_r5-c7_Region5_143208460.tif | 5- | 5 | | c7_R | 7 | | 5 | 7 | | | 67375 | 48125 | 0 | Tile_r5-c7_Region5_143208460.tif | | | 9625 | 9625 | Tile_r5-c7_Region5_143208460.tif 67375 48125 0 |
| 56 | Tile_r5-c8_Region5_143208460.tif | 5- | 5 | | c8_R | 8 | | 5 | 8 | | | 77000 | 48125 | 0 | Tile_r5-c8_Region5_143208460.tif | | | 9625 | 9625 | Tile_r5-c8_Region5_143208460.tif 77000 48125 0 |
| 57 | Tile_r5-c9_Region5_143208460.tif | 5- | 5 | | c9_R | 9 | | 5 | 9 | | | 86625 | 48125 | 0 | Tile_r5-c9_Region5_143208460.tif | | | 9625 | 9625 | Tile_r5-c9_Region5_143208460.tif 86625 48125 0 |
| 58 | Tile_r5-c10_Region5_143208460.tif | 5- | 5 | | c10_ | 10 | | 5 | 10 | | | 96250 | 48125 | 0 | Tile_r5-c10_Region5_143208460.tif | | | 9625 | 9625 | Tile_r5-c10_Region5_143208460.tif 96250 48125 0 |
| 59 | Tile_r5-c11_Region5_143208460.tif | 5- | 5 | | c11_ | 11 | | 5 | 11 | | | 105875 | 48125 | 0 | Tile_r5-c11_Region5_143208460.tif | | | 9625 | 9625 | Tile_r5-c11_Region5_143208460.tif 105875 48125 0 |
| 60 | Tile_r5-c12_Region5_143208460.tif | 5- | 5 | | c12_ | 12 | | 5 | 12 | | | 115500 | 48125 | 0 | Tile_r5-c12_Region5_143208460.tif | | | 9625 | 9625 | Tile_r5-c12_Region5_143208460.tif 115500 48125 0 |
| 61 | Tile_r5-c13_Region5_143208460.tif | 5- | 5 | | c13_ | 13 | | 5 | 13 | | | 125125 | 48125 | 0 | Tile_r5-c13_Region5_143208460.tif | | | 9625 | 9625 | Tile_r5-c13_Region5_143208460.tif 125125 48125 0 |
| 62 | Tile_r5-c14_Region5_143208460.tif | 5- | 5 | | c14_ | 14 | | 5 | 14 | | | 134750 | 48125 | 0 | Tile_r5-c14_Region5_143208460.tif | | | 9625 | 9625 | Tile_r5-c14_Region5_143208460.tif 134750 48125 0 |
| 63 | Tile_r5-c15_Region5_143208460.tif | 5- | 5 | | c15_ | 15 | | 5 | 15 | | | 144375 | 48125 | 0 | Tile_r5-c15_Region5_143208460.tif | | | 9625 | 9625 | Tile_r5-c15_Region5_143208460.tif 144375 48125 0 |

Copy the names of the image tiles in column A to column O

Note the values in column R,S; image tile size 10,240 pixels, overlap 6%; 10,240*0.94=9625,6

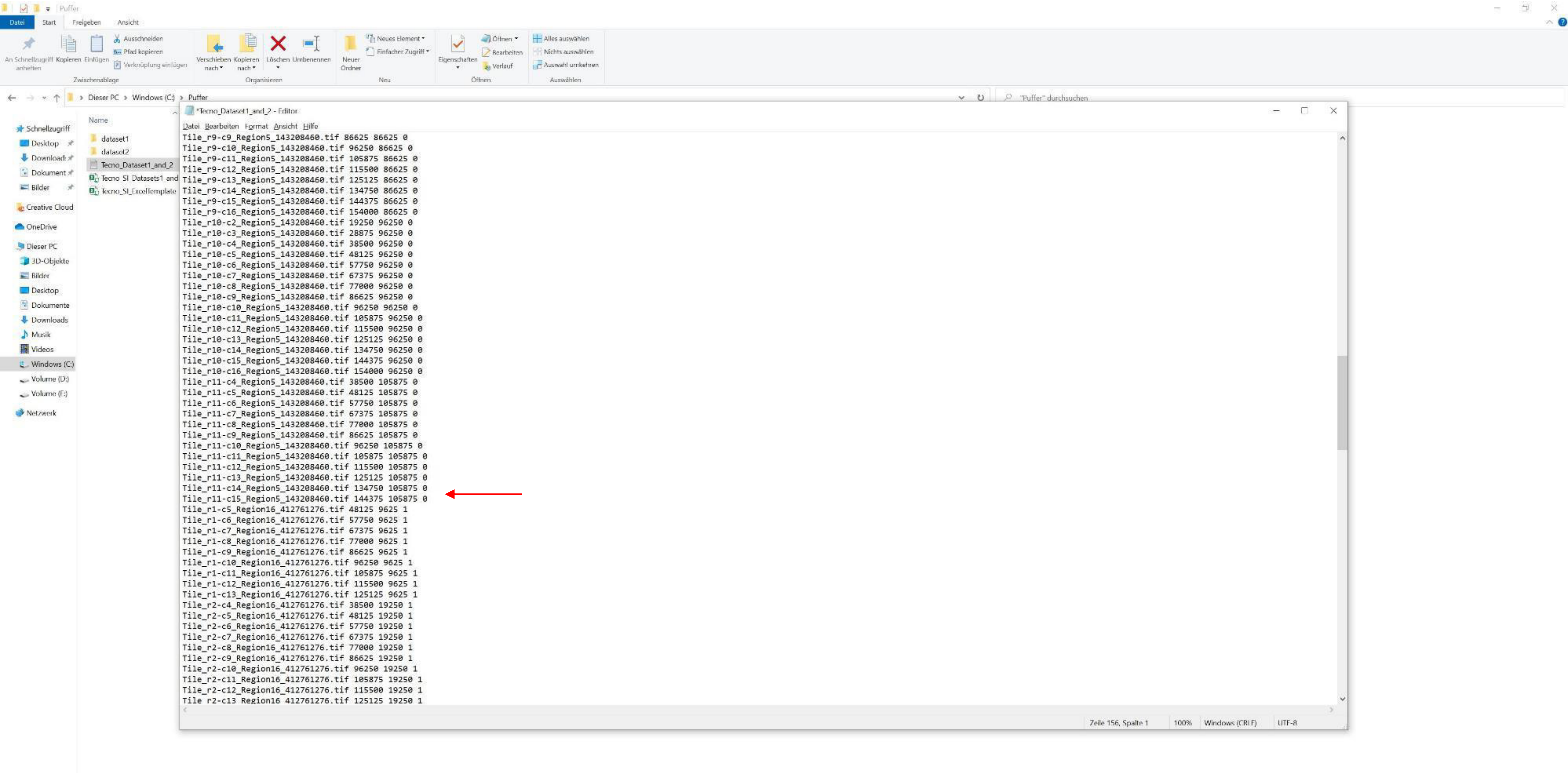


Copy all image tile coordinates into a separate text file

Delete the empty row between the coordinates of dataset 1 and dataset 2 (arrow; result)

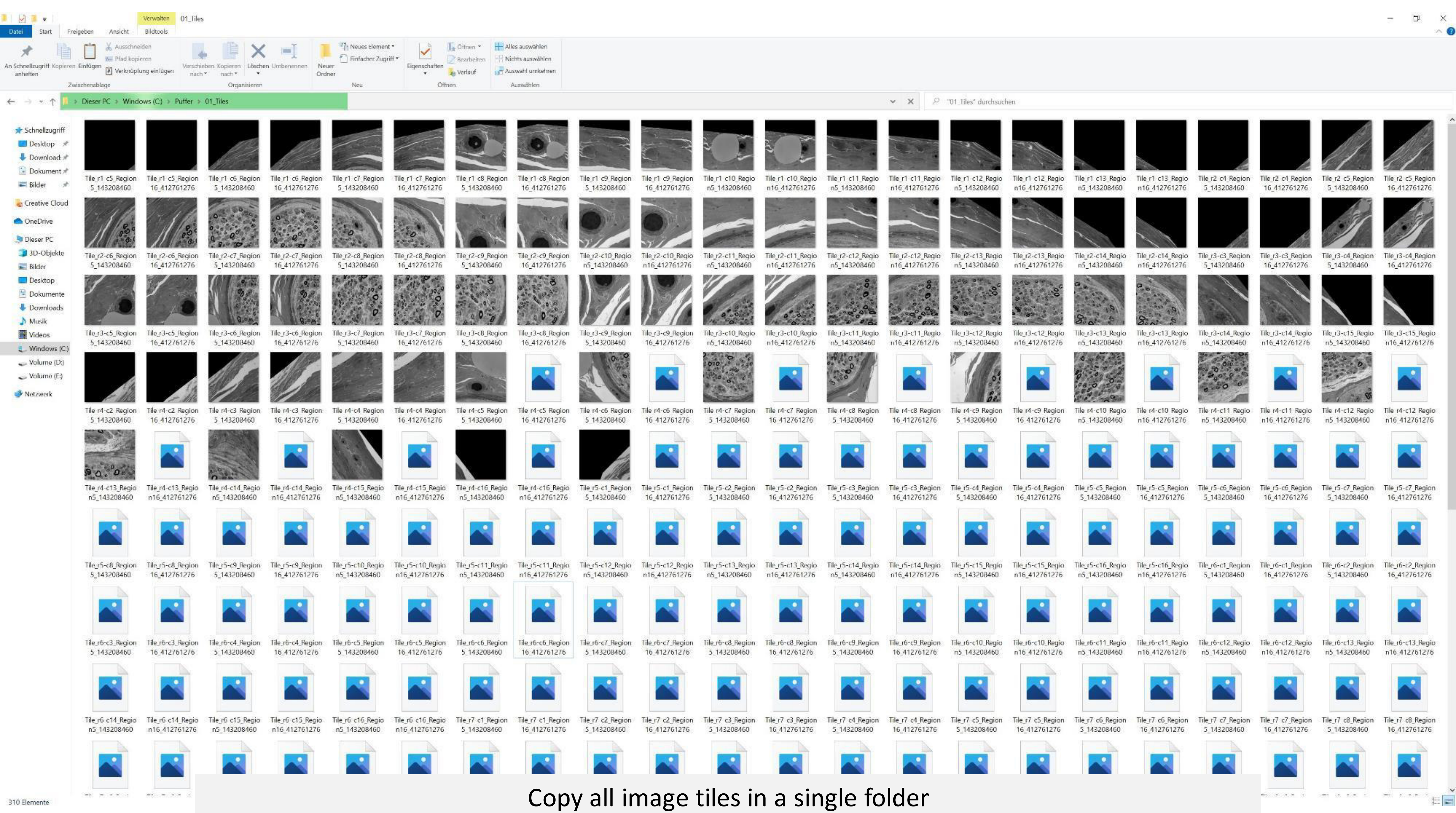


Copy all image tiles in a single folder

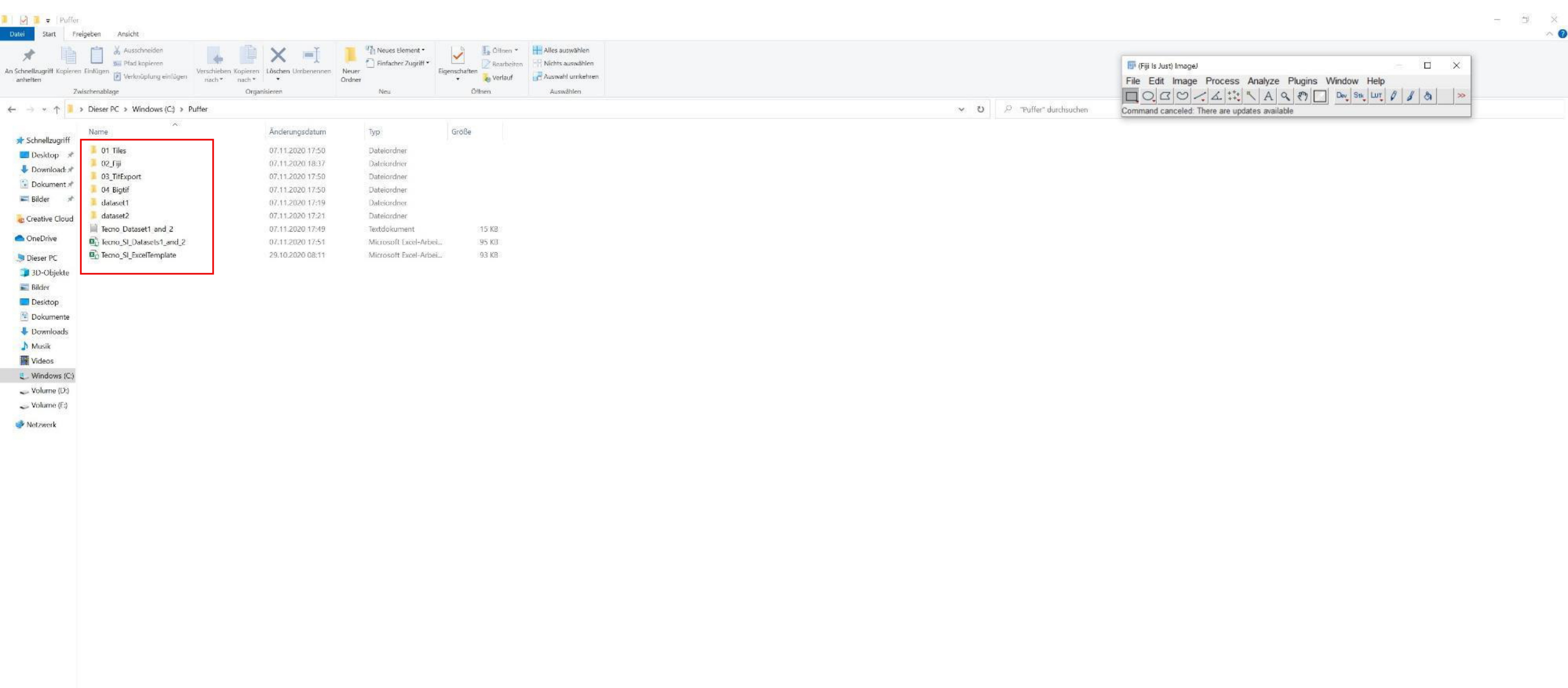


Prepare the following folders;
„01_Tiles“ with all image tiles of the different datasets (here dataset 1 and dataset 2)
„02_Fiji“ for the Fiji/ TrakEM2 project to be saved
„03_TifExport“ for the exported non-overlapping tif tiles (using TrakEM2)
„04_Bigtif“ for the export of the bigtif file (using nip2)

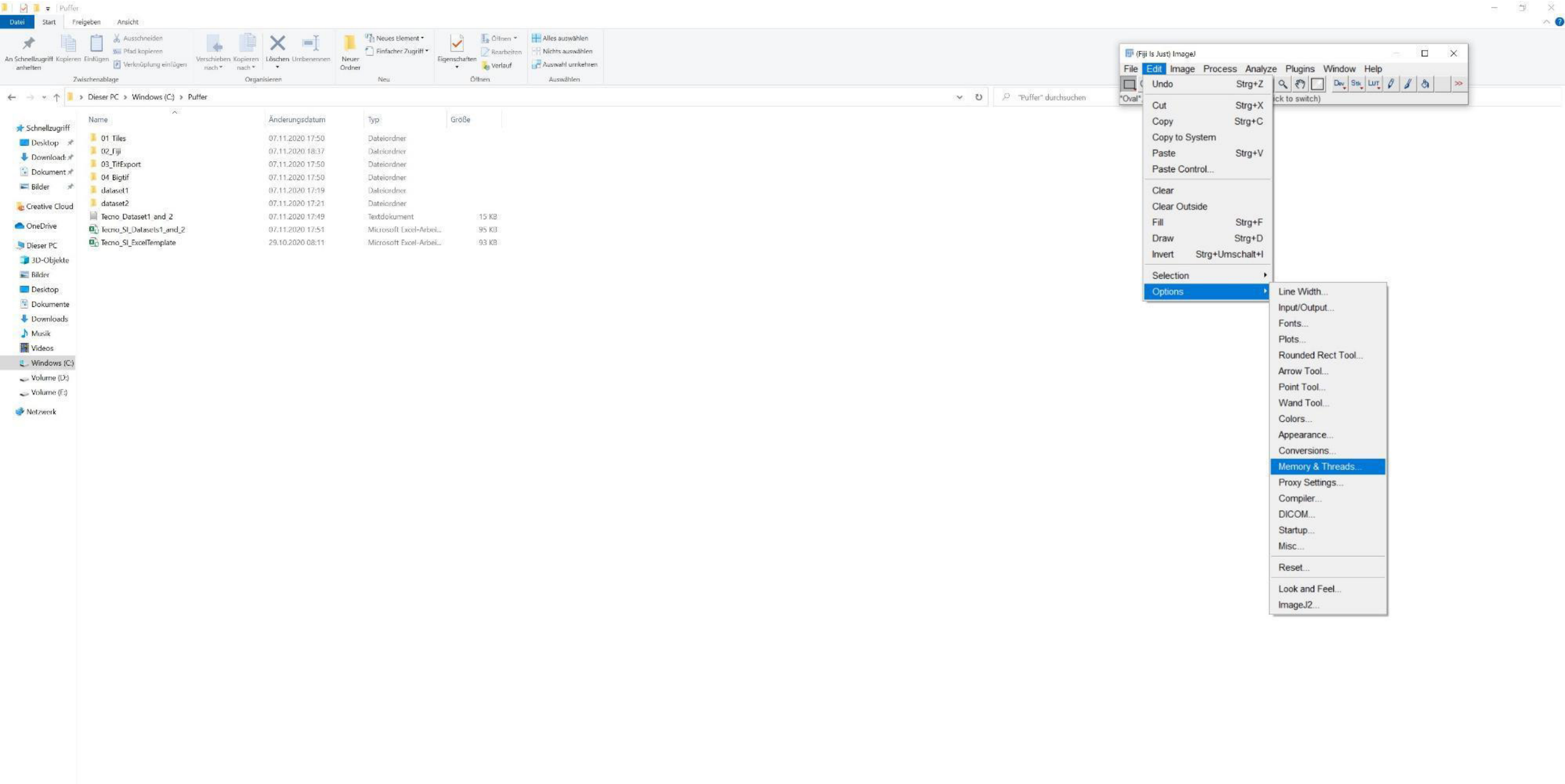


Open Fiji and adjust „Memory & Threads“ settings according to the computer parameters, restart afterwards

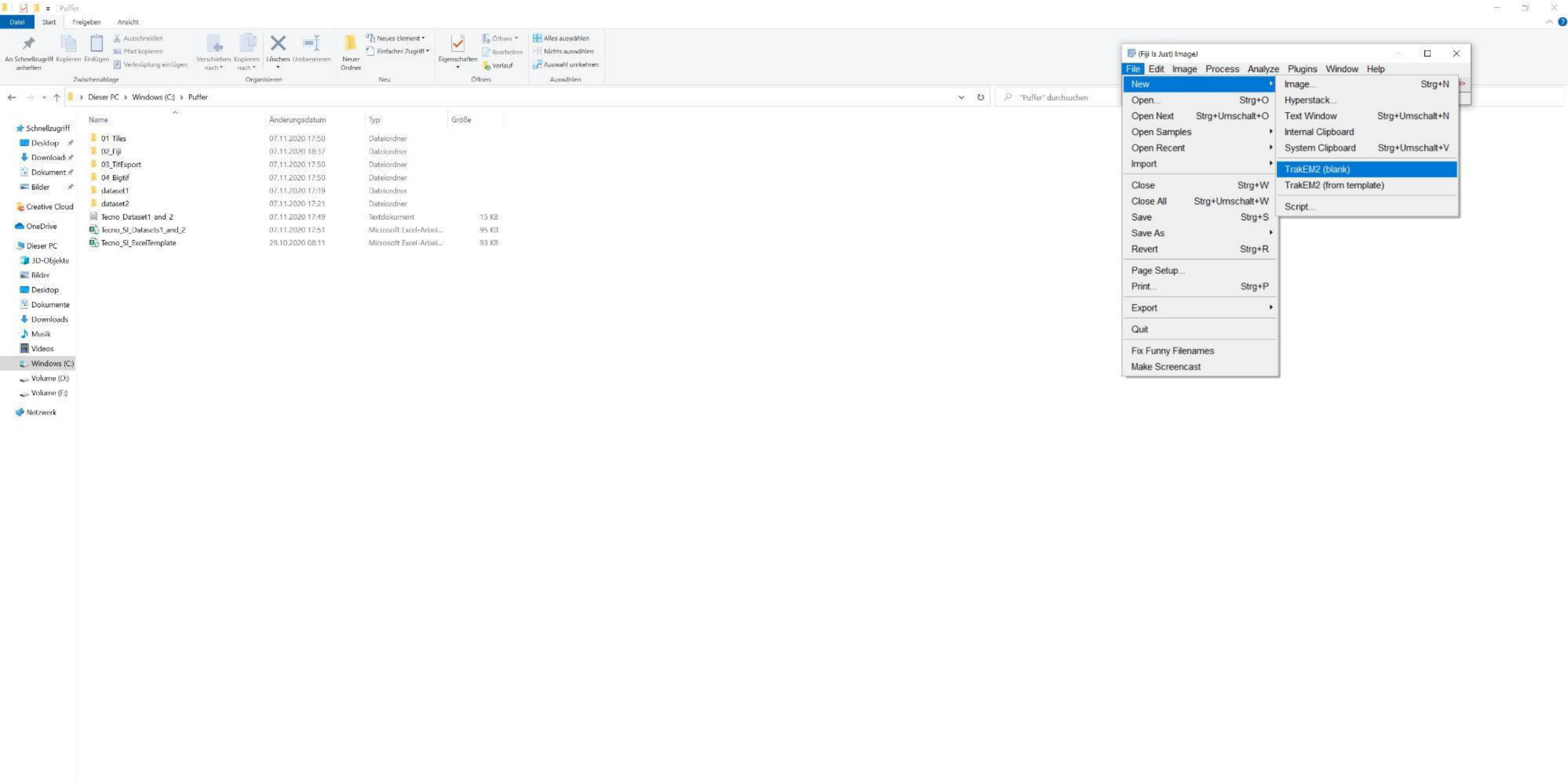


Open a new TrakEM2 project

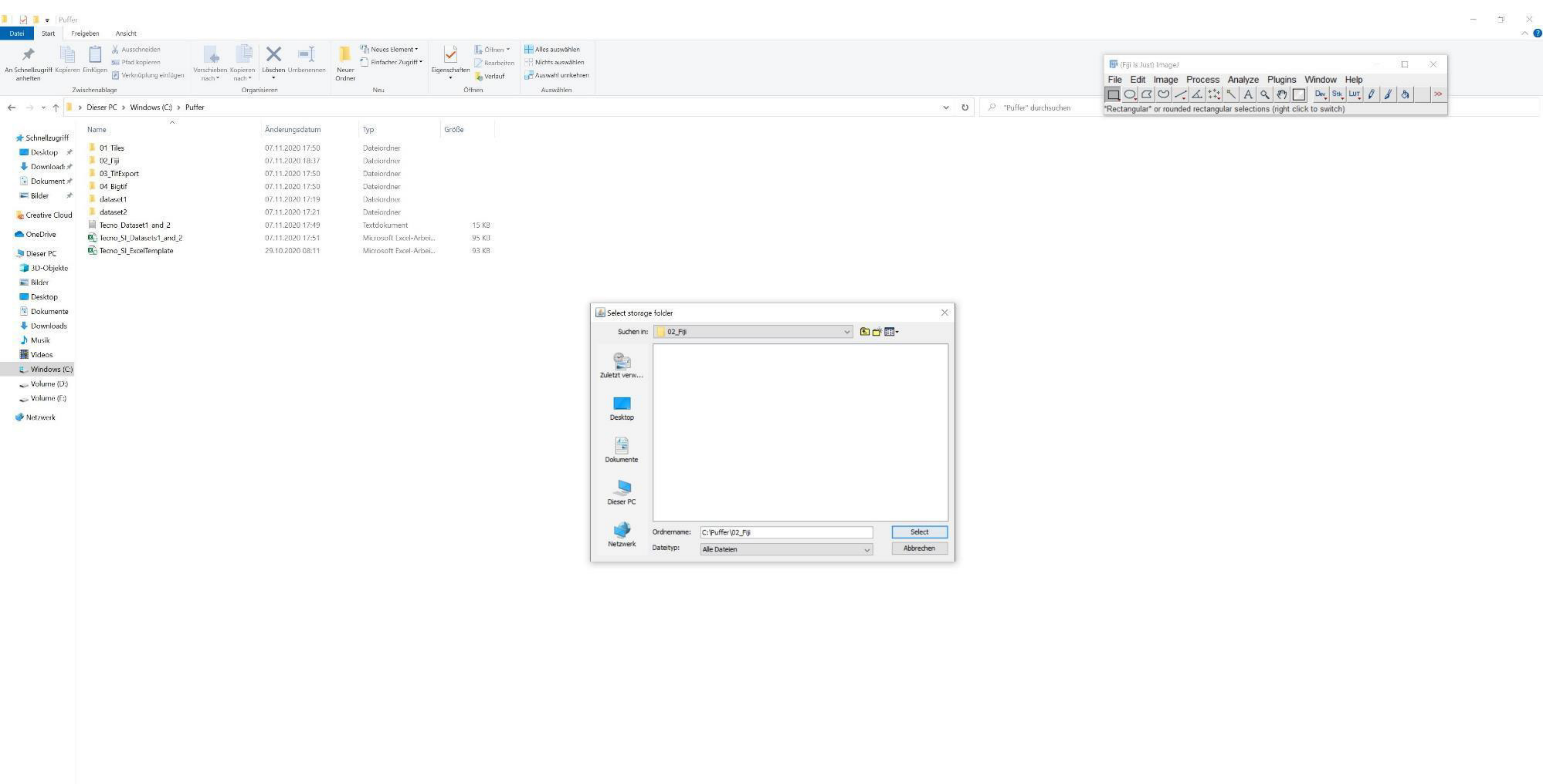


Choose „02_Fiji“ directory

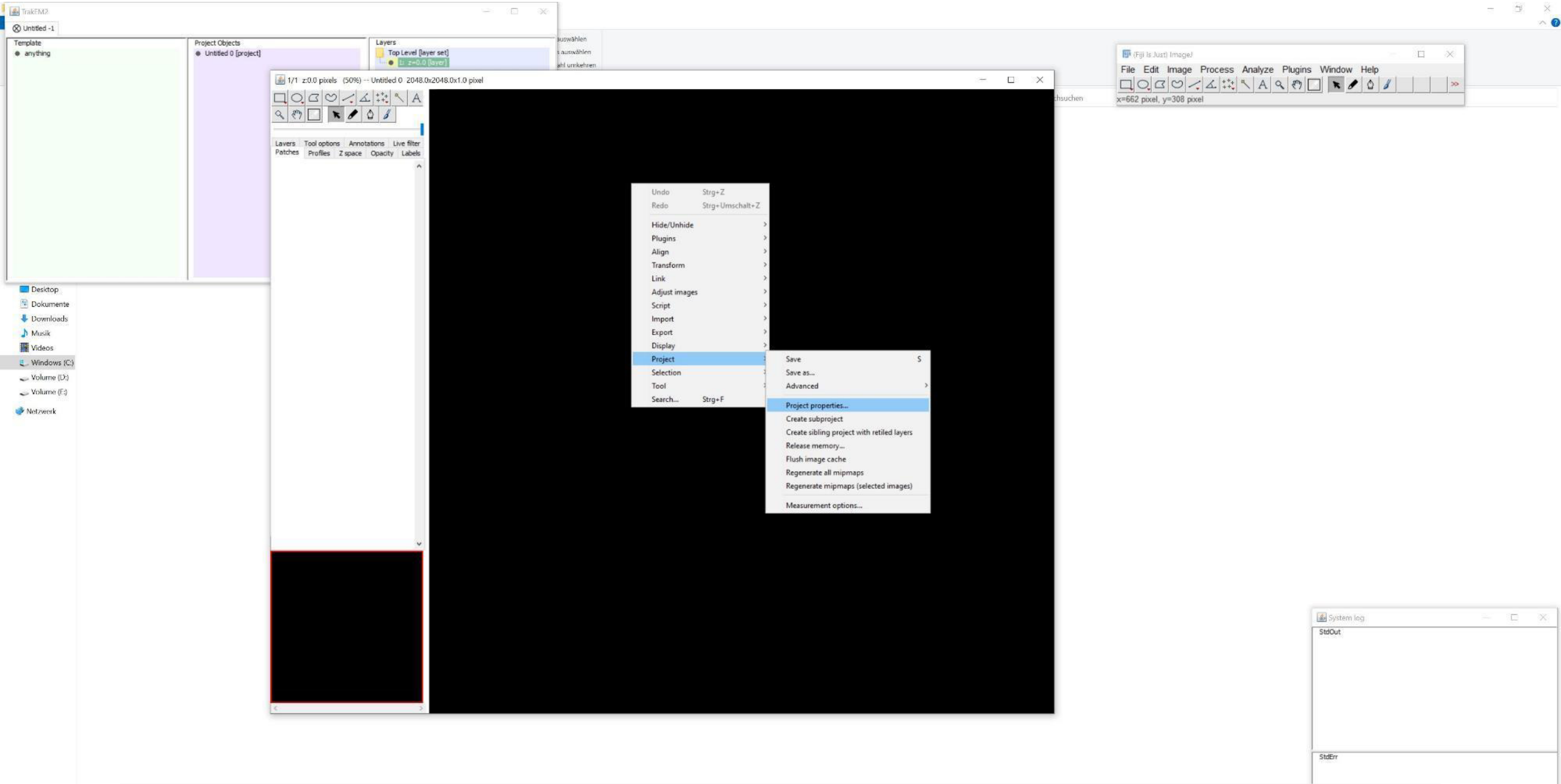


Right-click in the black canvas and adjust the „Project properties“

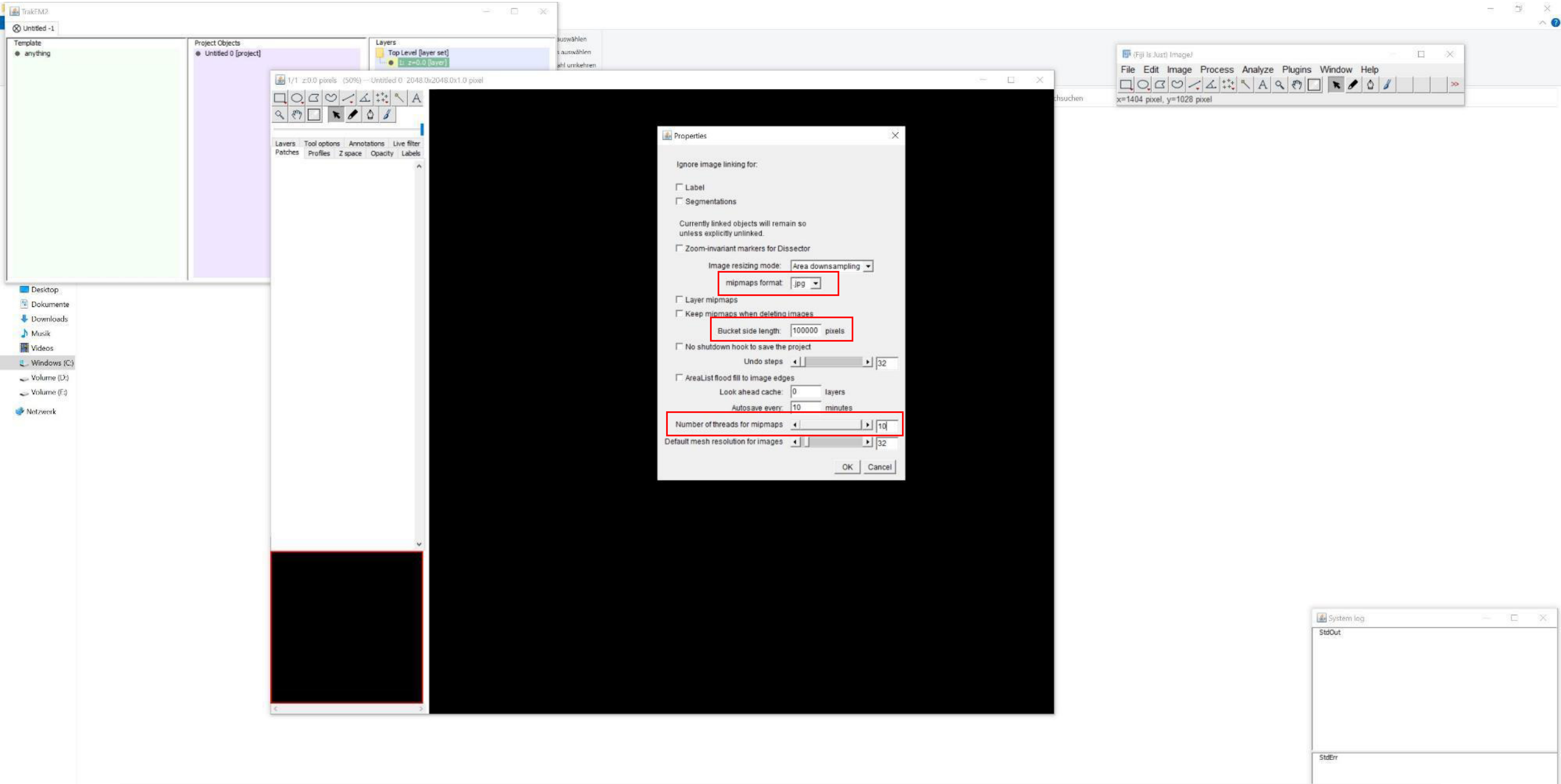


Change „mipmaps format“ to „.jpg“, the „Bucket side length“ to „100000“ and adjust the „Number of threads for mipmaps“

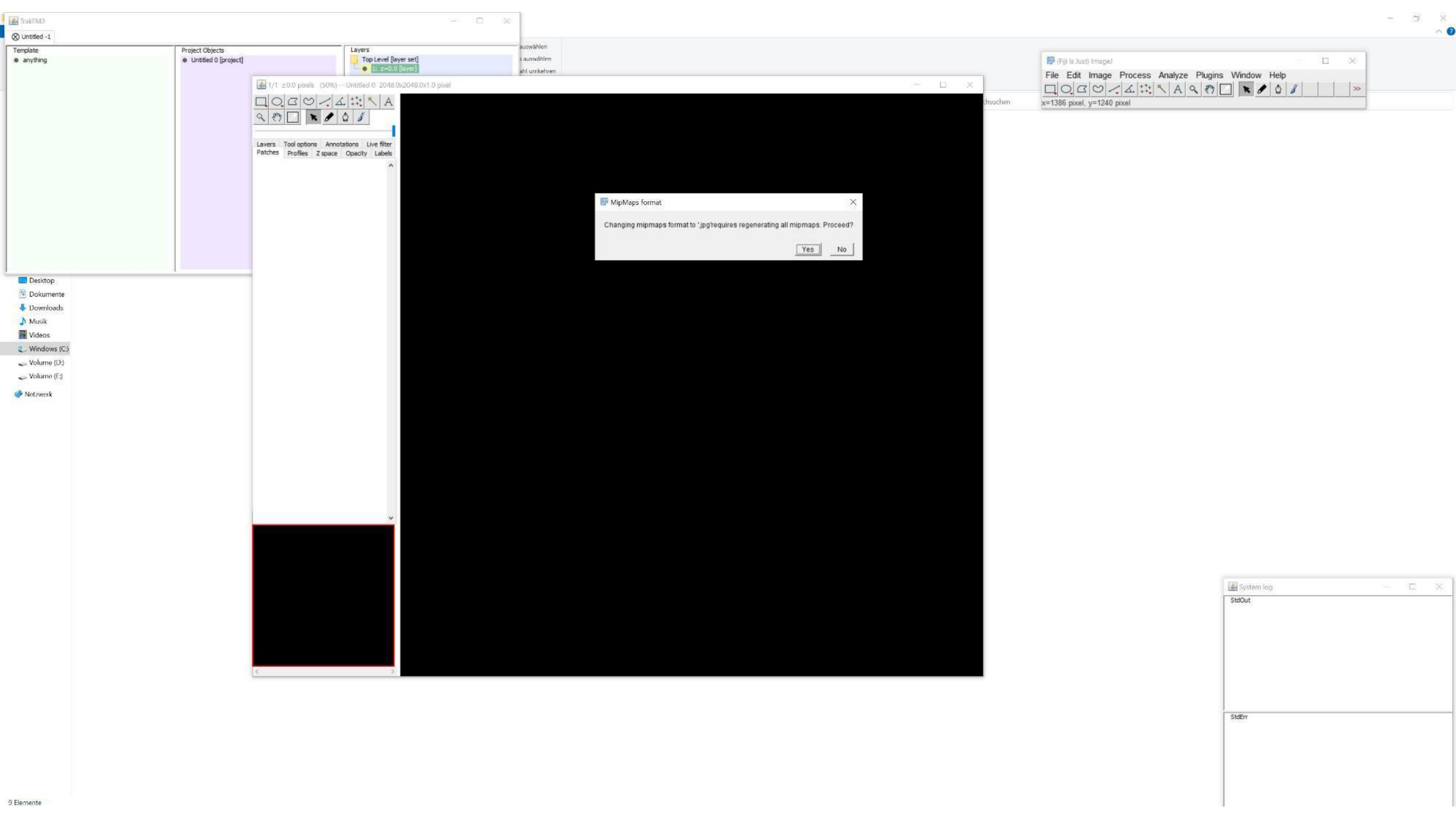
TrakEM2
Untitled -1
Template
anything
Project Objects
Untitled 0 [project]
Layers
Top Level [layer set]
1: z=0.0 [layer]
1/1 z:0.0 pixels (50%) — Untitled 0 2048.0x2048.0x1.0 pixel
Layers
Tool options
Annotations
Live filter
Patches
Profiles
Z space
Opacity
Labels
MipMaps format
Changing mipmaps format to '.jpg'requires regenerating all mipmaps. Proceed?
Yes
No
(Fiji Is Just) ImageJ
File
Edit
Image
Process
Analyze
Plugins
Window
Help
x=1386 pixel, y=1240 pixel
Desktop
Dokumente
Downloads
Musik
Videos
Windows (C:)
Volume (D:)
Volume (F:)
Netzwerk
9 Elemente
System log
StdOut
StdErr

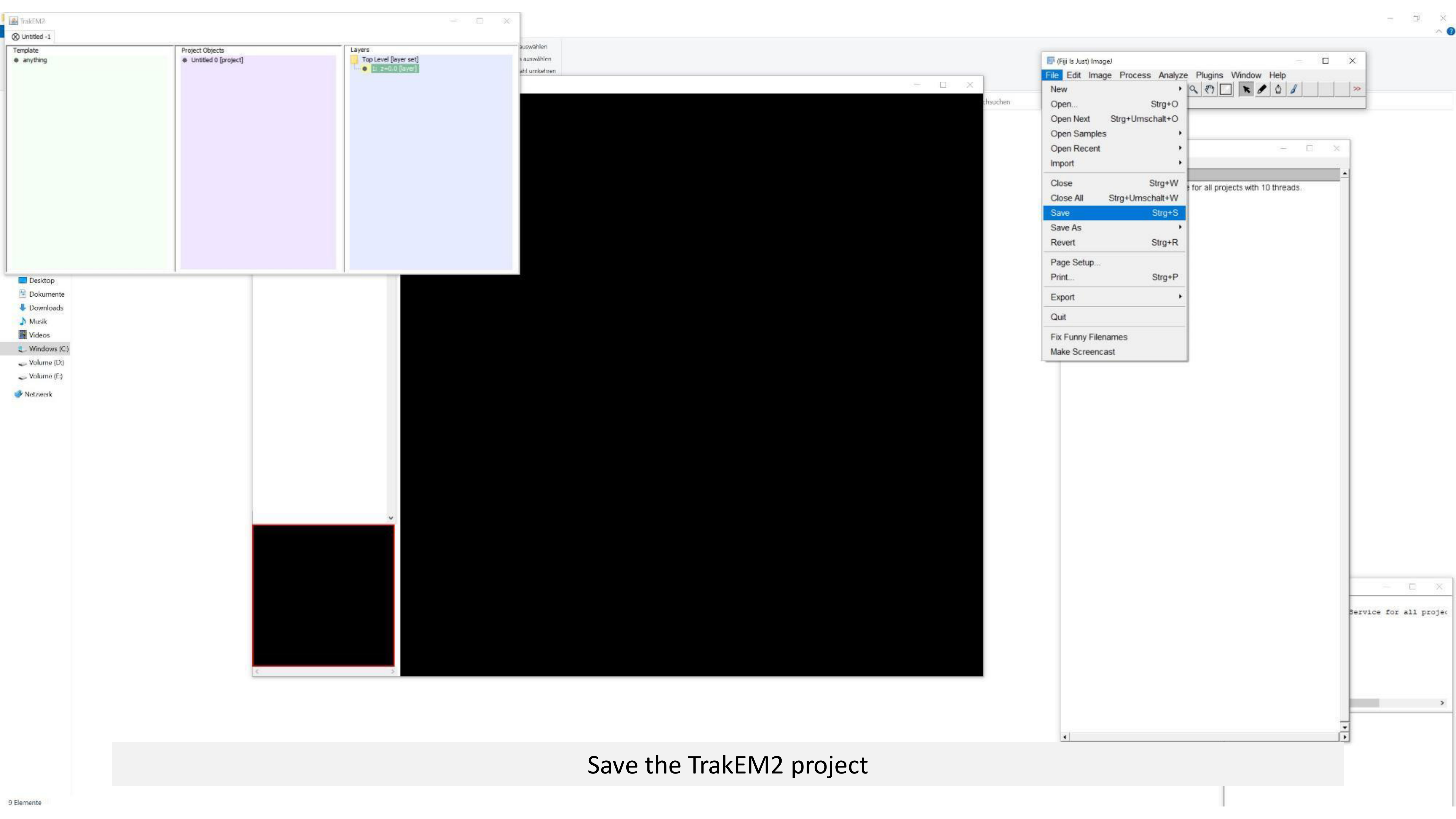


Save the TrakEM2 project

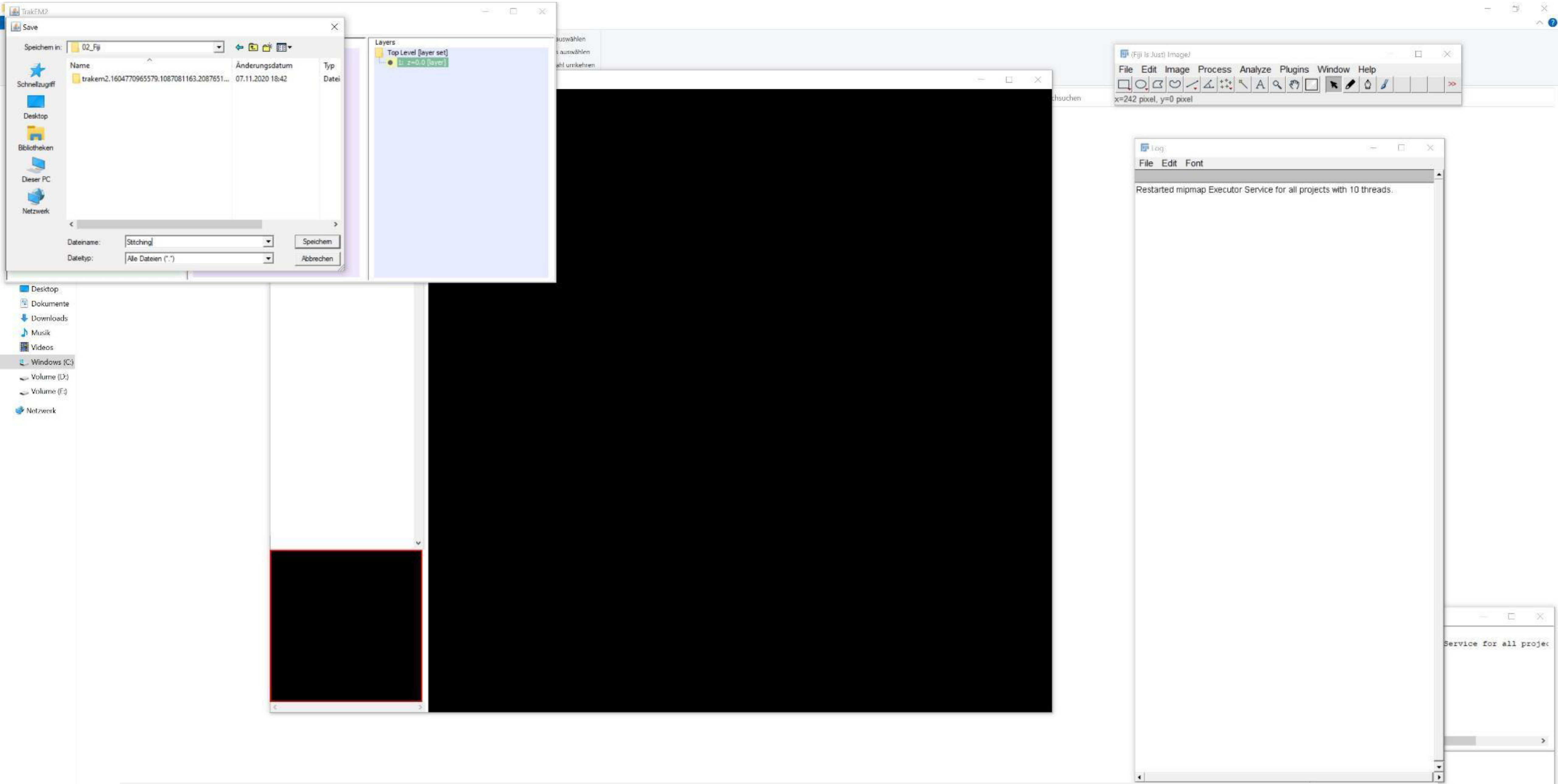


A separate file (.xml.gz) will be generated below the „trakem2. …“ folder
(The TrakEM2 project can be opened in a new session via drag&drop of this xml file on the small Fiji menu)

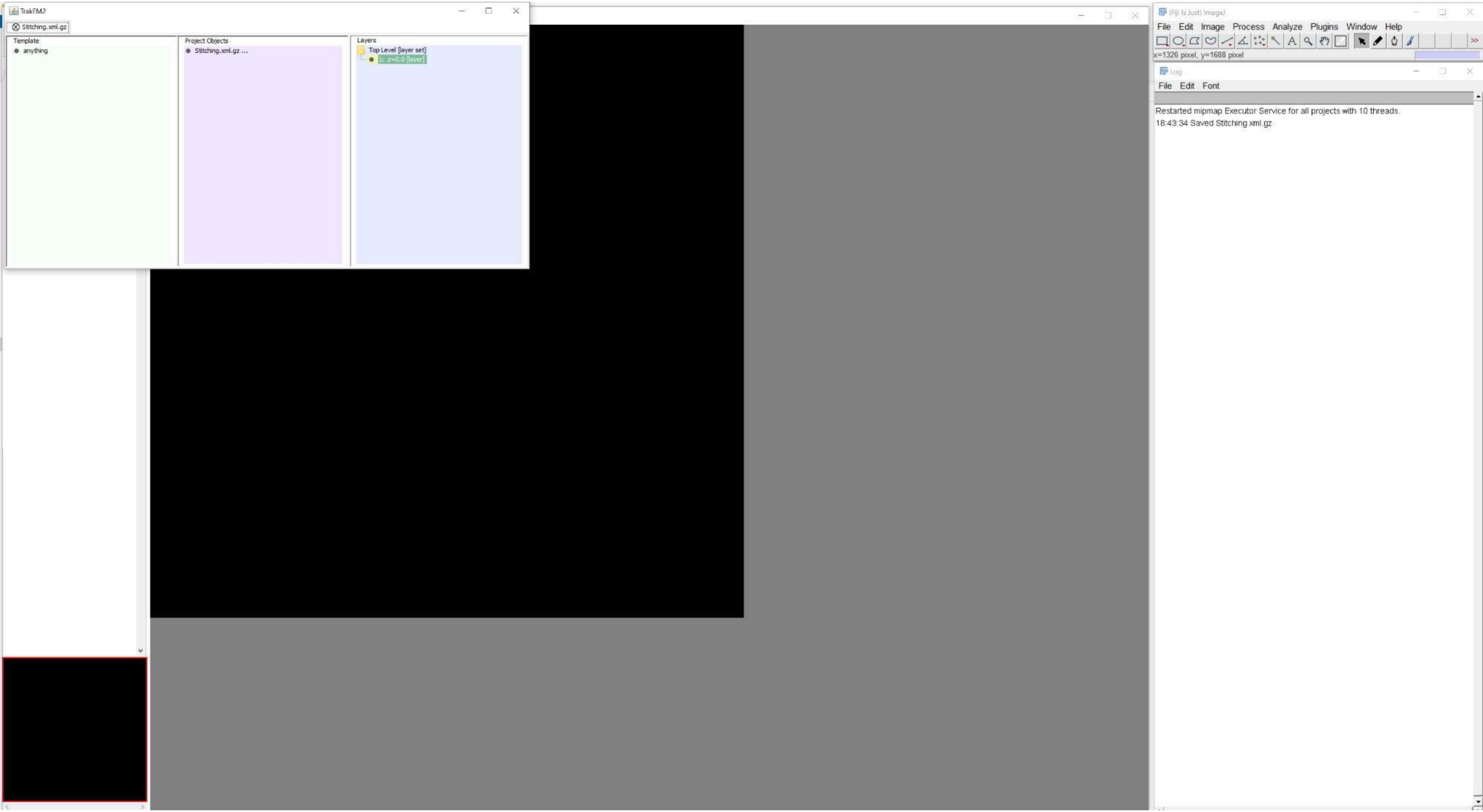
TrakEM2
Stitching.xml.gz
Template
anything
Project Objects
Stitching.xml.gz ...
Layers
Top Level [layer set]
1: z=0.0 [layer]
(Fiji Is Just) ImageJ
File Edit Image Process Analyze Plugins Window Help
x=1326 pixel, y=1688 pixel
Log
File Edit Font
Restarted mipmap Executor Service for all projects with 10 threads.
18:43:34 Saved Stitching.xml.gz

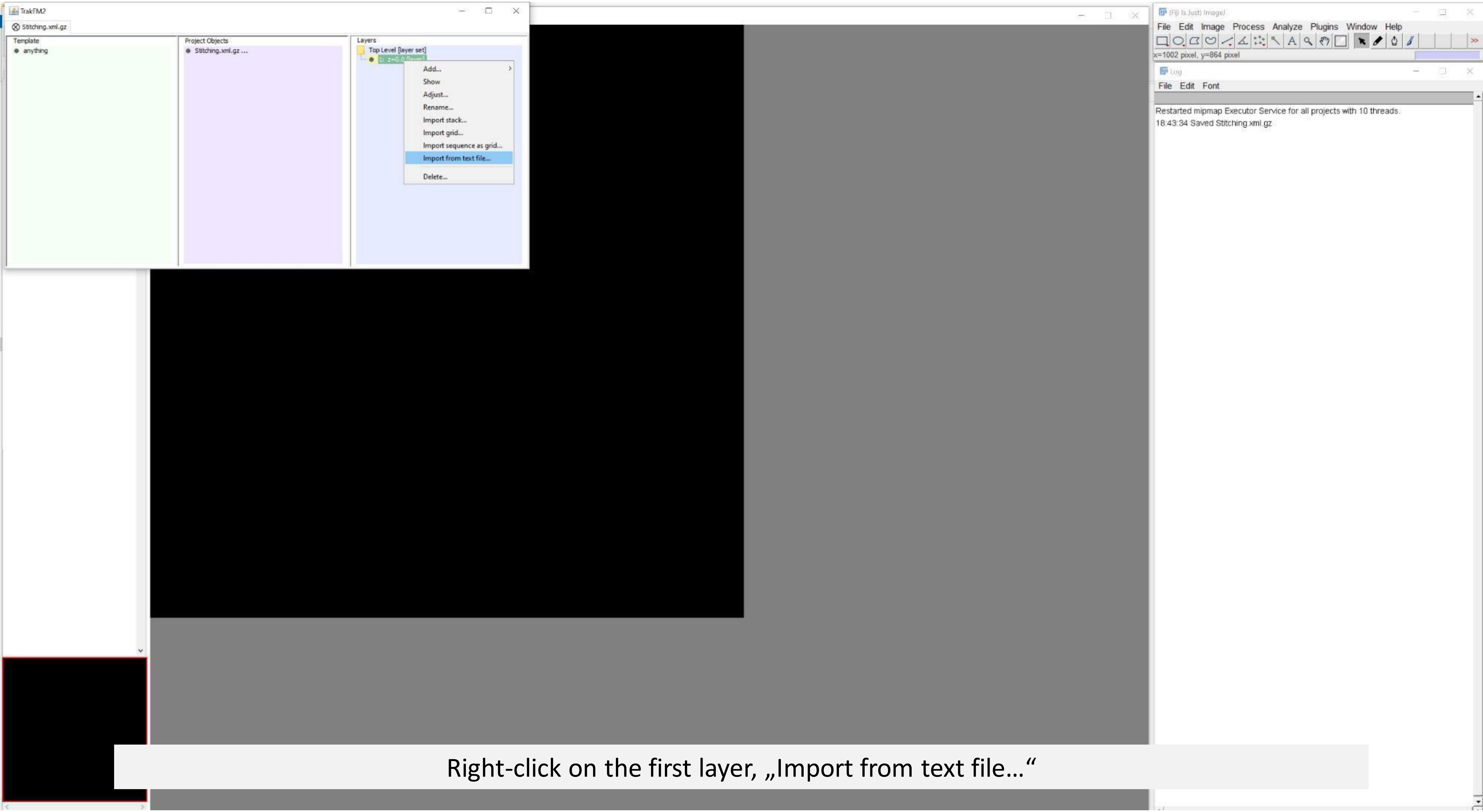


Right-click on the first layer, „Import from text file…“

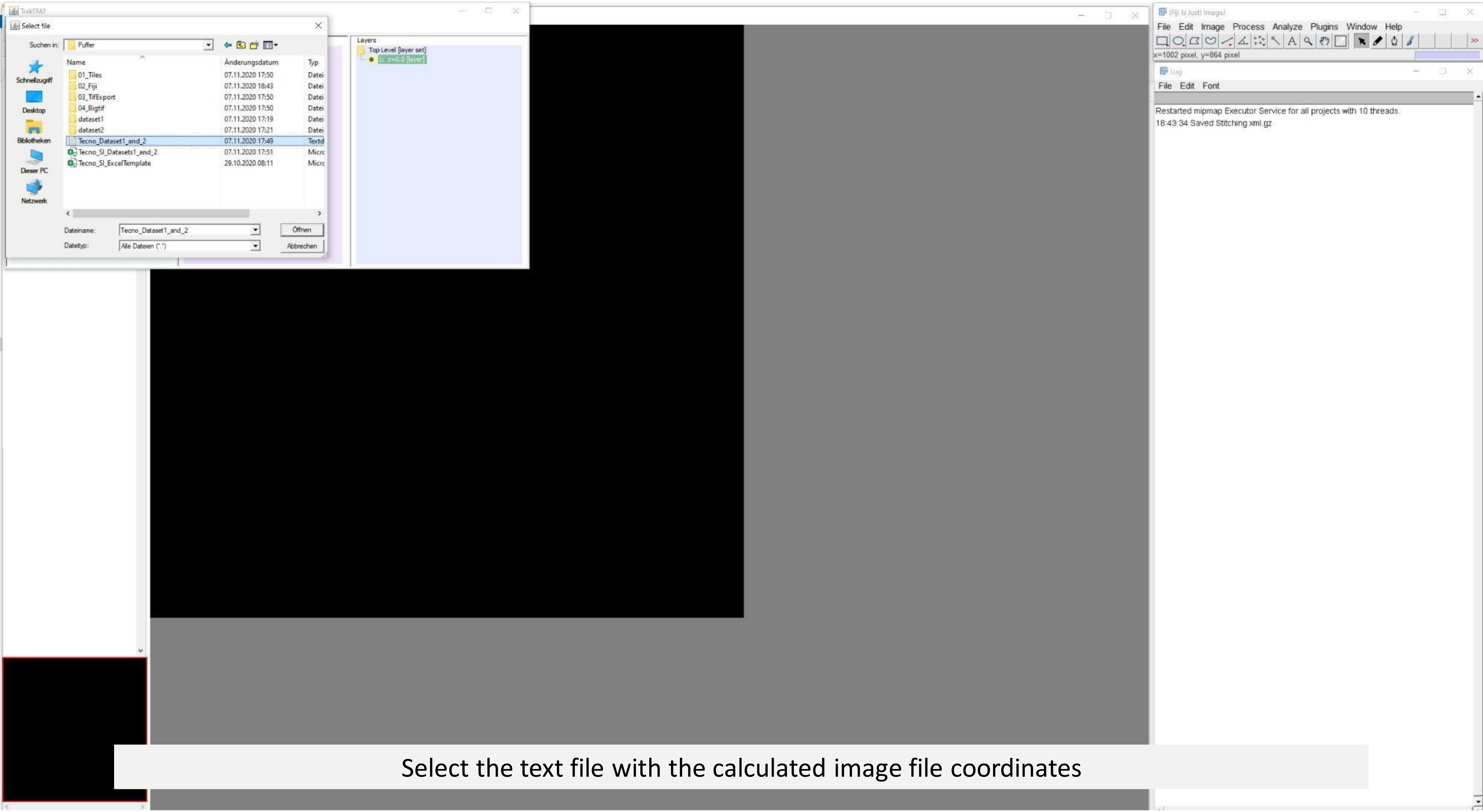


Select the text file with the calculated image file coordinates

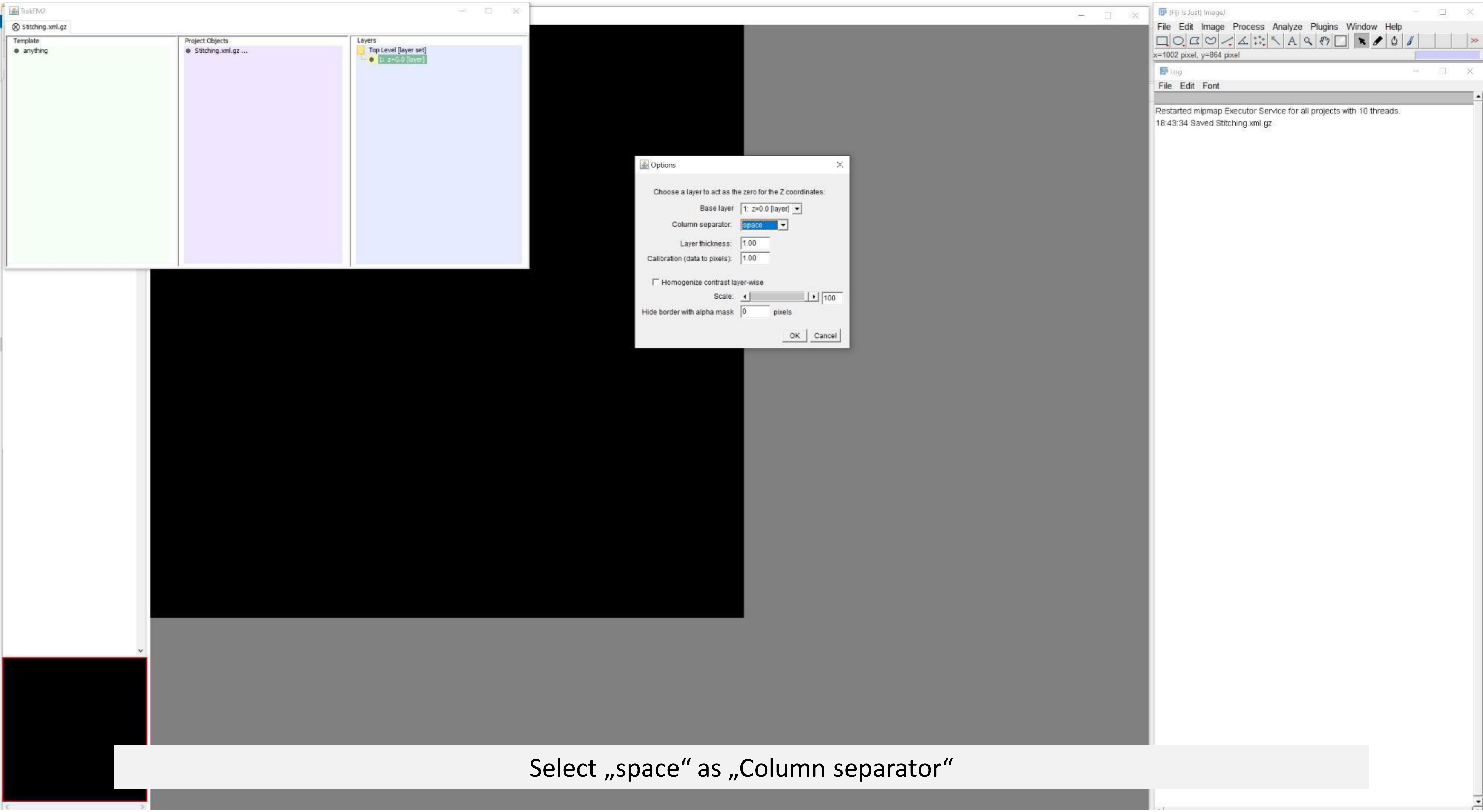


Select „space“ as „Column separator“

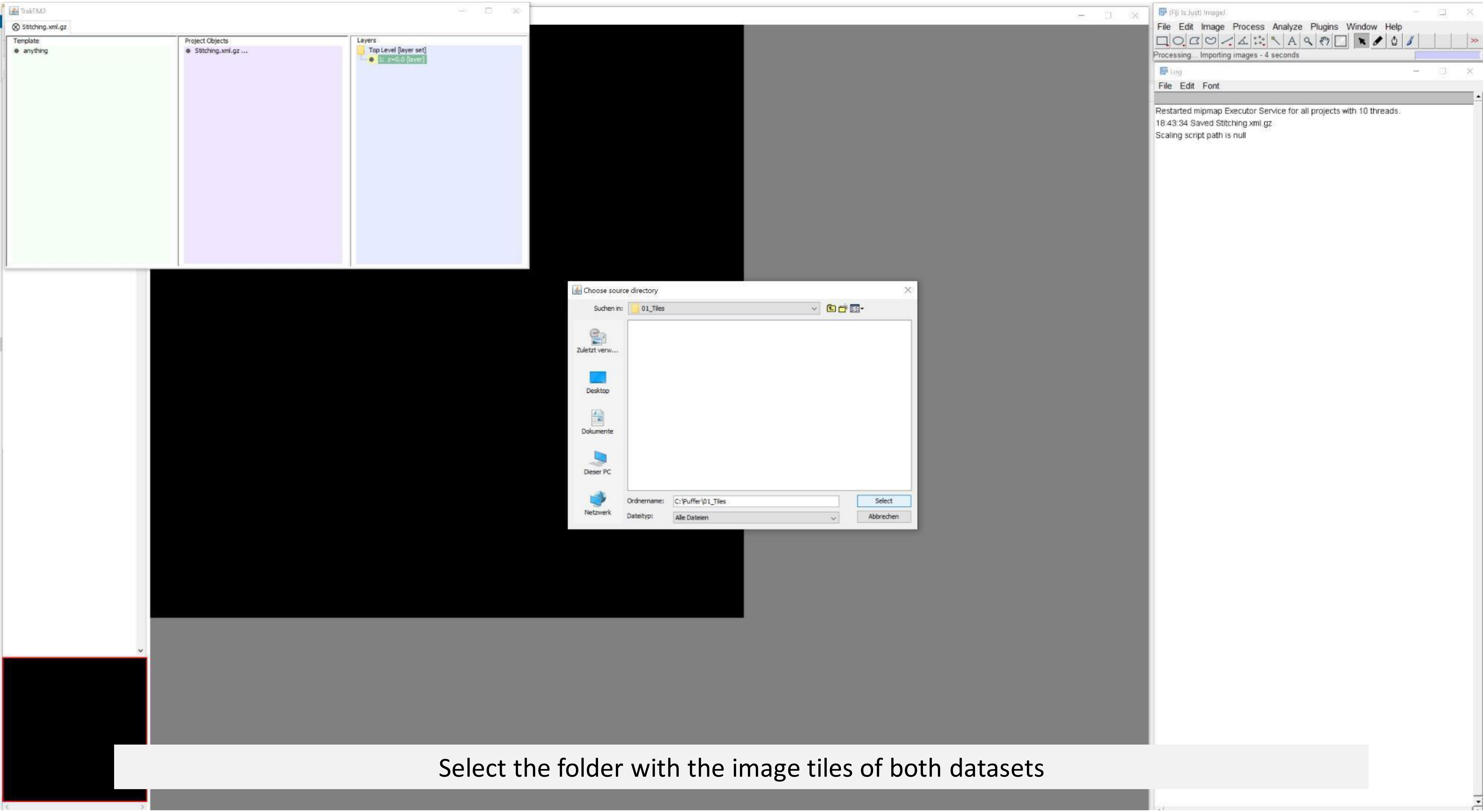


Select the folder with the image tiles of both datasets

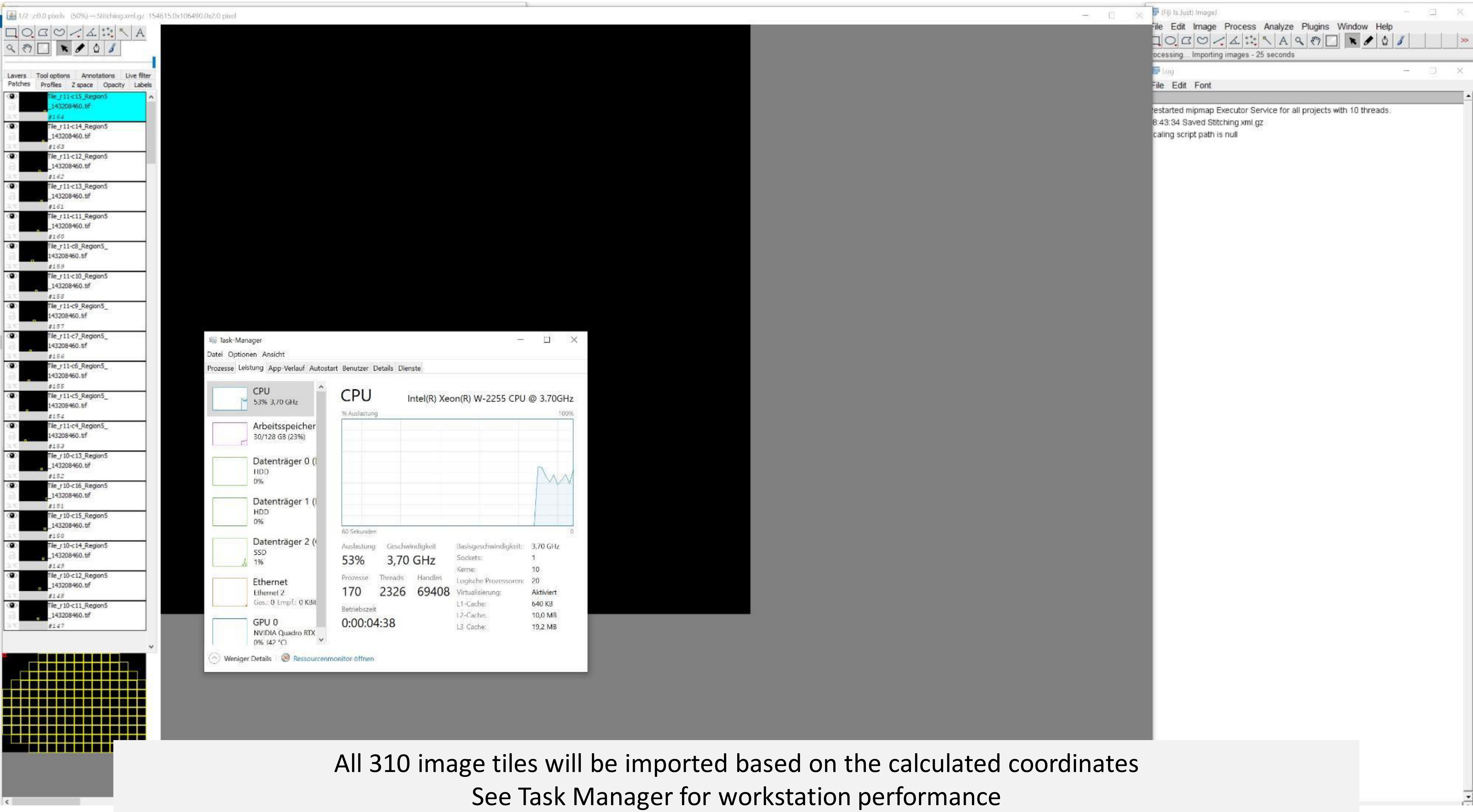


All 310 image tiles will be imported based on the calculated coordinates
See Task Manager for workstation performance

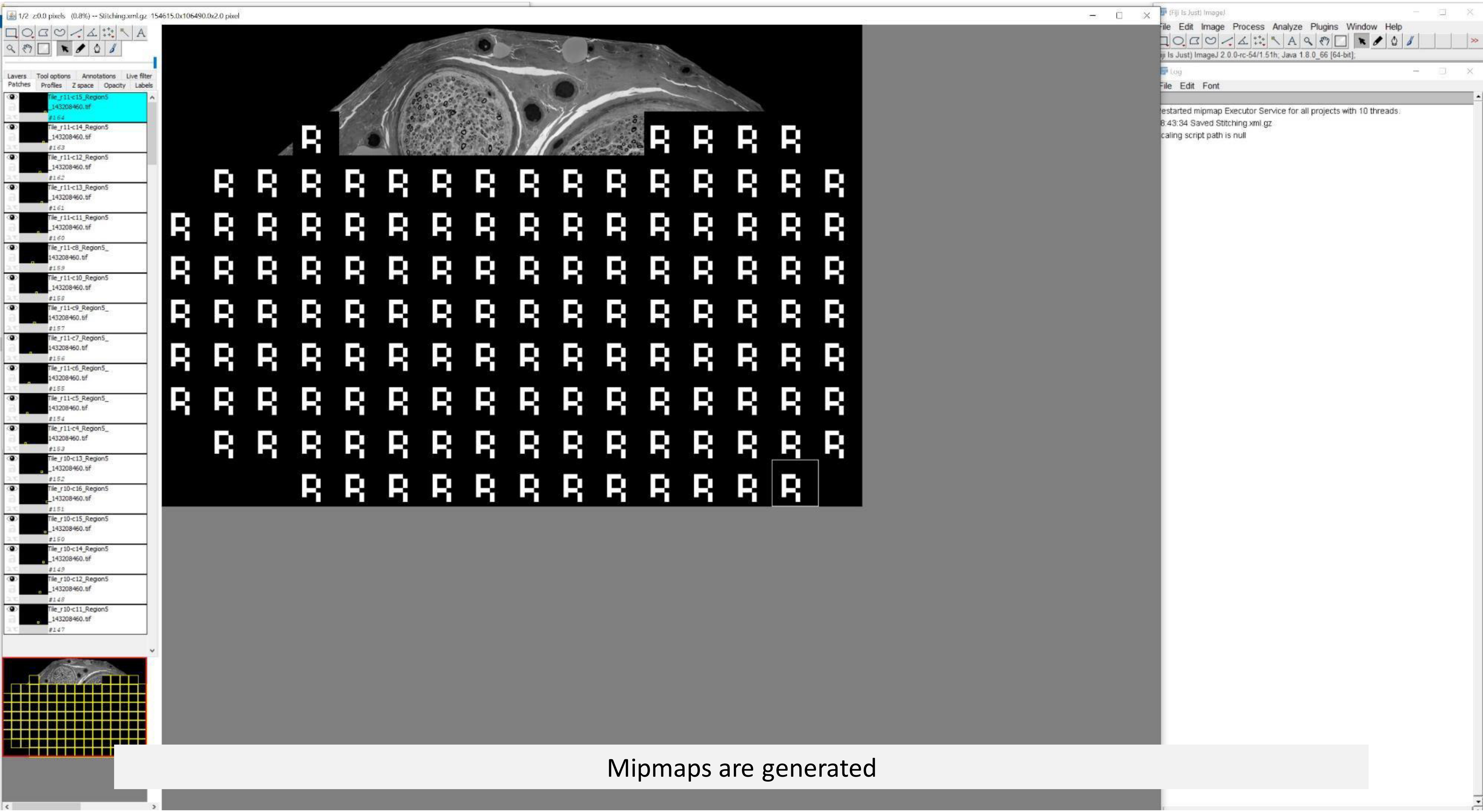


Mipmaps are generated

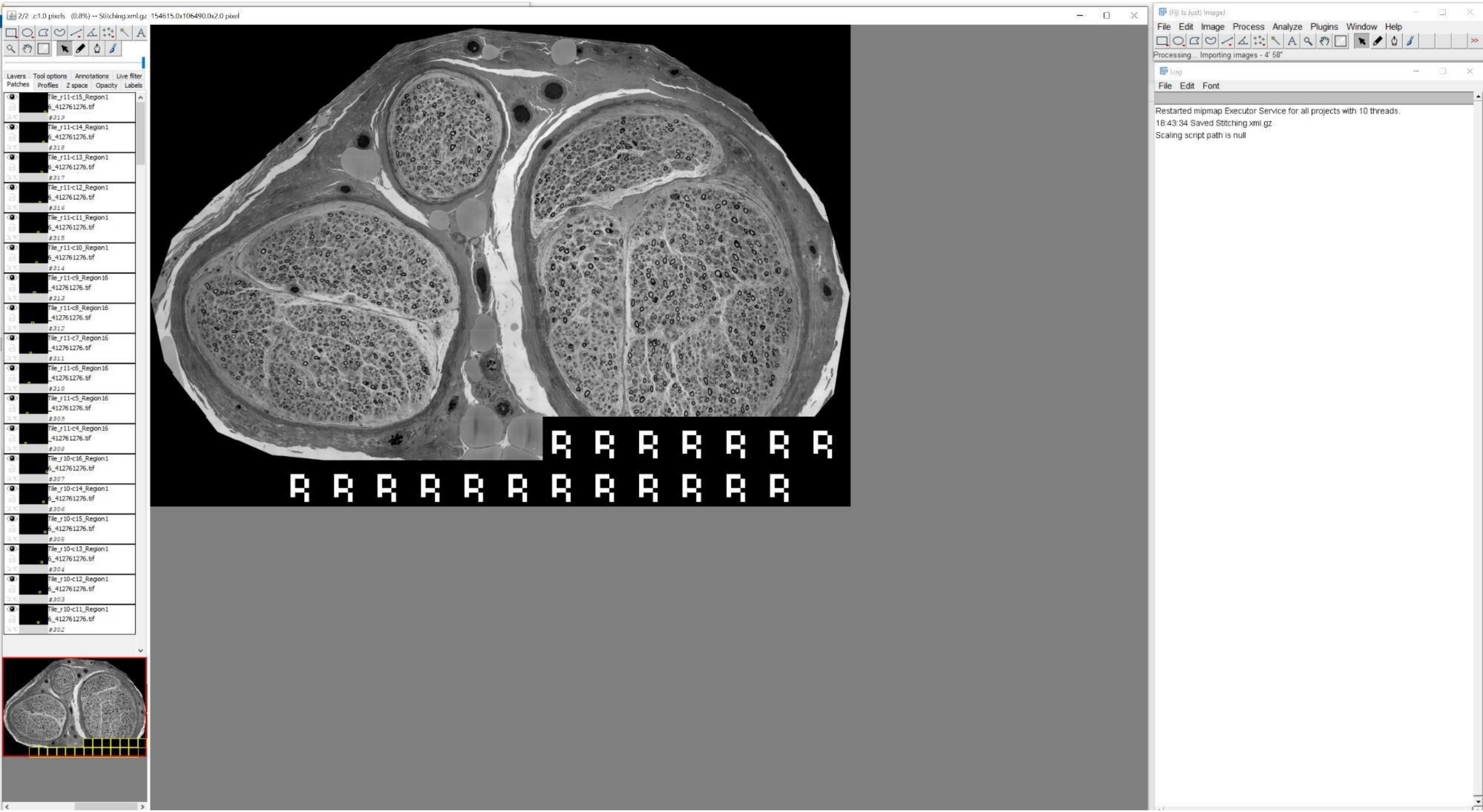
2/2 z:1.0 pixels (0.8%) -- Stitching.xml.gz 154615.0x106490.0x2.0 pixel
Layers Tool options Annotations Live filter
Patches Profiles Z space Opacity Labels
Tile_r11-c15_Region16_412761276.tif #319
Tile_r11-c14_Region16_412761276.tif #318
Tile_r11-c13_Region16_412761276.tif #317
Tile_r11-c12_Region16_412761276.tif #316
Tile_r11-c11_Region16_412761276.tif #315
Tile_r11-c10_Region16_412761276.tif #314
Tile_r11-c9_Region16_412761276.tif #313
Tile_r11-c8_Region16_412761276.tif #312
Tile_r11-c7_Region16_412761276.tif #311
Tile_r11-c6_Region16_412761276.tif #310
Tile_r11-c5_Region16_412761276.tif #309
Tile_r11-c4_Region16_412761276.tif #308
Tile_r10-c16_Region16_412761276.tif #307
Tile_r10-c14_Region16_412761276.tif #306
Tile_r10-c15_Region16_412761276.tif #305
Tile_r10-c13_Region16_412761276.tif #304
Tile_r10-c12_Region16_412761276.tif #303
Tile_r10-c11_Region16_412761276.tif #302
(Fiji Is Just) ImageJ
File Edit Image Process Analyze Plugins Window Help
Processing... Importing images - 4' 58"
Log
File Edit Font
Restarted mipmap Executor Service for all projects with 10 threads.
18:43:34 Saved Stitching.xml.gz
Scaling script path is null

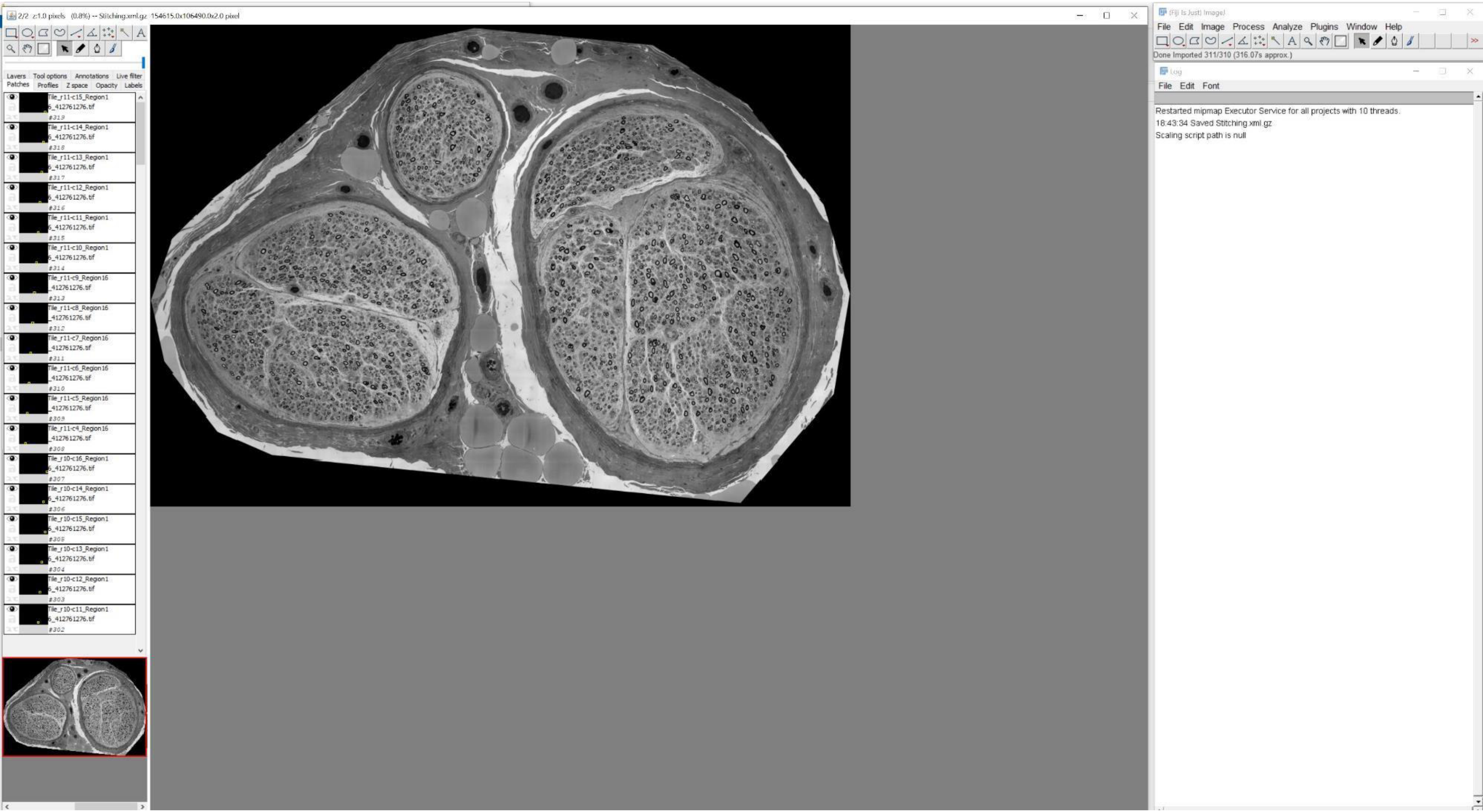
2/2 z:1.0 pixels (0.8%) -- Stitching.xml.gz 154615.0x106490.0x2.0 pixel
Layers Tool options Annotations Live filter
Patches Profiles Z space Opacity Labels
Tile_r11-c15_Region16_412761276.tif
#319
Tile_r11-c14_Region16_412761276.tif
#318
Tile_r11-c13_Region16_412761276.tif
#317
Tile_r11-c12_Region16_412761276.tif
#316
Tile_r11-c11_Region16_412761276.tif
#315
Tile_r11-c10_Region16_412761276.tif
#314
Tile_r11-c9_Region16_412761276.tif
#313
Tile_r11-c8_Region16_412761276.tif
#312
Tile_r11-c7_Region16_412761276.tif
#311
Tile_r11-c6_Region16_412761276.tif
#310
Tile_r11-c5_Region16_412761276.tif
#309
Tile_r11-c4_Region16_412761276.tif
#308
Tile_r10-c16_Region16_412761276.tif
#307
Tile_r10-c14_Region16_412761276.tif
#306
Tile_r10-c15_Region16_412761276.tif
#305
Tile_r10-c13_Region16_412761276.tif
#304
Tile_r10-c12_Region16_412761276.tif
#303
Tile_r10-c11_Region16_412761276.tif
#302
(Fiji Is Just) ImageJ
File Edit Image Process Analyze Plugins Window Help
Done Imported 311/310 (316.07s approx.)
Log
File Edit Font
Restarted mipmap Executor Service for all projects with 10 threads.
18:43:34 Saved Stitching.xml.gz
Scaling script path is null

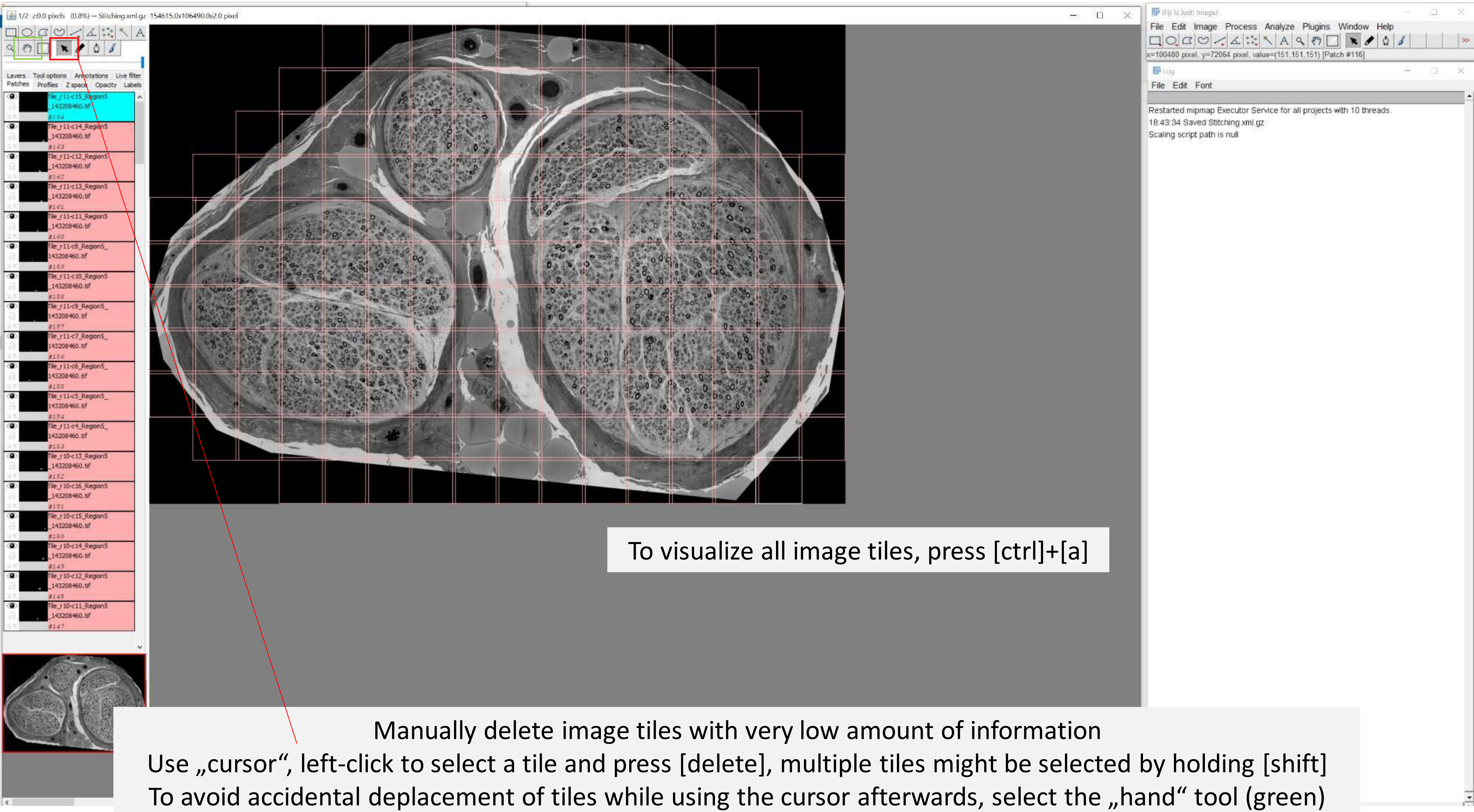

To visualize all image tiles, press [ctrl]+[a]

Manually delete image tiles with very low amount of information
Use „cursor“, left-click to select a tile and press [delete], multiple tiles might be selected by holding [shift]
To avoid accidental deplacement of tiles while using the cursor afterwards, select the „hand“ tool (green)

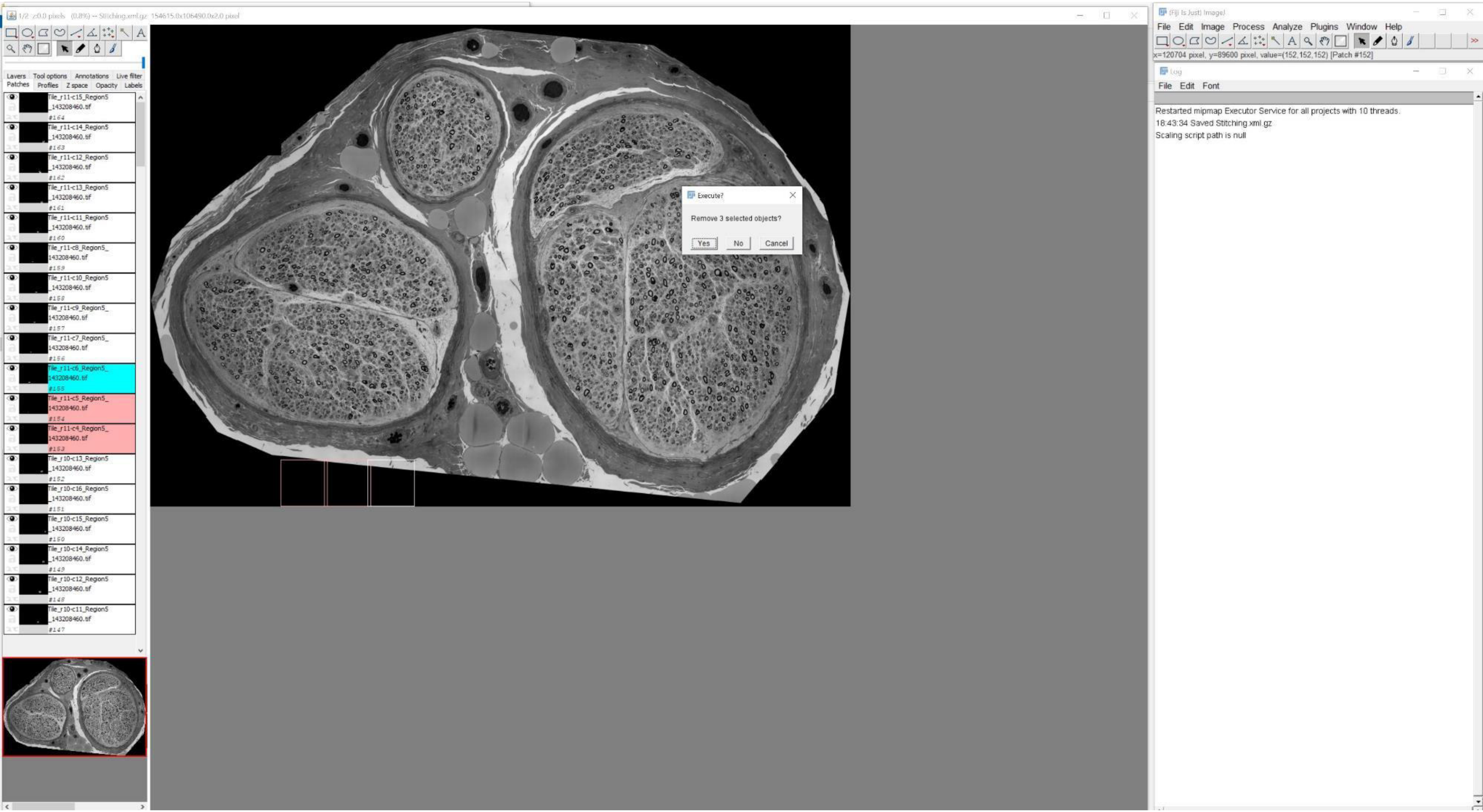

1/2 z:0.0 pixels (0.8%) -- Stitching.xml.gz 154615.0x106490.0x2.0 pixel
Layers Tool options Annotations Live filter
Patches Profiles Z space Opacity Labels
Tile_r11-c15_Region5_143208460.tif #164
Tile_r11-c14_Region5_143208460.tif #163
Tile_r11-c12_Region5_143208460.tif #162
Tile_r11-c13_Region5_143208460.tif #161
Tile_r11-c11_Region5_143208460.tif #160
Tile_r11-c8_Region5_143208460.tif #159
Tile_r11-c10_Region5_143208460.tif #158
Tile_r11-c9_Region5_143208460.tif #157
Tile_r11-c7_Region5_143208460.tif #156
Tile_r11-c6_Region5_143208460.tif #155
Tile_r11-c5_Region5_143208460.tif #154
Tile_r11-c4_Region5_143208460.tif #153
Tile_r10-c13_Region5_143208460.tif #152
Tile_r10-c16_Region5_143208460.tif #151
Tile_r10-c15_Region5_143208460.tif #150
Tile_r10-c14_Region5_143208460.tif #149
Tile_r10-c12_Region5_143208460.tif #148
Tile_r10-c11_Region5_143208460.tif #147
Execute?
Remove 3 selected objects?
Yes
No
Cancel
(Fiji Is Just) ImageJ
File Edit Image Process Analyze Plugins Window Help
x=120704 pixel, y=89600 pixel, value=(152,152,152) [Patch #152]
Log
File Edit Font
Restarted mipmap Executor Service for all projects with 10 threads.
18:43:34 Saved Stitching.xml.gz
Scaling script path is null

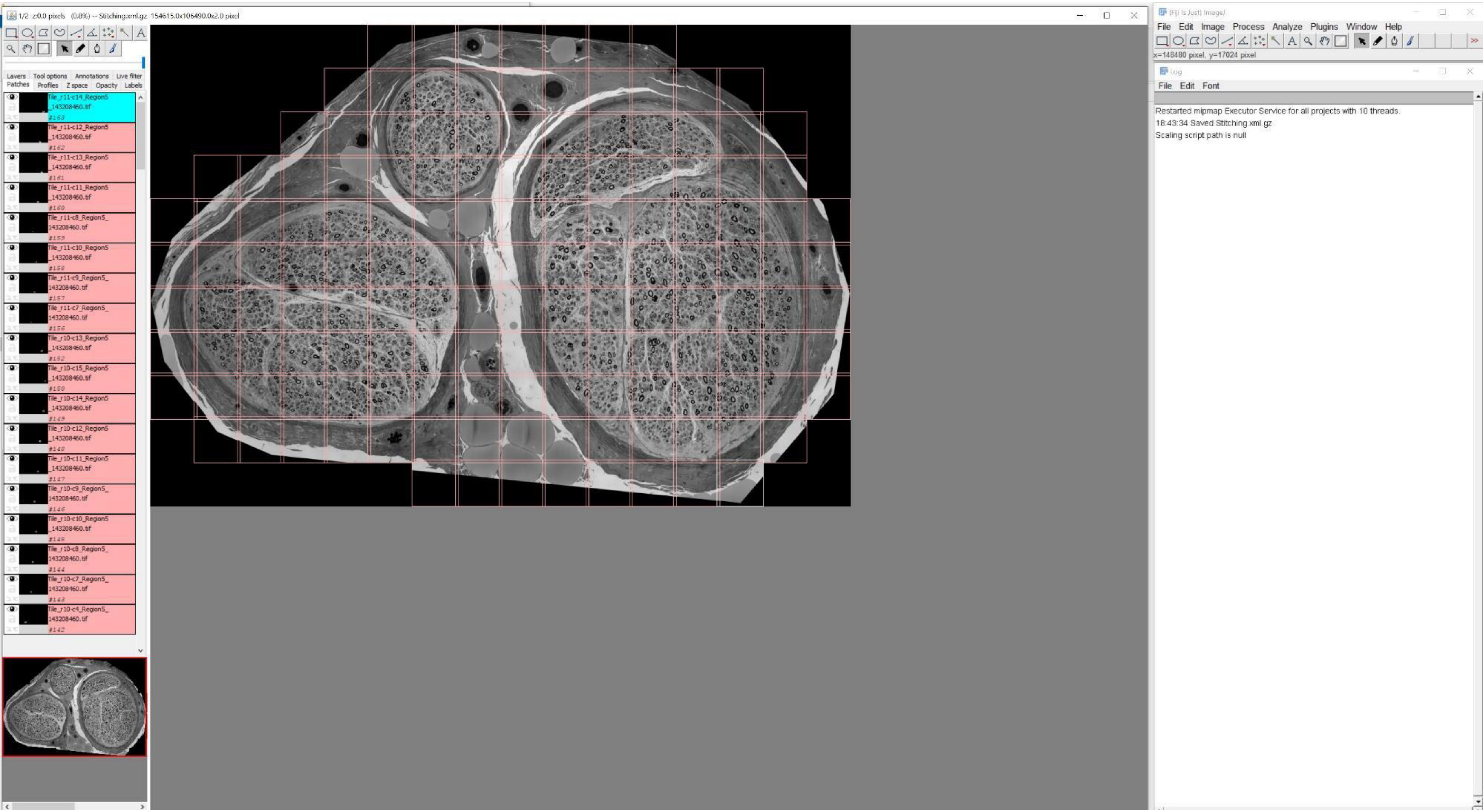
1/2 z:0.0 pixels (0.8%) -- Stitching.xml.gz 154615.0x106490.0x2.0 pixel
Layers Tool options Annotations Live filter
Patches Profiles Z space Opacity Labels
Tile_r11-c14_Region5_143208460.tif #163
Tile_r11-c12_Region5_143208460.tif #162
Tile_r11-c13_Region5_143208460.tif #161
Tile_r11-c11_Region5_143208460.tif #160
Tile_r11-c8_Region5_143208460.tif #159
Tile_r11-c10_Region5_143208460.tif #158
Tile_r11-c9_Region5_143208460.tif #157
Tile_r11-c7_Region5_143208460.tif #156
Tile_r10-c13_Region5_143208460.tif #152
Tile_r10-c15_Region5_143208460.tif #150
Tile_r10-c14_Region5_143208460.tif #149
Tile_r10-c12_Region5_143208460.tif #148
Tile_r10-c11_Region5_143208460.tif #147
Tile_r10-c9_Region5_143208460.tif #146
Tile_r10-c10_Region5_143208460.tif #145
Tile_r10-c8_Region5_143208460.tif #144
Tile_r10-c7_Region5_143208460.tif #143
Tile_r10-c4_Region5_143208460.tif #142
(Fiji Is Just) ImageJ
File Edit Image Process Analyze Plugins Window Help
x=148480 pixel, y=17024 pixel
Log
File Edit Font
Restarted mipmap Executor Service for all projects with 10 threads.
18:43:34 Saved Stitching.xml.gz
Scaling script path is null

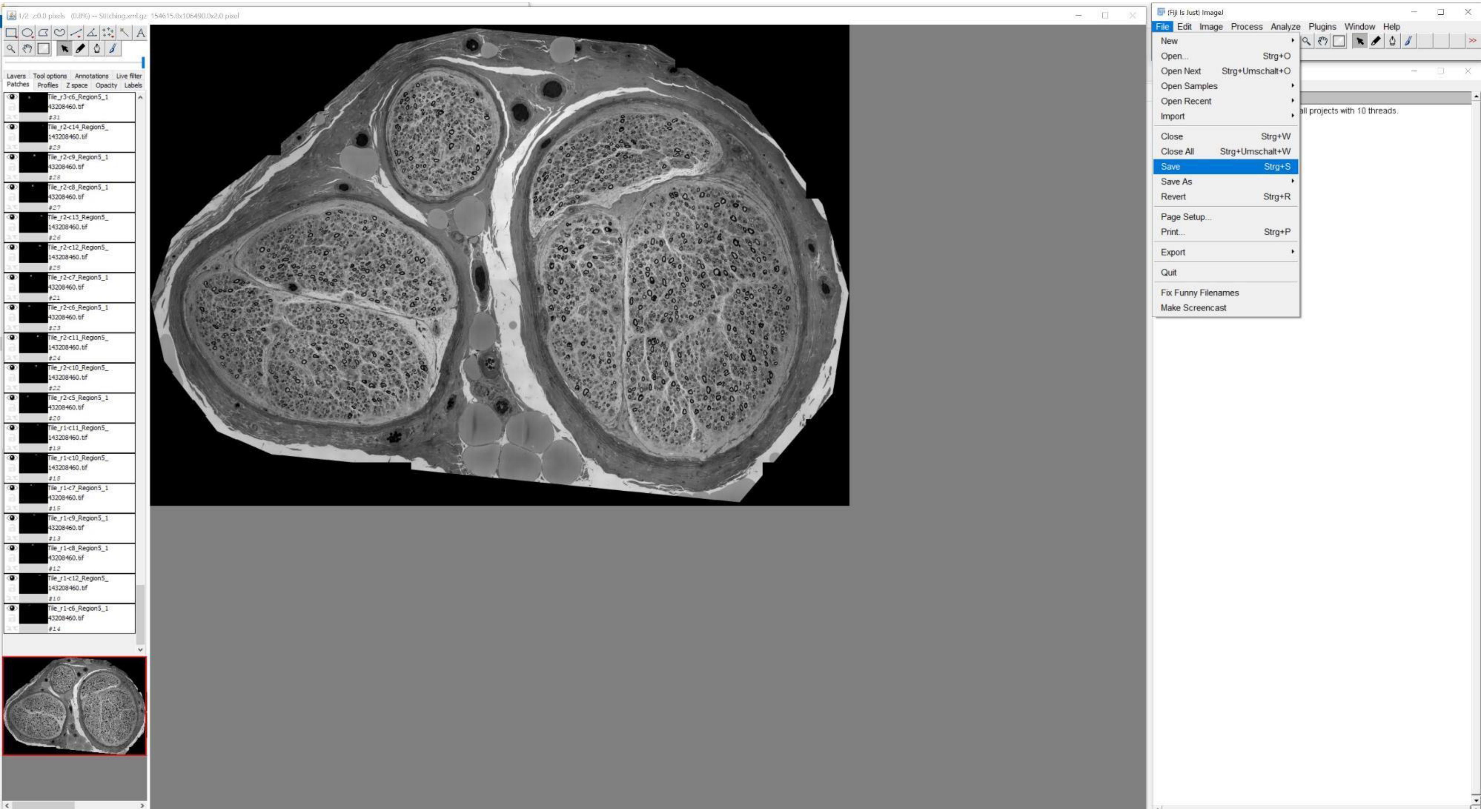

(Fiji Is Just) ImageJ
File
Edit
Image
Process
Analyze
Plugins
Window
Help
New
Open... Strg+O
Open Next Strg+Umschalt+O
Open Samples
Open Recent
Import
Close Strg+W
Close All Strg+Umschalt+W
Save Strg+S
Save As
Revert Strg+R
Page Setup...
Print... Strg+P
Export
Quit
Fix Funny Filenames
Make Screencast
all projects with 10 threads.
Layers
Tool options
Annotations
Live filter
Patches
Profiles
Z space
Opacity
Labels
Tile_r3-c6_Region5_143208460.tif
Tile_r2-c14_Region5_143208460.tif
Tile_r2-c9_Region5_143208460.tif
Tile_r2-c8_Region5_143208460.tif
Tile_r2-c13_Region5_143208460.tif
Tile_r2-c12_Region5_143208460.tif
Tile_r2-c7_Region5_143208460.tif
Tile_r2-c6_Region5_143208460.tif
Tile_r2-c11_Region5_143208460.tif
Tile_r2-c10_Region5_143208460.tif
Tile_r2-c5_Region5_143208460.tif
Tile_r1-c11_Region5_143208460.tif
Tile_r1-c10_Region5_143208460.tif
Tile_r1-c7_Region5_143208460.tif
Tile_r1-c9_Region5_143208460.tif
Tile_r1-c8_Region5_143208460.tif
Tile_r1-c12_Region5_143208460.tif
Tile_r1-c6_Region5_143208460.tif

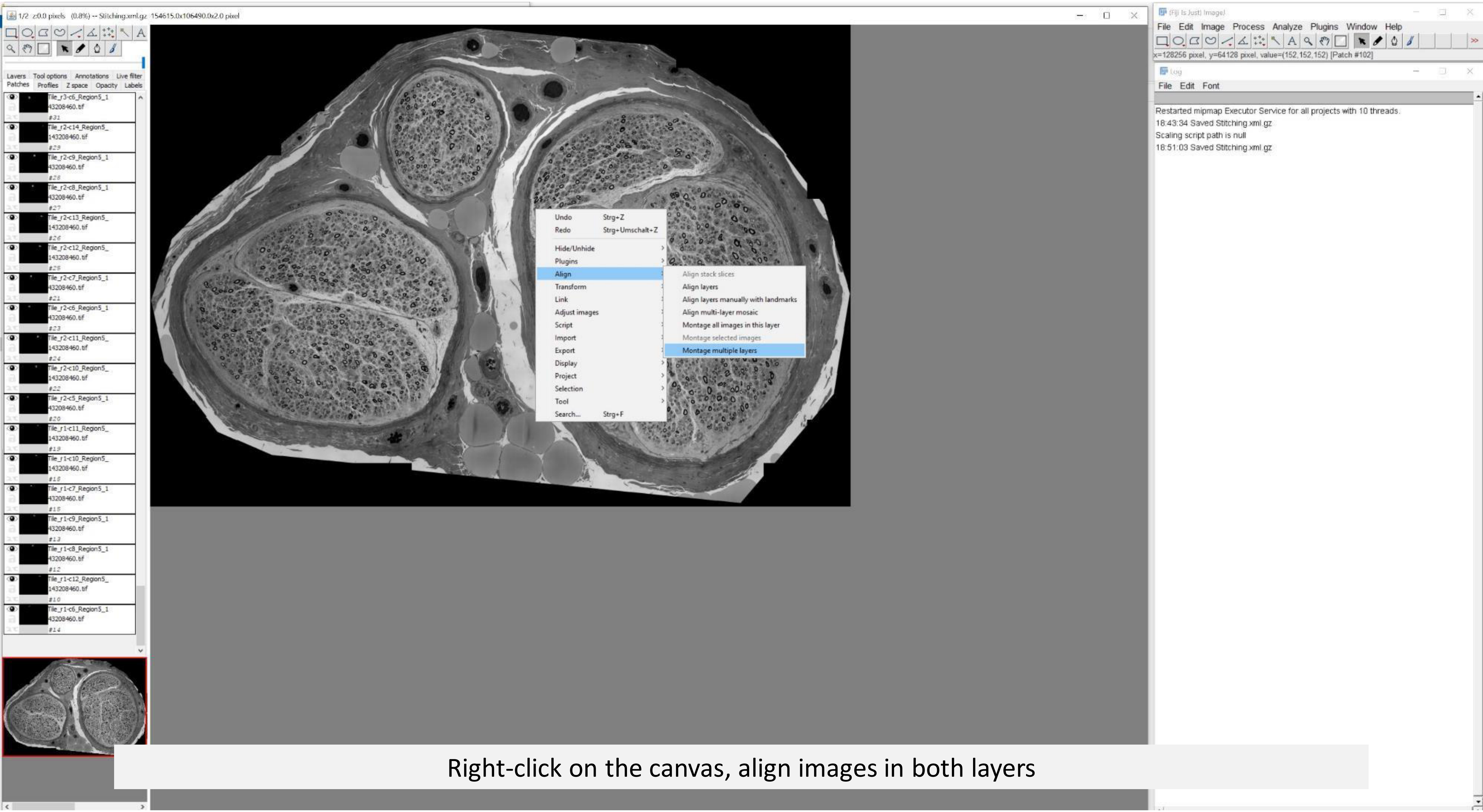


Right-click on the canvas, align images in both layers

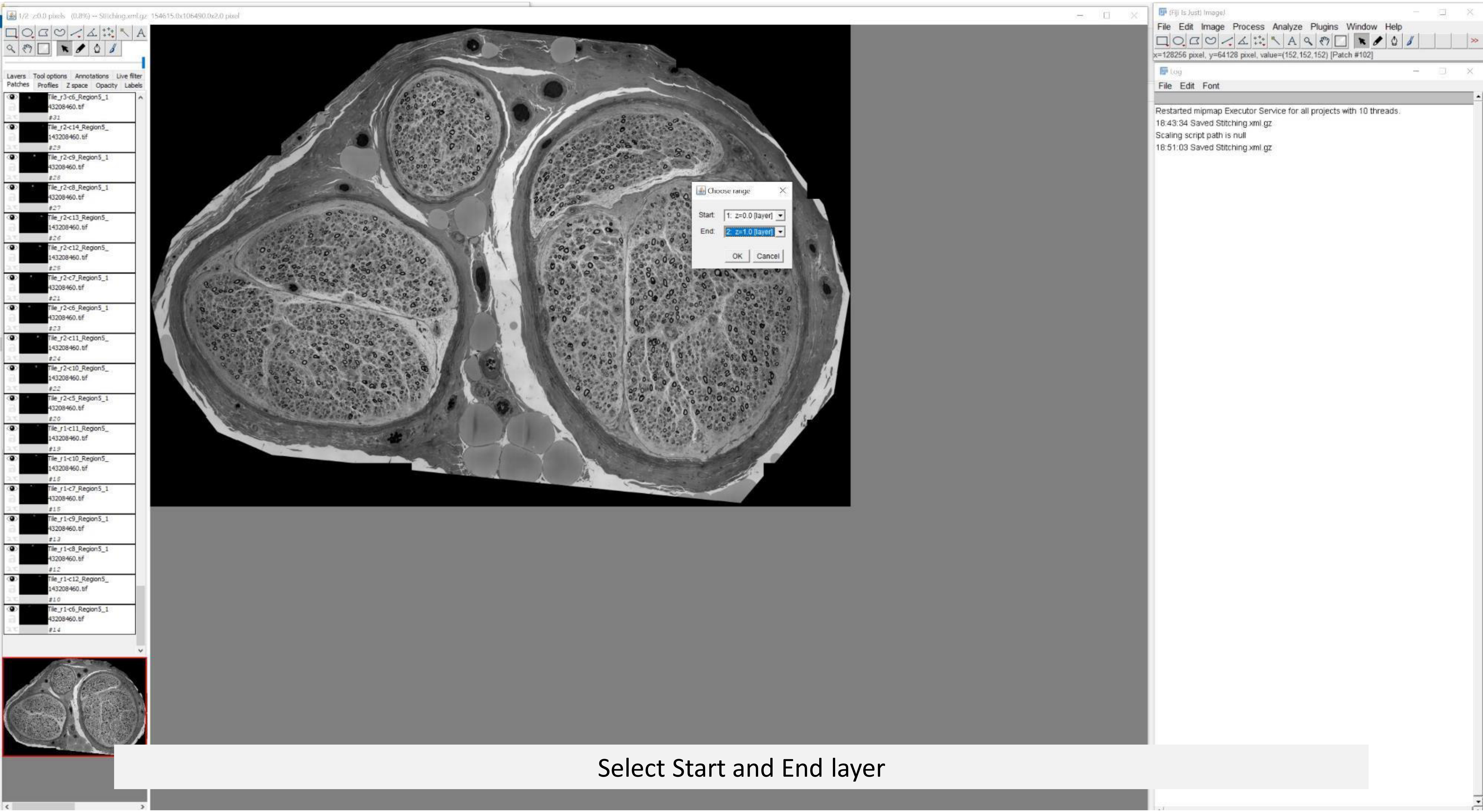


Select Start and End layer

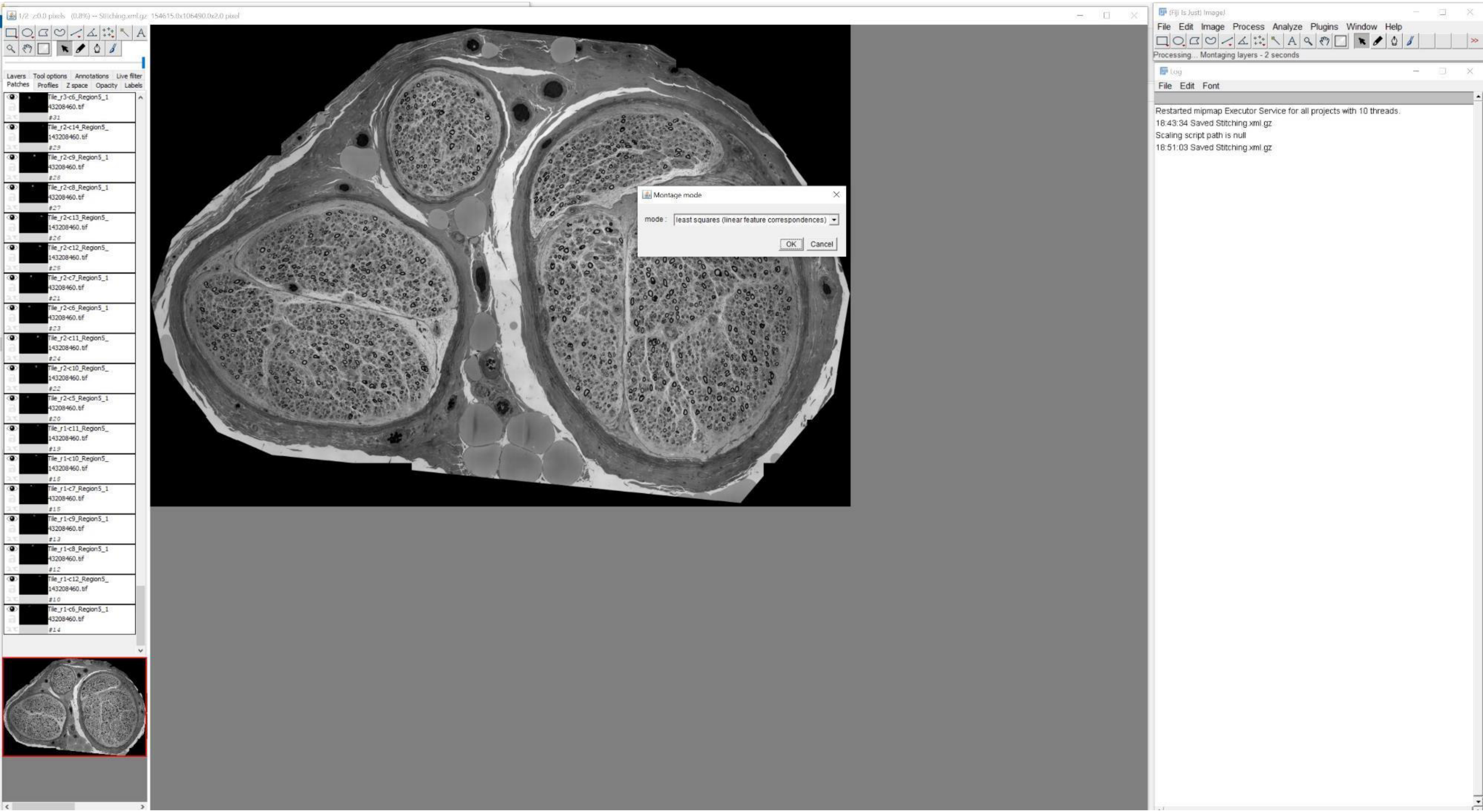
1/2 z:0.0 pixels (0.8%) -- Stitching.xml.gz 154615.0x106490.0x2.0 pixel
Layers Tool options Annotations Live filter
Patches Profiles Z space Opacity Labels
Tile_r3-c6_Region5_143208460.tif #31
Tile_r2-c14_Region5_143208460.tif #29
Tile_r2-c9_Region5_143208460.tif #28
Tile_r2-c8_Region5_143208460.tif #27
Tile_r2-c13_Region5_143208460.tif #26
Tile_r2-c12_Region5_143208460.tif #25
Tile_r2-c7_Region5_143208460.tif #21
Tile_r2-c6_Region5_143208460.tif #23
Tile_r2-c11_Region5_143208460.tif #24
Tile_r2-c10_Region5_143208460.tif #22
Tile_r2-c5_Region5_143208460.tif #20
Tile_r1-c11_Region5_143208460.tif #19
Tile_r1-c10_Region5_143208460.tif #18
Tile_r1-c7_Region5_143208460.tif #15
Tile_r1-c9_Region5_143208460.tif #13
Tile_r1-c8_Region5_143208460.tif #12
Tile_r1-c12_Region5_143208460.tif #10
Tile_r1-c6_Region5_143208460.tif #14
Montage mode
mode : least squares (linear feature correspondences)
OK
Cancel
(Fiji Is Just) ImageJ
File Edit Image Process Analyze Plugins Window Help
Processing... Montaging layers - 2 seconds
Log
File Edit Font
Restarted mipmap Executor Service for all projects with 10 threads.
18:43:34 Saved Stitching.xml.gz
Scaling script path is null
18:51:03 Saved Stitching.xml.gz

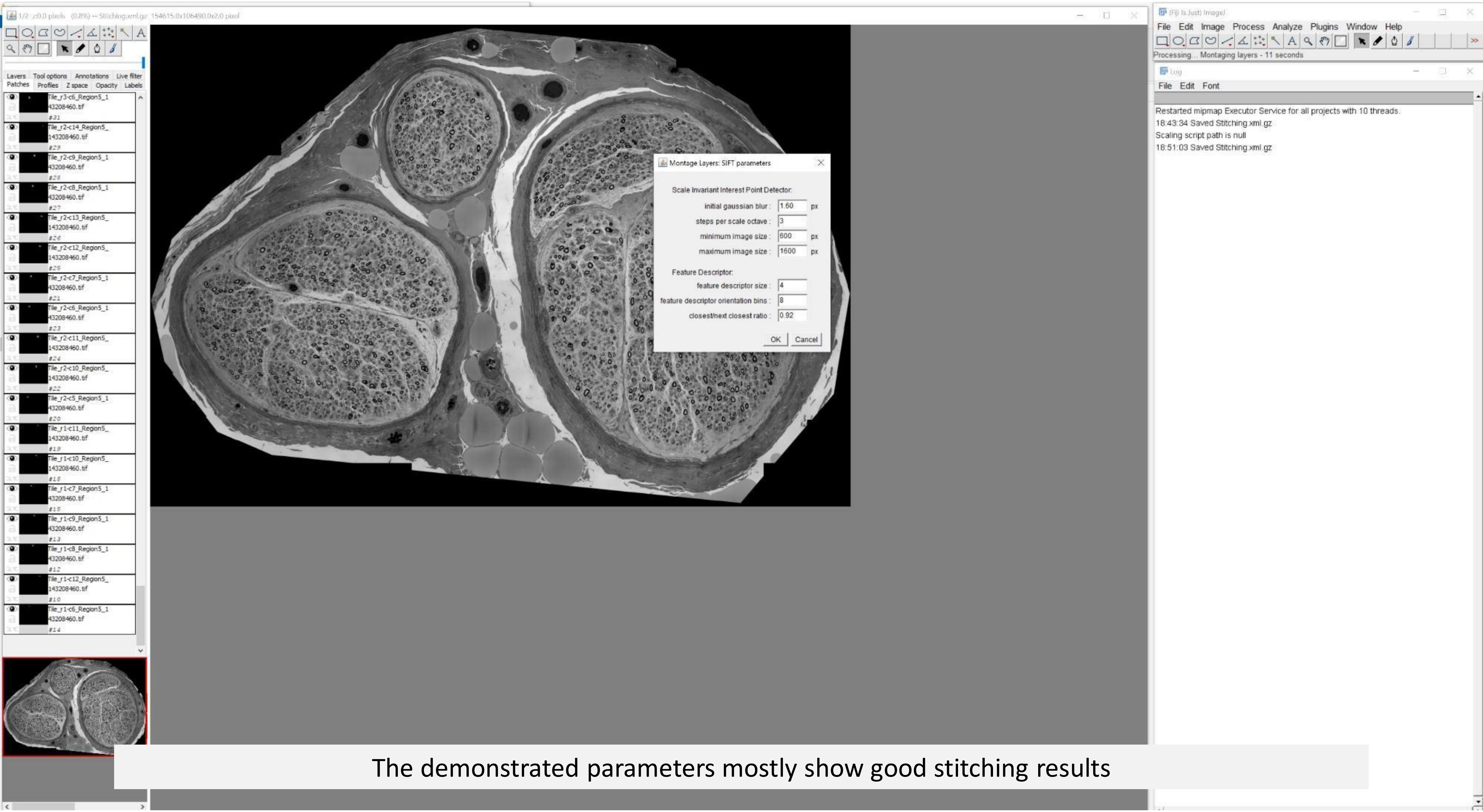


The demonstrated parameters mostly show good stitching results

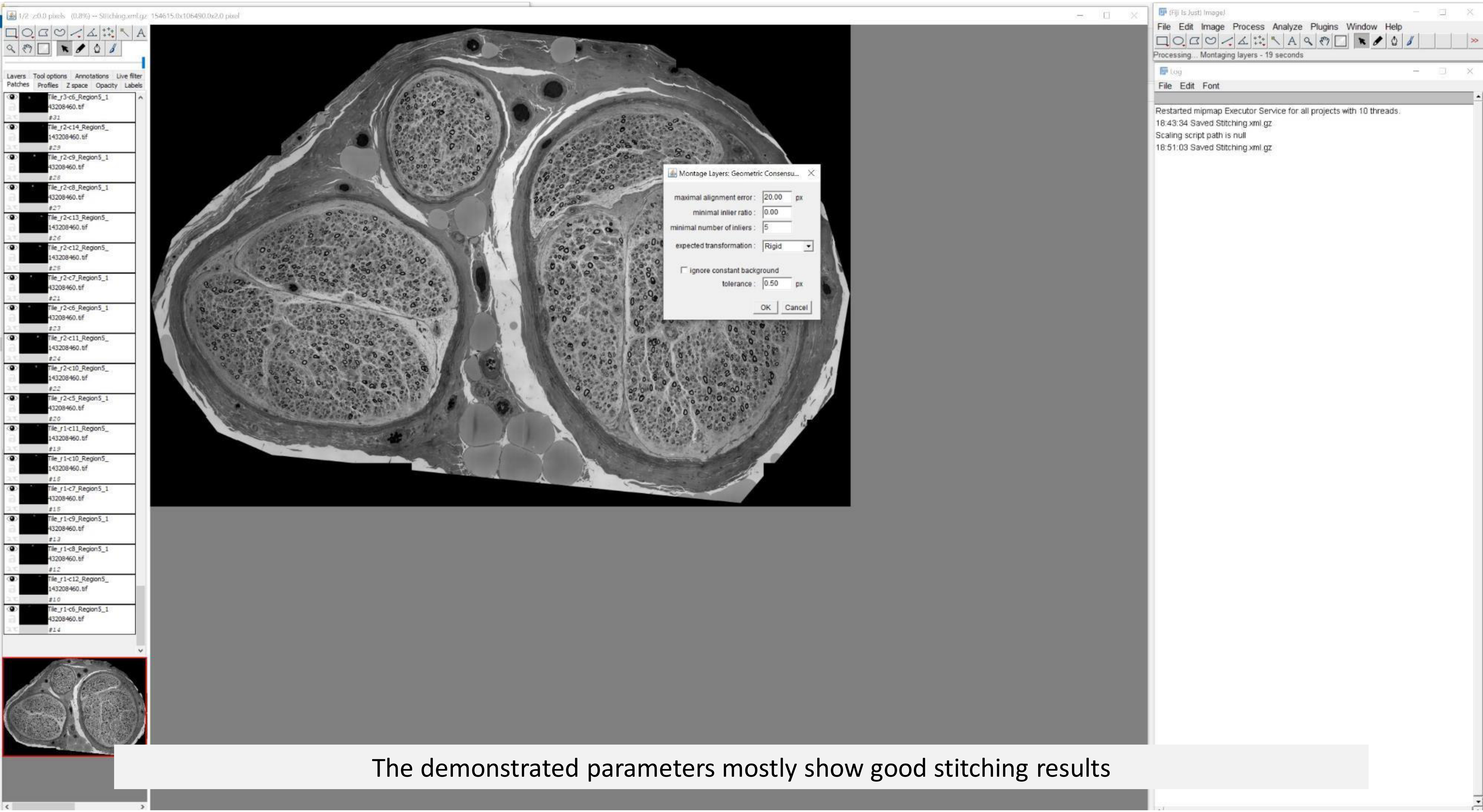


The demonstrated parameters mostly show good stitching results

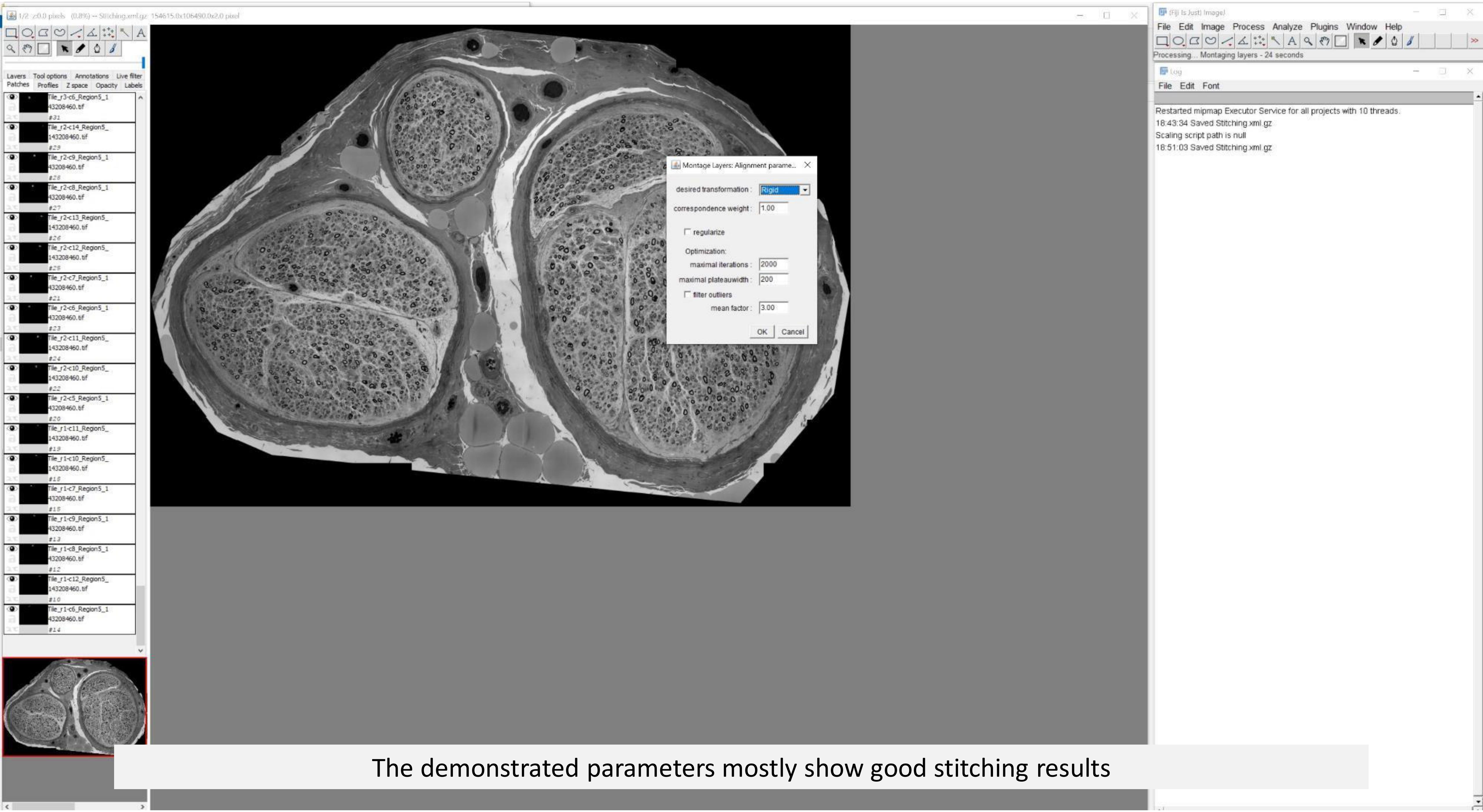


The demonstrated parameters mostly show good stitching results

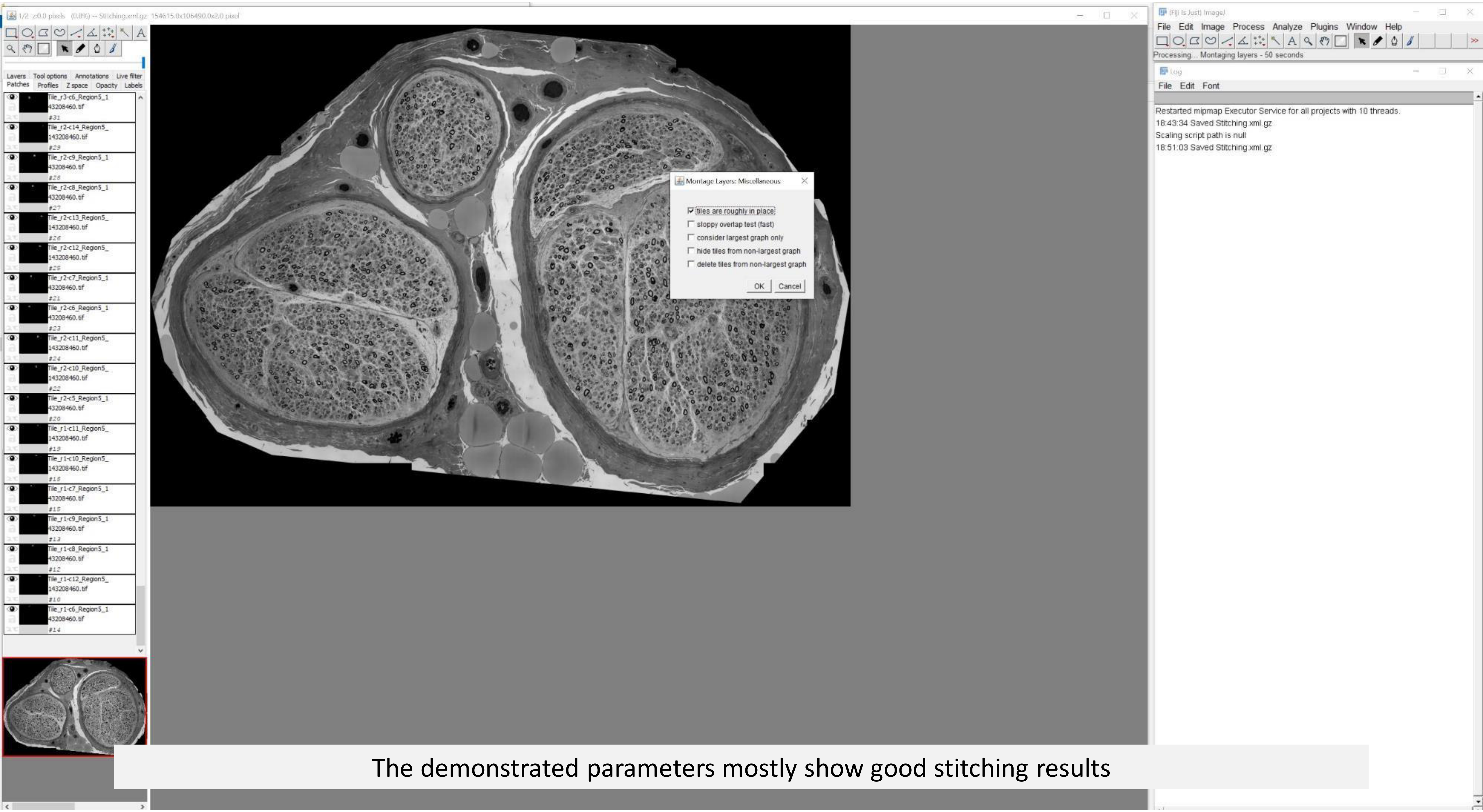


The demonstrated parameters mostly show good stitching results

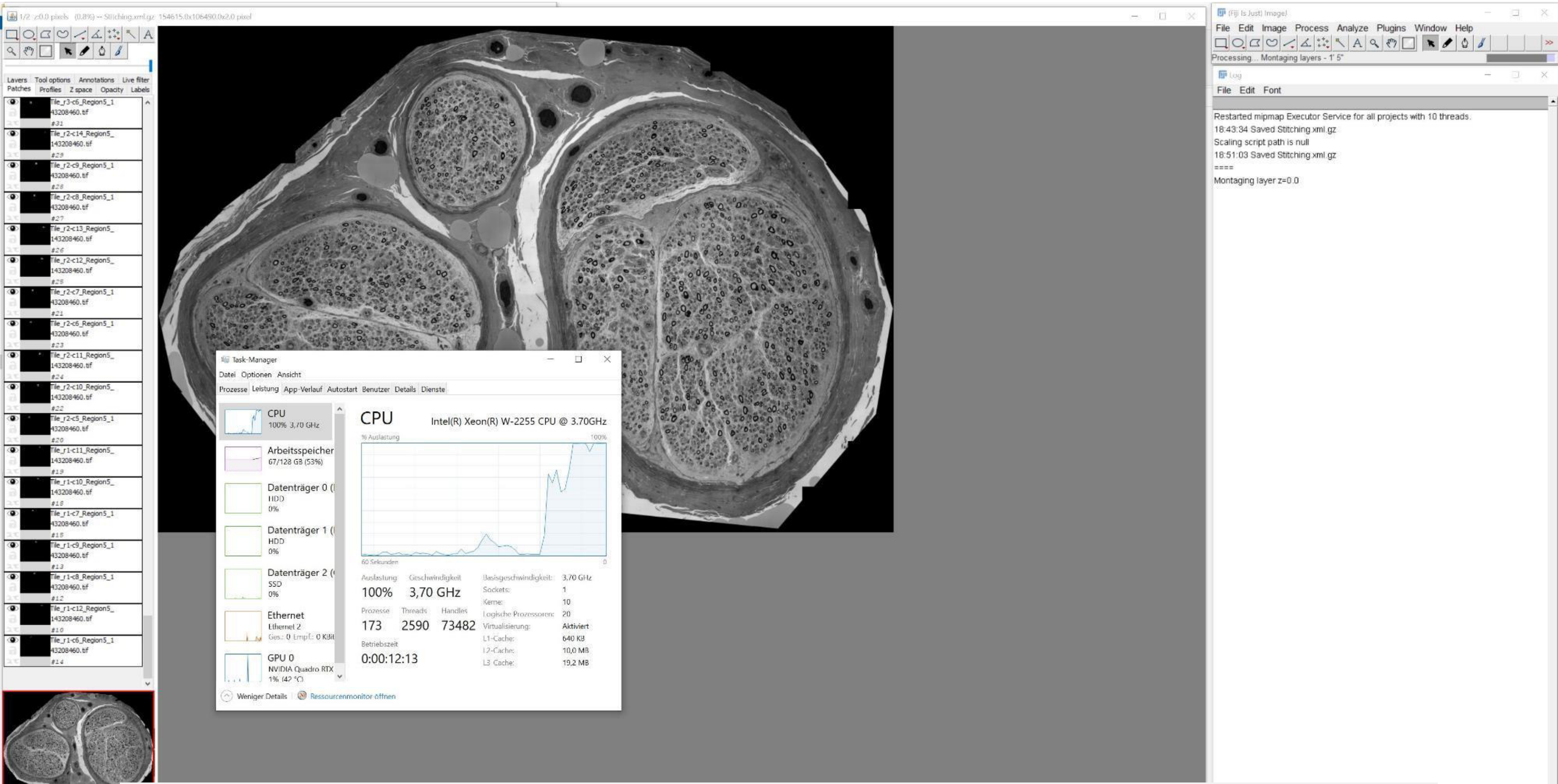


See information in Log

See Task Manager for workstation performance (multiple cores can be used for stitching)

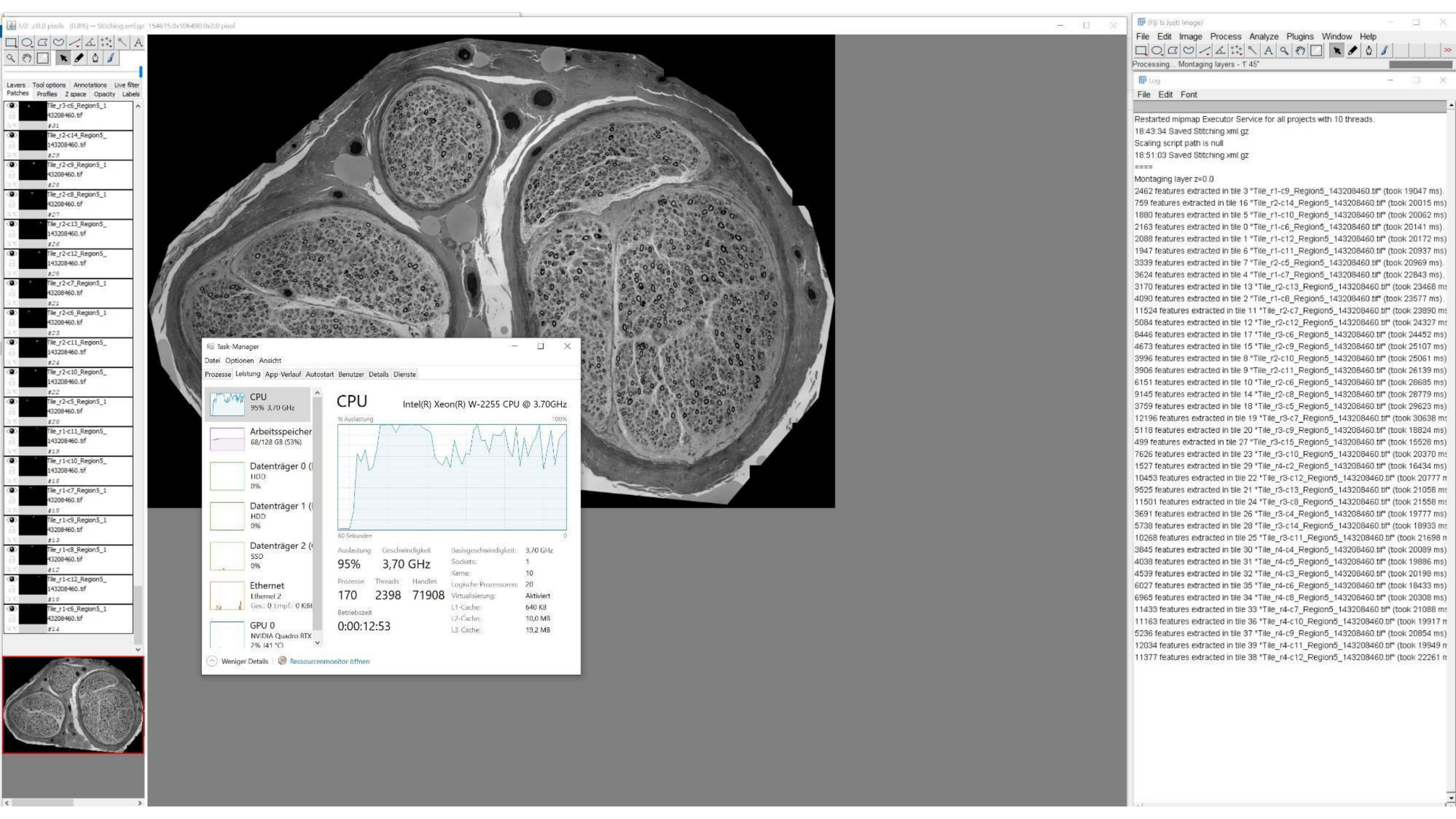

Task-Manager
Datei Optionen Ansicht
Prozesse Leistung App-Verlauf Autostart Benutzer Details Dienste
CPU
95% 3,70 GHz
Arbeitsspeicher
68/128 GB (53%)
Datenträger 0
HDD
0%
Datenträger 1
HDD
0%
Datenträger 2
SSD
0%
Ethernet
Ethernet 2
GPU 0
NVIDIA Quadro RTX
CPU
Intel(R) Xeon(R) W-2255 CPU @ 3.70GHz
% Auslastung
100%
60 Sekunden
Auslastung 95%
Geschwindigkeit 3,70 GHz
Prozesse 170
Threads 2398
Handles 71908
Betriebszeit 0:00:12:53
Basisgeschwindigkeit: 3,70 GHz
Sockets: 1
Kerne: 10
Logische Prozessoren: 20
Virtualisierung: Aktiviert
L1-Cache: 640 KB
L2-Cache: 10,0 MB
L3-Cache: 19,2 MB
Weniger Details
Ressourcenmonitor öffnen
(Fiji Is Just) ImageJ
File Edit Image Process Analyze Plugins Window Help
Processing... Montaging layers - 1' 45"
Log
File Edit Font
Restarted mipmap Executor Service for all projects with 10 threads.
18:43:34 Saved Stitching.xml.gz
Scaling script path is null
18:51:03 Saved Stitching.xml.gz
====
Montaging layer z=0.0
2462 features extracted in tile 3 "Tile_r1-c9_Region5_143208460.tif" (took 19047 ms).
759 features extracted in tile 16 "Tile_r2-c14_Region5_143208460.tif" (took 20015 ms)
1880 features extracted in tile 5 "Tile_r1-c10_Region5_143208460.tif" (took 20062 ms)
2163 features extracted in tile 0 "Tile_r1-c6_Region5_143208460.tif" (took 20141 ms).
2088 features extracted in tile 1 "Tile_r1-c12_Region5_143208460.tif" (took 20172 ms)
1947 features extracted in tile 6 "Tile_r1-c11_Region5_143208460.tif" (took 20937 ms)
3339 features extracted in tile 7 "Tile_r2-c5_Region5_143208460.tif" (took 20969 ms).
3624 features extracted in tile 4 "Tile_r1-c7_Region5_143208460.tif" (took 22843 ms).
3170 features extracted in tile 13 "Tile_r2-c13_Region5_143208460.tif" (took 23468 ms
4090 features extracted in tile 2 "Tile_r1-c8_Region5_143208460.tif" (took 23577 ms).
11524 features extracted in tile 11 "Tile_r2-c7_Region5_143208460.tif" (took 23890 ms
5084 features extracted in tile 12 "Tile_r2-c12_Region5_143208460.tif" (took 24327 ms
8446 features extracted in tile 17 "Tile_r3-c6_Region5_143208460.tif" (took 24452 ms)
4673 features extracted in tile 15 "Tile_r2-c9_Region5_143208460.tif" (took 25107 ms)
3996 features extracted in tile 8 "Tile_r2-c10_Region5_143208460.tif" (took 25061 ms)
3906 features extracted in tile 9 "Tile_r2-c11_Region5_143208460.tif" (took 26139 ms)
6151 features extracted in tile 10 "Tile_r2-c6_Region5_143208460.tif" (took 26685 ms)
9145 features extracted in tile 14 "Tile_r2-c8_Region5_143208460.tif" (took 28779 ms)
3759 features extracted in tile 18 "Tile_r3-c5_Region5_143208460.tif" (took 29623 ms)
12196 features extracted in tile 19 "Tile_r3-c7_Region5_143208460.tif" (took 30638 ms
5118 features extracted in tile 20 "Tile_r3-c9_Region5_143208460.tif" (took 16824 ms)
499 features extracted in tile 27 "Tile_r3-c15_Region5_143208460.tif" (took 15528 ms)
7626 features extracted in tile 23 "Tile_r3-c10_Region5_143208460.tif" (took 20370 ms
1527 features extracted in tile 29 "Tile_r4-c2_Region5_143208460.tif" (took 16434 ms)
10453 features extracted in tile 22 "Tile_r3-c12_Region5_143208460.tif" (took 20777 m
9525 features extracted in tile 21 "Tile_r3-c13_Region5_143208460.tif" (took 21058 ms
11501 features extracted in tile 24 "Tile_r3-c8_Region5_143208460.tif" (took 21558 ms
3691 features extracted in tile 26 "Tile_r3-c4_Region5_143208460.tif" (took 19777 ms)
5738 features extracted in tile 28 "Tile_r3-c14_Region5_143208460.tif" (took 18933 ms
10268 features extracted in tile 25 "Tile_r3-c11_Region5_143208460.tif" (took 21698 m
3845 features extracted in tile 30 "Tile_r4-c4_Region5_143208460.tif" (took 20089 ms)
4038 features extracted in tile 31 "Tile_r4-c5_Region5_143208460.tif" (took 19886 ms)
4539 features extracted in tile 32 "Tile_r4-c3_Region5_143208460.tif" (took 20198 ms)
6027 features extracted in tile 35 "Tile_r4-c6_Region5_143208460.tif" (took 18433 ms)
6965 features extracted in tile 34 "Tile_r4-c8_Region5_143208460.tif" (took 20308 ms)
11433 features extracted in tile 33 "Tile_r4-c7_Region5_143208460.tif" (took 21088 ms
11163 features extracted in tile 36 "Tile_r4-c10_Region5_143208460.tif" (took 19917 m
5236 features extracted in tile 37 "Tile_r4-c9_Region5_143208460.tif" (took 20854 ms)
12034 features extracted in tile 39 "Tile_r4-c11_Region5_143208460.tif" (took 19949 m
11377 features extracted in tile 38 "Tile_r4-c12_Region5_143208460.tif" (took 22261 m

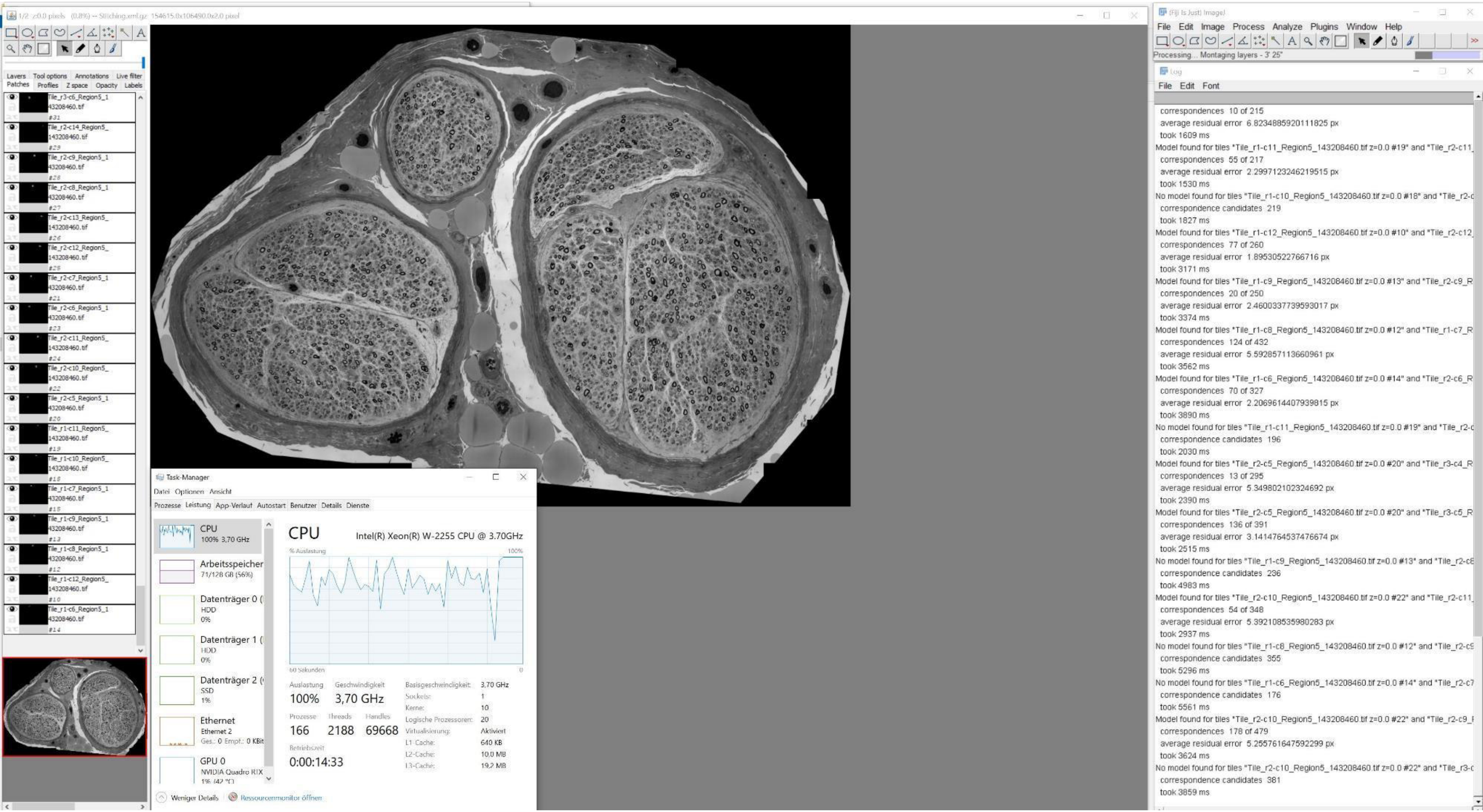
1/2 z:0.0 pixels (0.8%) -- Stitching.xml.gz 154615.0x106490.0x2.0 pixel
Layers Tool options Annotations Live filter
Patches Profiles Z space Opacity Labels
Tile_r3-c6_Region5_143208460.tif
Tile_r2-c14_Region5_143208460.tif
Tile_r2-c9_Region5_143208460.tif
Tile_r2-c8_Region5_143208460.tif
Tile_r2-c13_Region5_143208460.tif
Tile_r2-c12_Region5_143208460.tif
Tile_r2-c7_Region5_143208460.tif
Tile_r2-c6_Region5_143208460.tif
Tile_r2-c11_Region5_143208460.tif
Tile_r2-c10_Region5_143208460.tif
Tile_r2-c5_Region5_143208460.tif
Tile_r1-c11_Region5_143208460.tif
Tile_r1-c10_Region5_143208460.tif
Tile_r1-c7_Region5_143208460.tif
Tile_r1-c9_Region5_143208460.tif
Tile_r1-c8_Region5_143208460.tif
Tile_r1-c12_Region5_143208460.tif
Tile_r1-c6_Region5_143208460.tif
Task-Manager
Datei Optionen Ansicht
Prozesse Leistung App-Verlauf Autostart Benutzer Details Dienste
CPU 100% 3,70 GHz
Arbeitsspeicher 71/128 GB (56%)
Datenträger 0 HDD 0%
Datenträger 1 HDD 0%
Datenträger 2 SSD 1%
Ethernet Ethernet 2 Ges.: 0 Empf.: 0 KBit
GPU 0 NVIDIA Quadro RTX
CPU
Intel(R) Xeon(R) W-2255 CPU @ 3.70GHz
% Auslastung
100%
60 Sekunden
0
Auslastung 100%
Geschwindigkeit 3,70 GHz
Prozesse 166
Threads 2188
Handles 69668
Betriebszeit 0:00:14:33
Basisgeschwindigkeit: 3,70 GHz
Sockets: 1
Kerne: 10
Logische Prozessoren: 20
Virtualisierung: Aktiviert
L1-Cache: 640 KB
L2-Cache: 10,0 MB
L3-Cache: 19,2 MB
Weniger Details
Ressourcenmonitor öffnen
(Fiji Is Just) ImageJ
File Edit Image Process Analyze Plugins Window Help
Processing... Montaging layers - 3' 25"
Log
File Edit Font
correspondences 10 of 215
average residual error 6.8234885920111825 px
took 1609 ms
Model found for tiles "Tile_r1-c11_Region5_143208460.tif z=0.0 #19" and "Tile_r2-c11_
correspondences 55 of 217
average residual error 2.2997123246219515 px
took 1530 ms
No model found for tiles "Tile_r1-c10_Region5_143208460.tif z=0.0 #18" and "Tile_r2-c
correspondence candidates 219
took 1827 ms
Model found for tiles "Tile_r1-c12_Region5_143208460.tif z=0.0 #10" and "Tile_r2-c12_
correspondences 77 of 260
average residual error 1.89530522766716 px
took 3171 ms
Model found for tiles "Tile_r1-c9_Region5_143208460.tif z=0.0 #13" and "Tile_r2-c9_R
correspondences 20 of 250
average residual error 2.4600337739593017 px
took 3374 ms
Model found for tiles "Tile_r1-c8_Region5_143208460.tif z=0.0 #12" and "Tile_r1-c7_R
correspondences 124 of 432
average residual error 5.592857113660961 px
took 3562 ms
Model found for tiles "Tile_r1-c6_Region5_143208460.tif z=0.0 #14" and "Tile_r2-c6_R
correspondences 70 of 327
average residual error 2.2069614407939815 px
took 3890 ms
No model found for tiles "Tile_r1-c11_Region5_143208460.tif z=0.0 #19" and "Tile_r2-c
correspondence candidates 196
took 2030 ms
Model found for tiles "Tile_r2-c5_Region5_143208460.tif z=0.0 #20" and "Tile_r3-c4_R
correspondences 13 of 295
average residual error 5.349802102324692 px
took 2390 ms
Model found for tiles "Tile_r2-c5_Region5_143208460.tif z=0.0 #20" and "Tile_r3-c5_R
correspondences 136 of 391
average residual error 3.1414764537476674 px
took 2515 ms
No model found for tiles "Tile_r1-c9_Region5_143208460.tif z=0.0 #13" and "Tile_r2-c8
correspondence candidates 236
took 4983 ms
Model found for tiles "Tile_r2-c10_Region5_143208460.tif z=0.0 #22" and "Tile_r2-c11_
correspondences 54 of 348
average residual error 5.392108535980283 px
took 2937 ms
No model found for tiles "Tile_r1-c8_Region5_143208460.tif z=0.0 #12" and "Tile_r2-c9
correspondence candidates 355
took 5296 ms
No model found for tiles "Tile_r1-c6_Region5_143208460.tif z=0.0 #14" and "Tile_r2-c7
correspondence candidates 176
took 5561 ms
Model found for tiles "Tile_r2-c10_Region5_143208460.tif z=0.0 #22" and "Tile_r2-c9_
correspondences 178 of 479
average residual error 5.255761647592299 px
took 3624 ms
No model found for tiles "Tile_r2-c10_Region5_143208460.tif z=0.0 #22" and "Tile_r3-c
correspondence candidates 381
took 3859 ms

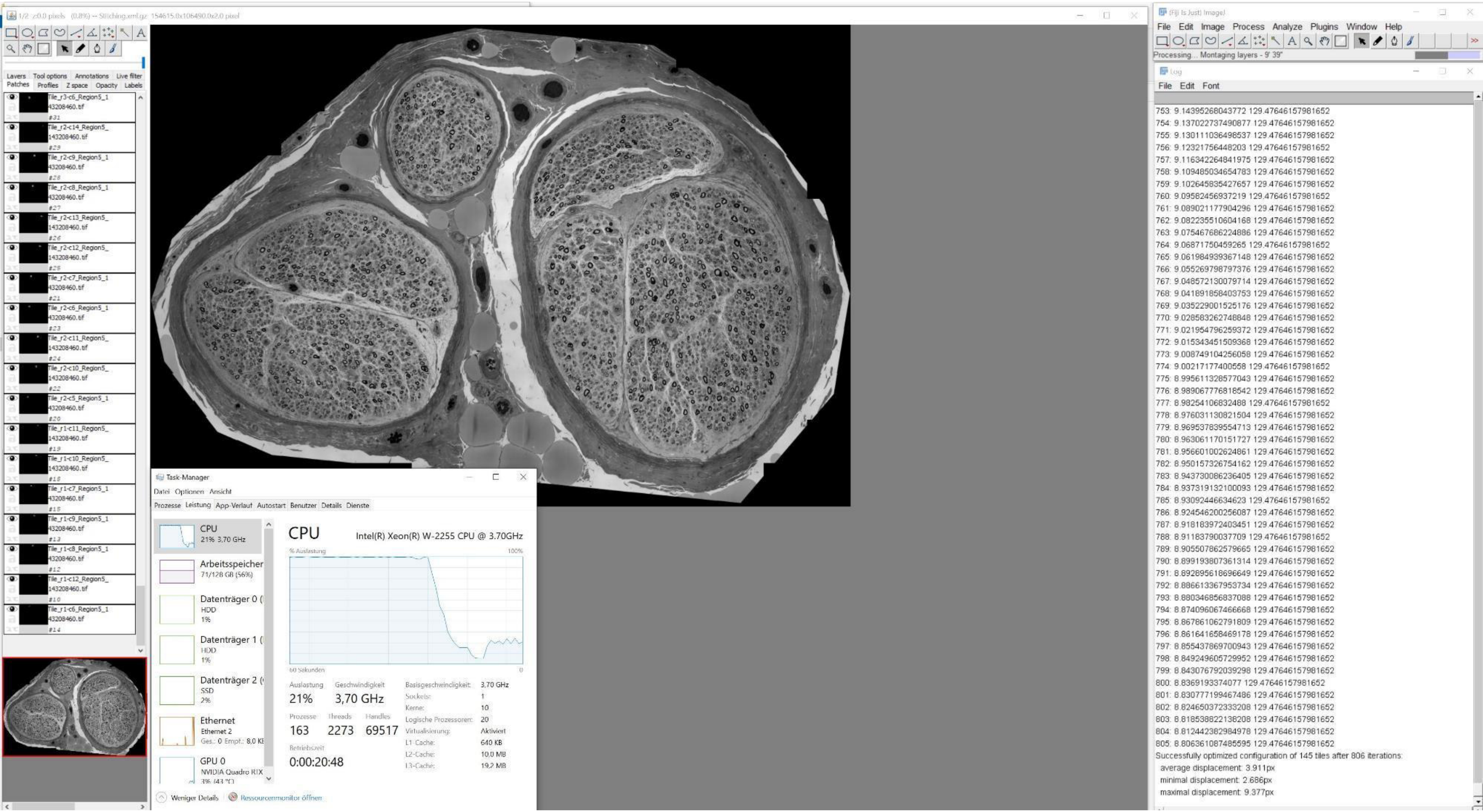
1/2 z:0.0 pixels (0.8%) -- Stitching.xml.gz 154615.0x106490.0x2.0 pixel
Layers Tool options Annotations Live filter
Patches Profiles Z space Opacity Labels
Tile_r3-c6_Region5_143208460.tif
Tile_r2-c14_Region5_143208460.tif
Tile_r2-c9_Region5_143208460.tif
Tile_r2-c8_Region5_143208460.tif
Tile_r2-c13_Region5_143208460.tif
Tile_r2-c12_Region5_143208460.tif
Tile_r2-c7_Region5_143208460.tif
Tile_r2-c6_Region5_143208460.tif
Tile_r2-c11_Region5_143208460.tif
Tile_r2-c10_Region5_143208460.tif
Tile_r2-c5_Region5_143208460.tif
Tile_r1-c11_Region5_143208460.tif
Tile_r1-c10_Region5_143208460.tif
Tile_r1-c7_Region5_143208460.tif
Tile_r1-c9_Region5_143208460.tif
Tile_r1-c8_Region5_143208460.tif
Tile_r1-c12_Region5_143208460.tif
Tile_r1-c6_Region5_143208460.tif
Task-Manager
Datei Optionen Ansicht
Prozesse Leistung App-Verlauf Autostart Benutzer Details Dienste
CPU
21% 3,70 GHz
Arbeitsspeicher
71/128 GB (56%)
Datenträger 0 (
HDD
1%
Datenträger 1 (
HDD
1%
Datenträger 2 (
SSD
2%
Ethernet
Ethernet 2
Ges.: 0 Empf.: 8,0 KB
GPU 0
NVIDIA Quadro RTX
CPU
Intel(R) Xeon(R) W-2255 CPU @ 3.70GHz
% Auslastung
100%
60 Sekunden
0
Auslastung
21%
Geschwindigkeit
3,70 GHz
Prozesse
163
Threads
2273
Handles
69517
Betriebszeit
0:00:20:48
Basisgeschwindigkeit: 3,70 GHz
Sockets: 1
Kerne: 10
Logische Prozessoren: 20
Virtualisierung: Aktiviert
L1-Cache: 640 KB
L2-Cache: 10,0 MB
L3-Cache: 19,2 MB
Weniger Details
Ressourcenmonitor öffnen
(Fiji Is Just) ImageJ
File Edit Image Process Analyze Plugins Window Help
Processing... Montaging layers - 9' 39"
Log
File Edit Font
753: 9.14395268043772 129.47646157981652
754: 9.137022737490877 129.47646157981652
755: 9.130111036498537 129.47646157981652
756: 9.12321756448203 129.47646157981652
757: 9.116342264841975 129.47646157981652
758: 9.109485034654783 129.47646157981652
759: 9.102645835427657 129.47646157981652
760: 9.09582456937219 129.47646157981652
761: 9.089021177904296 129.47646157981652
762: 9.082235510604168 129.47646157981652
763: 9.075467686224886 129.47646157981652
764: 9.06871750459265 129.47646157981652
765: 9.061984939367148 129.47646157981652
766: 9.055269798797376 129.47646157981652
767: 9.048572130079714 129.47646157981652
768: 9.041891858403753 129.47646157981652
769: 9.035229001525176 129.47646157981652
770: 9.028583262748848 129.47646157981652
771: 9.021954796259372 129.47646157981652
772: 9.015343451509368 129.47646157981652
773: 9.008749104256058 129.47646157981652
774: 9.00217177400558 129.47646157981652
775: 8.995611328577043 129.47646157981652
776: 8.989067776818542 129.47646157981652
777: 8.98254106832488 129.47646157981652
778: 8.976031130821504 129.47646157981652
779: 8.969537839554713 129.47646157981652
780: 8.963061170151727 129.47646157981652
781: 8.956601002624861 129.47646157981652
782: 8.950157326754162 129.47646157981652
783: 8.943730086236405 129.47646157981652
784: 8.937319132100093 129.47646157981652
785: 8.93092446634623 129.47646157981652
786: 8.924546200256087 129.47646157981652
787: 8.918183972403451 129.47646157981652
788: 8.91183790037709 129.47646157981652
789: 8.905507862579665 129.47646157981652
790: 8.899193807361314 129.47646157981652
791: 8.892895618696649 129.47646157981652
792: 8.886613367953734 129.47646157981652
793: 8.880346856837088 129.47646157981652
794: 8.874096067466668 129.47646157981652
795: 8.867861062791809 129.47646157981652
796: 8.861641658469178 129.47646157981652
797: 8.855437869700943 129.47646157981652
798: 8.849249605729952 129.47646157981652
799: 8.843076792039298 129.47646157981652
800: 8.8369193374077 129.47646157981652
801: 8.830777199467486 129.47646157981652
802: 8.824650372333208 129.47646157981652
803: 8.818538822138208 129.47646157981652
804: 8.812442382984978 129.47646157981652
805: 8.806361087485595 129.47646157981652
Successfully optimized configuration of 145 tiles after 806 iterations:
average displacement: 3.911px
minimal displacement: 2.686px
maximal displacement: 9.377px

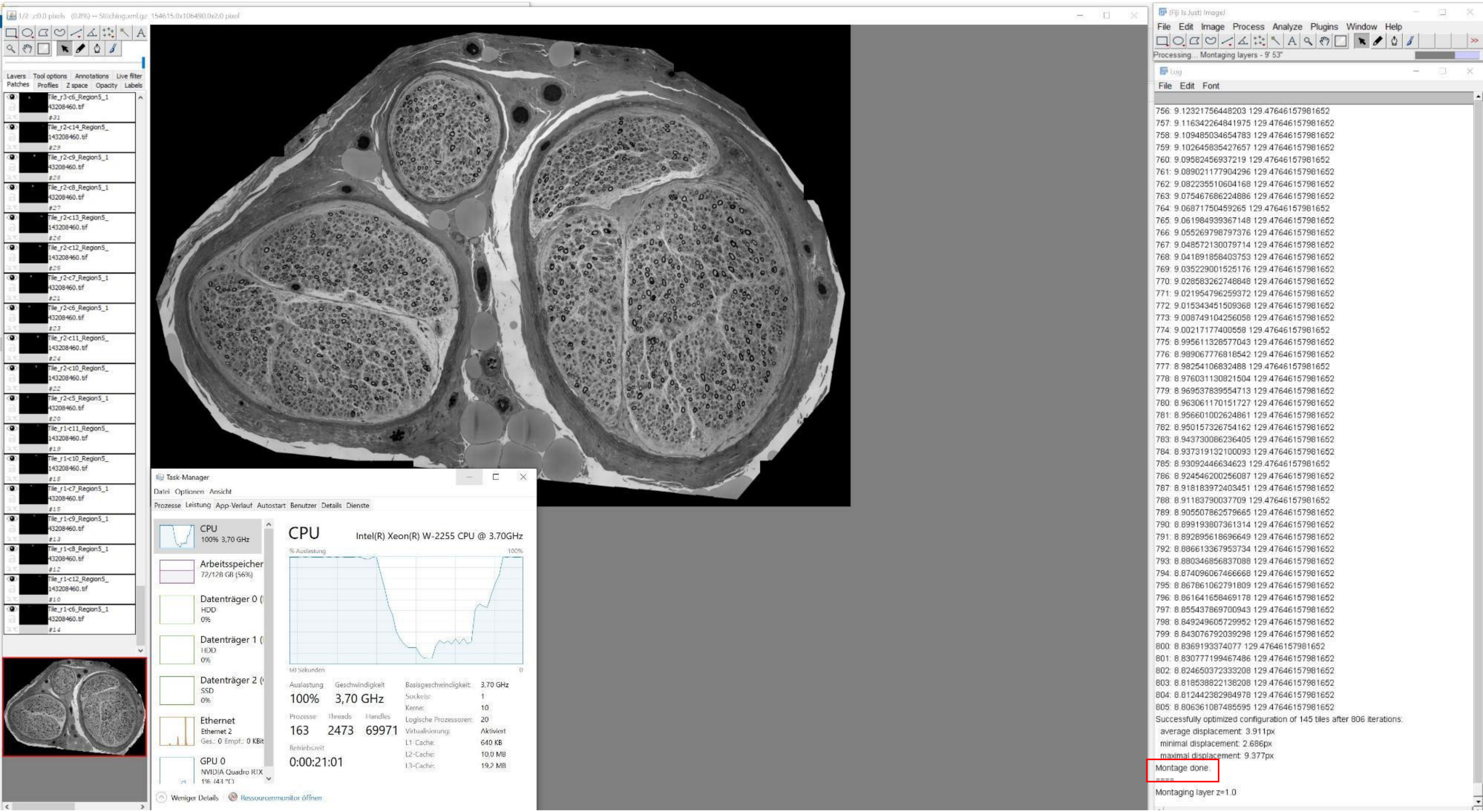

1/2 z:0.0 pixels (0.8%) -- Stitching.xml.gz 154615.0x106490.0x2.0 pixel
Layers Tool options Annotations Live filter
Patches Profiles Z space Opacity Labels
Tile_r3-c6_Region5_1 43208460.tif #31
Tile_r2-c14_Region5_143208460.tif #29
Tile_r2-c9_Region5_1 43208460.tif #28
Tile_r2-c8_Region5_1 43208460.tif #27
Tile_r2-c13_Region5_143208460.tif #26
Tile_r2-c12_Region5_143208460.tif #25
Tile_r2-c7_Region5_1 43208460.tif #23
Tile_r2-c6_Region5_1 43208460.tif #21
Tile_r2-c11_Region5_143208460.tif #24
Tile_r2-c10_Region5_143208460.tif #22
Tile_r2-c5_Region5_1 43208460.tif #20
Tile_r1-c11_Region5_143208460.tif #19
Tile_r1-c10_Region5_143208460.tif #18
Tile_r1-c7_Region5_1 43208460.tif #15
Tile_r1-c9_Region5_1 43208460.tif #13
Tile_r1-c8_Region5_1 43208460.tif #12
Tile_r1-c12_Region5_143208460.tif #10
Tile_r1-c6_Region5_1 43208460.tif #14
Task-Manager
Datei Optionen Ansicht
Prozesse Leistung App-Verlauf Autostart Benutzer Details Dienste
CPU 100% 3,70 GHz
Arbeitsspeicher 72/128 GB (56%)
Datenträger 0 HDD 0%
Datenträger 1 HDD 0%
Datenträger 2 SSD 0%
Ethernet Ethernet 2 Ges.: 0 Empf.: 0 KBit
GPU 0 NVIDIA Quadro RTX 1% (43 °C)
CPU Intel(R) Xeon(R) W-2255 CPU @ 3.70GHz
% Auslastung 100%
60 Sekunden 0
Auslastung 100%
Geschwindigkeit 3,70 GHz
Prozesse 163
Threads 2473
Handles 69971
Betriebszeit 0:00:21:01
Basisgeschwindigkeit: 3,70 GHz
Sockets: 1
Kerne: 10
Logische Prozessoren: 20
Virtualisierung: Aktiviert
L1-Cache: 640 KB
L2-Cache: 10,0 MB
L3-Cache: 19,2 MB
Weniger Details
Ressourcenmonitor öffnen
(Fiji Is Just) ImageJ
File Edit Image Process Analyze Plugins Window Help
Processing... Montaging layers - 9' 53"
Log
File Edit Font
756: 9.12321756448203 129.47646157981652
757: 9.116342264841975 129.47646157981652
758: 9.109485034654783 129.47646157981652
759: 9.102645835427657 129.47646157981652
760: 9.09582456937219 129.47646157981652
761: 9.089021177904296 129.47646157981652
762: 9.082235510604168 129.47646157981652
763: 9.075467686224886 129.47646157981652
764: 9.06871750459265 129.47646157981652
765: 9.061984939367148 129.47646157981652
766: 9.055269798797376 129.47646157981652
767: 9.048572130079714 129.47646157981652
768: 9.041891858403753 129.47646157981652
769: 9.035229001525176 129.47646157981652
770: 9.028583262748848 129.47646157981652
771: 9.021954796259372 129.47646157981652
772: 9.015343451509368 129.47646157981652
773: 9.008749104256058 129.47646157981652
774: 9.00217177400558 129.47646157981652
775: 8.995611328577043 129.47646157981652
776: 8.989067776818542 129.47646157981652
777: 8.98254106832488 129.47646157981652
778: 8.976031130821504 129.47646157981652
779: 8.969537839554713 129.47646157981652
780: 8.963061170151727 129.47646157981652
781: 8.956601002624861 129.47646157981652
782: 8.950157326754162 129.47646157981652
783: 8.943730086236405 129.47646157981652
784: 8.937319132100093 129.47646157981652
785: 8.93092446634623 129.47646157981652
786: 8.924546200256087 129.47646157981652
787: 8.918183972403451 129.47646157981652
788: 8.91183790037709 129.47646157981652
789: 8.905507862579665 129.47646157981652
790: 8.899193807361314 129.47646157981652
791: 8.892895618696649 129.47646157981652
792: 8.886613367953734 129.47646157981652
793: 8.880346856837088 129.47646157981652
794: 8.874096067466668 129.47646157981652
795: 8.867861062791809 129.47646157981652
796: 8.861641658469178 129.47646157981652
797: 8.855437869700943 129.47646157981652
798: 8.849249605729952 129.47646157981652
799: 8.843076792039298 129.47646157981652
800: 8.8369193374077 129.47646157981652
801: 8.830777199467486 129.47646157981652
802: 8.824650372333208 129.47646157981652
803: 8.818538822138208 129.47646157981652
804: 8.812442382984978 129.47646157981652
805: 8.806361087485595 129.47646157981652
Successfully optimized configuration of 145 tiles after 806 iterations:
average displacement: 3.911px
minimal displacement: 2.686px
maximal displacement: 9.377px
Montage done.
Montaging layer z=1.0

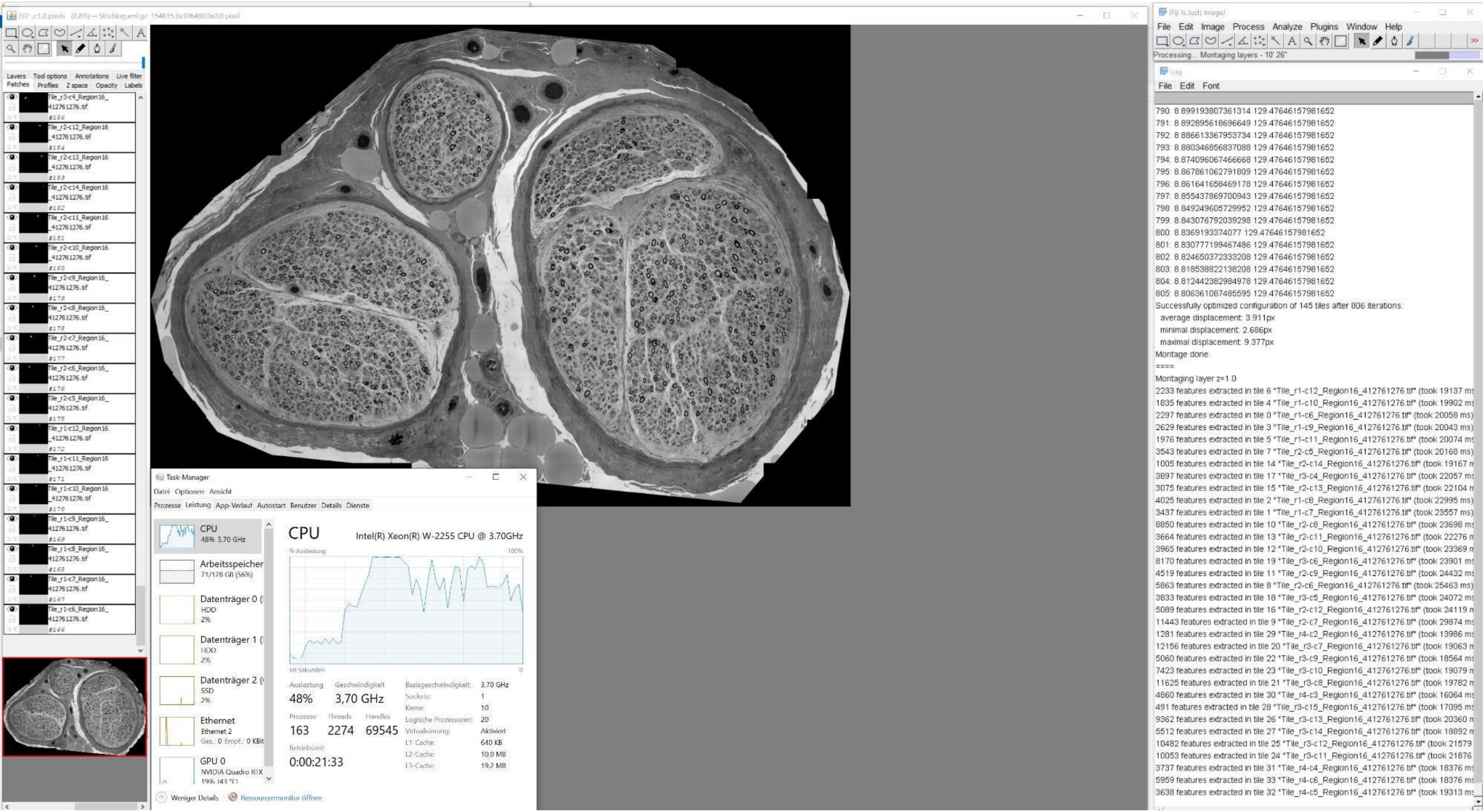
2/2 z:1.0 pixels (0.8%) -- Stitching.xml.gz 154615.0x106490.0x2.0 pixel
Layers Tool options Annotations Live filter
Patches Profiles Z space Opacity Labels
Tile_r3-c4_Region16_412761276.tif #186
Tile_r2-c12_Region16_412761276.tif #184
Tile_r2-c13_Region16_412761276.tif #183
Tile_r2-c14_Region16_412761276.tif #182
Tile_r2-c11_Region16_412761276.tif #181
Tile_r2-c10_Region16_412761276.tif #180
Tile_r2-c9_Region16_412761276.tif #179
Tile_r2-c8_Region16_412761276.tif #178
Tile_r2-c7_Region16_412761276.tif #177
Tile_r2-c6_Region16_412761276.tif #176
Tile_r2-c5_Region16_412761276.tif #175
Tile_r1-c12_Region16_412761276.tif #172
Tile_r1-c11_Region16_412761276.tif #171
Tile_r1-c10_Region16_412761276.tif #170
Tile_r1-c9_Region16_412761276.tif #169
Tile_r1-c8_Region16_412761276.tif #168
Tile_r1-c7_Region16_412761276.tif #167
Tile_r1-c6_Region16_412761276.tif #166
Task-Manager
Datei Optionen Ansicht
Prozesse Leistung App-Verlauf Autostart Benutzer Details Dienste
CPU 48% 3,70 GHz
Arbeitsspeicher 71/128 GB (56%)
Datenträger 0 HDD 2%
Datenträger 1 HDD 2%
Datenträger 2 SSD 2%
Ethernet Ethernet 2 Ges.: 0 Empf.: 0 KBit
GPU 0 NVIDIA Quadro RTX
CPU
Intel(R) Xeon(R) W-2255 CPU @ 3.70GHz
% Auslastung 100%
60 Sekunden 0
Auslastung 48%
Geschwindigkeit 3,70 GHz
Prozesse 163
Threads 2274
Handles 69545
Betriebszeit 0:00:21:33
Basisgeschwindigkeit: 3,70 GHz
Sockets: 1
Kerne: 10
Logische Prozessoren: 20
Virtualisierung: Aktiviert
L1-Cache: 640 KB
L2-Cache: 10,0 MB
L3-Cache: 19,2 MB
Weniger Details
Ressourcenmonitor öffnen
(Fiji Is Just) ImageJ
File Edit Image Process Analyze Plugins Window Help
Processing... Montaging layers - 10' 26"
Log
File Edit Font
790: 8.899193807361314 129.47646157981652
791: 8.892895618696649 129.47646157981652
792: 8.886613367953734 129.47646157981652
793: 8.880346856837088 129.47646157981652
794: 8.874096067466668 129.47646157981652
795: 8.867861062791809 129.47646157981652
796: 8.861641658469178 129.47646157981652
797: 8.855437869700943 129.47646157981652
798: 8.849249605729952 129.47646157981652
799: 8.843076792039298 129.47646157981652
800: 8.8369193374077 129.47646157981652
801: 8.830777199467486 129.47646157981652
802: 8.824650372333208 129.47646157981652
803: 8.818538822138208 129.47646157981652
804: 8.812442382984978 129.47646157981652
805: 8.806361087485595 129.47646157981652
Successfully optimized configuration of 145 tiles after 806 iterations:
average displacement: 3.911px
minimal displacement: 2.686px
maximal displacement: 9.377px
Montage done.
====
Montaging layer z=1.0
2233 features extracted in tile 6 "Tile_r1-c12_Region16_412761276.tif" (took 19137 ms
1835 features extracted in tile 4 "Tile_r1-c10_Region16_412761276.tif" (took 19902 ms
2297 features extracted in tile 0 "Tile_r1-c6_Region16_412761276.tif" (took 20058 ms)
2629 features extracted in tile 3 "Tile_r1-c9_Region16_412761276.tif" (took 20043 ms)
1976 features extracted in tile 5 "Tile_r1-c11_Region16_412761276.tif" (took 20074 ms
3543 features extracted in tile 7 "Tile_r2-c5_Region16_412761276.tif" (took 20168 ms)
1005 features extracted in tile 14 "Tile_r2-c14_Region16_412761276.tif" (took 19167 n
3897 features extracted in tile 17 "Tile_r3-c4_Region16_412761276.tif" (took 22057 ms
3075 features extracted in tile 15 "Tile_r2-c13_Region16_412761276.tif" (took 22104 n
4025 features extracted in tile 2 "Tile_r1-c8_Region16_412761276.tif" (took 22995 ms)
3437 features extracted in tile 1 "Tile_r1-c7_Region16_412761276.tif" (took 23557 ms)
8850 features extracted in tile 10 "Tile_r2-c8_Region16_412761276.tif" (took 23698 ms
3664 features extracted in tile 13 "Tile_r2-c11_Region16_412761276.tif" (took 22276 n
3965 features extracted in tile 12 "Tile_r2-c10_Region16_412761276.tif" (took 23369 n
8170 features extracted in tile 19 "Tile_r3-c6_Region16_412761276.tif" (took 23901 ms
4519 features extracted in tile 11 "Tile_r2-c9_Region16_412761276.tif" (took 24432 ms
5863 features extracted in tile 8 "Tile_r2-c6_Region16_412761276.tif" (took 25463 ms)
3833 features extracted in tile 18 "Tile_r3-c5_Region16_412761276.tif" (took 24072 ms
5089 features extracted in tile 16 "Tile_r2-c12_Region16_412761276.tif" (took 24119 n
11443 features extracted in tile 9 "Tile_r2-c7_Region16_412761276.tif" (took 29874 ms
1281 features extracted in tile 29 "Tile_r4-c2_Region16_412761276.tif" (took 13986 ms
12156 features extracted in tile 20 "Tile_r3-c7_Region16_412761276.tif" (took 19063 n
5060 features extracted in tile 22 "Tile_r3-c9_Region16_412761276.tif" (took 18564 ms
7423 features extracted in tile 23 "Tile_r3-c10_Region16_412761276.tif" (took 19079 n
11625 features extracted in tile 21 "Tile_r3-c8_Region16_412761276.tif" (took 19782 n
4860 features extracted in tile 30 "Tile_r4-c3_Region16_412761276.tif" (took 16064 ms
491 features extracted in tile 28 "Tile_r3-c15_Region16_412761276.tif" (took 17095 ms
9362 features extracted in tile 26 "Tile_r3-c13_Region16_412761276.tif" (took 20360 n
5512 features extracted in tile 27 "Tile_r3-c14_Region16_412761276.tif" (took 18892 n
10482 features extracted in tile 25 "Tile_r3-c12_Region16_412761276.tif" (took 21579
10053 features extracted in tile 24 "Tile_r3-c11_Region16_412761276.tif" (took 21876
3737 features extracted in tile 31 "Tile_r4-c4_Region16_412761276.tif" (took 18376 ms
5959 features extracted in tile 33 "Tile_r4-c6_Region16_412761276.tif" (took 18376 ms
3638 features extracted in tile 32 "Tile_r4-c5_Region16_412761276.tif" (took 19313 ms

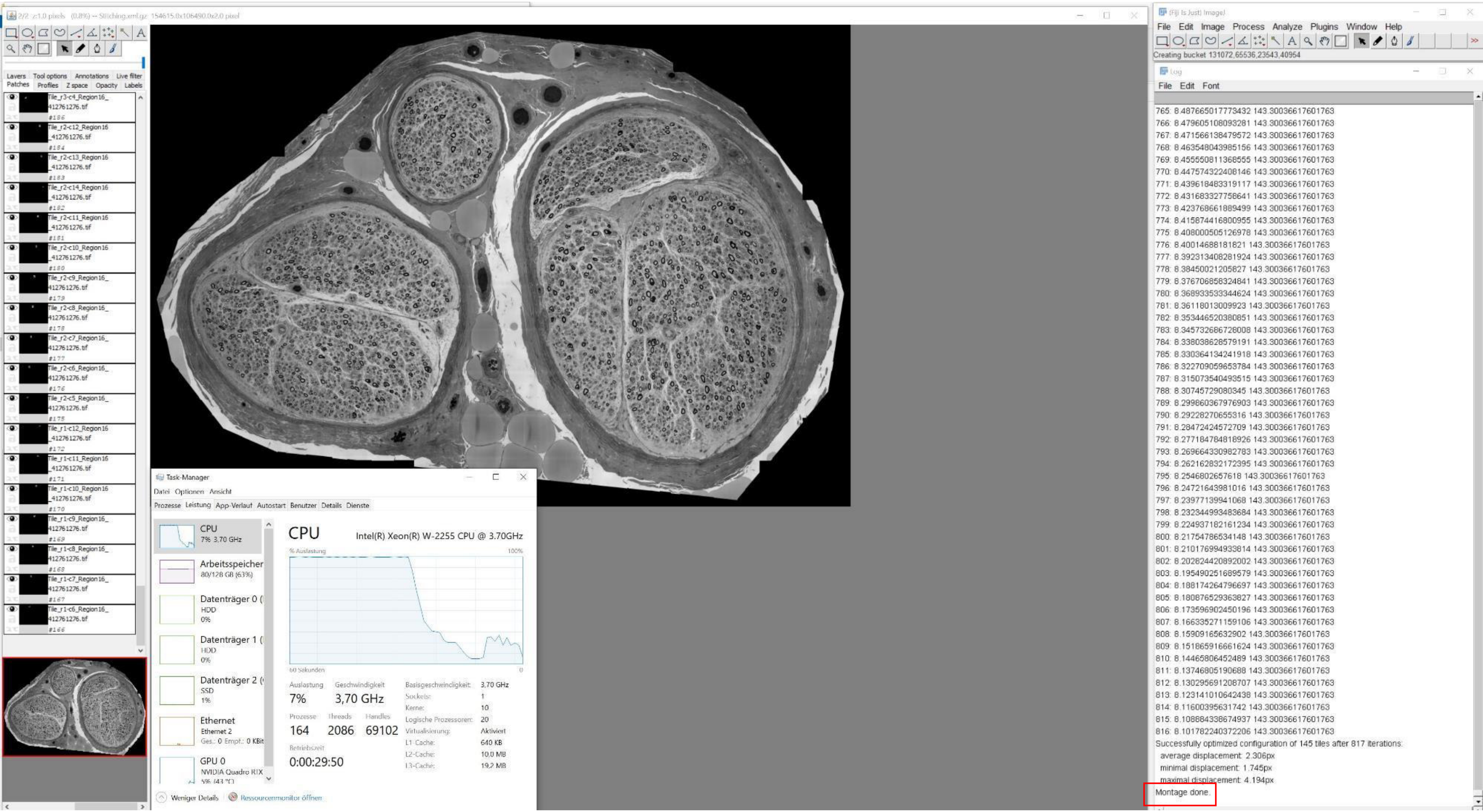
Task-Manager
Datei Optionen Ansicht
Prozesse Leistung App-Verlauf Autostart Benutzer Details Dienste
CPU
7% 3,70 GHz
Arbeitsspeicher
80/128 GB (63%)
Datenträger 0
HDD
0%
Datenträger 1
HDD
0%
Datenträger 2
SSD
1%
Ethernet
Ethernet 2
GPU 0
NVIDIA Quadro RTX
CPU
Intel(R) Xeon(R) W-2255 CPU @ 3.70GHz
% Auslastung
100%
60 Sekunden
0
Auslastung
7%
Geschwindigkeit
3,70 GHz
Prozesse
164
Threads
2086
Handles
69102
Betriebszeit
0:00:29:50
Basisgeschwindigkeit: 3,70 GHz
Sockets: 1
Kerne: 10
Logische Prozessoren: 20
Virtualisierung: Aktiviert
L1-Cache: 640 KB
L2-Cache: 10,0 MB
L3-Cache: 19,2 MB
Weniger Details
Ressourcenmonitor öffnen
(Fiji Is Just) ImageJ
File Edit Image Process Analyze Plugins Window Help
Creating bucket 131072,65536,23543,40954
Log
File Edit Font
765: 8.487665017773432 143.30036617601763
766: 8.479605108093281 143.30036617601763
767: 8.471566138479572 143.30036617601763
768: 8.463548043985156 143.30036617601763
769: 8.455550811368555 143.30036617601763
770: 8.447574322408146 143.30036617601763
771: 8.439618483319117 143.30036617601763
772: 8.431683327758641 143.30036617601763
773: 8.423768661889499 143.30036617601763
774: 8.415874416800955 143.30036617601763
775: 8.408000505126978 143.30036617601763
776: 8.40014688181821 143.30036617601763
777: 8.392313408281924 143.30036617601763
778: 8.38450021205827 143.30036617601763
779: 8.376706858324841 143.30036617601763
780: 8.368933533344624 143.30036617601763
781: 8.36118013009923 143.30036617601763
782: 8.353446520380851 143.30036617601763
783: 8.345732686728008 143.30036617601763
784: 8.338038628579191 143.30036617601763
785: 8.330364134241918 143.30036617601763
786: 8.322709059653784 143.30036617601763
787: 8.315073540493515 143.30036617601763
788: 8.30745729080345 143.30036617601763
789: 8.299860367976903 143.30036617601763
790: 8.29228270655316 143.30036617601763
791: 8.28472424572709 143.30036617601763
792: 8.277184784818926 143.30036617601763
793: 8.269664330982783 143.30036617601763
794: 8.262162832172395 143.30036617601763
795: 8.254680265761 8 143.30036617601763
796: 8.24721643981016 143.30036617601763
797: 8.23977139941068 143.30036617601763
798: 8.232344993483684 143.30036617601763
799: 8.224937182161234 143.30036617601763
800: 8.21754786534148 143.30036617601763
801: 8.210176994933814 143.30036617601763
802: 8.202824420892002 143.30036617601763
803: 8.195490251689579 143.30036617601763
804: 8.188174264796697 143.30036617601763
805: 8.180876529363827 143.30036617601763
806: 8.173596902450196 143.30036617601763
807: 8.166335271159106 143.30036617601763
808: 8.15909165632902 143.30036617601763
809: 8.151865916661624 143.30036617601763
810: 8.14465806452489 143.30036617601763
811: 8.13746805190688 143.30036617601763
812: 8.130295691208707 143.30036617601763
813: 8.123141010642438 143.30036617601763
814: 8.11600395631742 143.30036617601763
815: 8.108884338674937 143.30036617601763
816: 8.101782240372206 143.30036617601763
Successfully optimized configuration of 145 tiles after 817 iterations:
average displacement: 2.306px
minimal displacement: 1.745px
maximal displacement: 4.194px
Montage done.

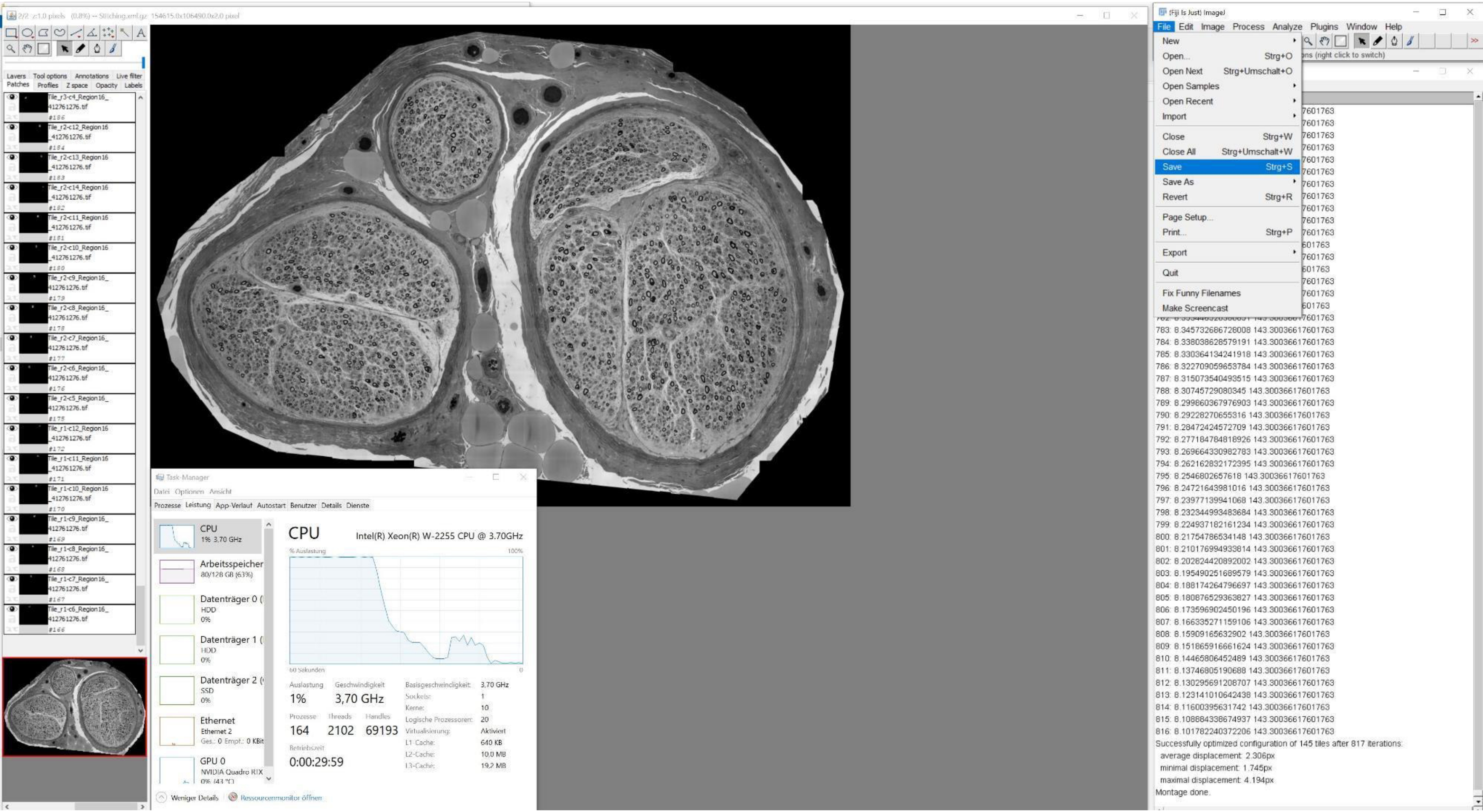
(Fiji Is Just) ImageJ
File Edit Image Process Analyze Plugins Window Help
New
Open... Strg+O
Open Next Strg+Umschalt+O
Open Samples
Open Recent
Import
Close Strg+W
Close All Strg+Umschalt+W
Save Strg+S
Save As
Revert Strg+R
Page Setup...
Print... Strg+P
Export
Quit
Fix Funny Filenames
Make Screencast
783: 8.345732686728008 143.30036617601763
784: 8.338038628579191 143.30036617601763
785: 8.330364134241918 143.30036617601763
786: 8.322709059653784 143.30036617601763
787: 8.315073540493515 143.30036617601763
788: 8.30745729080345 143.30036617601763
789: 8.299860367976903 143.30036617601763
790: 8.29228270655316 143.30036617601763
791: 8.28472424572709 143.30036617601763
792: 8.277184784818926 143.30036617601763
793: 8.269664330982783 143.30036617601763
794: 8.262162832172395 143.30036617601763
795: 8.2546802657618 143.30036617601763
796: 8.24721643981016 143.30036617601763
797: 8.23977139941068 143.30036617601763
798: 8.232344993483684 143.30036617601763
799: 8.224937182161234 143.30036617601763
800: 8.21754786534148 143.30036617601763
801: 8.210176994933814 143.30036617601763
802: 8.202824420892002 143.30036617601763
803: 8.195490251689579 143.30036617601763
804: 8.188174264796697 143.30036617601763
805: 8.180876529363827 143.30036617601763
806: 8.173596902450196 143.30036617601763
807: 8.166335271159106 143.30036617601763
808: 8.15909165632902 143.30036617601763
809: 8.151865916661624 143.30036617601763
810: 8.14465806452489 143.30036617601763
811: 8.13746805190688 143.30036617601763
812: 8.130295691208707 143.30036617601763
813: 8.123141010642438 143.30036617601763
814: 8.11600395631742 143.30036617601763
815: 8.108884338674937 143.30036617601763
816: 8.101782240372206 143.30036617601763
Successfully optimized configuration of 145 tiles after 817 iterations:
average displacement: 2.306px
minimal displacement: 1.745px
maximal displacement: 4.194px
Montage done.
Task-Manager
Datei Optionen Ansicht
Prozesse Leistung App-Verlauf Autostart Benutzer Details Dienste
CPU
1% 3,70 GHz
Arbeitsspeicher
80/128 GB (63%)
Datenträger 0
HDD
0%
Datenträger 1
HDD
0%
Datenträger 2
SSD
0%
Ethernet
Ethernet 2
GPU 0
NVIDIA Quadro RTX
CPU
Intel(R) Xeon(R) W-2255 CPU @ 3.70GHz
% Auslastung
100%
60 Sekunden
0
Auslastung 1%
Geschwindigkeit 3,70 GHz
Prozesse 164
Threads 2102
Handles 69193
Betriebszeit 0:00:29:59
Basisgeschwindigkeit: 3,70 GHz
Sockets: 1
Kerne: 10
Logische Prozessoren: 20
Virtualisierung: Aktiviert
L1-Cache: 640 KB
L2-Cache: 10,0 MB
L3-Cache: 19,2 MB
Weniger Details
Ressourcenmonitor öffnen
Layers Tool options Annotations Live filter
Patches Profiles Z space Opacity Labels
Tile_r3-c4_Region16_412761276.tif
Tile_r2-c12_Region16_412761276.tif
Tile_r2-c13_Region16_412761276.tif
Tile_r2-c14_Region16_412761276.tif
Tile_r2-c11_Region16_412761276.tif
Tile_r2-c10_Region16_412761276.tif
Tile_r2-c9_Region16_412761276.tif
Tile_r2-c8_Region16_412761276.tif
Tile_r2-c7_Region16_412761276.tif
Tile_r2-c6_Region16_412761276.tif
Tile_r2-c5_Region16_412761276.tif
Tile_r1-c12_Region16_412761276.tif
Tile_r1-c11_Region16_412761276.tif
Tile_r1-c10_Region16_412761276.tif
Tile_r1-c9_Region16_412761276.tif
Tile_r1-c8_Region16_412761276.tif
Tile_r1-c7_Region16_412761276.tif
Tile_r1-c6_Region16_412761276.tif

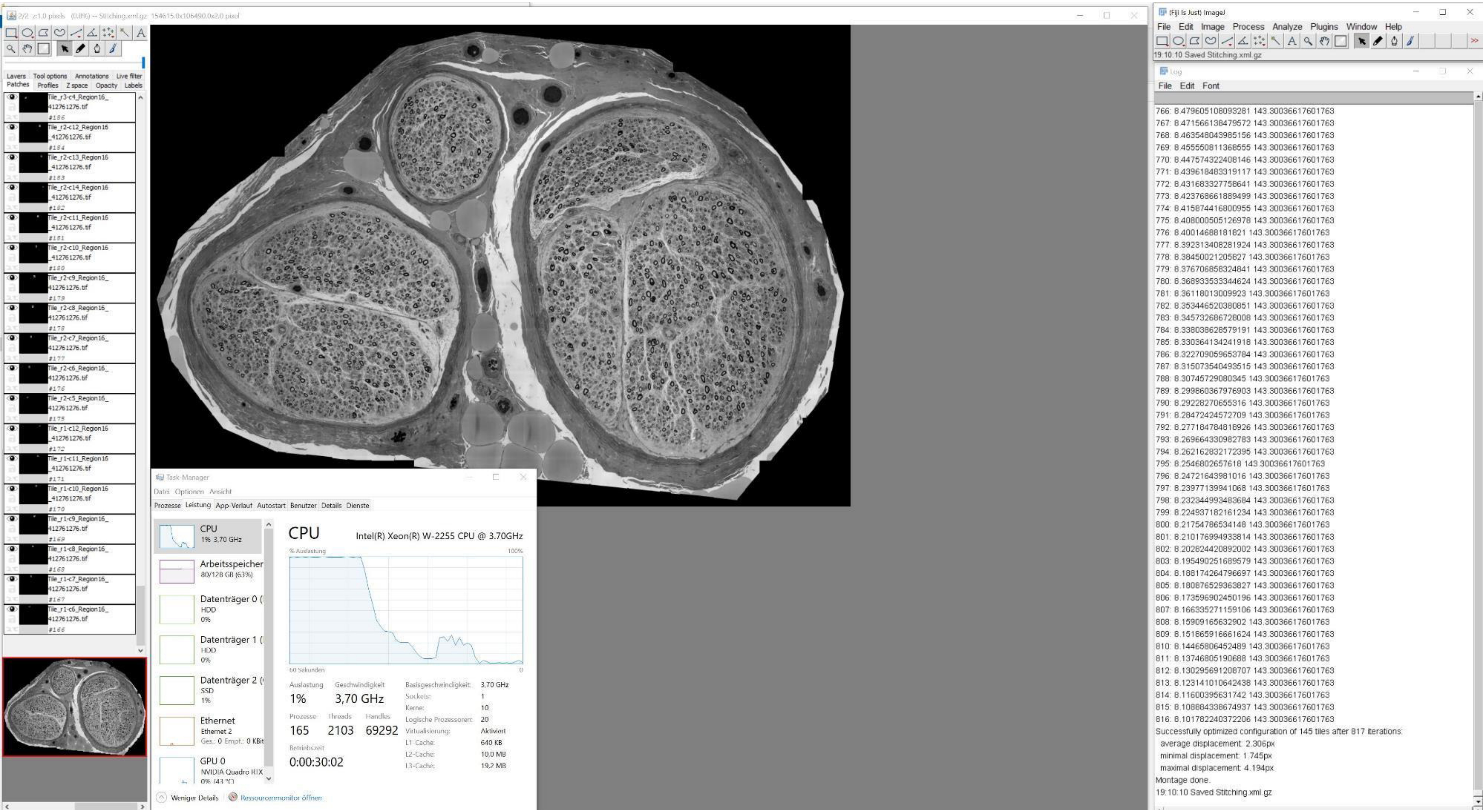
(Fiji Is Just) ImageJ
File Edit Image Process Analyze Plugins Window Help
19:10:10 Saved Stitching.xml.gz
Log
File Edit Font
766: 8.479605108093281 143.30036617601763
767: 8.471566138479572 143.30036617601763
768: 8.463548043985156 143.30036617601763
769: 8.455550811368555 143.30036617601763
770: 8.447574322408146 143.30036617601763
771: 8.439618483319117 143.30036617601763
772: 8.431683327758641 143.30036617601763
773: 8.423768661889499 143.30036617601763
774: 8.415874416800955 143.30036617601763
775: 8.408000505126978 143.30036617601763
776: 8.40014688181821 143.30036617601763
777: 8.392313408281924 143.30036617601763
778: 8.38450021205827 143.30036617601763
779: 8.376706858324841 143.30036617601763
780: 8.368933533344624 143.30036617601763
781: 8.36118013009923 143.30036617601763
782: 8.353446520380851 143.30036617601763
783: 8.345732686728008 143.30036617601763
784: 8.338038628579191 143.30036617601763
785: 8.330364134241918 143.30036617601763
786: 8.322709059653784 143.30036617601763
787: 8.315073540493515 143.30036617601763
788: 8.30745729080345 143.30036617601763
789: 8.299860367976903 143.30036617601763
790: 8.29228270655316 143.30036617601763
791: 8.28472424572709 143.30036617601763
792: 8.277184784818926 143.30036617601763
793: 8.269664330982783 143.30036617601763
794: 8.262162832172395 143.30036617601763
795: 8.2546802657618 143.30036617601763
796: 8.24721643981016 143.30036617601763
797: 8.23977139941068 143.30036617601763
798: 8.232344993483684 143.30036617601763
799: 8.224937182161234 143.30036617601763
800: 8.21754786534148 143.30036617601763
801: 8.210176994933814 143.30036617601763
802: 8.202824420892002 143.30036617601763
803: 8.195490251689579 143.30036617601763
804: 8.188174264796697 143.30036617601763
805: 8.180876529363827 143.30036617601763
806: 8.173596902450196 143.30036617601763
807: 8.166335271159106 143.30036617601763
808: 8.15909165632902 143.30036617601763
809: 8.151865916661624 143.30036617601763
810: 8.14465806452489 143.30036617601763
811: 8.13746805190688 143.30036617601763
812: 8.130295691208707 143.30036617601763
813: 8.123141010642438 143.30036617601763
814: 8.11600395631742 143.30036617601763
815: 8.108884338674937 143.30036617601763
816: 8.101782240372206 143.30036617601763
Successfully optimized configuration of 145 tiles after 817 iterations:
average displacement: 2.306px
minimal displacement: 1.745px
maximal displacement: 4.194px
Montage done.
19:10:10 Saved Stitching.xml.gz
Task-Manager
Datei Optionen Ansicht
Prozesse Leistung App-Verlauf Autostart Benutzer Details Dienste
CPU
1% 3,70 GHz
Arbeitsspeicher
80/128 GB (63%)
Datenträger 0
HDD
0%
Datenträger 1
HDD
0%
Datenträger 2
SSD
1%
Ethernet
Ethernet 2
GPU 0
NVIDIA Quadro RTX
CPU
Intel(R) Xeon(R) W-2255 CPU @ 3.70GHz
% Auslastung
100%
60 Sekunden
0
Auslastung
1%
Geschwindigkeit
3,70 GHz
Prozesse
165
Threads
2103
Handles
69292
Betriebszeit
0:00:30:02
Basisgeschwindigkeit: 3,70 GHz
Sockets: 1
Kerne: 10
Logische Prozessoren: 20
Virtualisierung: Aktiviert
L1-Cache: 640 KB
L2-Cache: 10,0 MB
L3-Cache: 19,2 MB
Weniger Details
Ressourcenmonitor öffnen
Layers Tool options Annotations Live filter
Patches Profiles Z space Opacity Labels

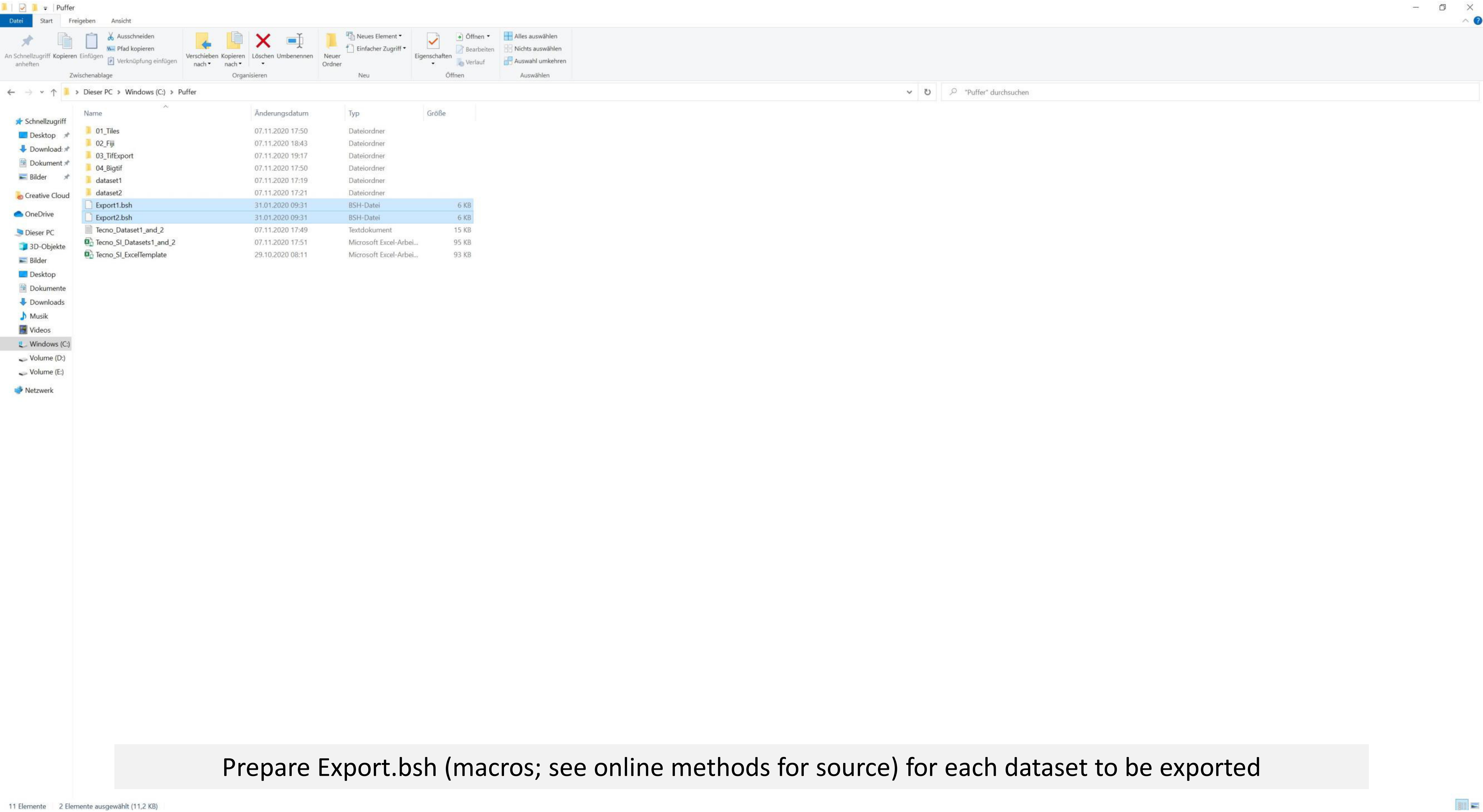


Prepare Export.bsh (macros; see online methods for source) for each dataset to be exported

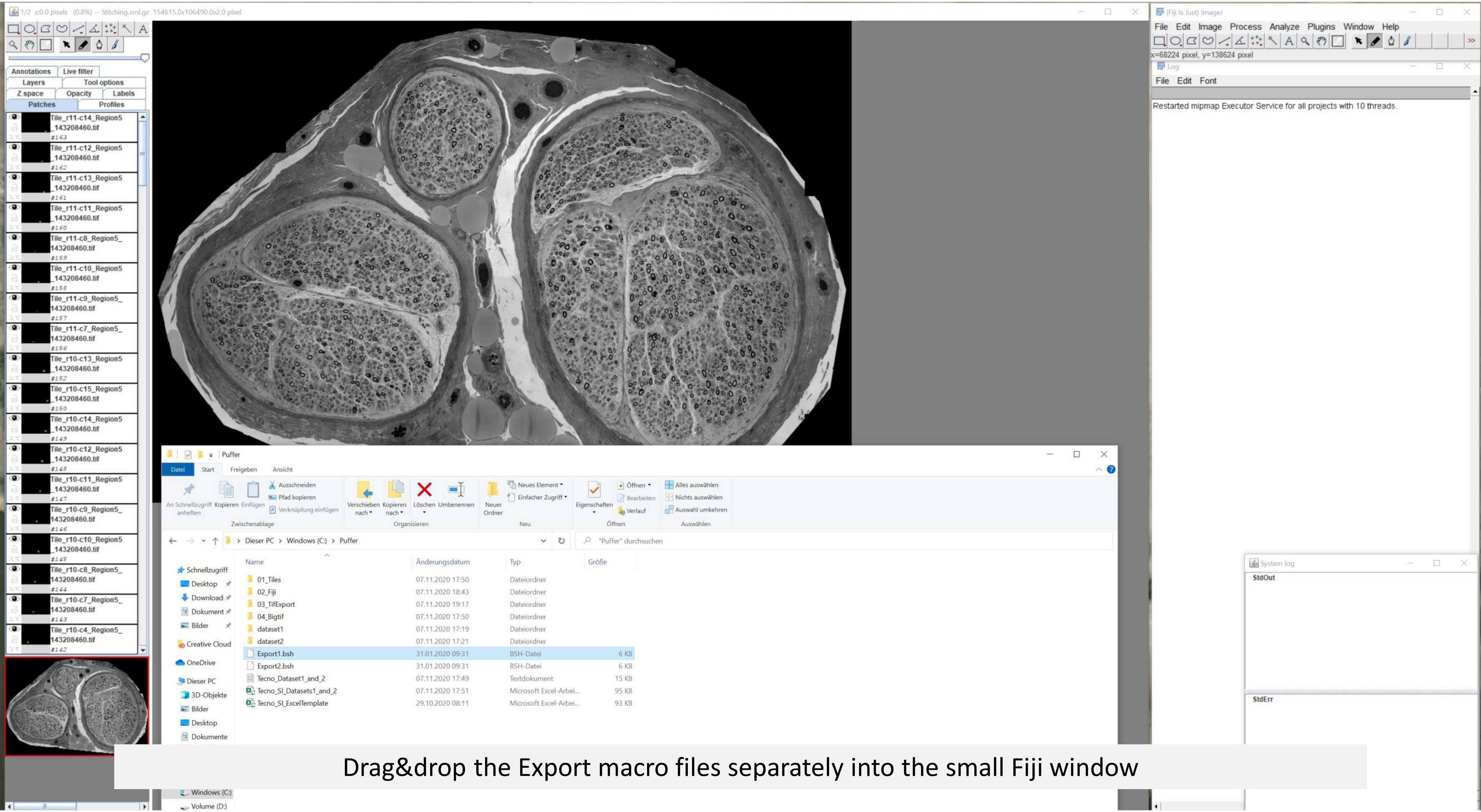


Drag&drop the Export macro files separately into the small Fiji window

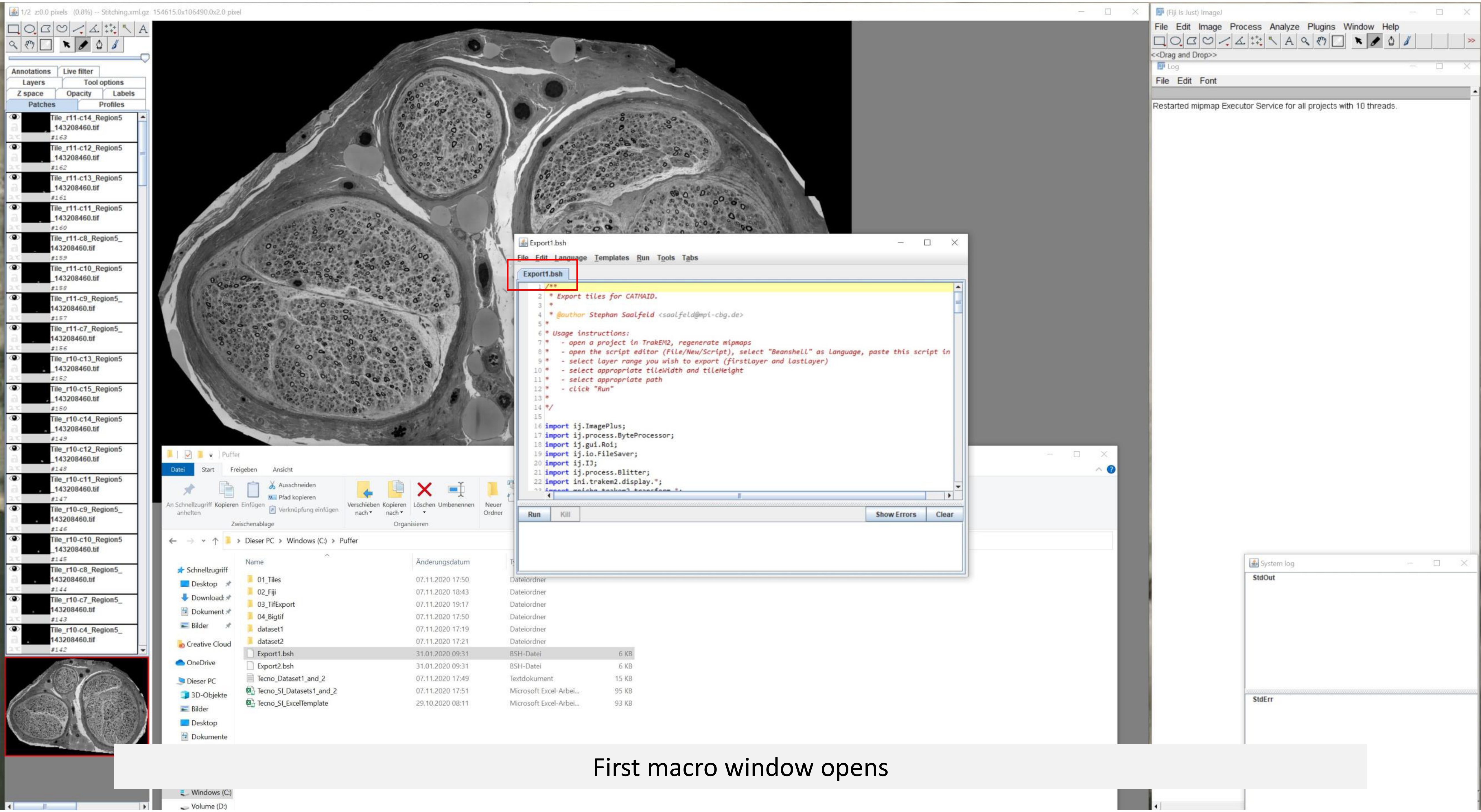


First macro window opens

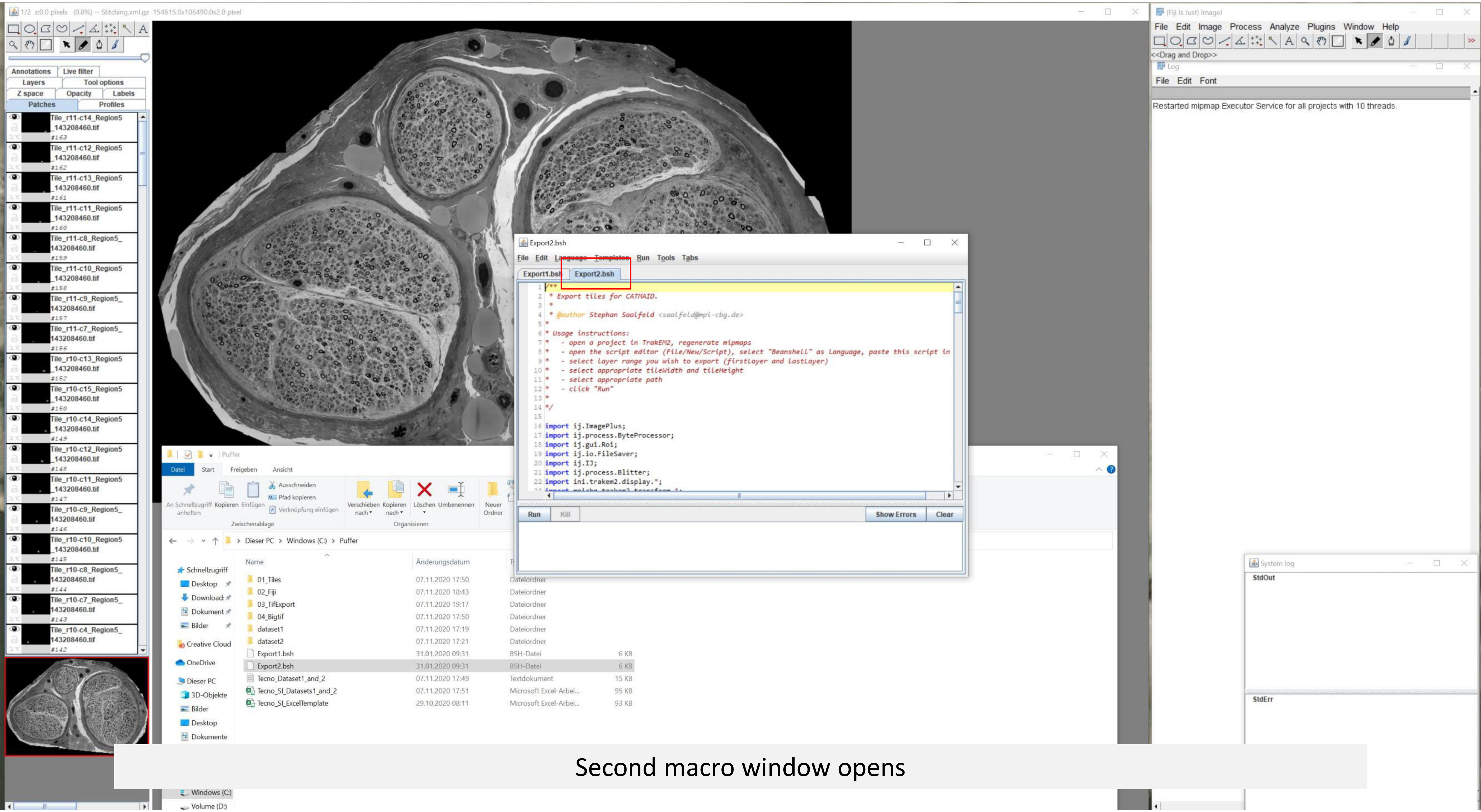


Second macro window opens

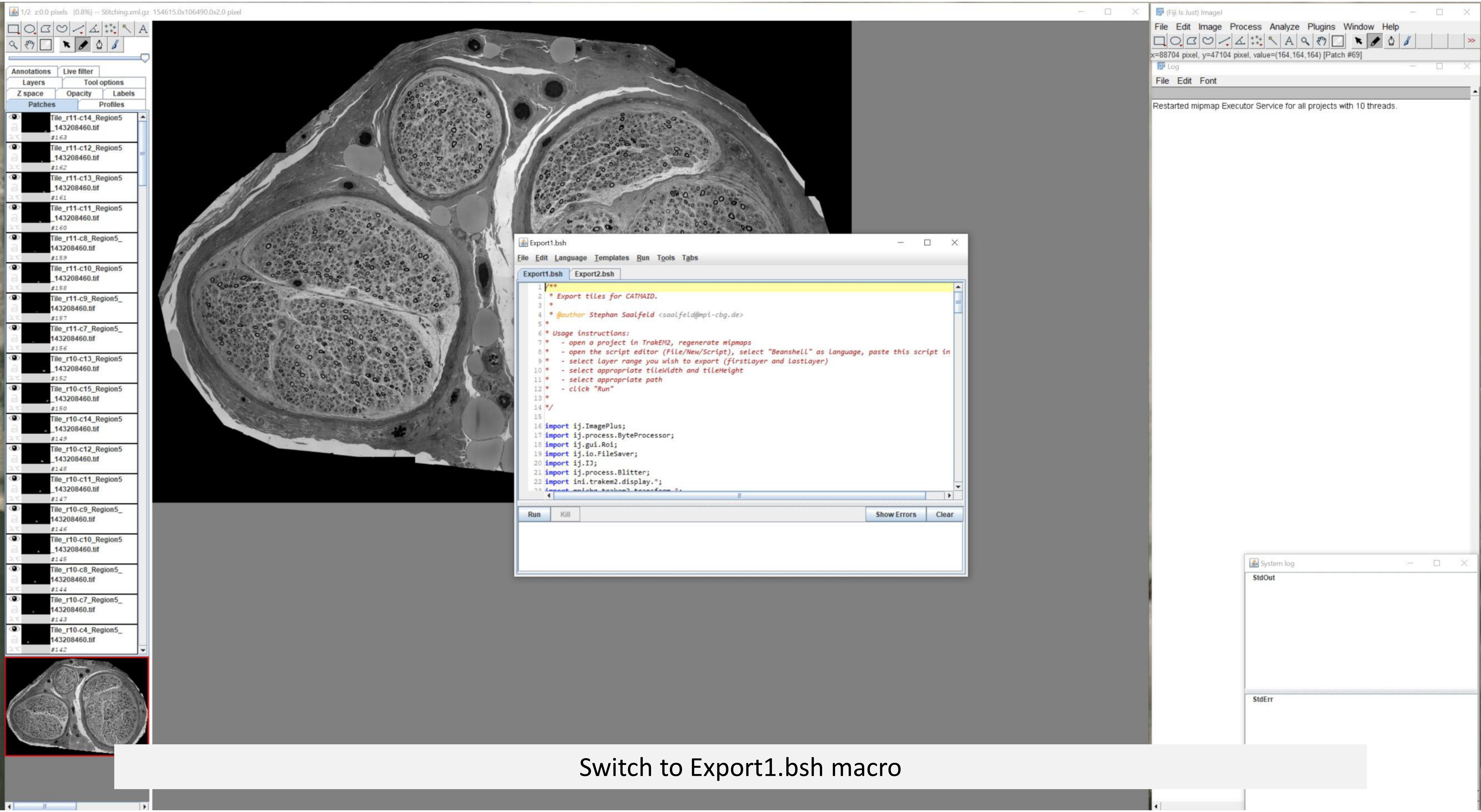


Switch to Export1.bsh macro

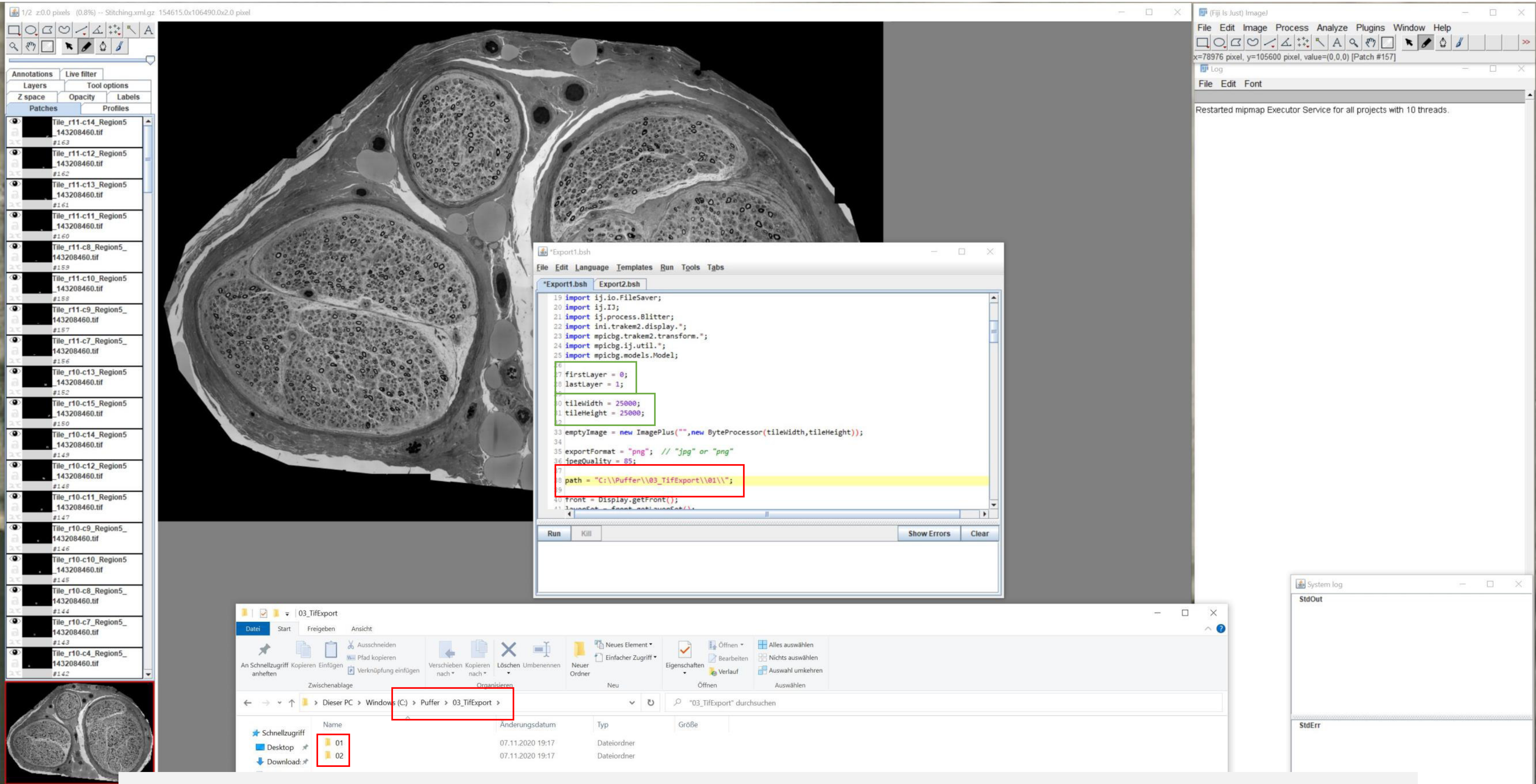


Prepare individual output folders for both datasets and select the respective path in the macros
See details for export parameter (green; layer, tile dimensions)

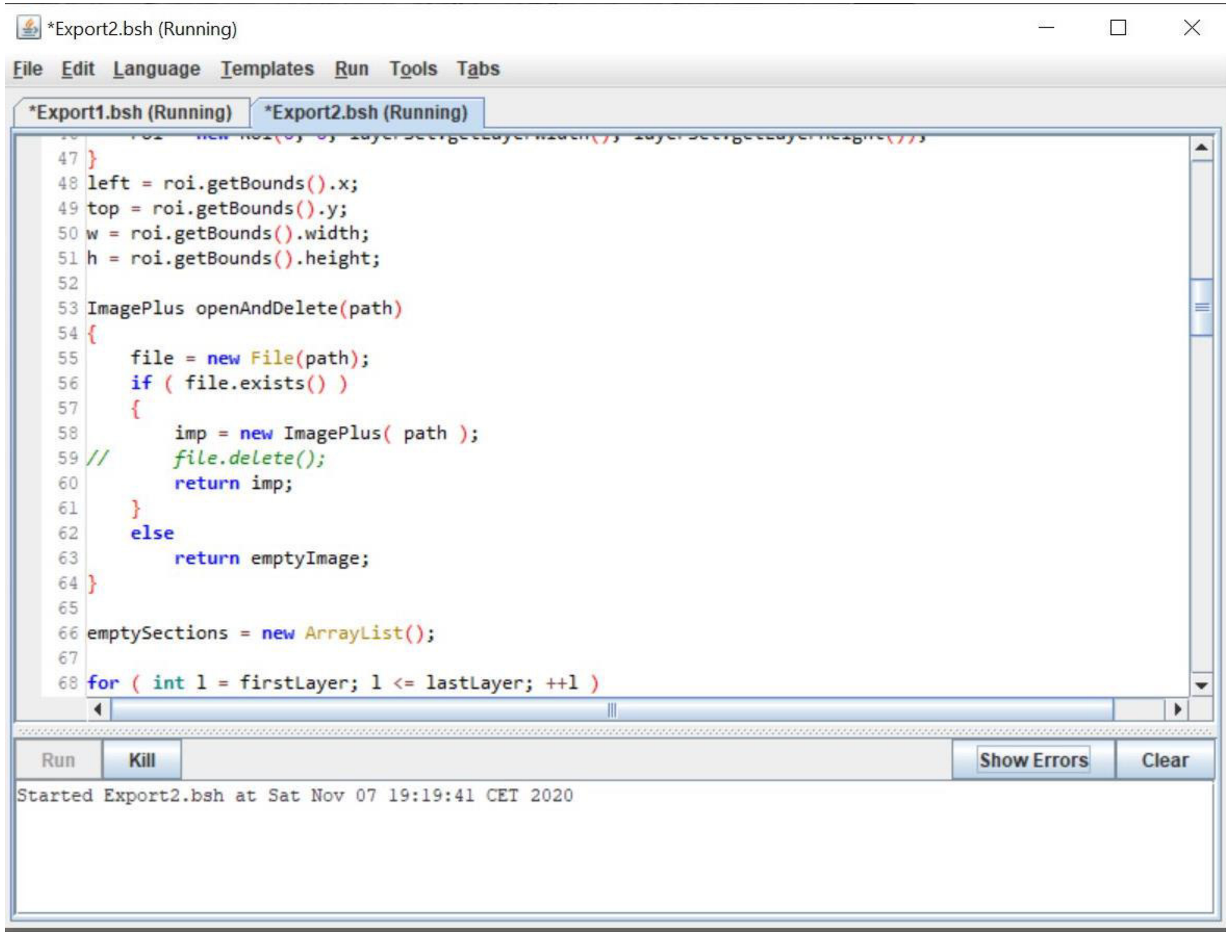


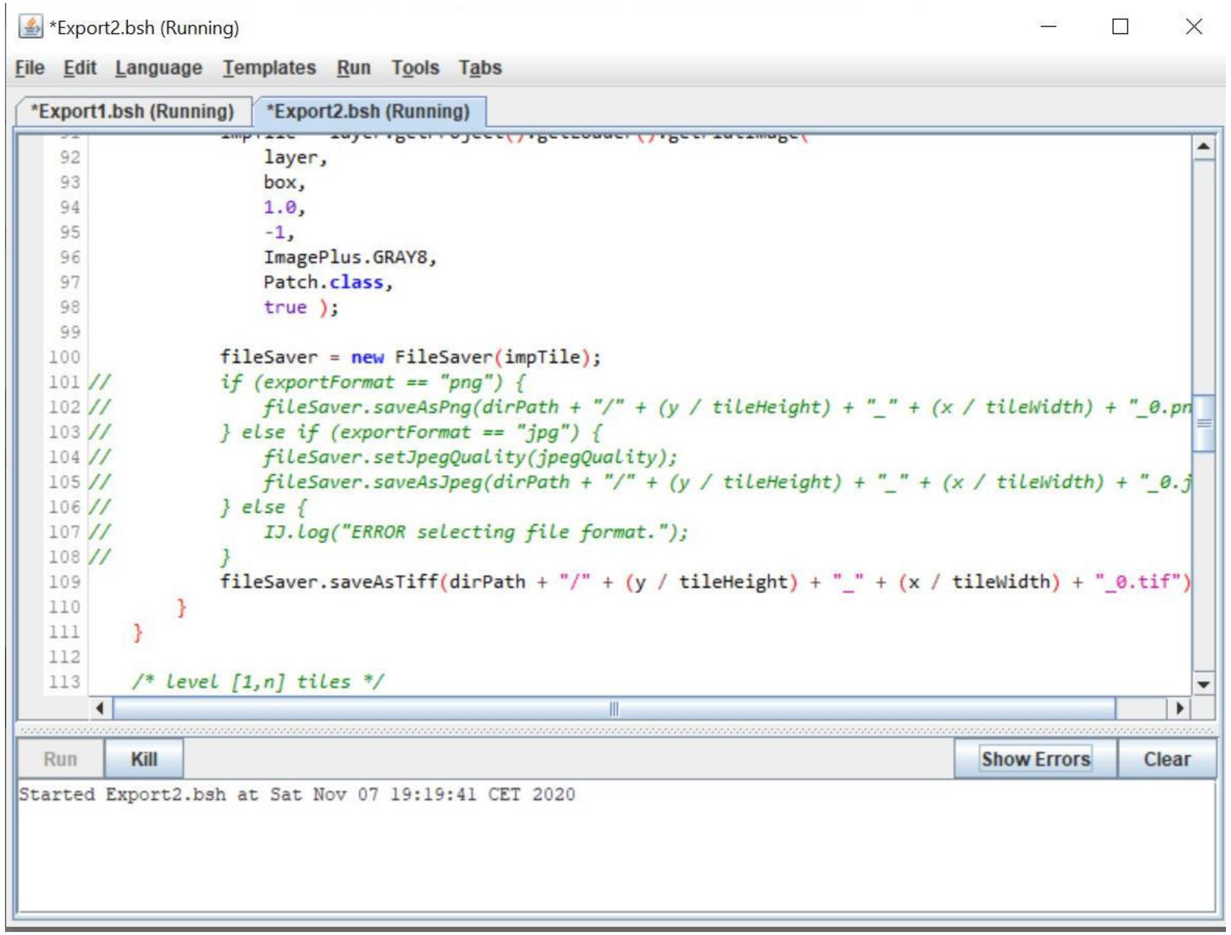


Modifications of the CATMAID macro to only export large tif tiles
Left; line 59 commented to avoid that the large tiles are deleted
Right; lines 101-108 commented to only export tif tiles

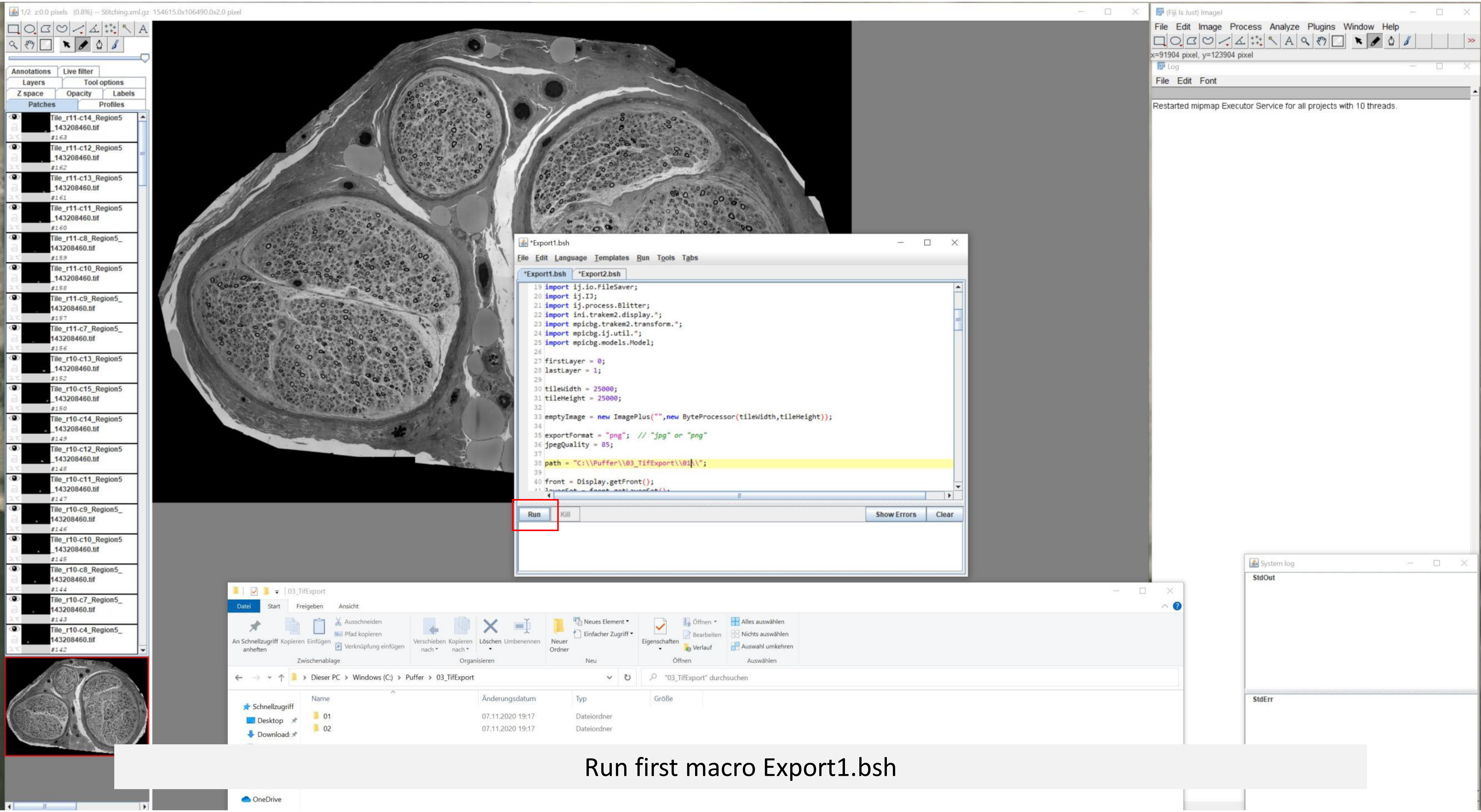


Run first macro Export1.bsh

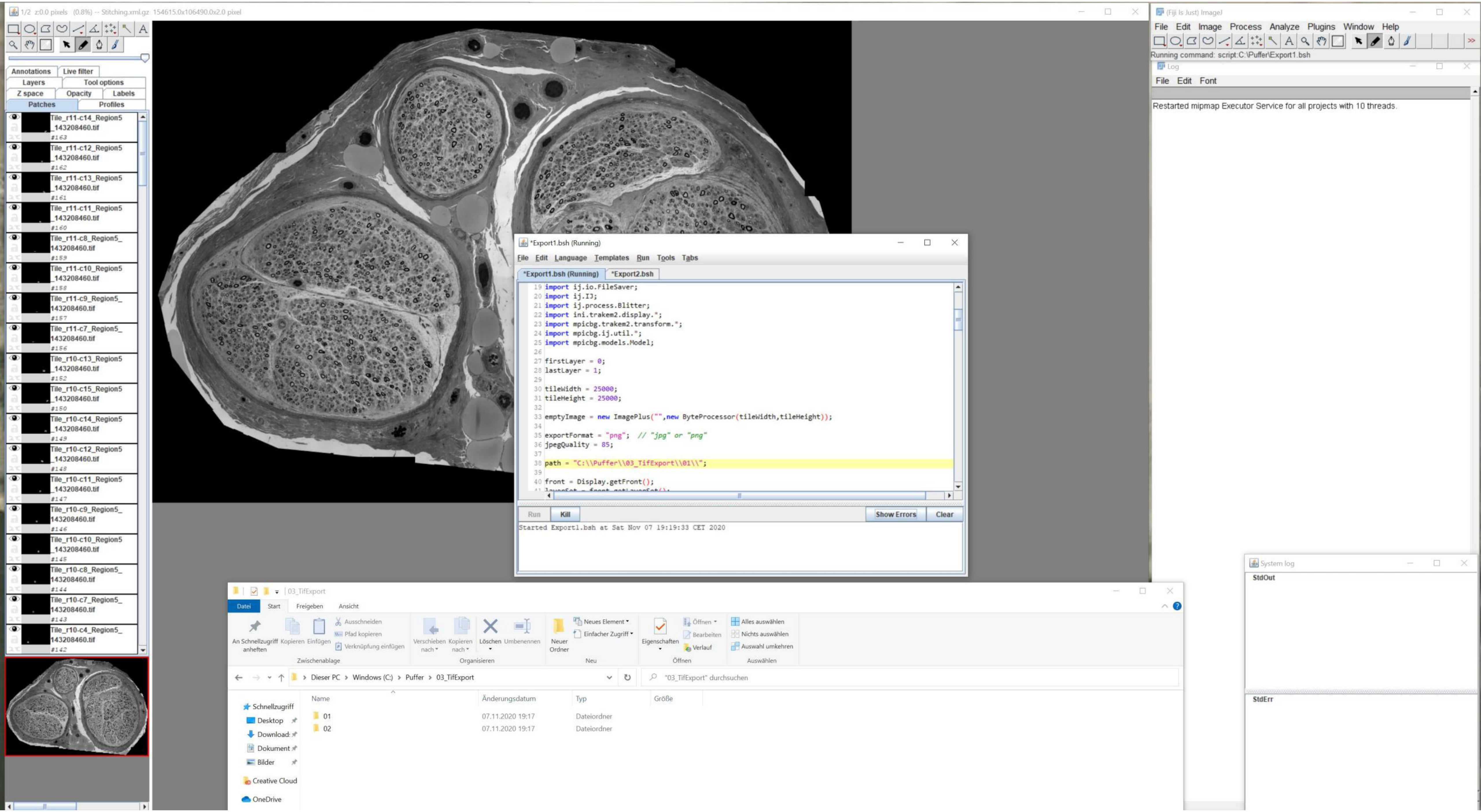

1/2 z:0.0 pixels (0.8%) -- Stitching.xml.gz 154615.0x106490.0x2.0 pixel
Annotations
Live filter
Layers
Tool options
Z space
Opacity
Labels
Patches
Profiles
Tile_r11-c14_Region5_143208460.tif #163
Tile_r11-c12_Region5_143208460.tif #162
Tile_r11-c13_Region5_143208460.tif #161
Tile_r11-c11_Region5_143208460.tif #160
Tile_r11-c8_Region5_143208460.tif #159
Tile_r11-c10_Region5_143208460.tif #158
Tile_r11-c9_Region5_143208460.tif #157
Tile_r11-c7_Region5_143208460.tif #156
Tile_r10-c13_Region5_143208460.tif #152
Tile_r10-c15_Region5_143208460.tif #150
Tile_r10-c14_Region5_143208460.tif #149
Tile_r10-c12_Region5_143208460.tif #148
Tile_r10-c11_Region5_143208460.tif #147
Tile_r10-c9_Region5_143208460.tif #146
Tile_r10-c10_Region5_143208460.tif #145
Tile_r10-c8_Region5_143208460.tif #144
Tile_r10-c7_Region5_143208460.tif #143
Tile_r10-c4_Region5_143208460.tif #142
*Export1.bsh (Running)
File Edit Language Templates Run Tools Tabs
*Export1.bsh (Running)
*Export2.bsh
19 import ij.io.FileSaver;
20 import ij.IJ;
21 import ij.process.Blitter;
22 import ini.trakem2.display.*;
23 import mpicbg.trakem2.transform.*;
24 import mpicbg.ij.util.*;
25 import mpicbg.models.Model;
27 firstLayer = 0;
28 lastLayer = 1;
30 tileWidth = 25000;
31 tileHeight = 25000;
33 emptyImage = new ImagePlus("",new ByteProcessor(tileWidth,tileHeight));
35 exportFormat = "png"; // "jpg" or "png"
36 jpegQuality = 85;
38 path = "C:\\Puffer\\03_TifExport\\01\\";
40 front = Display.getFront();
Run
Kill
Show Errors
Clear
Started Export1.bsh at Sat Nov 07 19:19:33 CET 2020
(Fiji Is Just) ImageJ
File Edit Image Process Analyze Plugins Window Help
Running command: script:C:\Puffer\Export1.bsh
Log
File Edit Font
Restarted mipmap Executor Service for all projects with 10 threads.
System log
StdOut
StdErr
03_TifExport
Datei Start Freigeben Ansicht
An Schnellzugriff anheften
Kopieren
Einfügen
Ausschneiden
Pfad kopieren
Verknüpfung einfügen
Zwischenablage
Verschieben nach
Kopieren nach
Löschen
Umbenennen
Organisieren
Neuer Ordner
Neues Element
Einfacher Zugriff
Neu
Eigenschaften
Öffnen
Bearbeiten
Verlauf
Öffnen
Alles auswählen
Nichts auswählen
Auswahl umkehren
Auswählen
Dieser PC > Windows (C:) > Puffer > 03_TifExport
"03_TifExport" durchsuchen
Schnellzugriff
Desktop
Download:
Dokument
Bilder
Creative Cloud
OneDrive
Name
Änderungsdatum
Typ
Größe
01
07.11.2020 19:17
Dateiordner
02
07.11.2020 19:17
Dateiordner

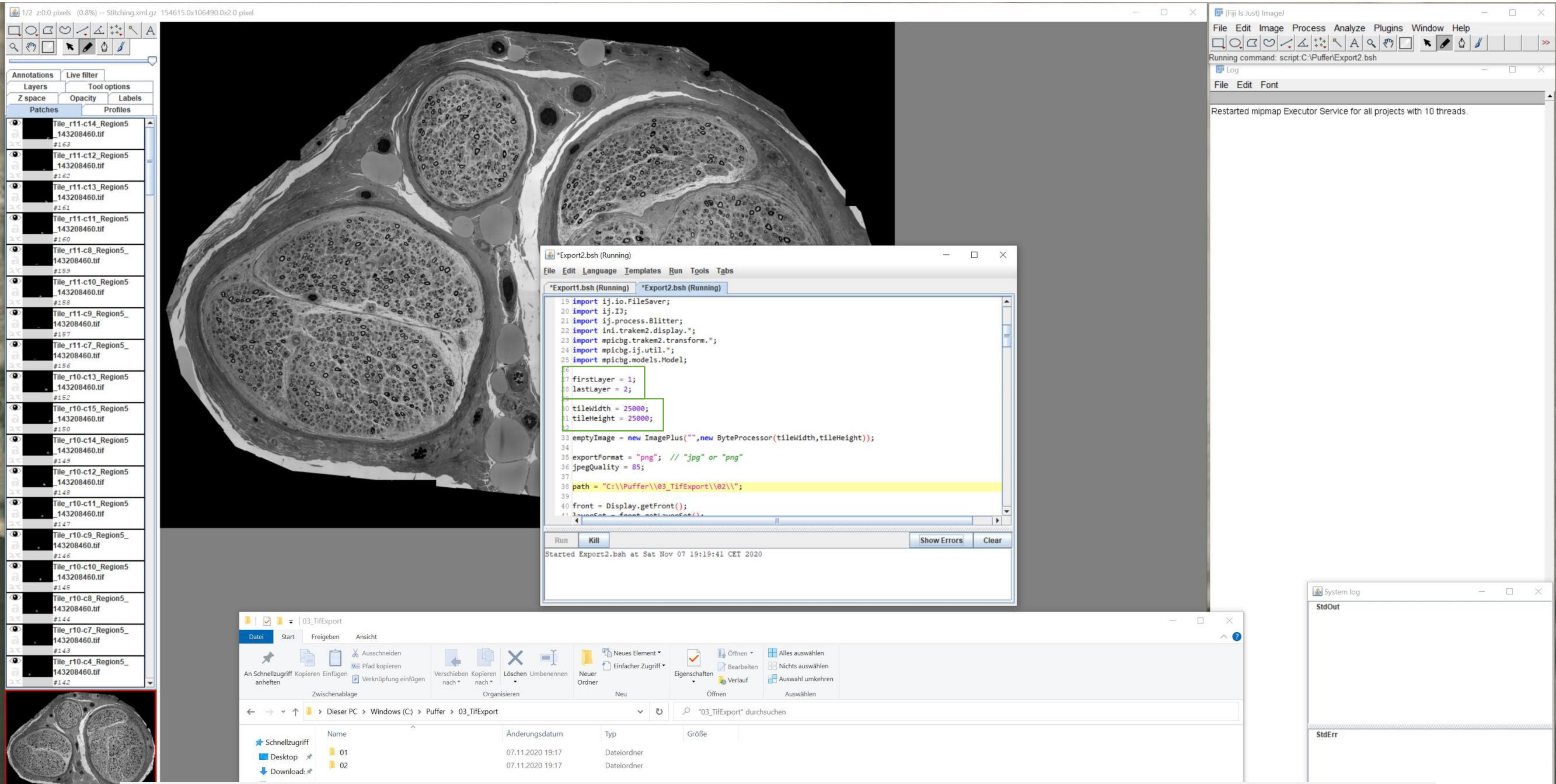


See details of the second macro
Run second macro Export2.bsh

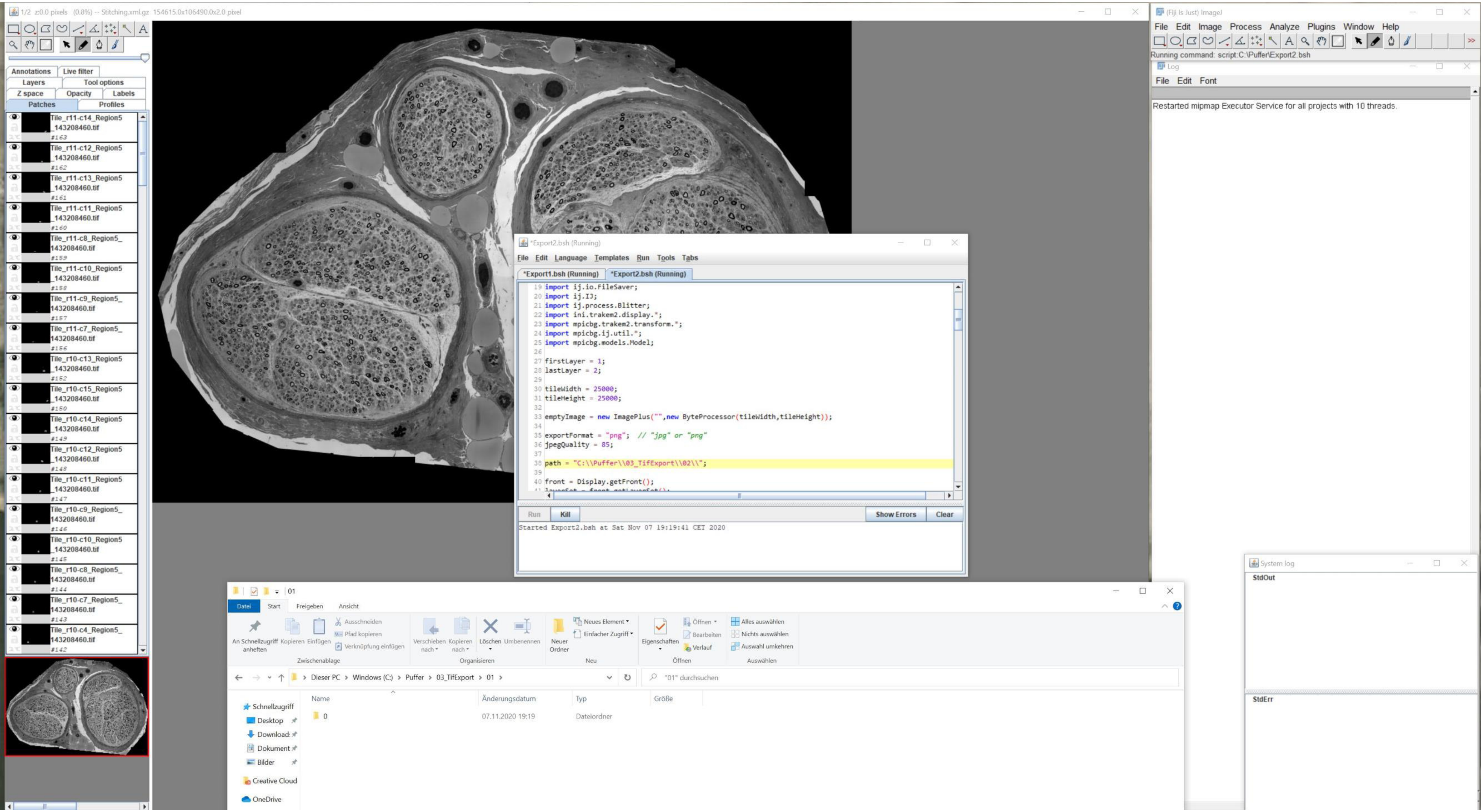
1/2 z:0.0 pixels (0.8%) -- Stitching.xml.gz 154615.0x106490.0x2.0 pixel
Annotations
Live filter
Layers
Tool options
Z space
Opacity
Labels
Patches
Profiles
Tile_r11-c14_Region5_143208460.tif
Tile_r11-c12_Region5_143208460.tif
Tile_r11-c13_Region5_143208460.tif
Tile_r11-c11_Region5_143208460.tif
Tile_r11-c8_Region5_143208460.tif
Tile_r11-c10_Region5_143208460.tif
Tile_r11-c9_Region5_143208460.tif
Tile_r11-c7_Region5_143208460.tif
Tile_r10-c13_Region5_143208460.tif
Tile_r10-c15_Region5_143208460.tif
Tile_r10-c14_Region5_143208460.tif
Tile_r10-c12_Region5_143208460.tif
Tile_r10-c11_Region5_143208460.tif
Tile_r10-c9_Region5_143208460.tif
Tile_r10-c10_Region5_143208460.tif
Tile_r10-c8_Region5_143208460.tif
Tile_r10-c7_Region5_143208460.tif
Tile_r10-c4_Region5_143208460.tif
*Export2.bsh (Running)
File Edit Language Templates Run Tools Tabs
*Export1.bsh (Running)
*Export2.bsh (Running)
import ij.io.FileSaver;
import ij.IJ;
import ij.process.Blitter;
import ini.trakem2.display.*;
import mpicbg.trakem2.transform.*;
import mpicbg.ij.util.*;
import mpicbg.models.Model;
firstLayer = 1;
lastLayer = 2;
tileWidth = 25000;
tileHeight = 25000;
emptyImage = new ImagePlus("",new ByteProcessor(tileWidth,tileHeight));
exportFormat = "png"; // "jpg" or "png"
jpegQuality = 85;
path = "C:\\Puffer\\03_TifExport\\02\\";
front = Display.getFront();
Run
Kill
Show Errors
Clear
Started Export2.bsh at Sat Nov 07 19:19:41 CET 2020
(Fiji Is Just) ImageJ
File Edit Image Process Analyze Plugins Window Help
Running command: script:C:\Puffer\Export2.bsh
Log
File Edit Font
Restarted mipmap Executor Service for all projects with 10 threads.
System log
StdOut
StdErr
01
Datei
Start
Freigeben
Ansicht
An Schnellzugriff anheften
Kopieren
Einfügen
Ausschneiden
Pfad kopieren
Verknüpfung einfügen
Zwischenablage
Verschieben nach
Kopieren nach
Löschen
Umbenennen
Organisieren
Neuer Ordner
Neues Element
Einfacher Zugriff
Neu
Eigenschaften
Öffnen
Bearbeiten
Verlauf
Öffnen
Alles auswählen
Nichts auswählen
Auswahl umkehren
Auswählen
Dieser PC › Windows (C:) › Puffer › 03_TifExport › 01 ›
"01" durchsuchen
Name
Änderungsdatum
Typ
Größe
Schnellzugriff
Desktop
Download
Dokument
Bilder
Creative Cloud
OneDrive
0
07.11.2020 19:19
Dateiordner

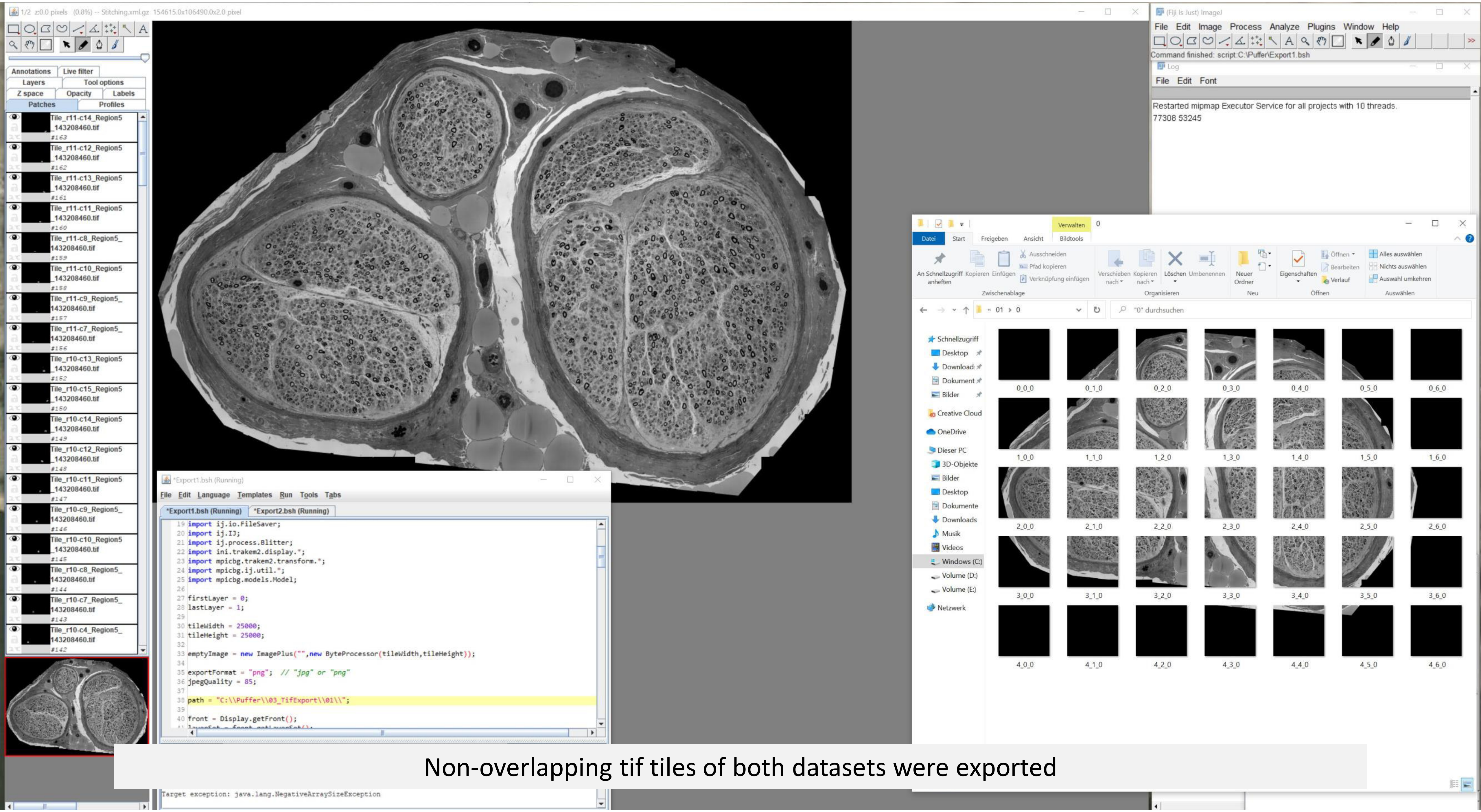


Non-overlapping tif tiles of both datasets were exported

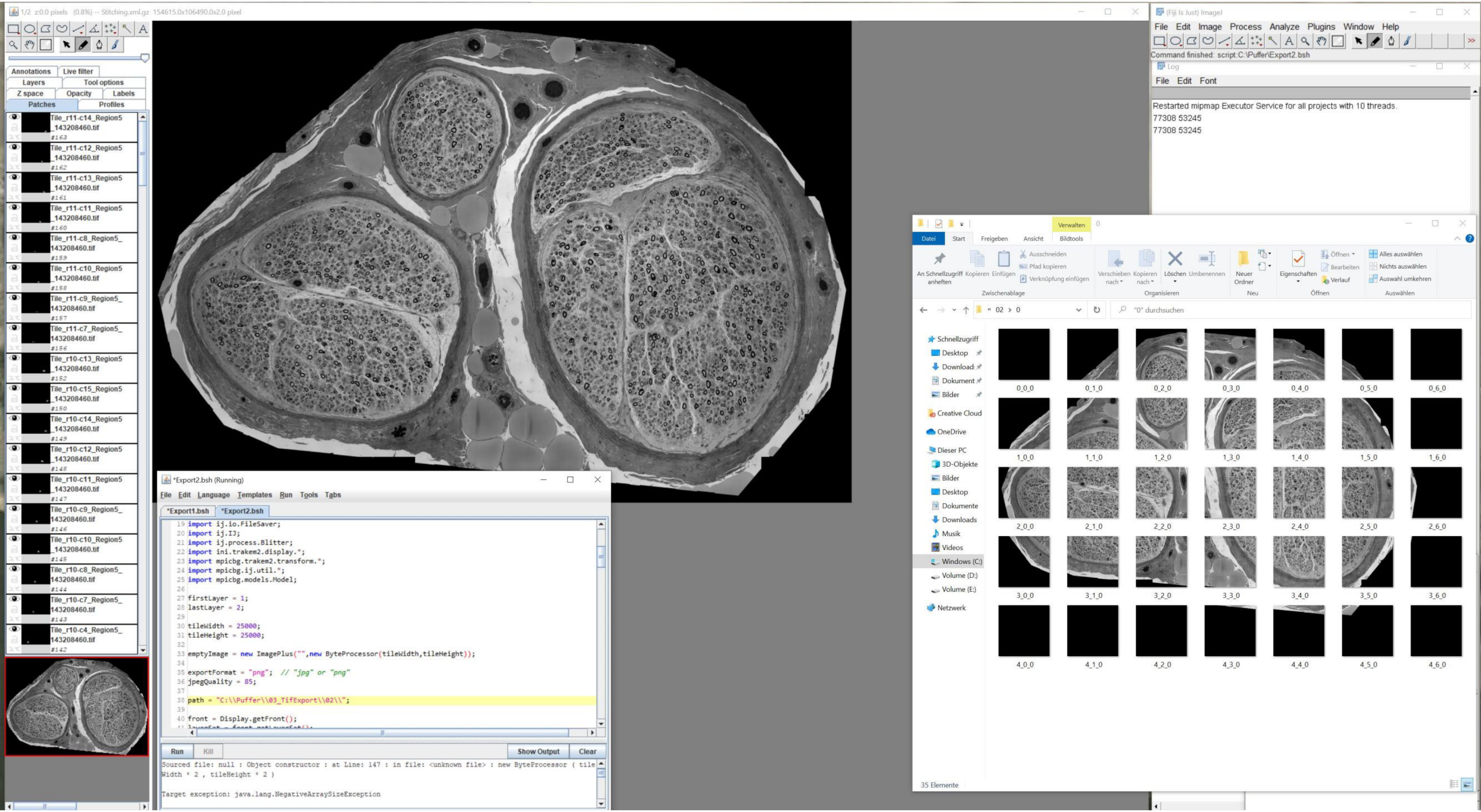
1/2 z:0.0 pixels (0.8%) -- Stitching.xml.gz 154615.0x106490.0x2.0 pixel
Annotations
Live filter
Layers
Tool options
Z space
Opacity
Labels
Patches
Profiles
Tile_r11-c14_Region5_143208460.tif
Tile_r11-c12_Region5_143208460.tif
Tile_r11-c13_Region5_143208460.tif
Tile_r11-c11_Region5_143208460.tif
Tile_r11-c8_Region5_143208460.tif
Tile_r11-c10_Region5_143208460.tif
Tile_r11-c9_Region5_143208460.tif
Tile_r11-c7_Region5_143208460.tif
Tile_r10-c13_Region5_143208460.tif
Tile_r10-c15_Region5_143208460.tif
Tile_r10-c14_Region5_143208460.tif
Tile_r10-c12_Region5_143208460.tif
Tile_r10-c11_Region5_143208460.tif
Tile_r10-c9_Region5_143208460.tif
Tile_r10-c10_Region5_143208460.tif
Tile_r10-c8_Region5_143208460.tif
Tile_r10-c7_Region5_143208460.tif
Tile_r10-c4_Region5_143208460.tif
(Fiji Is Just) ImageJ
File Edit Image Process Analyze Plugins Window Help
Command finished: script:C:\Puffer\Export2.bsh
Log
File Edit Font
Restarted mipmap Executor Service for all projects with 10 threads.
77308 53245
77308 53245
*Export2.bsh (Running)
File Edit Language Templates Run Tools Tabs
*Export1.bsh
*Export2.bsh
import ij.io.FileSaver;
import ij.IJ;
import ij.process.Blitter;
import ini.trakem2.display.*;
import mpicbg.trakem2.transform.*;
import mpicbg.ij.util.*;
import mpicbg.models.Model;
firstLayer = 1;
lastLayer = 2;
tileWidth = 25000;
tileHeight = 25000;
emptyImage = new ImagePlus("",new ByteProcessor(tileWidth,tileHeight));
exportFormat = "png"; // "jpg" or "png"
jpegQuality = 85;
path = "C:\\Puffer\\03_TifExport\\02\\";
front = Display.getFront();
Run
Kill
Show Output
Clear
Sourced file: null : Object constructor : at Line: 147 : in file: <unknown file> : new ByteProcessor ( tileWidth * 2 , tileHeight * 2 )
Target exception: java.lang.NegativeArraySizeException
Verwalten
Datei
Start
Freigeben
Ansicht
Bildtools
An Schnellzugriff anheften
Kopieren
Einfügen
Ausschneiden
Pfad kopieren
Verknüpfung einfügen
Zwischenablage
Verschieben nach
Kopieren nach
Löschen
Umbenennen
Organisieren
Neuer Ordner
Neu
Eigenschaften
Öffnen
Bearbeiten
Verlauf
Alles auswählen
Nichts auswählen
Auswahl umkehren
Auswählen
« 02 › 0
"0" durchsuchen
Schnellzugriff
Desktop
Download
Dokument
Bilder
Creative Cloud
OneDrive
Dieser PC
3D-Objekte
Bilder
Desktop
Dokumente
Downloads
Musik
Videos
Windows (C:)
Volume (D:)
Volume (E:)
Netzwerk
0_0_0
0_1_0
0_2_0
0_3_0
0_4_0
0_5_0
0_6_0
1_0_0
1_1_0
1_2_0
1_3_0
1_4_0
1_5_0
1_6_0
2_0_0
2_1_0
2_2_0
2_3_0
2_4_0
2_5_0
2_6_0
3_0_0
3_1_0
3_2_0
3_3_0
3_4_0
3_5_0
3_6_0
4_0_0
4_1_0
4_2_0
4_3_0
4_4_0
4_5_0
4_6_0
35 Elemente

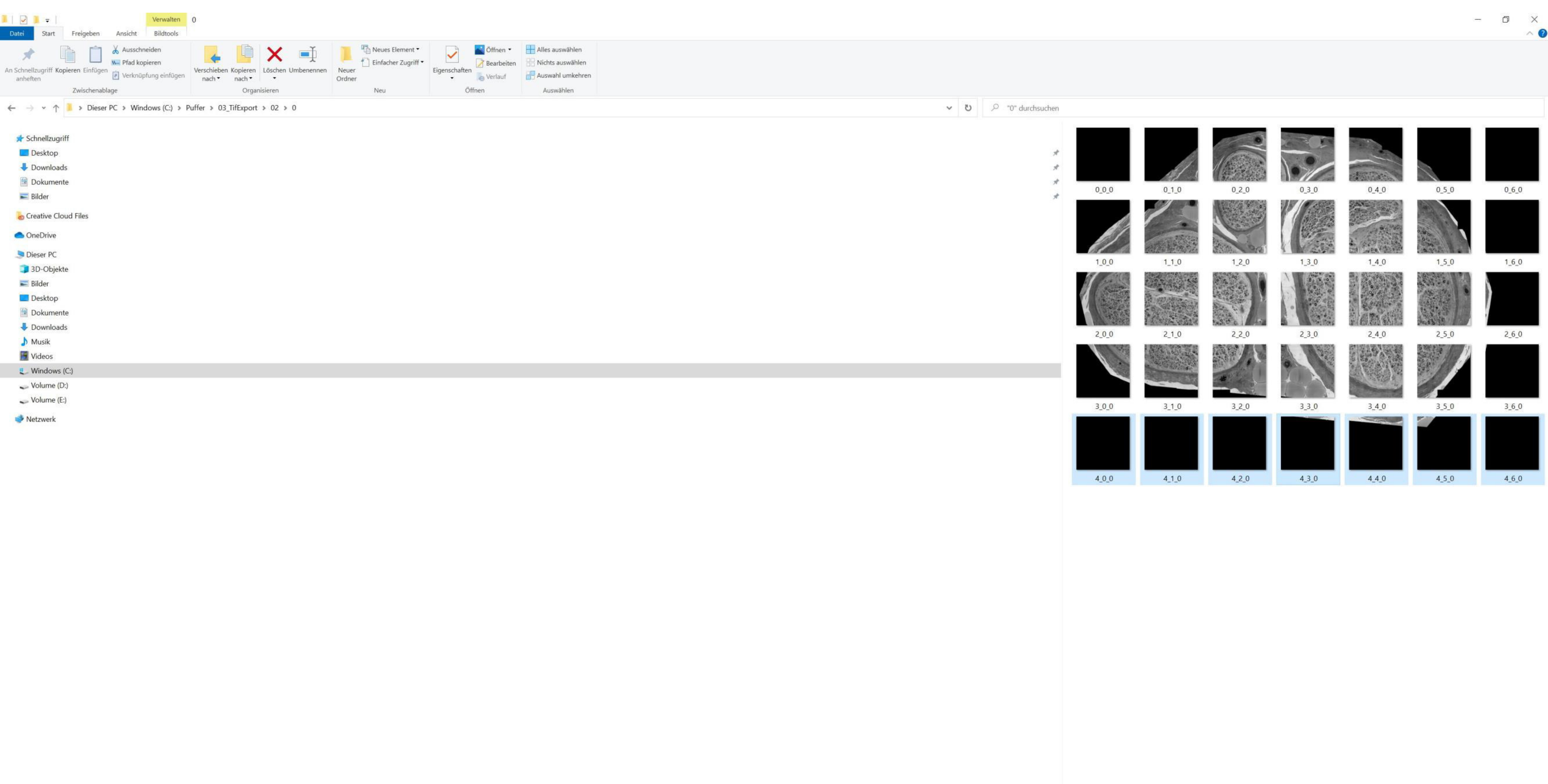


Tiles with low amounts of structural details might be deleted

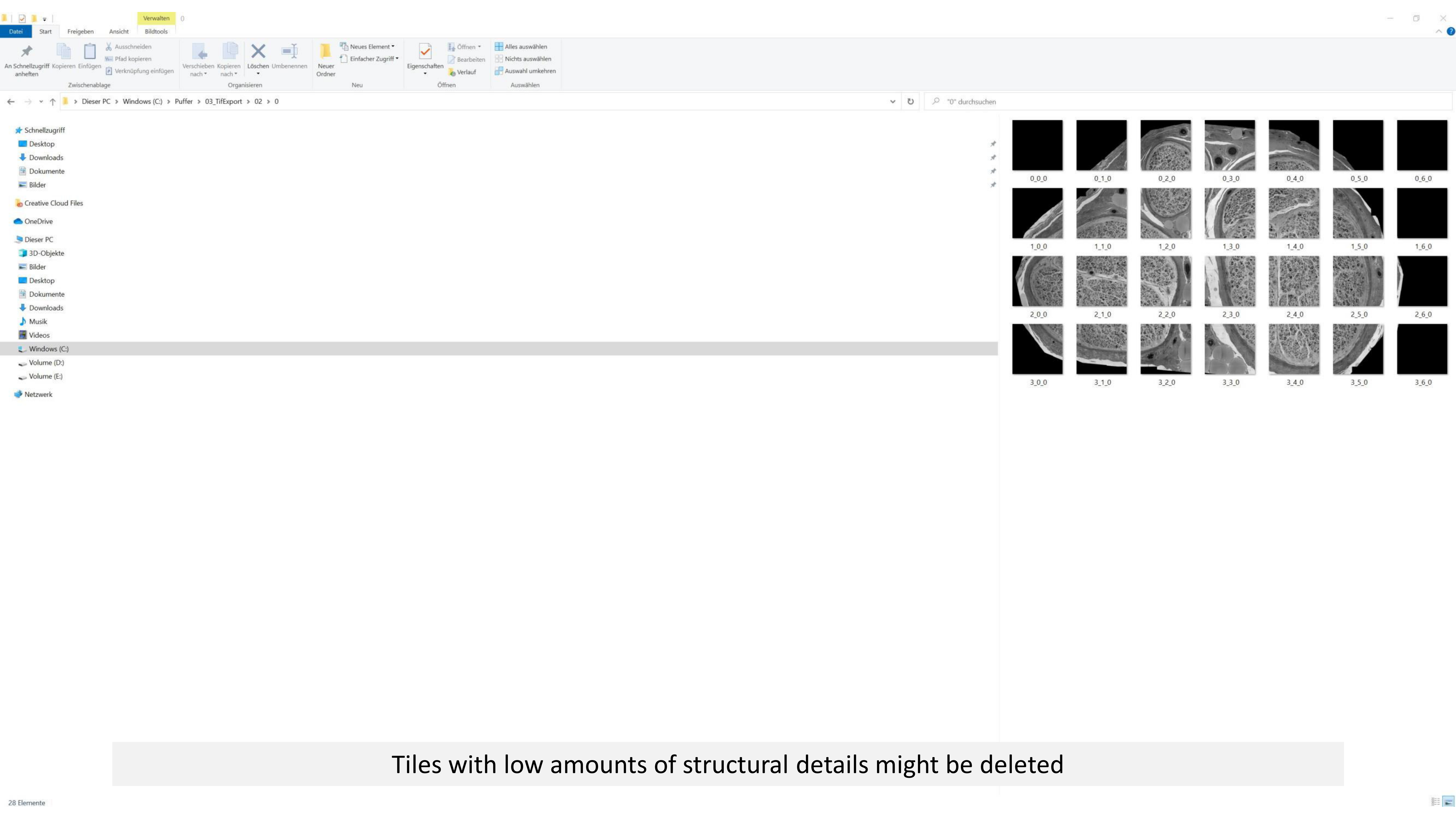


Tiles with low amounts of structural details might be deleted

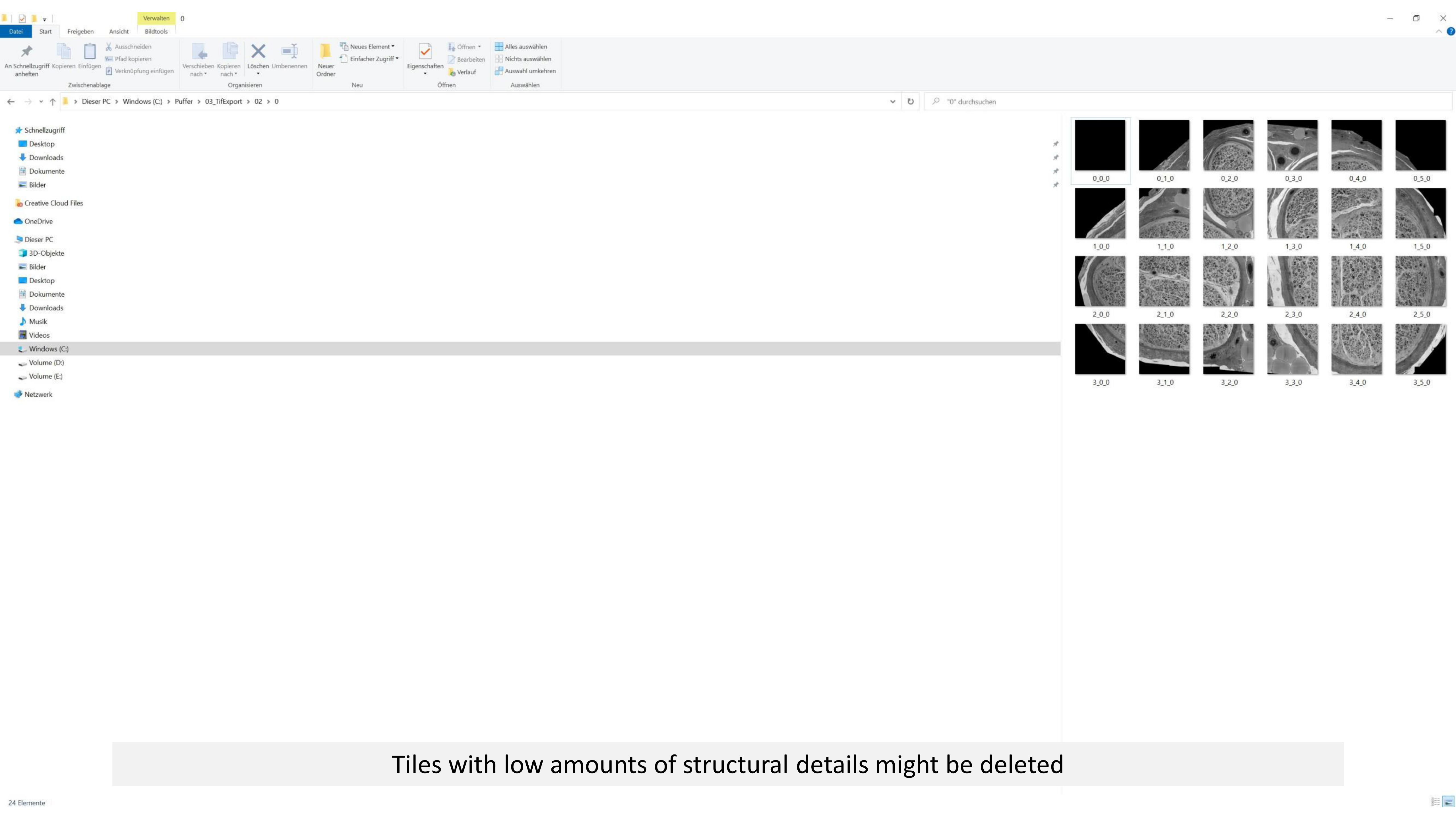


Tiles with low amounts of structural details might be deleted

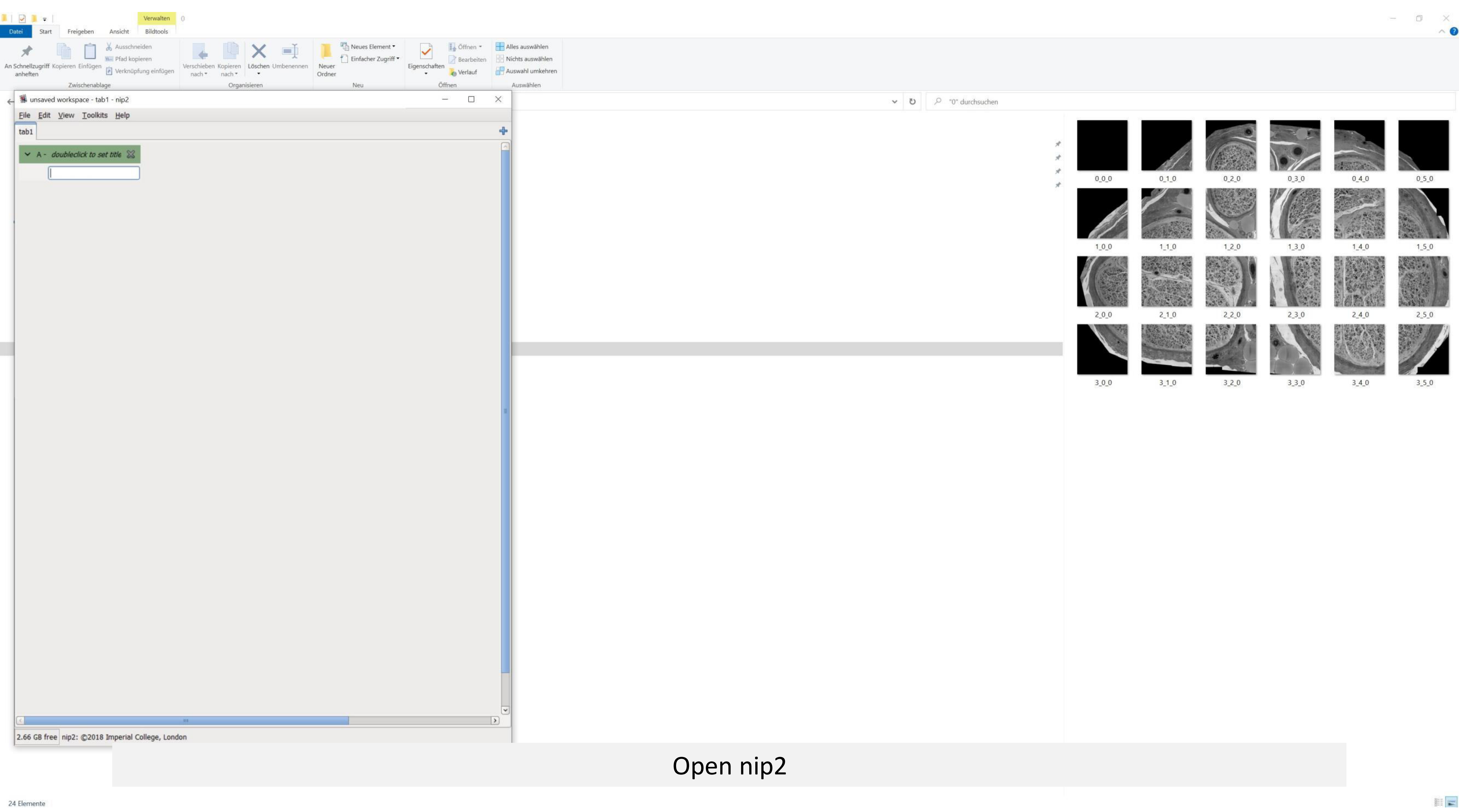
unsaved workspace - tab1 - nip2
File Edit View Toolkits Help
tab1
A - doubleclick to set title
2.66 GB free nip2: ©2018 Imperial College, London
"0" durchsuchen
0_0_0
0_1_0
0_2_0
0_3_0
0_4_0
0_5_0
1_0_0
1_1_0
1_2_0
1_3_0
1_4_0
1_5_0
2_0_0
2_1_0
2_2_0
2_3_0
2_4_0
2_5_0
3_0_0
3_1_0
3_2_0
3_3_0
3_4_0
3_5_0
24 Elemente

Open nip2

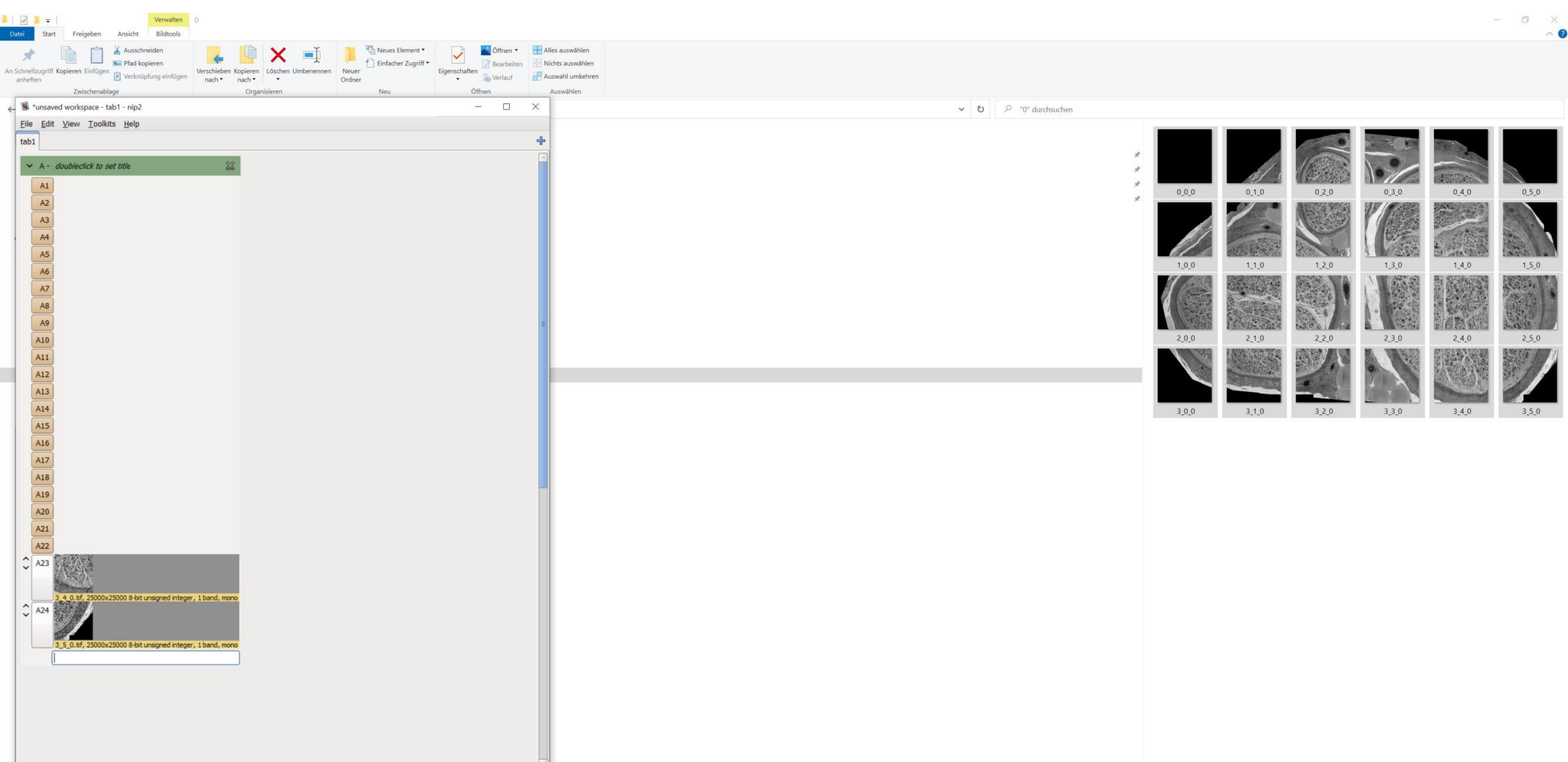


Select all tif tiles of one dataset and copy them via drag&drop into nip2
Ensure that the order (A1-A24) is correct; A1=0_0_0, …, A24= 3_5_0)
Image tiles might be renamed using Bulk rename utility

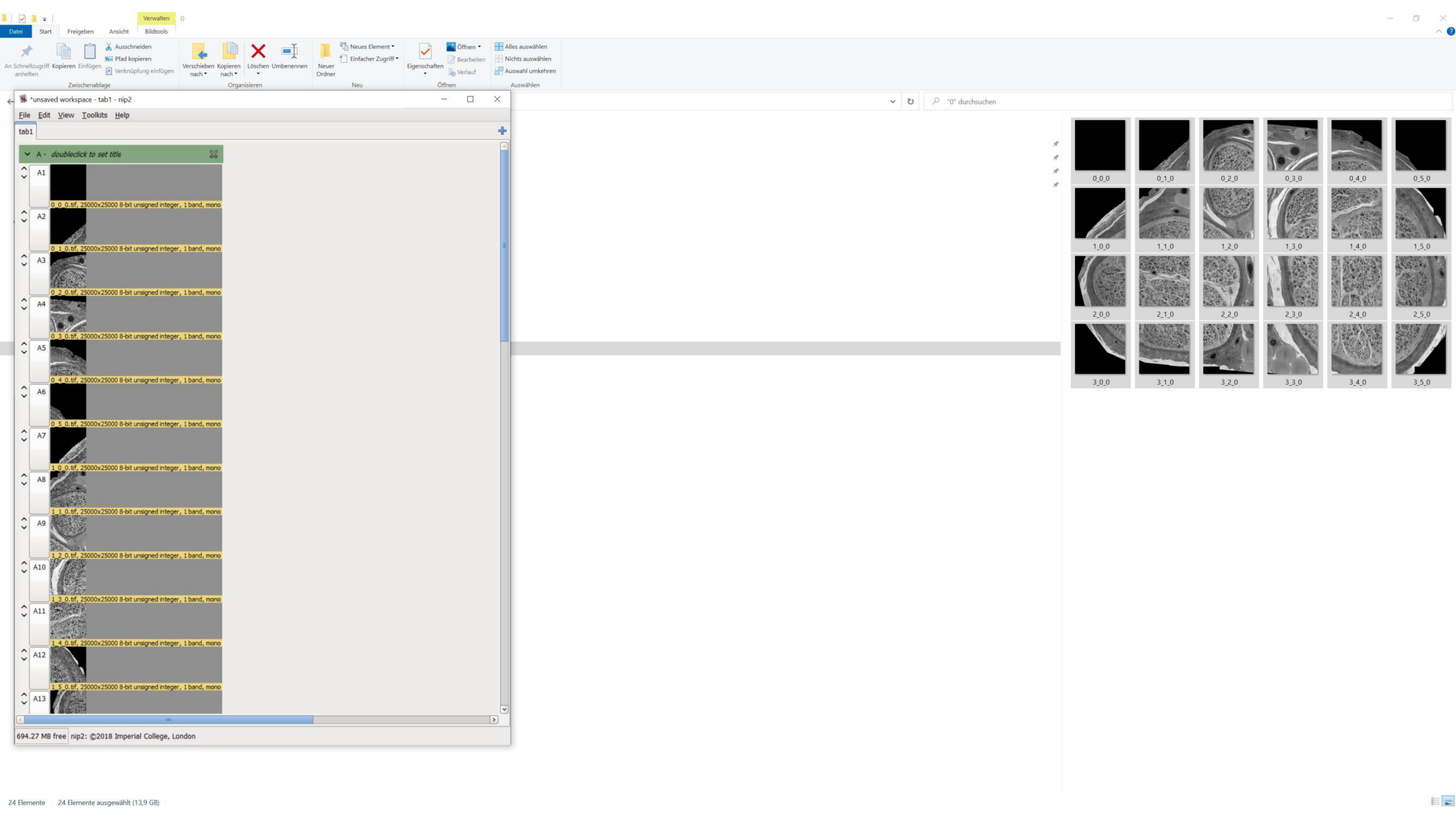
Verwalten
Datei
Start
Freigeben
Ansicht
Bildtools
An Schnellzugriff anheften
Kopieren
Einfügen
Ausschneiden
Pfad kopieren
Verknüpfung einfügen
Zwischenablage
Verschieben nach
Kopieren nach
Löschen
Umbenennen
Organisieren
Neuer Ordner
Neues Element
Einfacher Zugriff
Neu
Eigenschaften
Öffnen
Bearbeiten
Verlauf
Öffnen
Alles auswählen
Nichts auswählen
Auswahl umkehren
Auswählen
*unsaved workspace - tab1 - nip2
File
Edit
View
Toolkits
Help
tab1
A - doubleclick to set title
A1
0_0_0.tif, 25000x25000 8-bit unsigned integer, 1 band, mono
A2
0_1_0.tif, 25000x25000 8-bit unsigned integer, 1 band, mono
A3
0_2_0.tif, 25000x25000 8-bit unsigned integer, 1 band, mono
A4
0_3_0.tif, 25000x25000 8-bit unsigned integer, 1 band, mono
A5
0_4_0.tif, 25000x25000 8-bit unsigned integer, 1 band, mono
A6
0_5_0.tif, 25000x25000 8-bit unsigned integer, 1 band, mono
A7
1_0_0.tif, 25000x25000 8-bit unsigned integer, 1 band, mono
A8
1_1_0.tif, 25000x25000 8-bit unsigned integer, 1 band, mono
A9
1_2_0.tif, 25000x25000 8-bit unsigned integer, 1 band, mono
A10
1_3_0.tif, 25000x25000 8-bit unsigned integer, 1 band, mono
A11
1_4_0.tif, 25000x25000 8-bit unsigned integer, 1 band, mono
A12
1_5_0.tif, 25000x25000 8-bit unsigned integer, 1 band, mono
A13
694.27 MB free
nip2: ©2018 Imperial College, London
"0" durchsuchen
0_0_0
0_1_0
0_2_0
0_3_0
0_4_0
0_5_0
1_0_0
1_1_0
1_2_0
1_3_0
1_4_0
1_5_0
2_0_0
2_1_0
2_2_0
2_3_0
2_4_0
2_5_0
3_0_0
3_1_0
3_2_0
3_3_0
3_4_0
3_5_0
24 Elemente
24 Elemente ausgewählt (13,9 GB)

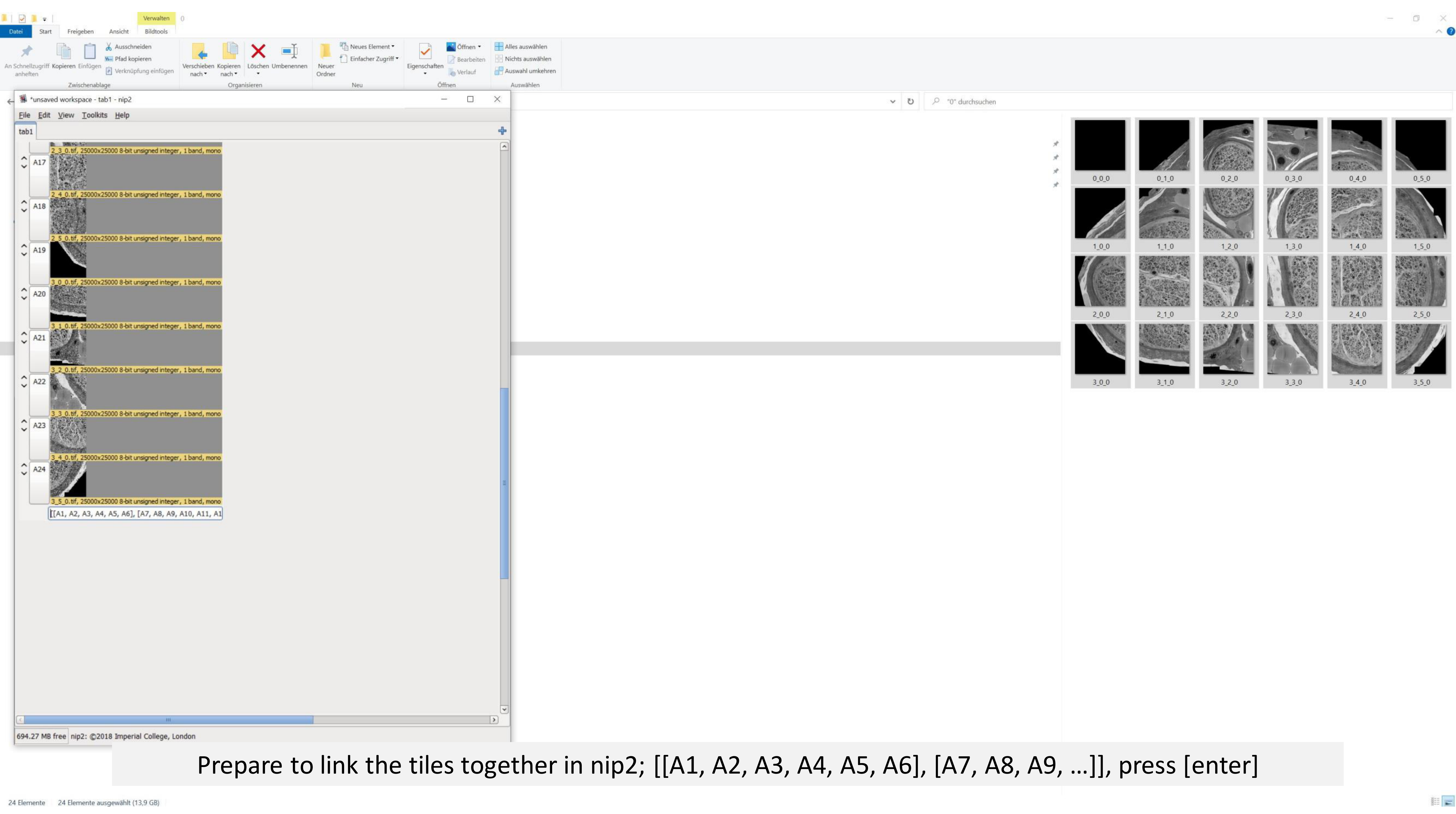


Prepare to link the tiles together in nip2; [[A1, A2, A3, A4, A5, A6], [A7, A8, A9, ...]], press [enter]

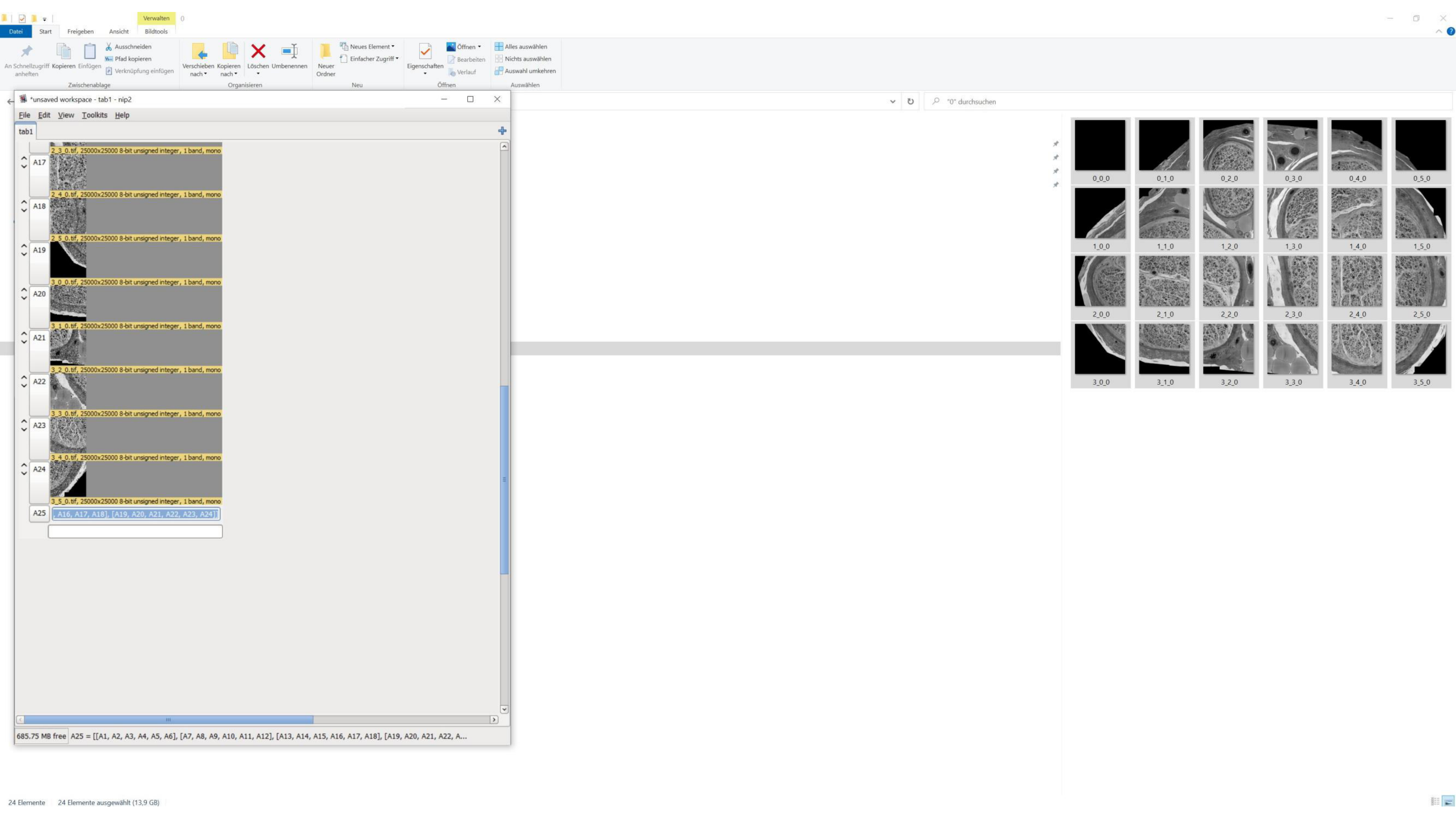
Verwalten
Datei
Start
Freigeben
Ansicht
Bildtools
An Schnellzugriff anheften
Kopieren
Einfügen
Ausschneiden
Pfad kopieren
Verknüpfung einfügen
Zwischenablage
Verschieben nach
Kopieren nach
Löschen
Umbenennen
Organisieren
Neuer Ordner
Neues Element
Einfacher Zugriff
Neu
Eigenschaften
Öffnen
Bearbeiten
Verlauf
Öffnen
Alles auswählen
Nichts auswählen
Auswahl umkehren
Auswählen
*unsaved workspace - tab1 - nip2
File
Edit
View
Toolkits
Help
tab1
"0" durchsuchen
2_3_0.tif, 25000x25000 8-bit unsigned integer, 1 band, mono
A17
2_4_0.tif, 25000x25000 8-bit unsigned integer, 1 band, mono
A18
2_5_0.tif, 25000x25000 8-bit unsigned integer, 1 band, mono
A19
3_0_0.tif, 25000x25000 8-bit unsigned integer, 1 band, mono
A20
3_1_0.tif, 25000x25000 8-bit unsigned integer, 1 band, mono
A21
3_2_0.tif, 25000x25000 8-bit unsigned integer, 1 band, mono
A22
3_3_0.tif, 25000x25000 8-bit unsigned integer, 1 band, mono
A23
3_4_0.tif, 25000x25000 8-bit unsigned integer, 1 band, mono
A24
3_5_0.tif, 25000x25000 8-bit unsigned integer, 1 band, mono
A25
, A16, A17, A18], [A19, A20, A21, A22, A23, A24]]
685.75 MB free
A25 = [[A1, A2, A3, A4, A5, A6], [A7, A8, A9, A10, A11, A12], [A13, A14, A15, A16, A17, A18], [A19, A20, A21, A22, A...
0_0_0
0_1_0
0_2_0
0_3_0
0_4_0
0_5_0
1_0_0
1_1_0
1_2_0
1_3_0
1_4_0
1_5_0
2_0_0
2_1_0
2_2_0
2_3_0
2_4_0
2_5_0
3_0_0
3_1_0
3_2_0
3_3_0
3_4_0
3_5_0
24 Elemente
24 Elemente ausgewählt (13,9 GB)

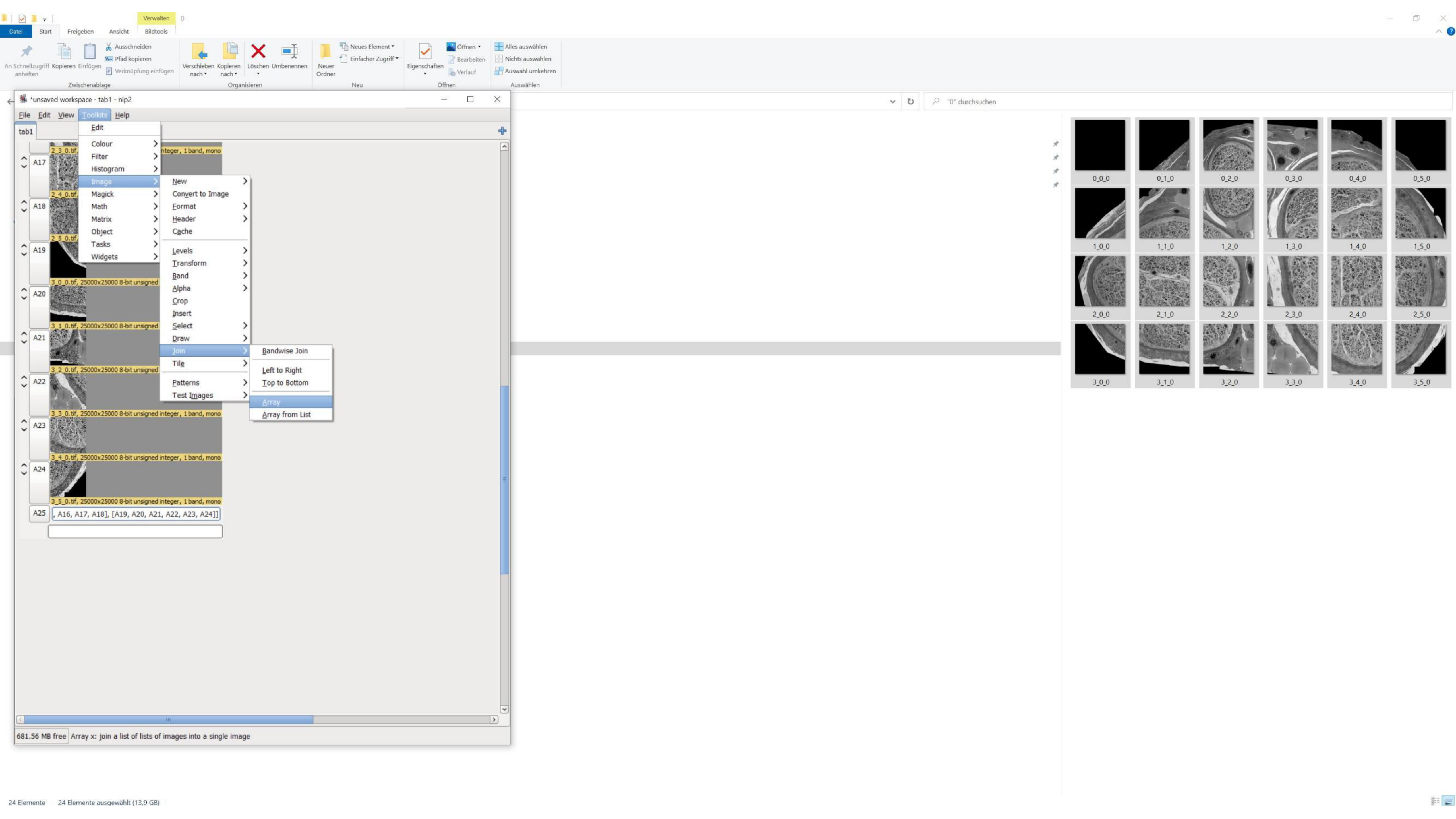
*unsaved workspace - tab1 - nip2
File
Edit
View
Toolkits
Help
tab1
Edit
Colour
Filter
Histogram
Image
Magick
Math
Matrix
Object
Tasks
Widgets
New
Convert to Image
Format
Header
Cache
Levels
Transform
Band
Alpha
Crop
Insert
Select
Draw
Join
Tile
Patterns
Test Images
Bandwise Join
Left to Right
Top to Bottom
Array
Array from List
A17
A18
A19
A20
A21
A22
A23
A24
A25
3_0_0.tif, 25000x25000 8-bit unsigned
3_1_0.tif, 25000x25000 8-bit unsigned
3_2_0.tif, 25000x25000 8-bit unsigned
3_3_0.tif, 25000x25000 8-bit unsigned integer, 1 band, mono
3_4_0.tif, 25000x25000 8-bit unsigned integer, 1 band, mono
3_5_0.tif, 25000x25000 8-bit unsigned integer, 1 band, mono
, A16, A17, A18], [A19, A20, A21, A22, A23, A24]]
681.56 MB free
Array x: join a list of lists of images into a single image
Datei
Start
Freigeben
Ansicht
Verwalten
Bildtools
Ausschneiden
Pfad kopieren
Verknüpfung einfügen
Kopieren
Einfügen
Zwischenablage
Verschieben nach
Kopieren nach
Löschen
Umbenennen
Organisieren
Neuer Ordner
Neues Element
Einfacher Zugriff
Neu
Eigenschaften
Öffnen
Bearbeiten
Verlauf
Alles auswählen
Nichts auswählen
Auswahl umkehren
Auswählen
"0" durchsuchen
0_0_0
0_1_0
0_2_0
0_3_0
0_4_0
0_5_0
1_0_0
1_1_0
1_2_0
1_3_0
1_4_0
1_5_0
2_0_0
2_1_0
2_2_0
2_3_0
2_4_0
2_5_0
3_0_0
3_1_0
3_2_0
3_3_0
3_4_0
3_5_0
24 Elemente
24 Elemente ausgewählt (13,9 GB)

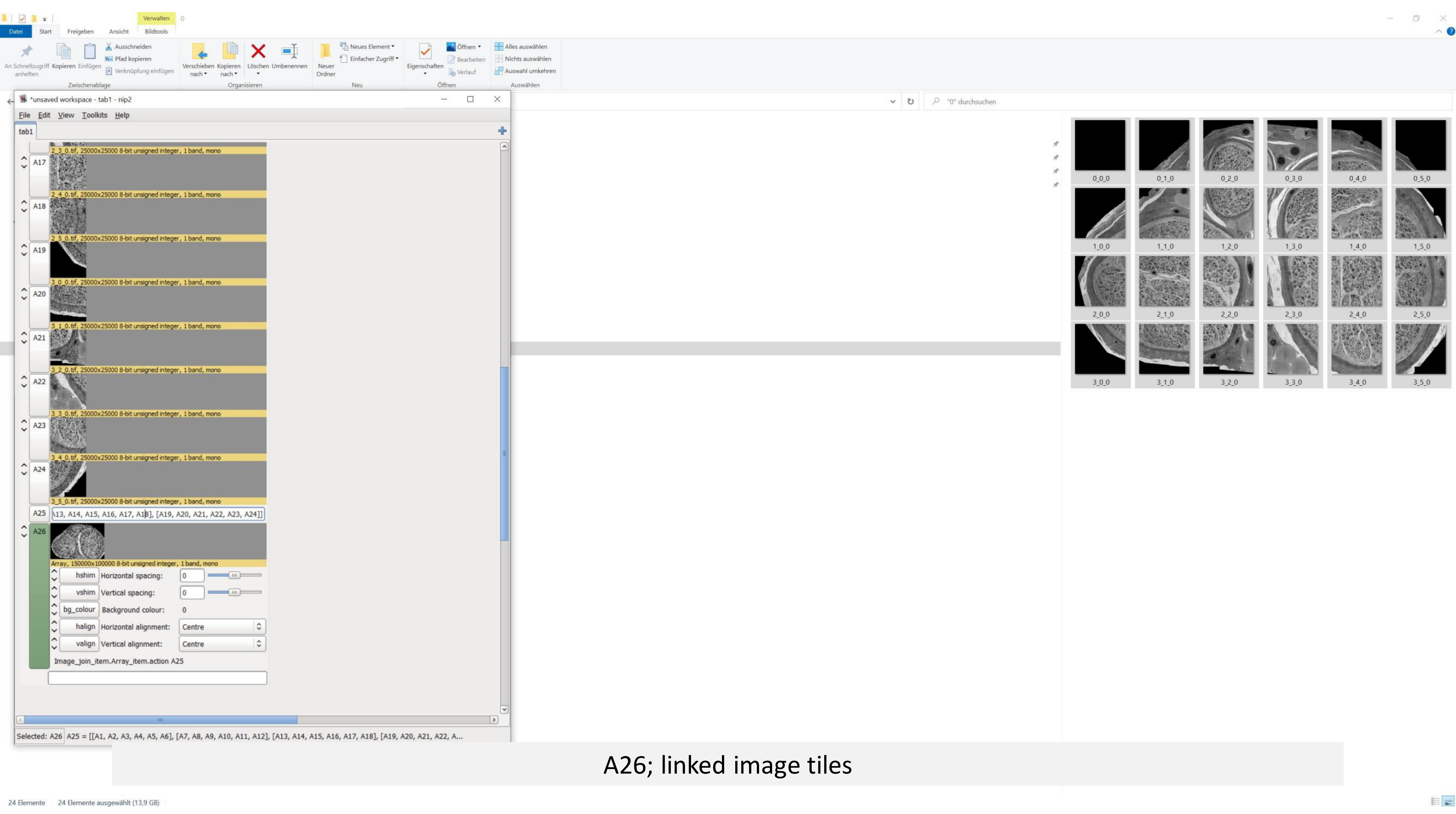


A26; linked image tiles

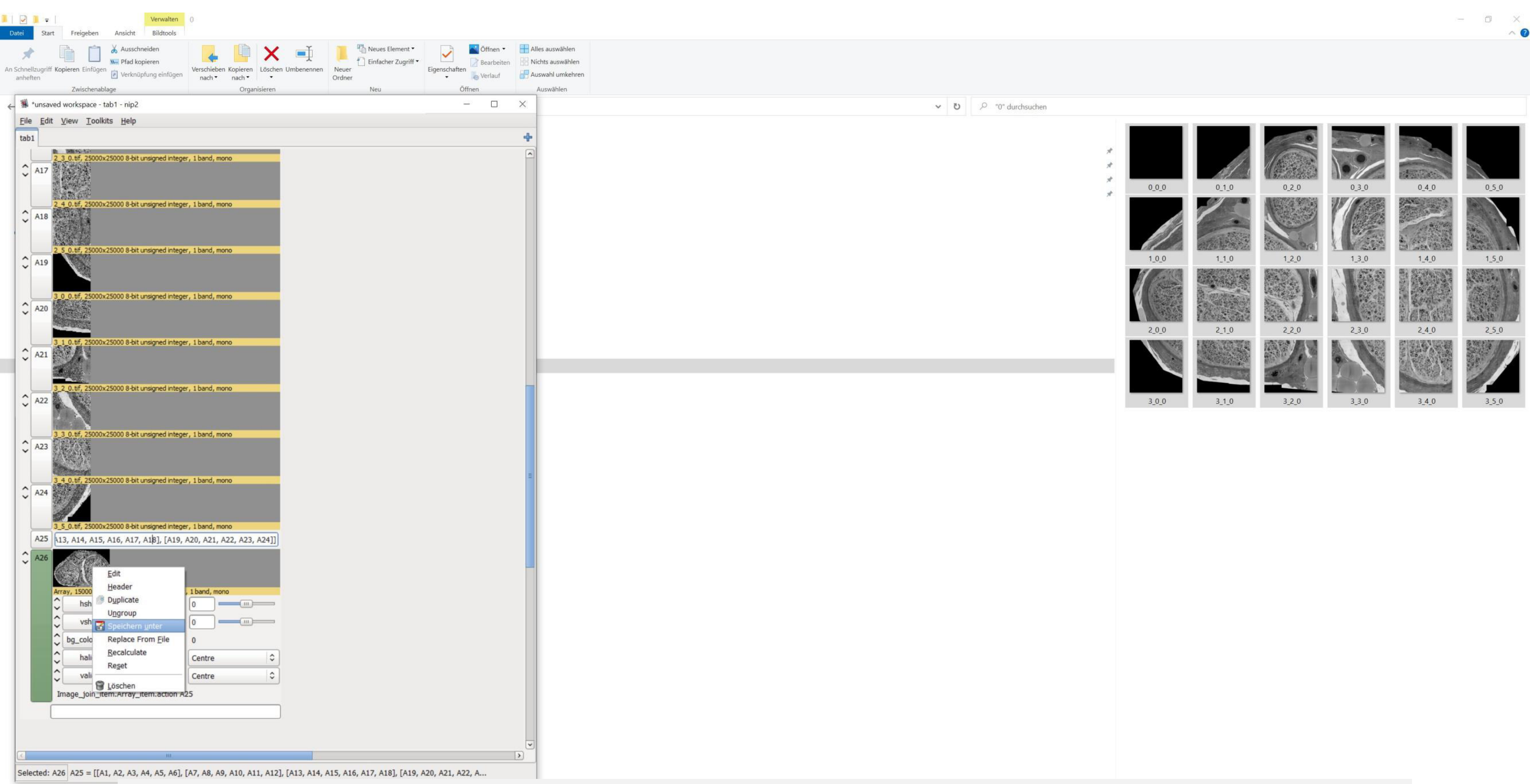


Right click on the preview-> save as

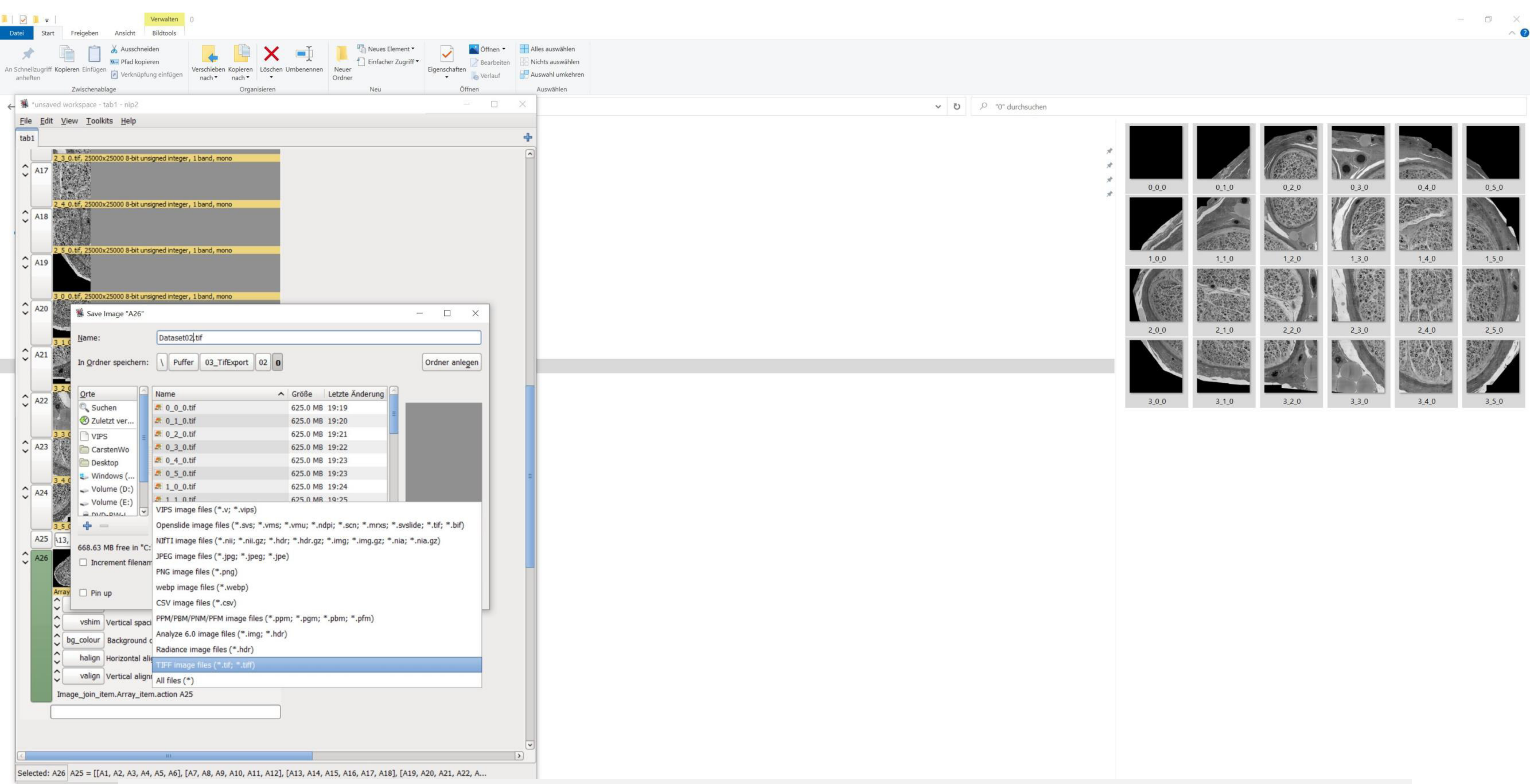
*unsaved workspace - tab1 - nip2
File Edit View Toolkits Help
2_3_0.tif, 25000x25000 8-bit unsigned integer, 1 band, mono
2_4_0.tif, 25000x25000 8-bit unsigned integer, 1 band, mono
2_5_0.tif, 25000x25000 8-bit unsigned integer, 1 band, mono
3_0_0.tif, 25000x25000 8-bit unsigned integer, 1 band, mono
Save Image "A26"
Name: Dataset02.tif
In Ordner speichern: \ Puffer 03_TifExport 02 0
Ordner anlegen
Orte
Suchen
Zuletzt ver...
VIPS
CarstenWo
Desktop
Windows (...
Volume (D:)
Volume (E:)
Name Größe Letzte Änderung
0_0_0.tif 625.0 MB 19:19
0_1_0.tif 625.0 MB 19:20
0_2_0.tif 625.0 MB 19:21
0_3_0.tif 625.0 MB 19:22
0_4_0.tif 625.0 MB 19:23
0_5_0.tif 625.0 MB 19:23
1_0_0.tif 625.0 MB 19:24
VIPS image files (*.v; *.vips)
Openslide image files (*.svs; *.vms; *.vmu; *.ndpi; *.scn; *.mrxs; *.svslide; *.tif; *.bif)
NIfTI image files (*.nii; *.nii.gz; *.hdr; *.hdr.gz; *.img; *.img.gz; *.nia; *.nia.gz)
JPEG image files (*.jpg; *.jpeg; *.jpe)
PNG image files (*.png)
webp image files (*.webp)
CSV image files (*.csv)
PPM/PBM/PNM/PFM image files (*.ppm; *.pgm; *.pbm; *.pfm)
Analyze 6.0 image files (*.img; *.hdr)
Radiance image files (*.hdr)
TIFF image files (*.tif; *.tiff)
All files (*)
668.63 MB free in "C:
Increment filenam
Pin up
Image_join_item.Array_item.action A25
Selected: A26 A25 = [[A1, A2, A3, A4, A5, A6], [A7, A8, A9, A10, A11, A12], [A13, A14, A15, A16, A17, A18], [A19, A20, A21, A22, A...
"0" durchsuchen
0_0_0 0_1_0 0_2_0 0_3_0 0_4_0 0_5_0
1_0_0 1_1_0 1_2_0 1_3_0 1_4_0 1_5_0
2_0_0 2_1_0 2_2_0 2_3_0 2_4_0 2_5_0
3_0_0 3_1_0 3_2_0 3_3_0 3_4_0 3_5_0

Save as TIFF

24 Elemente 24 Elemente ausgewählt (13,9 GB)

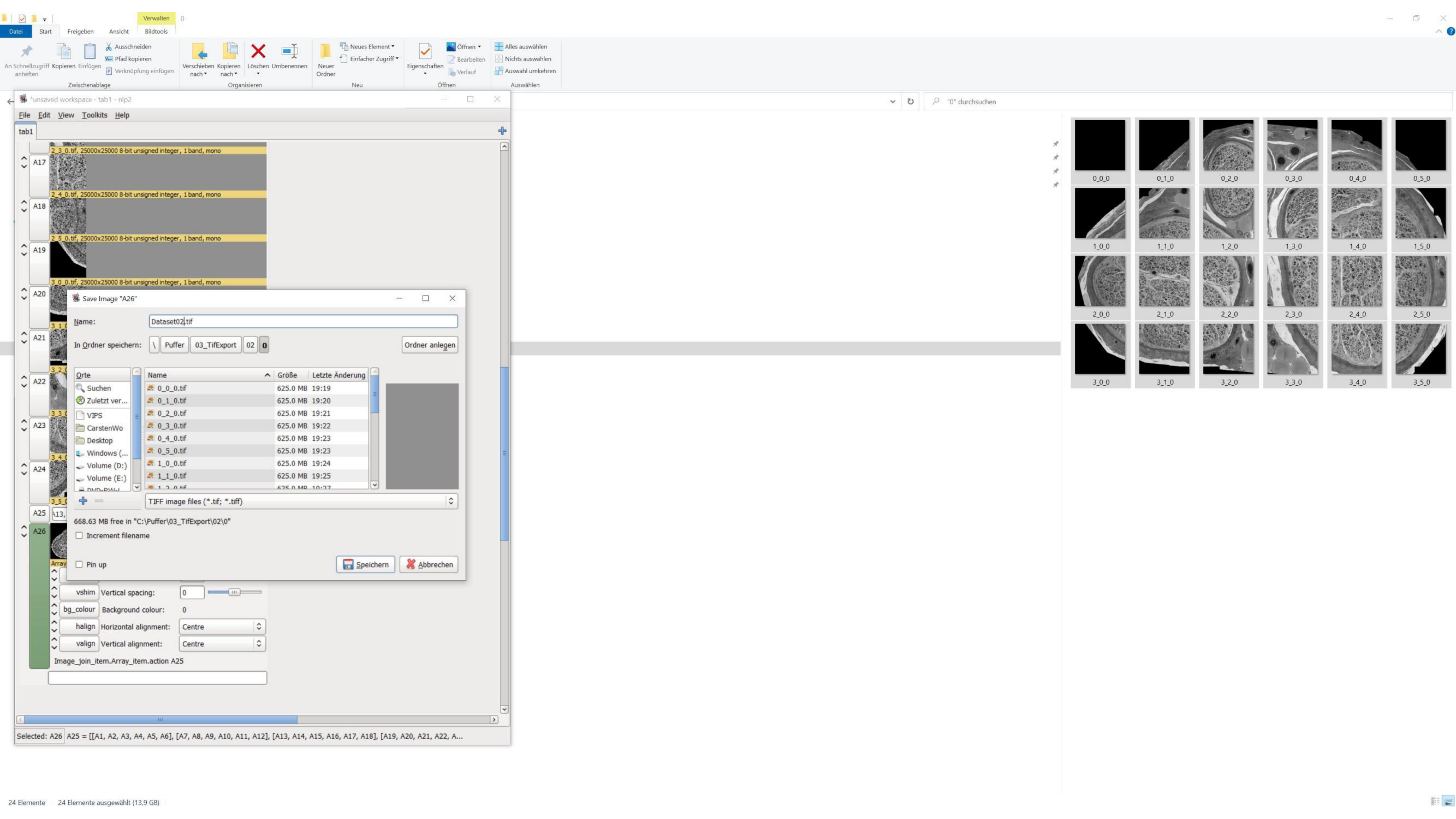

Verwalten
Datei
Start
Freigeben
Ansicht
Bildtools
Zwischenablage
Organisieren
Neu
Öffnen
Auswählen
*unsaved workspace - tab1 - nip2
File Edit View Toolkits Help
tab1
2_3_0.tif, 25000x25000 8-bit unsigned integer, 1 band, mono
2_4_0.tif, 25000x25000 8-bit unsigned integer, 1 band, mono
2_5_0.tif, 25000x25000 8-bit unsigned integer, 1 band, mono
3_0_0.tif, 25000x25000 8-bit unsigned integer, 1 band, mono
Save Image "A26"
Name: Dataset02.tif
In Ordner speichern: \ Puffer 03_TifExport 02 0
Ordner anlegen
Orte
Suchen
Zuletzt ver...
VIPS
CarstenWo
Desktop
Windows (...
Volume (D:)
Volume (E:)
Name Größe Letzte Änderung
0_0_0.tif 625.0 MB 19:19
0_1_0.tif 625.0 MB 19:20
0_2_0.tif 625.0 MB 19:21
0_3_0.tif 625.0 MB 19:22
0_4_0.tif 625.0 MB 19:23
0_5_0.tif 625.0 MB 19:23
1_0_0.tif 625.0 MB 19:24
1_1_0.tif 625.0 MB 19:25
TIFF image files (*.tif; *.tiff)
668.63 MB free in "C:\Puffer\03_TifExport\02\0"
Increment filename
Pin up
Speichern
Abbrechen
Vertical spacing:
Background colour:
Horizontal alignment: Centre
Vertical alignment: Centre
Image_join_item.Array_item.action A25
Selected: A26 A25 = [[A1, A2, A3, A4, A5, A6], [A7, A8, A9, A10, A11, A12], [A13, A14, A15, A16, A17, A18], [A19, A20, A21, A22, A...
"0" durchsuchen
0_0_0
0_1_0
0_2_0
0_3_0
0_4_0
0_5_0
1_0_0
1_1_0
1_2_0
1_3_0
1_4_0
1_5_0
2_0_0
2_1_0
2_2_0
2_3_0
2_4_0
2_5_0
3_0_0
3_1_0
3_2_0
3_3_0
3_4_0
3_5_0
24 Elemente
24 Elemente ausgewählt (13,9 GB)

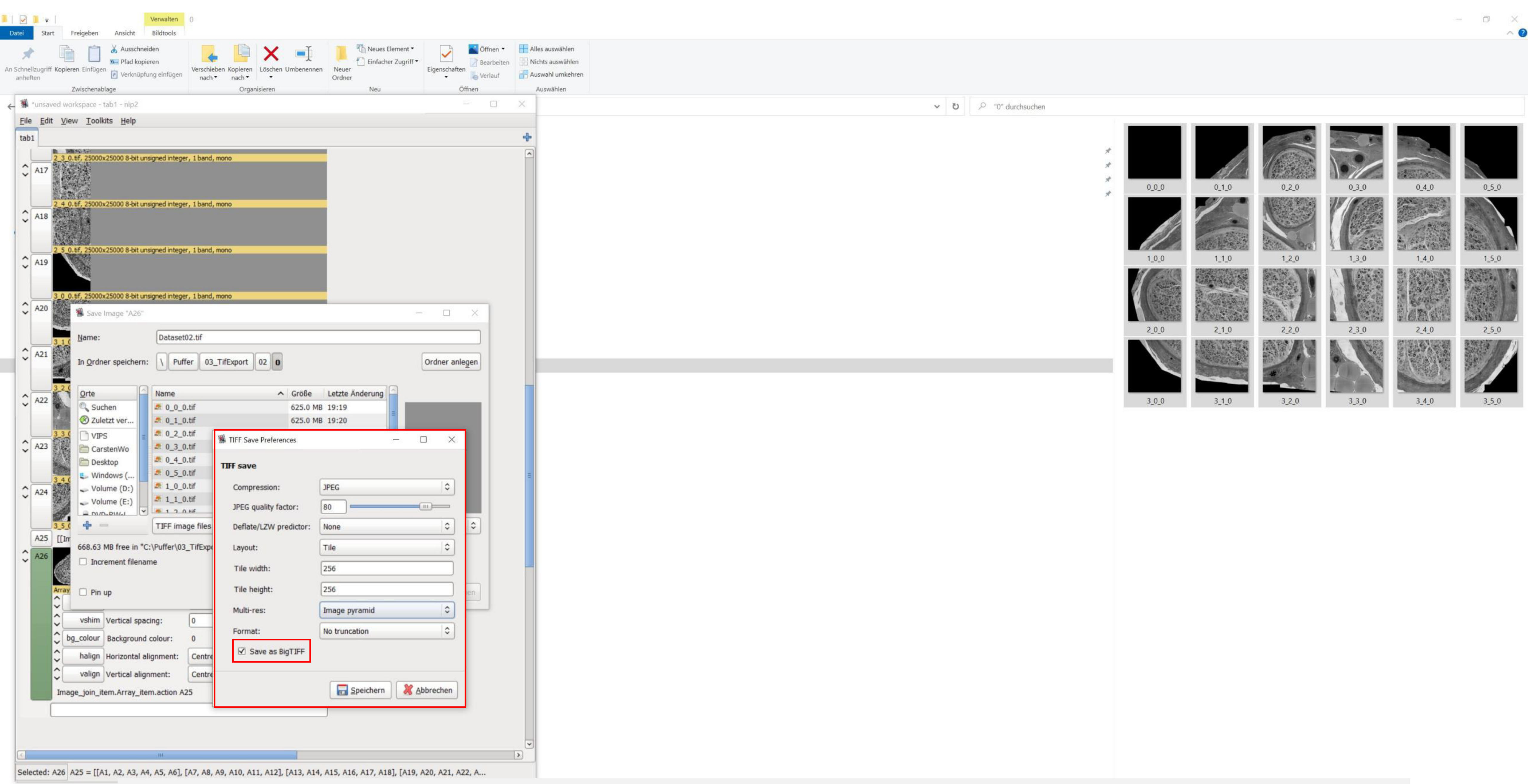


See „TIFF Save Preferences“ for detailed export parameters

Check „Save as BigTIFF“

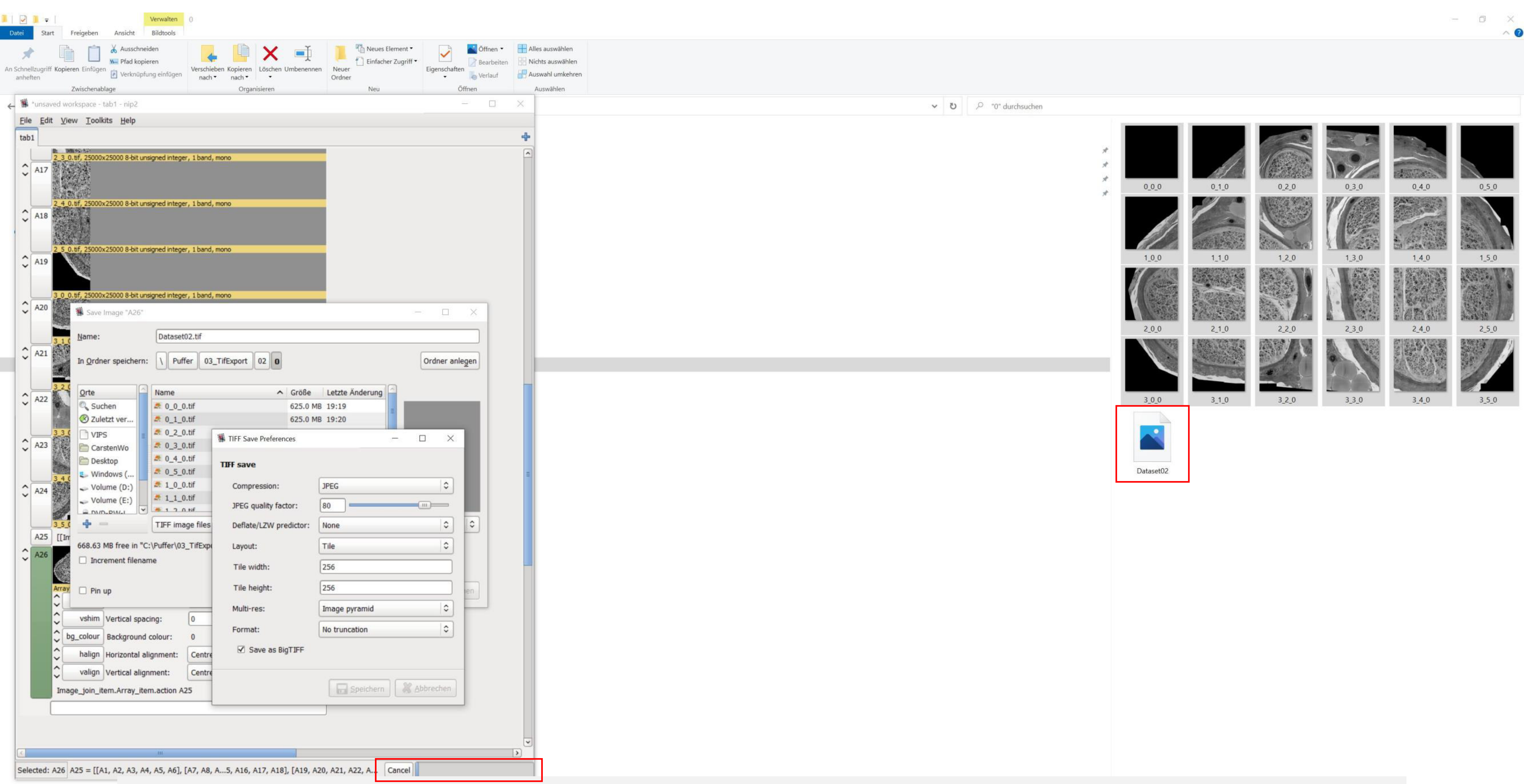


Press save; a new file is prepared (export might take some minutes)

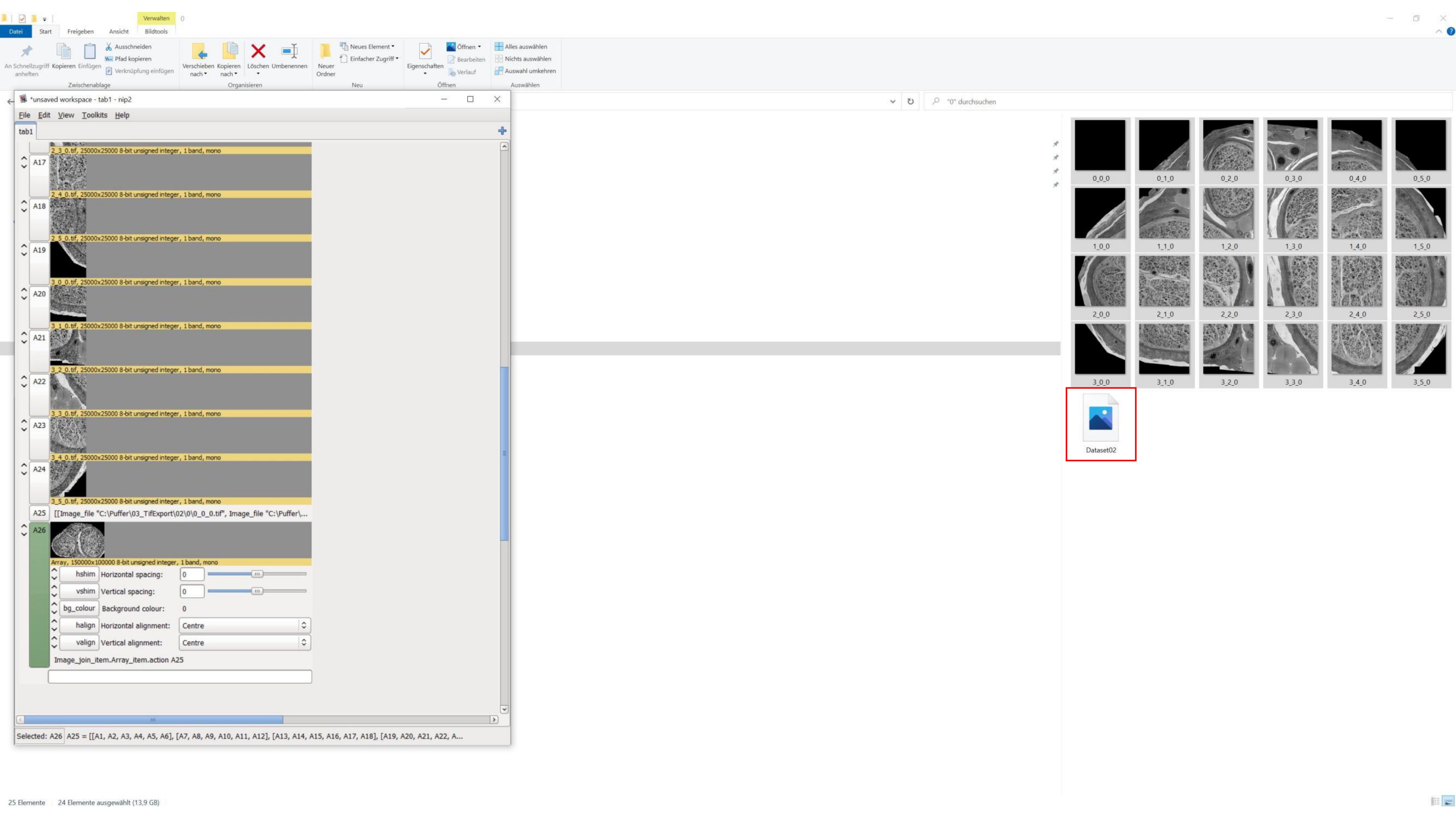
*unsaved workspace - tab1 - nip2
File Edit View Toolkits Help
tab1
2_3_0.tif, 25000x25000 8-bit unsigned integer, 1 band, mono
A17
2_4_0.tif, 25000x25000 8-bit unsigned integer, 1 band, mono
A18
2_5_0.tif, 25000x25000 8-bit unsigned integer, 1 band, mono
A19
3_0_0.tif, 25000x25000 8-bit unsigned integer, 1 band, mono
A20
3_1_0.tif, 25000x25000 8-bit unsigned integer, 1 band, mono
A21
3_2_0.tif, 25000x25000 8-bit unsigned integer, 1 band, mono
A22
3_3_0.tif, 25000x25000 8-bit unsigned integer, 1 band, mono
A23
3_4_0.tif, 25000x25000 8-bit unsigned integer, 1 band, mono
A24
3_5_0.tif, 25000x25000 8-bit unsigned integer, 1 band, mono
A25 [[Image_file "C:\Puffer\03_TifExport\02\0\0_0_0.tif", Image_file "C:\Puffer\...
A26
Array, 150000x100000 8-bit unsigned integer, 1 band, mono
hshim Horizontal spacing: 0
vshim Vertical spacing: 0
bg_colour Background colour: 0
halign Horizontal alignment: Centre
valign Vertical alignment: Centre
Image_join_item.Array_item.action A25
Selected: A26 A25 = [[A1, A2, A3, A4, A5, A6], [A7, A8, A9, A10, A11, A12], [A13, A14, A15, A16, A17, A18], [A19, A20, A21, A22, A...
Datei Start Freigeben Ansicht Verwalten Bildtools
An Schnellzugriff anheften Kopieren Einfügen Ausschneiden Pfad kopieren Verknüpfung einfügen
Zwischenablage
Verschieben nach Kopieren nach Löschen Umbenennen
Organisieren
Neuer Ordner Neues Element Einfacher Zugriff
Neu
Eigenschaften Öffnen Bearbeiten Verlauf
Öffnen
Alles auswählen Nichts auswählen Auswahl umkehren
Auswählen
"0" durchsuchen
0_0_0 0_1_0 0_2_0 0_3_0 0_4_0 0_5_0
1_0_0 1_1_0 1_2_0 1_3_0 1_4_0 1_5_0
2_0_0 2_1_0 2_2_0 2_3_0 2_4_0 2_5_0
3_0_0 3_1_0 3_2_0 3_3_0 3_4_0 3_5_0
Dataset02
25 Elemente 24 Elemente ausgewählt (13,9 GB)

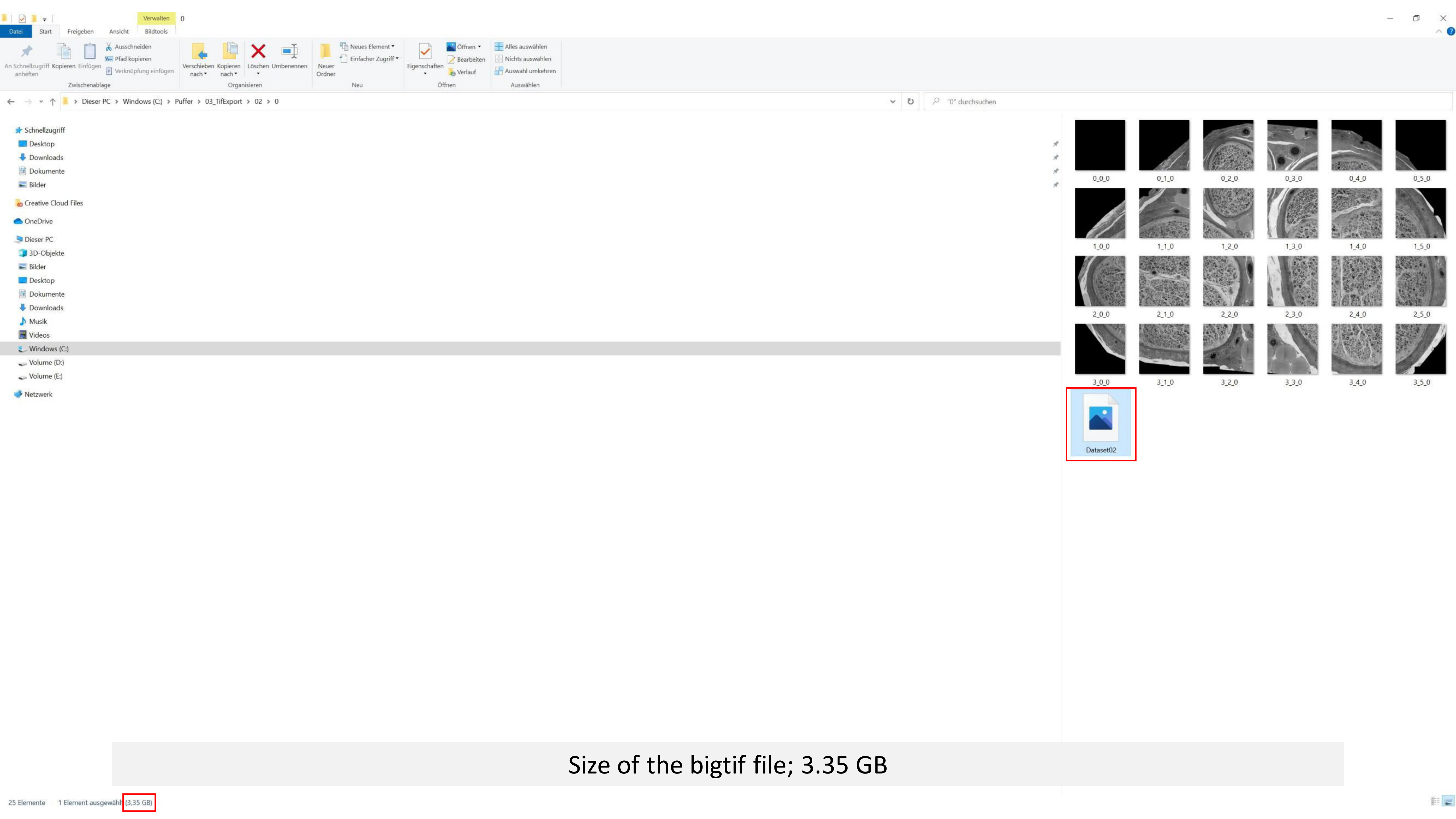


Size of the bigtif file; 3.35 GB

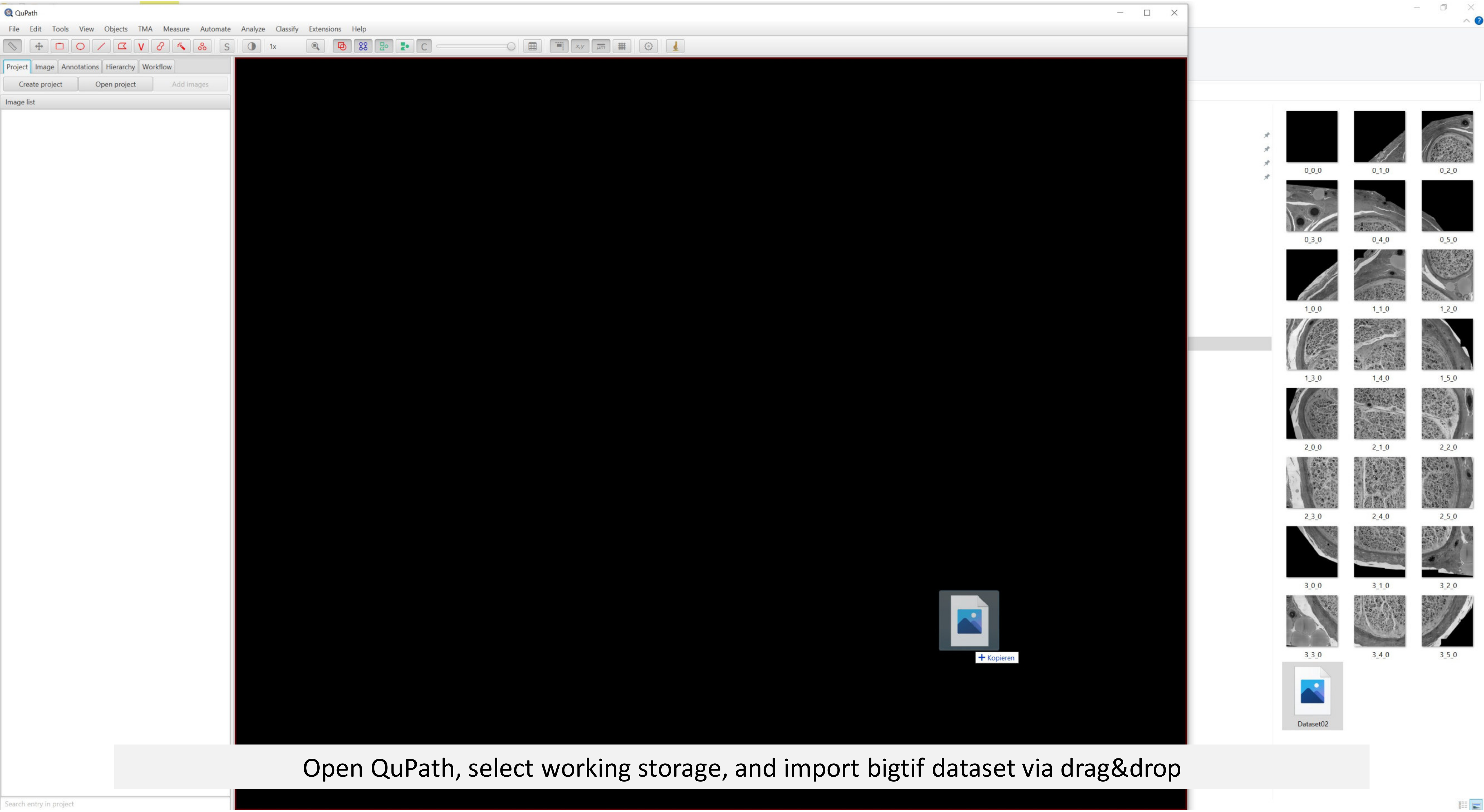


Open QuPath, select working storage, and import bigtif dataset via drag&drop

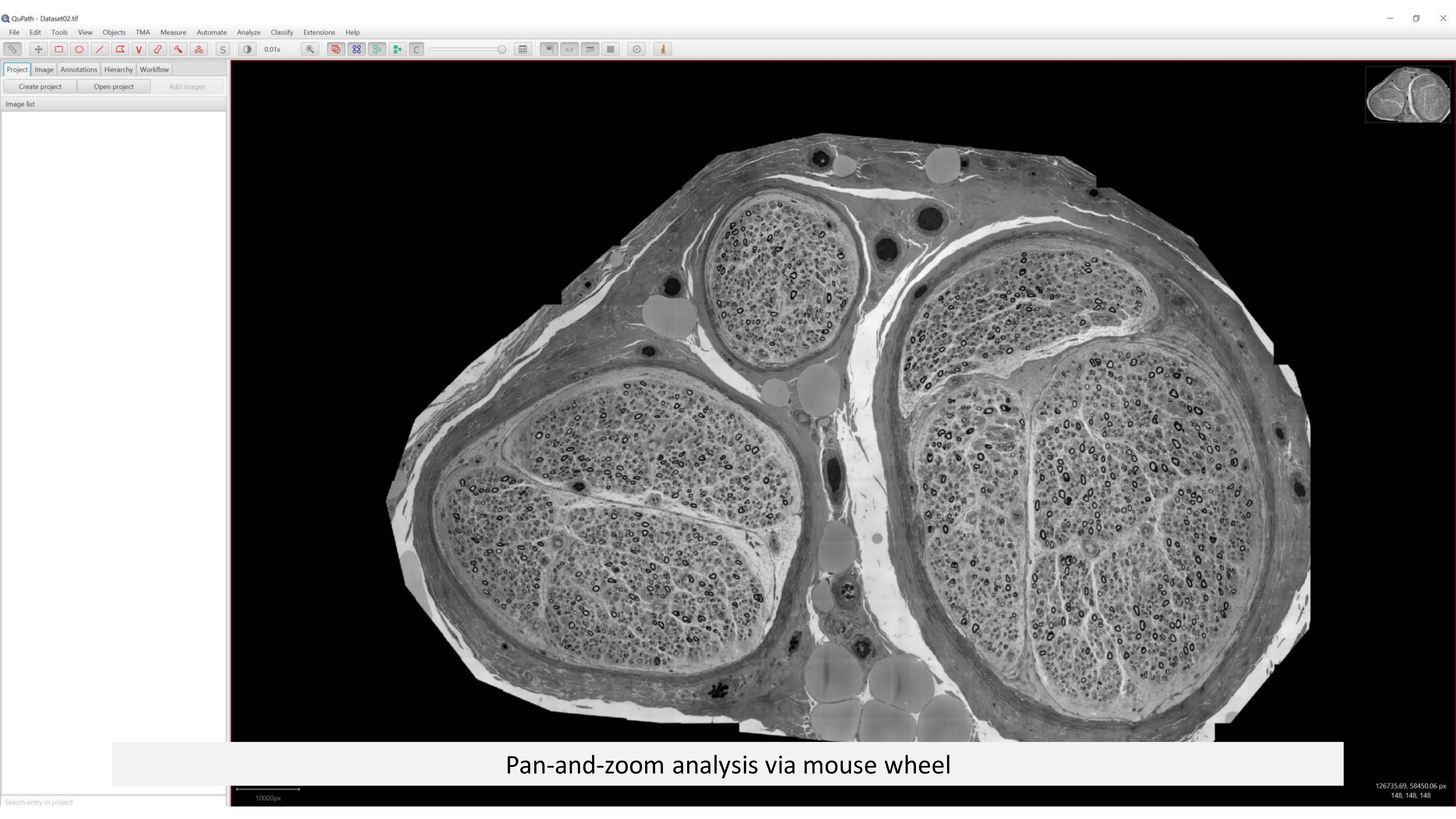
QuPath - Dataset02.tif
File Edit Tools View Objects TMA Measure Automate Analyze Classify Extensions Help
0.01x
Project Image Annotations Hierarchy Workflow
Create project
Open project
Add images
Image list
Search entry in project
10000px
126735.69, 58450.06 px
148, 148, 148
Pan-and-zoom analysis via mouse wheel

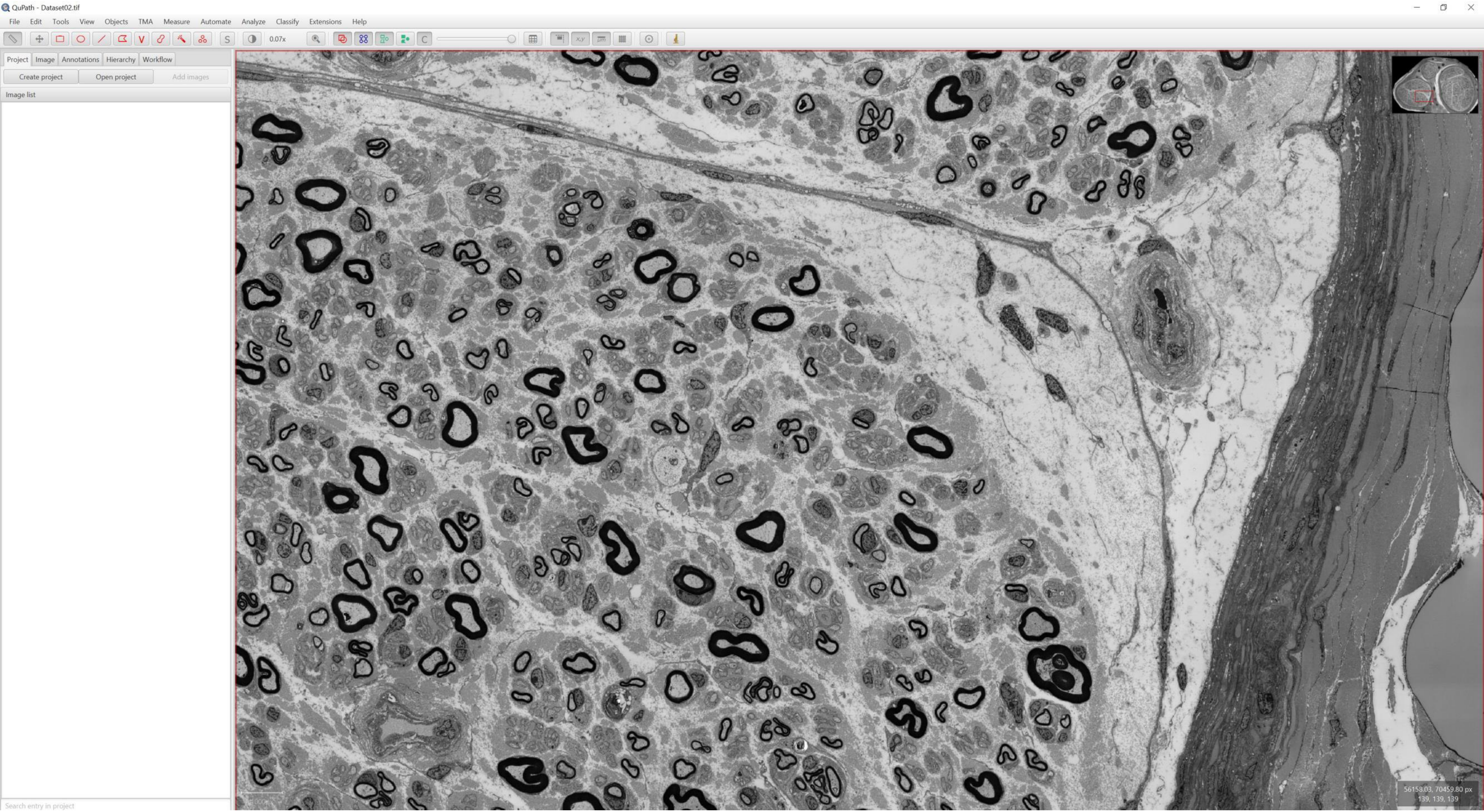
QuPath - Dataset02.tif
File
Edit
Tools
View
Objects
TMA
Measure
Automate
Analyze
Classify
Extensions
Help
0.07x
Project
Image
Annotations
Hierarchy
Workflow
Create project
Open project
Add images
Image list
Search entry in project
56158.03, 70459.80 px
139, 139, 139

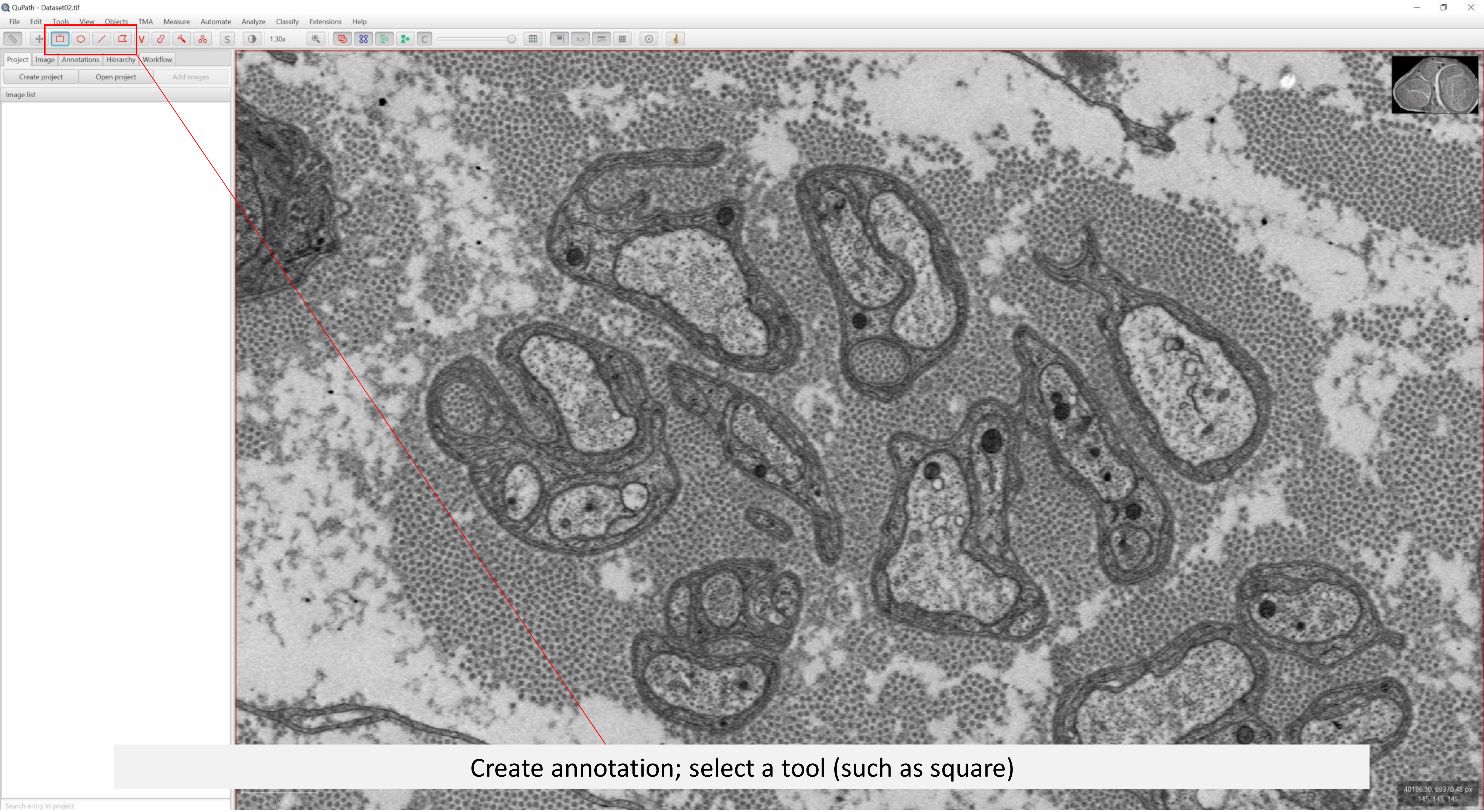


Create annotation; select a tool (such as square)

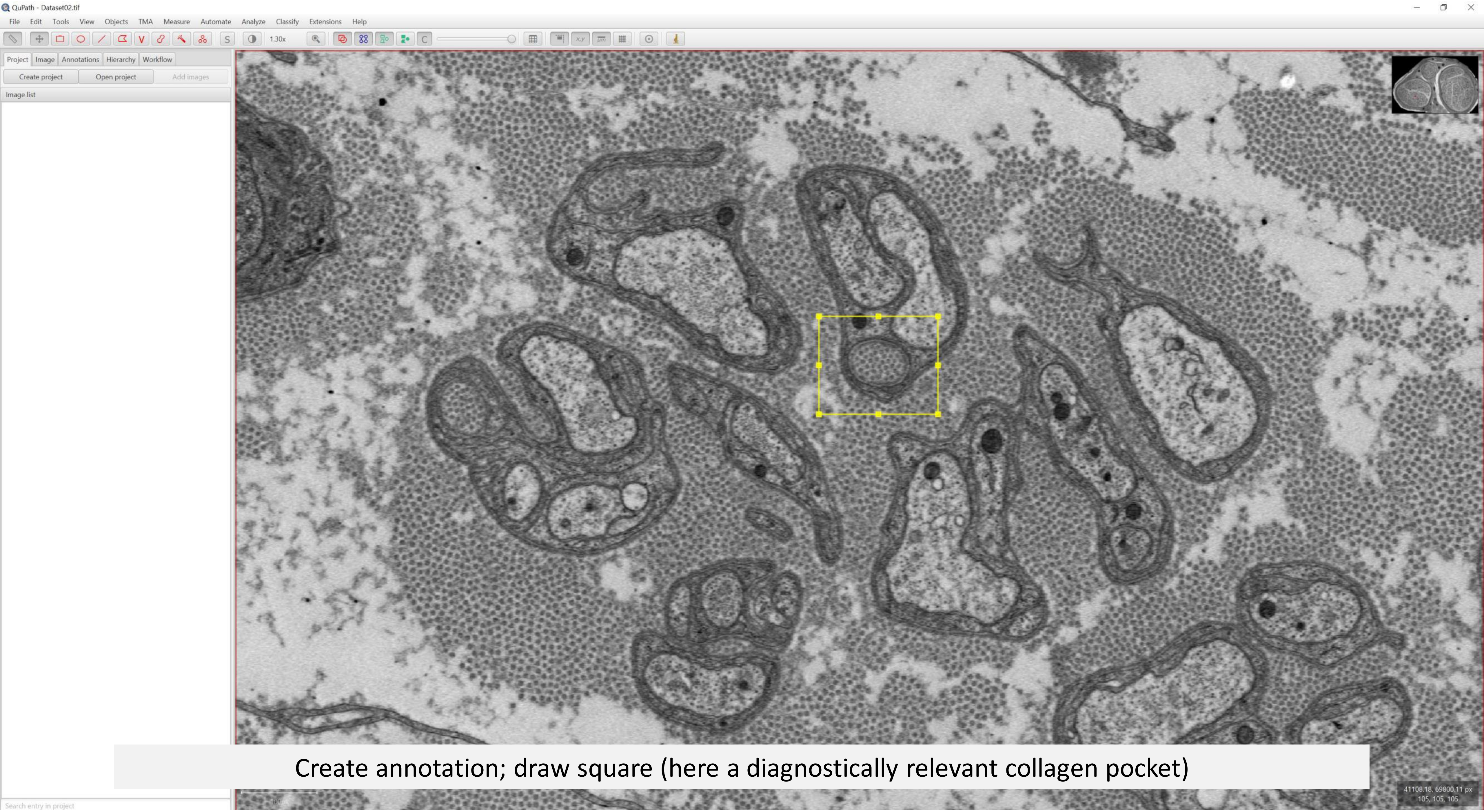


Create annotation; draw square (here a diagnostically relevant collagen pocket)

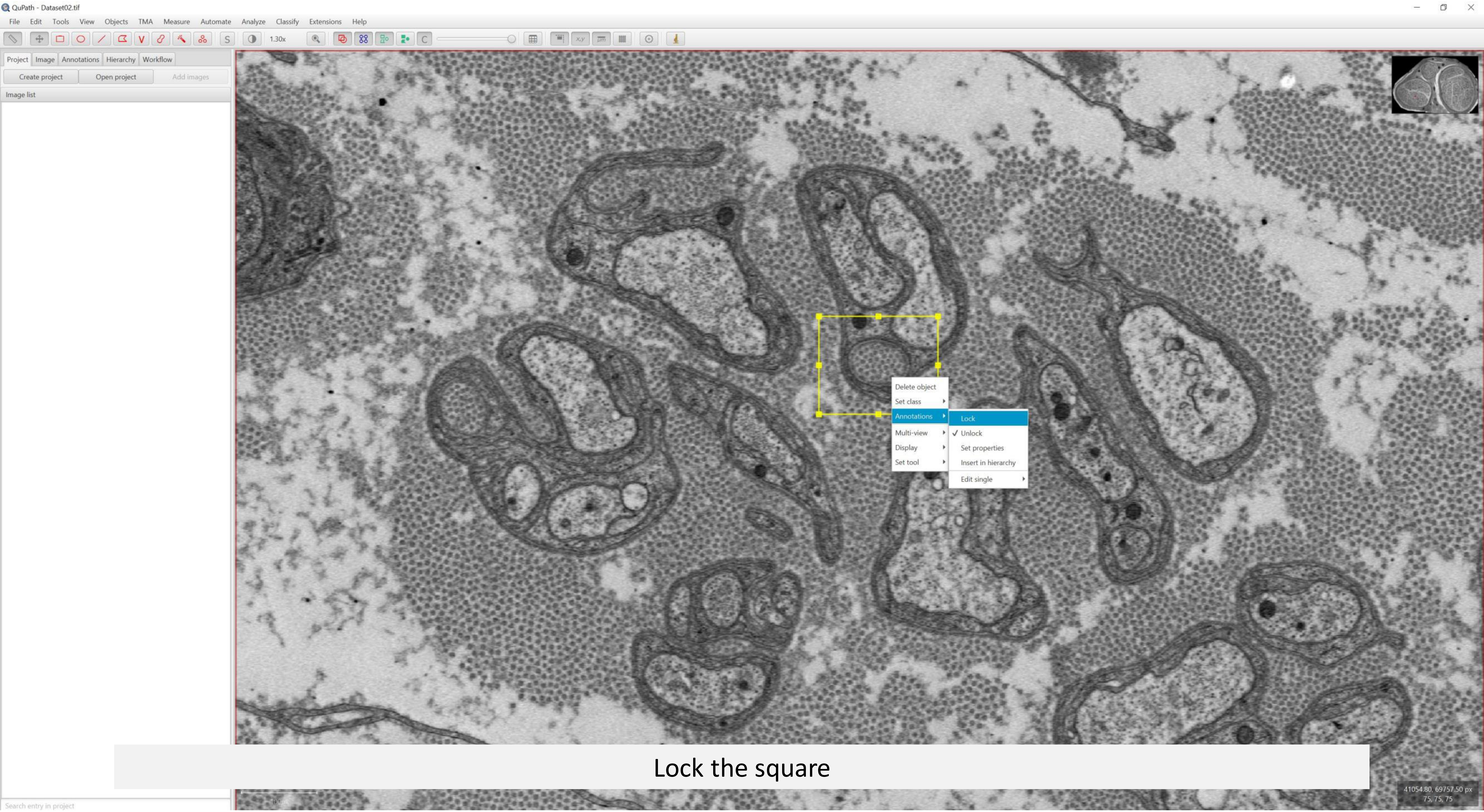


Lock the square

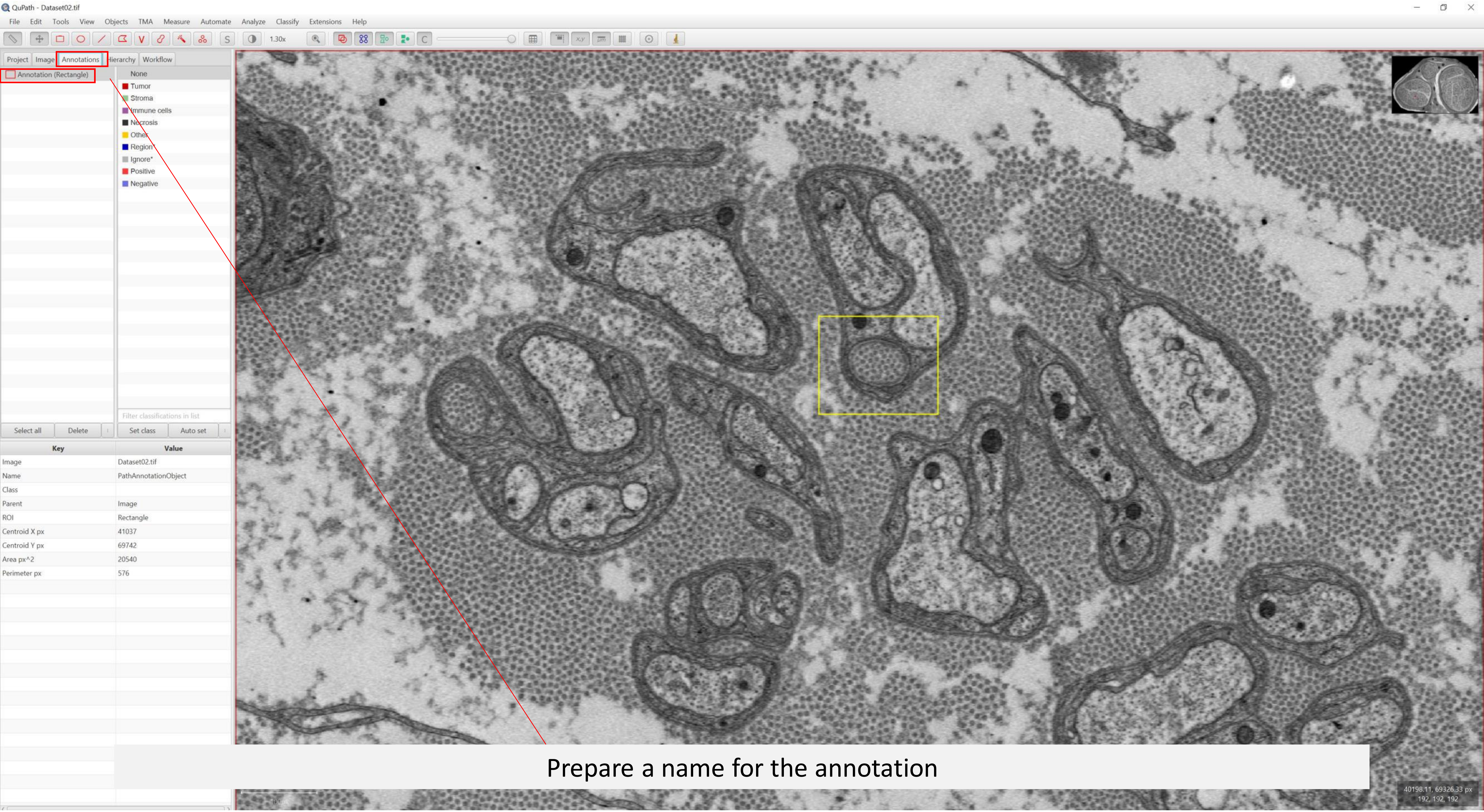


Prepare a name for the annotation

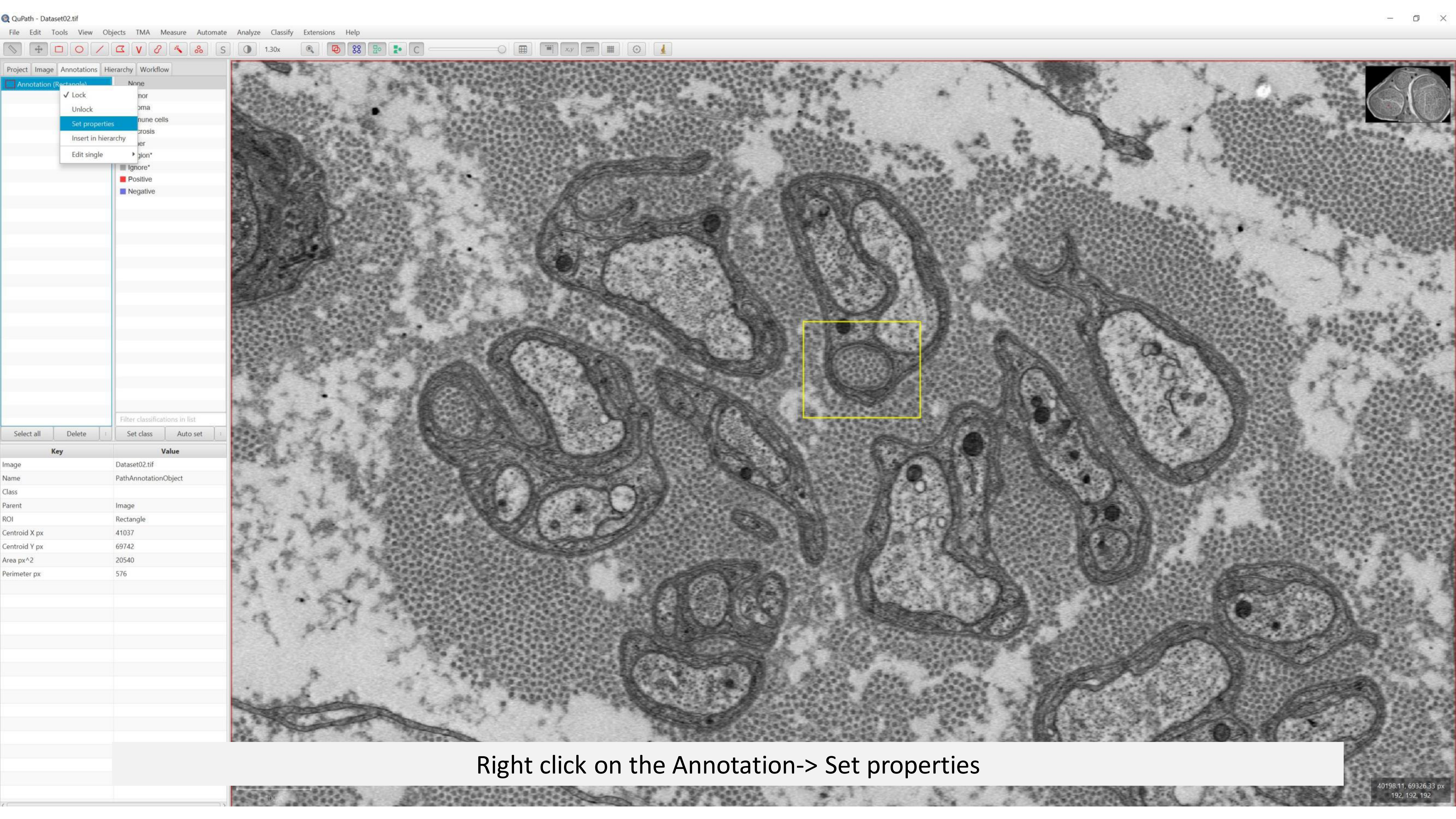
QuPath - Dataset02.tif
File Edit Tools View Objects TMA Measure Automate Analyze Classify Extensions Help
Project Image Annotations Hierarchy Workflow
Lock
Unlock
Set properties
Insert in hierarchy
Edit single
Ignore*
Positive
Negative
Select all
Delete
Set class
Auto set
Key
Value
Image
Dataset02.tif
Name
PathAnnotationObject
Class
Parent
Image
ROI
Rectangle
Centroid X px
41037
Centroid Y px
69742
Area px^2
20540
Perimeter px
576
Right click on the Annotation-> Set properties

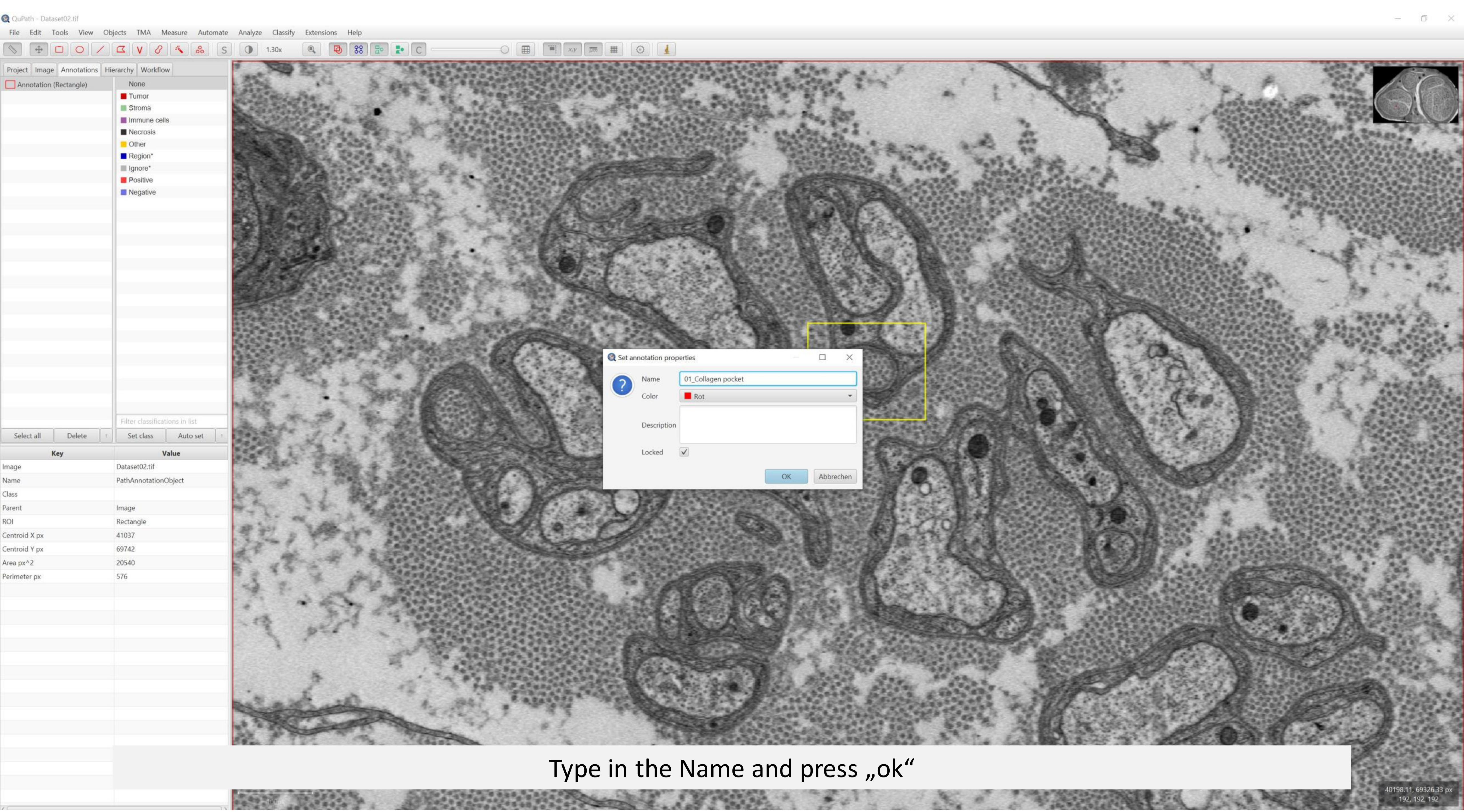


Type in the Name and press „ok“

QuPath - Dataset02.tif

File Edit Tools View Objects TMA Measure Automate Analyze Classify Extensions Help

1.30x

Project | Image | Annotations | Hierarchy | Workflow

01_Collagen pocket (Rectangle)

None
- Tumor
- Stroma
- Immune cells
- Necrosis
- Other
- Region*
- Ignore*
- Positive
- Negative

Filter classifications in list

Select all | Delete | Set class | Auto set

| Key | Value |
|---|---|
| Image | Dataset02.tif |
| Name | 01_Collagen pocket |
| Class | |
| Parent | Image |
| ROI | Rectangle |
| Centroid X px | 41037 |
| Centroid Y px | 69742 |
| Area px^2 | 20540 |
| Perimeter px | 576 |

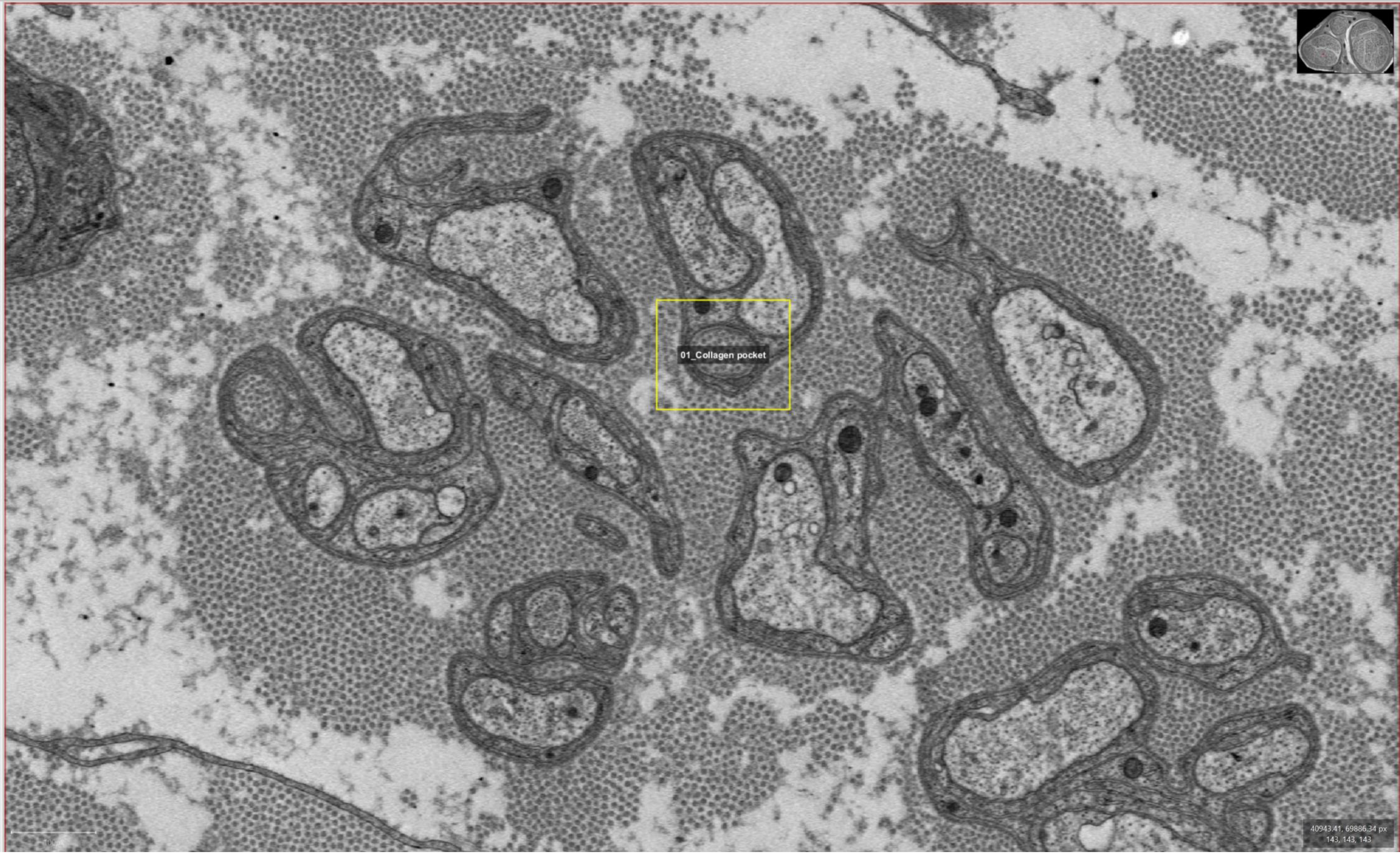

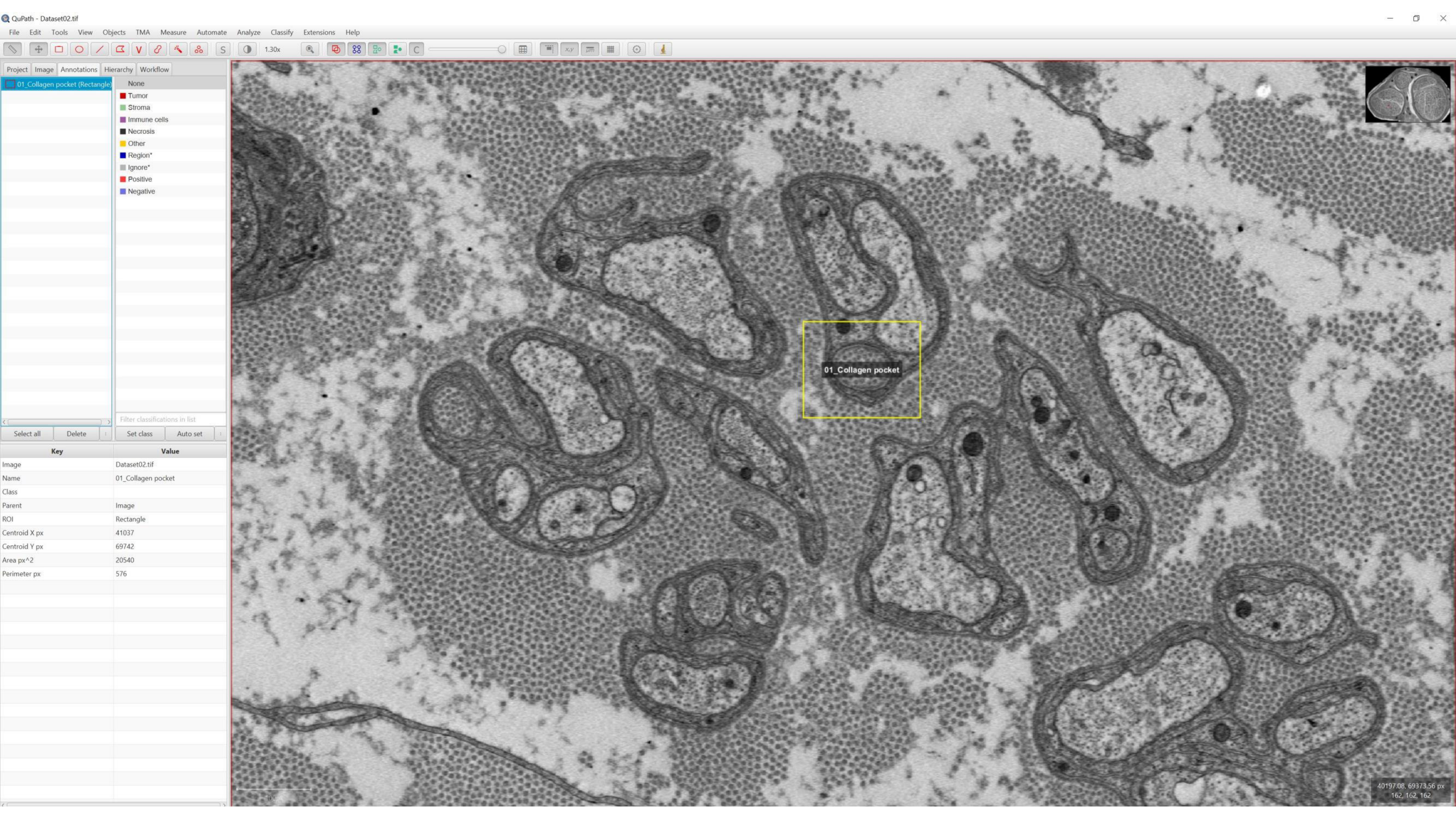
QuPath - Dataset02.tif
File
Edit
Tools
View
Objects
TMA
Measure
Automate
Analyze
Classify
Extensions
Help
1.30x
Project
Image
Annotations
Hierarchy
Workflow
01_Collagen pocket (Rectangle)
None
Tumor
Stroma
Immune cells
Necrosis
Other
Region*
Ignore*
Positive
Negative
Filter classifications in list
Select all
Delete
Set class
Auto set
Key
Value
Image
Dataset02.tif
Name
01_Collagen pocket
Class
Parent
Image
ROI
Rectangle
Centroid X px
41037
Centroid Y px
69742
Area px^2
20540
Perimeter px
576
01_Collagen pocket
40197.08, 69373.56 px
162, 162, 162

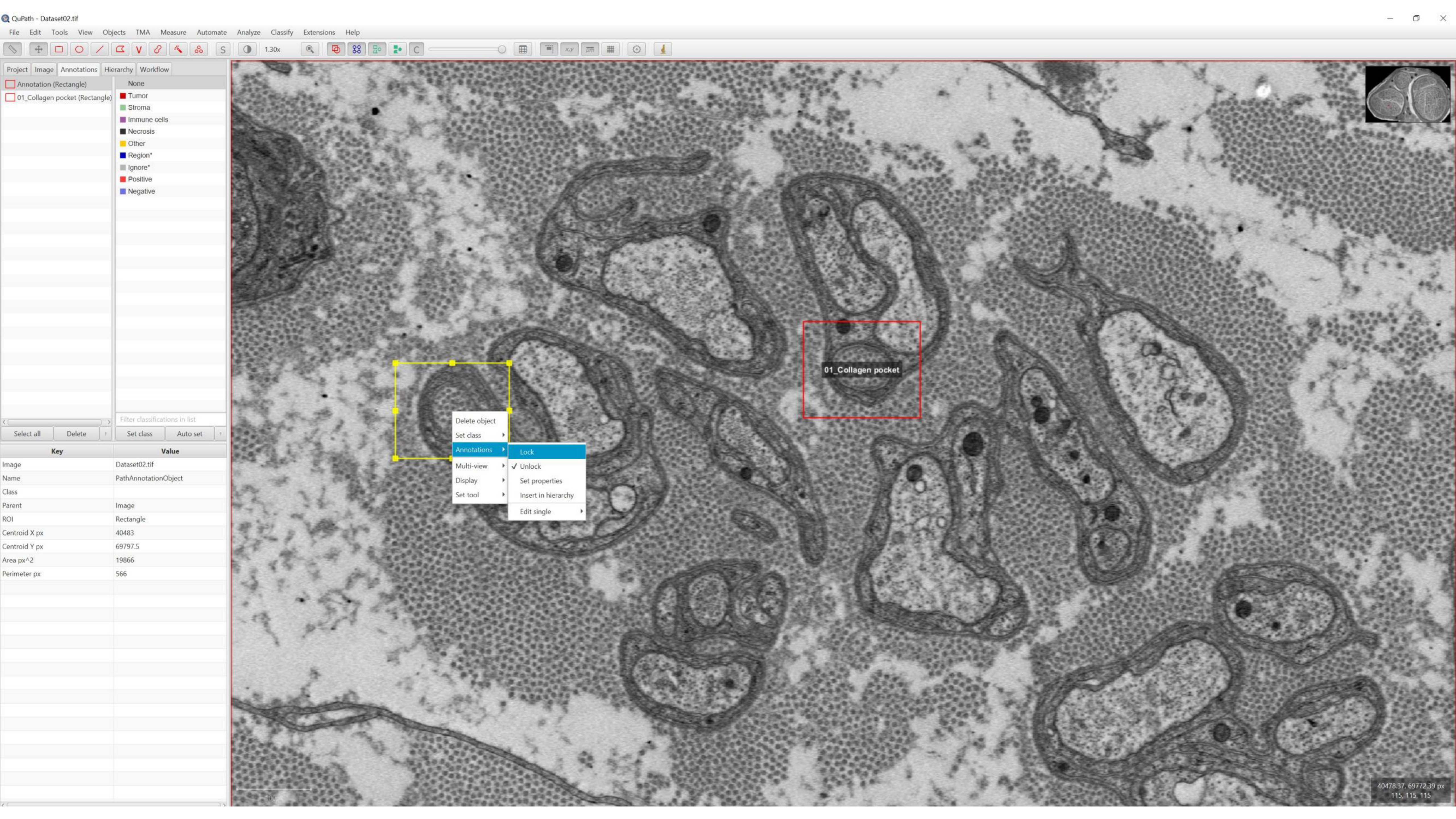
QuPath - Dataset02.tif
File Edit Tools View Objects TMA Measure Automate Analyze Classify Extensions Help
1.30x
Project Image Annotations Hierarchy Workflow
Annotation (Rectangle)
01_Collagen pocket (Rectangle)
None
Tumor
Stroma
Immune cells
Necrosis
Other
Region*
Ignore*
Positive
Negative
Filter classifications in list
Select all
Delete
Set class
Auto set
Key
Value
Image
Dataset02.tif
Name
PathAnnotationObject
Class
Parent
Image
ROI
Rectangle
Centroid X px
40483
Centroid Y px
69797.5
Area px^2
19866
Perimeter px
566
01_Collagen pocket
Delete object
Set class
Annotations
Multi-view
Display
Set tool
Lock
✓ Unlock
Set properties
Insert in hierarchy
Edit single
40478.37, 69772.39 px
115, 115, 115

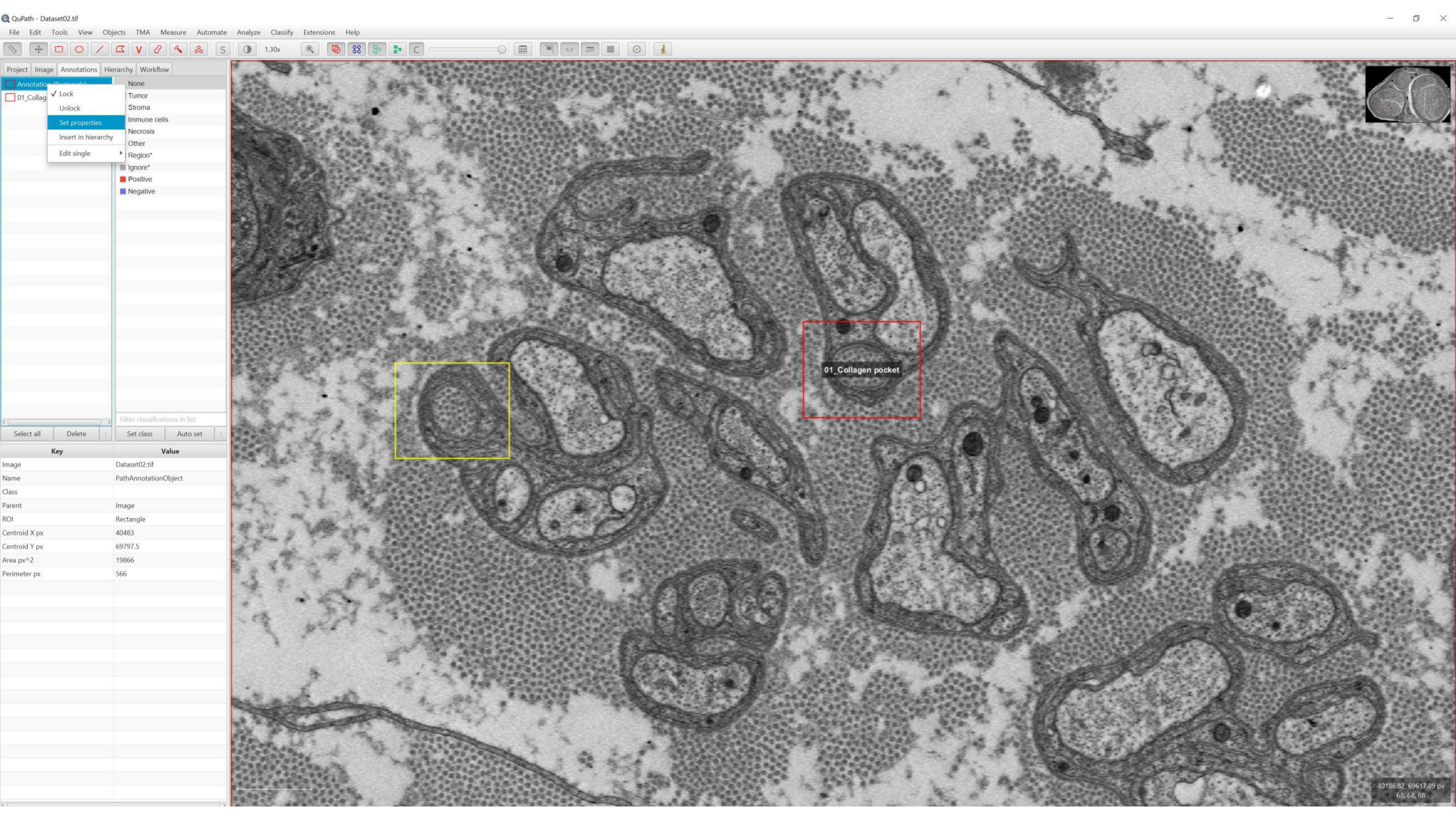
QuPath - Dataset02.tif
File Edit Tools View Objects TMA Measure Automate Analyze Classify Extensions Help
1.30x
Project Image Annotations Hierarchy Workflow
01_Collag
Lock
Unlock
Set properties
Insert in hierarchy
Edit single
None
Tumor
Stroma
Immune cells
Necrosis
Other
Region*
Ignore*
Positive
Negative
Filter classifications in list
Select all
Delete
Set class
Auto set
Key
Value
Image
Dataset02.tif
Name
PathAnnotationObject
Class
Parent
Image
ROI
Rectangle
Centroid X px
40483
Centroid Y px
69797.5
Area px^2
19866
Perimeter px
566
01_Collagen pocket
40186.82, 69617.89 px
68, 68, 68

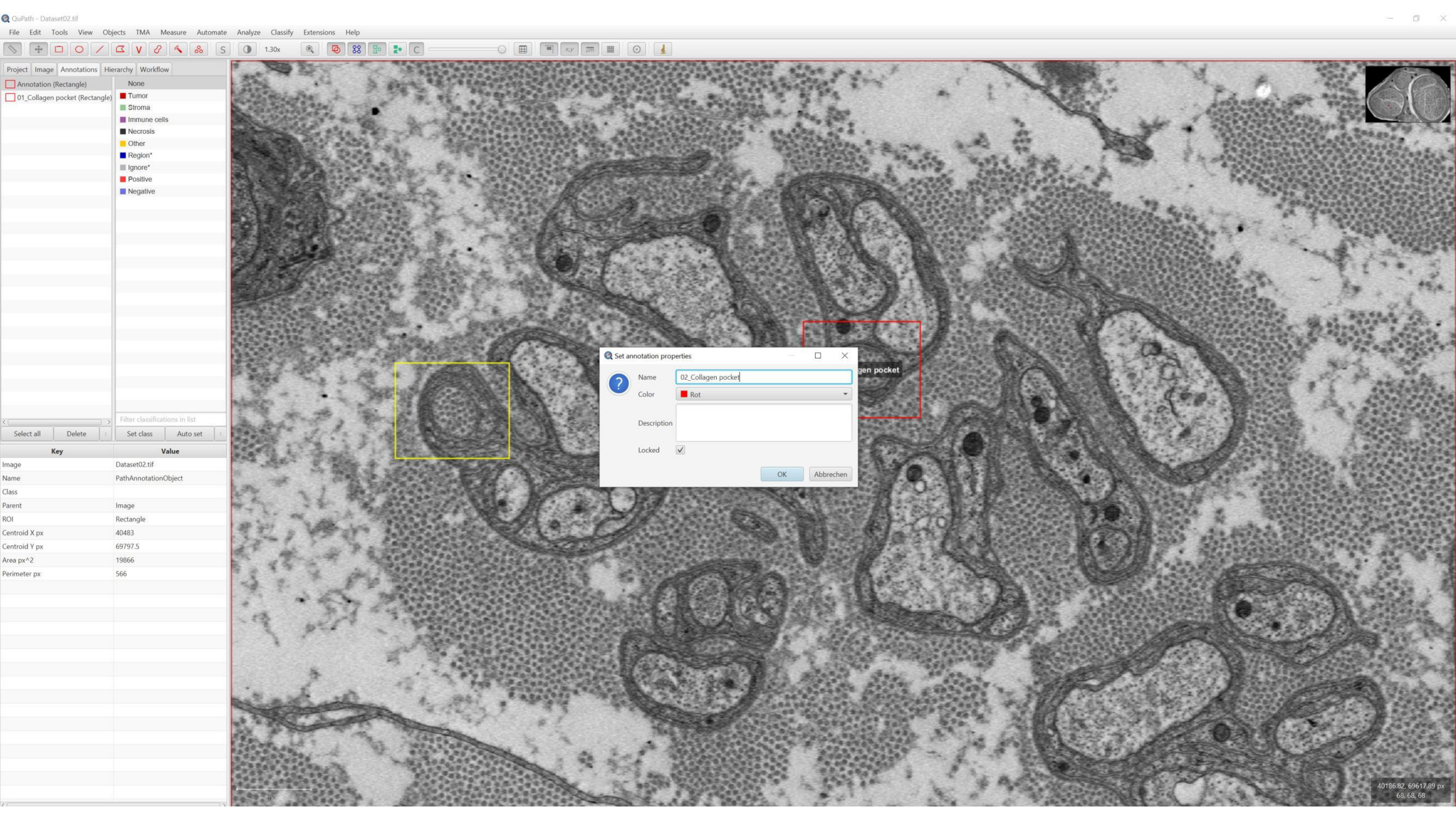
QuPath - Dataset02.tif
File Edit Tools View Objects TMA Measure Automate Analyze Classify Extensions Help
1.30x
Project Image Annotations Hierarchy Workflow
Annotation (Rectangle)
01_Collagen pocket (Rectangle)
None
Tumor
Stroma
Immune cells
Necrosis
Other
Region*
Ignore*
Positive
Negative
Filter classifications in list
Select all
Delete
Set class
Auto set
Key
Value
Image
Dataset02.tif
Name
PathAnnotationObject
Class
Parent
Image
ROI
Rectangle
Centroid X px
40483
Centroid Y px
69797.5
Area px^2
19866
Perimeter px
566
Set annotation properties
Name
02_Collagen pocket
Color
Rot
Description
Locked
OK
Abbrechen
gen pocket
40186.82, 69617.89 px
68, 68, 68

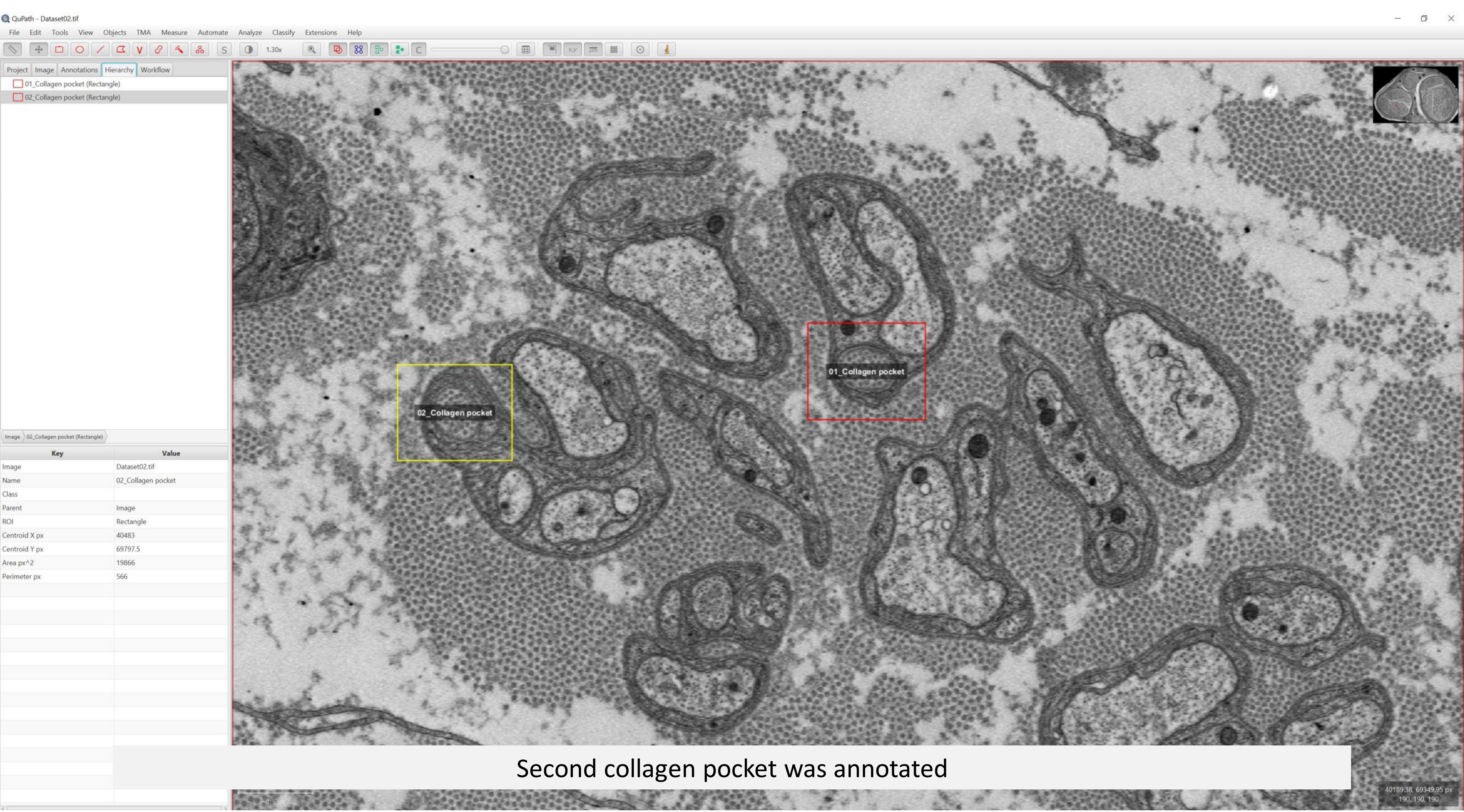


Second collagen pocket was annotated

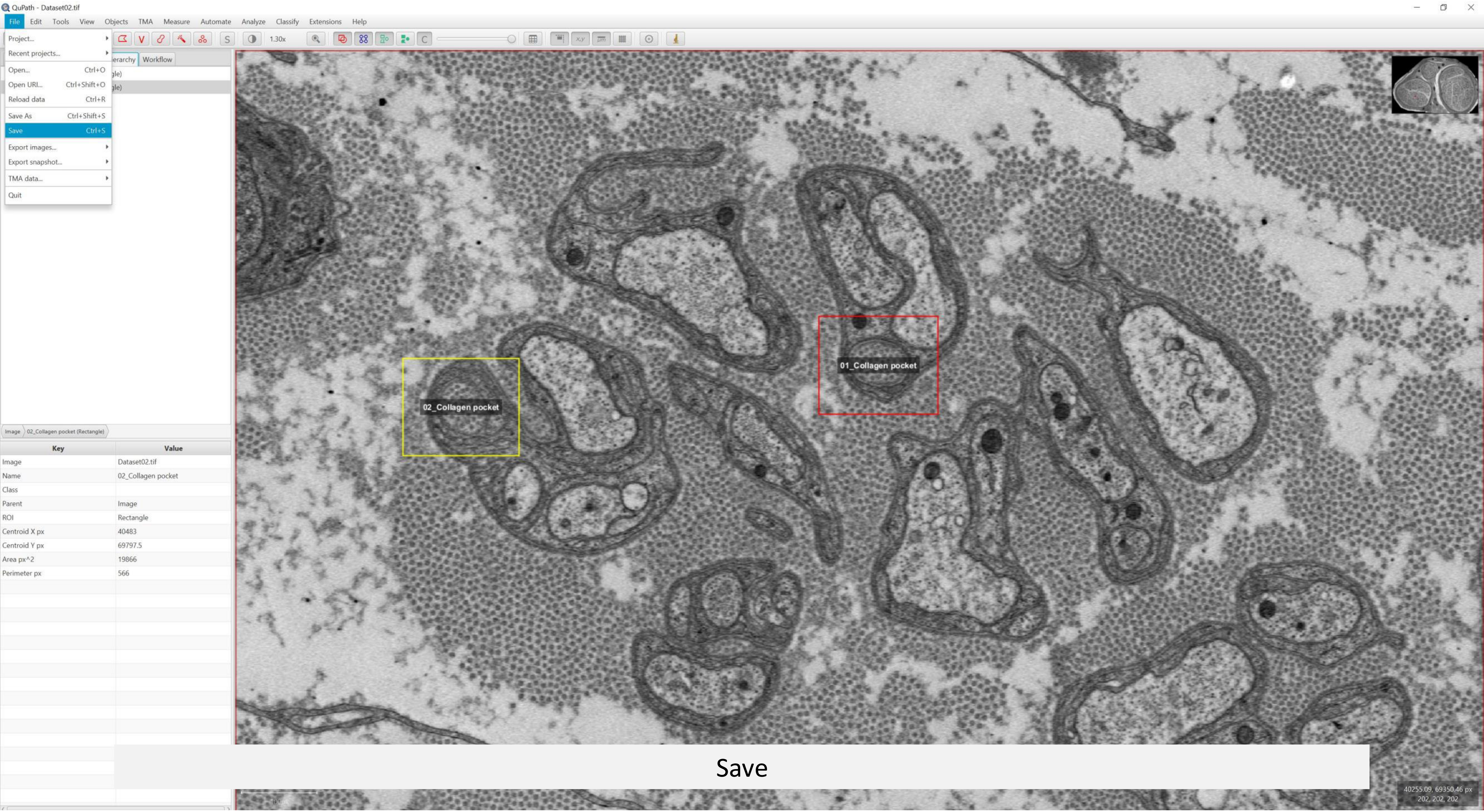
QuPath - Dataset02.tif
File
Edit
Tools
View
Objects
TMA
Measure
Automate
Analyze
Classify
Extensions
Help
Project...
Recent projects...
Open... Ctrl+O
Open URI... Ctrl+Shift+O
Reload data Ctrl+R
Save As Ctrl+Shift+S
Save Ctrl+S
Export images...
Export snapshot...
TMA data...
Quit
1.30x
Workflow
Image
02_Collagen pocket (Rectangle)
Key
Value
Image
Dataset02.tif
Name
02_Collagen pocket
Class
Parent
Image
ROI
Rectangle
Centroid X px
40483
Centroid Y px
69797.5
Area px^2
19866
Perimeter px
566
02_Collagen pocket
01_Collagen pocket
40255.09, 69350.46 px
202, 202, 202

Save

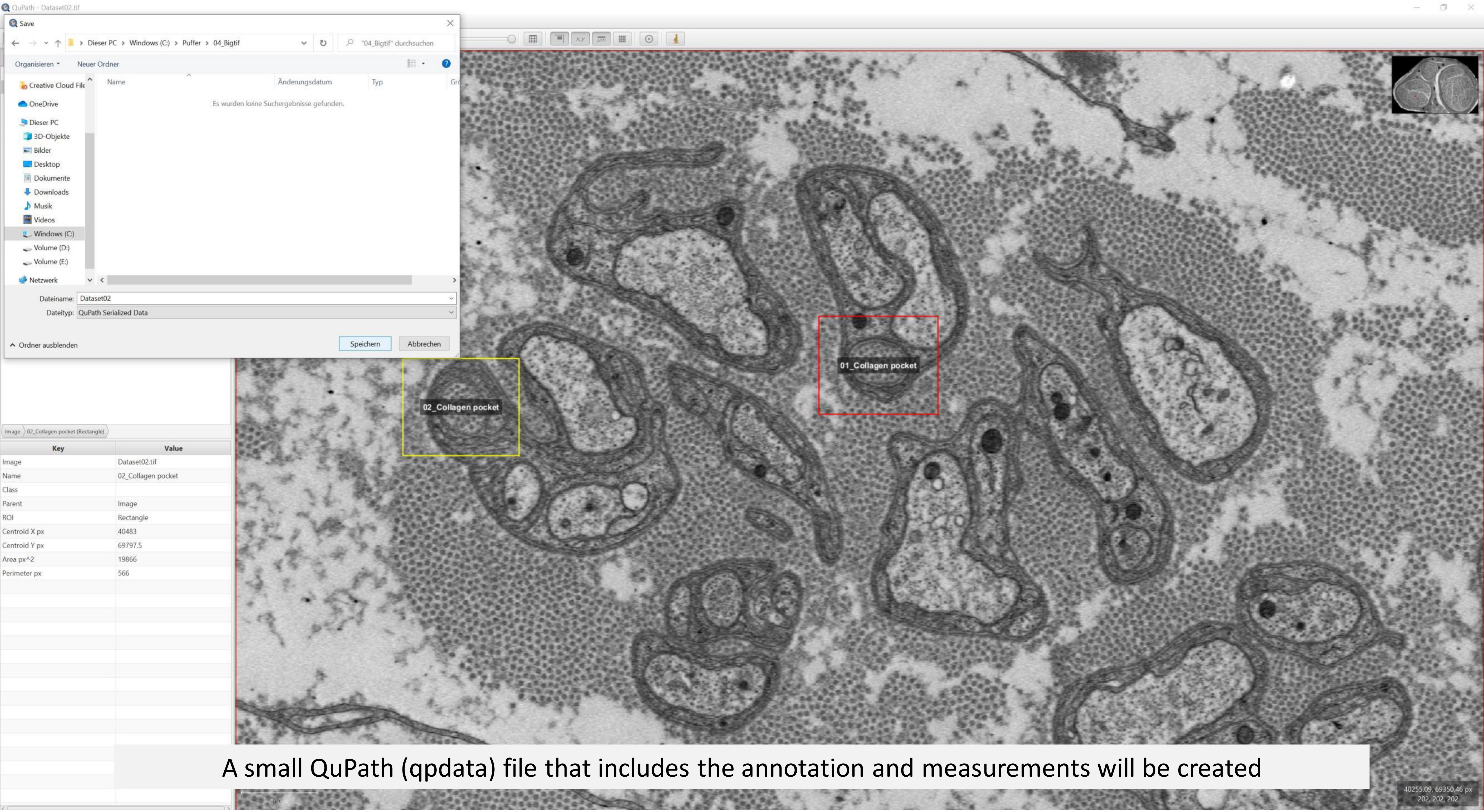


A small QuPath (qpdata) file that includes the annotation and measurements will be created

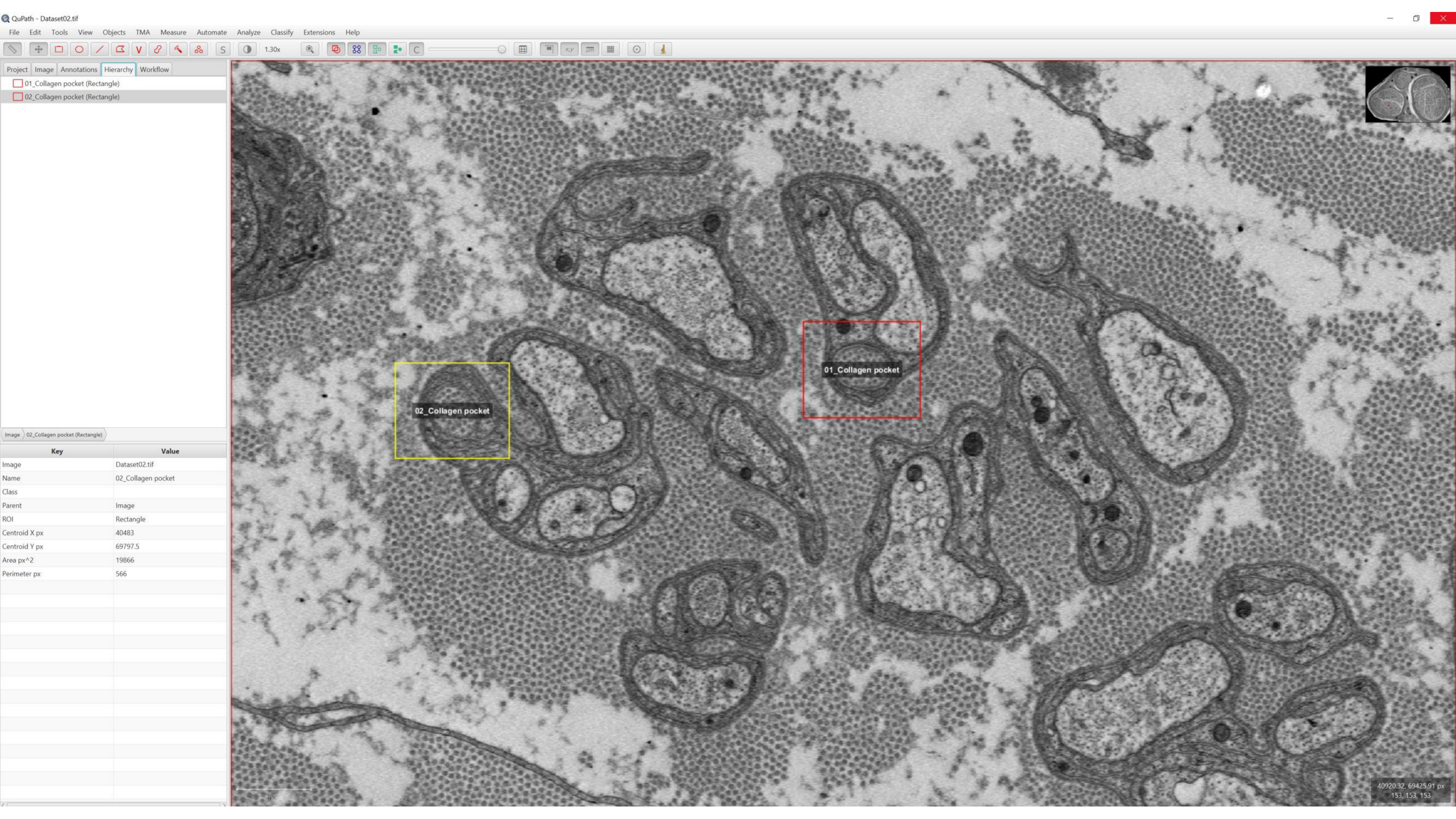
QuPath - Dataset02.tif
File Edit Tools View Objects TMA Measure Automate Analyze Classify Extensions Help
1.30x
Project Image Annotations Hierarchy Workflow
01_Collagen pocket (Rectangle)
02_Collagen pocket (Rectangle)
Image 02_Collagen pocket (Rectangle)
Key Value
Image Dataset02.tif
Name 02_Collagen pocket
Class
Parent Image
ROI Rectangle
Centroid X px 40483
Centroid Y px 69797.5
Area px^2 19866
Perimeter px 566
02_Collagen pocket
01_Collagen pocket
40920.32, 69425.91 px
153, 153, 153

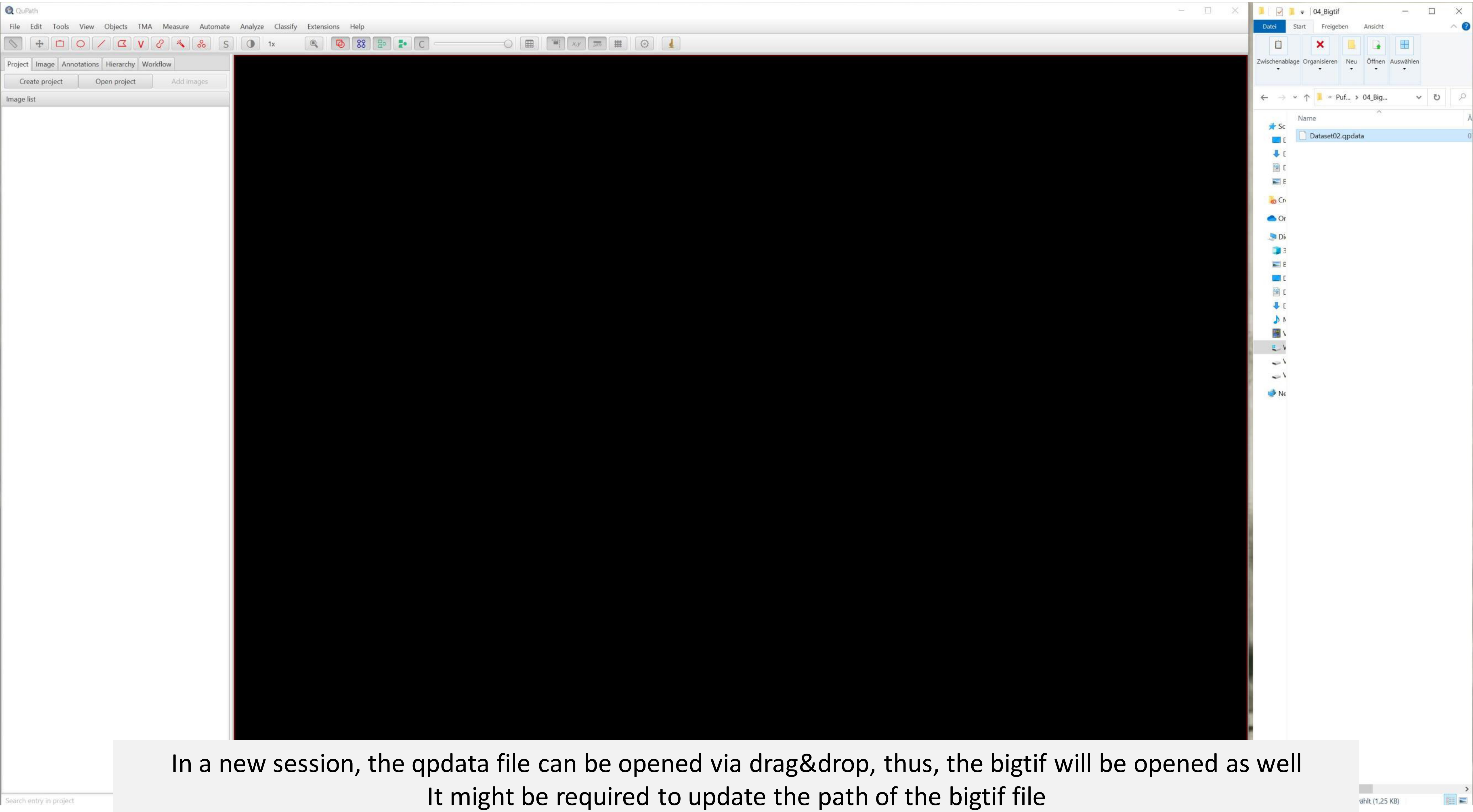


In a new session, the qpdata file can be opened via drag&drop, thus, the bigtif will be opened as well
It might be required to update the path of the bigtif file

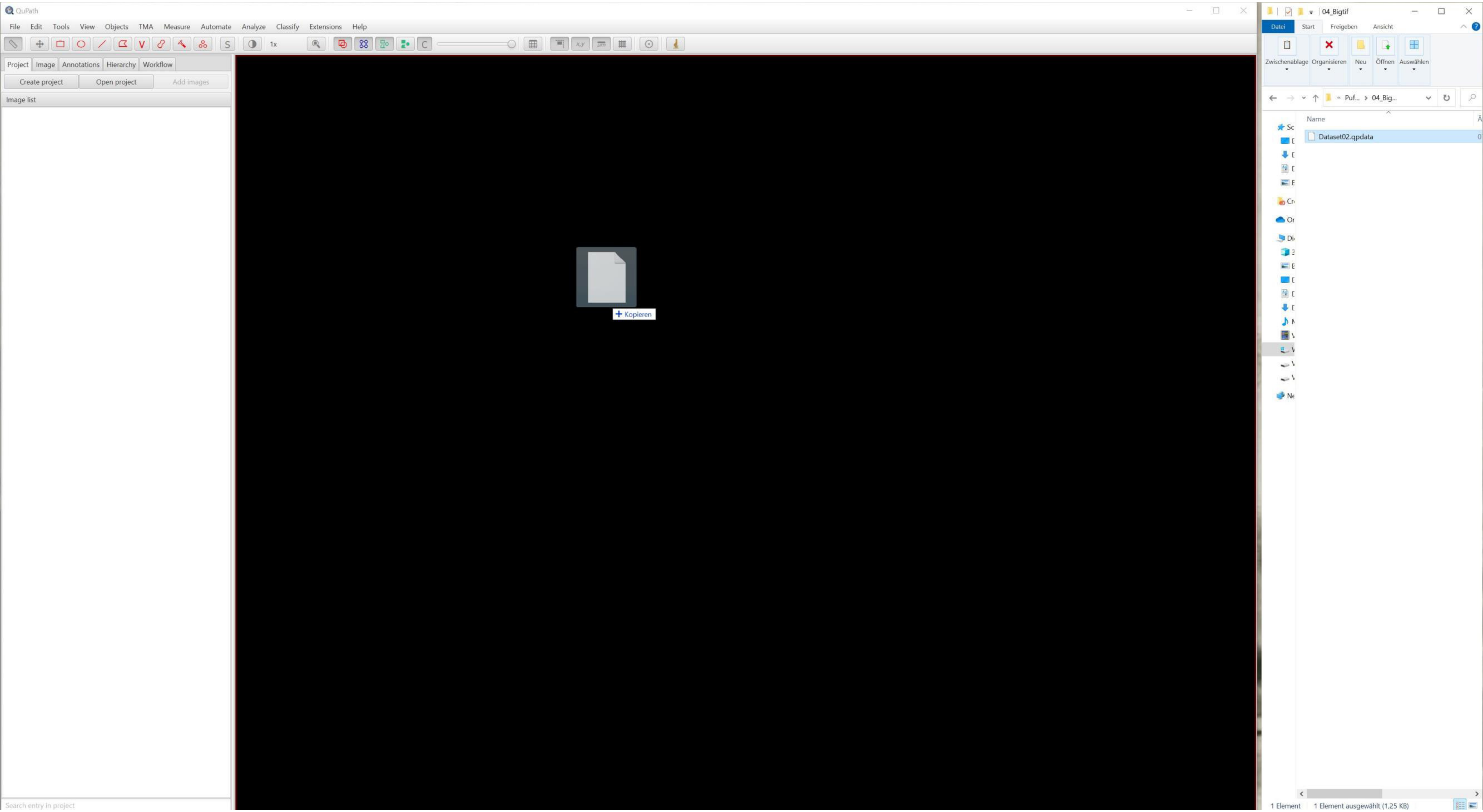
QuPath
File
Edit
Tools
View
Objects
TMA
Measure
Automate
Analyze
Classify
Extensions
Help
1x
Project
Image
Annotations
Hierarchy
Workflow
Create project
Open project
Add images
Image list
Search entry in project
+ Kopieren
04_Bigtif
Datei
Start
Freigeben
Ansicht
Zwischenablage
Organisieren
Neu
Öffnen
Auswählen
Puf...
04_Big...
Name
Dataset02.qpdata
1 Element
1 Element ausgewählt (1,25 KB)

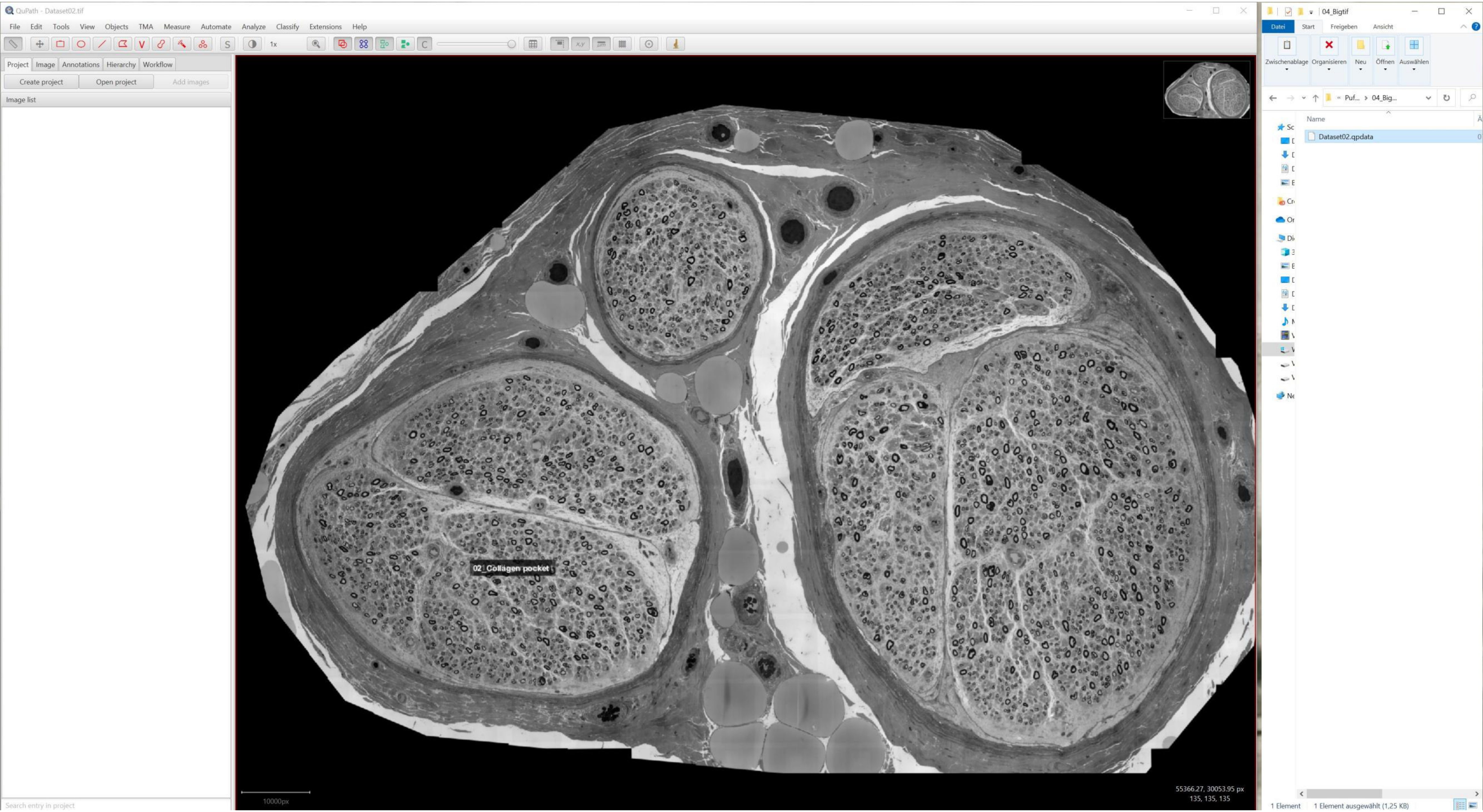
QuPath - Dataset02.tif
File Edit Tools View Objects TMA Measure Automate Analyze Classify Extensions Help
Project Image Annotations Hierarchy Workflow
Create project
Open project
Add images
Image list
02_Collagen pocket
10000px
55366.27, 30053.95 px
135, 135, 135
Search entry in project
04_Bigtif
Datei Start Freigeben Ansicht
Zwischenablage Organisieren Neu Öffnen Auswählen
Puf... › 04_Big...
Name
Dataset02.qpdata
1 Element
1 Element ausgewählt (1,25 KB)

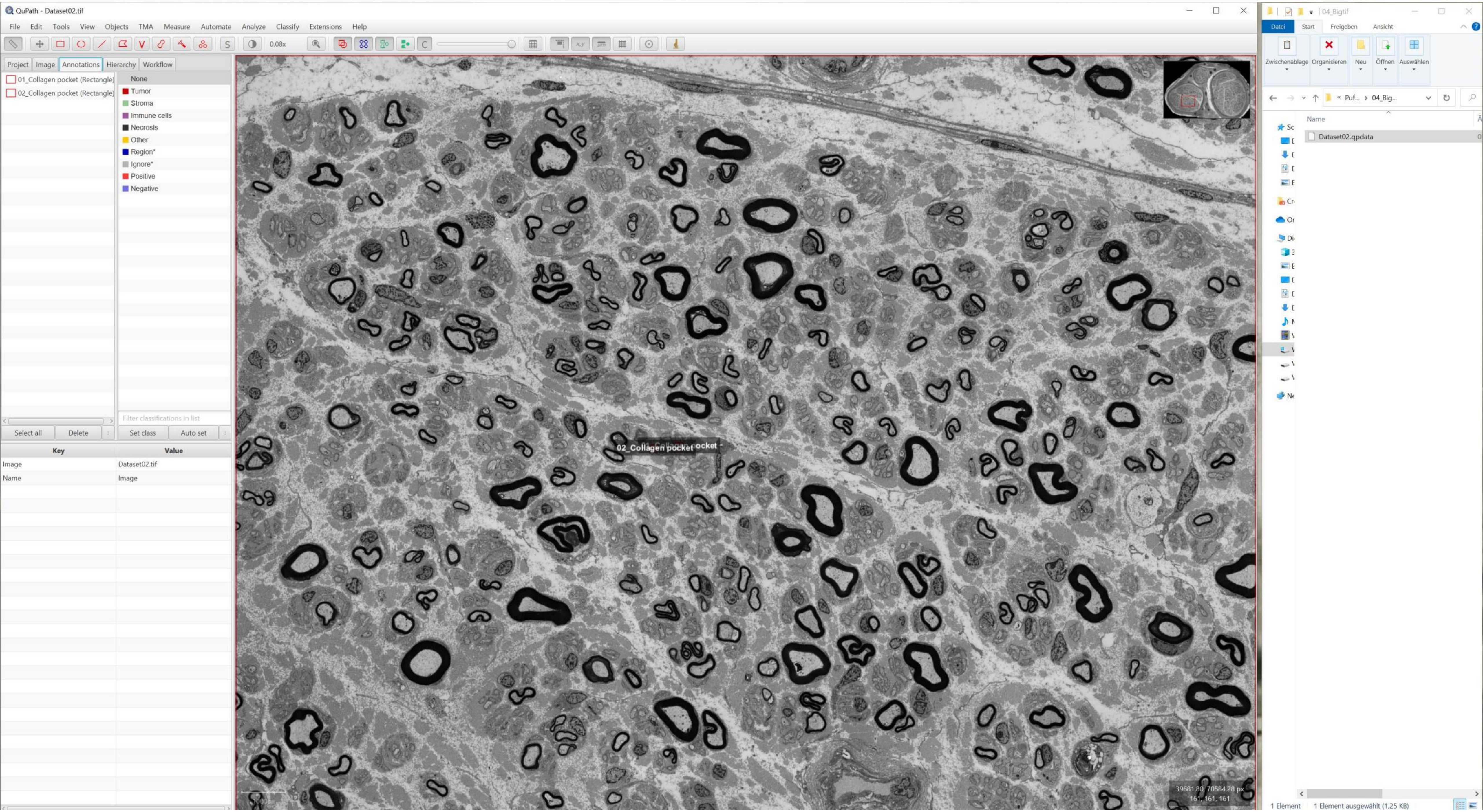
QuPath - Dataset02.tif
File Edit Tools View Objects TMA Measure Automate Analyze Classify Extensions Help
0.08x
Project Image Annotations Hierarchy Workflow
01_Collagen pocket (Rectangle)
02_Collagen pocket (Rectangle)
None
Tumor
Stroma
Immune cells
Necrosis
Other
Region*
Ignore*
Positive
Negative
Filter classifications in list
Select all
Delete
Set class
Auto set
Key
Value
Image
Dataset02.tif
Name
Image
02_Collagen pocket
39681.80, 70584.28 px
161, 161, 161
04_Bigtif
Datei Start Freigeben Ansicht
Zwischenablage Organisieren Neu Öffnen Auswählen
Puf... › 04_Big...
Name
Dataset02.qpdata
1 Element
1 Element ausgewählt (1,25 KB)

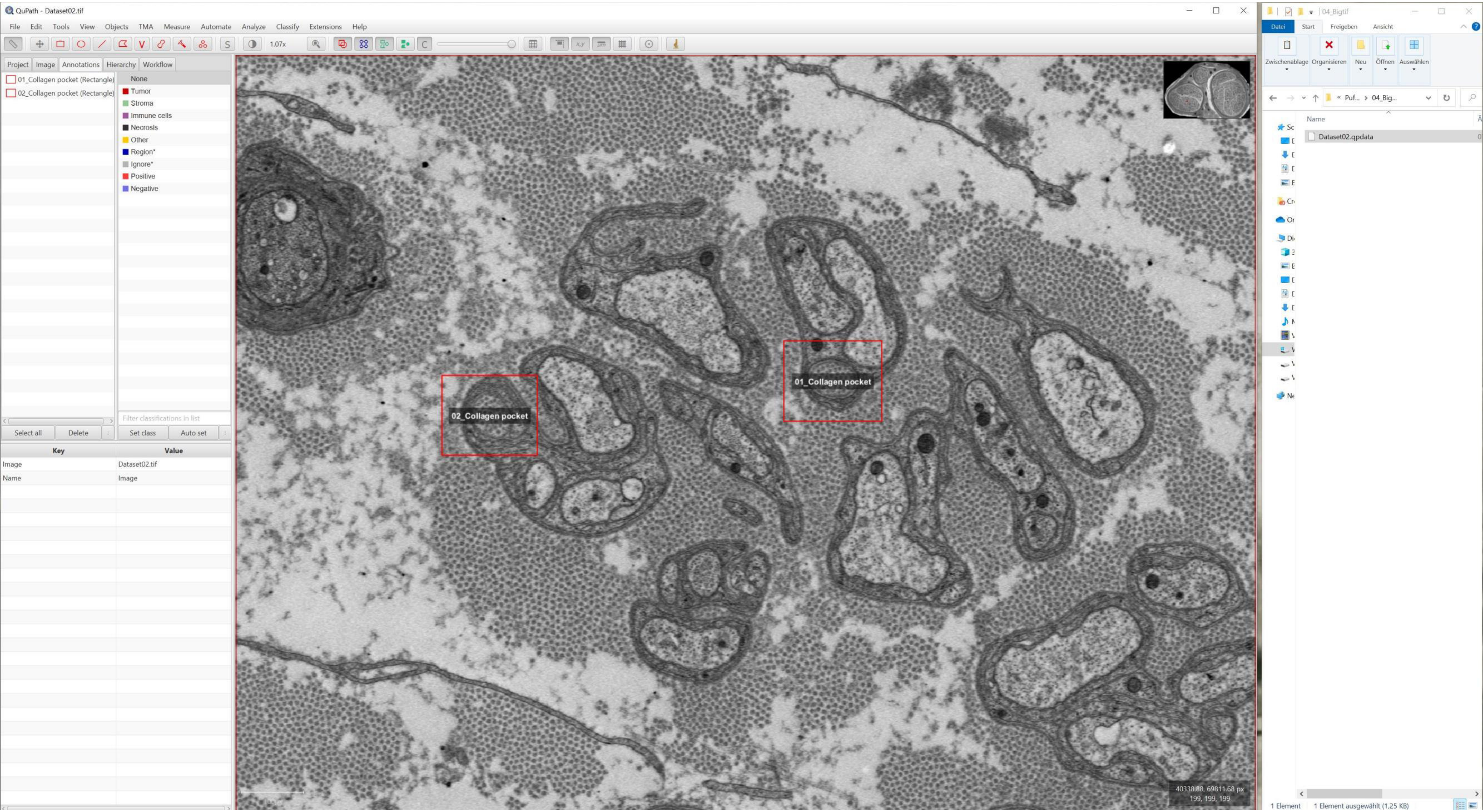
QuPath - Dataset02.tif
File Edit Tools View Objects TMA Measure Automate Analyze Classify Extensions Help
1.07x
Project Image Annotations Hierarchy Workflow
01_Collagen pocket (Rectangle)
02_Collagen pocket (Rectangle)
None
Tumor
Stroma
Immune cells
Necrosis
Other
Region*
Ignore*
Positive
Negative
Filter classifications in list
Select all
Delete
Set class
Auto set
Key
Value
Image
Dataset02.tif
Name
Image
02_Collagen pocket
01_Collagen pocket
40338.88, 69811.68 px
199, 199, 199
04_Bigtif
Datei Start Freigeben Ansicht
Zwischenablage Organisieren Neu Öffnen Auswählen
Puf... › 04_Big...
Name
Dataset02.qpdata
1 Element
1 Element ausgewählt (1,25 KB)

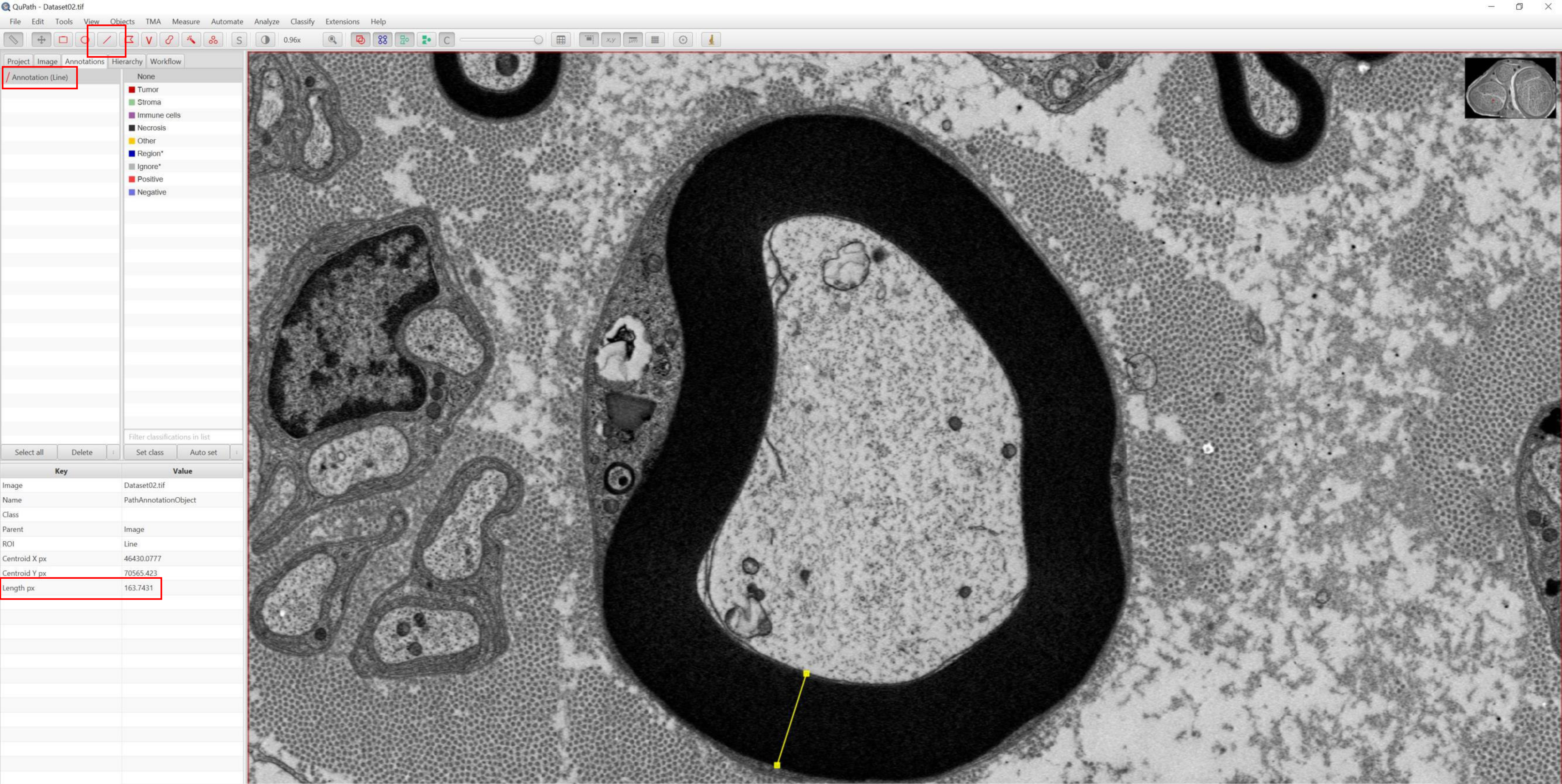


A line can be used to measure distances; 163 pixels * 9 nm pixel size= 1,467 nm

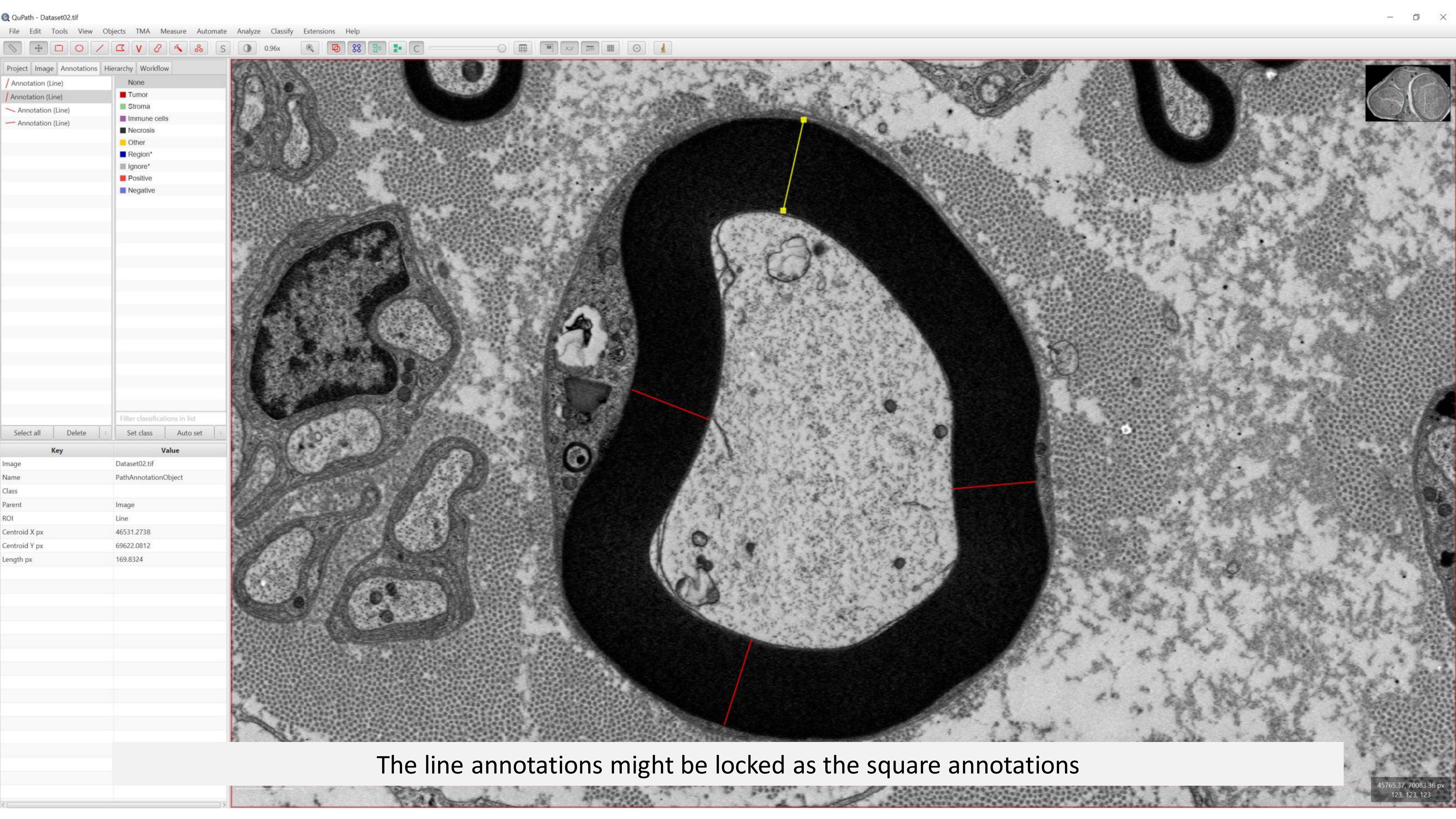


The line annotations might be locked as the square annotations

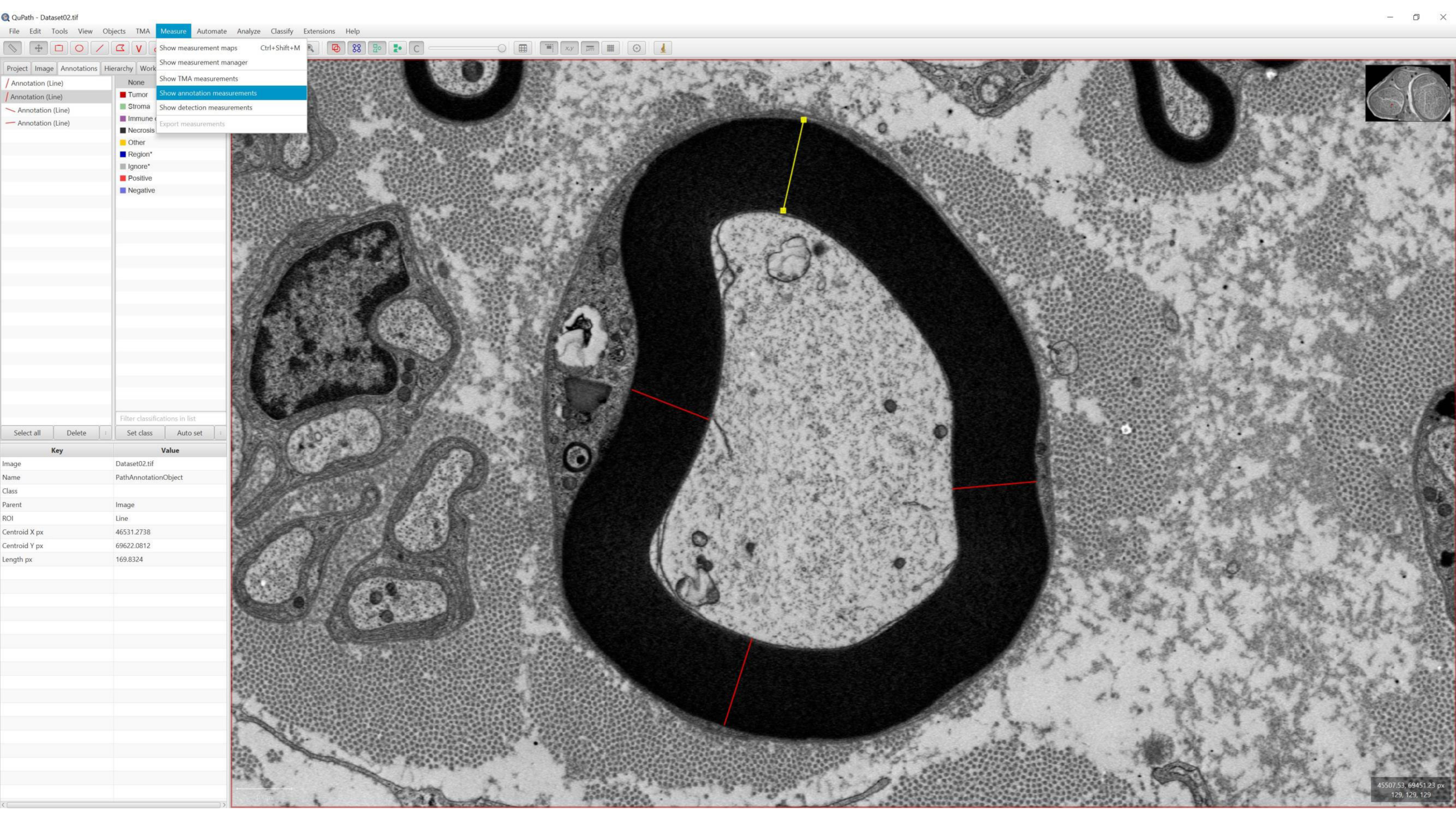
QuPath - Dataset02.tif
File Edit Tools View Objects TMA Measure Automate Analyze Classify Extensions Help
Show measurement maps Ctrl+Shift+M
Show measurement manager
Show TMA measurements
Show annotation measurements
Show detection measurements
Export measurements
Project Image Annotations Hierarchy Work
Annotation (Line)
Annotation (Line)
Annotation (Line)
Annotation (Line)
None
Tumor
Stroma
Immune
Necrosis
Other
Region*
Ignore*
Positive
Negative
Filter classifications in list
Select all Delete
Set class Auto set
Key Value
Image Dataset02.tif
Name PathAnnotationObject
Class
Parent Image
ROI Line
Centroid X px 46531.2738
Centroid Y px 69622.0812
Length px 169.8324
45507.53, 69451.23 px
129, 129, 129

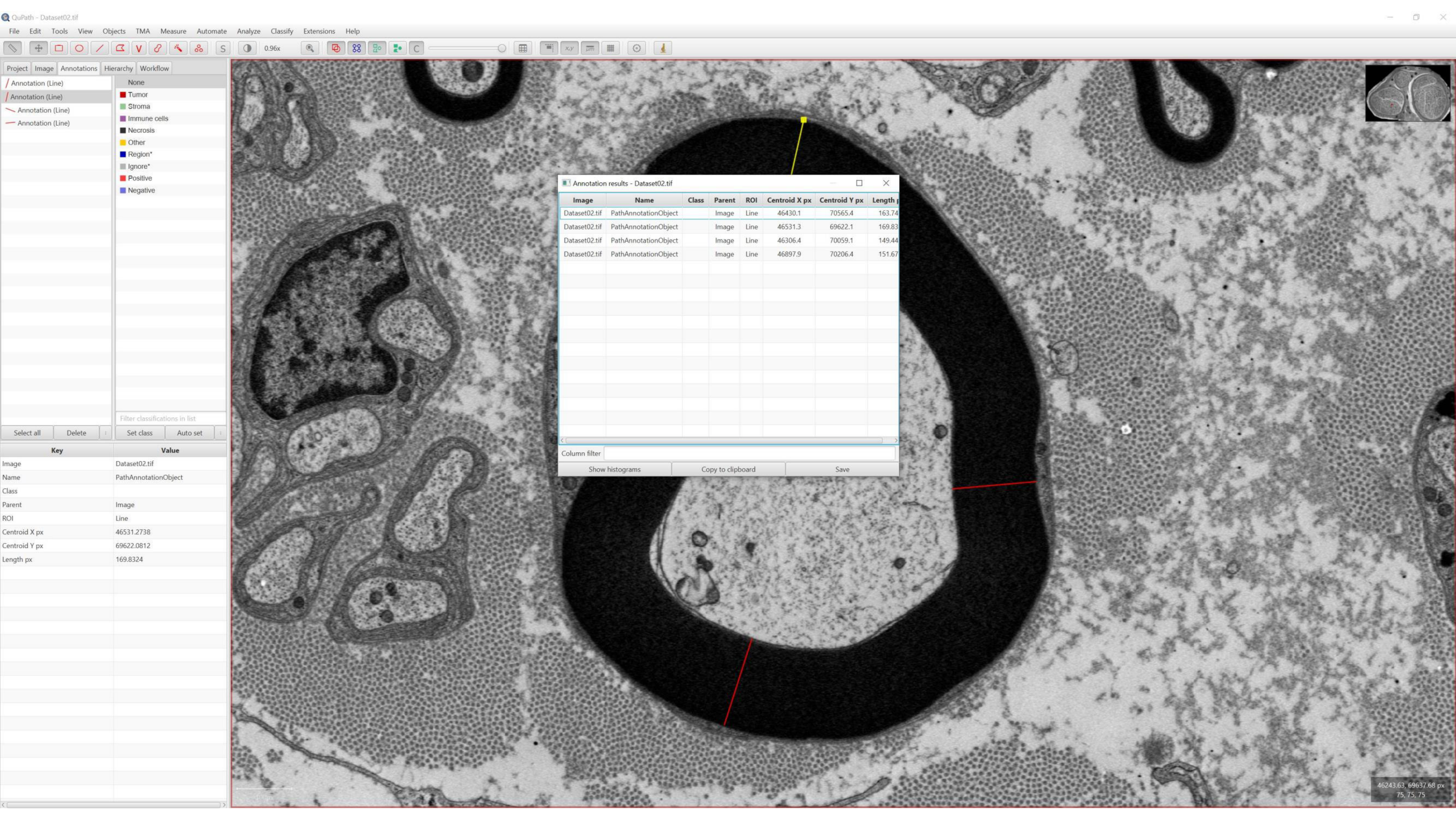
QuPath - Dataset02.tif
File Edit Tools View Objects TMA Measure Automate Analyze Classify Extensions Help
0.96x
Project Image Annotations Hierarchy Workflow
Annotation (Line)
Annotation (Line)
Annotation (Line)
Annotation (Line)
None
Tumor
Stroma
Immune cells
Necrosis
Other
Region*
Ignore*
Positive
Negative
Filter classifications in list
Select all Delete
Set class Auto set
Key Value
Image Dataset02.tif
Name PathAnnotationObject
Class
Parent Image
ROI Line
Centroid X px 46531.2738
Centroid Y px 69622.0812
Length px 169.8324
Annotation results - Dataset02.tif
Image Name Class Parent ROI Centroid X px Centroid Y px Length
Dataset02.tif PathAnnotationObject Image Line 46430.1 70565.4 163.74
Dataset02.tif PathAnnotationObject Image Line 46531.3 69622.1 169.83
Dataset02.tif PathAnnotationObject Image Line 46306.4 70059.1 149.44
Dataset02.tif PathAnnotationObject Image Line 46897.9 70206.4 151.67
Column filter
Show histograms Copy to clipboard Save
46243.63, 69637.68 px
75, 75, 75

EM Macro Editor -- carsten_SK_III_30s_02

File Edit View Insert Debug Help

Macro Name : carsten_SK_III_30s_02
Version
Stage goto X= 61.471 mm    Stage goto Y= 45.453 mm
Stage goto X= 61.471 mm    Stage goto Y= 45.453 mm
Stage goto Z= 29.880 mm
Stage goto R= 169.3 °
Scan Rotation= 356.1 °
Delay For 30 seconds
Stage goto X= 61.803 mm    Stage goto Y= 45.538 mm
Stage goto X= 61.803 mm    Stage goto Y= 45.538 mm
Stage goto R= 138.2 °
Scan Rotation= 358.6 °
Delay For 30 seconds
Stage goto X= 61.368 mm    Stage goto Y= 45.605 mm
Stage goto X= 61.368 mm    Stage goto Y= 45.605 mm
Stage goto R= 109.4 °
Scan Rotation= 18.4 °
Delay For 30 seconds
Stage goto X= 61.666 mm    Stage goto Y= 45.530 mm
Stage goto X= 61.666 mm    Stage goto Y= 45.530 mm
Stage goto R= 78.1 °
Scan Rotation= 4.2 °
Delay For 30 seconds
Stage goto X= 61.161 mm    Stage goto Y= 45.593 mm
Stage goto X= 61.161 mm    Stage goto Y= 45.593 mm
Stage goto R= 49.4 °
Scan Rotation= 358.5 °
Delay For 30 seconds
Stage goto X= 61.433 mm    Stage goto Y= 45.462 mm
Stage goto X= 61.433 mm    Stage goto Y= 45.462 mm
Stage goto R= 17.8 °
Scan Rotation= 10.3 °
Delay For 30 seconds
Stage goto X= 61.319 mm    Stage goto Y= 45.468 mm
Stage goto X= 61.319 mm    Stage goto Y= 45.468 mm
Stage goto R= 348.2 °
Scan Rotation= 348.7 °
Delay For 30 seconds
Stage goto X= 61.314 mm    Stage goto Y= 45.463 mm
Stage goto X= 61.314 mm    Stage goto Y= 45.463 mm
Stage goto R= 318.5 °
Scan Rotation= 356.5 °
Delay For 30 seconds
Stage goto X= 61.339 mm    Stage goto Y= 45.292 mm
Stage goto X= 61.339 mm    Stage goto Y= 45.292 mm
Stage goto R= 288.6 °
Scan Rotation= 0.1 °
Delay For 30 seconds
Stage goto X= 61.852 mm    Stage goto Y= 45.276 mm
Stage goto X= 61.852 mm    Stage goto Y= 45.276 mm
Stage goto R= 257.8 °
Scan Rotation= 348.4 °
Delay For 30 seconds
Stage goto X= 61.698 mm    Stage goto Y= 45.308 mm
Stage goto X= 61.698 mm    Stage goto Y= 45.308 mm
Stage goto R= 228.2 °
Scan Rotation= 357.3 °
Delay For 30 seconds
Stage goto X= 62.060 mm    Stage goto Y= 45.274 mm
Stage goto X= 62.060 mm    Stage goto Y= 45.274 mm
Stage goto R= 197.1 °
Scan Rotation= 355.9 °
Delay For 30 seconds
Delay For 3600 seconds
EHT Off
